# Chenodeoxycholic Acid Modulation via *Bacteroides intestinalis* AM1 underscores a Novel Approach in Acute Liver Failure

**DOI:** 10.1101/2025.06.11.659225

**Authors:** Sushmita Pandey, Neha Sharma, Nupur Sharma, Sadam H Bhat, Vipul Sharma, Manish Kushwaha, Anil Kumar, Yash Magar, Abhishak Gupta, Babu Mathew, Gaurav Tripathi, Vasundhra Bindal, Sanju Yadav, Manisha Yadav, Anupama Kumari, Shvetank Sharma, Chhagan Bihari, Anju Katyal, Rakhi Maiwall, Shiv Kumar Sarin, Jaswinder Singh Maras

## Abstract

**Background:** Acute liver failure (ALF) is associated with rapid and progressive hepatocellular injury, and severe metabolic-microbial derangements. We investigated early metabolic markers of non-survival, and a potential microbial intervention using Bacteroides intestinalis-AM1, to improve outcomes in ALF.

**Method:** Plasma metabolomics and meta-proteomics were performed in 40 ALF patients and 5 healthy controls (training cohort). A non-survival marker panel was identified and validated in 270 ALF patients (test cohort) using high resolution mass spectrometry and machine learning. It was functionally validated in acetaminophen-induced ALF mouse model. B. intestinalis-AM1 was used to study alteration of gut bacteria and amelioration of liver injury.

**Results:** ALF non-survivors showed a distinct metabolomic signature with elevated primary bile acids {chenodeoxycholic acid (CDCA), cholic acid (CA)}, tryptophan, tyrosine, and enrichment of pathways linked to inflammation, cell death, and stress response (p<0.01, FDR<0.01, FC>1.5). Non-survivors had higher alpha/beta diversity (p<0.05) with increase in Proteobacteria, Firmicutes, Actinobacteria (p<0.05); functionally associated with energy, amino acid and xenobiotic metabolism (p<0.05). A gut microbiota derangement in converting primary to secondary bile acids was evident as CDCA and cytotoxic metabolites (4-(2-Amino phenyl)-2,4-dioxobutanoate, L-Tyrosine) were higher. Elevated CDCA (logFC>10) levels correlated with mortality in ALF patients as well as in mouse model. In the later, administration of *B. intestinalis*-AM1 bacteria, (10^9) reduced CDCA and CA levels by enhancing *FXR*, *FGF15*, *SLC10A1 gene expression,* attenuating inflammation (IL-1beta, TLR4-signalling), necroptosis, and modulating glutathione(oxidative-repair), tryptophan(inflammation), and histidine (tissue repair) metabolism.

**Conclusion:** High levels of chenodeoxycholic acid (CDCA) represent a poor prognostic indicator in ALF patients. B. intestinalis-AM1, a primary-to-secondary bile acid converter, effectively reduce CDCA levels, activated FXR, reduced inflammation and protected hepatocytes, highlighting its therapeutic potential in ALF.

## INTRODUCTION

Acute liver failure (ALF) is a rapidly progressive, life-threatening condition marked by sudden liver dysfunction in individuals without pre-existing liver disease^1^. It is most often triggered by acute viral hepatitis or drug toxicity, especially acetaminophen overdose^1^. Histologically, ALF involves massive hepatocellular necrosis and mitochondrial dysfunction^2^

Despite advances in the management, ALF remains a condition with high mortality^3^. Interventions such as plasma exchange, renal replacement therapy, and artificial liver support systems provide temporary support and serve as a bridge to transplantation^4^. However, the success of these therapies or liver transplant largely depends on the timely identification of patients who are likely to benefit^4^.

This underscores the urgent need for reliable prognostic indicators to stratify ALF patients at high-risk of early mortality. Early identification of such patients could prompt the use of aggressive medical therapies and guide transplant decisions. One promising approach is to analyze the circulating plasma of ALF patients^2^. Plasma is easily accessible and reflects systemic changes occurring due to liver failure, including alterations in the metabolome and microbiome both of which are crucial components of ALF pathogenesis^2^.

Recent research emphasizes that the interplay between metabolic disturbances and gut-derived microbial products contributes to the hyper-inflammatory state^5^. Changes in the plasma metabolome such as elevated bile acids, amino acids, and oxidative stress markers and circulating bacterial peptides can potentially serve as biomarkers for disease severity and outcome.

Our investigation focused on baseline plasma microbiome (bacterial peptides) and metabolome alterations in order to identify markers predicting early mortality in ALF patients. Multiomics profile of ALF plasma was analysed and significant increase in primary bile acid; Chenodeoxycholic acid was validated in 270 ALF patients using high-resolution mass spectrometry (HRMS) and machine learning. Impaired bile acid metabolism and primary bile acid buildup in ALF indicated a gut microbiota deficiency in converting primary to secondary bile acids. To restore this, we used Bacteroides intestinalis-AM1 (10^9^ cfu/ml), human intestinal isolate, which contributes to the conversion of primary bile acids into secondary bile acids□ and plays a role in FXR activation in various diseases□ in the APAP-induced acute liver failure mouse model. We studied its efficacy in ameliorating liver injury and hepatocyte death and restorative alterations in bile acid composition, plasma proteomics, and metabolomics

## RESULTS

### Plasma metabolome and meta-proteome diversity could stratify ALF non-survivor

Baseline plasma metabolome and meta-proteome analyses^2,6^ were conducted to identify markers predicting early mortality in ALF patients (Supplementary Figures 1 and 2). Untargeted metabolomics of plasma from ALF survivors (n=8), non-survivors (n=32), and healthy controls (n=5) revealed 9,814 features across positive and negative ionization modes, with 548 annotated and compared among groups.

Compared to healthy controls, ALF patients showed 321 differentially expressed metabolites (DEMs; 153 upregulated, 168 downregulated; FC>1.5; p<0.05; FDR<0.05). Additionally, 233 DEMs (114 up, 119 downregulated) distinguished ALF non-survivors (ALF-NS) from survivors (ALF-S; Supplementary Figures 3 and 4). Partial least square discriminant analysis (PLS-DA) and hierarchical clustering distinctly separated ALF-NS from ALF-S and healthy controls (Figure 1A, Supplementary Figures 3 and 4). Metabolic pathways such as tryptophan and histidine metabolism were significantly increased in ALF-NS, while lysine degradation, purine metabolism, and glycerophospholipid metabolism were downregulated (Figure 1B). Key metabolites including chenodeoxycholic acid and L-tyrosine were most influential in distinguishing ALF-NS (Supplementary Figure 5). The top five metabolites showed a high probability of detection (POD) for non-survival with AUROC >80% (Figure 1C), indicating that ALF-NS exhibit a unique metabolic profile associated with inflammation and oxidative stress.

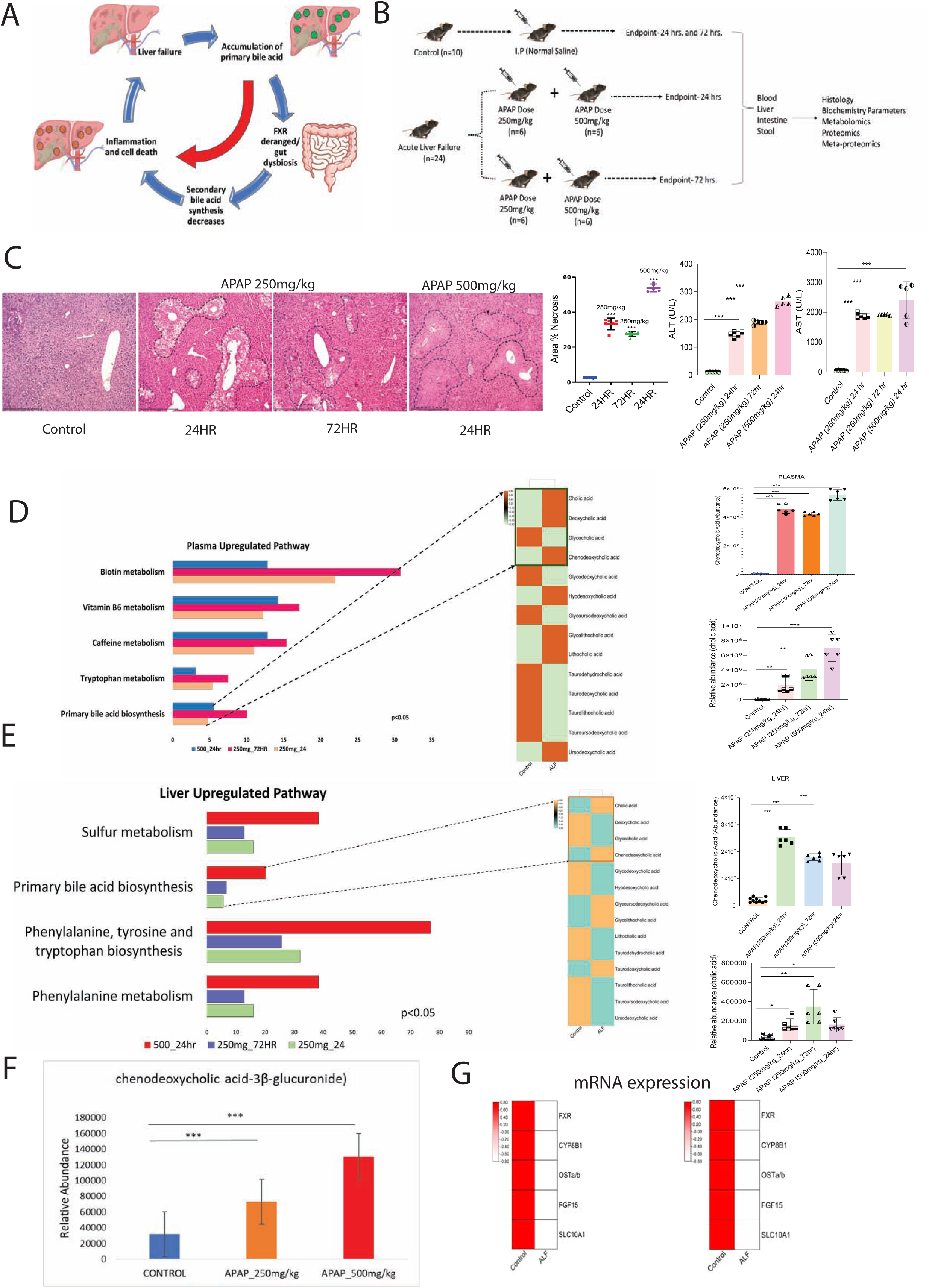
Plasma metabolome and meta-proteome diversity could stratify ALF non-survivor. **Figure 1A-** Partial least square discriminant analysis (PLSDA) showing clear segregation of Healthy (red dot), ALF-S (blue dot) and ALF-NS (green dot) **Figure 1B-** Heatmap and hierarchical clustering analysis capable to segregate Healthy (red), ALF-S (blue) and ALF-NS (green) (Red-upregulated; Green-downregulated; Black-unregulated) **Figure 1C-** AUC value for prediction of ALF-NS (POD) in the training cohort, panel of the five best metabolites (AUC=1 POD > 80%) **Figure 1D-** Alpha-diversity indices (Shannon/Simpson index) and Principal coordinate analysis (beta-diversity) in metaproteome study groups HC, ALF-S, ALF-NS (*p* < 0.05, Kruskal–Wallis test) at class level. **Figure 1E-** Correlation of clinical parameters linked to different meta-proteome phylum’s along with their expression status (Red bar-upregulated; blue-downregulated) **Figure 1F-** Bacterial peptides linked to positively and negatively correlated metabolites along with individual meta-proteome expression (red-positive correlation, green-negative correlation R^2^<0.5 p<0.05) (red bar-upregulated; blue bar-downregulated) **Figure 1G-** AUC value for prediction of ALF-NS (POD) in the training cohort, panel of the five best metabolites (AUC=1 POD > 80%)

Plasma meta-proteome analysis identified >15 phyla and >25 genera dysregulated in ALF-NS, with increased Firmicutes, Cyanobacteria, and Tenericutes (p<0.05; Figure 1D, Supplementary Figures 6-8). Alpha diversity was highest in ALF-NS (p<0.05), and beta diversity analysis distinguished ALF-NS from ALF-S and healthy controls (Figure 1D). Specific bacterial-peptide clusters were identified via linear discriminant analysis and random forest models (Supplementary Figures 6 and 7). Microbial metabolism linked to defence mechanisms, xenobiotic degradation, and amino acid metabolism were increased in ALF-NS (Supplementary Figure 8). Notably, Actinobacteria, Cyanobacteria, and Proteobacteria peptides correlated with liver and kidney function, ALF severity indices, and circulating DEMs (Figures 1E and 1F). The top five meta-proteome markers showed a high POD for non-survival (AUROC >80%; Figure 1G). These findings suggest that gut dysbiosis in ALF-NS influences the circulating metabolome and clinical phenotype, highlighting the interplay between microbiome and systemic metabolism.

### Circulating Chenodeoxycholic acid levels correlate with severity and early mortality in ALF

Integrated analysis of plasma metabolome and meta-proteome using Weighted Metabolite and Meta-protein Correlation Network Analyses (WMCNA/WMpCNA) revealed ALF non-survivor–specific clusters. These unsupervised methods grouped biomolecules into modules based on shared expression patterns.

#### WMCNA

A total of 548 metabolites identified 8 modules (soft threshold>9, scale-free topology fit index>0.85; *p*<0.05; **Figure-2A**). Metabolites in the red, yellow and turquoise module were ALF-Non-Survivor specific modules^7^.

#### WMpCNA

A total of 579 bacterial peptides identified 5 modules (soft threshold>9, scale-free topology fit index>0.85; *p*<0.05; **Figure-2A)**. The module trait relationship analysis identified Mp_Grey, Mp_Turquoise and Mp_yellow modules (**Supplementary Figure-9)** associated with LCA; Streptomyces, Streptococcus pneumoniae and others functionally linked with dysregulated lipid metabolism, amino acids-metabolism and others, specific to ALF-NS^7^ **(Supplementary Figure-10).**

#### Multi-modular Correlation Network Analysis

Mean module intensity of metabolites modules (WMCNA) and the meta-proteome module (WMpCNA) were cross-correlated to obtain multi-modular correlation network (MMCN). The MMCN showed striking association between meta-proteome modules and metabolic pathway (p<0.05; R^2^>0.5**; Figure-2B).** Additionally, linear regression analysis performed between 1) metabolome and meta-proteome cluster with the clinical complication **(Figure-2C)**, 2) Phylum’s of NS specific meta-proteome cluster with the clinical complication **(Supplementary Figure-11)** and 3) NS specific metabolome clusters associated metabolic pathways with the clinical complication **(Figure-2D)** showed significant association of meta-proteome cluster, metabolic pathway and meta-proteome phylum in the prediction of infection at baseline, sepsis, necrosis and multi-organ failure whereas the metabolome module showed striking association with AKI, and hepatic encephalopathy seen in ALF patients (**Figure-2C, Supplementary Figure-11).** Finally, ALF-NS specific meta-proteomic and metabolic modules along with the top 5 metabolites showed direct correlation with the clinical profiles specifically with the severity of ALF **(Figure-2D).** Suggesting that increase in the pathogenic bacteria associated with metabolic pathway could be directly linked to the development of ALF severity.

The POD metabolome showed the highest diagnostic efficiency for predicting non survival as compared to POD Meta-proteome **(AUC 0.986; Fig. 2E).**

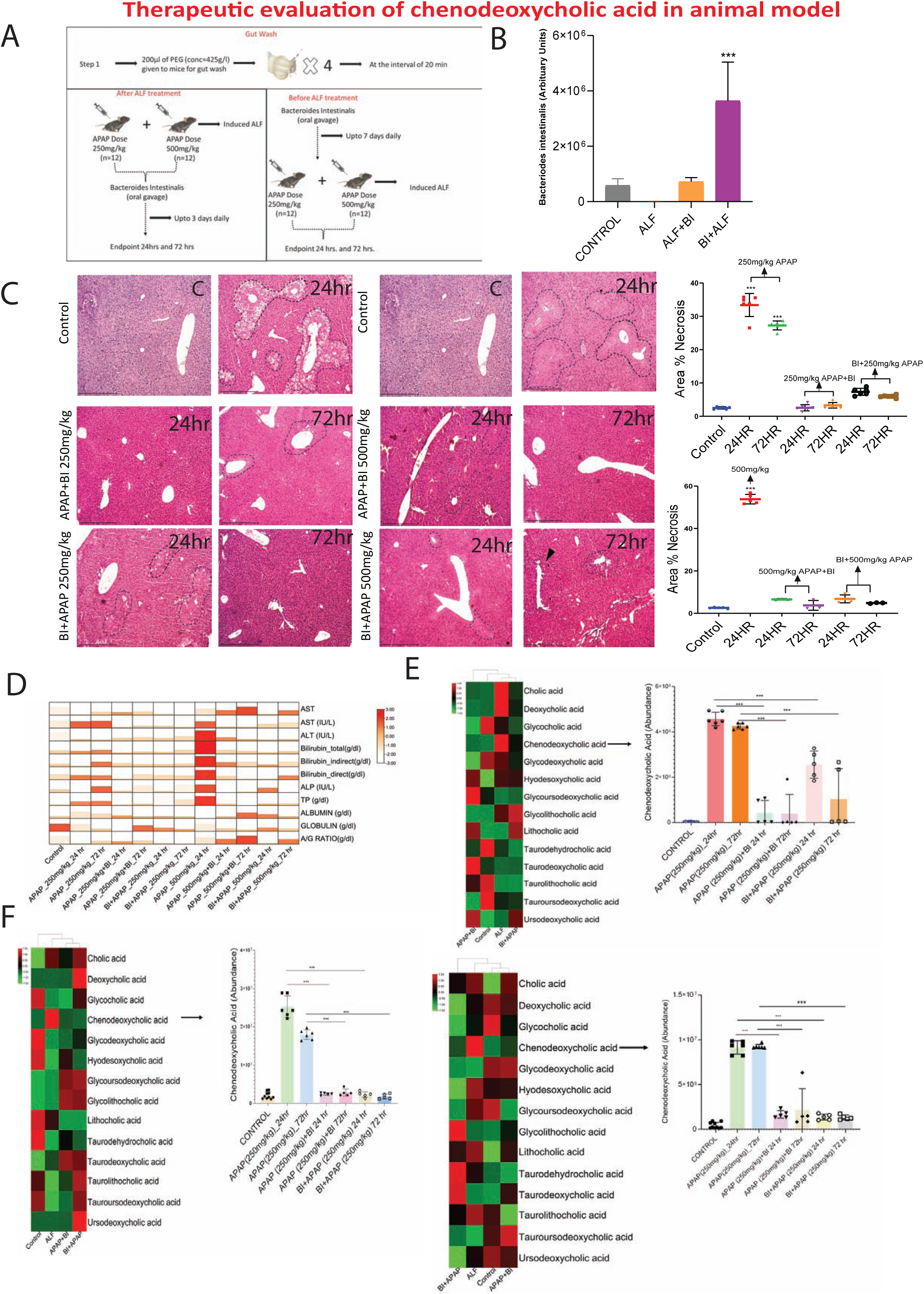
Circulating Chenodeoxycholic acid levels correlates with severity and early mortality in ALF. **Figure 2A-** Weighted Metabolome Correlation Network Analysis Heatmap showing module– trait relationship represented as mean values for each group. The colour scale on the right indicates correlations from −1 (green) to 1 (red) and Weighted Meta-proteome Correlation Network Analysis Heatmap showing module–trait relationship represented as mean values for each group. The colour scale on the right indicates correlations from −1 (green) to 1 (red). **Figure 2B-** Multi-modular Correlation Cluster (MMCC) plot of different metabolome and Meta-proteome modules (Red; R^2^>0.4; p<0.05) **Figure 2C-** Linear regression of the meta-proteome and metabolome module cluster with the clinical-complication of ALF patients; Linear regression of meta-proteome module (Mp yellow and Mp grey) with the clinical complications of ALF-patients; Linear regression of metabolic pathway with the clinical-complications of ALF-patients. **Figure 2D-** Correlation of clinical parameters linked to different meta-proteome phylum’s along with their expression status (Red bar-upregulated; blue-downregulated) **Figure 2E-** Comparison between POD metabolome and POD meta-proteome, AUROC is high for POD metabolome (>0.95; p<0.05) **Figure 2F-** Schematic representation of the study conducted in the validation cohort (TC1-plasma and paired one drop blood; TC2-External-validation; TC3-disease control-SAH) **Figure 2G-** Panel of best 5 metabolite selected based on MDA, FC, and p value and their comparison between the study group (non-survival ALF and survival ALF) in different cohorts (Training, Test cohort 1 (plasma 100µl and one drop blood), Test cohort 2-External Validation. **Figure 2H-** AUROC analysis and Hazard Ratio of MELD score, KCH score and metabolic indicator (C02528, HMDB0028699, C00386, C00082 and C01252) shows best result for C02528 with HR of 1.7 **Figure 2I-** -Assessment of survival and patients on risk based on POD metabolome in ALF patients. (C02528 Chenodeoxycholic acid >log 10FC) **Figure 2J-** Heatmap showing the bile acid profile in ALF-NS, ALF-S and Healthy control, showing increase in primary bile acid

To assess the influence of different etiologies, we compared viral and DILI survivors and non-survivors. There were no significant differences in plasma signatures (metabolome and meta-proteome) across different etiologies, indicating the robustness and specificity of biomolecular signatures in stratifying ALF-NS. (Supplementary-Figures-12, 13 and 14).

Finally, the metabolite indicator panel (L-Tyrosine;4-(2-Amino phenyl)-2,4-90 dioxobutanoate; chenodeoxycholic acid; carnosine; alanyl-tyrosine) was significantly increased in ALF-NS as compared to ALF-S irrespective of etiology, making the identified panel as a pan ALF marker **(Supplementary Figure-14).**

On the basis of FC, MDA, p-value and AUROC values top 5 meta-proteomic LCA and metabolites were selected **(Figure-2G).** AUROC analysis of metabolome panel superseded the meta-proteome panel and thus was chosen for validation.

The metabolome panel was validated in three distinct validation cohorts comprising of TC-1; plasma and one drop blood, for 160 ALF samples (ALF-S;53, ALF-NS;107), TC-2; External Validation cohort; 70 ALF samples (ALF-S;25, ALF-NS;45) and TC-3; Disease cohort; 70 SAH samples **(Figure-2F).** HRMS quantitation and validation showed significant increase in the top 5 metabolite indicators in ALF-NS as compared to ALF-S **(Figure-2G and Supplementary** Figure 14**, 15 and 16).**

Benchmarking of top 5 metabolomics indicator panel against ALF clinical scores showed highest AUROC for the metabolomics indicator panel **(Figure-2H)**.

Next, the identified top 5 metabolite panel: (4-(2-Aminophenyl)-2,4-dioxobutanoate, Carnosine, Chenodeoxycholic acid, Alanyl tyrosine and L-Tyrosine) was subjected to validation using machine learning.

A total of 270 ALF patients were subjected to machine learning analysis using 5 ML algorithms in the CARET package using a fixed allocation of the dataset (80% training set) and then validate using the holdout (20% of the whole data) set^8-10^. The five ML algorithms used and model performance was assessed using kappa score; anything above K=0.7 was considered significant. Overall accuracy of the model was also calculated based on which the best model was selected. Evaluation of the model was performed by assessing its accuracy, sensitivity, specificity, p-value, positive predictive value, negative predictive value and balanced accuracy. In total, 30 trained and tested ML models were generated using 5 ML algorithms across 5 metabolites combination. Fourfold (outer) nested repeated (five times) tenfold (inner) cross-validation (With randomized stratified splitting) was used to train and test ML models and the hyperparameters of each algorithm were optimized. Accuracy and the kappa of model development (training cohort) were significant for all metabolites and comparison for the test cohort (plasma and paired one drop blood). Positive predictive value, negative predictive value and balanced accuracy of plasma (100µl) were found to be highest for RF model. Prediction capability of the identified top 5 metabolites in combination showed highest accuracy, sensitivity, specificity and p-value making the identified panel candidate indicators and the random forest model as the preferred ML algorithm for segregation of ALF patients predisposed to early mortality using plasma samples (**Supplementary** Figure 14**, 15 and 16).**

Interestingly, AUROC analysis along with the COX univariate and multivariate regression analysis showed significant association of Chenodeoxycholic acid (C02528; HR=1.7; p<0.05) with early mortality in ALF patients **(Figure-2I and Supplementary Figure-17)**. Acute liver failure patients with more than 10 FC plasma value for Chenodeoxycholic acid showed significant lower survival **(Figure-2I)** and increase in primary bile acid was validated (bile acid analysis) and was higher in ALF-NS **(Figure-2I).**

### Increase of mortality is associated to increase in chenodeoxycholic acid in pre-clinical mouse model of Acute liver failure

To investigate whether the accumulation of primary bile acids also contributes to APAP-induced hepatotoxicity, leading to inflammation and cell death (Figure-3A), we developed a mouse model of APAP-induced acute liver failure (ALF). Mice were administered two doses (250 mg/kg and 500 mg/kg) and sacrificed at 24 and 72 hours (Figure-3B). Liver injury was assessed by measuring serum ALT and AST levels and scoring necrosis in H&E-stained paraffin sections (Figure-3C). A significant difference in necrosis, AST, and ALT levels was observed between the control and APAP-treated mice (250 mg/kg and 500 mg/kg).

Plasma metabolome analysis showed significant increase in primary bile acid metabolism, tryptophan metabolism, and other pathways in preclinical model of ALF. Total bile acid profiling showed significant increase in primary bile acids, particularly chenodeoxycholic acid and cholic acid, in APAP-treated mice across all study groups (Figure-3D). Similar results were observed in liver metabolome analysis, showing elevated levels of primary bile acids, specifically chenodeoxycholic acid and cholic acid (Figure-3E).

Increase in Chenodeoxycholic acid is known to produce CDCA-3β-glucuronide^11^ and results in deactivation of FXR indicating impaired bile acid transport. Expression of CDCA-3β-glucuronide was higher in preclinical model of ALF (Figure 3F).

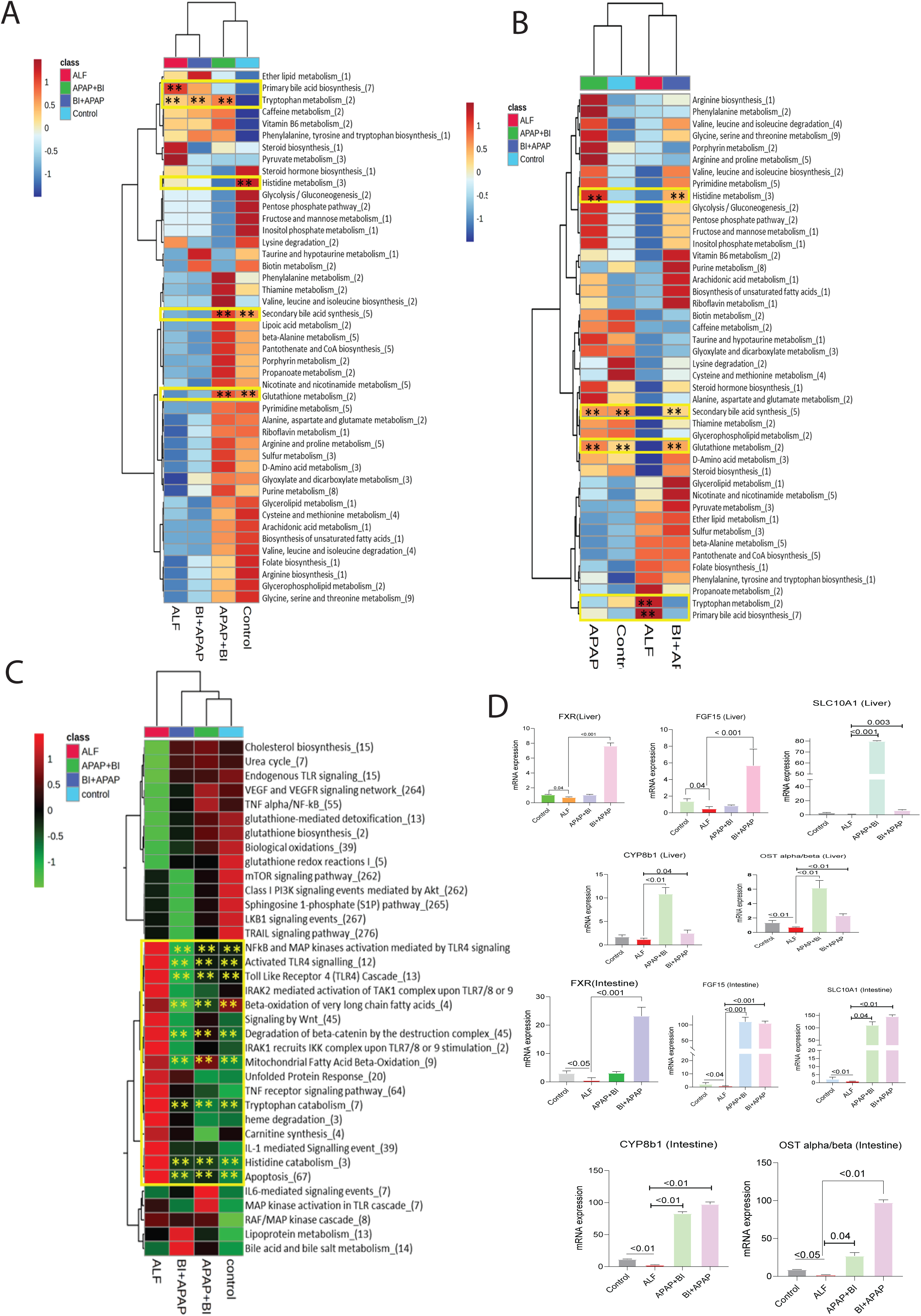
Increase of mortality is associated to increase in chenodeoxycholic acid in mice model of APAP induced ALF. **Figure 3A-** Hypothesis-Acute liver failure---primary bile acid accumulation---gut dysbiosis and FXR deranged---decrease in secondary bile acid---inflammation and cell death. **Figure 3B-** Liver injury after N-acetyl-p-aminophenol (APAP) overdose. Male C57BL/6 mice (10 to 11 weeks old) received two intraperitoneal injection of 250□mg APAP/kg body and 500mg/kg weight after a fasting period of 12□h **Figure 3C-** Livers from control mice (ctr, treated with NaCl 0.9%) and APAP-treated mice stained with haematoxylin and eosin (H&E) show progressing necrotic areas (lighter), 24□h and 72□h after APAP overdose. Scale bar: 500□µm; The liver damage is also reflected by the levels of transaminase activities (aspartate aminotransferase [AST] and alanine aminotransferase [ALT]) in the serum. **Figure 3D-** Plasma metabolome analysis revealed increase in primary bile acid metabolism specifically chenodeoxycholic acid and cholic acid in APAP induced mice (p<0.05) **Figure 3E-** Liver analysis revealed increase in primary bile acid metabolism specifically chenodeoxycholic acid and cholic acid in APAP induced mice (p<0.05) **Figure 3F-** Relative abundance of chenodeoxycholic acid beta glucuronide in APAP 250mg/kg and APAP 500mg/kg (p<0.05) **Figure 3G-** Relative mRNA Expression of bile transporter gene between control and APAP induce mice group/ALF p<0.05

Further, gene expression analysis of the gut and liver for various bile acid transport-related genes showed significant downregulation of key bile acid transporter genes (fxr, fgf15, SLC10A1, and others) in both the liver and intestine, indicating impaired bile acid transport and subsequent accumulation of primary bile acids.

### Bacteroides Intestinalis AM-1 therapy ameliorates liver injury, necrosis, and inflammation in preclinical model of ALF

To further investigate the potential of gut modulation in mice for improving liver failure and reducing primary bile acid accumulation, we examined the effects of *Bacteroides intestinalis* AM-1 treatment. This human intestinal isolate contributes to the conversion of primary bile acids into secondary bile acids□ and plays a role in FXR activation in various diseases□. *Bacteroides intestinalis* AM-1 was cultured in an anaerobic medium and a suspension containing 1 × 10□ CFU/mL colonies was administered to the mice (Supplementary Figure 18). In brief, *Bacteroides intestinalis* AM-1 was given following gut wash, in mice as stated below

1. Before APAP injection (7 days-2 dose per day for gut colonization)
2. After APAP injection (3 days-2 dose per day) to mitigate ALF (Figure 4A) and both the groups were compared for alleviation of liver injury, necrosis, and inflammation in preclinical model of ALF

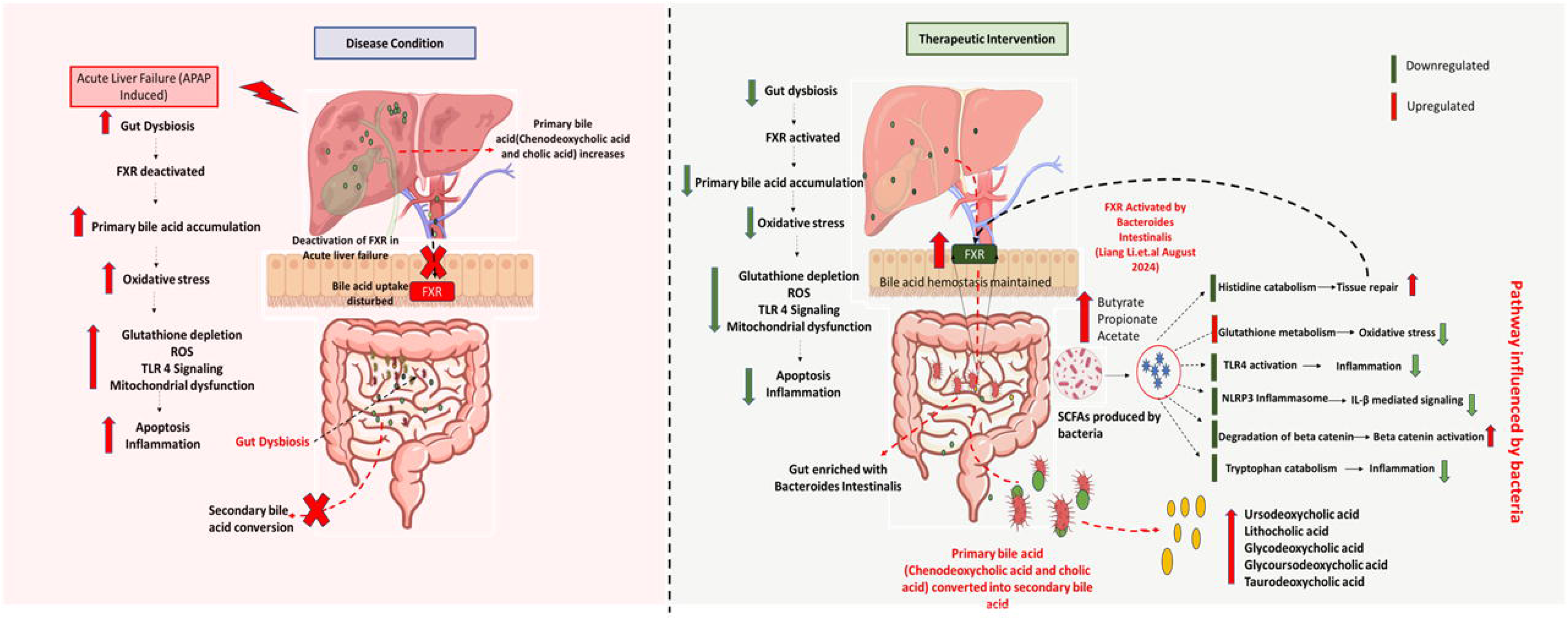
Bacteroides Intestinalis AM-1 therapy in APAP induced mice reverse ALF in mice. **Figure 4A-** Schematic representation of the study conducted in the mice for therapeutic evaluation of the Bacteroides intestinalis AM-1. **Figure 4B-** Relative abundance of Bacteroides intestinalis AM-1 in control group, APAP group/ALF, APAP+BI and BI+APAP, showing abundance of bacteria in the stool samples. **Figure 4C-** -Livers from control mice (ctr, treated with NaCl 0.9%) and APAP-treated mice stained and treatment group with haematoxylin and eosin (H&E) show progressing necrotic areas (lighter), 24□h and 72□h after APAP overdose and treatment. Scale bar: 500□µm; **Figure 4D-** The liver recovery is also reflected by the levels of transaminase activities (aspartate aminotransferase [AST] and alanine aminotransferase [ALT]) in the serum in treatment group **Figure 4E-** Relative Abundance heat map showing increase bile acid profile in plasma and in different study groups (Control; APAP/ALF, BI+APAP, APAP+BI) (red; upregulated, green; downregulated) and relative abundance of chenodeoxycholic acid. **Figure 4F-** Relative Abundance heat map showing increase bile acid profile in liver and in different study groups (Control; APAP/ALF, BI+APAP, APAP+BI) (red; upregulated, green; downregulated) and relative abundance of chenodeoxycholic acid.; Relative Abundance heat map showing increase bile acid profile in intestine and in different study groups (Control; APAP/ALF, BI+APAP, APAP+BI) (red; upregulated, green; downregulated) and relative abundance of chenodeoxycholic acid.

To confirm colonization in the mice’s gut, we analysed the metaproteome of their stool samples and detected the bacterium in both groups (Figure 4B). Mice receiving *Bacteroides intestinalis* AM-1 prior to APAP injection showed better results but in both the groups *Bacteroides intestinalis* AM-1 was able to reduce primary bile acid accumulation, reduced liver injury, necrosis, and inflammation (Figure 4C). We also observed significant reduction of AST, ALT, and other liver function markers (Figure 4D). Interestingly, *Bacteroides intestinalis* AM-1 showed significant reduction in chenodeoxycholic acid levels in the plasma, liver, and intestine of ALF model (Figures 4D, 4E, and 4F).

Together these results highlight the utility of *Bacteroides intestinalis AM-1* in mitigation of ALF in a preclinical model of acute liver failure.

### *Bacteroides intestinalis* AM-1 colonization modulates molecular pathways linked to liver injury and inflammation pre-clinical model of ALF

To assess the effects of bacteria targeting primary bile acids, we analyzed plasma and liver metabolomes along with liver proteomics to identify factors driving liver damage reversal across all mouse groups.

Plasma and liver metabolome analysis highlighted key metabolic changes associated with reduced inflammation and oxidative stress, enhanced glutathione metabolism and antioxidant defence respectively.

Significant reduction of primary bile acid accumulation alleviates liver stress and inflammatory responses. These metabolic shifts suggest that modulating gut microbiota and bile acid metabolism may contribute to liver protection. (Figure-5A and 5B and Supplementary Figures-19, 20, 21, 22, 24 and 24).

Liver proteomics analysis further highlights significant reduction in NF-κB and MAP kinase pathways, TLR4 signalling, histidine catabolism and mitochondrial beta-oxidation suggesting a metabolic shift that alleviates inflammation, apoptosis, oxidative stress and cellular damage. Together these findings emphasize the potential of gut microbiota modulation in influencing key signalling pathways and metabolic processes that contribute to liver protection (Figure-5C and Supplementary Figure-25, 26 and 27).

As Bacteroides intestinalis AM-1 modulates bile acid metabolism, the genes linked to bile acid homeostasis including *FXR*, *FGF15*, *SLC10A1*, and others, were significantly increased in the liver and intestine of bacteria-treated mice further suggesting the recalibration of bile acid metabolism, which may contribute to reduced bile acid accumulation and improved liver function (Figure-5D and Supplementary Figure-29). Together these findings suggest that Bacteroides intestinalis AM-1 treatment exerts a protective effect in preclinical model of ALF respectively. Our result paves a way for modulation of gut microbiome for treatment of ALF this novel approach warrants human validation

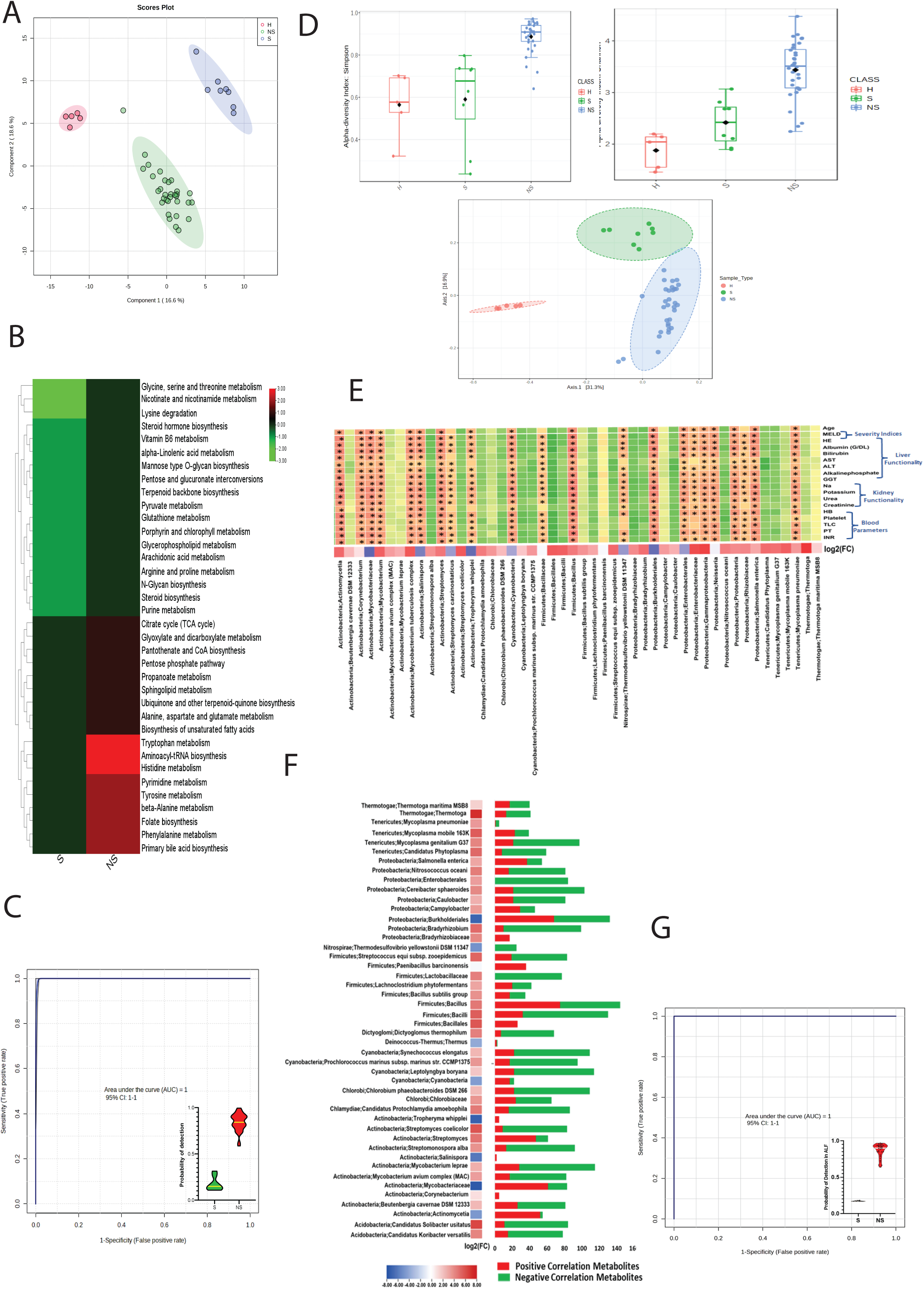
*Bacteroides intestinalis* AM-1 colonization modulates various pathways to alleviate APAP-induced injury in a mouse model. **Figure 5A-** Unsupervised hierarchical clustering with different group showing plasma metabolome pathway of liver is capable to segregate Treatment and ALF (250_24hrs) and control group in mice model **Figure 5B-** Unsupervised hierarchical clustering with different group showing liver metabolome pathway of liver is capable to segregate Treatment and ALF (250_24hrs) and control group in mice model **Figure 5C-** Unsupervised hierarchical clustering with different group showing liver proteome pathway of liver is capable to segregate Treatment and ALF (250_24hrs) and control group in mice model **Figure 5D-** Relative mRNA Expression of bile transporter gene between control and APAP induce and treatment group mice group/ALF p<0.05

**Figure 6A.**
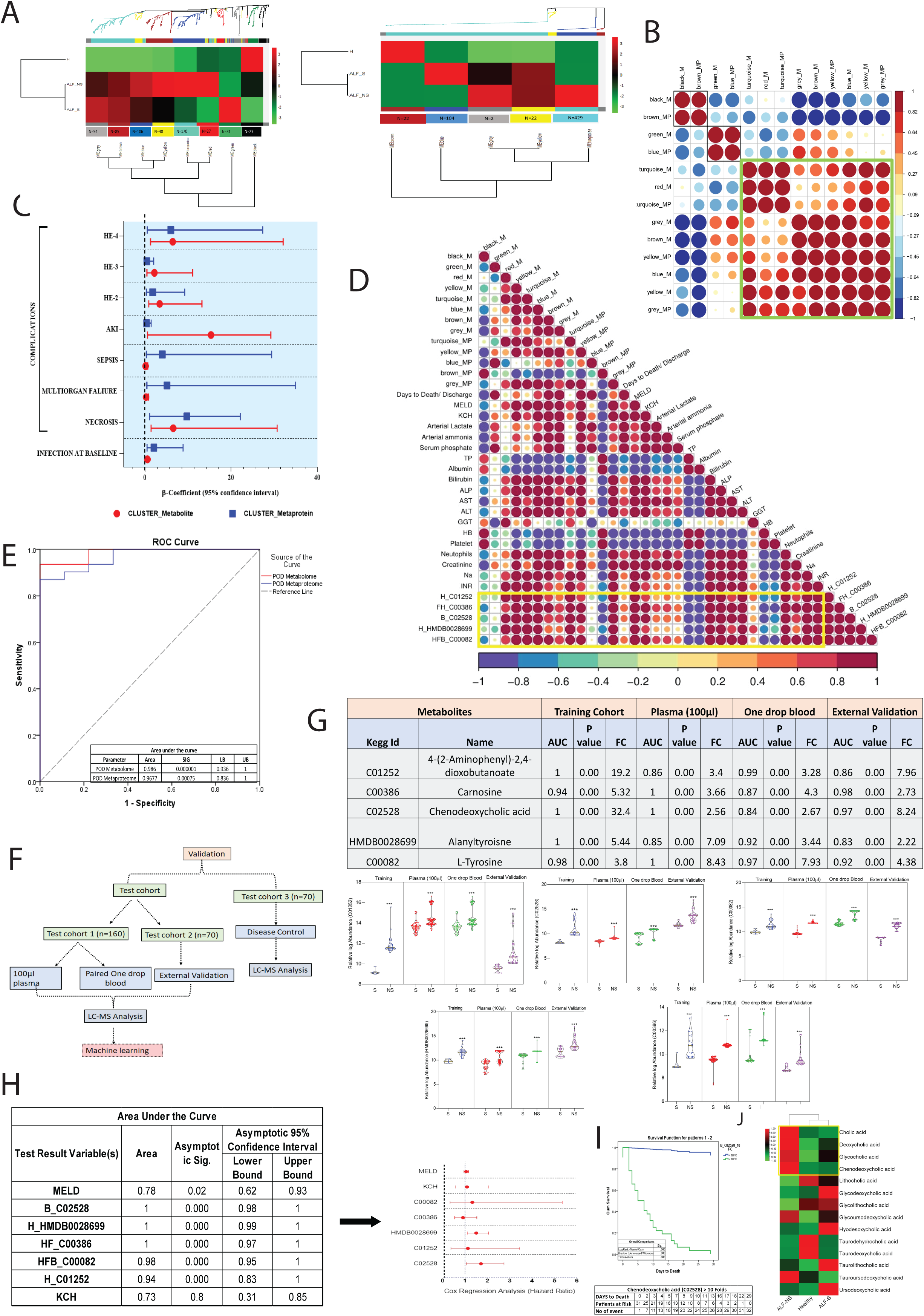
Graphical Abstract. In this pilot study, we identified circulating chenodeoxycholic acid, a hydrophobic bile acid, as a highly hazardous metabolite in ALF patients, particularly those at risk of early mortality. Our findings suggest that elevated levels of chenodeoxycholic acid may contribute to hyperinflammation and oxidative stress in the context of ALF. To address this, we targeted the gut microbiota using Bacteroides intestinalis AM-1 in mice, a bacterium known to convert primary bile acids into less toxic secondary bile acids. This intervention not only facilitated bile acid conversion but also activated FXR signaling, which is essential for maintaining bile acid homeostasis. As a result, we observed a marked reduction in oxidative stress, inflammation, and hepatocellular damage, highlighting the therapeutic potential of microbial modulation in mitigating liver injury in ALF.

## DISCUSSION

This study investigated the plasma metabolome and bacterial peptides in ALF patients to identify potential diagnostic markers. We also evaluated *Bacteroides intestinalis AM-1* as a novel therapeutic intervention in a preclinical mouse model. A panel of metabolites and microbial peptides were identified, with the metabolite panel demonstrating superior diagnostic accuracy, particularly in predicting early mortality. Notably, primary bile acids, especially chenodeoxycholic acid, were significantly elevated in non-survivors. ALF patients also showed substantial gut microbiota dysbiosis, which was associated with increased inflammation and impaired detoxification. These findings suggest that accumulation of primary bile acids, along with gut microbial imbalance, may contribute to inflammation and necro-apoptosis in ALF. In the mouse model, elevated chenodeoxycholic acid levels were validated, supporting the relevance of treatment with *B. intestinalis AM-1*, known for converting primary to secondary bile acids. *B. intestinalis* effectively modulated the FXR axis, reduced necro-apoptotic and inflammatory responses, enhanced glutathione activity, and improved bile acid homeostasis. These results highlight the therapeutic potential of *B. intestinalis AM-1* in modulating bile acid metabolism and gut-liver axis inflammation in ALF.

Plasma analysis of ALF particularly ALF non-survivors showed higher levels of chenodeoxycholic acid and cholic acid in the circulation and lower levels of secondary bile acids and associated bacteria which convert primary to secondary bile acid. Disrupted amino acid metabolism was observed, including decreased branched-chain and increased aromatic amino acids indicators of oxidative stress and impaired protein synthesis ^12,13^ in ALF non-survivor. Elevated tryptophan and activation of the kynurenine pathway correlated with immune dysfunction^14-16^. Increased bile acid synthesis, particularly chenodeoxycholic acid, was linked to early mortality, likely due to gut microbiota imbalance impairing detoxification^17^. Higher carnosine levels suggested a neuroprotective response to ammonia-induced stress^18,19^. Moreover, ALF non-survivors exhibited elevated microbial diversity and circulating bacterial peptides, indicating gut dysbiosis and compromised intestinal integrity^5,20^. These findings suggest the importance of plasma metabolomic and microbial signatures for early prognostication and highlight their role in ALF progression.

A strong correlation between plasma metabolome and meta-proteome was observed in ALF non-survivors, as revealed by MMCC network analysis. Module-trait relationships identified key metabolic pathways tryptophan, bile acid, and tyrosine metabolism linked to specific microbial modules, especially the turquoise module, highlighting microbial influence in ALF pathogenesis. Tryptophan and histidine metabolism, modulated by gut microbiota, were notably associated with inflammation and immune responses. The turquoise meta-proteome module, enriched in Actinobacteria, Proteobacteria, and Firmicutes, corresponded to elevated aromatic amino acids, indicative of disease severity. These microbial shifts reflect gut dysbiosis and increased intestinal permeability, promoting systemic inflammation via altered bile acid metabolism^21^. The meta-proteome diversity predicted infection and multi-organ failure, while metabolome changes associated with AKI and hepatic encephalopathy.

Comparative analysis against SAH (diseases controls) and across ALF etiologies showed that microbial and metabolic signatures were independent of disease origin, making them potential universal predictors.

Five plasma metabolites, including chenodeoxycholic acid, were validated across three distinct validation cohorts, with random forest models achieving high predictive accuracy for early mortality. This pilot study proposes a robust, mass spectrometry-based plasma metabolite panel for early ALF risk stratification, offering significant clinical potential.

Our study demonstrates that the accumulation of primary bile acids, particularly chenodeoxycholic acid (CDCA) and cholic acid, plays a significant role in acetaminophen (APAP)-induced hepatotoxicity, contributing to inflammation and hepatocellular death in preclinical mouse model of acute liver failure (ALF). The dose-dependent elevation of serum ALT and AST levels, along with histological evidence of necrosis, confirmed liver injury in APAP-treated groups.

Integrated metabolomic analyses of plasma and liver tissues revealed consistent upregulation of primary bile acid metabolism, along with perturbations in tryptophan metabolism suggestive of inflammation and systemic metabolic dysregulation during ALF.

A critical finding was the marked increase in CDCA and its metabolite CDCA-3β-glucuronide, which is known to inhibit farnesoid X receptor (FXR) signaling^11^. FXR is a key nuclear receptor that regulates bile acid homeostasis and protects against bile acid-mediated liver injury^22^. Its deactivation likely exacerbates bile acid toxicity, contributing to hepatocellular damage and inflammation. This was further supported by significant downregulation of FXR and other bile acid transport genes (e.g., FGF15, SLC10A1) in both hepatic and intestinal tissues of preclinical mice model, indicating impaired enterohepatic circulation and bile acid clearance^22^. These results suggest that bile acid accumulation and transport dysfunction are central to the pathogenesis of APAP-induced liver injury.

Targeting bile acid metabolism and FXR signalling could represent a promising therapeutic strategy for mitigating ALF progression^23^. Future studies were carried out which showed that modulation of bile acid profiles or enhancement of FXR activity can improve outcomes in APAP-induced hepatotoxicity.

This was achieved by using *Bacteroides intestinalis AM-1*, a bacterium involved in primary bile acid metabolism, and observed a significant reversal of liver damage in preclinical model of APAP-induced acute liver failure (ALF)^24^. Metabolomic profiling of plasma and liver supported the beneficial effect of *Bacteroides intestinalis* AM-1 and showed reduced inflammation and oxidative stress.

Interestingly, *Bacteroides intestinalis* AM-1 enhanced glutathione metabolism supports detoxification, while increased secondary bile acid synthesis helps restore bile acid homeostasis^25^.

Additionally, we observed concurrent reduction in primary bile acid accumulation, particularly hepatotoxic species like chenodeoxycholic acid, and alleviated bile acid-induced liver stress. These findings suggest that gut microbial modulation directly impacts liver resilience through metabolic reprogramming.

Proteomic analysis further corroborated these protective effects of *Bacteroides intestinalis* AM-1, showing reduced activation of pro-inflammatory NF-κB and MAPK signaling pathways, along with suppression of TLR4-mediated immune activation^26^. These changes indicate a dampened inflammatory response and reduced apoptosis in the liver. Additionally, the downregulation of histidine catabolism and mitochondrial beta-oxidation points to decreased oxidative stress and mitochondrial burden, further supporting hepatoprotection^27,28^.

Importantly, upregulation of bile acid transport and regulatory genes (FXR, FGF15, SLC10A1) in both liver and intestine suggests restoration of bile acid signalling and improved enterohepatic circulation^29^. These findings highlight the therapeutic potential of modulating gut microbiota to influence host metabolic and immune pathways, offering a promising strategy for mitigating liver injury and promoting recovery in preclinical model of ALF.

In conclusion, bacterial gut modulation has predominately been studied in patients with alcoholic hepatitis, COPD, and the other organ system^29^. The results of our study strongly indicate that bacterial treatment could mitigate liver injury and could be evaluated in ALF patients ^30,31^. However, safety studies would be required in patients with HAV and HEV induced ALF, as bacteria can induce strong immune reactions^32^. Further work is required to determine whether use of *Bacteroides intestinalis* AM-1targeting to reduce primary bile acid accumulation might be used to treat patients with ALF. Our data also suggest that high levels of chenodeoxycholic acid may be used as a predictive indicator for segregation of ALF patients predisposed to early mortality.

## METHODS

### Patient cohorts

We evaluated, a total of 270 ALF-patients along with 25 healthy-controls prospectively seen (December 2019 to April 2023) at the Institute of Liver and Biliary Sciences, New Delhi, India. Etiological distribution of 270 ALF patients; 15% -HEV (Hepatitis E Virus), 10%-DILI (Drug Induced Liver Injury), 65% -HAV (Hepatitis A Virus), 10%-Others (indeterminate and others). ALF was diagnosed as; presence of jaundice with hepatic encephalopathy within 4 weeks with laboratory evidence of increased INR (>1.5)^1^. Patients with underlying features of chronic liver disease, such as splenomegaly, clinical ascites or known hepatitis B or C infection or regular alcohol intake, were excluded from the study. All patients underwent plain CT scan of the brain to screen the presence and severity of cerebral-edema in the emergency before being shifted to the liver intensive care unit (L-ICU). All the ALF patients with presence of hepatic encephalopathy (Grade 1-4) were enrolled in this study and were sampled at the time of admission (baseline). The patients were managed by intubation and ventilation according to their standard indications. Fentanyl and propofol was used along with atracurium for paralysis wherever required during ventilation. Volume revival need was determined by the clinical parameters like heart rate, blood pressure, urine output, arterial blood gas and stroke volume variation^33^. Patients who do not respond initially were managed by norepinephrine with the parallel use of low dose of hydrocortisone and vasopressin. Intravenous N-acetylcysteine was given to all the patients for 5 days. Patients were administered with antibiotics and antifungals as per institutional protocols. Continuous renal replacement therapy (CRRT) was performed in patients with suspected cerebral edema on CRRT ^34^. Liver transplantation was available both as living related or availability of deceased donor liver status. Further, patients were clinically followed up for 90 days or death from the date of admission. If a patient died due to progressive liver failure or liver related complications, such as cerebral edema, renal failure, infection, sepsis or multi-organ failure during the ICU stay or within 30 days it was considered as death due to liver failure (ALF-Non-Survivor; ALF-NS). The patients who improved and survived were categorized as ALF survivors (ALF-S). All procedures involved in the study were conducted by the institute ethical committees (IRB approval no-IEC/2019/70/NA06) and written informed consent was obtained from all subjects enrolled in this study.

### Sample collection

A total of 10 mL of blood was drawn from ALF patients using a sterile syringe at the time of enrollment, adhering to the predefined inclusion and exclusion criteria. Additionally, a single drop (∼35 µL) of blood was obtained from the index finger of each <u>patient.</u>

#### Processing of One-Drop Blood Samples

The collected one-drop blood samples (∼35 µL) were subjected to centrifugation at 2,000 rpm for 20 minutes at 4°C to facilitate plasma isolation. The resulting plasma fraction (∼15–17 µL) was carefully extracted and processed for metabolomic analysis, as outlined in our recent publication^2^.

#### Plasma depletion

Blood (10ml) samples were collected at baseline and subjected to centrifuge for 20 minutes at 2,000 rpm at 4 °C for plasma isolation. Further plasma samples were first subjected to plasma depletion followed by preparation for metabolomics and meta-proteomics) using mass spectrometry^6,35,36^.

#### Separation of Low Abundant Protein (LAP) and Albuminous fraction

Albumin was removed from plasma samples of the study cohort of ALF Patients (n=40; ALF Non-Survivors; ALF-NS=32, ALF-survivors; ALF-S=8) and Healthy Control (HC) (n=5) using a **dye-binding affinity chromatographic technique** which employs the use of HiTrap Blue HP 1ml (Cat no. 17-041201, GE Healthcare Life Sciences, Sweden) on AKTA pure purification system which is prepacked with **Blue Sepharose beads immobilized by**

### Cibacron Blue F3G-A dye in which

Albumin attaches to the Cibacron blue dye due to its albumin-dye binding property. The high trap blue column (Cibacron blue dye) was used as it has a specific binding with albumin^37-40^. Baseline 100µl plasma was diluted 1:4 times using Binding buffer (20mM Na_2_HPO_4_) followed by loading onto a 1 ml HiTrap Blue column in which the sample was loaded for a brief period of 2 min followed by washing of the column using 20mM Na_2_HPO_4_ for 5 min in which we observe a sharp peak of the unbounded low abundant proteins (LAP), this is followed by elution phase of 7 min with an increase in the percentage (5-98%) of 2M NaCl + 20mM Na_2_HPO_4_ which results in the elution of the dye bounded albumin as mentioned in the previous paper for SAH^41^.

### Sample Preparation for Metabolomics

For metabolomic analysis, an organic phase extraction protocol was employed to isolate metabolites from 100 µL of low-abundant protein (LAP) samples from both the training cohort (ALF = 40, HC = 5) and test cohort (ALF = 160, HC = 20), using 100 µL of plasma and paired one-drop blood (∼16 µL plasma). Sample preparation involved a methanol-to-sample ratio of 4:1, with 100 µL of LAP from the training cohort and 100 µL of baseline plasma with paired one-drop blood from the test cohort incubated overnight at −20°C to facilitate protein precipitation. Samples were then centrifuged at 13,000 rpm for 10 minutes, and the resulting protein pellet was discarded while the supernatant was collected and vacuum-dried. The dried samples were reconstituted in 100 µL of a standardized solution (90% water, 5% acetonitrile, 5% internal and external standards) to ensure optimal metabolite stability. A total of 80 µL of each prepared sample was transferred into a high-performance liquid chromatography (HPLC) tube, while the remaining 25 µL was pooled across all samples to create a quality control sample. This pooled sample was analyzed at serial dilutions (1:1, 1:2, 1:4, and 1:8) to assess analytical reproducibility. Internal standards included Dinoseb (1 mg/mL), MCPA (1 mg/mL), and Dimetrazole (1 mg/mL), whereas external standards comprised Cholesterol (0.1 mg/mL), Colchicine (4 mg/mL), Imparinine (2 mg/mL), Roxithromycin (2 mg/mL), Amiloride (1 mg/mL), and Atropine (2 mg/mL).

### For mass spectrometry-based metabolomics

15 µL of each prepared sample was subjected to reverse-phase chromatography on a C18 column (Thermo Scientific™ 25003102130; 3 µm, 2.1 mm, 100 mm) using an ultra-high-performance liquid chromatography (UHPLC) system. The mobile phase consisted of 0.1% formic acid (Phase A) and 100% acetonitrile (Phase B). The samples were introduced into the heated electrospray ionization (HESI) source of a Q-Exactive mass spectrometer (Thermo Scientific, San Jose, CA) and analyzed. The spray voltage was set to +3.7 kV in positive ionization mode and −3.1 kV in negative ionization mode. The heated capillary temperature was maintained at 360°C, with sheath and auxiliary gas flow rates set to 15 and 10 arbitrary units, respectively.

Metabolite identification was conducted using Compound Discoverer 3.0 (Thermo Fisher Scientific, Waltham, United States), with annotation performed through multiple databases, including mass list databases, ChemSpider, mzVault, Metabolika, and mzCloud™. The identified and annotated metabolite features underwent log normalization and Pareto scaling using MetaboAnalyst 5.0 (http://metaboanalyst.ca), which facilitated subsequent analyses such as principal component analysis (PCA), partial least squares discriminant analysis (PLS-DA), heat map generation, and random forest classification. Additionally, pathway enrichment analysis was carried out using MetaboAnalyst 5.0, while IBM SPSS Statistics version 2.0 was employed for multi-variate correlation and univariate regression analyses, as detailed in our recent publication.

### Sample Preparation for Meta-proteomics

Following albumin depletion from baseline plasma, low-abundant protein (LAP) samples from all study groups, including acute liver failure (ALF) patients (n=40), ALF non-survivors (ALF-NS, n=32), ALF survivors (ALF-S, n=8), and healthy controls (HC, n=5), underwent protein quantification using the Bradford assay. For in-solution protein digestion, 50 µg of protein was vacuum-dried and reconstituted in 200 µL of ammonium bicarbonate (ABC) buffer. To reduce disulfide bonds, 20 µL of 10 mM dithiothreitol (DTT) was added, and the mixture was incubated at 60°C for 1 hour in a water bath. Following reduction, 15 µL of 10 mM iodoacetamide (IAA) was introduced for protein alkylation, with incubation in the dark for 30 minutes. Protein digestion was carried out using 10 µL of sequencing-grade trypsin (0.1 µg/µL), with samples incubated at 37°C for 20–24 hours. Trypsin activity was subsequently inhibited by adding 5 µL of concentrated formic acid.

For C18 column-mediated peptide purification, the digested LAP was purified using the Thermo Scientific Pierce C18 Spin column (Catalog No. 89873). Initially, the column was preconditioned with 200 µL of 100% acetonitrile (ACN) for 5 minutes, followed by centrifugation for 2 minutes, and the flowthrough was discarded. To activate the column, 200 µL of wash buffer (50% ACN + 50% water) was added, incubated for 5 minutes, and then centrifuged, with the flowthrough removed. Equilibration was performed by adding 200 µL of equilibration buffer, incubating for 2 minutes, and discarding the flowthrough. The digested protein sample (50 µg) was then loaded onto the column and incubated for 5 minutes, with repeated loading for a total of four cycles to enhance peptide binding. The flowthrough was discarded, and the column was washed with 200 µL of equilibration buffer (5% ACN + 95% water) for 2 minutes, followed by centrifugation and removal of the flowthrough. Peptide elution was carried out in three sequential steps using an elution buffer (90% ACN + 10% water): first, 30 µL was incubated for 2 minutes, followed by another 30 µL for 2 minutes, and finally, 40 µL was incubated and centrifuged. The eluate was collected, and the used column was discarded. The samples were lyophilized at 4°C under 60 millibar pressure until dry, ensuring they were not over-dried to prevent difficulties in reconstitution. The dried peptides were then reconstituted in 40 µL of 0.1% (v/v) formic acid, centrifuged at 15,000g, and 20 µL of the supernatant was loaded into an HPLC vial.

For mass spectrometry-based meta-proteomics analysis, proteins were isolated from plasma samples across all study groups. These proteins were subjected to reduction, alkylation, and trypsin digestion, followed by analysis via mass spectrometry as described in the proteome analysis section. MS/MS data were acquired and analyzed using Proteome Discoverer (version 2.0, Thermo Fisher Scientific, Waltham, MA, United States), employing bacterial and fungal sequences from UniprotSwP_20170609 (sequences: 467231) and MG_BG_UPSP (sequences: 2019194). The Mascot algorithm (Mascot 2.4, Matrix Science) was used for cross-validation of microbial species identification. Peptide identification was considered significant at p < 0.05, with q-values p < 0.05 and a false discovery rate (FDR) of 0.01. Only rank-1 peptides with peptide-spectrum matches (PSMs) greater than 3 were included for biodiversity and functional analyses using Unipept. Peptides that mapped to eukaryotic, fungal, and viral databases were excluded, ensuring that only bacterial species-associated peptides were retained for statistical, functional, and biodiversity analyses, as detailed in our recent publications.

**Weighted Metabolome Co-Expression Network (WMCNA) and Weighted Metabolome Co-Expression Network (WMpCNA);** was also performed upon identified Metabolites using ‘WGCNA’ R package in Perseus software. In this, the correlation matrix between the metabolites or Meta-proteins in the dataset describes a fully connected, weighted network, in which the weight on each edge denotes the correlation between the quantitative profiles of the two metabolites or Meta-proteins^42^. Soft-threshold=9, signed network and Bi-weight mid-correlation, a robust alternative to Pearson correlation, were chosen to calculate correlations between all Metabolite signatures. In order to obtain a scale-free co-expression network, a power parameter of 12 was selected, leading to an approximately scale-free network with a scale-free topology of 0.85. Hierarchical clustering of the co-expression network identified 8 and 5 modules of identified metabolites and meta-proteins respectively.

#### Probability of death (POD) calculation

In the current study Probability of death (POD) for Acute liver failure patients (ALF) was assessed to determine the putative indicator of non-survival in ALF patients. POD was calculated using the Top meta-proteins and metabolites identified based on p-value, AUC, and mean decrease in accuracy in each comparison (ALF VS HC and ALF-Non survivor VS ALF-Survivor) using IBM SPSS 20.

#### Machine Learning

approach^9^ was employed using five different ML (LDA, RF, SVM, KNN, CART) algorithms to validate the plausible biomarker metabolomic species. In total, we implemented 30 ML models comprising 5 ML algorithms along with 5 metabolomic species individually as well as combinedly. Fourfold (outer) nested repeated (five times) tenfold (inner) cross-validation (with randomized stratified splitting) was done on training and test cohort in R with the Caret package. In this way, repeated tenfold cross-validation was performed 20 times and the models obtained the best results. Besides, the overall cross-validation prediction performance was summarized by the accuracy, sensitivity, specificity, and -log10p performance measures. The equations used to quantify these performance measures are presented below (in which TP represents true positives, TN represents true negatives, FP represents false positives and FN represents false negatives):

Accuracy=TP+TN/ TP+TN+FP+FN

Sensitivity=TP/TP+FN

Specificity=TN/TN+FP

### Bacterial culture and cell culture

*Bacteroides intestinalis AM-1* strain DSM 17393 (*B. intestinalis*) was purchased from Leibniz-Institute DSMZ (Germany) and cultured in Columbia agar medium supplemented with 5% defibrinated sheep blood. *B. intestinalis* was cultured anaerobically at 37°C for 24□h in an anaerobic cabinet (Whitley DG250 Anaerobic workstation).

### Mouse Model of ALF

Six-week-old male C57BL/6 germ-free mice were bred and maintained at the in-house animal facility of ILBS. All experimental procedures were approved by the Institutional Animal Ethics Committee (Approval No: IAEC/ILBS/23/1). The mice were housed under controlled conditions, with free access to food and water, a 12-hour light/dark cycle, and a stable temperature of 25°C ± 2°C. After a one-week acclimatization period, mice were randomly divided into twelve groups (Supplementary Figure-20).

Acetaminophen (APAP) was dissolved in warm saline (0.9% NaCl at 55°C), then cooled to 37°C before administration. Non-overnight-fasted mice received intraperitoneal (i.p.) injections of APAP at doses of 250 mg/kg or 500 mg/kg, while control groups were injected with normal saline.

In the bacteria pre-treatment group (BI+APAP), mice underwent gut cleansing using polyethylene glycol (PEG) dissolved in normal saline (425 g/L), administered four times at 20-minute intervals. Following gut washing, B. intestinalis was delivered intragastrically at a dose of 1 × 10□ CFU per 200 µL daily for seven days, administered twice, before i.p. injection of APAP at either 250 mg/kg or 500 mg/kg. These mice were sacrificed at 24 hours and 72 hours post-APAP administration.

In another group, non-overnight-fasted mice were first injected i.p. with APAP (250 mg/kg or 500 mg/kg), followed by intragastric administration of B. intestinalis at the same dose (1 × 10□ CFU per 200 µL) daily for three days, administered twice. These mice were also sacrificed at 24 hours and 72 hours post-APAP treatment.

After sacrifice, blood, liver, intestine, and stool samples were collected and stored at −80°C for further analysis.

### Staining procedures

Formalin-fixed tissue samples were embedded in paraffin and stained with H & E. To determine necrosis and cell damage. Five-micrometre frozen sections were then cut and stained with H&E. Representative images from each group of mice are shown in each figure.

### Biochemical analysis

Serum levels of ALT and AST and other liver function test were determined using Infinity ALT kit (Abkine).

### Real-time qPCR

RNA was extracted from mouse liver and intestine and cDNAs were generated as per Verso cDNA Synthesis Kit (Thermoscientific-AB1453A). Primer sequences for mouse genes were obtained from the NCBI and Primer 3. All primers used in this study are listed in Supplementary Table (4). Mouse gene expression and amplification of bacterial genes were determined with Sybr Green (Bio-Rad Laboratories) using Applied Biosystems ViiA 7 Real-Time PCR System. The qPCR value of mouse genes was normalized to GAPDH.

### Statistical Analysis

Results are shown as mean and standard deviation unless indicated otherwise. Statistical analysis was performed using Graph Pad Prism v6, SPSS V20, and P-values of < 0.05 using Benjamini & Hochberg correction were considered statistically significant. Unpaired (two-tail) Student’s t-test, and Mann-Whitney U test were performed for comparison of two groups. For comparison among more than two groups, a one-way analysis of variance, the Kruskal-Walli’s test was performed. All correlations were performed using Spearman correlation analysis and r^2^>0.5, p<0.05 was considered as statistically significant. Annotated features were subjected to different statistical software platforms. First, missing value imputation was applied to data in which half the minimum positive value was estimated for meta-proteome that were undetected in the samples. Subsequently, data were filtered based on non-parametric relative standard deviation (MAD/median) and were subjected to log normalization and Pareto-scaling using Metaboanalyst 5.0 (http://metaboanalyst.ca.) server. PCA and PLS-DA, Heat map, Random-forest analysis, and other statistical analyses were performed. To determine the best Metabolites among the top 5 metabolites univariate and multivariate AUROC analysis was performed by IBM SPSS and Hazard ratio were analyzed. Additionally, survival analysis was performed by Kaplan-Meier curve in IBM SPSS on the basis of panel of 5 metabolites.

## Authors’ contribution

JSM conceptualized the work. SP, and NS helped in sample enrolment, processing, and experimental work and were helped by NS, SB, VS, MK, AK, YM, AG, BM, GT, VB, SY, MY, AK, SS, CB, AK, RM and SKS provided the samples. Data analysis was performed by SP and NS under the guidance of JSM. The manuscript was drafted by SP, NS, and JSM. RM and SKS read the manuscript and provided expert advice. This manuscript has been seen approved by all authors.

## Abbreviations

ALF: Acute Liver Failure
NS: Non-Survivors
S: Survivors
HC: Healthy Control
ALT: Alanine transaminases
AST: Aspartate transaminase
ALP: Alkaline Phosphatase
GGT: Gamma-glutamyl Transferase
MELD: Model for End-Stage Liver Disease
CTP: Child-Turcotte-Pugh
POD: Probability of detection
PLS-DA: Partial least squares-discriminant analysis
LDA: Linear discriminant analysis
AUROC: Area under the Receiver Operating Characteristics
SVM: Support Vector Machine
KNN: k-nearest neighbours
CART: Classification and Regression Tree
RF: Random Forest
HR: Hazard Ratio
ML: Machine Learning
DEM: Differentially Expressed Metabolite
DEMP: Differentially Expressed Meta-protein
ALS: Artificial Liver Support
CDCA: chenodeoxycholic acid
ROS: Reactive Oxygen Species
AKT/PKB: Protein kinase
B JNK: Jun N-terminal kinase
ERK: Extracellular signal-regulated kinases
IL-1β: Interleukin-1-beta
NLRP3: Nucleotide-binding domain, leucine-rich–containing family, pyrin domain containing-3
TRAIL: TNF-related apoptosis-inducing ligand (TRAIL)
SAH: Severe Alcoholic Hepatitis
KCH: King’s College Criteria
SLED: Sustained low-efficiency dialysis
CRRT: Continuous Renal Replacement Therapy
CT: Computed Tomography Scan
HAV: Hepatitis A virus
HEV: Hepatitis E virus
DILI: Drug Induced Liver Injury
TC: Test Cohort
NLDB: National Liver disease Biobank

## Figures and Table legends

**Table 1:**
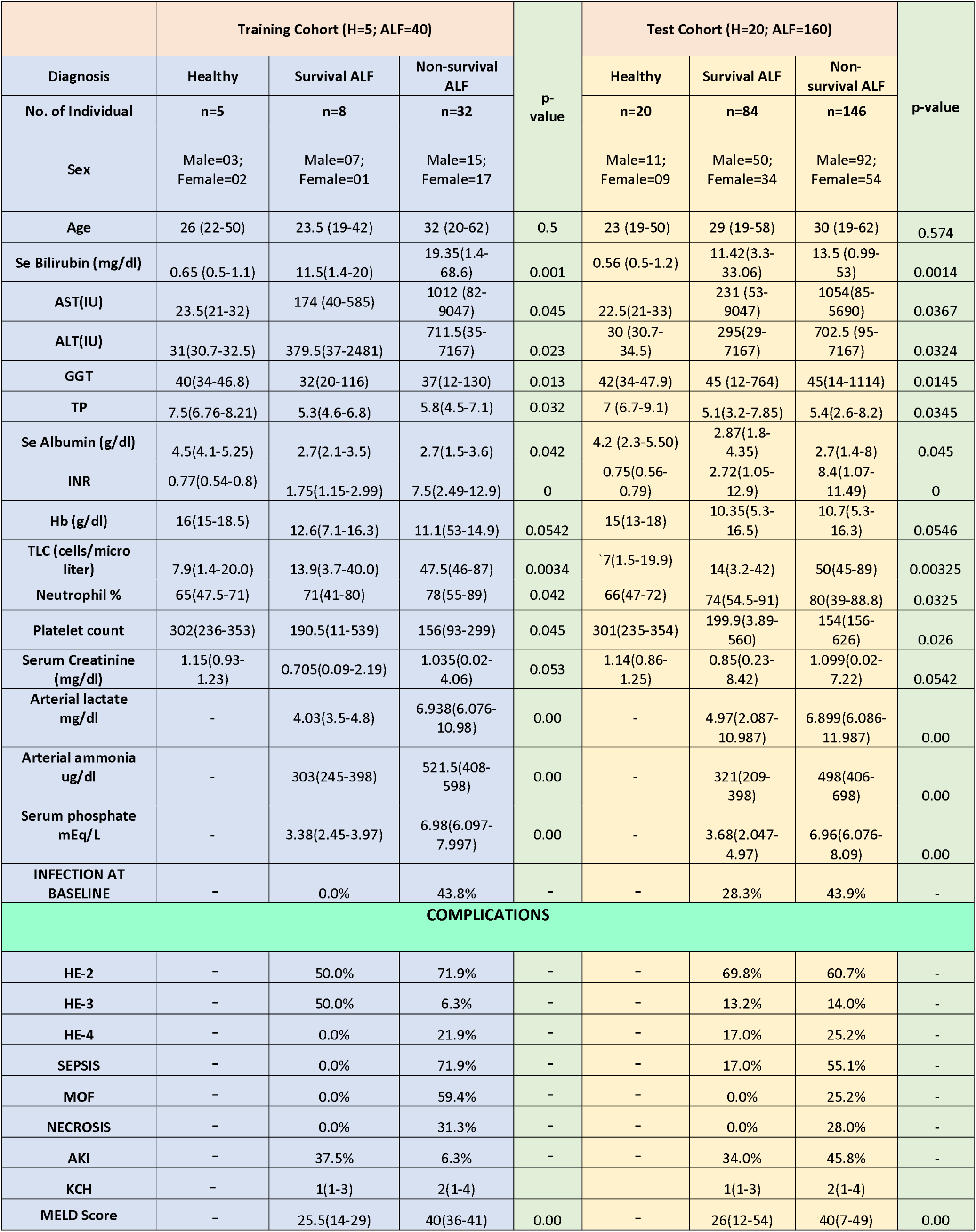
Demographic profile of the Training (HC=5, ALF=40) and test cohort (HC=20, ALF=270) along with complications.

|  | Training Cohort (H=5; ALF=40) |  |  | p-value | Test Cohort (H=20; ALF=160) |  |  | p-value |
| --- | --- | --- | --- | --- | --- | --- | --- | --- |
| Diagnosis | Healthy | Survival ALF | Non-survival ALF |  | Healthy | Survival ALF | Non-survival ALF |  |
| No. of Individual | n=5 | n=8 | n=32 |  | n=20 | n=84 | n=146 |  |
| Sex | Male=03;<br>Female=02 | Male=07;<br>Female=01 | Male=15;<br>Female=17 |  | Male=11;<br>Female=09 | Male=50;<br>Female=34 | Male=92;<br>Female=54 |  |
| Age | 26 (22-50) | 23.5 (19-42) | 32 (20-62) | 0.5 | 23 (19-50) | 29 (19-58) | 30 (19-62) | 0.574 |
| Se Bilirubin (mg/dl) | 0.65 (0.5-1.1) | 11.5(1.4-20) | 19.35(1.4-68.6) | 0.001 | 0.56 (0.5-1.2) | 11.42(3.3-33.06) | 13.5 (0.99-53) | 0.0014 |
| AST(IU) | 23.5(21-32) | 174 (40-585) | 1012 (82-9047) | 0.045 | 22.5(21-33) | 231 (53-9047) | 1054(85-5690) | 0.0367 |
| ALT(IU) | 31(30.7-32.5) | 379.5(37-2481) | 711.5(35-7167) | 0.023 | 30 (30.7-34.5) | 295(29-7167) | 702.5 (95-7167) | 0.0324 |
| GGT | 40(34-46.8) | 32(20-116) | 37(12-130) | 0.013 | 42(34-47.9) | 45 (12-764) | 45(14-1114) | 0.0145 |
| TP | 7.5(6.76-8.21) | 5.3(4.6-6.8) | 5.8(4.5-7.1) | 0.032 | 7 (6.7-9.1) | 5.1(3.2-7.85) | 5.4(2.6-8.2) | 0.0345 |
| Se Albumin (g/dl) | 4.5(4.1-5.25) | 2.7(2.1-3.5) | 2.7(1.5-3.6) | 0.042 | 4.2 (2.3-5.50) | 2.87(1.8-4.35) | 2.7(1.4-8) | 0.045 |
| INR | 0.77(0.54-0.8) | 1.75(1.15-2.99) | 7.5(2.49-12.9) | 0 | 0.75(0.56-0.79) | 2.72(1.05-12.9) | 8.4(1.07-11.49) | 0 |
| Hb (g/dl) | 16(15-18.5) | 12.6(7.1-16.3) | 11.1(53-14.9) | 0.0542 | 15(13-18) | 10.35(5.3-16.5) | 10.7(5.3-16.3) | 0.0546 |
| TLC(cells/micro liter) | 7.9(1.4-20.0) | 13.9(3.7-40.0) | 47.5(46-87) | 0.0034 | 7(1.5-19.9) | 14(3.2-42) | 50(45-89) | 0.00325 |
| Neutrophil % | 65(47.5-71) | 71(41-80) | 78(55-89) | 0.042 | 66(47-72) | 74(54.5-91) | 80(39-88.8) | 0.0325 |
| Platelet count | 302(236-353) | 190.5(11-539) | 156(93-299) | 0.045 | 301(235-354) | 199.9(3.89-560) | 154(156-626) | 0.026 |
| Serum Creatinine (mg/dl) | 1.15(0.93-1.23) | 0.705(0.09-2.19) | 1.035(0.02-4.06) | 0.053 | 1.14(0.86-1.25) | 0.85(0.23-8.42) | 1.099(0.02-7.22) | 0.0542 |
| Arterial lactate mg/dl | - | 4.03(3.5-4.8) | 6.938(6.076-10.98) | 0.00 | - | 4.97(2.087-10.987) | 6.899(6.086-11.987) | 0.00 |
| Arterial ammonia ug/dl | - | 303(245-398) | 521.5(408-598) | 0.00 | - | 321(209-398) | 498(406-698) | 0.00 |
| Serum phosphate mEq/L | - | 3.38(2.45-3.97) | 6.98(6.097-7.997) | 0.00 | - | 3.68(2.047-4.97) | 6.96(6.076-8.09) | 0.00 |
| INFECTION AT BASELINE | - | 0.0% | 43.8% | - | - | 28.3% | 43.9% | - |
| COMPLICATIONS |  |  |  |  |  |  |  |  |
| HE-2 | - | 50.0% | 71.9% | - | - | 69.8% | 60.7% | - |
| HE-3 | - | 50.0% | 6.3% | - | - | 13.2% | 14.0% | - |
| HE-4 | - | 0.0% | 21.9% | - | - | 17.0% | 25.2% | - |
| SEPSIS | - | 0.0% | 71.9% | - | - | 17.0% | 55.1% | - |
| MOF | - | 0.0% | 59.4% | - | - | 0.0% | 25.2% | - |
| NECROSIS | - | 0.0% | 31.3% | - | - | 0.0% | 28.0% | - |
| AKI | - | 37.5% | 6.3% | - | - | 34.0% | 45.8% | - |
| KCH | - | 1(1-3) | 2(1-4) |  |  | 1(1-3) | 2(1-4) |  |
| MELD Score | - | 25.5(14-29) | 40(36-41) | 0.00 | - | 26(12-54) | 40(7-49) | 0.00 |

**Supplementary Table 1:** Description of the metabolites identified along with their expression status in ALF vs H and NS vs S vs H (p<0.05, FC> 1.5)

**Supplementary Table 2:** Description of the Meta-Proteins (Bacterial peptides) identified along with their taxonomic classification and expression values in in ALF vs H and NS vs S vs H (p<0.05, FC> 1.5)

**Supplementary Table 3:** Correlation of bacterial peptides (WMpCNA) with the metabolomic pathways (R^2^>0.3; p<0.05)

**Supplementary Table 4:** List of primers of bile acid transport

