## Supplementary Figures for "Chenodeoxycholic Acid Modulation via *Bacteroides intestinalis* AM1 underscores a Novel Approach in Acute Liver Failure"

**Supplementary Figure 1:** Overview of the study design for metaproteome (with red arrows) (ALF; n=40), and metabolome (with green arrows) (ALF; n=40) (ALF; n=160) done in training cohort and test cohort (plasma and one drop blood)

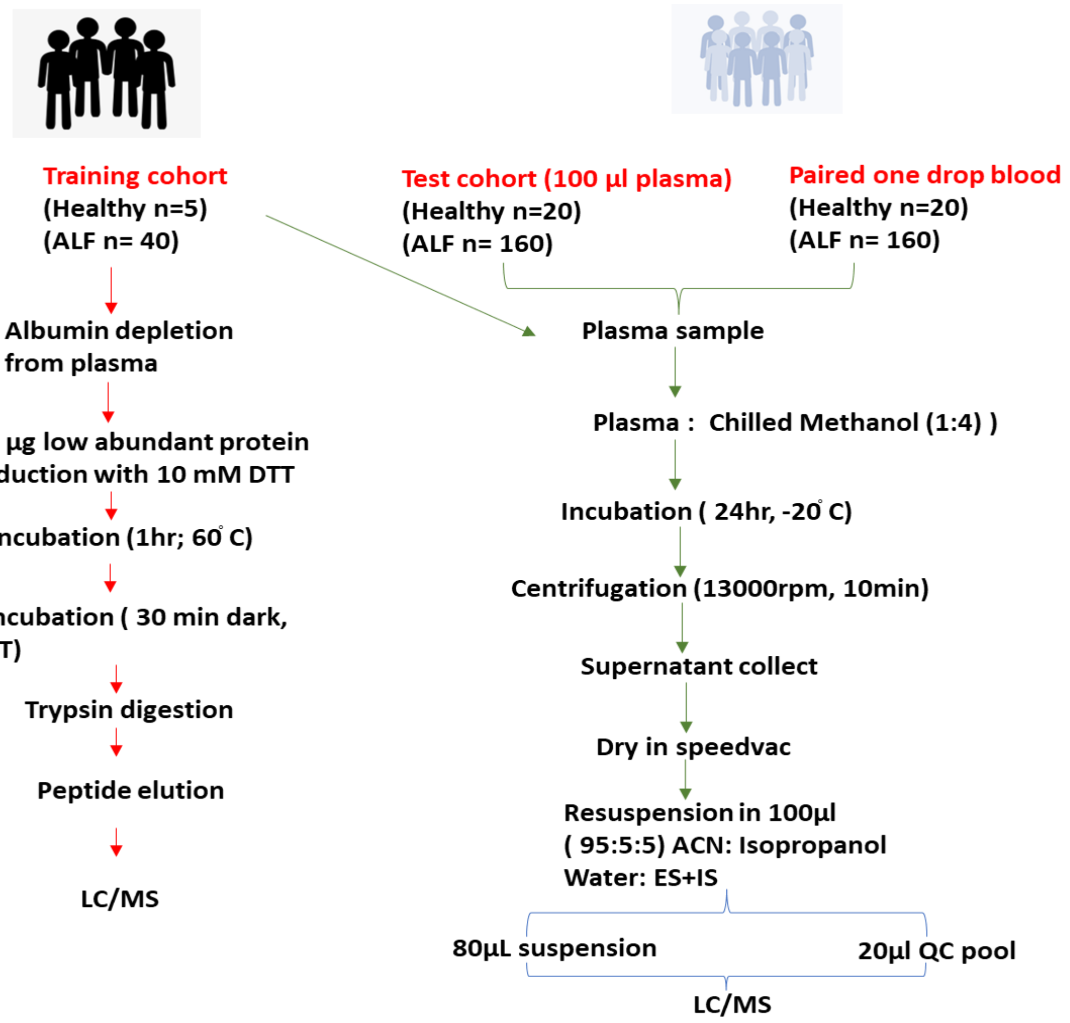

**Supplementary Figure 2**: Flow chart of statistical analysis performed for Integrated Analysis of Plasma Metabolome and Meta-Proteome in Baseline Plasma and also to elucidate key Metabolome Indicators of Poor outcomes in Acute Liver Failure Patients.

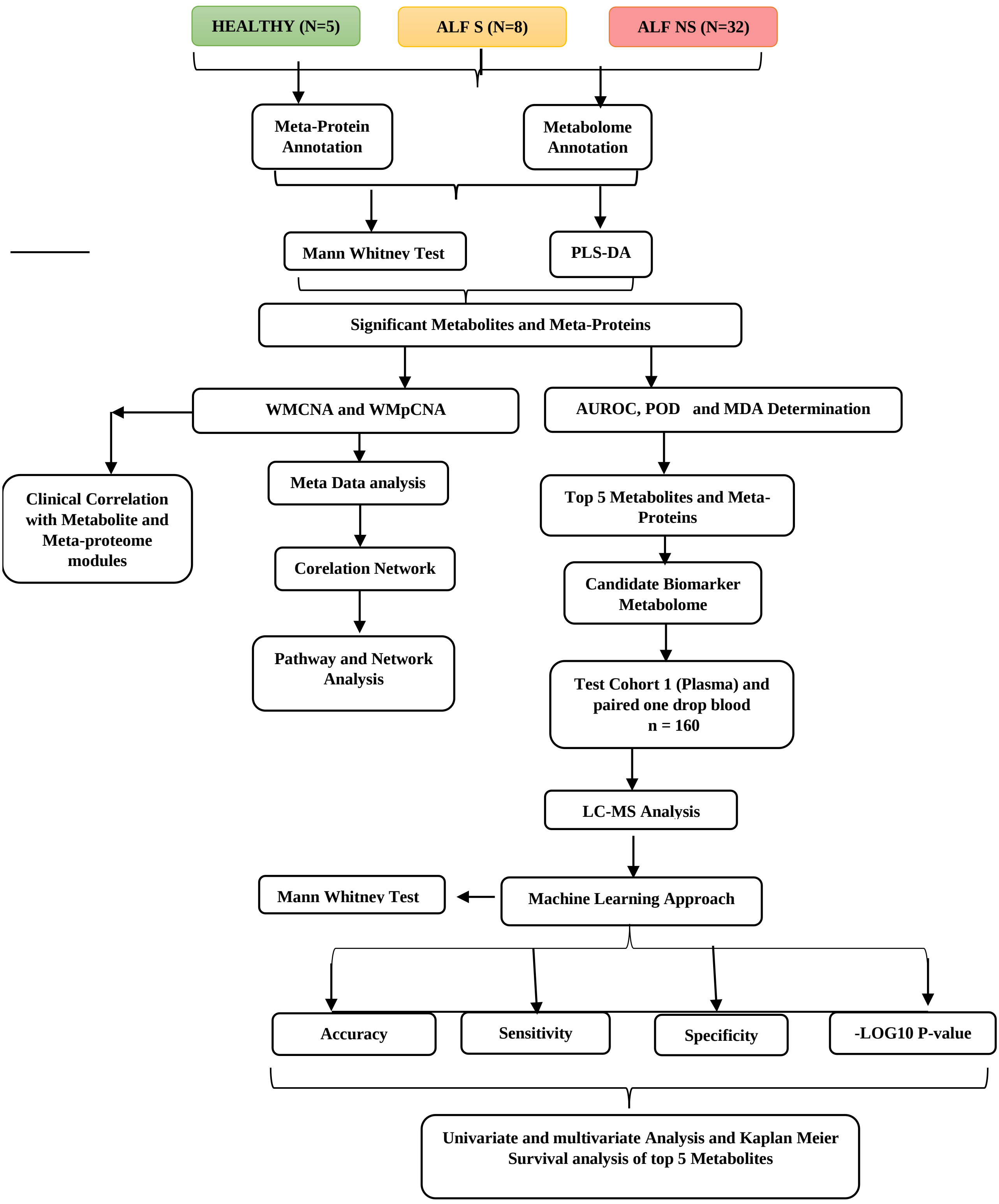

**Supplementary Figure 3:** Metabolome profile could help to stratify non-survivors of ALF and healthy control detailed in Panel-1

**Panel 1- Metabolomics profile of different study groups**

- **Metabolomics profile of ALF VS Healthy**

A) Volcano plot of ALF vs. Healthy baseline plasma samples in training cohort. green dot; downregulated and red dot; upregulated (FC>1.5, $p < 0.05$; Mann-Whitney-U-test, FDR<0.01; Benjamini-Hochberg-correction).

B) Partial Least Squares - Discriminant Analysis (PLS-DA) Plot showing clear separation of protein species in Acute liver failure and healthy control samples with ALF; HC revealing component variance

C) Loading plot between the ALF and healthy samples.

D) VIP scores of important features identified by PLS-DA in ALF and healthy samples.

E) Heat map showing hierarchical clustering of 50 different metabolite species from ALF vs Healthy (distance measure using Euclidean, and clustering algorithm using ward D.

F) 15 different metabolite features identified by Random Forest and ranked by the mean decrease in classification accuracy from ALF VS H.

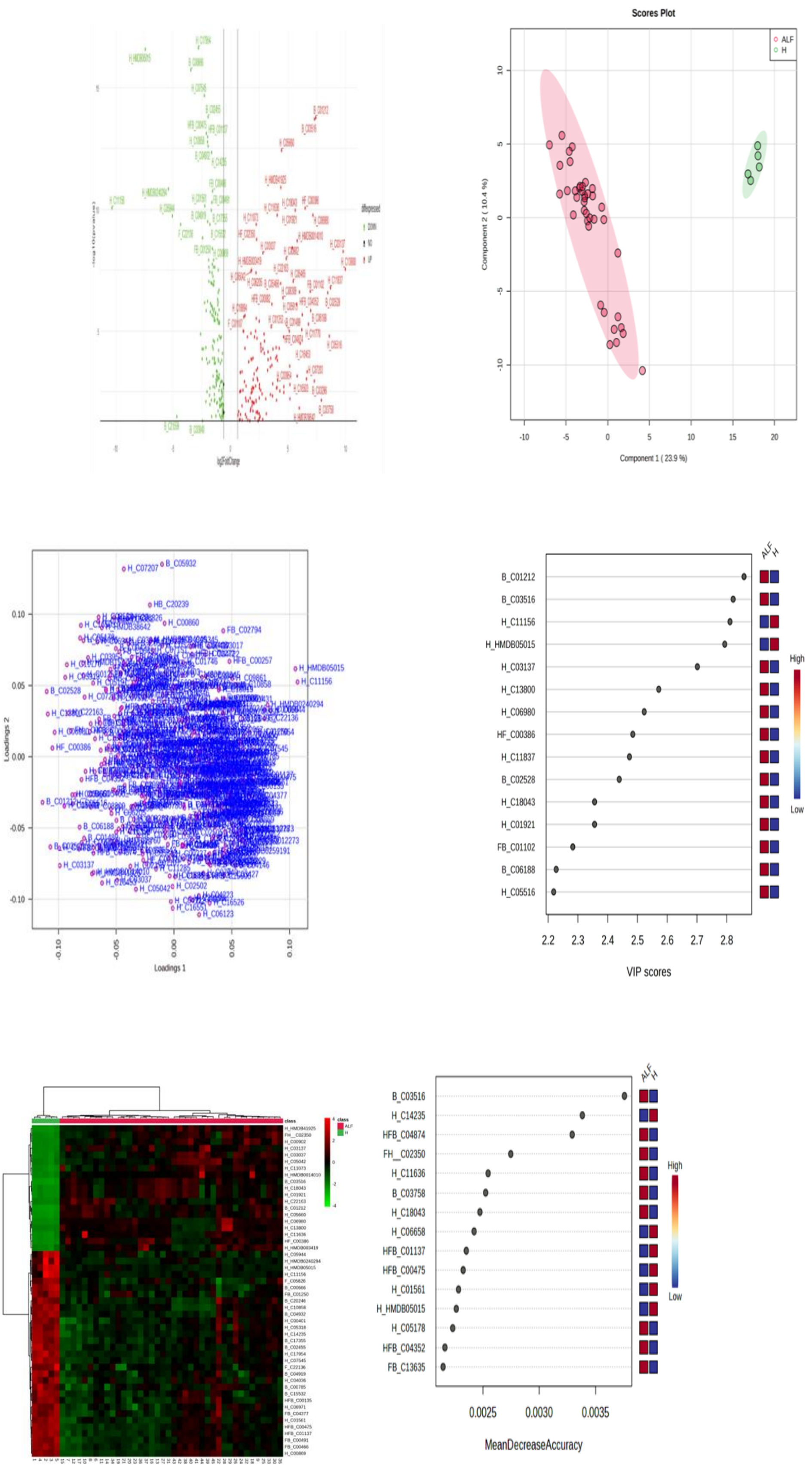

log2FoldChange
Scores Plot
ALF
H
Component 1 ( 23.9 %)
Component 2 ( 10.4 %)
Loadings 1
Loadings 2
B_C01212
B_C03516
H_C11156
H_HMDB05015
H_C03137
H_C13800
H_C06980
HF_C00386
H_C11837
B_C02528
H_C18043
H_C01921
FB_C01102
B_C06188
H_C05516
High
Low
VIP scores
B_C03516
H_C14235
HFB_C04874
FH_C02350
H_C11636
B_C03758
H_C18043
H_C06658
HFB_C01137
HFB_C00475
H_C01561
H_HMDB05015
H_C05178
HFB_C04352
FB_C13635
MeanDecreaseAccuracy

**Supplementary Figure 4: Metabolite profile could stratify non-survivors of Acute liver failure**

Panel 3- A) Volcano plot of NS vs. S baseline plasma samples in training cohort. green dot; downregulated and red dot; upregulated (FC>1.5, $p < 0.05$; Mann-Whitney-U-test, FDR<0.01; Benjamini-Hochberg-correction).

B) Partial Least Squares - Discriminant Analysis (PLS-DA) Plot showing clear separation of metabolite species in with NS vs. S

C) Loading plot between the selected components with NS vs. S

D) VIP scores of important features identified by PLS-DA with NS vs. S

E) Heat map showing hierarchical clustering of 50 different metabolite species from NS VS S (distance measure using Euclidean, and clustering algorithm using ward D.

F) 15 different metabolite features identified by Random Forest and ranked by the mean decrease in classification accuracy from NS vs. S

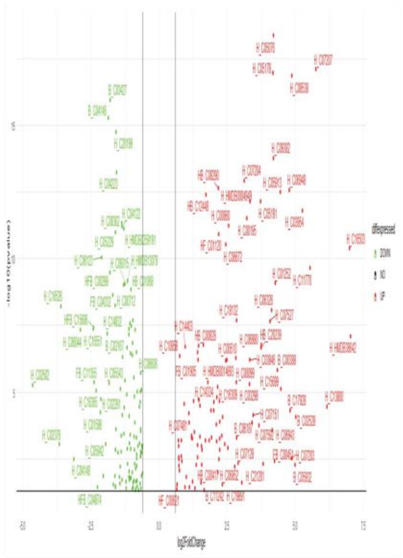

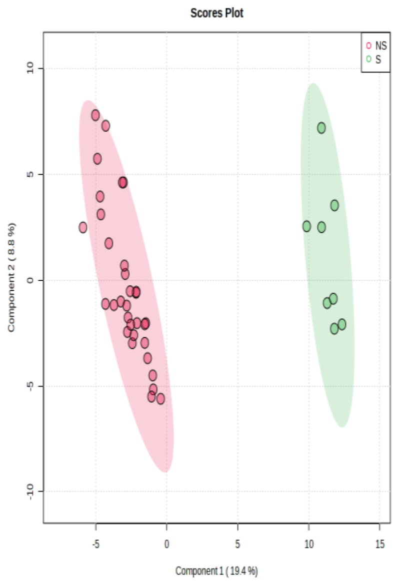

Scores Plot
NS
S
Component 2 ( 8.8 %)
Component 1 ( 19.4 %)


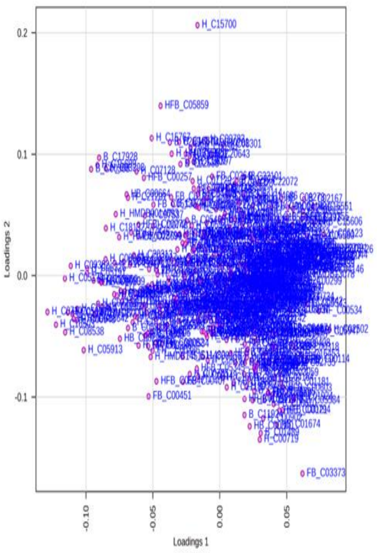

Loadings 2
Loadings 1


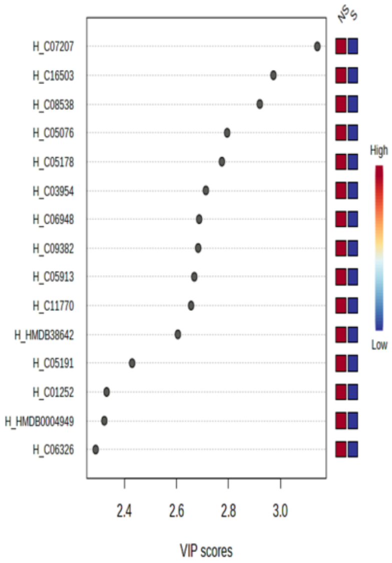

NS
S
H_C07207
H_C16503
H_C08538
H_C05076
H_C05178
H_C03954
H_C06948
H_C09382
H_C05913
H_C11770
H_HMDB38642
H_C05191
H_C01252
H_HMDB0004949
H_C06326
High
Low
VIP scores


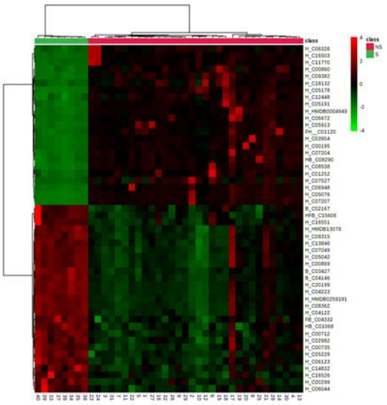

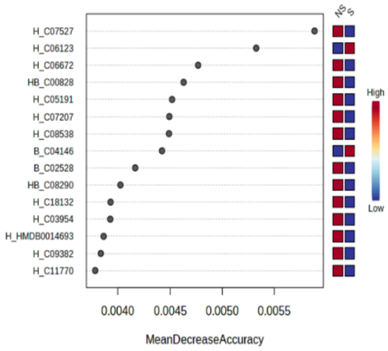

NS
S
H_C07527
H_C06123
H_C06672
HB_C00828
H_C05191
H_C07207
H_C08538
B_C04146
B_C02528
HB_C08290
H_C18132
H_C03954
H_HMDB0014693
H_C09382
H_C11770
High
Low
MeanDecreaseAccuracy

**Supplementary Figure 5:** Heat map showing hierarchical clustering of 50 different metabolome from ALF vs Healthy and NS VS S (distance measure using Euclidean, and clustering algorithm using ward D and 15 different metabolite features identified by Random Forest and ranked by the mean decrease in classification accuracy from ALF vs Healthy and NS VS S

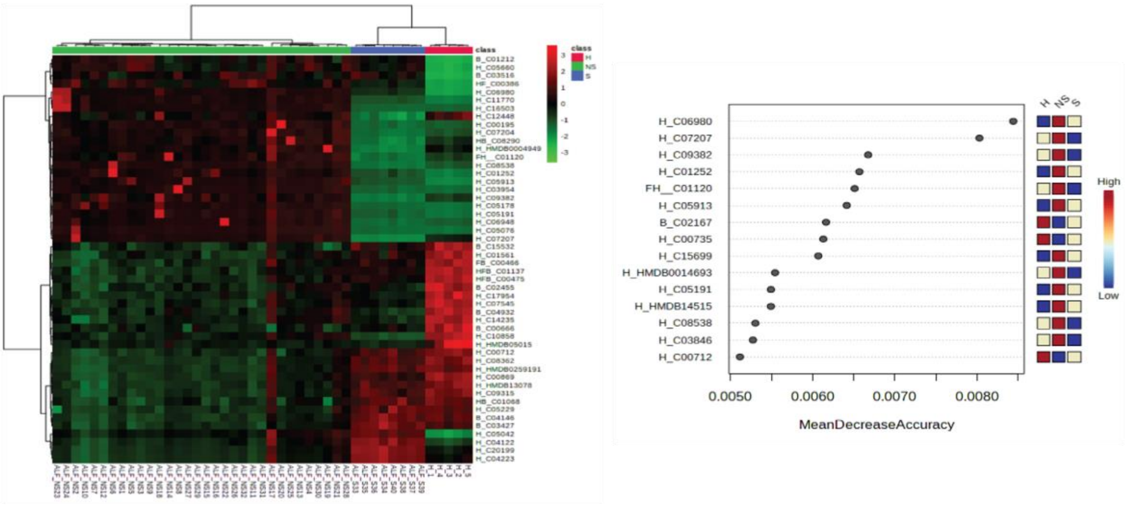

**Supplementary figure 6: Meta-proteomics profile of different study groups**

A) Metaproteome abundance in the study group ALF VS H at the phylum level ($p<0.05$).

B) Alpha diversity indices (Shannon/Simpson index) in metaproteome of ALF VS Healthy.

C) Beta Diveristy plot plot showing clear separation of metaproteome in ALF VS H.

D) Partial Least Squares - Discriminant Analysis (PLS-DA) Plot showing clear separation of metabolite species in with ALF Vs H

E) 15 different Bacterial features identified by Random Forest and ranked by the mean decrease in classification accuracy from ALF Vs H

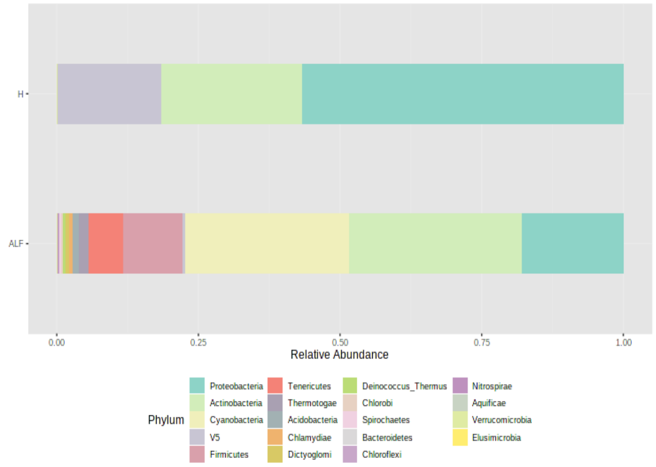
H
ALF
0.00
0.25
0.50
0.75
1.00
Relative Abundance
Phylum
Proteobacteria
Actinobacteria
Cyanobacteria
V5
Firmicutes
Tenericutes
Thermotogae
Acidobacteria
Chlamydiae
Dictyoglomi
Deinococcus_Thermus
Chlorobi
Spirochaetes
Bacteroidetes
Chloroflexi
Nitrospirae
Aquificae
Verrucomicrobia
Elusimicrobia

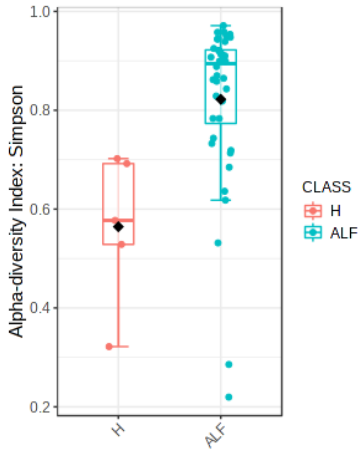
Alpha-diversity Index: Simpson
1.0
0.8
0.6
0.4
0.2
H
ALF
CLASS
H
ALF

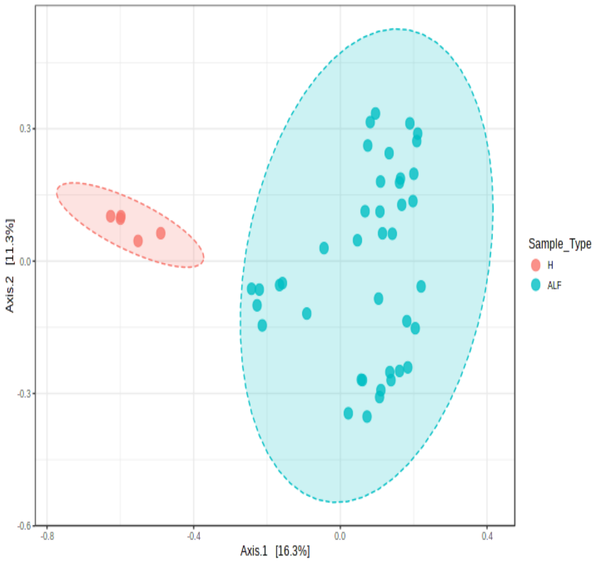
0.3
0.0
-0.3
-0.6
Axis.2 [11.3%]
-0.8
-0.4
0.0
0.4
Axis.1 [16.3%]
Sample_Type
H
ALF

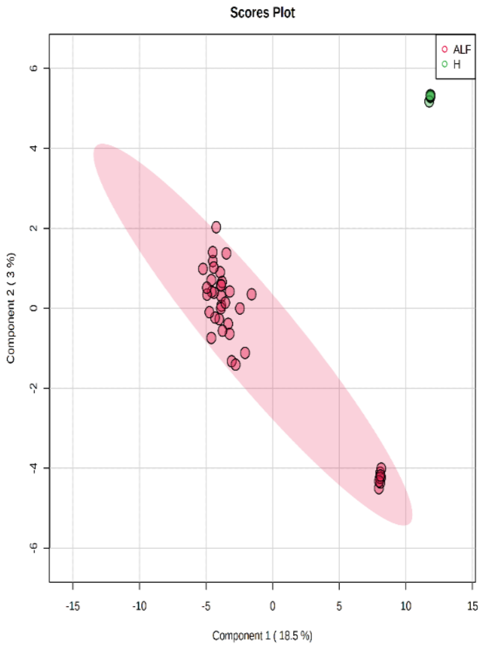
Scores Plot
ALF
H
6
4
2
0
-2
-4
-6
Component 2 ( 3 %)
-15
-10
-5
0
5
10
15
Component 1 ( 18.5 %)

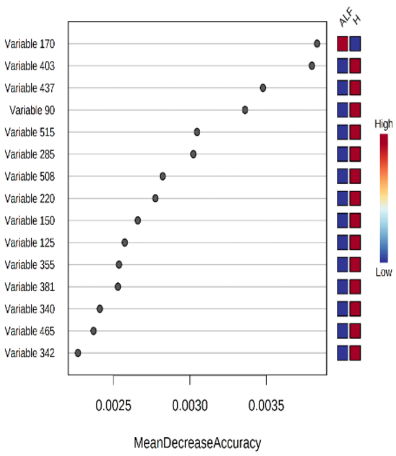
ALF
H
Variable 170
Variable 403
Variable 437
Variable 90
Variable 515
Variable 285
Variable 508
Variable 220
Variable 150
Variable 125
Variable 355
Variable 381
Variable 340
Variable 465
Variable 342
High
Low
0.0025
0.0030
0.0035
MeanDecreaseAccuracy

**Supplementary Figure 7: Meta-proteomics profile of different study groups**

A) Metaproteome abundance in the study group NS VS S at the phylum level ($p<0.05$).

B) Alpha diversity indices (Shannon/Simpson index) in metaproteome of NS VS S.

C) Partial Least Squares - Discriminant Analysis (PLS-DA) Plot showing clear separation of metabolite species in with NS VS S

D) Beta Diveristy plot plot showing clear separation of metaproteome in NS VS S.

E) LDA scores for bacterial peptides in NS VS S

F) 15 different Bacterial features identified by Random Forest and ranked by the mean decrease in classification accuracy from NS VS S

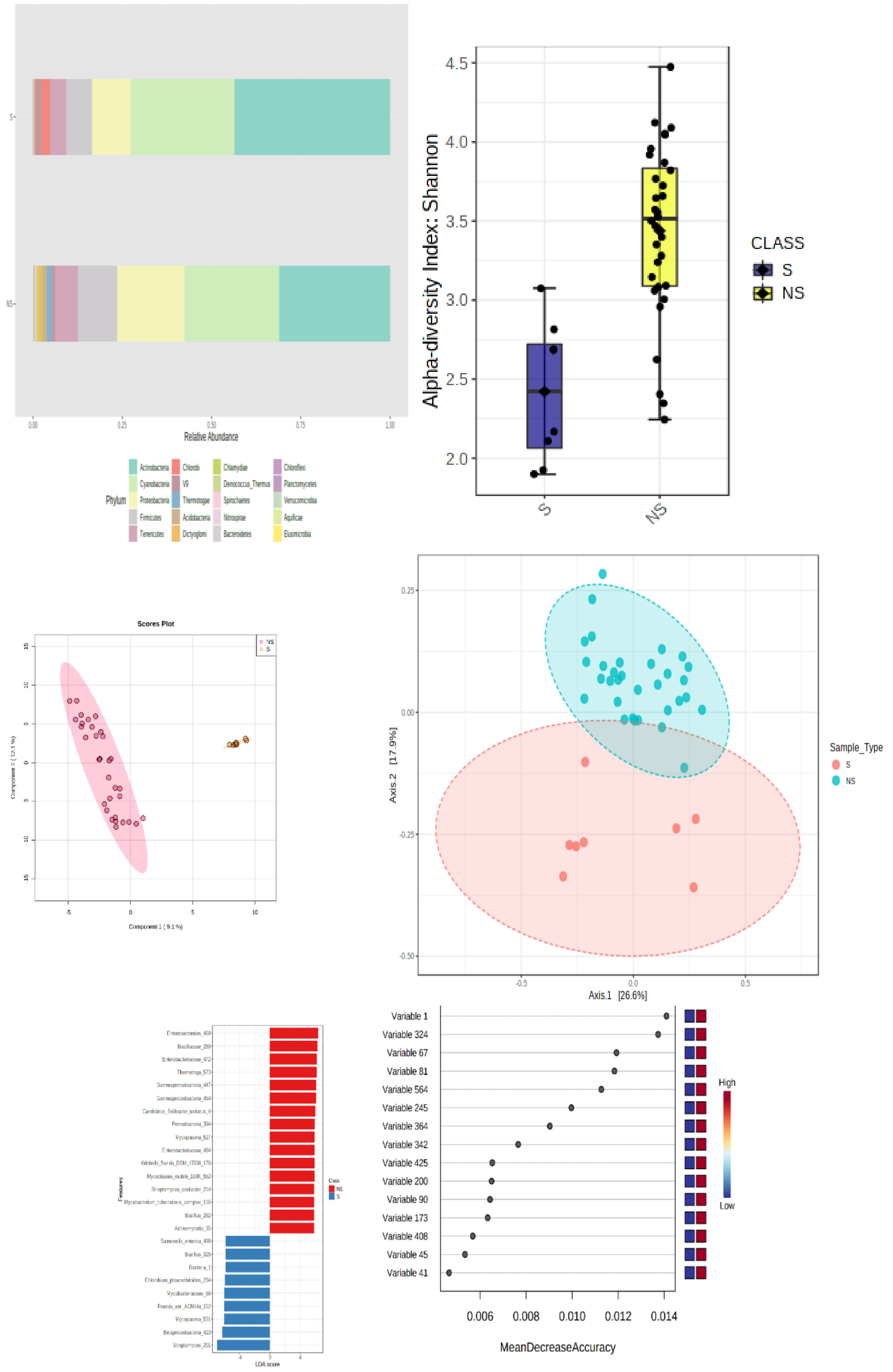

Relative Abundance
0.00
0.25
0.50
0.75
1.00
S
NS
Phylum
Actinobacteria
Chlorobi
Chlamydiae
Chloroflexi
Cyanobacteria
Deinococcus_Thermus
Planctomycetes
Proteobacteria
Thermotogae
Spirochaetes
Verrucomicrobia
Firmicutes
Acidobacteria
Nitrospirae
Aquificae
Tenericutes
Dictyoglomi
Bacteroidetes
Elusimicrobia
Alpha-diversity Index: Shannon
4.5
4.0
3.5
3.0
2.5
2.0
S
NS
CLASS
S
NS
Scores Plot
Axis.2 [17.9%]
Axis.1 [26.6%]
0.25
0.00
-0.25
-0.50
-0.5
0.0
0.5
Sample_Type
S
NS
Variable 1
Variable 324
Variable 67
Variable 81
Variable 564
Variable 245
Variable 364
Variable 342
Variable 425
Variable 200
Variable 90
Variable 173
Variable 408
Variable 45
Variable 41
0.006
0.008
0.010
0.012
0.014
MeanDecreaseAccuracy
High
Low

**Supplementary Figure 8: (A)** Relative metaproteome abundance (ALF-Survivor; ALF-Non-Survivor, Healthy-control) at phylum and genus levels ($p < 0.05$)

(B) Cluster of Orthologous Groups function analysis of bacterial peptides in study groups HC, ALF-S, and ALF-NS.

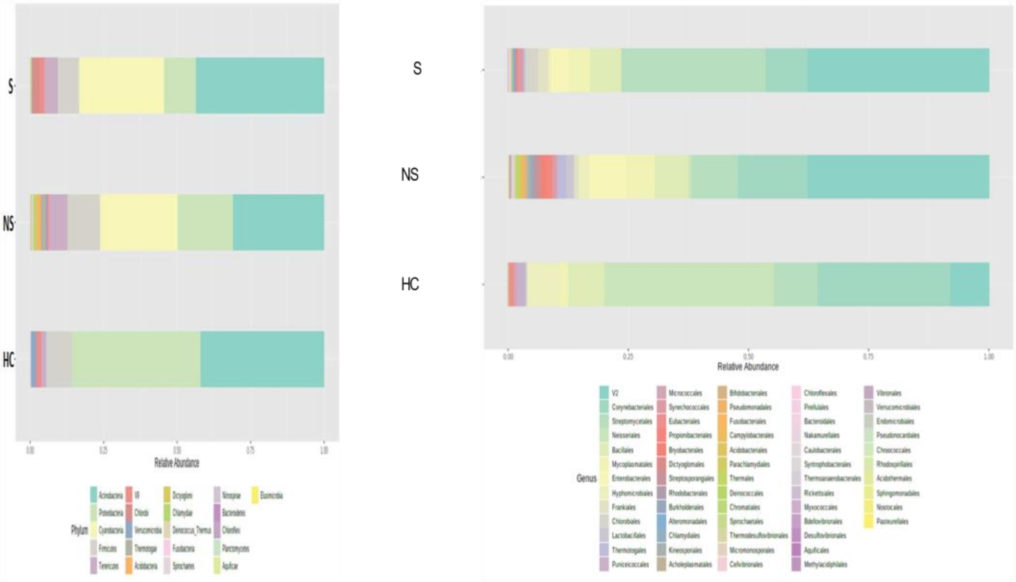


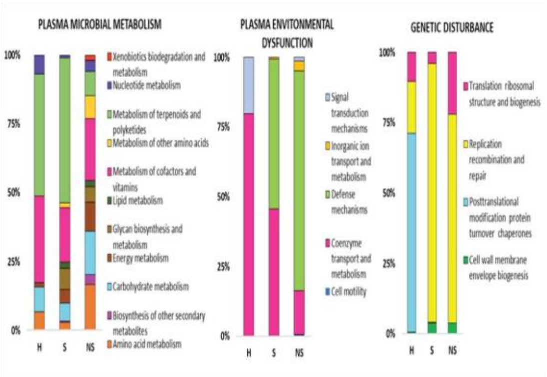

**Supplementary figure 9:** Bacterial features associated with different WMpCNA module

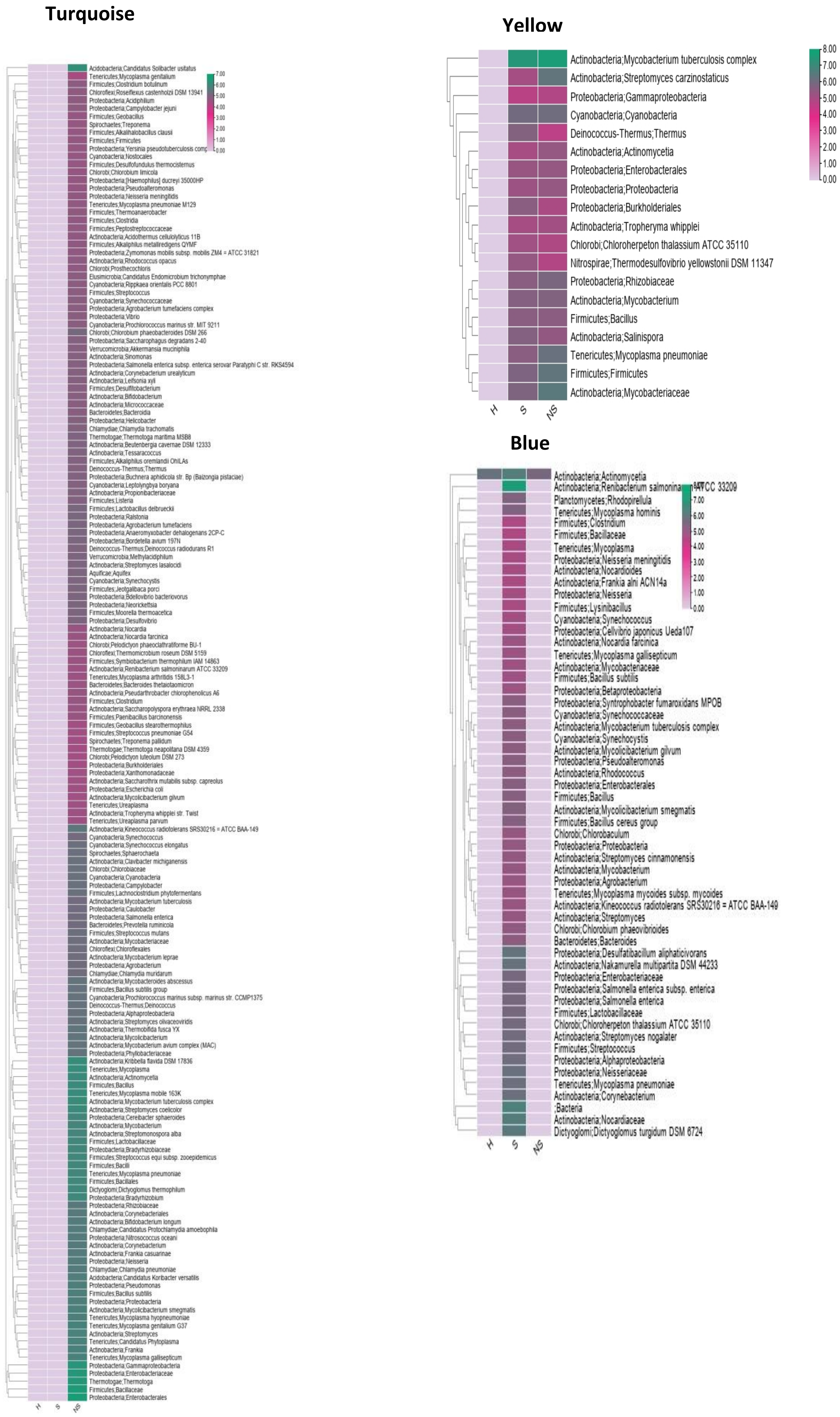

**Brown**

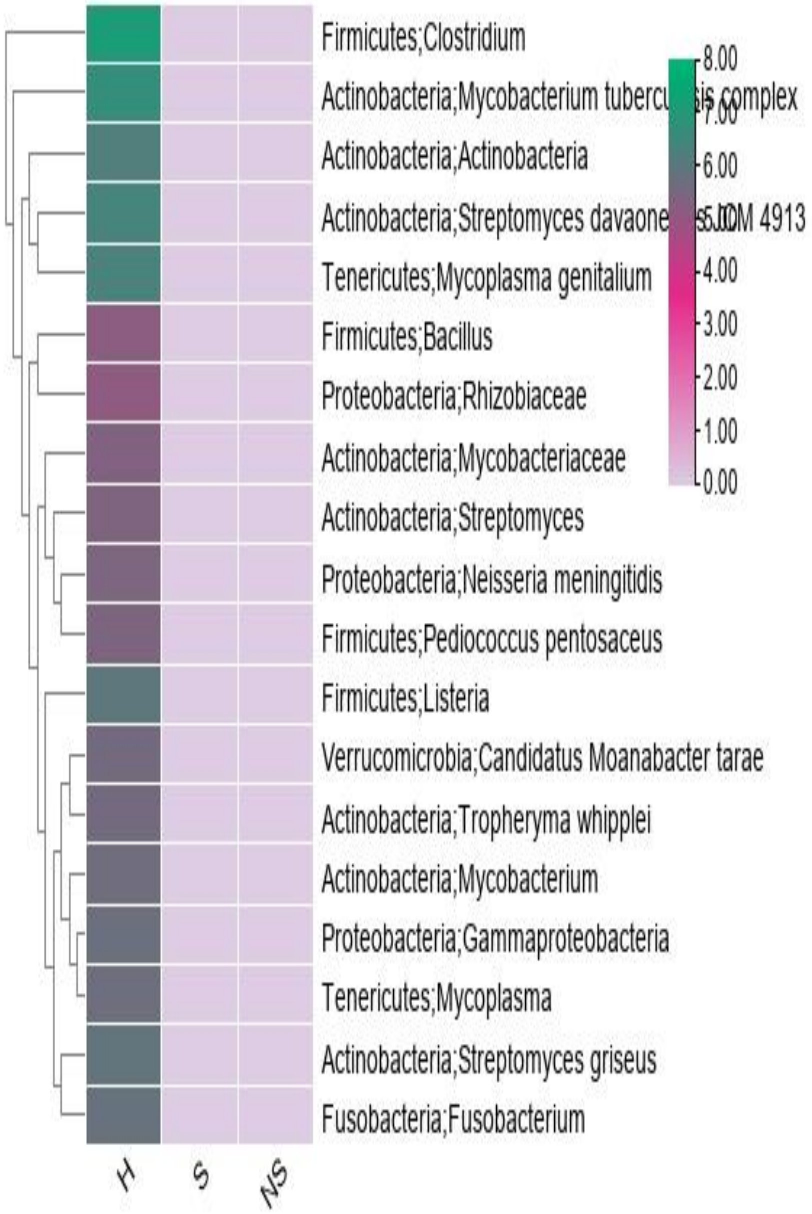
Firmicutes;Clostridium
Actinobacteria;Mycobacterium tuberculosis complex
Actinobacteria;Actinobacteria
Actinobacteria;Streptomyces davaonensis JCM 4913
Tenericutes;Mycoplasma genitalium
Firmicutes;Bacillus
Proteobacteria;Rhizobiaceae
Actinobacteria;Mycobacteriaceae
Actinobacteria;Streptomyces
Proteobacteria;Neisseria meningitidis
Firmicutes;Pediococcus pentosaceus
Firmicutes;Listeria
Verrucomicrobia;Candidatus Moanabacter tarae
Actinobacteria;Tropheryma whipplei
Actinobacteria;Mycobacterium
Proteobacteria;Gammaproteobacteria
Tenericutes;Mycoplasma
Actinobacteria;Streptomyces griseus
Fusobacteria;Fusobacterium
H
S
NS
8.00
7.00
6.00
5.00
4.00
3.00
2.00
1.00
0.00

**Supplementary Figure 10:** Correlation plot of functions annotated from directly correlating Mp functionality and metabolome modules

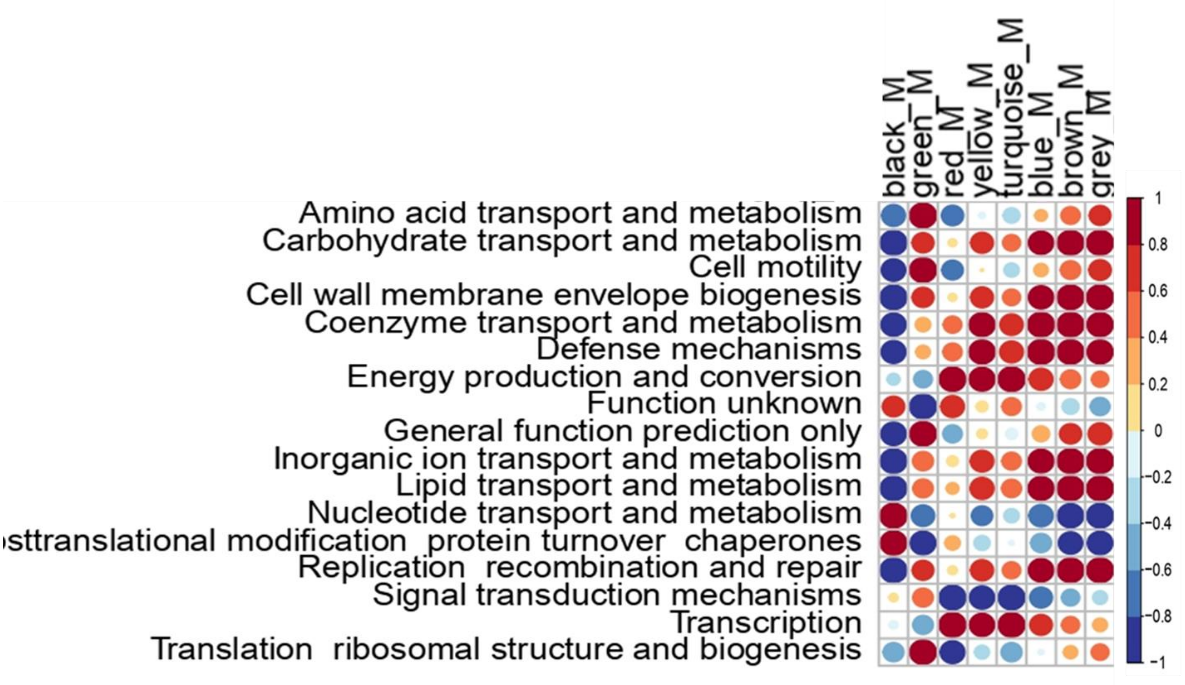

**Supplementary Figure 11**: Linear regression of the meta-protein's turquoise module with the clinical complication of ALF

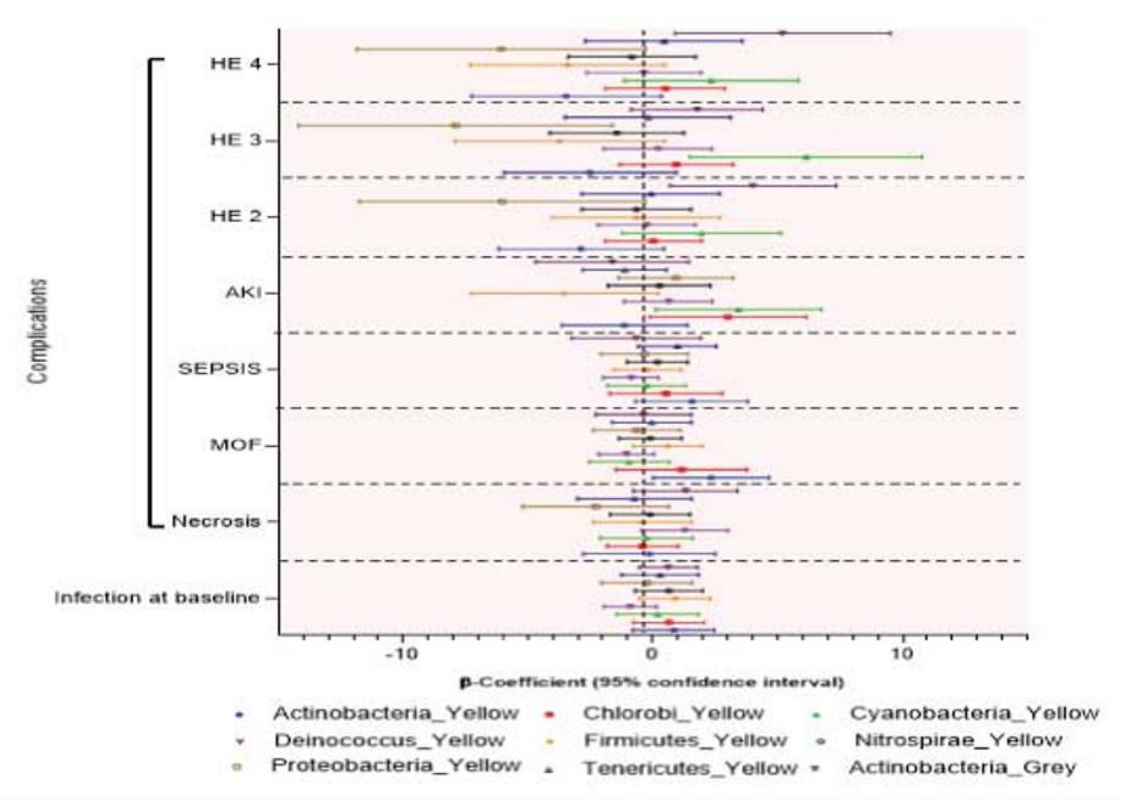

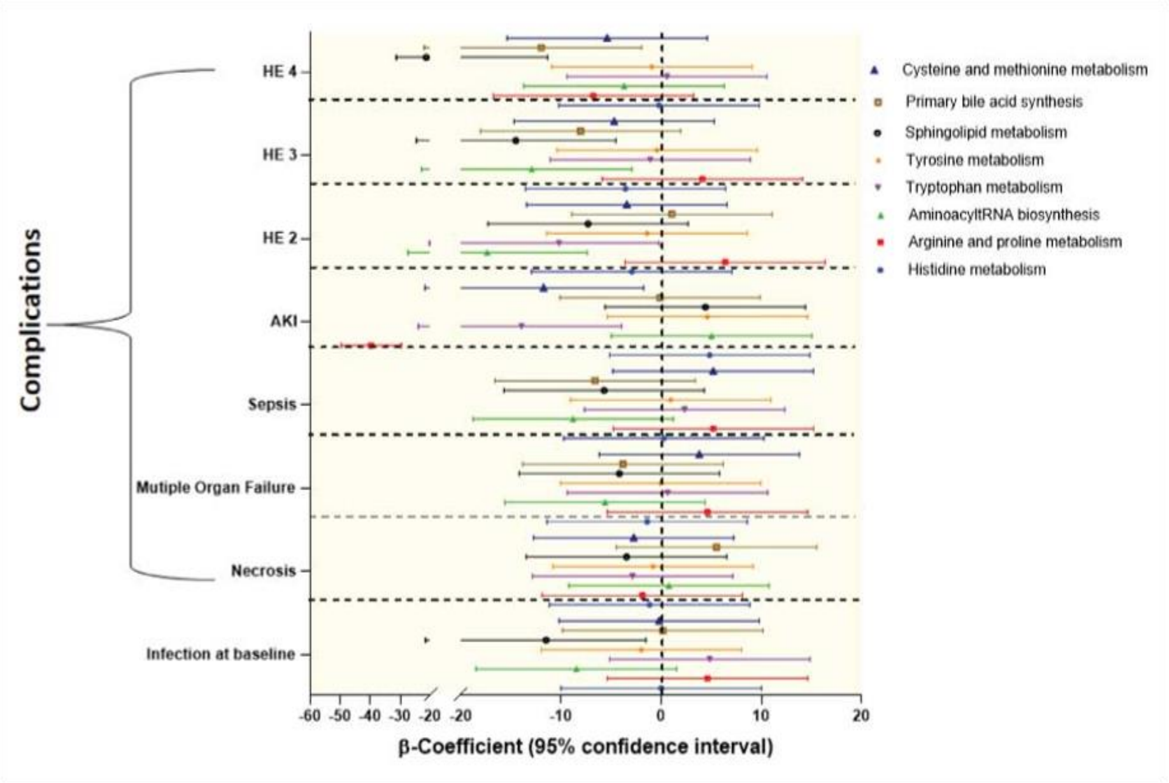
Complications
HE 4
HE 3
HE 2
AKI
Sepsis
Mutiple Organ Failure
Necrosis
Infection at baseline
-60 -50 -40 -30 -20 -20 -10 0 10 20
β-Coefficient (95% confidence interval)
Cysteine and methionine metabolism
Primary bile acid synthesis
Sphingolipid metabolism
Tyrosine metabolism
Tryptophan metabolism
AminoacyltRNA biosynthesis
Arginine and proline metabolism
Histidine metabolism

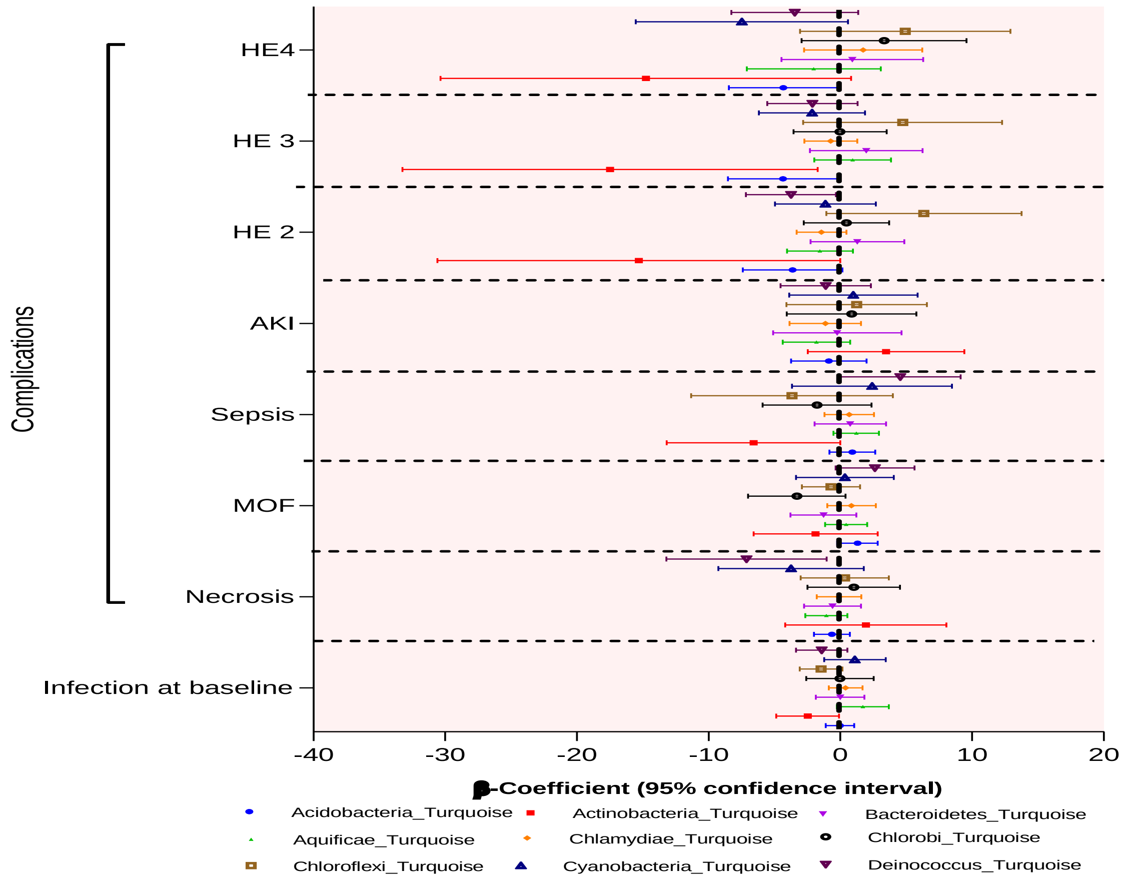
Complications
HE4
HE 3
HE 2
AKI
Sepsis
MOF
Necrosis
Infection at baseline
-40 -30 -20 -10 0 10 20
β-Coefficient (95% confidence interval)
Acidobacteria_Turquoise
Actinobacteria_Turquoise
Bacteroidetes_Turquoise
Aquificae_Turquoise
Chlamydiae_Turquoise
Chlorobi_Turquoise
Chloroflexi_Turquoise
Cyanobacteria_Turquoise
Deinococcus_Turquoise

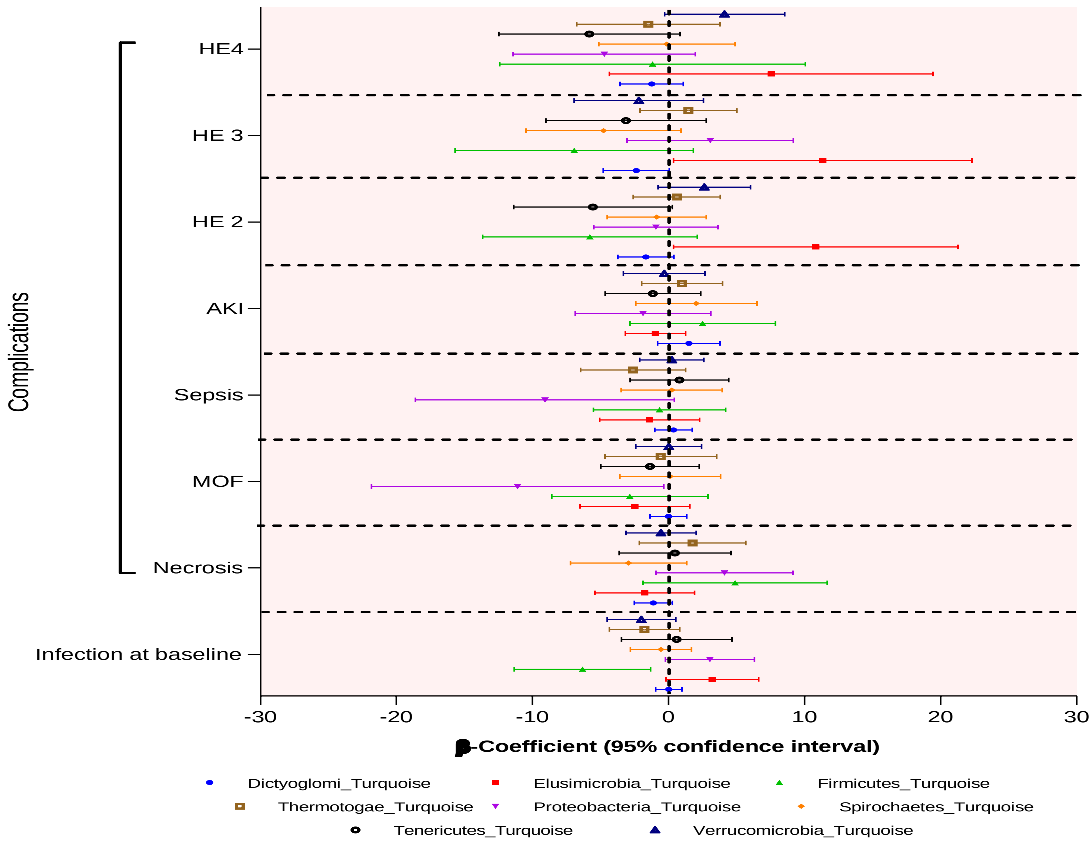

Complications
HE4
HE 3
HE 2
AKI
Sepsis
MOF
Necrosis
Infection at baseline
-30
-20
-10
0
10
20
30
β-Coefficient (95% confidence interval)
Dictyoglomi_Turquoise
Elusimicrobia_Turquoise
Firmicutes_Turquoise
Thermotogae_Turquoise
Proteobacteria_Turquoise
Spirochaetes_Turquoise
Tenericutes_Turquoise
Verrucomicrobia_Turquoise

**Supplementary Figure 12:** Meta-proteomic analysis on the basis of etiology (HEV, DILI, HAV and Others) **Figure 1A-** Unsupervised hierarchical clustering with different etiology showing meta-proteomic phylum LCA signatures is capable to segregate ALF-Non-Survivor. The expression is given as Red; upregulated, Green; downregulated, and Black; unregulated ($p<0.05$; Kruskal-Wallis-test) **Figure 1B-** Linear Discriminating Analysis (LDA) of bacterial peptides in different etiology study group, ALF-survivor, ALF-non-survivor ($p < 0.05$; Kruskal-Wallis-test) **Figure 1C-**Mean Decrease in Accuracy showing top 15 bacterial peptides along (Red; upregulated and blue; downregulated)

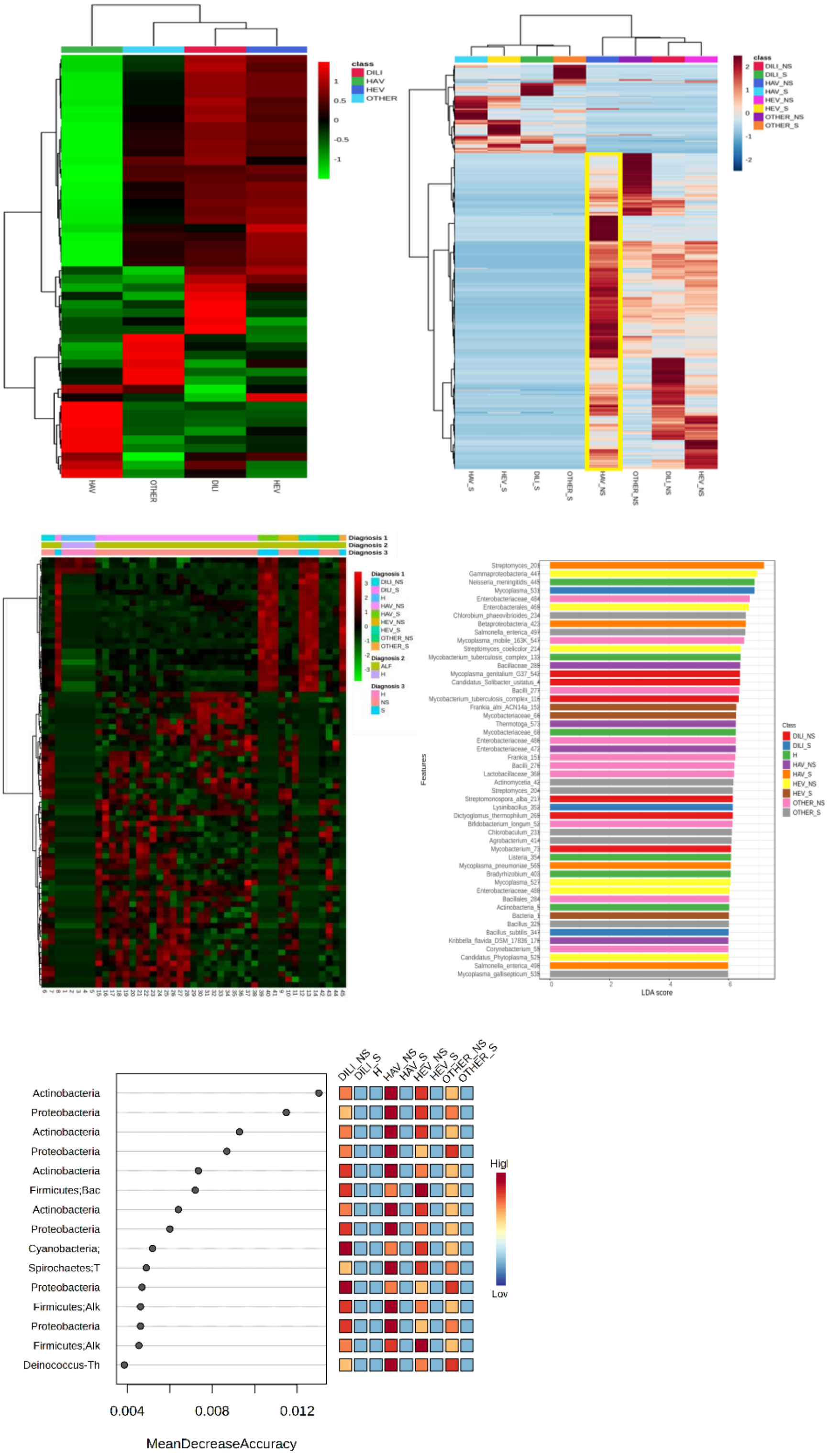
class
DILI
HAV
HEV
OTHER
HAV
OTHER
DILI
HEV
DILI_NS
DILI_S
HAV_NS
HAV_S
HEV_NS
HEV_S
OTHER_NS
OTHER_S
Diagnosis 1
Diagnosis 2
Diagnosis 3
Features
LDA score
Class
Actinobacteria
Proteobacteria
Actinobacteria
Proteobacteria
Actinobacteria
Firmicutes;Bac
Actinobacteria
Proteobacteria
Cyanobacteria;
Spirochaetes;T
Proteobacteria
Firmicutes;Alk
Proteobacteria
Firmicutes;Alk
Deinococcus-Th
0.004
0.008
0.012
MeanDecreaseAccuracy
DILI_NS
DILI_S
H
HAV_NS
HAV_S
HEV_NS
HEV_S
OTHER_NS
OTHER_S
Low

**Supplementary Figure13 -** Heat map and mean decrease accuracy according to the etiology of the samples. **Figure 2A-** Unsupervised hierarchical clustering with different etiology showing metabolome signatures is capable to segregate ALF-Non-Survivor. The expression is given as Red; upregulated, Green; downregulated, and Black; unregulated ($p<0.05$; Kruskal-Wallis-test) **Figure 2B-** Mean Decrease in Accuracy of different etiologies showing top metabolome (Red; upregulated and blue; downregulated) **Figure 2C-** Unsupervised hierarchical clustering with DILI (Drug Induced liver injury) showing metabolome signatures is capable to segregate ALF-Non-Survivor. The expression is given as brown; upregulated, blue; downregulated, and white; unregulated ($p<0.05$; Kruskal-Wallis-test) **Figure 2D-** Mean Decrease in Accuracy of DILI (Drug Induced liver injury) showing top metabolome (Red; upregulated and blue; downregulated). **Figure 2E-** Unsupervised hierarchical clustering with HAV showing metabolome signatures is capable to segregate ALF-Non-Survivor. The expression is given as brown; upregulated, blue; downregulated, and white; unregulated ($p<0.05$; Kruskal-Wallis-test) **Figure 2F-** Mean Decrease in Accuracy HAV showing top metabolome (Red; upregulated and blue; downregulated). **Figure 2G-** Unsupervised hierarchical clustering with HEV showing metabolome signatures is capable to segregate ALF-Non-Survivor. The expression is given as brown; upregulated, blue; downregulated, and white; unregulated ($p<0.05$; Kruskal-Wallis-test) **Figure 2H-** Mean Decrease in Accuracy HEV showing top metabolome (Red; upregulated and blue; downregulated). **Figure 2G-** Unsupervised hierarchical clustering with other etiologies (Wilson disease, autoimmune and etc) showing metabolome signatures is capable to segregate ALF-Non-Survivor. The expression is given as brown; upregulated, blue; downregulated, and white; unregulated ($p<0.05$; Kruskal-Wallis-test) **Figure 2H-** Mean Decrease in Accuracy Others etiologies (Wilson disease, autoimmune and etc) showing top metabolome (Red; upregulated and blue; downregulated).

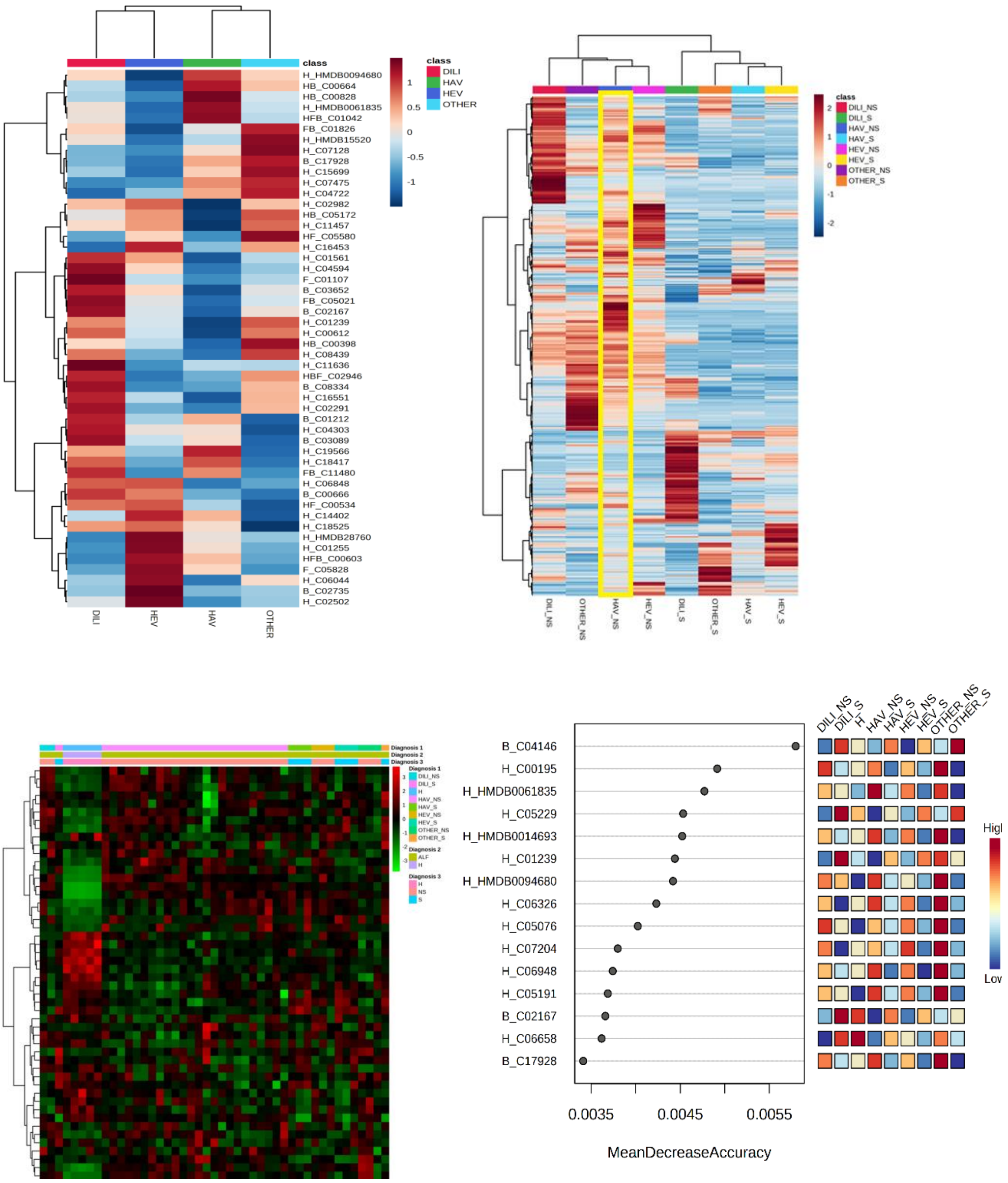
class
DILI
HAV
HEV
OTHER
H_HMDB0094680
HB_C00664
HB_C00828
H_HMDB0061835
HFB_C01042
FB_C01826
H_HMDB15520
H_C07128
B_C17928
H_C15699
H_C07475
H_C04722
H_C02982
HB_C05172
H_C11457
HF_C05580
H_C16453
H_C01561
H_C04594
F_C01107
B_C03652
FB_C05021
B_C02167
H_C01239
H_C00612
HB_C00398
H_C08439
H_C11636
HBF_C02946
B_C08334
H_C16551
H_C02291
B_C01212
H_C04303
B_C03089
H_C19566
H_C18417
FB_C11480
H_C06848
B_C00666
HF_C00534
H_C14402
H_C18525
H_HMDB28760
H_C01255
HFB_C00603
F_C05828
H_C06044
B_C02735
H_C02502
DILI
HEV
HAV
OTHER
class
DILI_NS
DILI_S
HAV_NS
HAV_S
HEV_NS
HEV_S
OTHER_NS
OTHER_S
Diagnosis 1
Diagnosis 2
Diagnosis 3
ALF
H
NS
S
B_C04146
H_C00195
H_HMDB0061835
H_C05229
H_HMDB0014693
H_C01239
H_HMDB0094680
H_C06326
H_C05076
H_C07204
H_C06948
H_C05191
B_C02167
H_C06658
B_C17928
0.0035
0.0045
0.0055
MeanDecreaseAccuracy
DILI_NS
DILI_S
H
HAV_NS
HAV_S
HEV_NS
HEV_S
OTHER_NS
OTHER_S
High
Low

**Supplementary Figure 14:** Top 5 metabolites correlation with the variability in the etiology of ALF (a) DILI (Drug Induced Liver Injury) (b) HAV (Hepatitis A Virus) (c) HEV (Hepatitis E virus) (d) Other etiologies (Wilson disease, autoimmune and etc)

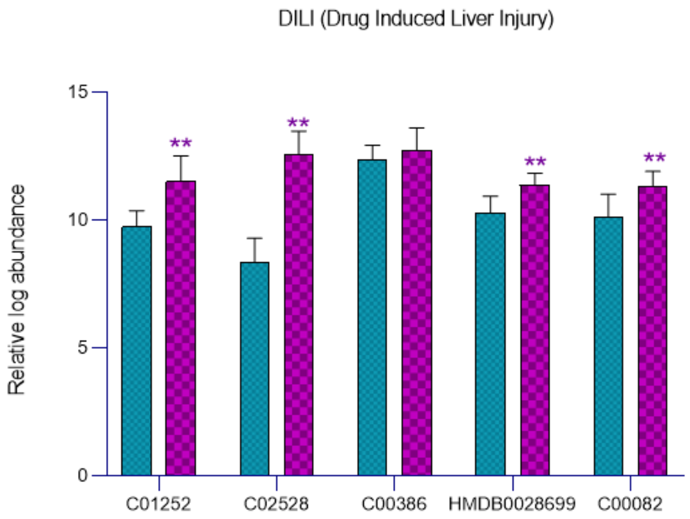


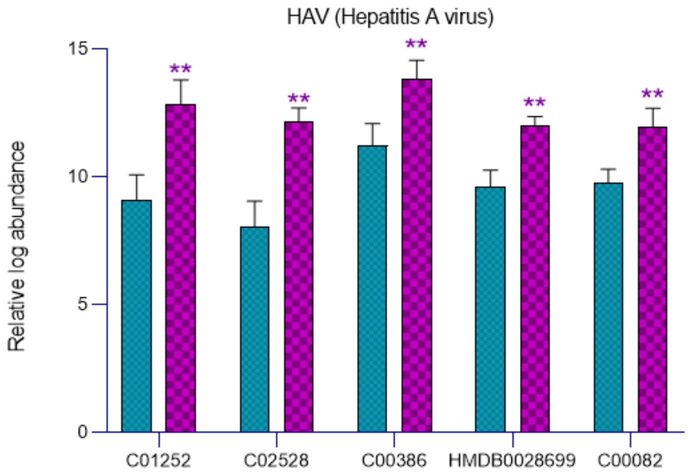

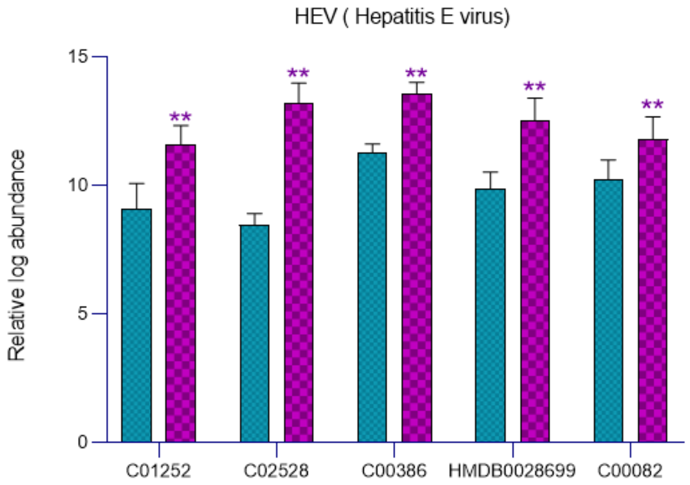
HEV ( Hepatitis E virus)
Relative log abundance
15
10
5
0
**
C01252
C02528
C00386
HMDB0028699
C00082

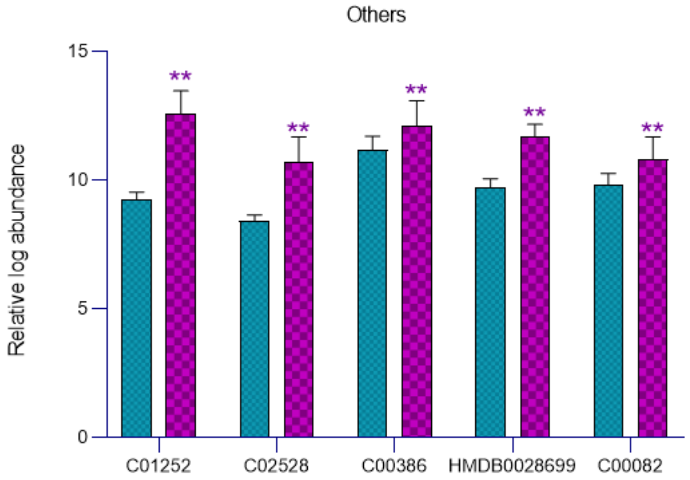
Others
Relative log abundance
15
10
5
0
**
C01252
C02528
C00386
HMDB0028699
C00082

**Supplementary Figure 15:** Quantitative Assessment of top 5 metabolite in Test-Cohort 2-Disease control (Baseline Plasma-Severe Alcoholic Hepatitis-SAH; n=70) with ALF training-Cohort (p<0.05; Mann-Whitney-U-test)

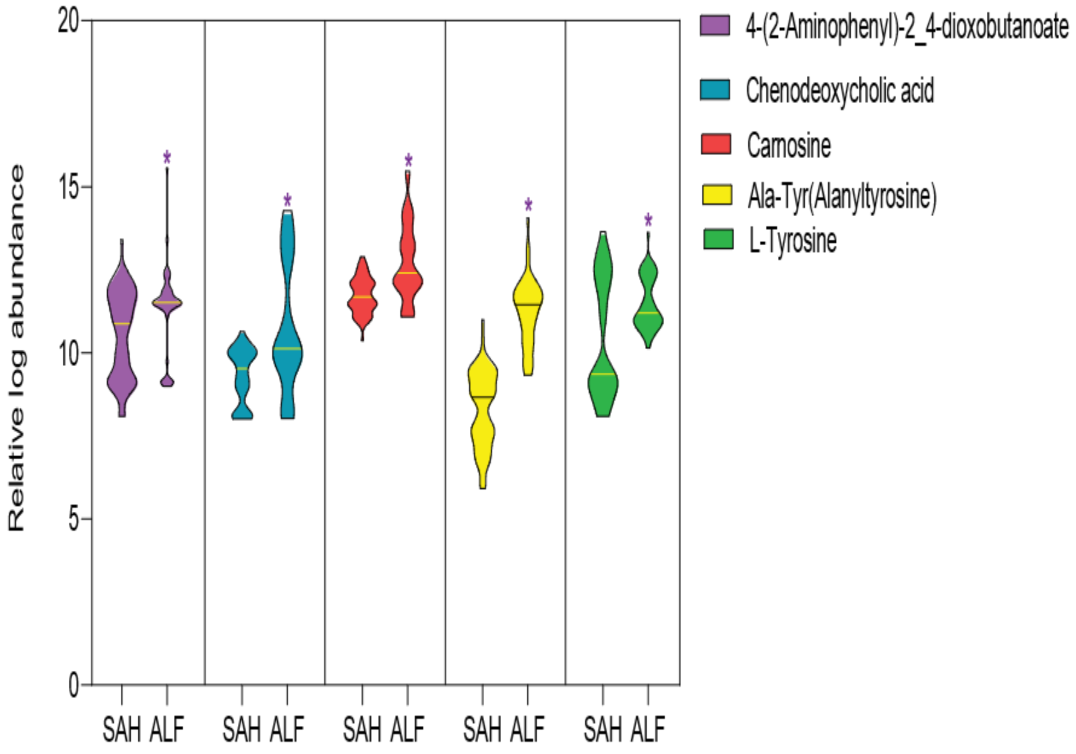

**Supplementary Figure 16:** Accuracy and kappa value of the different ML models for indicator identification in study group and Accuracy, specificity, sensitivity, and p value of five different metabolome species individually and altogether in test cohort plasma (100µl) and one drop blood along with confusion matrix for different ML models.

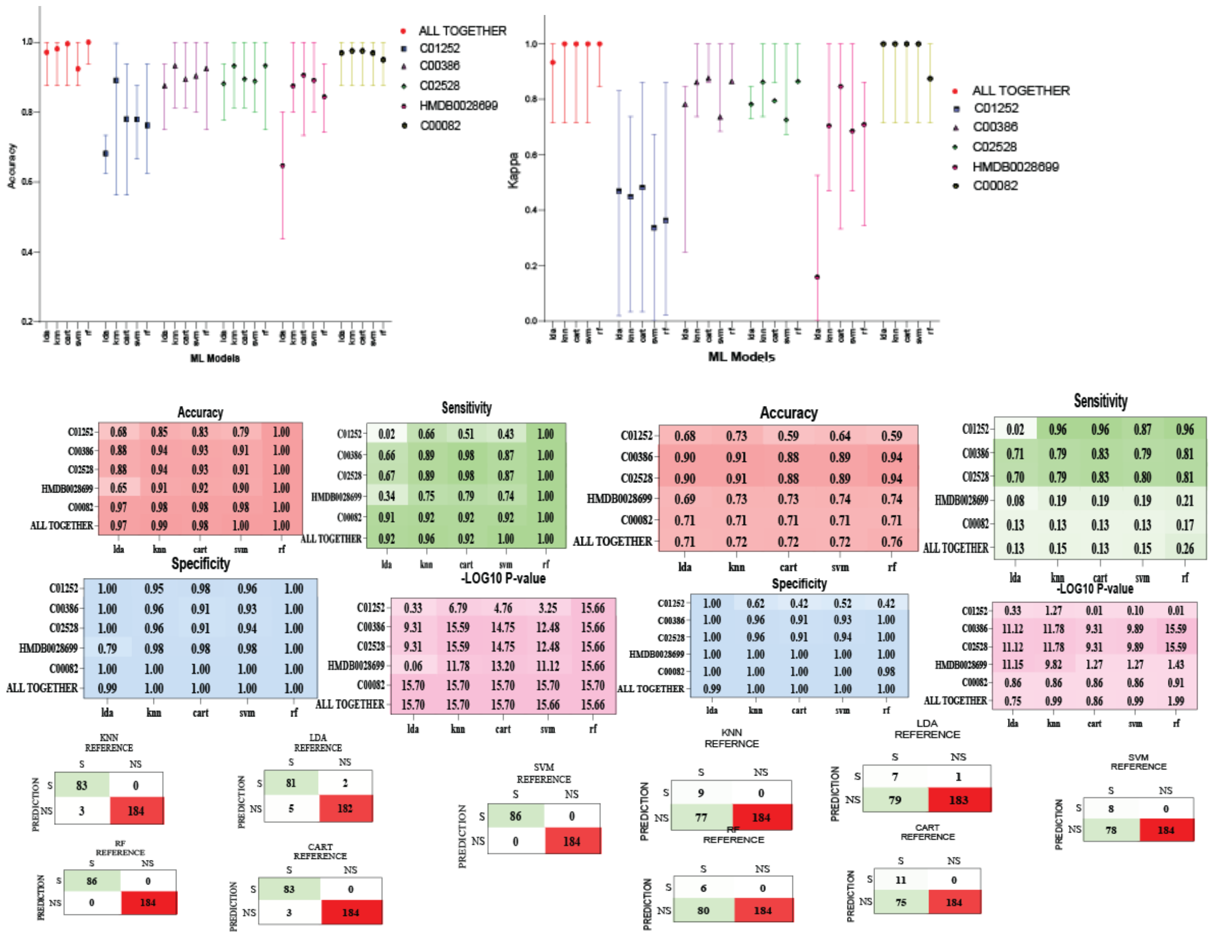

**Supplementary Figure 17:** Positive predictive value (PPV), Negative predictive value (NPV) and balanced accuracy for their respective models of top 5 metabolites and all together.

| Plasma (100µl) | | | | | | |
|---|---|---|---|---|---|---|
| | | KNN | LDA | SVM | RF | CART |
| **C00082** | Pos Pred Value | 1 | 1 | 1.00 | 1 | 1 |
| | Neg Pred Value | 0.964 | 0.9554 | 0.96 | 1 | 0.964 |
| | Balanced Accuracy | 0.9623 | 0.9528 | 0.96 | 1 | 0.9623 |
| **C00386** | Pos Pred Value | 0.9216 | 1 | 0.8679 | 1 | 0.8387 |
| | Neg Pred Value | 0.945 | 0.8492 | 0.9346 | 1 | 0.9898 |
| | Balanced Accuracy | 0.9247 | 0.8208 | 0.9013 | 1 | 0.9438 |
| **C01252** | Pos Pred Value | 0.8537 | 1 | 0.8519 | 1 | 0.9286 |
| | Neg Pred Value | 0.8487 | 0.673 | 0.7744 | 1 | 0.7955 |
| | Balanced Accuracy | 0.8022 | 0.5094 | 0.6983 | 1 | 0.7359 |
| **C02528** | Pos Pred Value | 0.9216 | 1 | 0.8679 | 1 | 0.8387 |
| | Neg Pred Value | 0.945 | 0.8492 | 0.9346 | 1 | 0.9898 |
| | Balanced Accuracy | 0.9247 | 0.8208 | 0.9013 | 1 | 0.9438 |
| **HMDB0028699** | Pos Pred Value | 0.9524 | 0.4286 | 0.9512 | 1 | 0.9545 |
| | Neg Pred Value | 0.8898 | 0.7034 | 0.8824 | 1 | 0.9052 |
| | Balanced Accuracy | 0.868 | 0.5577 | 0.8586 | 1 | 0.8869 |
| **ALL TOGETHER** | Pos Pred Value | 1 | 0.98 | 1 | 1 | 1 |
| | Neg Pred Value | 0.9817 | 0.9636 | 1 | 1 | 0.964 |
| | Balanced Accuracy | 0.9811 | 0.9576 | 1 | 1 | 0.9623 |

| One drop blood | | | | | | |
|---|---|---|---|---|---|---|
| | | KNN | LDA | SVM | RF | CART |
| **C00082** | Pos Pred Value | 1 | 1 | 1 | 0.818 | 1 |
| | Neg Pred Value | 0.69935 | 0.699 | 0.699 | 0.704 | 0.6994 |
| | Balanced Accuracy | 0.5664 | 0.56604 | 0.566 | 0.5755 | 0.5664 |
| **C00386** | Pos Pred Value | 0.913 | 1 | 0.8571 | 1 | 0.8148 |
| | Neg Pred Value | 0.9035 | 0.8699 | 0.9009 | 0.9145 | 0.9152 |
| | Balanced Accuracy | 0.8755 | 0.8491 | 0.8635 | 0.9057 | 0.8684 |
| **C01252** | Pos Pred Value | 0.5543 | 1 | 0.4742 | 0.4474 | 0.4474 |
| | Neg Pred Value | 0.9706 | 0.67296 | 0.8889 | 0.9565 | 0.9565 |
| | Balanced Accuracy | 0.7895 | 0.50943 | 0.6956 | 0.6867 | 0.6867 |
| **C02528** | Pos Pred Value | 0.913 | 1 | 0.8571 | 1 | 0.8148 |
| | Neg Pred Value | 0.9035 | 0.8699 | 0.9009 | 0.9145 | 0.9151 |
| | Balanced Accuracy | 0.8775 | 0.8491 | 0.8635 | 0.9057 | 0.8684 |
| **HMDB0028699** | Pos Pred Value | 1 | 1 | 1 | 1 | 1 |
| | Neg Pred Value | 0.7133 | 0.6859 | 0.7133 | 0.7181 | 0.7133 |
| | Balanced Accuracy | 0.5943 | 0.53774 | 0.5943 | 0.6038 | 0.5943 |
| **ALL TOGETHER** | Pos Pred Value | 1 | 0.875 | 1 | 1 | 1 |
| | Neg Pred Value | 0.7039 | 0.69737 | 0.7039 | 0.7329 | 0.6994 |
| | Balanced Accuracy | 0.5755 | 0.56136 | 0.5755 | 0.6321 | 0.566 |

**Supplementary Figure 18:** Univariate and multivariate cox regression analysis

| Univariate COX regression analysis | | | | | | | |
|---|---|---|---|---|---|---|---|
| | | | | | | 95.0% CI for Exp(B) | |
| | Hazard | SE | Wald | Sig. | Exp(Hazard) | Lower | Upper |
| B_C02528 | .674 | .141 | 22.980 | **.000** | 1.962 | 1.489 | 2.584 |
| H_C01252 | .618 | .172 | 12.986 | **.000** | 1.855 | 1.326 | 2.597 |
| H_HMDB0028699 | .453 | .094 | 23.135 | **.000** | 1.573 | 1.308 | 1.891 |
| HF_C00386 | .388 | .129 | 9.096 | **.003** | 1.474 | 1.145 | 1.896 |
| HFB_C00082 | .771 | .192 | 16.144 | **.000** | 2.161 | 1.484 | 3.147 |
| MELD | .075 | .024 | 10.202 | **.001** | 1.078 | 1.029 | 1.129 |
| TotalProtein | .088 | .195 | .205 | .651 | 1.092 | .746 | 1.600 |
| AlbuminGDL | -.346 | .290 | 1.428 | .232 | .707 | .401 | 1.248 |
| Bilirubin | .055 | .021 | 6.683 | **.010** | 1.056 | 1.013 | 1.101 |
| Alkalinephosphate | .005 | .002 | 3.955 | **.047** | 1.005 | 1.000 | 1.009 |
| AST | .000 | .000 | .115 | .734 | 1.000 | 1.000 | 1.000 |
| ALT | .000 | .000 | .032 | .859 | 1.000 | 1.000 | 1.000 |
| GGT | .002 | .006 | .063 | .801 | 1.002 | .990 | 1.014 |
| HB | .017 | .058 | .083 | .774 | 1.017 | .907 | 1.140 |
| Platelet | .000 | .002 | .021 | .884 | 1.000 | .997 | 1.004 |
| Neutophils | .038 | .017 | 4.714 | **.030** | 1.039 | 1.004 | 1.075 |
| Lymphocytes | -.042 | .021 | 3.832 | **.050** | .959 | .919 | 1.000 |
| Creatinine | .042 | .189 | .050 | .824 | 1.043 | .720 | 1.512 |
| Na | .069 | .034 | 4.139 | **.042** | 1.072 | 1.003 | 1.146 |
| INR | .212 | .059 | 12.731 | **.000** | 1.236 | 1.100 | 1.388 |

| Multivariate cox regression analysis | | | | | |
|---|---|---|---|---|---|
| | | | | 95.0% CI for Exp(B) | |
| | Hazard | Sig. | Exp(Hazard) | Lower | Upper |
| **B_C02528** | **.537** | **.026** | **1.710** | **1.067** | **2.741** |
| H_C01252 | .119 | .835 | 1.126 | .368 | 3.442 |
| H_HMDB0028699 | .406 | **.011** | 1.500 | 1.097 | 2.052 |
| HF_C00386 | -.099 | .709 | .906 | .538 | 1.525 |
| HFB_C00082 | .281 | .692 | 1.325 | .329 | 5.337 |
| MELD | .046 | .369 | 1.047 | .947 | 1.157 |
| TotalProtein | - | - | - | - | - |
| AlbuminGDL | - | - | - | - | - |
| Bilirubin | .055 | .114 | 1.057 | .987 | 1.131 |
| Alkalinephosphate | .009 | **.018** | 1.009 | 1.002 | 1.017 |
| AST | - | - | - | - | - |
| ALT | - | - | - | - | - |
| GGT | - | - | - | - | - |
| HB | - | - | - | - | - |
| Platelet | - | - | - | - | - |
| Neutophils | .082 | .436 | 1.086 | .883 | 1.334 |
| Lymphocytes | .010 | .937 | 1.010 | .793 | 1.285 |
| Creatinine | | - | | | |
| Na | .000 | .994 | 1.000 | .916 | 1.091 |
| INR | -.188 | .218 | .828 | .614 | 1.118 |

**Supplementary Figure 19:** Flow diagram for bacterial culture methodology

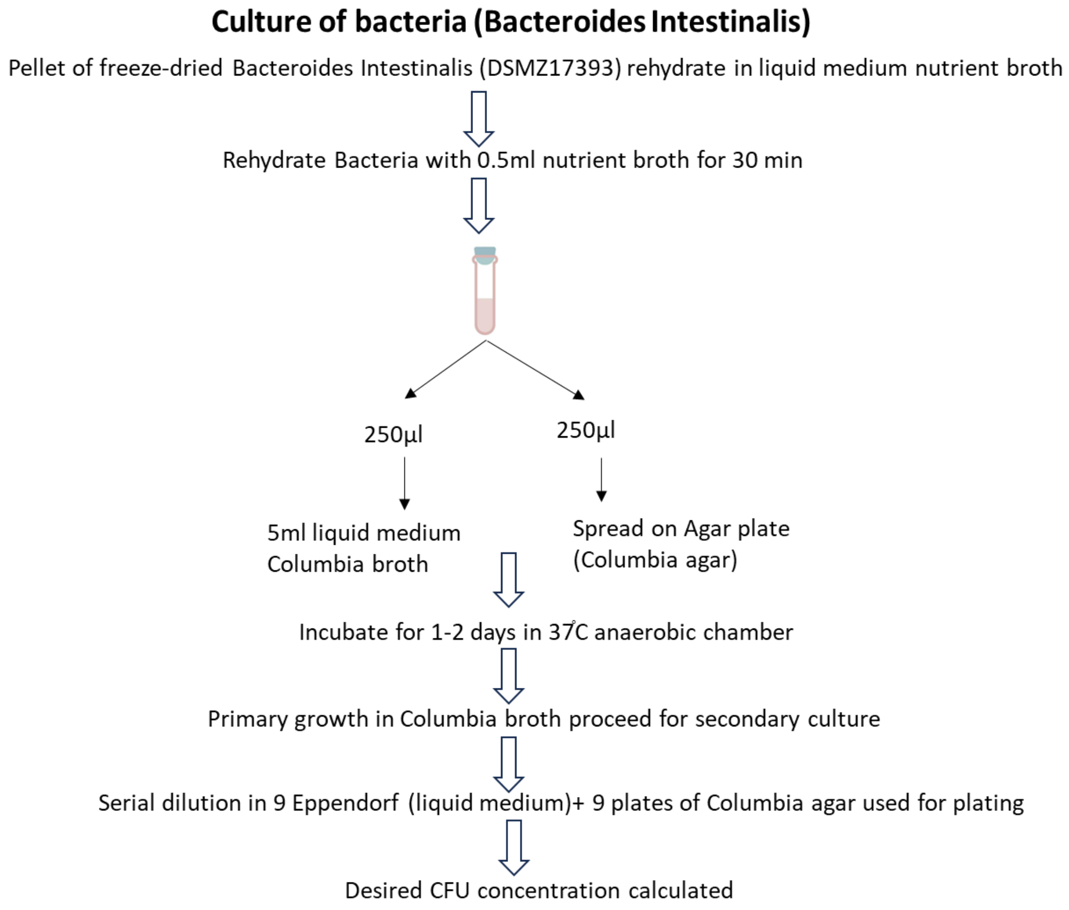

**Supplementary Figure 20:** Number of animals per group in animal model

| No. of animal | Treatment group |
|---|---|
| 10 | Control |
| 5 | APAP_250_24 |
| 5 | APAP_250_72 |
| 5 | APAP_500_24 |
| 5 | APAP_500_72 |
| 5 | BI+APAP_250_24 |
| 5 | BI+APAP_250_72 |
| 5 | BI+APAP_500_24 |
| 5 | BI+APAP_500_72 |
| 5 | APAP+BI_250_24 |
| 5 | APAP+BI_250_72 |
| 5 | APAP+BI_500_24 |
| 5 | APAP+BI_500_72 |

**Supplementary Figure 21:** Unsupervised hierarchical clustering with different group showing metabolome pathway of plasma is capable to segregate Treatment and ALF (250_72hrs) and control group in mice model

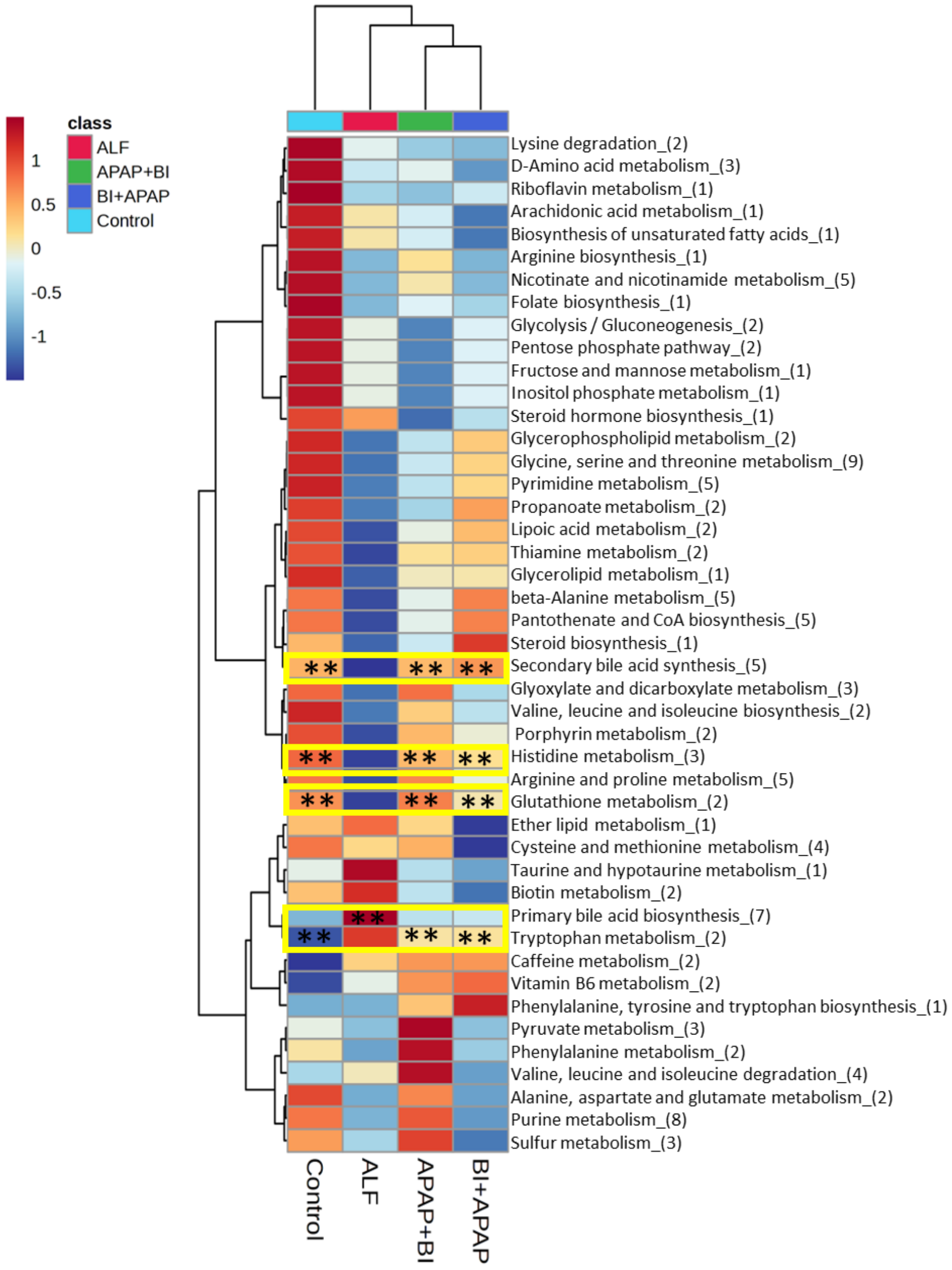

**Supplementary Figure 22:** Unsupervised hierarchical clustering with different group showing metabolome pathway of plasma is capable to segregate Treatment and ALF (500_24hrs) and control group in mice model

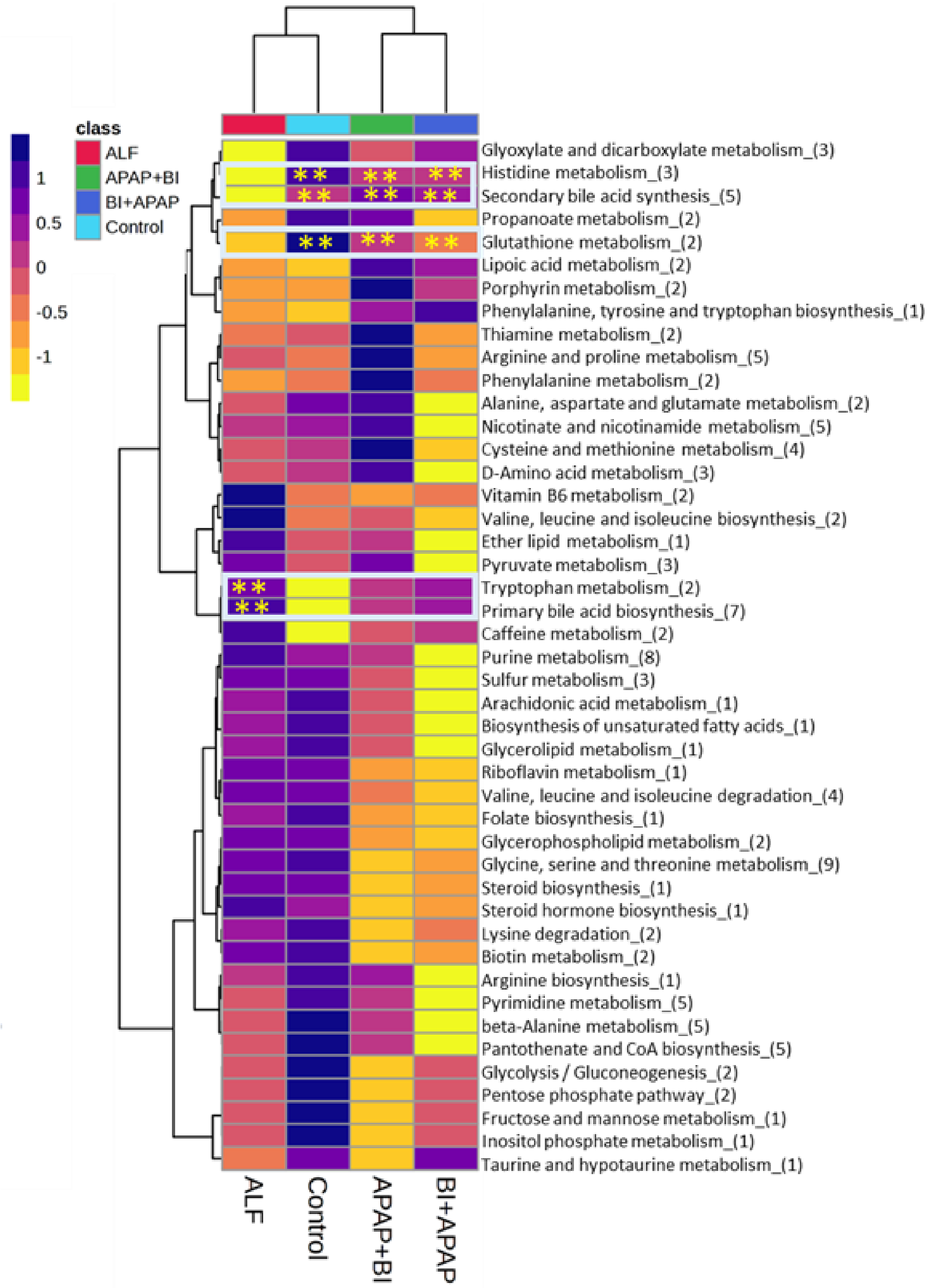

**Supplementary Figure 22:** Unsupervised hierarchical clustering with different group showing metabolome pathway of plasma is capable to segregate Treatment and ALF (500_72hrs) and control group in mice model

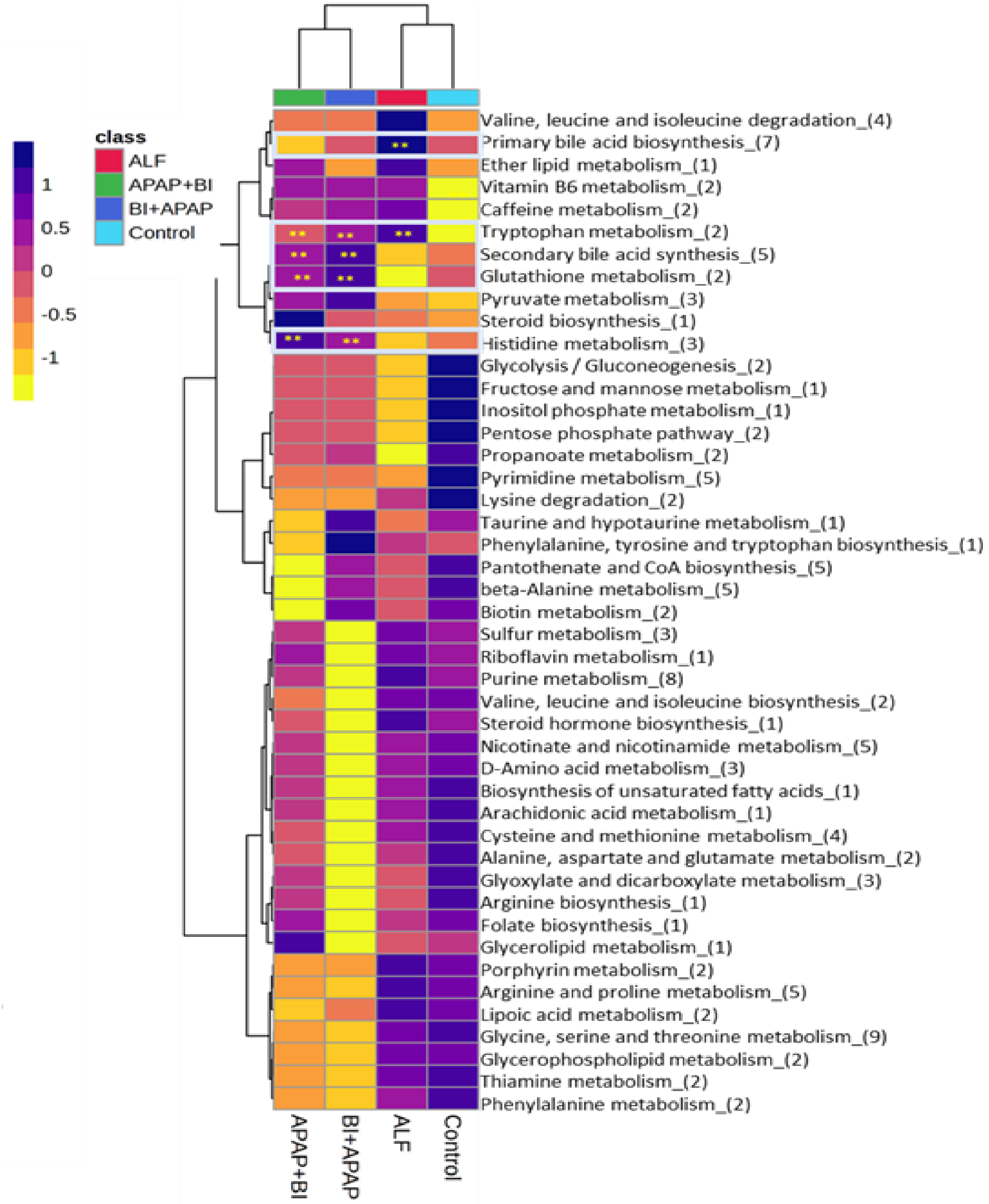

**Supplementary figure 23:** Unsupervised hierarchical clustering with different group showing metabolome pathway of liver is capable to segregate Treatment and ALF (250_72hrs) and control group in mice model

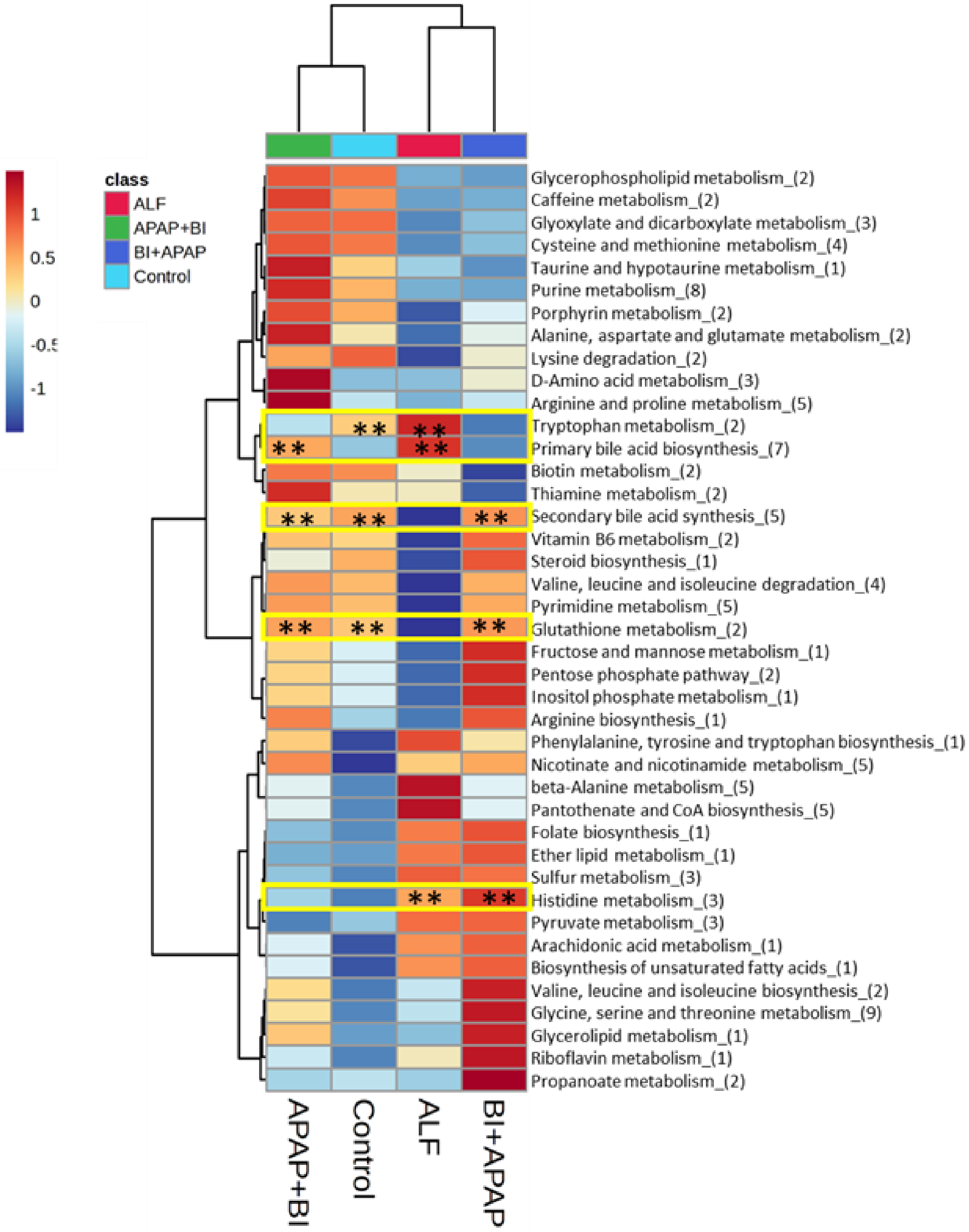

**Supplementary Figure 24:** Unsupervised hierarchical clustering with different group showing metabolome pathway of liver is capable to segregate Treatment and ALF (500_24hrs) and control group in mice model

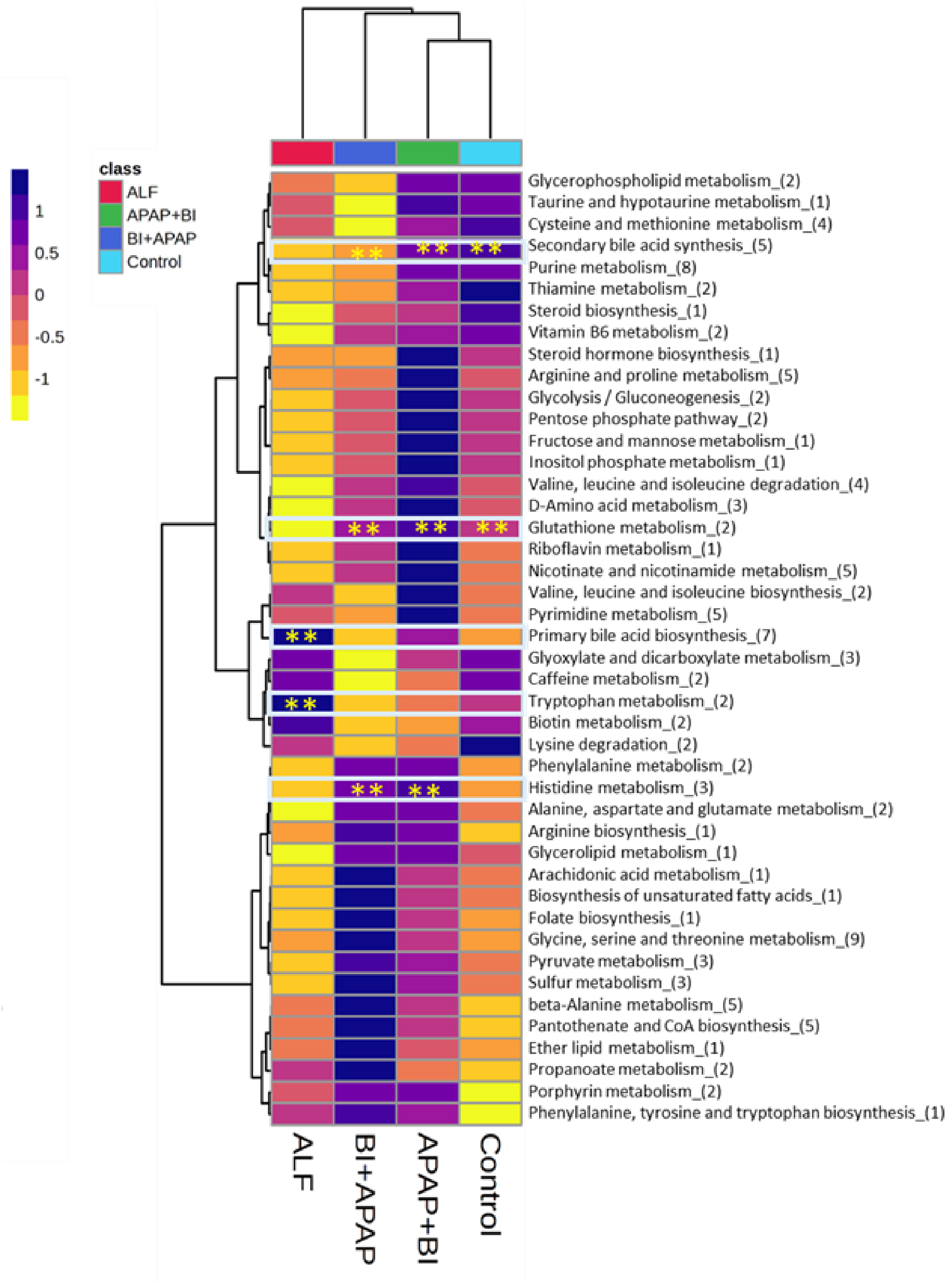

**Supplementary Figure 24:** Unsupervised hierarchical clustering with different group showing metabolome pathway of liver is capable to segregate Treatment and ALF (500_72hrs) and control group in mice model

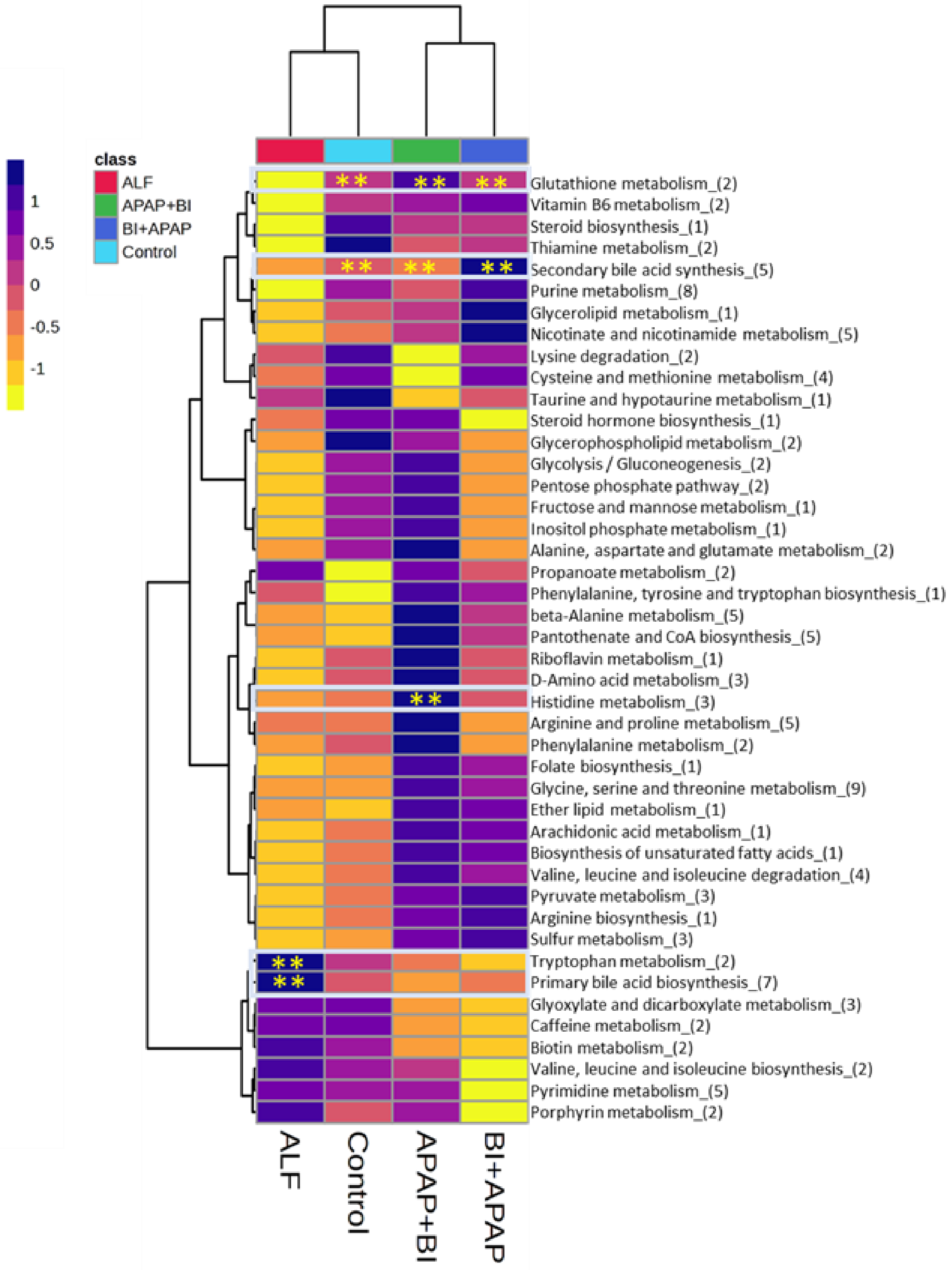

**Supplementary Figure 25:** Unsupervised hierarchical clustering with different group showing proteomics pathway of liver is capable to segregate Treatment and ALF (250_24hrs) and control group in mice model

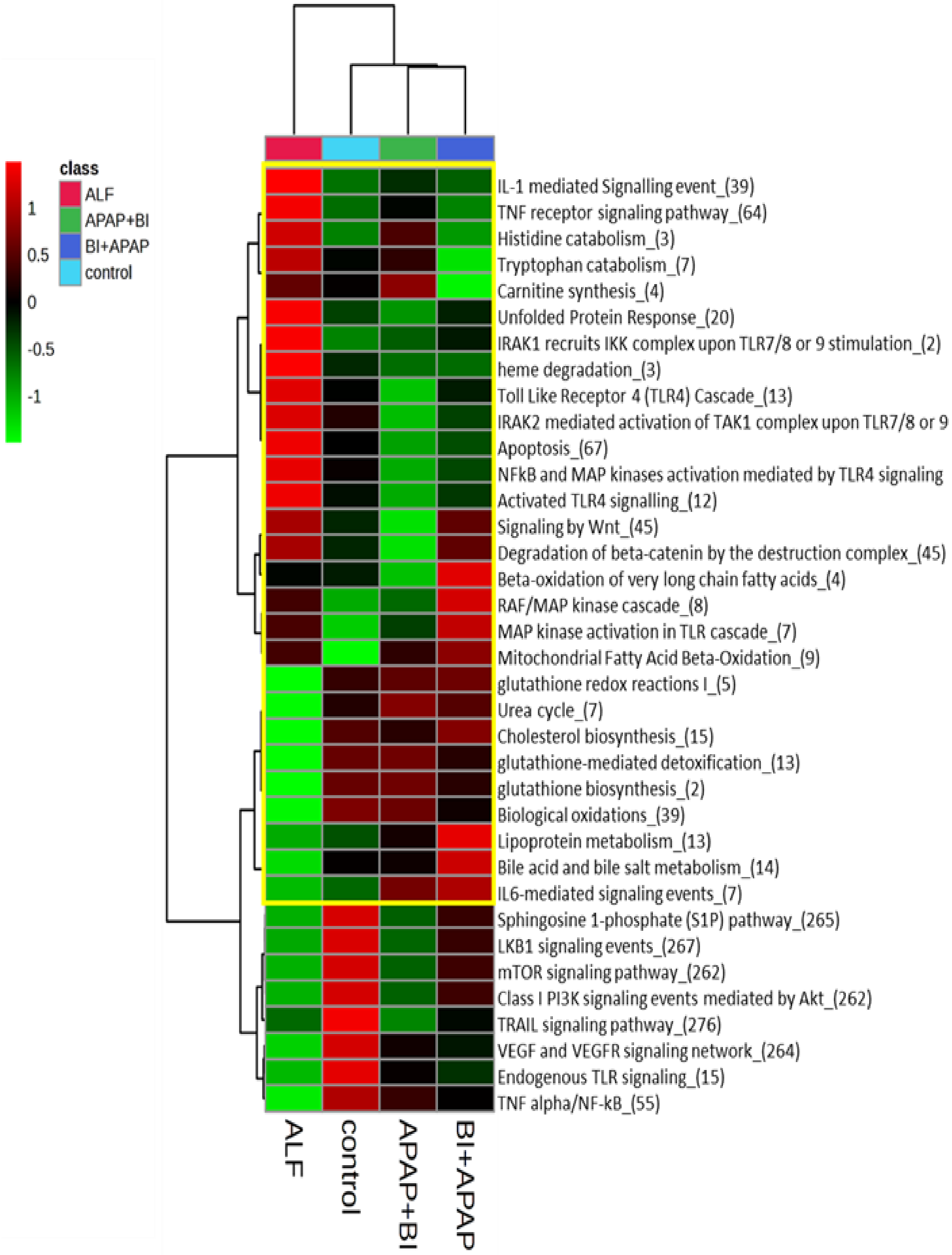

**Supplementary Figure 26:** Unsupervised hierarchical clustering with different group showing proteomics pathway of liver is capable to segregate Treatment and ALF (500_24hrs) and control group in mice model

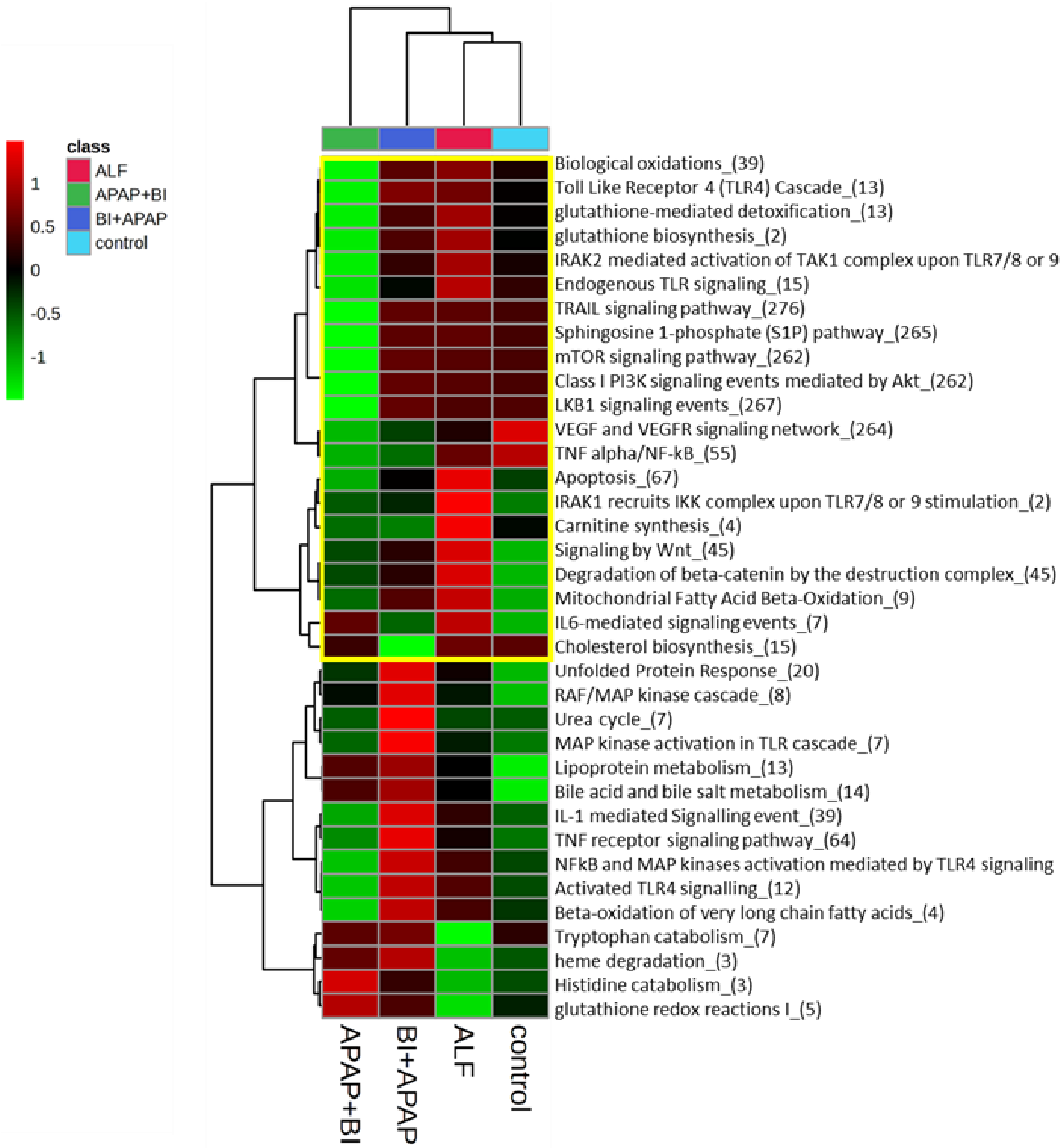

**Supplementary Figure 27:** Unsupervised hierarchical clustering with different group showing proteomics pathway of liver is capable to segregate Treatment and ALF (250_72hrs) and control group in mice model

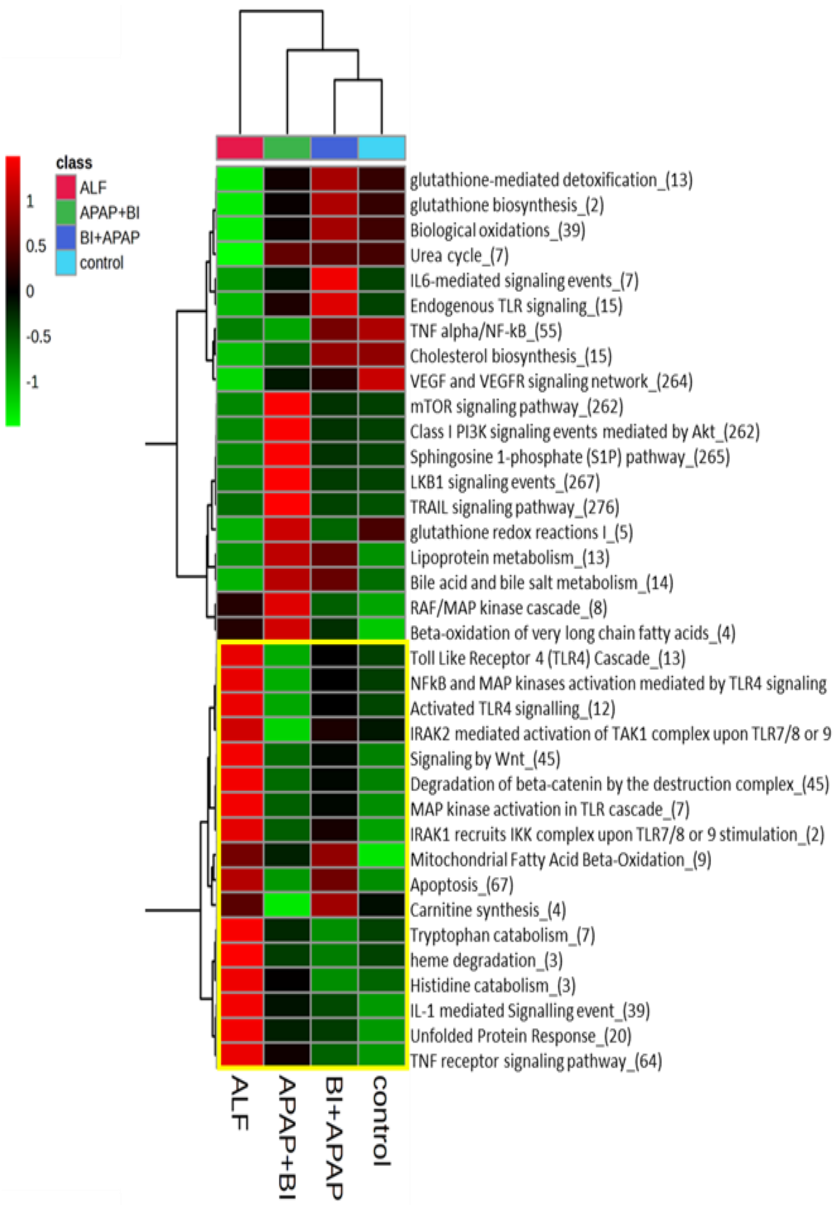

**Supplementary Figure 28:** MRNA Expression of different bile transported gene in different study groups in liver and intestine.

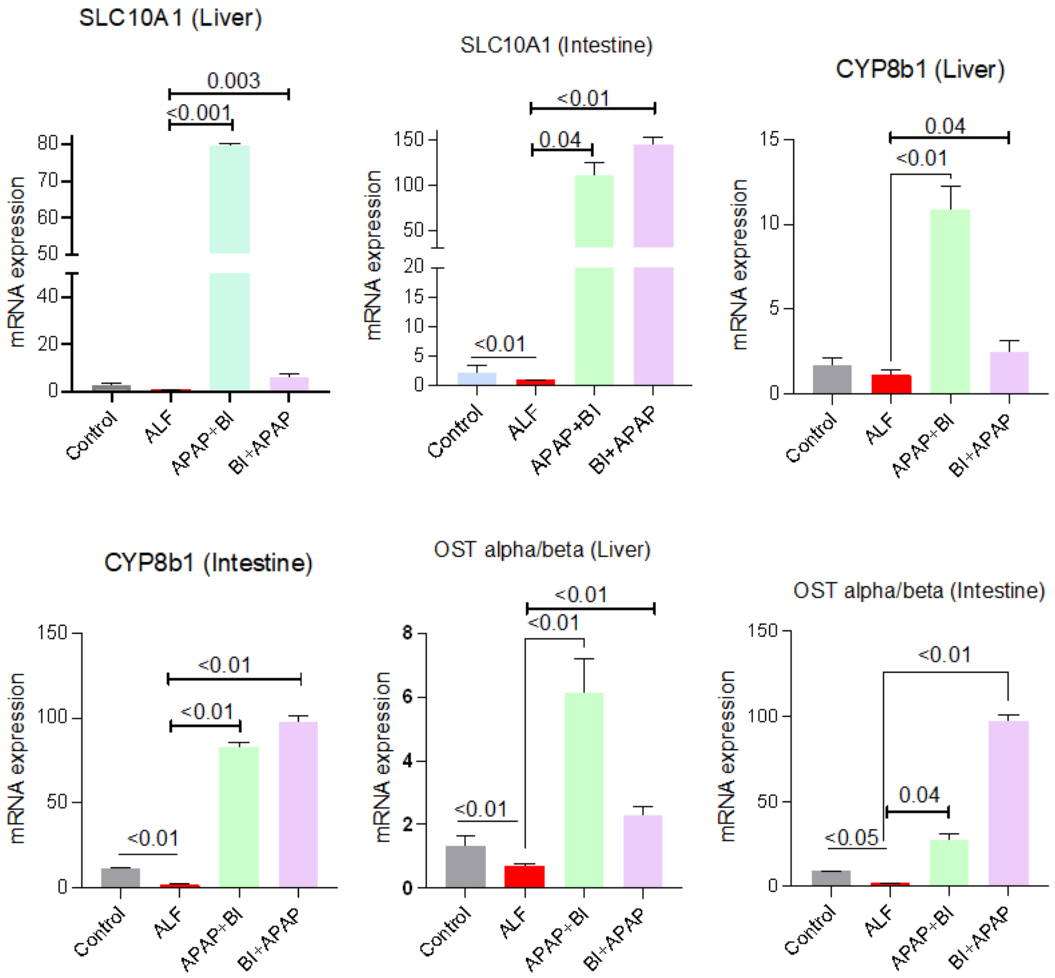
