## Supplementray Tables for "Chenodeoxycholic Acid Modulation via *Bacteroides intestinalis* AM1 underscores a Novel Approach in Acute Liver Failure"

**Supplementary table 1- Total metabolites identified in Acute liver failure Plasma samples in Healthy control , Survival ALF and Non-survival ALF**

| S.no | KEGG/HMDB ID | Type of Metabolite | Checked | Name | Molecular Weight | RT [min] | RT Tolerance [min] | Area (Max.) | # BioCyc Pathways | mzCloud Best Match | mzCloud Best Sim. Match | Identification status | Metabolika Pathways | Average Expression HEALTHY | Average Expression ALF | Average Expression ALF S | Average Expression ALF NS | FC (ALF VS HC) | pval (ALF VS HC) | Mean Decrease Accuracy (Random forest: (ALF VS HC) | AUC(ALF VS HC) | Pval AUC(ALF VS HC) | FC (NS VS S) | pval (NS VS S) | MeanDecrease Accuracy (Random Forrest (NS VS S) | AUC (NS VS S) | Pval (NS VS S) | FVALUE (HC VS NS VS S) | pval (HC VS NS VS S) | MeanDecrease Accuracy (Random Forrest (HC VS NS VS S) |
|---|---|---|---|---|---|---|---|---|---|---|---|---|---|---|---|---|---|---|---|---|---|---|---|---|---|---|---|---|---|---|
| **519** | HMDB0028699 | H | FALSE | Ala-Tyr(Alanyltyrosine) | 252.11 | 1.091 | 0.2 | 930507.3 | 0 | | | Spectral Match | | 10938.96 | 135516.5 | 184446.72 | 164783.9 | 4.1258 | 0.0096 | 0.000222 | 0.99 | 0.0001 | 5.4418 | 6.46E-07 | 0.002914 | 1 | 0.0000 | 6.9764 | 0.000529 | 0.00012 |
| **161** | C02528 | B | FALSE | Chenodeoxycholic acid | 392.2901 | 11.013 | 0.2 | 933425.5 | 0 | | | Spectral Match | | 161.4736 | 170797.5 | 4020.865 | 212491.6 | 349.1 | 0.0000 | 0.000819 | 1 | 0.0000 | 32.398 | 8.20E-05 | 0.004165 | 1 | 0.0000 | 19.181 | 1.21E-06 | 0.001792 |
| **122** | C01252 | H | FALSE | 4-(2-Aminophenyl)-2_4-dioxobutanoate | 207.0523 | 0.994 | 0.2 | 1852504 | 0 | | | Spectral Match | Other | 2444.682 | 258903.4 | 10024.95 | 321123 | 34.503 | 0.0000 | 0.001557 | 0.975 | 0.0000 | 19.203 | 8.35E-10 | 0.00194 | 1 | 0.0000 | 8.6899 | 0.000695 | 0 |
| **38** | C00386 | HF | FALSE | Carnosine | 225.0993 | 7.521 | 0.2 | 3761385 | 0 | | | Spectral Match | | 2534.769 | 699682.8 | 952662.94 | 850787.8 | 89.83 | 0.0000 | 0.001719 | 0.925 | 0.0000 | 5.3223 | 0.000898 | 0.002374 | 0.953125 | 0.0000 | 89.981 | 6.55E-16 | 0.00062 |

| | | | | | | | | | | | | | | | | | | | | | | | | | | | | | | |
|---|---|---|---|---|---|---|---|---|---|---|---|---|---|---|---|---|---|---|---|---|---|---|---|---|---|---|---|---|---|---|
| **8** | C00082 | HFB | FALSE | L-Tyrosine | 181.0734 | 7.259 | 0.2 | 718233.7 | 0 | | | Spectral Match | Superpathway of chorismate metabolism;Superpathway of aromatic amino acid biosynthesis;Other | 3204.714 | 112362.1 | 22440.42 | 134842.5 | 11.988 | 0.0000 | 0.001732 | 0.53 | 0.0000 | 3.7969 | 0.00049 | 0.000821 | 0.984375 | 0.0000 | 23.775 | 1.24E-07 | 0.000997 |
| **1** | C00009 | H | FALSE | Orthophosphate | 97.97707 | 1.121 | 0.116 | 35688795 | 0 | | 61 | Spectral Match | Superpathway of L-lysine degradation;Superpathway of chorismate metabolism;Superpathway of aromatic amino acid biosynthesis;Superpathway of L-threonine metabolism;Superpathway of NAD biosynthesis in eukaryotes;Anaerobic aromatic compound degradation (Thauera aromatica);Arginine, ornithine and proline interconversion;Superpathway of L-arginine and L-ornithine degradation;Superpathway of L-isoleucine biosynthesis I;Superpathway of butirocin biosynthesis;S | 1197347 | 1154930 | 1025 0762 | 11874097 | NS | NS | NS | NS | NS | NS | NS | NS | NS | NS | 5.2921 | 0.008918 | 0 |

| | | | | | | | | | | | | | | | | | | | | | | | | | | | | | | |
|---|---|---|---|---|---|---|---|---|---|---|---|---|---|---|---|---|---|---|---|---|---|---|---|---|---|---|---|---|---|---|
| | | | | | | | | | | | | | uperpathway of L-arginine, putrescine, and 4-aminobutanoate degradation;Biotin biosynthesis II;Adenosylcobalamin biosynthesis I (anaerobic);Other | | | | | | | | | | | | | | | | | |
| **2** | C00021 | HFB | FALSE | S-Adenosyl-L-homocysteine | 191.054 | 0.676 | 0.2 | 7452886.4 | 0 | | | Spectral Match | | 1051521.3 | 1306036.2 | 1182890.2 | 1337322.6 | 0.4655 | 0.0196 | 0 | 1 | 0.6034 | NS | NS | NS | NS | NS | 11.201 | 0.000126 | 0.000272 |
| **3** | C00025 | FB | FALSE | Glutamic Acid | 147.0527 | 0.918 | 0.2 | 20886364 | 0 | | | Spectral Match | Superpathway of L-lysine degradation;Superpathway of chorismate metabolism;Superpathway of aromatic amino acid biosynthesis;Superpathway of L-threonine metabolism;Superpathway of NAD biosynthesis in eukaryotes;Anaerobic aromatic compound degradation (Thauera aromatica);Arginine, ornithine and proline interconversio | 1470998 | 4994736 | 2727244 | 5561609 | NS | NS | NS | NS | NS | NS | NS | NS | NS | NS | 12.329 | 6.13E-05 | 0.000279 |

| | | | | | | | | | | | | | | | | | | | | | | | | | | | | | | |
|---|---|---|---|---|---|---|---|---|---|---|---|---|---|---|---|---|---|---|---|---|---|---|---|---|---|---|---|---|---|---|
| | | | | | | | | | | | | | n;Superpathway of indole-3-acetate conjugate biosynthesis;Superpathway of L-arginine and L-ornithine degradation;Superpathway of branched chain amino acid biosynthesis;Superpathway of L-isoleucine biosynthesis I;Superpathway of butirocin biosynthesis;Superpathway of L-arginine, putrescine, and 4-aminobutanoate degradation;Adenosylcobalamin biosynthesis I (anaerobic);Other | | | | | | | | | | | | | | | | | |
| 4 | C00037 | B | FALSE | ?-(Methylene cyclopropyl)glycine | 127.0631 | 0.979 | 0.2 | 58993776 | 0 | | | Spectral Match | Superpathway of L-lysine degradation | 1329448.7 | 1542272 | 1228357 | 1620751 | 4.0291 | 0.0023 | 0 | 0.965 | 0.0000 | NS | NS | NS | NS | NS | 5.6702 | 0.006608 | 0 |
| 5 | C00073 | FB | FALSE | L-Methionine | 149.0506 | 1.066 | 0.2 | 34425502 | 0 | | 85.3 | Spectral Match | Superpathway of butirocin biosynthesis;Biotin biosynthesis II;Other | 2615597.8 | 9059140 | 12830235 | 8116366 | 12.933 | 0.0001 | 0 | 0.985 | 0.0000 | NS | NS | NS | NS | NS | 30.491 | 6.61E-09 | 0.002346 |
| 6 | C00078 | HFB | FALS | L-Tryptophan | 204.0 | 1.3 | 0.2 | 3584 | 0 | | | Spectra | Superpathway of chorismate | 6020. | 6059 | 2432 | 6966 | 3.42 | 0.0 | 0 | 0.67 | 0.00 | NS | NS | NS | NS | NS | 9.0 | 0.00 | 0.00024 |

| | | | | | | | | | | | | | | | | | | | | | | | | | | | | | | |
|---|---|---|---|---|---|---|---|---|---|---|---|---|---|---|---|---|---|---|---|---|---|---|---|---|---|---|---|---|---|---|
| | | | E | | 891 | 68 | | 30.7 | | | | l Match | metabolism;Superpathway of aromatic amino acid biosynthesis;Superpathway of NAD biosynthesis in eukaryotes;Superpathway of aromatic compound degradation via 2-oxopent-4-enoate;Superpathway of aromatic compound degradation via 3-oxoadipate;Other | 947 | 3.73 | 8.01 | 0.16 | 69 | 001 | | | 00 | | | | | | 092 | 0555 | 4 |
| **7** | C00079 | FB | FALSE | 3-(2,6-Dioxocyclohexyl)propanenitrile | 165.0788 | 2.11 | 0.2 | 1473103 | 0 | 58.6 | | Spectral Match | Superpathway of chorismate metabolism;Superpathway of aromatic amino acid biosynthesis;Anaerobic aromatic compound degradation (Thauera aromatica);Superpathway of indole-3-acetate conjugate biosynthesis;Superpathway of jasmonoyl-amino acid conjugates biosynthesis;O | 25795.53 | 261037 | 72859.99 | 308081.3 | 3.2906 | 0.0194 | 0.000105 | 0.98 | 0.0002 | 2.4996 | 0.041094 | 0 | 0.875 | 0.0041 | 7.7446 | 0.001371 | 0.000435 |

| | | | | | | | | | | | | | | | | | | | | | | | | | | | | | | |
|---|---|---|---|---|---|---|---|---|---|---|---|---|---|---|---|---|---|---|---|---|---|---|---|---|---|---|---|---|---|---|
| | | | | | | | | | | | | | ther | | | | | | | | | | | | | | | | | |
| 9 | C00090 | B | FALSE | Catechol;1_2-Dihydroxybenzene | 110.0367 | 1.008 | 0.2 | 6224953 | 0 | | 90.2 | Spectral Match | Superpathway of aromatic compound degradation via 2-oxopent-4-enoate;Superpathway of aromatic compound degradation via 3-oxoadipate;Other | 153776.8 | 1517870 | 1212577 | 1594193 | 3.5225 | 0.0002 | 0 | 0.975 | 0.0000 | NS | NS | NS | NS | NS | 20.7558 | 5.38E-07 | 7.72E-05 |
| 10 | C00095 | HFB | FALSE | D-Fructose | 179.0563 | 4.527 | 0.2 | 12556648 | 0 | | | Spectral Match | | 5001.613 | 211750.6 | 450343.1 | 152102.4 | 16.986 | 0.0011 | 0.000132 | 1 | 0.0000 | NS | NS | NS | NS | NS | 6.9584 | 0.002454 | 0 |
| 11 | C00099 | H | FALSE | Beta-Alanine | 89.04794 | 0.93 | 0.2 | 19003758 | 0 | | | Spectral Match | Superpathway of L-lysine degradation;Superpathway of NAD biosynthesis in eukaryotes;Superpathway of aromatic compound degradation via 2-oxopent-4-enoate;Superpathway of aromatic compound degradation via 3-oxoadipate;Arginine, ornithine and proline interconversion;Superpathw | 138908.8 | 3007112 | 288001 | 3686889 | 6.6661 | 0.0039 | 0.000533 | 0.965 | 0.0002 | 7.1407 | 4.39E-06 | 0.000267 | 0.9375 | 0.0000 | 3.6794 | 0.033696 | 0 |

| | | | | | | | | | | | | | | | | | | | | | | | | | | | | | | |
|---|---|---|---|---|---|---|---|---|---|---|---|---|---|---|---|---|---|---|---|---|---|---|---|---|---|---|---|---|---|---|
| | | | | | | | | | | | | | ay of indole-3-acetate conjugate biosynthesis;Biotin biosynthesis II;Other | | | | | | | | | | | | | | | | | |
| 12 | C00110 | H | FALSE | Dolichyldiphosphate | 344.117 | 10.939 | 0.2 | 5678593 | 0 | | | Spectral Match | | 25212.01 | 31082.29 | 18388.21 | 34255.81 | 0.476 | 0.0000 | 0.000397 | 0.975 | 0.1735 | NS | NS | NS | NS | NS | 37.328 | 4.82E-10 | 0 |
| 13 | C00111 | H | FALSE | [Similar to: Tris(2,3-dibromopropyl) phosphate; ΔMass: 594.4915 Da] | 97.08932 | 2.913 | 0.2 | 868306 | 0 | | 71.2 | Spectral Match | | 3021.699 | 185078.7 | 298308 | 156771.4 | 24.608 | 0.0034 | 0.001898 | 0.75 | 0.0000 | NS | NS | NS | NS | NS | 8.802 | 0.000642 | 0.000472 |
| 14 | C00114 | FB | FALSE | Choline | 103.0999 | 0.729 | 0.075 | 14803529 | 0 | 82.2 | | Spectral Match | | 386993.7 | 3149257 | 3808524 | 2984440 | NS | NS | NS | NS | NS | 0.49095 | 0.016091 | 5.00E-04 | 0.664063 | 0.0633 | 73.369 | 1.97E-14 | 0.002562 |
| 15 | C00121 | HFB | FALSE | [Similar to: Isoprenaline; ΔMass: 60.0605 Da] | 151.0604 | 0.842 | 0.2 | 17887162 | 0 | | 89.9 | Spectral Match | | 664066.7 | 3854422 | 5721747 | 3387591 | NS | NS | NS | NS | NS | NS | NS | NS | NS | NS | 4.4574 | 0.017559 | 0 |
| 16 | C00135 | HFB | FALSE | L-Histidine | 154.0625 | 7.477 | 0.2 | 126944.7 | 0 | | | Spectral Match | | 25597.92 | 20944.09 | 21303.32 | 20854.28 | 0.33019 | 0.0000 | 0.000728 | 1 | 0.1337 | 0.64213 | 0.001008 | 0.000762 | 0.582031 | 0.8912 | 9.7399 | 0.000335 | 0.000118 |
| 17 | C00148 | HBF | FALSE | D-(+)-Proline | 115.0633 | 1.124 | 0.1 | 8947024 | 0 | 68.4 | | Spectral Match | Arginine, ornithine and proline interconversion;Other | 313074.7 | 1930077 | 874381 | 2194001 | NS | NS | NS | NS | NS | NS | NS | NS | NS | NS | 7.4949 | 0.001646 | -7.35E-06 |
| 18 | C00153 | FB | FALSE | Nicotinamide | 122.0479 | 0.383 | 0.2 | 1367417 | 0 | 59.8 | | Spectral Match | Superpathway of NAD biosynthesis in eukaryotes;Other | 114483.6 | 188717.5 | 208330.7 | 183814.2 | 0.6601 | 0.0247 | 0 | 0.96 | 0.0153 | 0.56849 | 0.002996 | 0.000308 | 0.683594 | 0.5096 | 8.2602 | 0.000944 | 0.000511 |

| | | | | | | | | | | | | | | | | | | | | | | | | | | | | | | |
|---|---|---|---|---|---|---|---|---|---|---|---|---|---|---|---|---|---|---|---|---|---|---|---|---|---|---|---|---|---|---|
| 19 | C00155 | FB | FALSE | Homocysteine | 135.035 | 1.022 | 0.2 | 3246176 | 0 | | 93.9 | Spectral Match | Other | 39174.24 | 652607.1 | 272226 | 747702.4 | 5.5889 | 0.0260 | 0 | 0.885 | 0.0004 | NS | NS | NS | NS | NS | 21.512 | 3.70E-07 | 0.000693 |
| 20 | C00195 | H | FALSE | [SP(20:0)]N-(eicosanoyl)-sphing-4-enine | 593.5719 | 18.947 | 0.2 | 2204515 | 0 | | | Spectral Match | | 18035.54 | 291794.1 | 26459.96 | 358127.7 | 6.1149 | 0.0028 | 0 | 1 | 0.0000 | 9.3256 | 3.10E-12 | 0.001453 | 1 | 0.0000 | 13.485 | 3.00E-05 | 0.000686 |
| 21 | C00212 | HFB | FALSE | Adenosine | 268.1044 | 1.072 | 0.2 | 405153.3 | 0 | | | Spectral Match | | 929955.35 | 91793.74 | 59376.34 | 998898.09 | 0.3755 | 0.0042 | 0 | 1 | 0.8929 | NS | NS | NS | NS | NS | 25.443 | 5.77E-08 | 0.001111 |
| 22 | C00221 | HFB | FALSE | beta-D-Glucose | 181.0706 | 0.681 | 0.2 | 2534381 | 0 | | | Spectral Match | | 270626.4 | 416920.9 | 287969.5 | 449158.7 | 0.56016 | 0.0118 | 0 | 0.695 | 0.2153 | NS | NS | NS | NS | NS | 11.772 | 8.73E-05 | 0.000375 |
| 23 | C00227 | HB | FALSE | Acetylphosphate | 139.9881 | 20.415 | 0.2 | 1000373 | 0 | | 66 | Spectral Match | Superpathway of L-lysine degradation;Other | 9109.378 | 31760.67 | 28257.88 | 32636.37 | NS | NS | NS | NS | NS | 0.53173 | 0.000477 | 0 | 0.675781 | 0.4888 | 4.2574 | 0.020723 | 0.000251 |
| 24 | C00242 | FB | FALSE | Guanine | 151.0491 | 1.068 | 0.2 | 708599.2 | 0 | | 67.3 | Spectral Match | Other | 9390.421 | 128081.8 | 62887.8 | 144380.3 | 4.724 | 0.0004 | 0.000235 | 0.98 | 0.0000 | NS | NS | NS | NS | NS | 6.9841 | 0.002407 | 0.000235 |
| 25 | C00253 | FB | FALSE | Nicotinicacid | 123.0317 | 1.057 | 0.2 | 916855.3 | 0 | | 94.1 | Spectral Match | Superpathway of NAD biosynthesis in eukaryotes;Adenosylcobalamin biosynthesis I (anaerobic);Other | 37716.99 | 181587 | 141894.8 | 191510.1 | 1.8023 | 0.0005 | 0 | 0.995 | 0.0000 | NS | NS | NS | NS | NS | 5.1546 | 0.009956 | 0 |
| 26 | C00257 | HFB | FALSE | D-Gluconic acid | 197.0657 | 1.382 | 0.2 | 473589 | 0 | | | Spectral Match | | 139702.2 | 70858.65 | 18094.54 | 84049.68 | 0.16449 | 0.0000 | 0 | 0.58 | 0.0925 | 2.8294 | 0.000461 | 0 | 0.964844 | 0.0000 | 3.9228 | 0.02742 | 0.000133 |
| 2 | C00 | H | FA | Benzaldehy | 10 | 1. | 0. | 93 | 0 | | 88 | Spe | Superpathway | 24 | 15 | 53 | 17 | 2. | 0. | 0. | 1 | 0. | 2. | 0. | 0.00 | 1 | 0. | 5 | 2. | 0.00 |

| | | | | | | | | | | | | | | | | | | | | | | | | | | | | | | |
|---|---|---|---|---|---|---|---|---|---|---|---|---|---|---|---|---|---|---|---|---|---|---|---|---|---|---|---|---|---|---|
| 7 | 261 | | LSE | de | 6.0419 | 119 | 2 | 0061.6 | | | .8 | ctral Match | of aromatic compound degradation via 2-oxopent-4-enoate;Superpathway of aromatic compound degradation via 3-oxoadipate;Other | 265.84 | 3366.1 | 119 | 8427.9 | 453 | 0057 | 000125 | | 0000 | 3333 | 000555 | 0595 | | 0000 | 4.701 | 02E-12 | 3944 |
| 28 | C00272 | HFB | FALSE | [Similar to: Methyl 3-[2-[(2,2-dichloroacetyl)amino]-4-(trifluoromethyl)phenoxy]thiophene-2-carboxylate; ΔMass: 186.8561 Da] | 240.1099 | 1.07 | 0.2 | 327594.7 | 0 | | 60.6 | Spectral Match | | 34053.93 | 59609.14 | 16314.83 | 70432.72 | 0.61836 | 0.02002 | 0 | 0.655 | 0.2581 | 2.6573 | 3.22E-07 | 2.80E-05 | 0.992188 | 0.00000 | 19.313 | 1.13E-06 | 0.00311 |
| 29 | C00295 | FB | FALSE | Oroticacid(vitaminB13) | 156.0171 | 0.886 | 0.2 | 33198506 | 0 | | | Spectral Match | Other | 5578926 | 7958417 | 6353544 | 8359636 | 0.51956 | 0.02992 | 0 | 0.99 | 0.6361 | NS | NS | NS | NS | NS | 7.8225 | 0.001295 | 1.11E-05 |
| 30 | C00299 | HFB | FALSE | [Similar to: Aflatoxin G2; ΔMass: 86.0037 Da] | 244.0703 | 11.18 | 0.2 | 17818777 | 0 | | 70.2 | Spectral Match | Other | 140916.8 | 274440.5 | 415537.2 | 239166.4 | NS | NS | NS | NS | NS | 0.28717 | 3.40E-10 | 0.000143 | 0.789063 | 0.0075 | 22.268 | 2.55E-07 | 0.000507 |
| 31 | C00300 | HB | FALSE | Creatine | 132.0767 | 5.985 | 0.2 | 1598726 | 0 | | | Spectral Match | | 141133.64 | 304813.4 | 263786 | 315070.3 | 8.2688 | 0.0063 | 0 | 1 | 0.0001 | NS | NS | NS | NS | NS | 11.461 | 0.000107 | 0.000479 |
| 32 | C00318 | HB | FALSE | DL-Carnitine | 161.104 | 0.88 | 0.089 | 403628 | 0 | 88.9 | | Spectral | | 162928 | 806228 | 132947 | 675417 | NS | NS | NS | NS | NS | 0.3301 | 0.0014 | 0 | 0.6718 | 0.09 | 6.59 | 0.0032 | 0 |

| | | | | | | | | | | | | | | | | | | | | | | | | | | | | | | |
|---|---|---|---|---|---|---|---|---|---|---|---|---|---|---|---|---|---|---|---|---|---|---|---|---|---|---|---|---|---|---|
| | | | | | 7 | 8 | | 72 | | | | Match | | 9 | 2 | 17 | 3 | | | | | | 9 | 23 | | 75 | 81 | 87 | 21 | |
| 33 | C00319 | F | FALSE | D-Sphingosine | 299.2812 | 11.906 | 0.054 | 9243665 | 0 | 72.1 | | Spectral Match | Other | 53091.47 | 114242.8 | 81283.31 | 122482.7 | NS | NS | NS | NS | NS | NS | NS | NS | NS | NS | 5.22251 | 0.009409 | 0 |
| 34 | C00327 | FB | FALSE | L(+)-Citrulline;2-Amino-5-uredovalerate | 175.0951 | 0.913 | 0.2 | 861824.8 | 0 | | 79.9 | Spectral Match | Arginine, ornithine and proline interconversion;Other | 442554.18 | 174956.5 | 154105.8 | 180169.2 | NS | NS | NS | NS | NS | NS | NS | NS | NS | NS | 8.65559 | 0.000712 | 0.000429 |
| 35 | C00328 | FB | FALSE | L-Kynurenine | 208.0841 | 1.136 | 0.2 | 1812094 | 0 | | | Spectral Match | Superpathway of NAD biosynthesis in eukaryotes;Superpathway of aromatic compound degradation via 2-oxopent-4-enoate;Superpathway of aromatic compound degradation via 3-oxoadipate;Other | 421351.5 | 288255.3 | 6269.211 | 358751.8 | 3.469 | 0.0023 | 0 | 0.65 | 0.5447 | 2.835 | 0.004284 | 0 | 0.703125 | 0.1455 | 4.6044 | 0.015559 | 0.000125 |
| 36 | C00334 | H | FALSE | gamma-Aminobutyric Acid | 103.0634 | 0.926 | 0.138 | 7701399 | 0 | | 92.6 | Spectral Match | Superpathway of L-arginine and L-ornithine degradation;Superpathway of L-arginine, putrescine, and 4-aminobutanoate degradation;Other | 121943.3 | 125383.8 | 10326.21 | 130914.2 | 3.62 | 0.0081 | 0 | 1 | 0.0000 | NS | NS | NS | NS | NS | 7.1494 | 0.002127 | 0.000286 |
| 37 | C00341 | HFB | FALS | Geranyl diphosphat | 158.0 | 7.1 | 0.2 | 2457 | 0 | | | Spectra | | 5634 | 4534 | 2087 | 5146 | NS | NS | NS | NS | NS | NS | NS | NS | NS | NS | 65. | 1.35 | 0.00064 |

| | | | | | | | | | | | | | | | | | | | | | | | | | | | | | | |
|---|---|---|---|---|---|---|---|---|---|---|---|---|---|---|---|---|---|---|---|---|---|---|---|---|---|---|---|---|---|---|
| | | | E | e | 417 | 09 | | 823 | | | | l Match | | 5.08 | 55 | 14.6 | 40.1 | | | | | | | | | | | 11 | E-13 | 2 |
| 39 | C00388 | B | FALSE | Histamine | 111.0794 | 1.076 | 0.2 | 3056393 | 0 | | | Spectral Match | | 31596.69 | 519682.7 | 20442.18 | 644492.8 | NS | NS | NS | NS | NS | 20.339 | 1.46E-06 | 0.002156 | 1 | 0.0000 | 48.727 | 1.14E-11 | 0.001138 |
| 40 | C00398 | HB | FALSE | Tryptamine | 79.0426 | 0.824 | 0.2 | 2141449 | 0 | | | Spectral Match | | 116861.9 | 176968.9 | 182703.2 | 175535.3 | 0.5999 | 0.0100 | 0 | 0.96 | 0.1509 | 0.64063 | 0.013586 | 0.000533 | 0.515625 | 0.7830 | 14.868 | 1.31E-05 | 0.000471 |
| 41 | C00401 | H | FALSE | Chondroitin | 378.1047 | 20.4 | 0.2 | 2619018 | 0 | | | Spectral Match | | 207172.8 | 253046.4 | 180519.5 | 271178.1 | 0.47354 | 0.0000 | 0.001219 | 0.99 | 0.0655 | NS | NS | NS | NS | NS | 21.402 | 3.90E-07 | 0.001627 |
| 42 | C00407 | FB | FALSE | L-Isoleucine | 131.0943 | 0.983 | 0.2 | 4955048 | 0 | | | Spectral Match | Superpathway of linamarin and lotaustralin biosynthesis;Superpathway of L-threonine metabolism;Superpathway of indole-3-acetate conjugate biosynthesis;Superpathway of jasmonoyl-amino acid conjugates biosynthesis;Superpathway of branched chain amino acid biosynthesis;Superpathway of L-isoleucine biosynthesis I | 33296.85 | 1051307 | 261864.4 | 1248668 | 11.157 | 0.0001 | 0.000706 | 0.59 | 0.0000 | NS | NS | NS | NS | NS | 22.233 | 2.60E-07 | 0.000404 |
| 4 | C00 | HB | FA | L- | 12 | 0. | 0. | 37 | 0 | 88 | | Spe | Superpathway | 41 | 96 | 44 | 10 | NS | N | NS | N | N | 0. | 0. | 0.00 | 0. | 0. | 4. | 0. | 0 |

| | | | | | | | | | | | | | | | | | | | | | | | | | | | | | | |
|---|---|---|---|---|---|---|---|---|---|---|---|---|---|---|---|---|---|---|---|---|---|---|---|---|---|---|---|---|---|---|
| 3 | 408 | F | LSE | Pipecolate | 9.0785 | 899 | 074 | 505012 | | .9 | | ctral Match | of L-lysine degradation | 797.18 | 598.46 | 413.23 | 9644.8 | | S | | S | S | 35163 | 001912 | 0133 | 75 | 0306 | 4648 | 017453 | |
| 44 | C00417 | HB | FALSE | cis-Aconitic acid | 88.01506 | 1.14 | 0.2 | 2417937 | 0 | | | Spectral Match | | 290359.8 | 348208.7 | 57141.43 | 420975.5 | NS | NS | NS | NS | NS | 4.4689 | 0.009779 | 0 | 0.902344 | 0.0008 | 16.414 | 5.41E-06 | 0.000133 |
| 45 | C00423 | B | FALSE | trans-Cinnamate | 148.052 | 1.589 | 0.2 | 1.11E+08 | 0 | 72.7 | | Spectral Match | Other | 1921159 | 20072501 | 4724087 | 23909605 | 3.3519 | 0.0156 | 0.000125 | 0.97 | 0.0003 | 2.9618 | 0.005119 | 0 | 0.929688 | 0.0010 | 17.434 | 3.07E-06 | 0.000154 |
| 46 | C00438 | HFB | FALSE | N-Carbamoyl-L-aspartate | 177.0508 | 0.79 | 0.2 | 1140568 | 0 | | | Spectral Match | | 67327.09 | 195746.9 | 125196.4 | 213384.6 | NS | NS | NS | NS | NS | NS | NS | NS | NS | NS | 12.807 | 4.54E-05 | 0.001154 |
| 47 | C00448 | HFB | FALSE | Farnesyl diphosphate | 190.0587 | 5.077 | 0.2 | 660119 | 0 | | | Spectral Match | | 96519.87 | 145732.3 | 125297.3 | 150841 | 0.57877 | 0.0113 | 0 | 0.69 | 0.1780 | NS | NS | NS | NS | NS | 11.013 | 0.000143 | 0.001102 |
| 48 | C00451 | FB | FALSE | (1R_2S)-1-Hydroxypropane-1_2_3-tricarboxylate | 192.0263 | 1.07 | 0.2 | 75074511 | 0 | | 90.6 | Spectral Match | Other | 268662 | 12158298 | 4624669.9 | 15082255 | NS | NS | NS | NS | NS | 18.892 | 0.003686 | -0.00013 | 0.921875 | 0.0006 | 17.54 | 2.90E-06 | 0.000712 |
| 49 | C00466 | FB | FALSE | (R)-Acetoin | 88.0526 | 14.349 | 0.2 | 1417072 | 0 | | | Spectral Match | Superpathway of L-lysine degradation;Other | 553448.26 | 470883.5 | 444667.12 | 474736.5 | 0.33531 | 0.0000 | 0.001879 | 0.975 | 0.0134 | NS | NS | NS | NS | NS | 90.725 | 5.69E-16 | 0.000526 |
| 50 | C00475 | HFB | FALSE | Cytidine | 242.0784 | 10.927 | 0.2 | 3857975 | 0 | | | Spectral Match | | 58423.74 | 33907.25 | 33188.27 | 34087 | 0.23249 | 0.0000 | 0.002323 | 0.69 | 0.0000 | NS | NS | NS | NS | NS | 3.7152 | 0.032685 | 0.001553 |
| 51 | C00483 | H | FALSE | Tyramine | 137.0814 | 1.026 | 0.2 | 242077.6 | 0 | | 66.1 | Spectral Match | | 3569.985 | 12401.19 | 6644.067 | 13840.47 | NS | NS | NS | NS | NS | NS | NS | NS | NS | NS | 76.564 | 9.81E-15 | 0.005114 |

| | | | | | | | | | | | | | | | | | | | | | | | | | | | | | | |
|---|---|---|---|---|---|---|---|---|---|---|---|---|---|---|---|---|---|---|---|---|---|---|---|---|---|---|---|---|---|---|
| **52** | C00491 | FB | FALSE | L-Dicysteine | 81.01552 | 10.943 | 0.2 | 34035567 | 0 | | | Spectral Match | | 42493.15 | 35011.39 | 31287.92 | 35942.26 | 0.32482 | 0.0000 | 0.001715 | 0.815 | 0.0028 | NS | NS | NS | NS | NS | 41.857 | 1.00E-10 | 0.006129 |
| **53** | C00502 | B | FALSE | D-Xylonate | 167.0554 | 0.828 | 0.2 | 972561.1 | 0 | | | Spectral Match | | 139099.57 | 218815.3 | 194856.9 | 224804.9 | 5.5688 | 0.0000 | 0.001552 | 1 | 0.0000 | NS | NS | NS | NS | NS | 46.032 | 2.60E-11 | 0.001206 |
| **54** | C00510 | H | FALSE | [SP(18:0)]N-(9Z-octadecenoyl)-sphing-4-enine | 563.5253 | 18.009 | 0.2 | 673735.2 | 0 | | | Spectral Match | | 11991.73 | 93063.93 | 11851.26 | 113367.1 | NS | NS | NS | NS | NS | 7.0155 | 5.37E-07 | 0.002549 | 1 | 0.0000 | 27.871 | 1.98E-08 | 0.000656 |
| **55** | C00534 | HF | FALSE | Pyridoxamine | 83.03746 | 1.069 | 0.2 | 9476277 | 0 | | | Spectral Match | | 1408033 | 1421553 | 2569957 | 1134452 | 2.6019 | 0.0359 | 0 | 1 | 0.6138 | 2.938 | 0.001036 | 0.001972 | 0.898438 | 0.0028 | 22.205 | 2.63E-07 | 0.002031 |
| **56** | C00547 | FB | FALSE | Norepinephrine | 169.0734 | 1.675 | 0.2 | 345343.9 | 0 | | | Spectral Match | Other | 15078.06 | 68815.94 | 32357.72 | 77930.49 | NS | NS | NS | NS | NS | NS | NS | NS | NS | NS | 23.776 | 1.24E-07 | 0.003579 |
| **57** | C00552 | B | FALSE | meso-Tartaric acid | 151.024 | 0.89 | 0.2 | 709991.2 | 0 | | | Spectral Match | | 34950.63 | 187923.6 | 145988.6 | 198407.4 | NS | NS | NS | NS | NS | NS | NS | NS | NS | NS | 42.668 | 7.66E-11 | 0.00104 |
| **58** | C00565 | H | FALSE | N-Dimethylethanolamine | 89.08426 | 0.888 | 0.2 | 370740.2 | 0 | | | Spectral Match | Other | 45618.82 | 55134.45 | 39396.92 | 59068.83 | 0.4602 | 0.0000 | 0 | 0.785 | 0.3770 | NS | NS | NS | NS | NS | 51.484 | 5.03E-12 | 0.000695 |
| **59** | C00581 | H | FALSE | Guanidinoacetate | 118.0611 | 1.605 | 0.2 | 2554589 | 0 | | | Spectral Match | | 27987.22 | 388105.8 | 165634.4 | 443723.7 | NS | NS | NS | NS | NS | NS | NS | NS | NS | NS | 82.342 | 2.93E-15 | 0.002134 |
| **60** | C00583 | H | FALSE | Propane-1_2-diol1-phosphate | 156.0184 | 15.195 | 0.2 | 97448.37 | 0 | | | Spectral Match | | 8171.925 | 6277.851 | 5031.627 | 6589.408 | 0.30873 | 0.0000 | 0.00025 | 0.66 | 0.2351 | NS | NS | NS | NS | NS | 28.295 | 1.65E-08 | 0 |

| | | | | | | | | | | | | | | | | | | | | | | | | | | | | | | |
|---|---|---|---|---|---|---|---|---|---|---|---|---|---|---|---|---|---|---|---|---|---|---|---|---|---|---|---|---|---|---|
| **61** | C00601 | FB | FALSE | Phenylacetaldehyde | 120.0574 | 1.098 | 0.2 | 7492219.7 | 0 | | 77.1 | Spectral Match | Anaerobic aromatic compound degradation (Thauera aromatica);Other | 304432.93 | 878898.86 | 655537.28 | 934849.26 | NS | NS | NS | NS | NS | NS | NS | NS | NS | NS | 5.8531 | 0.005724 | 0 |
| **62** | C00603 | HFB | FALSE | [Similar to: 4-Hydroxyphenylacetic acid; ΔMass: 17.0068 Da] | 135.0405 | 0.853 | 0.2 | 11732952 | 0 | | 92.8 | Spectral Match | | 628033 | 2408257 | 3503662 | 2134406 | NS | NS | NS | NS | NS | 0.38895 | 0.018631 | 0 | 0.65625 | 0.2522 | 4.3474 | 0.019231 | 0 |
| **63** | C00612 | H | FALSE | N1-Acetylspermidine | 187.1679 | 0.74 | 0.2 | 58252241 | 0 | | | Spectral Match | | 259421.9 | 561866 | 507378.2 | 575488 | NS | NS | NS | NS | NS | NS | NS | NS | NS | NS | 9.9316 | 0.000294 | 0.000267 |
| **64** | C00621 | HF | FALSE | [Similar to: Amiloride; ΔMass: 57.9976 Da] | 171.0503 | 1.593 | 0.2 | 3576193 | 0 | | 83.3 | Spectral Match | | 274428.21 | 50350.5 | 598660.76 | 479729.24 | NS | NS | NS | NS | NS | 1.5719 | 0.04156 | 0.000333 | 0.777344 | 0.05002 | 8.3928 | 0.000858 | 0.000251 |
| **65** | C00633 | B | FALSE | 4-Hydroxybenzaldehyde | 122.0366 | 1.074 | 0.2 | 1057743 | 0 | | | Spectral Match | Superpathway of aromatic compound degradation via 2-oxopent-4-enoate;Anaerobic aromatic compound degradation (Thauera aromatica);Superpathway of aromatic compound degradation via 3-oxoadipate;Other | 402667.03 | 248618 | 1086647.3 | 283610.6 | NS | NS | NS | NS | NS | NS | NS | NS | NS | NS | 4.2028 | 0.021685 | 0 |
| **6** | C00 | H | FA | 4- | 15 | 1. | 0. | 11 | 0 | | | Spe | | 37 | 23 | 25 | 22 | 0. | 0. | 0. | 0. | 0. | 0. | 0. | 0 | 0. | 0. | 4. | 0. | 0 |

| | | | | | | | | | | | | | | | | | | | | | | | | | | | | | | |
|---|---|---|---|---|---|---|---|---|---|---|---|---|---|---|---|---|---|---|---|---|---|---|---|---|---|---|---|---|---|---|
| 6 | 651 | | LSE | Methylene-L-glutamate | 9.0527 | 004 | 2 | 61935 | | | | ctral Match | | 1067 | 4755.8 | 7361.6 | 9104.3 | 24952 | 0002 | 000125 | 64 | 1524 | 5532 | 043571 | | 59375 | 5574 | 9437 | 011801 | |
| 67 | C00664 | HB | FALSE | 5-Formiminotetrahydrofolate | 237.0983 | 7.752 | 0.2 | 780662.6 | 0 | | | Spectral Match | | 6443.976 | 116516.2 | 18261.85 | 141079.8 | 5.6352 | 0.0017 | 0 | 0.785 | 0.0000 | 4.2754 | 0.000159 | 0.000375 | 0.960938 | 0.0000 | 4.0319 | 0.025018 | 0 |
| 68 | C00666 | B | FALSE | 2_6-Diaminoheptanedioicacid | 190.0947 | 0.962 | 0.2 | 1794914 | 0 | | | Spectral Match | Other | 520343.3 | 127743.8 | 106932.5 | 132946.6 | 0.091323 | 0.0000 | 0.000444 | 1 | 0.0000 | NS | NS | NS | NS | NS | 6.1275 | 0.004624 | 0 |
| 69 | C00670 | FB | FALSE | sn-glycero-3-Phosphocholine | 257.1019 | 0.935 | 0.2 | 5173067 | 0 | | 64.5 | Spectral Match | | 116856.4 | 1266618 | 1225683 | 1276852 | 4.345 | 0.0381 | 0 | 0.865 | 0.0002 | NS | NS | NS | NS | NS | 13.888 | 2.35E-05 | 0.001287 |
| 70 | C00706 | H | FALSE | 4-Piperidinecarboxamide | 128.0947 | 3.497 | 0.2 | 829115.9 | 0 | 52.2 | 78.6 | Spectral Match | | 32845.18 | 146048.9 | 79309.88 | 162733.7 | 1.6997 | 0.0470 | 0 | 0.68 | 0.0000 | NS | NS | NS | NS | NS | 8.5732 | 0.000755 | 0 |
| 71 | C00712 | H | FALSE | (6Z)-Octadecenoicacid | 282.2547 | 15.797 | 0.2 | 174255.5 | 0 | | | Spectral Match | Other | 16692.42 | 13434.41 | 19754.53 | 11854.38 | 0.33652 | 0.0000 | 0.002 | 0.975 | 0.0693 | 0.40518 | 9.25E-10 | 0.001251 | 0.933594 | 0.0000 | 3.7637 | 0.031366 | 0 |
| 72 | C00719 | H | FALSE | Betaine | 117.078 | 0.803 | 0.2 | 154481.4 | 0 | | | Spectral Match | | 2494.538 | 37256.56 | 33531.1 | 38187.92 | 5.6255 | 0.0401 | 0 | 0.805 | 0.0015 | NS | NS | NS | NS | NS | 16.61 | 4.85E-06 | 0.000635 |
| 73 | C00729 | H | FALSE | Tropine | 141.115 | 0.944 | 0.2 | 46655021 | 0 | 87.7 | | Spectral Match | Other | 12976232 | 20765196 | 14200405 | 22406394 | NS | NS | NS | NS | NS | NS | NS | NS | NS | NS | 54.765 | 1.99E-12 | 0.00429 |
| 74 | C00735 | H | FALSE | Cortisol | 361.2025 | 15.013 | 0.2 | 1248060 | 0 | | | Spectral Match | | 13894.44 | 16082.07 | 21524.59 | 14721.44 | 0.4731 | 0.0001 | 0 | 0.975 | 0.3686 | 0.46378 | 6.87E-08 | 0.001879 | 0.875 | 0.0004 | 6.3415 | 0.003921 | 0.001244 |
| 7 | C00 | FB | FA | D-Serine | 10 | 0. | 0. | 62 | 0 | | | Spe | Superpathway | 57 | 12 | 85 | 14 | NS | N | NS | N | N | NS | NS | NS | NS | N | 1 | 0. | 0.00 |

| | | | | | | | | | | | | | | | | | | | | | | | | | | | | | | |
|---|---|---|---|---|---|---|---|---|---|---|---|---|---|---|---|---|---|---|---|---|---|---|---|---|---|---|---|---|---|---|
| 5 | 740 | | LSE | | 5.0426 | 856 | 2 | 1178.9 | | | | ctral Match | of chorismate metabolism;Superpathway of aromatic amino acid biosynthesis;Other | 965.97 | 9403.4 | 226.15 | 0447.7 | | S | | S | S | | | | | S | 0.457 | 000206 | 0403 |
| 76 | C00750 | H | FALSE | N,N'-Bis(3-aminopropyl)-1,4-butanediamine | 201.2086 | 9.784 | 0.052 | 1614868 | 0 | | | Spectral Match | | 178752.1 | 170073.9 | 214532.7 | 158959.2 | 0.38944 | 0.0000 | 0.000618 | 0.675 | 0.5311 | 0.5048 | 4.03E-07 | 0.000571 | 0.898438 | 0.0002 | 24.792 | 7.76E-08 | 0.000744 |
| 77 | C00783 | H | FALSE | Tropinone | 139.0993 | 1.367 | 0.2 | 1399848 | 0 | | 70.3 | Spectral Match | Other | 55481.83 | 221123.7 | 115268.8 | 247587.4 | NS | NS | NS | NS | NS | NS | NS | NS | NS | NS | 19.218 | 1.19E-06 | 0.000667 |
| 78 | C00785 | BH | FALSE | Urocanic acid | 138.0426 | 1.06 | 0.2 | 7874073 | 0 | 56.8 | | Spectral Match | | 1952398 | 1249758 | 926935 | 1330464 | 3.8379 | 0.0087 | 0.000772 | 0.605 | 0.0143 | NS | NS | NS | NS | NS | 4.907 | 0.012157 | 0 |
| 79 | C00791 | FB | FALSE | Creatinine | 113.0589 | 1.248 | 0.2 | 2394354 | 0 | | | Spectral Match | | 195090.1 | 599870.3 | 785544.3 | 553451.8 | NS | NS | NS | NS | NS | 0.43442 | 0.038402 | 0 | 0.601563 | 0.3447 | 37.86 | 3.99E-10 | 0.000333 |
| 80 | C00792 | H | FALSE | D-Arginine | 174.1111 | 0.83 | 0.119 | 7658518 | 0 | | | Spectral Match | Arginine, ornithine and proline interconversion;Superpathway of L-arginine and L-ornithine degradation;Superpathway of L-arginine, putrescine, and 4-aminobutanoate degradation;Other | 273122.4 | 193938.2 | 252442.5 | 179312.1 | NS | NS | NS | NS | NS | NS | NS | NS | NS | NS | 17.562 | 2.87E-06 | 0.000125 |
| 8 | C00 | HF | FA | D-Sorbitol | 18 | 0. | 0. | 17 | 0 | | | Spe | | 79 | 37 | 59 | 31 | NS | N | NS | N | N | NS | NS | NS | NS | N | 5 | 1. | 0.00 |

| | | | | | | | | | | | | | | | | | | | | | | | | | | | | | | |
|---|---|---|---|---|---|---|---|---|---|---|---|---|---|---|---|---|---|---|---|---|---|---|---|---|---|---|---|---|---|---|
| 1 | 794 | B | LSE | | 3.0866 | 84 | 2 | 842319 | | | | ctral Match | | 6520.9 | 15107 | 60284 | 53813 | | S | | S | S | | | | | S | 6.516 | 23E-12 | 651 |
| 82 | C00828 | HB | FALSE | Menaquinone | 444.3044 | 13.153 | 0.2 | 414907.8 | 0 | | | Spectral Match | | 17677.39 | 63672.87 | 15862.77 | 75625.4 | NS | NS | NS | NS | NS | 3.082 | 1.74E-07 | 0.004629 | 0.996094 | 0.0000 | 95.293 | 2.45E-16 | 0.000712 |
| 83 | C00829 | H | FALSE | 1_3_8-Naphthalenertriol | 176.0468 | 10.381 | 0.2 | 3.19E+08 | 0 | | 97 | Spectral Match | | 184021.8 | 230385.2 | 230546.8 | 230344.8 | 0.50404 | 0.0000 | 0.00025 | 0.61 | 0.0016 | NS | NS | NS | NS | NS | 10.322 | 0.000226 | -7.35E-06 |
| 84 | C00836 | F | FALSE | Sphinganine;2-Amion-1_3-octadecanediol | 301.2968 | 11.719 | 0.2 | 7072920 | 0 | | | Spectral Match | Other | 993508.4 | 597800.1 | 872896.5 | 529026 | 0.24707 | 0.0000 | 0.000667 | 0.78 | 0.0103 | 0.42003 | 3.60E-06 | 0.000286 | 0.777344 | 0.0041 | 10.248 | 0.000237 | 0.000408 |
| 85 | C00860 | H | FALSE | L-Histidinol | 141.0902 | 1.066 | 0.2 | 246894.5 | 0 | | 79.8 | Spectral Match | Other | 10461.54 | 33825.49 | 4314.024 | 41203.36 | NS | NS | NS | NS | NS | 5.8893 | 5.78E-12 | 0.002367 | 1 | 0.0000 | 64.074 | 1.74E-13 | 0.001464 |
| 86 | C00869 | H | FALSE | 2-Oxooctadecanoicacid | 298.2498 | 12.464 | 0.2 | 67594.75 | 0 | | 74.3 | Spectral Match | | 1089.964 | 886.6158 | 1040.514 | 848.1412 | 0.32984 | 0.0000 | 0.001289 | 0.92 | 0.0002 | 0.54385 | 4.54E-08 | 0.00075 | 0.984375 | 0.0000 | 10.5 | 0.0002 | 0.000258 |
| 87 | C00902 | H | FALSE | (4E)-2-Oxohexenoicacid | 128.0472 | 0.994 | 0.2 | 5394458 | 0 | | | Spectral Match | Other | 10304.53 | 1022238 | 1017379 | 1023453 | 41.06 | 0.0000 | 0 | 0.95 | 0.0000 | NS | NS | NS | NS | NS | 29.295 | 1.08E-08 | 0.000611 |
| 88 | C00906 | H | FALSE | Dihydrothymine | 128.0583 | 0.893 | 0.081 | 2779912 | 0 | | 86.3 | Spectral Match | | 76239.14 | 579463.8 | 740500.7 | 539204.6 | 2.8507 | 0.0480 | 0 | 1 | 0.0001 | 0.46498 | 0.015567 | 0.000286 | 0.597656 | 0.3703 | 69.156 | 5.15E-14 | 0.006568 |
| 89 | C00922 | H | FALSE | 2_3-Dimethylmaleate | 144.0418 | 1.048 | 0.2 | 2411094 | 0 | | 81.8 | Spectral Match | Other | 138732.9 | 529288.2 | 312313.6 | 583531.9 | NS | NS | NS | NS | NS | NS | NS | NS | NS | NS | 8.7252 | 0.000678 | 0 |
| 9 | C00 | FB | FA | Porphobilin | 22 | 1. | 0. | 27 | 0 | | | Spe | Adenosylcobal | 46 | 49 | 24 | 55 | 0. | 0. | 0 | 0. | 0. | NS | NS | NS | NS | N | 1 | 3. | 0.00 |

| | | | | | | | | | | | | | | | | | | | | | | | | | | | | | | |
|---|---|---|---|---|---|---|---|---|---|---|---|---|---|---|---|---|---|---|---|---|---|---|---|---|---|---|---|---|---|---|
| 0 | 931 | | LSE | ogen | 6.0946 | 123 | 11 | 88048 | | | | ctral Match | amin biosynthesis I (anaerobic);Other | 5206.9 | 4155.6 | 7776.3 | 5750.5 | 38589 | 0068 | | 975 | 9957 | | | | | S | 7.383 | 16E-06 | 0251 |
| 91 | C00954 | H | FALSE | 3-Indoleacetic Acid | 175.0629 | 1.145 | 0.2 | 531160 | 0 | | | Spectral Match | Superpathway of indole-3-acetate conjugate biosynthesis | 7759.799 | 74485 | 16682.36 | 88935.66 | 4.6623 | 0.02331 | 0 | 0.965 | 0.0000 | 4.678 | 0.001401 | 0.000749 | 0.992188 | 0.00000 | 9.7696 | 0.000328 | 0.00062 |
| 92 | C00990 | B | FALSE | 5-Aminopentanamide | 116.0946 | 0.855 | 0.2 | 235231.4 | 0 | | | Spectral Match | Superpathway of L-lysine degradation | 13663.2 | 42719.82 | 43264.69 | 42583.6 | NS | NS | NS | NS | NS | NS | NS | NS | NS | NS | 15.322 | 1.01E-05 | 0.000125 |
| 93 | C00992 | B | FALSE | Aquacob(III)alamin | 546.2216 | 20.608 | 0.2 | 1798800 | 0 | | | Spectral Match | | 87403.63 | 222528.2 | 797509.58 | 258282.9 | NS | NS | NS | NS | NS | 2.2914 | 0.00043 | 0.000541 | 1 | 0.0000 | 8.5799 | 0.000751 | 0.000674 |
| 94 | C01004 | FB | FALSE | 3-Carboxy-1-methylpyridinium;N-Methylnicotinicacid | 137.0473 | 1.008 | 0.2 | 770904.7 | 0 | | 66.1 | Spectral Match | Superpathway of chorismate metabolism;Superpathway of aromatic amino acid biosynthesis;Superpathway of aromatic compound degradation via 2-oxopent-4-enoate;Anaerobic aromatic compound degradation (Thauera aromatica);Superpathway of aromatic compound degradation via 3-oxoadipate;Other | 96500.1 | 186427.7 | 233379.8 | 174689.7 | NS | NS | NS | NS | NS | 0.47516 | 0.016327 | 0 | 0.640625 | 0.3252 | 54.809 | 1.96E-12 | 0.001224 |

| | | | | | | | | | | | | | | | | | | | | | | | | | | | | | | |
|---|---|---|---|---|---|---|---|---|---|---|---|---|---|---|---|---|---|---|---|---|---|---|---|---|---|---|---|---|---|---|
| 95 | C01012 | FB | FALSE | (R)-Pantolactone | 130.0628 | 0.422 | 0.2 | 601538.9 | 0 | | | Spectral Match | Superpathway of L-threonine metabolism;Superpathway of branched chain amino acid biosynthesis;Superpathway of L-isoleucine biosynthesis I | 39699.33 | 61357.9 | 45435.8 | 65338.42 | 0.59103 | 0.0038 | 0 | 0.51 | 0.0311 | NS | NS | NS | NS | NS | 12.584 | 5.22E-05 | 0 |
| 96 | C01019 | HB | FALSE | L-Fucose | 165.0759 | 20.576 | 0.2 | 563223 | 0 | | | Spectral Match | | 13288.24 | 42365.73 | 35217.72 | 44152.73 | NS | NS | NS | NS | NS | NS | NS | NS | NS | NS | 3.8719 | 0.028623 | 0 |
| 97 | C01035 | FB | FALSE | 4-Guanidinobutanoate | 145.0846 | 1.01 | 0.2 | 965664.1 | 0 | | | Spectral Match | | 35812.42 | 217969.6 | 165745.2 | 231025.7 | 2.3112 | 0.0012 | 0 | 0.82 | 0.0000 | NS | NS | NS | NS | NS | 7.4174 | 0.001743 | 0 |
| 98 | C01042 | HFB | FALSE | N-Acetyl-L-aspartate | 176.0553 | 0.871 | 0.2 | 214252.6 | 0 | | | Spectral Match | Other | 8361.933 | 34705.55 | 9918.568 | 40902.3 | NS | NS | NS | NS | NS | 2.7438 | 1.80E-05 | 0.001755 | 1 | 0.0000 | 19.514 | 1.02E-06 | 0.000728 |
| 99 | C01047 | FB | FALSE | N5-Ethyl-L-glutamine | 174.1 | 0.976 | 0.2 | 868176.8 | 0 | | 79.9 | Spectral Match | Other | 138366.8 | 168159.3 | 88327.18 | 188117.3 | 0.44398 | 0.0005 | 0 | 1 | 0.5938 | NS | NS | NS | NS | NS | 22.594 | 2.18E-07 | 0.001766 |
| 100 | C01053 | FB | FALSE | (R)-4-Dehydropantoate | 146.0575 | 0.985 | 0.2 | 7879067 | 0 | | 77.4 | Spectral Match | Superpathway of L-threonine metabolism;Superpathway of branched chain amino acid biosynthesis;Superpathway of L-isoleucine biosynthesis I;Other | 323564.7 | 1354810 | 1567856 | 1301548 | NS | NS | NS | NS | NS | 0.52645 | 0.020925 | 0 | 0.558594 | 0.5595 | 34.839 | 1.20E-09 | 0.000444 |
| 10 | C01059 | B | FALS | 2_5-Dihydroxyp | 111.0 | 1.1 | 0.2 | 1540 | 0 | | 90.2 | Spectra | Other | 2252 | 3210 | 1296 | 3688 | 4.68 | 0.0 | 0.00 | 1 | 0.00 | NS | NS | NS | NS | NS | 9.3 | 0.00 | 0.00022 |

| | | | | | | | | | | | | | | | | | | | | | | | | | | | | | | |
|---|---|---|---|---|---|---|---|---|---|---|---|---|---|---|---|---|---|---|---|---|---|---|---|---|---|---|---|---|---|---|
| 1 | | | E | yridine | 319 | 45 | | 964 | | | | l Match | | 4.69 | 19.8 | 52.7 | 61.6 | 92 | 000 | 0236 | | 00 | | | | | | 855 | 0427 | 9 |
| 102 | C01068 | HB | FALSE | D-Ribitol 5-phosphate | 231.0279 | 20.532 | 0.2 | 2903358 | 0 | | | Spectral Match | | 282910.7 | 377857.7 | 476442.6 | 353211.5 | 0.53607 | 0.00002 | 4.00E-04 | 1 | 0.0312 | 0.48797 | 1.38E-09 | 0.0025 | 0.835938 | 0.0025 | 10.15 | 0.000254 | 0.000739 |
| 103 | C01079 | HFB | FALSE | Protoporphyrinogen IX | 568.3047 | 7.677 | 0.2 | 4076464 | 0 | | | Spectral Match | Other | 44927.58 | 219757.1 | 122257.6 | 244132 | 1.8301 | 0.0030 | 0.000308 | 0.72 | 0.0000 | NS | NS | NS | NS | NS | 21.743 | 3.30E-07 | 0.00025 |
| 104 | C01087 | FB | FALSE | (R)-2-Hydroxyglutarate | 148.0371 | 0.657 | 0.2 | 181253.5 | 0 | | 63.4 | Spectral Match | Other | 30189.07 | 29062.43 | 28447.15 | 29216.25 | 0.37632 | 0.0002 | 0 | 0.575 | 0.7919 | NS | NS | NS | NS | NS | 12.234 | 6.51E-05 | 0.000458 |
| 105 | C01092 | H | FALSE | 8-Amino-7-oxononanoate | 187.1202 | 7.1 | 0.2 | 368931 | 0 | | | Spectral Match | Biotin biosynthesis II;Other | 4498.296 | 49912.72 | 16011.39 | 58388.05 | 4.2427 | 0.0000 | 0.001115 | 0.985 | 0.0000 | 2.4557 | 0.000314 | 0.0000671 | 1 | 0.0000 | 36.655 | 6.15E-10 | 0.0006163 |
| 106 | C01102 | FB | FALSE | O-Phospho-L-homoserine | 98.50518 | 0.293 | 0.2 | 935475.5 | 0 | | | Spectral Match | | 1120.7 | 359362.4 | 354284.1 | 360632 | 122.68 | 0.0000 | 0.00025 | 0.585 | 0.0000 | NS | NS | NS | NS | NS | 11.395 | 0.000111 | 0 |
| 107 | C01104 | H | FALSE | Nonyltrimethylammonium | 185.2138 | 10.873 | 0.2 | 9613937 | 0 | 73.2 | | Spectral Match | | 34170.23 | 65147.32 | 538884.66 | 67962.99 | NS | NS | NS | NS | NS | NS | NS | NS | NS | NS | 14.819 | 1.35E-05 | 0.000289 |
| 108 | C01107 | F | FALSE | Amiloride | 229.0471 | 1.859 | 0.2 | 71084387 | 0 | 89.2 | | Spectral Match | | 17420.57 | 99249.31 | 10347608 | 98192.62 | 2.235 | 0.0000 | 0.001533 | 0.845 | 0.0000 | 0.62501 | 9.26E-06 | 0.00103 | 0.601563 | 0.5347 | 29.156 | 1.15E-08 | 0.000244 |
| 109 | C01109 | H | FALSE | 4-Methylene-L-glutamine | 158.0688 | 1.319 | 0.2 | 133627.3 | 0 | | | Spectral Match | | 302229.65 | 262232.81 | 235451.72 | 269055.58 | 0.32374 | 0.0000 | 0.000536 | 1 | 0.6424 | NS | NS | NS | NS | NS | 11.165 | 0.000129 | 0.0000682 |
| 11 | C01118 | FB | FALS | O-Succinyl-L- | 219.0 | 0.9 | 0.2 | 3245 | 0 | | | Spectra | Other | 1446 | 5589 | 3284 | 6165 | NS | NS | NS | NS | NS | NS | NS | NS | NS | NS | 13. | 2.78 | 0.00058 |

| | | | | | | | | | | | | | | | | | | | | | | | | | | | | | | |
|---|---|---|---|---|---|---|---|---|---|---|---|---|---|---|---|---|---|---|---|---|---|---|---|---|---|---|---|---|---|---|
| 0 | | | E | homoserine | 735 | 62 | | 34.4 | | | | l Match | | 7.83 | 1.8 | 5.52 | 3.37 | | | | | | | | | | | 609 | E-05 | 3 |
| 111 | C01120 | HF | FALSE | [Similar to: R-Palmitoyl-(1-methyl) Ethanolamide; ΔMass: -68.9721 Da] | 382.2702 | 13.291 | 0.2 | 3581395 | 0 | | 85 | Spectral Match | | 511442.97 | 358032 | 342029.7 | 362032.5 | 2.7611 | 0.00000 | 0.001863 | 0.52 | 0.0000 | 4.5455 | 1.16E-11 | 0.001719 | 0.632813 | 0.8622 | 63.649 | 1.93E-13 | 0.001173 |
| 112 | C01137 | HFB | FALSE | S-Adenosylmethioninamine | 356.1634 | 15.729 | 0.2 | 1389295 | 0 | | | Spectral Match | | 212221.8 | 132778.5 | 130781.2 | 133277.8 | 0.24999 | 0.0000 | 0.002351 | 1 | 0.0000 | NS | NS | NS | NS | NS | 16.025 | 6.73E-06 | 0.002196 |
| 113 | C01142 | FB | FALSE | (3S)-3_6-Diaminohexanoate | 146.1051 | 0.791 | 0.2 | 2662716 | 0 | | 72.4 | Spectral Match | Superpathway of L-lysine degradation;Biotin biosynthesis II;Other | 186927.6 | 593280.7 | 581707.7 | 596173.9 | NS | NS | NS | NS | NS | 0.63115 | 0.038749 | 0 | 0.558594 | 0.9293 | 38.149 | 3.60E-10 | 0.004695 |
| 114 | C01180 | FB | FALSE | 2-keto-4-methylthiobutyricacid(KMBA) | 148.019 | 0.934 | 0.2 | 1251520 | 0 | | | Spectral Match | Other | 180048.73 | 289069.7 | 287057.7 | 289572.7 | 6.2395 | 0.0001 | 0 | 1 | 0.0000 | NS | NS | NS | NS | NS | 11.798 | 8.59E-05 | 0.000105 |
| 115 | C01181 | HFB | FALSE | 4-Trimethylammoniobutanoate | 147.1254 | 0.866 | 0.2 | 1006414 | 0 | | | Spectral Match | | 53030.85 | 220856.4 | 305208.5 | 199768.4 | NS | NS | NS | NS | NS | 0.43175 | 0.021918 | 0.000154 | 0.609375 | 0.3175 | 23.488 | 1.42E-07 | 0.000729 |
| 116 | C01183 | F | FALSE | N,N-Dimethyldecylamine N-oxide | 201.2086 | 9.72 | 0.052 | 5786562 | 0 | 84.2 | | Spectral Match | | 111469.8 | 640201.41 | 466505.41 | 683362.91 | 0.22687 | 0.0000 | 0 | 1 | 0.0781 | NS | NS | NS | NS | NS | 11.964 | 7.72E-05 | 0.000118 |
| 117 | C01212 | B | FALSE | UDP-N-acetylmuramoyl-L-alanine | 376.0778 | 7.961 | 0.2 | 237379.5 | 0 | | | Spectral Match | | 115.9413 | 553839.7 | 230047.31 | 634668.14 | 171.15 | 0.0000 | 0.000711 | 1 | 0.0000 | NS | NS | NS | NS | NS | 25.468 | 5.71E-08 | 0.001118 |
| 118 | C01214 | FB | FALSE | 1-Amino-1-deoxy-scyllo- | 179.078 | 0.97 | 0.2 | 238533 | 0 | | 86.3 | Spectral | Other | 180722. | 420571. | 260584. | 460567. | NS | NS | NS | NS | NS | NS | NS | NS | NS | NS | 7.90 | 0.0012 | 0 |

| | | | | | | | | | | | | | | | | | | | | | | | | | | | | | | |
|---|---|---|---|---|---|---|---|---|---|---|---|---|---|---|---|---|---|---|---|---|---|---|---|---|---|---|---|---|---|---|
| | | | | inositol | 7 | 7 | | 7 | | | | Match | | 9 | 2 | 8 | 8 | | | | | | | | | | | 69 | 18 | |
| 119 | C01234 | H | FALSE | 1-Aminocyclopropane-1-carboxylate | 101.048 | 0.937 | 0.111 | 4424176 | 0 | | | Spectral Match | Superpathway of L-threonine metabolism;Superpathway of jasmonoyl-amino acid conjugates biosynthesis;Superpathway of branched chain amino acid biosynthesis;Superpathway of L-isoleucine biosynthesis I;Other | 169998.3 | 929378.3 | 633246 | 1003411 | 1.9966 | 0.0121 | 0 | 0.595 | 0.0000 | NS | NS | NS | NS | NS | 15.204 | 1.08E-05 | 0.000125 |
| 120 | C01239 | H | FALSE | N-Acetyl-beta-D-glucosaminylamine | 220.1052 | 0.987 | 0.2 | 1537664 | 0 | | | Spectral Match | | 133016.9 | 310232.1 | 352563 | 299649.3 | NS | NS | NS | NS | NS | 0.54006 | 0.000143 | 0.000308 | 0.578125 | 0.3436 | 18.514 | 1.72E-06 | 0.000952 |
| 121 | C01250 | FB | FALSE | N-Acetyl-L-glutamate5-semialdehyde | 173.0681 | 0.981 | 0.2 | 1750182 | 0 | | | Spectral Match | Other | 503647 | 248754.7 | 186455.4 | 264329.6 | 0.18486 | 0.0000 | 0 | 0.975 | 0.0046 | NS | NS | NS | NS | NS | 49.253 | 9.70E-12 | 0.001126 |
| 123 | C01255 | H | FALSE | N-(6-Aminohexanoyl)-6-aminohexanoate | 244.1779 | 6.783 | 0.2 | 958108.3 | 0 | | 64.5 | Spectral Match | | 16887.79 | 243406.7 | 222035.4 | 248749.5 | 5.1191 | 0.0003 | 0.000267 | 1 | 0.0000 | NS | NS | NS | NS | NS | 7.2224 | 0.002014 | 0 |
| 124 | C01284 | HFB | FALSE | 1D-myo-Inositol 1,3,4,5,6-pentakisphosphate | 290.9545 | 21.036 | 0.2 | 1883014 | 0 | | | Spectral Match | | 160641.8 | 283490.6 | 206511.7 | 302735.4 | NS | NS | NS | NS | NS | NS | NS | NS | NS | NS | 5.4347 | 0.007961 | 0.000353 |
| 125 | C01297 | H | FALSE | 6-Hydroxypseudooxynicotine | 194.1065 | 15.99 | 0.2 | 107136.7 | 0 | | 89.1 | Spectral Match | | 31618.29 | 17278.91 | 14315.15 | 18019.85 | 0.20812 | 0.0000 | 0.000619 | 0.985 | 0.0303 | NS | NS | NS | NS | NS | 27.346 | 2.48E-08 | 0.000583 |
| 1 | C01 | HF | FA | 2'- | 19 | 0. | 0. | 16 | 0 | | | Spe | | 63 | 21 | 44 | 25 | NS | N | NS | N | N | 4. | 0. | 0.00 | 0. | 0. | 1 | 0. | 0.00 |

| | | | | | | | | | | | | | | | | | | | | | | | | | | | | | | |
|---|---|---|---|---|---|---|---|---|---|---|---|---|---|---|---|---|---|---|---|---|---|---|---|---|---|---|---|---|---|---|
| 26 | 346 | B | LSE | Deoxyuridine 5'-diphosphate | 5.01 | 671 | 2 | 780901 | | | | ctral Match | | 9690.4 | 13182 | 6380.8 | 29882 | | S | | S | S | 6798 | 000129 | 1165 | 988281 | 0000 | 0.291 | 000231 | 0125 |
| 127 | C01489 | B | FALSE | D-Iditol | 182.0786 | 0.983 | 0.2 | 18399140 | 0 | | | Spectral Match | Other | 18761.44 | 336590 | 3429794 | 3349663 | 74.648 | 0.0000 | 0.000308 | 1 | 0.0000 | NS | NS | NS | NS | NS | 3.7701 | 0.031198 | 0 |
| 128 | C01516 | H | FALSE | Morphine | 285.1355 | 11.34 | 0.2 | 1135419 | 0 | | | Spectral Match | | 22194.88 | 49030.72 | 21047.43 | 56026.54 | NS | NS | NS | NS | NS | 1.7629 | 0.000645 | 0.001532 | 1 | 0.0000 | 7.448 | 0.001704 | 0.000639 |
| 129 | C01546 | H | FALSE | 2-Furoate | 112.0156 | 0.999 | 0.2 | 948700.6 | 0 | | 81.8 | Spectral Match | | 110554.5 | 191793 | 124701.8 | 208565.8 | 0.6085 | 0.0223 | 0 | 0.8 | 0.2298 | NS | NS | NS | NS | NS | 14.861 | 1.32E-05 | 0.002049 |
| 130 | C01561 | H | FALSE | [ST(2:0)](5Z_7E)-(3S)-9_10-seco-5_7_10(19)-cholestatriene-3_25-diol | 400.3322 | 16.438 | 0.2 | 189151.7 | 0 | | | Spectral Match | Other | 27104.94 | 14245.24 | 15261.2 | 13991.25 | 0.21225 | 0.0000 | 0.002282 | 0.69 | 0.0000 | 0.60969 | 0.000824 | 0 | 0.609375 | 0.5230 | 6.1465 | 0.004556 | -0.00013 |
| 131 | C01586 | H | FALSE | Hippuric Acid | 179.0577 | 4.544 | 0.2 | 455073.9 | 0 | | 68.9 | Spectral Match | Other | 12576.55 | 74441.99 | 158931 | 53319.72 | NS | NS | NS | NS | NS | 0.22562 | 5.80E-05 | 0.000241 | 0.753906 | 0.0117 | 8.4303 | 0.000835 | 0.000575 |
| 132 | C01595 | HF | FALSE | Linoleate | 280.2388 | 16.873 | 0.2 | 175135.2 | 0 | | | Spectral Match | Other | 37339.63 | 71349.48 | 49287.37 | 76865.01 | NS | NS | NS | NS | NS | NS | NS | NS | NS | NS | 4.1513 | 0.022638 | 0 |
| 133 | C01606 | H | FALSE | Phthalic Acid, Bis-Hexyl Ester | 334.214 | 7.924 | 0.2 | 532528.7 | 0 | | | Spectral Match | Other | 24072.96 | 61335.41 | 48504.21 | 64543.21 | NS | NS | NS | NS | NS | NS | NS | NS | NS | NS | 123.82 | 2.45E-18 | 0.003656 |
| 134 | C01674 | HB | FALSE | Chitobiose | 212.066 | 0.79 | 0.2 | 650072 | 0 | | | Spectral | | 169526. | 141560. | 183504. | 131074. | 0.3206 | 0.03 | 0 | 0.86 | 0.5091 | NS | NS | NS | NS | NS | 62.8 | 2.34E- | 0.000297 |

| | | | | | | | | | | | | | | | | | | | | | | | | | | | | | | |
|---|---|---|---|---|---|---|---|---|---|---|---|---|---|---|---|---|---|---|---|---|---|---|---|---|---|---|---|---|---|---|
| | | | | | 8 | 4 | | .8 | | | | Match | | 4 | 6 | 2 | 7 | 8 | 30 | | | | | | | | | 92 | 13 | |
| 135 | C01717 | B | FALSE | 4-Hydroxy-2-quinolinecarboxylic acid | 189.042 | 16.644 | 0.2 | 332363.4 | 0 | | 81.6 | Spectral Match | Other | 10776.97 | 29179.55 | 21984.15 | 30978.4 | NS | NS | NS | NS | NS | NS | NS | NS | NS | NS | 31.6446 | 4.15E-09 | 0.002514 |
| 136 | C01732 | B | FALSE | Mesaconate | 128.0119 | 0.677 | 0.2 | 208548.7 | 0 | | | Spectral Match | | 29343.95 | 43162.39 | 36399.77 | 44853.05 | 0.54999 | 0.03557 | 0 | 0.595 | 0.3846 | NS | NS | NS | NS | NS | 19.877 | 8.43E-07 | 5.00E-04 |
| 137 | C01746 | H | FALSE | Piperidine | 85.08947 | 1.213 | 0.102 | 23778273 | 0 | | | Spectral Match | | 2124489 | 3929681 | 1988604 | 4414950 | 0.63669 | 0.0290 | 0 | 0.945 | 0.2270 | NS | NS | NS | NS | NS | 5.8419 | 0.005775 | 0.00019 |
| 138 | C01770 | FB | FALSE | 1_4-Lactone | 86.03707 | 0.382 | 0.2 | 787858.8 | 0 | | | Spectral Match | Other | 92581.9 | 144655 | 154503.3 | 142192.9 | NS | NS | NS | NS | NS | NS | NS | NS | NS | NS | 12.841 | 4.45E-05 | -2.81E-05 |
| 139 | C01772 | FB | FALSE | 2-coumarate | 164.0468 | 1.073 | 0.2 | 25205782 | 0 | | | Spectral Match | Superpathway of chorismate metabolism;Superpathway of aromatic amino acid biosynthesis;Anaerobic aromatic compound degradation (Thauera aromatica);Other | 446118.9 | 5168957 | 2232434 | 5903088 | 3.9118 | 0.0010 | 0 | 0.86 | 0.0000 | NS | NS | NS | NS | NS | 7.745 | 0.00137 | 0 |
| 140 | C01794 | H | FALSE | Choloyl-CoA | 577.6908 | 10.096 | 0.2 | 548228.4 | 0 | | | Spectral Match | | 17014.5 | 24594.82 | 37586.85 | 21346.82 | 0.61382 | 0.0183 | 0 | 0.77 | 0.1536 | 0.37552 | 7.51E-06 | 0 | 0.839844 | 0.0014 | 8.5344 | 0.000776 | -0.00022 |
| 141 | C01817 | H | FALSE | Homocystine | 268.054 | 1.074 | 0.2 | 359781.3 | 0 | | | Spectral Mat | | 24490.09 | 65862.24 | 71109.81 | 64550.35 | NS | NS | NS | NS | NS | NS | NS | NS | NS | NS | 11.04 | 0.00014 | 0.00036 |

| | | | | | | | | | | | | | | | | | | | | | | | | | | | | | | |
|---|---|---|---|---|---|---|---|---|---|---|---|---|---|---|---|---|---|---|---|---|---|---|---|---|---|---|---|---|---|---|
| | | | | | | | | | | | | ch | | | | | | | | | | | | | | | | 8 | | |
| 142 | C01826 | FB | FALSE | [FAamino(5:0)]2S-amino-pentanoicacid | 117.079 | 0.779 | 0.112 | 73860013 | 0 | | | Spectral Match | Superpathway of L-lysine degradation;Superpathway of linamarin and lotaustralin biosynthesis;Arginine, ornithine and proline interconversion;Superpathway of indole-3-acetate conjugate biosynthesis;Superpathway of jasmonoyl-amino acid conjugates biosynthesis;Superpathway of branched chain amino acid biosynthesis;Other | 2856526 | 14022133 | 3172049 | 16734654 | NS | NS | NS | NS | NS | NS | NS | NS | NS | NS | 17.589 | 2.82E-06 | 4.00E-04 |
| 143 | C01877 | FB | FALSE | 4-Oxoproline | 129.0406 | 1.074 | 0.046 | 34294897 | 0 | 50.7 | | Spectral Match | | 1551621 | 8701698 | 56840003 | 9456121 | 2.0186 | 0.0464 | 0 | 0.73 | 0.0001 | NS | NS | NS | NS | NS | 4.9689 | 0.011564 | 4.00E-04 |
| 144 | C01905 | FB | FALSE | D-Asparagine | 132.0533 | 1.024 | 0.2 | 9593349.8 | 0 | | | Spectral Match | Other | 32124.78 | 170783.4 | 452241.4 | 202168.9 | NS | NS | NS | NS | NS | 2.6923 | 6.28E-07 | 0.001682 | 0.96875 | 0.0000 | 26.445 | 3.69E-08 | 0.000316 |
| 145 | C01909 | H | FALSE | Dethiobiotin | 214.1311 | 1.116 | 0.2 | 312533.7 | 0 | | 73.6 | Spectral Match | Biotin biosynthesis II;Other | 455549.65 | 634437.27 | 358332.28 | 703338.52 | NS | NS | NS | NS | NS | NS | NS | NS | NS | NS | 7.8333 | 0.001285 | -4.85E-05 |

| | | | | | | | | | | | | | | | | | | | | | | | | | | | | | | |
|---|---|---|---|---|---|---|---|---|---|---|---|---|---|---|---|---|---|---|---|---|---|---|---|---|---|---|---|---|---|---|
| 146 | C01921 | H | FALSE | Glycocholic acid | 465.3066 | 10.172 | 0.2 | 20517838 | 0 | | | Spectral Match | | 35583.88 | 4769261 | 3457076 | 5097307 | 49.451 | 0.0000 | 0.000222 | 0.96 | 0.0000 | NS | NS | NS | NS | NS | 22.918 | 1.87E-07 | 0.005274 |
| 147 | C01924 | H | FALSE | Homoarginine | 188.1267 | 0.823 | 0.2 | 377284.5 | 0 | | | Spectral Match | | 6422.156 | 73334.23 | 111682.1 | 63747.26 | 4.5967 | 0.0072 | 0 | 1 | 0.0000 | 0.37209 | 0.006189 | 0.000154 | 0.664063 | 0.1554 | 87.343 | 1.09E-15 | 0.003224 |
| 148 | C02046 | F | FALSE | Atropine | 289.1666 | 6.952 | 0.2 | 64577908 | 0 | 86.6 | | Spectral Match | Other | 2380802 | 5054064 | 3397726 | 5468149 | NS | NS | NS | NS | NS | NS | NS | NS | NS | NS | 10.596 | 0.000188 | 0.000267 |
| 149 | C02050 | H | FALSE | Linoleoyl-CoA | 344.1214 | 10.943 | 0.2 | 5037112 | 0 | | | Spectral Match | | 108451.3 | 179438.3 | 158158.6 | 184758.2 | 0.6578 | 0.0015 | 0.000143 | 0.71 | 0.0000 | NS | NS | NS | NS | NS | 15.562 | 8.77E-06 | 0 |
| 150 | C02067 | FB | FALSE | Pseudouridine | 244.2022 | 13.191 | 0.2 | 363512.1 | 0 | | | Spectral Match | | 6483.198 | 5601.111 | 3628.317 | 6094.31 | 0.32833 | 0.0000 | 0.00025 | 0.975 | 0.3591 | NS | NS | NS | NS | NS | 32.25 | 3.26E-09 | 0.000765 |
| 151 | C02115 | B | FALSE | 2-Methylserine | 119.0582 | 1.059 | 0.134 | 3577846 | 0 | | 77.1 | Spectral Match | Superpathway of L-threonine metabolism;Superpathway of branched chain amino acid biosynthesis;Superpathway of L-isoleucine biosynthesis I;Adenosylcobalamin biosynthesis I (anaerobic);Other | 67672.79 | 871782.5 | 528587 | 957581.3 | 4.5023 | 0.0000 | 0 | 0.975 | 0.0000 | NS | NS | NS | NS | NS | 60.9 | 3.87E-13 | 0.000667 |
| 152 | C02167 | HB | FALSE | Maleylpyruvate | 182.995 | 0.672 | 0.2 | 228342.3 | 0 | | | Spectral Mat | | 34539.81 | 35767.03 | 66934.94 | 27975.05 | 0.43289 | 0.000 | 0.000125 | 0.6 | 0.6862 | 0.26864 | 6.41E-08 | 0.002003 | 0.890625 | 0.000 | 129.9 | 1.03E-18 | 0.000886 |

| | | | | | | | | | | | | | | | | | | | | | | | | | | | | | |
|---|---|---|---|---|---|---|---|---|---|---|---|---|---|---|---|---|---|---|---|---|---|---|---|---|---|---|---|---|---|
| | | | | | | | | | | | | ch | | | | | | | 7 | | | | | | | | 8 | 3 | | |
| 153 | C02266 | H | FALSE | D-Xylonolactone | 149.0448 | 0.832 | 0.2 | 1108347 | 0 | | | Spectral Match | | 14119.4 | 245521 | 228346.3 | 249814.7 | 6.1729 | 0.0001 | 0 | 0.59 | 0.0000 | NS | NS | NS | NS | NS | 5.8728 | 0.005637 | 0.000375 |
| 154 | C02291 | HF | FALSE | L-Cystathionine | 222.0598 | 0.673 | 0.2 | 242252.9 | 0 | | | Spectral Match | | 23762.38 | 37048.82 | 76571.05 | 27168.26 | NS | NS | NS | NS | NS | 0.22799 | 1.89E-05 | -0.000031 | 0.785156 | 0.0059 | 12.066 | 7.24E-05 | 0.000526 |
| 155 | C02350 | H | FALSE | (S)-Allantoin | 78.01435 | 1.051 | 0.2 | 32393656 | 0 | | | Spectral Match | | 831140.9 | 5890313 | 6053223 | 5849585 | 4.8273 | 0.0000 | 0.002746 | 1 | 0.0000 | 0.62761 | 0.002448 | 0.0000286 | 0.628906 | 0.5027 | 15.738 | 7.93E-06 | 0.000133 |
| 156 | C02372 | H | FALSE | 4-Aminophenol | 109.0528 | 1.996 | 0.2 | 448893.7 | 0 | | | Spectral Match | | 5882.464 | 68834.62 | 16150.12 | 82005.75 | 3.8613 | 0.0093 | 0 | 0.605 | 0.0001 | 2.982 | 0.001494 | 0.0000154 | 0.933594 | 0.0001 | 71.755 | 2.83E-14 | 0.001427 |
| 157 | C02376 | B | FALSE | 5-Methylcytosine | 125.0588 | 1.223 | 0.2 | 1615657 | 0 | | | Spectral Match | | 3998.255 | 169819.3 | 558636 | 726150.06 | 19.369 | 0.0029 | 0.001471 | 0.955 | 0.0000 | 0.079121 | 0.000901 | -0.000011 | 0.71875 | 0.0236 | 9.2783 | 0.00046 | 0 |
| 158 | C02455 | H | FALSE | 1-Phenylethylamine | 121.089 | 1.335 | 0.2 | 464090.4 | 0 | | | Spectral Match | Other | 915330.75 | 608860.34 | 438868.9 | 651082.2 | 0.25563 | 0.0000 | 0.001867 | 0.845 | 0.0021 | NS | NS | NS | NS | NS | 5.2605 | 0.009146 | 0.001125 |
| 159 | C02502 | FB | FALSE | 2-Hydroxypyridine | 95.03736 | 1.083 | 0.2 | 4208183 | 0 | | | Spectral Match | | 25384.41 | 373025.6 | 1452284 | 103211.1 | NS | NS | NS | NS | NS | 0.040191 | 5.29E-06 | 0.0001736 | 0.898438 | 0.0002 | 10.026 | 0.000276 | 0.000889 |
| 160 | C02505 | B | FALSE | 2-Aminoacetophenone; O-Acetylaniline | 135.0682 | 6.551 | 0.2 | 298140.8 | 0 | | 72.5 | Spectral Match | | 7855.221 | 22664.89 | 27081.88 | 215660.64 | NS | NS | NS | NS | NS | 0.51728 | 0.024947 | 0.0000154 | 0.640625 | 0.5090 | 36.023 | 7.75E-10 | 0.001511 |
| 162 | C02538 | H | FALSE | Estrone 3-sulfate | 350.169 | 7.64 | 0.2 | 535981 | 0 | | | Spectral | | 8849.94 | 90595.8 | 29455.0 | 105881 | NS | NS | NS | NS | NS | NS | NS | NS | NS | NS | 28.3 | 1.63E- | 7.29E-05 |

| | | | | | | | | | | | | | | | | | | | | | | | | | | | | | | |
|---|---|---|---|---|---|---|---|---|---|---|---|---|---|---|---|---|---|---|---|---|---|---|---|---|---|---|---|---|---|---|
| | | | | | 2 | 5 | | .6 | | | | Match | | 9 | 1 | 5 | | | | | | | | | | | | 2 | 08 | |
| 163 | C02571 | H | FALSE | Acetyl-L-carnitine | 203.115 | 1.061 | 0.049 | 8884617 | 0 | | 72.4 | Spectral Match | | 481993.9 | 1839525 | 2972194 | 1556357 | NS | NS | NS | NS | NS | 0.33045 | 0.035624 | 0 | 0.578125 | 0.3830 | 9.0999 | 0.000521 | 0.001364 |
| 164 | C02636 | FB | FALSE | 3-Dehydrocarnitine | 159.0891 | 0.981 | 0.2 | 1121017 | 0 | | 90.6 | Spectral Match | Superpathway of L-lysine degradation | 116257.5 | 243177.6 | 147327.1 | 267140.3 | NS | NS | NS | NS | NS | NS | NS | NS | NS | NS | 5.861 | 0.005689 | -0.00011 |
| 165 | C02648 | H | FALSE | 4-Hydroxy-2-butynal | 84.02154 | 1.512 | 0.2 | 1344450 | 0 | | | Spectral Match | | 165909.1 | 234006.6 | 152958.6 | 254268.6 | 0.51582 | 0.01330 | 0 | 1 | 0.4186 | NS | NS | NS | NS | NS | 10.57 | 0.000191 | 0.000133 |
| 166 | C02659 | B | FALSE | Acetonecyanohydrin | 85.05311 | 1.216 | 0.2 | 1663601 | 0 | | | Spectral Match | Superpathway of linamarin and lotaustralin biosynthesis | 101433.9 | 326116.5 | 250420.9 | 345040.4 | NS | NS | NS | NS | NS | NS | NS | NS | NS | NS | 17.938 | 2.34E-06 | 0.001253 |
| 167 | C02735 | B | FALSE | 2-Hydroxyphenethylamine | 137.0838 | 1.428 | 0.2 | 285376.2 | 0 | | | Spectral Match | | 30727.59 | 58706.43 | 84624.99 | 52226.78 | NS | NS | NS | NS | NS | 0.39368 | 7.27E-05 | -0.000031 | 0.75 | 0.0238 | 25.997 | 4.50E-08 | 0.000964 |
| 168 | C02774 | FB | FALSE | 2-hydroxycaproicacid | 132.0785 | 5.086 | 0.2 | 2947536 | 0 | | 67.5 | Spectral Match | | 449639.8 | 695901.6 | 636465.6 | 710760.6 | 0.62194 | 0.03449 | 0 | 0.725 | 0.2134 | NS | NS | NS | NS | NS | 16.451 | 5.30E-06 | 0.000222 |
| 169 | C02794 | HB | FALSE | 3-Hydroxykynurenine | 224.0788 | 1.509 | 0.2 | 6922383 | 0 | | | Spectral Match | Superpathway of NAD biosynthesis in eukaryotes;Other | 2214779 | 1167347 | 206129.5 | 1407652 | 0.17871 | 0.00001 | 0 | 0.605 | 0.1230 | 4.0191 | 4.83E-05 | 0.002648 | 0.972656 | 0.00000 | 34.009 | 1.65E-09 | 0.000235 |
| 170 | C02835 | H | FALSE | Imidazoleaceticacid | 126.0428 | 1.421 | 0.2 | 109440.8 | 0 | | | Spectral Match | Other | 11316.26 | 21618.33 | 20779.57 | 21828.02 | 0.39658 | 0.00002 | 0 | 0.975 | 0.1353 | 0.57239 | 0.021234 | 0.000667 | 0.570313 | 0.7431 | 60.17 | 4.67E-13 | 0.000235 |
| 171 | C02862 | H | FALSE | Butyryl-L-carnitine | 231.146 | 2.23 | 0.054 | 419184 | 0 | | | Spectral | | 24749.4 | 89361.9 | 144570. | 75559.8 | NS | NS | NS | NS | NS | 0.3291 | 0.0350 | 0 | 0.5742 | 0.39 | 9.29 | 0.0004 | 0 |

| | | | | | | | | | | | | | | | | | | | | | | | | | | | | | | |
|---|---|---|---|---|---|---|---|---|---|---|---|---|---|---|---|---|---|---|---|---|---|---|---|---|---|---|---|---|---|---|
| | | | | | 3 | 8 | | .2 | | | | Match | | 1 | 4 | 3 | 5 | | | | | | 7 | 33 | | 19 | 47 | 16 | 56 | |
| 172 | C02923 | HBF | FALSE | 2,3-Dihydroxytoluene | 124.0304 | 0.681 | 0.2 | 1971284 | 0 | | | Spectral Match | | 220521.5 | 311121.7 | 201500.4 | 338527 | 0.51573 | 0.0116 | 0 | 1 | 0.2706 | NS | NS | NS | NS | NS | 77.23 | 8.50E-15 | 0.000901 |
| 173 | C02946 | H | FALSE | 4-Acetamidobutanoic acid | 145.0736 | 0.405 | 0.2 | 4384968 | 0 | 62.9 | | Spectral Match | Superpathway of L-lysine degradation;Biotin biosynthesis II | 228880.2 | 380340.2 | 317365.5 | 396083.9 | NS | NS | NS | NS | NS | NS | NS | NS | NS | NS | 226.25 | 3.24E-23 | 0.003085 |
| 174 | C02982 | FB | FALSE | L-2-Hydroxyphytanate | 328.2963 | 16.598 | 0.2 | 620070.9 | 0 | | 70.1 | Spectral Match | | 115066.23 | 298332.34 | 399331.8 | 273307.47 | NS | NS | NS | NS | NS | 0.46254 | 4.36E-08 | 0 | 0.835938 | 0.0001 | 7.9761 | 0.001158 | 0.000133 |
| 175 | C02989 | H | FALSE | L-Methionine S-oxide | 165.0454 | 0.93 | 0.2 | 1596526 | 0 | | | Spectral Match | Other | 341719.45 | 371027.8 | 386310.2 | 367207.2 | 4.133 | 0.0006 | 0 | 0.675 | 0.0000 | NS | NS | NS | NS | NS | 195.43 | 5.31E-22 | 0.003999 |
| 176 | C02990 | H | FALSE | L-Palmitoylcarnitine | 399.3524 | 13.376 | 0.2 | 8302970 | 0 | | 85 | Spectral Match | | 359349 | 555817.1 | 483847.7 | 573809.5 | 0.61054 | 0.0001 | 8.57E-05 | 0.64 | 0.0000 | NS | NS | NS | NS | NS | 100.71 | 9.43E-17 | 0.005493 |
| 177 | C03017 | H | FALSE | O-Propanoylcarnitine | 217.1309 | 6.672 | 0.2 | 136629.7 | 0 | | | Spectral Match | | 29782.4 | 31431.43 | 6346.598 | 37702.63 | 0.38708 | 0.0021 | 0 | 1 | 0.6963 | 3.754 | 4.83E-05 | 0.003028 | 1 | 0.0000 | 20.77 | 5.35E-07 | 0.001336 |
| 178 | C03037 | B | FALSE | (+)-Bornane-2_5-dione | 166.0991 | 7.912 | 0.2 | 197467.5 | 0 | | | Spectral Match | | 616.3267 | 10506.37 | 14078.53 | 9613.33 | 7.0547 | 0.0000 | 0.001 | 0.545 | 0.0000 | 0.46744 | 7.64E-05 | 0 | 0.753906 | 0.0171 | 68.316 | 6.27E-14 | 0.003708 |
| 179 | C03045 | H | FALSE | 2-Oxindole | 133.0523 | 1.115 | 0.2 | 955367 | 0 | | 93.9 | Spectral Match | Other | 118448 | 147718.3 | 74844.62 | 165936.8 | 0.45023 | 0.0007 | 0 | 0.625 | 0.5499 | NS | NS | NS | NS | NS | 24.049 | 1.09E-07 | 0.000353 |
| 180 | C03059 | H | FALSE | 2-Hydroxymethylserine | 135.053 | 0.99 | 0.2 | 404008 | 0 | | 93.9 | Spectral | | 120081 | 862628 | 100710 | 826509 | 0.2792 | 0.00 | 0 | 1 | 0.3002 | 0.5061 | 0.0432 | 0 | 0.6210 | 0.34 | 4.96 | 0.0116 | 0 |

| | | | | | | | | | | | | | | | | | | | | | | | | | | | | | | |
|---|---|---|---|---|---|---|---|---|---|---|---|---|---|---|---|---|---|---|---|---|---|---|---|---|---|---|---|---|---|---|
| | | | | | 8 | 8 | | 05 | | | | Match | | 35 | 2 | 47 | 0 | 9 | 44 | | | | 6 | 41 | | 94 | 68 | 02 | 45 | |
| 181 | C03078 | B | FALSE | 4-Guanidinobutanamide | 144.10006 | 0.851 | 0.2 | 2786941 | 0 | | 81.8 | Spectral Match | | 42584.97 | 631362.7 | 249712.3 | 726775.3 | 5.081 | 0.0205 | 0 | 0.67 | 0.0006 | NS | NS | NS | NS | NS | 7.756 | 0.001359 | 0.000118 |
| 182 | C03089 | B | FALSE | 5-Methylthio-D-ribose | 179.0378 | 0.452 | 0.2 | 22961203 | 0 | | | Spectral Match | | 11622957 | 10103568 | 10119184 | 10099664 | NS | NS | NS | NS | NS | NS | NS | NS | NS | NS | 8.5553 | 0.000764 | 0.000235 |
| 183 | C03114 | H | FALSE | Dimethylbenzimidazole | 146.0839 | 1.567 | 0.2 | 1000029 | 0 | | | Spectral Match | Adenosylcobalamin biosynthesis I (anaerobic);Other | 49305.33 | 184412.6 | 159621.5 | 190610.4 | NS | NS | NS | NS | NS | NS | NS | NS | NS | NS | 4.4815 | 0.017215 | 0 |
| 184 | C03137 | FB | FALSE | N-Acetyl-D-tryptophan | 246.0991 | 7.726 | 0.2 | 15693384 | 0 | | | Spectral Match | | 907.2589 | 1841690 | 5472075 | 934093.2 | 886.02 | 0.0000 | 0.000421 | 0.945 | 0.0000 | NS | NS | NS | NS | NS | 4.418 | 0.01814 | 0.000267 |
| 185 | C03175 | H | FALSE | [Similar to: 3-Iodo-L-tyrosine; ΔMass: 180.9682 Da] | 126.0023 | 22.035 | 0.2 | 4355670 | 0 | | 63.2 | Spectral Match | | 835377.5 | 857531.4 | 741759.7 | 886474.3 | 0.38226 | 0.00001 | 0.001235 | 0.975 | 0.8576 | NS | NS | NS | NS | NS | 20.651 | 5.68E-07 | 0.000118 |
| 186 | C03194 | FB | FALSE | (R)-1-Aminopropan-2-ol | 75.06841 | 0.874 | 0.2 | 227491 | 0 | | | Spectral Match | Other | 16862.02 | 27069.24 | 16253.36 | 29773.22 | NS | NS | NS | NS | NS | NS | NS | NS | NS | NS | 21.283 | 4.14E-07 | 0.001195 |
| 187 | C03210 | B | FALSE | 2_2'-Iminodipropanoate | 161.0685 | 0.379 | 0.2 | 1466673 | 0 | | | Spectral Match | Superpathway of L-lysine degradation;Superpathway of butirocin biosynthesis;Other | 62355.81 | 107300.3 | 85240.77 | 112815.1 | 0.65649 | 0.0193 | 0 | 0.505 | 0.0194 | NS | NS | NS | NS | NS | 6.8205 | 0.002722 | 0.000154 |
| 188 | C03228 | H | FALSE | 3-Hydroxycyclohexanone | 114.0681 | 5.088 | 0.2 | 2306873 | 0 | | | Spectral Match | | 322354.9 | 544567.9 | 479962.4 | 560719.3 | NS | NS | NS | NS | NS | NS | NS | NS | NS | NS | 11.525 | 0.000102 | 0.000484 |

| | | | | | | | | | | | | | | | | | | | | | | | | | | | | | | |
|---|---|---|---|---|---|---|---|---|---|---|---|---|---|---|---|---|---|---|---|---|---|---|---|---|---|---|---|---|---|---|
| 189 | C03232 | B | FALSE | 3-Phosphonooxypyruvate | 183.9779 | 0.651 | 0.2 | 305262.8 | 0 | | | Spectral Match | Other | 17697.62 | 51898.67 | 22760.8 | 59183.14 | NS | NS | NS | NS | NS | NS | NS | NS | NS | NS | 4.47226 | 0.017341 | 0 |
| 190 | C03296 | H | FALSE | N2-Succinyl-L-arginine | 274.1274 | 2.903 | 0.2 | 2247653 | 0 | | 90.7 | Spectral Match | | 1347.408 | 535273.7 | 806323.6 | 467511.2 | 155.91 | 0.0011 | 0.001594 | 1 | 0.0000 | NS | NS | NS | NS | NS | 23.986 | 1.13E-07 | 0.000125 |
| 191 | C03298 | B | FALSE | Nalpha_Nalpha-Dimethyl-L-histidine | 183.1001 | 1.037 | 0.2 | 1907794 | 0 | | | Spectral Match | | 70522.99 | 326928.8 | 24273.84 | 402592.5 | NS | NS | NS | NS | NS | 9.9509 | 4.81E-06 | 0.002261 | 1 | 0.0000 | 15.426 | 9.48E-06 | 0.000667 |
| 192 | C03338 | FB | FALSE | 2_3_5-Trihydroxytoluene | 140.0469 | 1.073 | 0.2 | 22339676 | 0 | | 79.7 | Spectral Match | | 442788.4 | 451990.7 | 189008.5 | 517736.3 | 3.4373 | 0.0036 | 0.000211 | 0.955 | 0.0000 | NS | NS | NS | NS | NS | 6.8862 | 0.002591 | 0.00037 |
| 193 | C03341 | FB | FALSE | (2R_4S)-2_4-Diaminopentanoate | 132.0898 | 0.908 | 0.2 | 53242295 | 0 | | | Spectral Match | Arginine, ornithine and proline interconversion;Superpathway of L-arginine and L-ornithine degradation;Other | 1026642 | 11801124 | 3736909 | 13817178 | 3.76 | 0.0022 | 0.000118 | 0.89 | 0.0000 | 2.1611 | 0.03656 | 0 | 0.871094 | 0.0060 | 31.776 | 3.94E-09 | 0.000433 |
| 194 | C03373 | H | FALSE | Aminoimidazoleribotide | 295.0575 | 0.802 | 0.2 | 7764473 | 0 | | | Spectral Match | Other | 1510411 | 1644848 | 3145055 | 1269796 | NS | NS | NS | NS | NS | NS | NS | NS | NS | NS | 15.682 | 8.19E-06 | -0.00013 |
| 195 | C03410 | B | FALSE | N-Glycoloyl-neuraminate | 326.1086 | 16.97 | 0.2 | 1080140 | 0 | | | Spectral Match | | 144906.4 | 172462.5 | 121929.5 | 185095.7 | 0.51368 | 0.0061 | 0 | 1 | 0.8085 | NS | NS | NS | NS | NS | 52.219 | 4.07E-12 | 0.000222 |
| 196 | C03427 | B | FALSE | T-2 Triol | 360.2131 | 11.244 | 0.2 | 1257314 | 0 | 54.9 | | Spectral Match | | 31355.91 | 47268.51 | 85883.99 | 37614.64 | 0.63645 | 0.0167 | 0.000143 | 0.885 | 0.0527 | 0.28815 | 1.10E-16 | 0.00314 | 1 | 0.0000 | 4.3602 | 0.019029 | 0 |

| | | | | | | | | | | | | | | | | | | | | | | | | | | | | | | |
|---|---|---|---|---|---|---|---|---|---|---|---|---|---|---|---|---|---|---|---|---|---|---|---|---|---|---|---|---|---|---|
| 197 | C03516 | B | FALSE | Magnesium protoporphyrin | 583.2196 | 10.17 | 0.2 | 125441.7 | 0 | | | Spectral Match | | 69.39795 | 28484.96 | 23805.67 | 29654.78 | 157.48 | 0.0000 | 0.003756 | 1 | 0.0000 | NS | NS | NS | NS | NS | 23.177 | 1.65E-07 | -7.35E-06 |
| 198 | C03539 | H | FALSE | S-Ribosyl-L-homocysteine | 267.0771 | 8.226 | 0.2 | 271900.8 | 0 | | | Spectral Match | Other | 30959.25 | 42820.09 | 58282.26 | 38954.55 | 0.57415 | 0.0027 | 0 | 0.78 | 0.0632 | 0.44424 | 2.13E-07 | 5.00E-04 | 0.808594 | 0.0023 | 10.714 | 0.000174 | 0 |
| 199 | C03590 | H | FALSE | 2-(4-Hydroxyphenyl)-2-oxoacetate | 166.0262 | 16.644 | 0.2 | 1624971 | 0 | | 82.1 | Spectral Match | Superpathway of aromatic compound degradation via 2-oxopent-4-enoate;Superpathway of aromatic compound degradation via 3-oxoadipate | 301889.8 | 249255.8 | 177963.4 | 267078.9 | 0.32438 | 0.0000 | 0.001875 | 0.54 | 0.1836 | NS | NS | NS | NS | NS | 8.4783 | 0.000807 | 0.00062 |
| 200 | C03642 | B | FALSE | Taurolithocholate sulfate | 563.7401 | 0.828 | 0.2 | 240203.9 | 0 | | | Spectral Match | | 90835.03 | 51269.25 | 59214 | 49283.06 | 0.22496 | 0.0084 | 0.000133 | 0.705 | 0.7138 | 0.54957 | 0.047018 | 4.00E-04 | 0.609375 | 0.5819 | 4.7119 | 0.014249 | 0 |
| 201 | C03652 | H | FALSE | (2R_3S)-2_3-Dimethylmalate | 162.0522 | 0.943 | 0.2 | 1081626 | 0 | | 86.3 | Spectral Match | Superpathway of butirocin biosynthesis;Other | 179577 | 206520.2 | 332490.9 | 175027.5 | 0.44396 | 0.0088 | -9.52E-05 | 0.51 | 0.8446 | 0.33456 | 0.000239 | 0 | 0.675781 | 0.0805 | 3.9879 | 0.025959 | 0.000118 |
| 202 | C03656 | B | FALSE | [FAoxo_amino(6:0)]3-oxo-5S-amino-hexanoicacid | 145.0733 | 1.219 | 0.2 | 1480837 | 0 | | | Spectral Match | Superpathway of L-lysine degradation;Biotin biosynthesis II | 35784.77 | 149963 | 192611.4 | 139300.9 | NS | NS | NS | NS | NS | 0.48694 | 0.001321 | 0.000211 | 0.632813 | 0.1800 | 131.64 | 8.12E-19 | 0.006414 |
| 203 | C03657 | H | FALSE | 1,4-Dihydroxy-2-naphthoate | 101.0142 | 0.367 | 0.2 | 494889.2 | 0 | | | Spectral Match | | 174844 | 169313 | 127241.8 | 179830.8 | NS | NS | NS | NS | NS | NS | NS | NS | NS | NS | 16.655 | 4.73E-06 | 0.001334 |
| 2 | C03 | HF | FA | 3-(4- | 18 | 2. | 0. | 37 | 0 | | | Spe | | 26 | 74 | 73 | 75 | 9. | 0. | 0. | 0. | 0. | NS | NS | NS | NS | N | 2 | 4. | 0 |

| | | | | | | | | | | | | | | | | | | | | | | | | | | | | | | |
|---|---|---|---|---|---|---|---|---|---|---|---|---|---|---|---|---|---|---|---|---|---|---|---|---|---|---|---|---|---|---|
| 04 | 672 | B | LSE | Hydroxyphenyl)lactate | 2.0559 | 641 | 2 | 08933 | | | | ctral Match | | 232.88 | 7501.8 | 0211 | 1824.5 | 6106 | 0001 | 000286 | 54 | 0000 | | | | | S | 1.032 | 69E-07 | |
| 205 | C03684 | H | FALSE | 6-Pyruvoyl-5,6,7,8-tetrahydropterin | 236.0787 | 1.192 | 0.2 | 188866.1 | 0 | | | Spectral Match | Superpathway of NAD biosynthesis in eukaryotes;Superpathway of aromatic compound degradation via 2-oxopent-4-enoate;Superpathway of aromatic compound degradation via 3-oxoadipate;Other | 28010.09 | 37463.76 | 43168.58 | 36037.55 | 0.53042 | 0.0067 | 0 | 1 | 0.2862 | 0.53453 | 0.001404 | 0 | 0.558594 | 0.4295 | 5.1958 | 0.009633 | 0.000489 |
| 206 | C03690 | B | FALSE | Bis(2-ethylhexyl) phthalate | 390.2753 | 16.817 | 0.082 | 2471094 | 0 | 78.4 | | Spectral Match | | 296665.4 | 550955.5 | 495006.8 | 564942.7 | NS | NS | NS | NS | NS | NS | NS | NS | NS | NS | 35.126 | 1.08E-09 | 0.000133 |
| 207 | C03752 | B | FALSE | [Similar to: Isoprenaline; ΔMass: 17.0548 Da] | 194.0661 | 0.875 | 0.2 | 221143 | 0 | | 81.5 | Spectral Match | | 23748.98 | 21838.02 | 18630.4 | 22639.92 | 0.36118 | 0.0000 | 0.000333 | 0.62 | 0.4976 | NS | NS | NS | NS | NS | 15.294 | 1.02E-05 | 0.001838 |
| 208 | C03758 | HB | FALSE | Dopamine | 153.0785 | 2.925 | 0.2 | 27436933 | 0 | | | Spectral Match | Other | 10225.22 | 6012124 | 9629540 | 5107770 | 236.77 | 0.0071 | 0.002523 | 1 | 0.0005 | NS | NS | NS | NS | NS | 6.996 | 0.002386 | 0 |
| 209 | C03771 | HB | FALSE | 5-Guanidino-2-oxopentanoate | 172.0734 | 7.49 | 0.2 | 478435.6 | 0 | | | Spectral Match | | 92377.4 | 86063.92 | 79444.71 | 87718.72 | 0.36324 | 0.0000 | 0.000267 | 0.8 | 0.5551 | NS | NS | NS | NS | NS | 23.391 | 1.49E-07 | 0.001453 |
| 210 | C03827 | B | FALSE | 2-Dehydro-3-deoxy-L-fuconate | 163.060 | 0.68 | 0.2 | 173477 | 0 | | | Spectral | | 114627 | 247452 | 140957 | 274076 | NS | NS | NS | NS | NS | NS | NS | NS | NS | NS | 13.8 | 2.36E- | 0.001194 |

| | | | | | | | | | | | | | | | | | | | | | | | | | | | | | |
|---|---|---|---|---|---|---|---|---|---|---|---|---|---|---|---|---|---|---|---|---|---|---|---|---|---|---|---|---|---|
| | | | | | 3 | | | 14 | | | Match | | 2 | 4 | 4 | 2 | | | | | | | | | | | 8 | 05 | |
| 211 | C03840 | H | FALSE | 5-Hydroxy-1_4-naphthoquinone | 174.0323 | 25.039 | 0.2 | 1043409 | 0 | | Spectral Match | | 324008.3 | 159211.6 | 22859.67 | 193299.6 | 0.18298 | 0.0487 | 0.000558 | 0.58 | 0.7062 | 5.7629 | 0.001245 | 0 | 0.875 | 0.0000 | 42.799 | 7.34E-11 | 0.001897 |
| 212 | C03846 | FB | FALSE | 6-Acetamido-3-aminohexanoate | 188.1154 | 1.08 | 0.2 | 7211479 | 0 | | Spectral Match | Superpathway of L-lysine degradation | 65919.52 | 1009280 | 76200.91 | 1242550 | 5.2331 | 0.0494 | 0 | 1 | 0.0014 | 10.05 | 5.99E-07 | 0.002581 | 0.992188 | 0.0000 | 6.2343 | 0.004258 | 0.000308 |
| 213 | C03912 | H | FALSE | (S)-1-Pyrroline-5-carboxylate | 113.0476 | 1.043 | 0.2 | 3721943 | 0 | | Spectral Match | Arginine, ornithine and proline interconversion;Other | 357511.1 | 904017.2 | 774217.1 | 936467.3 | NS | NS | NS | NS | NS | NS | NS | NS | NS | NS | 20.507 | 6.11E-07 | 0.000133 |
| 214 | C03954 | H | FALSE | N-Hydroxy-2-acetamidofluorene | 239.0937 | 10.11 | 0.2 | 740474.8 | 0 | 65.2 | Spectral Match | | 1001.264 | 103235.3 | 2104.529 | 128517.9 | 35.272 | 0.0003 | 0.000286 | 0.92 | 0.0000 | 38.386 | 1.52E-12 | 0.003927 | 1 | 0.0000 | 17.417 | 3.10E-06 | 0.000967 |
| 215 | C03982 | H | FALSE | 2-MethylpropanalO-methyloxime | 101.0842 | 3.599 | 0.2 | 528105 | 0 | 74.5 | Spectral Match | Superpathway of linamarin and lotaustralin biosynthesis;Other | 39593.49 | 67923.29 | 88507.36 | 62777.28 | NS | NS | NS | NS | NS | 0.47972 | 0.0004 | 0.000333 | 0.648438 | 0.0789 | 15.229 | 1.06E-05 | 0.000222 |
| 216 | C03986 | H | FALSE | 3-Hydroxy-4-methylanthranilate | 167.0578 | 0.348 | 0.2 | 194383 | 0 | | Spectral Match | Other | 8755.505 | 8852.367 | 6893.456 | 9342.095 | 0.4027 | 0.0000 | 0.001505 | 1 | 0.9981 | NS | NS | NS | NS | NS | 31.1 | 5.16E-09 | 0.000618 |
| 217 | C03990 | H | FALSE | [Similar to: Lithocholic acid; ΔMass: -15.9545 Da] | 392.2522 | 13.189 | 0.2 | 239460.7 | 0 | 77.3 | Spectral Match | | 14902.93 | 36036.58 | 29690.47 | 37623.11 | NS | NS | NS | NS | NS | NS | NS | NS | NS | NS | 4.7768 | 0.013514 | 0.00023 |
| 218 | C04036 | H | FALSE | 1-palmitoylglycerol;MAG (16:0) | 330.2757 | 15.198 | 0.2 | 3713538 | 0 | | Spectral Match | | 7871.768 | 5378.703 | 5280.843 | 5403.168 | 0.26808 | 0.0000 | 0.000767 | 0.53 | 0.0230 | NS | NS | NS | NS | NS | 32.313 | 3.18E-09 | 0.003061 |
| 2 | C04 | H | FA | 1- | 49 | 1 | 0. | 92 | 0 | 61 | Spe | | 73 | 18 | 64 | 21 | NS | N | NS | N | N | NS | NS | NS | NS | N | 7. | 0. | 0 |

| | | | | | | | | | | | | | | | | | | | | | | | | | | | | | | |
|---|---|---|---|---|---|---|---|---|---|---|---|---|---|---|---|---|---|---|---|---|---|---|---|---|---|---|---|---|---|---|
| 19 | 102 | | LSE | Palmitoylglycerophosphocholine | 5.303 | 3.842 | 2 | 73321 | | | .6 | ctral Match | | 3547.8 | 21944 | 2369.4 | 16837 | | S | | S | S | | | | | S | 3563 | 001824 | |
| 220 | C04122 | B | FALSE | D-1-Aminopropan-2-olO-phosphate | 155.0344 | 20.1388 | 0.2 | 1626625 | 0 | | | Spectral Match | Superpathway of L-threonine metabolism;Adenosylcobalamin biosynthesis I (anaerobic);Other | 1017.766 | 4415.866 | 6634.206 | 3861.281 | 1.8138 | 0.02776 | 0 | 0.765 | 0.0000 | 0.39103 | 6.42E-12 | 8.00E-04 | 1 | 0.0000 | 17.179 | 3.53E-06 | 0.000267 |
| 221 | C04146 | H | FALSE | all-trans-Octaprenyldiphosphate | 360.2131 | 13.533 | 0.2 | 6169174 | 0 | | | Spectral Match | | 91754.18 | 101208.5 | 190353.2 | 78922.39 | 0.47222 | 0.0009 | 0.000143 | 0.655 | 0.9033 | 0.27534 | 5.36E-16 | 0.004423 | 1 | 0.0000 | 4.61 | 0.015488 | 0 |
| 222 | C04148 | H | FALSE | Phenylacetylglutamine | 264.1096 | 5.44 | 0.2 | 3860719 | 0 | | | Spectral Match | | 47628.89 | 445906.8 | 1324611 | 226230.7 | NS | NS | NS | NS | NS | 0.11473 | 0.00314 | 0 | 0.65625 | 0.0596 | 17.089 | 3.71E-06 | 0.000143 |
| 223 | C04164 | H | FALSE | trans-Cinnamoylbeta-D-glucoside | 310.1048 | 0.859 | 0.2 | 206030.8 | 0 | | | Spectral Match | | 2801.065 | 54266.96 | 60134.27 | 52800.14 | 7.614 | 0.0001 | 0.001286 | 1 | 0.0000 | NS | NS | NS | NS | NS | 22.649 | 2.12E-07 | 0.000493 |
| 224 | C04166 | H | FALSE | (+/-)-6-Hydroxy-3-oxo-alpha-ionol | 224.1404 | 10.664 | 0.2 | 502433.7 | 0 | | | Spectral Match | | 19377.69 | 23267.54 | 19250.84 | 24271.71 | 0.48084 | 0.0000 | 0 | 0.975 | 0.0525 | NS | NS | NS | NS | NS | 46.598 | 2.18E-11 | 0.001952 |
| 225 | C04210 | H | FALSE | N5-(L-1-Carboxyethyl)-L-ornithine | 204.1103 | 0.972 | 0.2 | 4078940 | 0 | | | Spectral Match | | 328743.6 | 615697.9 | 380791.6 | 674424.5 | NS | NS | NS | NS | NS | NS | NS | NS | NS | NS | 3.6406 | 0.034828 | 0 |
| 226 | C04223 | FB | FALSE | (+/-)-6-Hydroxy-3-oxo-alpha-ionone | 222.1249 | 7.69 | 0.2 | 430924 | 0 | | 65.3 | Spectral Match | | 602.2742 | 1566.247 | 2596.453 | 1308.695 | NS | NS | NS | NS | NS | 0.33792 | 5.66E-14 | 0.002484 | 1 | 0.0000 | 4.3816 | 0.018693 | 0 |
| 227 | C04236 | H | FALSE | [Similar to: Choline; ΔMass: | 86.0191 | 1.06 | 0.2 | 231834 | 0 | | 87.2 | Spectral | | 23585.1 | 582918. | 875422. | 509792 | 9.3222 | 0.00 | 0.0002 | 0.965 | 0.0000 | NS | NS | NS | NS | NS | 6.10 | 0.0047 | 0.000421 |

| | | | | | | | | | | | | | | | | | | | | | | | | | | | | | | |
|---|---|---|---|---|---|---|---|---|---|---|---|---|---|---|---|---|---|---|---|---|---|---|---|---|---|---|---|---|---|---|
| | | | | 17.0806 Da] | 1 | 2 | | 7 | | | | Match | | 2 | 2 | 8 | | | 04 | 86 | | | | | | | | 04 | 22 | |
| **228** | C04277 | FB | FALSE | 1_8-Diazacyclotetradecane-2_9-dione | 226.1673 | 3.659 | 0.2 | 3322280 | 0 | | | Spectral Match | | 4135276 | 6374614 | 3034426 | 7209661 | NS | NS | NS | NS | NS | NS | NS | NS | NS | NS | 7.3017 | 0.001899 | 0.000375 |
| **229** | C04282 | H | FALSE | 1-Pyrroline-4-hydroxy-2-carboxylate | 129.0423 | 1.074 | 0.049 | 43510869 | 0 | | | Spectral Match | Superpathway of butirocin biosynthesis;Other | 1456343 | 11238912 | 11385958 | 11202150 | 2.9308 | 0.0078 | 0 | 0.975 | 0.0000 | NS | NS | NS | NS | NS | 10 | 0.000281 | 0.000471 |
| **230** | C04284 | H | FALSE | 2-Heptyl-4-hydroxyquinoline-N-oxide | 259.1563 | 6.096 | 0.2 | 346082.2 | 0 | | | Spectral Match | Other | 3033.967 | 66479.25 | 31882.08 | 75128.55 | 7.0281 | 0.0001 | 0.00145 | 0.77 | 0.0000 | NS | NS | NS | NS | NS | 9.9929 | 0.000282 | 0.001503 |
| **231** | C04303 | FB | FALSE | N-Benzoyl-D-arginine-4-nitroanilide | 398.1686 | 9.312 | 0.2 | 137039.9 | 0 | | | Spectral Match | | 11293.29 | 22729.08 | 14788.9 | 24714.13 | NS | NS | NS | NS | NS | NS | NS | NS | NS | NS | 87.758 | 1.00E-15 | 0.004446 |
| **232** | C04332 | HFB | FALSE | [Similar to: Dodecyl sulfate; ΔMass: -58.9604 Da] | 325.1156 | 20.404 | 0.2 | 392844.1 | 0 | | 75.2 | Spectral Match | | 34681.98 | 51688.54 | 89630.22 | 42203.12 | 0.61556 | 0.0300 | 0 | 0.97 | 0.1241 | 0.30828 | 2.25E-09 | 0.003441 | 0.964844 | 0.0000 | 23.572 | 1.37E-07 | 0.000353 |
| **233** | C04352 | FB | FALSE | N-[(R)-4'-Phosphopantothenoyl]-L-cysteine | 200.0362 | 1.087 | 0.2 | 4459100 | 0 | | | Spectral Match | | 3242.478 | 688542.5 | 220055.2 | 805664.3 | 63.295 | 0.0000 | 0.002161 | 0.995 | 0.0000 | NS | NS | NS | NS | NS | 92.153 | 4.36E-16 | 0.008441 |
| **234** | C04377 | B | FALSE | 5,10-Methylenetetrahydromethanopterin | 393.1246 | 9.264 | 0.2 | 2400589 | 0 | | | Spectral Match | | 67558.75 | 44896.53 | 42532.5 | 45487.54 | 0.2715 | 0.0000 | 0.000308 | 0.75 | 0.0037 | NS | NS | NS | NS | NS | 26.784 | 3.18E-08 | 0 |
| **235** | C04405 | B | FALSE | (2S,3S)-3-Hydroxy-2-methylbutanoyl-CoA | 290.0627 | 10.936 | 0.2 | 6379688 | 0 | | | Spectral Match | | 80467.54 | 240466 | 239407.1 | 240730.7 | NS | NS | NS | NS | NS | NS | NS | NS | NS | NS | 12.929 | 4.21E-05 | 0.000432 |
| **23** | C04431 | B | FALS | [Similar to: Tropine; | 139.0 | 0.9 | 0.11 | 5171 | 0 | | 88.2 | Spectra | | 3806 | 9348 | 8246 | 9624 | NS | NS | NS | NS | NS | NS | NS | NS | NS | NS | 11. | 8.25 | 0 |

| | | | | | | | | | | | | | | | | | | | | | | | | | | | | | | |
|---|---|---|---|---|---|---|---|---|---|---|---|---|---|---|---|---|---|---|---|---|---|---|---|---|---|---|---|---|---|---|
| 6 | | | E | ΔMass: 2.1123 Da] | 031 | 72 | 7 | 326 | | | | l Match | | 97.7 | 66.9 | 25.7 | 27.1 | | | | | | | | | | | 861 | E-05 | |
| 237 | C04483 | H | FALSE | Deoxycholic acid | 392.2904 | 12.144 | 0.2 | 436096.3 | 0 | 79.5 | | Spectral Match | | 7063.581 | 50751.81 | 28221.28 | 56384.44 | 2.5323 | 0.0018 | 0.000105 | 1 | 0.0000 | NS | NS | NS | NS | NS | 6.90884 | 0.002548 | 0.000316 |
| 238 | C04594 | H | FALSE | (9Z)-(13S)-12_13-Epoxyoctadeca-9_11-dienoicacid | 294.2185 | 10.424 | 0.2 | 177781.7 | 0 | | | Spectral Match | Other | 3521.611 | 5678.748 | 6025.322 | 5592.105 | 0.65652 | 0.0117 | 0 | 1 | 0.0013 | 0.629 | 0.001741 | 0 | 0.519531 | 0.5926 | 10.472 | 0.000204 | 0 |
| 239 | C04646 | H | FALSE | 6-Thioinosine-5'-monophosphate | 183.0202 | 5.074 | 0.2 | 840478.7 | 0 | | | Spectral Match | | 119196.2 | 195405.9 | 172041.1 | 201247.1 | NS | NS | NS | NS | NS | NS | NS | NS | NS | NS | 3.9187 | 0.027515 | 0 |
| 240 | C04670 | H | FALSE | (1S_4S)-4-Hydroxy-3-oxocyclohexane-1-carboxylate | 158.0575 | 15.89 | 0.2 | 332515.6 | 0 | | | Spectral Match | Superpathway of branched chain amino acid biosynthesis | 1426.159 | 2885.462 | 1714.293 | 3178.254 | NS | NS | NS | NS | NS | NS | NS | NS | NS | NS | 5.868 | 0.005658 | 0.000211 |
| 241 | C04722 | FB | FALSE | 3alpha,7alpha,12alpha-Trihydroxy-5beta-cholestanoate | 448.3179 | 12.999 | 0.2 | 447199.4 | 0 | | | Spectral Match | | 63655.66 | 76930.24 | 12801.45 | 92962.43 | 0.435 | 0.0092 | 0 | 0.925 | 0.8059 | 4.6617 | 2.12E-05 | 0.001107 | 0.96875 | 0.0000 | 5.2697 | 0.009079 | 0 |
| 242 | C04732 | HFB | FALSE | [Similar to: 3-Pyridylamidoxime; ΔMass: -2.0017 Da] | 139.0606 | 20.261 | 0.2 | 1259477 | 0 | | 82.7 | Spectral Match | | 24241.57 | 22608.43 | 31853.54 | 20297.15 | 0.38123 | 0.0002 | 0 | 0.85 | 0.6141 | 0.43912 | 1.24E-05 | 0.0000774 | 0.769531 | 0.0073 | 14.351 | 1.78E-05 | 0.0001109 |
| 243 | C04874 | H | FALSE | [Similar to: 4-Amino-7-phenylpyrazolo[5,1-c][1,2,4]triazine-3-carboxamide; ΔMass: 0.0030 Da] | 254.0886 | 12.287 | 0.2 | 31059376 | 0 | | 77.8 | Spectral Match | | 26469.01 | 48185.48 | 10599597 | 3373286 | 73.124 | 0.0000 | 0.003292 | 0.69 | 0.0000 | 0.20886 | 0.038026 | 0 | 0.605469 | 0.2081 | 85.466 | 1.57E-15 | 0.0003161 |
| 2 | C04 | B | FA | 1-(4- | 26 | 2. | 0. | 16 | 0 | | 76 | Spe | | 93 | 33 | 22 | 36 | NS | N | NS | N | N | NS | NS | NS | NS | N | 1 | 9. | 0.00 |

| | | | | | | | | | | | | | | | | | | | | | | | | | | | | | | |
|---|---|---|---|---|---|---|---|---|---|---|---|---|---|---|---|---|---|---|---|---|---|---|---|---|---|---|---|---|---|---|
| 44 | 908 | | LSE | hydroxy-2-methylpyrimid-5-ylmethyl)-3-(β-hydroxyethyl)-2-methylpyridiniumbromide | 0.1402 | 455 | 2 | 6942.6 | | | .5 | ctral Match | | 42.002 | 950.86 | 218.23 | 884.02 | | S | | S | S | | | | | S | 30.65 | 31E-19 | 8027 |
| 245 | C04919 | B | FALSE | Lipid IVA | 1404.849 | 13.653 | 0.2 | 26160098 | 0 | | | Spectral Match | Other | 82832.88 | 53553.71 | 42106.76 | 56415.45 | 0.25029 | 0.0000 | 0.001711 | 0.805 | 0.0079 | NS | NS | NS | NS | NS | 5.2063 | 0.009552 | 0 |
| 246 | C04932 | FB | FALSE | Lipid A disaccharide | 661.437 | 14.427 | 0.2 | 2323396 | 0 | | | Spectral Match | | 60066 | 39453.02 | 22412.23 | 43713.22 | 0.25584 | 0.0000 | 5.00E-04 | 0.915 | 0.0039 | NS | NS | NS | NS | NS | 36.41 | 6.73E-10 | 0.001412 |
| 247 | C05021 | H | FALSE | N-Acetyl-beta-D-galactosamine | 221.0893 | 1.016 | 0.2 | 1111736 | 0 | | | Spectral Match | Other | 122226.6 | 220483.6 | 237073.8 | 216336.1 | 0.66663 | 0.0223 | 5.68E-05 | 0.57 | 0.0796 | 0.56609 | 0.002211 | 0 | 0.53125 | 0.6465 | 4.6461 | 0.015036 | 0 |
| 248 | C05042 | H | FALSE | Phosphinothricin | 181.0501 | 20.211 | 0.2 | 972238.4 | 0 | | | Spectral Match | | 307.031 | 2545.437 | 3139.255 | 2396.982 | 3.389 | 0.0000 | 0.000941 | 0.525 | 0.0000 | 0.51321 | 2.02E-08 | 0.002341 | 0.996094 | 0.0000 | 12.058 | 7.27E-05 | 0.000915 |
| 249 | C05076 | H | FALSE | Imazaquin | 311.1269 | 5.956 | 0.2 | 3648631 | 0 | | | Spectral Match | | 989.8702 | 47987.62 | 1979.213 | 59489.73 | 16.137 | 0.0001 | 0 | 1 | 0.0000 | 18.348 | 4.28E-19 | 0.003338 | 1 | 0.0000 | 11.185 | 0.000128 | 0.001094 |
| 250 | C05100 | H | FALSE | 3-Ureidoisobutyrate | 146.0687 | 1.16 | 0.2 | 1998261 | 0 | | | Spectral Match | Superpathway of chorismate metabolism;Superpathway of aromatic amino acid biosynthesis;Superpathway of NAD biosynthesis in | 95670.25 | 511364.2 | 360796.2 | 549006.2 | NS | NS | NS | NS | NS | NS | NS | NS | NS | NS | 7.7604 | 0.001355 | 0.000125 |

| | | | | | | | | | | | | | | | | | | | | | | | | | | | | | | |
|---|---|---|---|---|---|---|---|---|---|---|---|---|---|---|---|---|---|---|---|---|---|---|---|---|---|---|---|---|---|---|
| | | | | | | | | | | | | | eukaryotes;Superpathway of indole-3-acetate conjugate biosynthesis;Superpathway of jasmonoyl-amino acid conjugates biosynthesis;Superpathway of butirocin biosynthesis;Adenosylcobalamin biosynthesis I (anaerobic);Other | | | | | | | | | | | | | | | | | |
| 251 | C05122 | FB | FALSE | [Similar to: 3,4-Dimethyl-2,5-bis(3,4,5-trimethoxyphenyl)tetrahydrofuran; ΔMass: -83.0774 Da] | 515.2922 | 7.307 | 0.2 | 447631.5 | 0 | | 72 | Spectral Match | | 382825.51 | 980339.34 | 965529.83 | 984116.72 | NS | NS | NS | NS | NS | NS | NS | NS | NS | NS | 32.0299 | 3.56E-09 | 0.000679 |
| 252 | C05130 | HB | FALSE | Imidazole-4-acetaldehyde | 110.048 | 1.041 | 0.135 | 1326031 | 0 | | | Spectral Match | | 84076.83 | 226772 | 697555.97 | 266026 | NS | NS | NS | NS | NS | 2.2911 | 0.000484 | 0 | 0.964844 | 0.0000 | 9.4595 | 0.000406 | 3.33E-05 |
| 253 | C05172 | H | FALSE | Selenophosphoric acid | 79.94221 | 0.926 | 0.2 | 396003.3 | 0 | | | Spectral Match | | 639999.52 | 1111162.9 | 895570.53 | 116561 | NS | NS | NS | NS | NS | NS | NS | NS | NS | NS | 73.4774 | 1.93E-14 | 0.0006443 |
| 254 | C05178 | H | FALSE | (R)-Reticuline | 329.1615 | 6.853 | 0.2 | 91043.54 | 0 | | | Spectral Match | | 241.7814 | 141241.22 | 564.5722 | 176660.39 | 18.614 | 0.0001 | 0.00223 | 1 | 0.0000 | 18.07 | 1.10E-17 | 0.00089 | 1 | 0.0000 | 15.197 | 1.08E-05 | 0.0002886 |

| | | | | | | | | | | | | | | | | | | | | | | | | | | | | | | |
|---|---|---|---|---|---|---|---|---|---|---|---|---|---|---|---|---|---|---|---|---|---|---|---|---|---|---|---|---|---|---|
| 255 | C05191 | H | FALSE | Dihydrosanguinarine | 333.0994 | 9.839 | 0.2 | 348330.6 | 0 | | | Spectral Match | | 1027.558 | 47519.48 | 2720.137 | 58719.32 | 15.482 | 0.0000 | 0.000211 | 1 | 0.0000 | 13.217 | 7.19E-13 | 0.004519 | 1 | 0.0000 | 7.7753 | 0.00134 | 0.000143 |
| 256 | C05204 | H | FALSE | [FA(16:2)]N-hexadecyl-ethanolamine | 299.2811 | 14.556 | 0.2 | 18392907 | 0 | | | Spectral Match | Other | 283500.7 | 444387.9 | 477603.2 | 436084 | 0.632219 | 0.0022 | 0 | 1 | 0.0000 | 0.61934 | 2.26E-05 | 0.001685 | 0.691406 | 0.0834 | 8.9939 | 0.000561 | -0.00011 |
| 257 | C05229 | B | FALSE | Phaseollidin hydrate | 342.1453 | 13.152 | 0.2 | 1293917 | 0 | | | Spectral Match | | 45288.17 | 55832.41 | 94070.78 | 46272.81 | 0.51981 | 0.0018 | -0.00012 | 1 | 0.3930 | 0.32953 | 1.04E-11 | 0.002014 | 0.996094 | 0.0000 | 14.785 | 1.38E-05 | 0.000646 |
| 258 | C05306 | H | FALSE | Chlorophyll a | 891.5301 | 13.348 | 0.2 | 167039.3 | 0 | | | Spectral Match | | 7949.008 | 22937.17 | 18320.48 | 24091.34 | NS | NS | NS | NS | NS | NS | NS | NS | NS | NS | 21.838 | 3.15E-07 | 0.00075 |
| 259 | C05318 | H | FALSE | (S)-7_8_13_14-Tetrahydroprotoberberine | 235.1351 | 9.525 | 0.2 | 494887.4 | 0 | | | Spectral Match | | 46117.98 | 41962.59 | 29739.89 | 45018.27 | 0.35869 | 0.0000 | 0 | 0.635 | 0.3135 | NS | NS | NS | NS | NS | 6.8025 | 0.00276 | 0 |
| 260 | C05328 | HB | FALSE | D-2-Amino-hexano-6-lactam | 128.0947 | 0.808 | 0.2 | 650033.4 | 0 | | | Spectral Match | | 86407.9 | 143723.5 | 148906.8 | 142427.6 | NS | NS | NS | NS | NS | 0.61683 | 0.02669 | 0 | 0.511719 | 0.6600 | 6.2071 | 0.004348 | 0.000887 |
| 261 | C05341 | HFB | FALSE | beta-Alanyl-L-lysine | 216.1344 | 9.568 | 0.2 | 2862883 | 0 | | | Spectral Match | | 49262.24 | 93564.5 | 86895.65 | 95231.72 | NS | NS | NS | NS | NS | NS | NS | NS | NS | NS | 69.918 | 4.31E-14 | 0.003876 |
| 262 | C05345 | HFB | FALSE | beta-D-Fructose 6-phosphate | 260.0287 | 0.791 | 0.2 | 632283.7 | 0 | | | Spectral Match | Superpathway of butirocin biosynthesis;Other | 42875.5 | 116911.6 | 29890.02 | 138667 | NS | NS | NS | NS | NS | NS | NS | NS | NS | NS | 17.335 | 3.24E-06 | 0.001 |
| 263 | C05382 | HFB | FALSE | Sedoheptulose 7-phosphate | 144.0128 | 21.768 | 0.2 | 895111.8 | 0 | | | Spectral Match | | 81051.59 | 153700.7 | 122621.4 | 161470.5 | NS | NS | NS | NS | NS | NS | NS | NS | NS | NS | 23.427 | 1.47E-07 | 0.00118 |

| | | | | | | | | | | | | | | | | | | | | | | | | | | | | | | |
|---|---|---|---|---|---|---|---|---|---|---|---|---|---|---|---|---|---|---|---|---|---|---|---|---|---|---|---|---|---|---|
| 264 | C05399 | B | FALSE | Melibiitol | 343.1252 | 1.066 | 0.2 | 807723.6 | 0 | | | Spectral Match | | 13178.37 | 138990.8 | 51895.68 | 160764.6 | 3.621 | 0.0126 | 0 | 1 | 0.0001 | NS | NS | NS | NS | NS | 100.13 | 1.04E-16 | 0.001401 |
| 265 | C05427 | H | FALSE | [Similar to: Amiloride; ΔMass: 1.9352 Da] | 227.1127 | 1.921 | 0.2 | 1107863 | 0 | | 82.9 | Spectral Match | | 2731.971 | 274872.8 | 181051.8 | 298328.1 | 35.638 | 0.0062 | 0.001141 | 1 | 0.0001 | NS | NS | NS | NS | NS | 9.54 | 0.000384 | 0.000471 |
| 266 | C05465 | B | FALSE | Taurochenodeoxycholate | 498.2914 | 16.111 | 0.2 | 185473.5 | 0 | | | Spectral Match | | 9550.747 | 111097.8 | 1158095 | 1099198 | 43.404 | 0.0000 | 0.000571 | 0.635 | 0.0000 | NS | NS | NS | NS | NS | 11.988 | 7.61E-05 | 0.000111 |
| 267 | C05466 | H | FALSE | Glycochenodeoxycholic acid | 449.3115 | 11.259 | 0.07 | 3284532 | 0 | 79.3 | | Spectral Match | | 10310.53 | 594522.5 | 212571.5 | 690010.3 | 20.291 | 0.0000 | 0.00025 | 1 | 0.0000 | NS | NS | NS | NS | NS | 153.85 | 4.68E-20 | 0.005304 |
| 268 | C05472 | H | FALSE | [Similar to: Everolimus; ΔMass: 591.3398 Da] | 366.2416 | 19.787 | 0.2 | 18262476 | 0 | | 69.1 | Spectral Match | | 2859585 | 3765983 | 2123669 | 4176562 | 0.50142 | 0.0009 | 0.00025 | 1 | 0.3352 | NS | NS | NS | NS | NS | 24.409 | 9.26E-08 | 0.001507 |
| 269 | C05516 | HFB | FALSE | 5-Amino-4-imidazolecarboxylate | 127.038 | 2.062 | 0.2 | 4710592 | 0 | | | Spectral Match | | 847.3296 | 928659.4 | 451487.8 | 1047952 | 385.26 | 0.0000 | 0.000756 | 0.62 | 0.0000 | NS | NS | NS | NS | NS | 36.033 | 7.73E-10 | 0.001565 |
| 270 | C05527 | H | FALSE | 3-Sulfinylpyruvate | 152.9847 | 0.668 | 0.2 | 196887.7 | 0 | | | Spectral Match | | 24802.41 | 45557.65 | 72238.59 | 38887.41 | NS | NS | NS | NS | NS | 0.36371 | 0.014596 | 0 | 0.609375 | 0.1987 | 72.089 | 2.63E-14 | 0.002326 |
| 271 | C05543 | HF | FALSE | 3-Dehydroxycarnitine | 145.1098 | 0.895 | 0.071 | 3126030 | 0 | | | Spectral Match | | 55194.48 | 505051.8 | 933257.2 | 398000.5 | 3.6795 | 0.0032 | 0.000154 | 1 | 0.0000 | 0.28107 | 4.48E-06 | 0.001137 | 0.773438 | 0.0078 | 87.453 | 1.06E-15 | 0.006674 |
| 272 | C05580 | HFB | FALSE | 3,4-Dihydroxymandelate | 183.0307 | 0.686 | 0.2 | 1324856 | 0 | | | Spectral Match | | 100851.9 | 198010.2 | 216578.7 | 193368.1 | NS | NS | NS | NS | NS | 0.55189 | 0.007195 | 0 | 0.613281 | 0.4098 | 34.801 | 1.22E-09 | 0.000693 |

| | | | | | | | | | | | | | | | | | | | | | | | | | | | | | | |
|---|---|---|---|---|---|---|---|---|---|---|---|---|---|---|---|---|---|---|---|---|---|---|---|---|---|---|---|---|---|---|
| **273** | C05584 | H | FALSE | 3-Methoxy-4-hydroxymandelate | 199.0605 | 0.84 | 0.2 | 3932252 | 0 | | | Spectral Match | | 315693.1 | 738941.4 | 1239758 | 613737.3 | NS | NS | NS | NS | NS | NS | NS | NS | NS | NS | 35.341 | 9.98E-10 | 0.000731 |
| **274** | C05585 | H | FALSE | 2_5-Dihydroxybenzaldehyde | 138.0315 | 0.999 | 0.2 | 409947 | 0 | | 77.3 | Spectral Match | Superpathway of chorismate metabolism;Superpathway of aromatic compound degradation via 2-oxopent-4-enoate;Anaerobic aromatic compound degradation (Thauera aromatica);Superpathway of aromatic compound degradation via 3-oxoadipate;Other | 17832.74 | 57801.8 | 16945.98 | 68015.76 | NS | NS | NS | NS | NS | 2.672 | 0.000926 | 0 | 0.996094 | 0.0000 | 6.7119 | 0.002955 | 0 |
| **275** | C05588 | H | FALSE | Metanephrine | 197.1047 | 5.024 | 0.2 | 279531.3 | 0 | | | Spectral Match | | 8737.079 | 8218.28 | 3705.159 | 9346.561 | 0.35 | 0.0000 | 0 | 0.975 | 0.5548 | 1.6496 | 0.001007 | 0.000471 | 1 | 0.0000 | 120.45 | 4.02E-18 | 0.000633 |
| **276** | C05596 | H | FALSE | 4-Hydroxyphenylacetylglycine | 209.068 | 1.052 | 0.2 | 928457.9 | 0 | | | Spectral Match | | 43624.64 | 157622.6 | 96814.8 | 172824.5 | NS | NS | NS | NS | NS | NS | NS | NS | NS | NS | 20.684 | 5.59E-07 | 0.000222 |
| **277** | C05658 | H | FALSE | 1H-Indol-3-ol | 133.11 | 0.847 | 0.2 | 638915 | 0 | | 92.8 | Spectral Match | | 94965.65 | 95370.68 | 139967.3 | 84221.52 | 0.41748 | 0.0013 | 0 | 0.535 | 0.8818 | 0.40793 | 0.000368 | 0 | 0.636719 | 0.0588 | 15.147 | 1.11E-05 | 0.000677 |
| **278** | C05660 | HB | FALSE | 5-Methoxyindoleacetate | 205.073 | 8.90 | 0.2 | 245032 | 0 | | 67.8 | Spectral | Other | 887.938 | 53020.2 | 29049.8 | 59012.8 | 21.843 | 0.00 | 0.0011 | 0.505 | 0.0000 | NS | NS | NS | NS | NS | 37.3 | 4.81E- | 0.000735 |

| | | | | | | | | | | | | | | | | | | | | | | | | | | | | | | |
|---|---|---|---|---|---|---|---|---|---|---|---|---|---|---|---|---|---|---|---|---|---|---|---|---|---|---|---|---|---|---|
| | | | | | 4 | 4 | | .7 | | | | Match | | 1 | 6 | | 8 | | 00 | 64 | | | | | | | | 34 | 10 | |
| 279 | C05774 | F | FALSE | Cobinamide | 495.7528 | 0.838 | 0.2 | 102479.8 | 0 | | | Spectral Match | | 28577.53 | 22791.7 | 18001.86 | 23989.16 | 0.31582 | 0.00070 | 0 | 0.56 | 0.7522 | NS | NS | NS | NS | NS | 4.208 | 0.021593 | 0 |
| 280 | C05828 | H | FALSE | Methylimidazoleacetic acid | 140.058 | 0.978 | 0.2 | 880521.4 | 0 | | 94.1 | Spectral Match | | 184440.6 | 176797.1 | 126481 | 189376.1 | 0.35601 | 0.00000 | 0 | 1 | 0.6808 | NS | NS | NS | NS | NS | 35.441 | 9.62E-10 | 0.000235 |
| 281 | C05842 | HFB | FALSE | N1-Methyl-2-pyridone-5-carboxamide | 152.0582 | 1.161 | 0.2 | 3630891 | 0 | | 67.3 | Spectral Match | | 618886.41 | 770248.2 | 1421364 | 607469.3 | 5.1886 | 0.02442 | 0 | 1 | 0.0001 | 0.25781 | 0.000887 | 0 | 0.71875 | 0.0358 | 16.276 | 5.84E-06 | 0.000677 |
| 282 | C05844 | H | FALSE | 5-L-Glutamyl-taurine | 254.057 | 2.903 | 0.2 | 1758913 | 0 | | | Spectral Match | Other | 2928.928 | 412992.4 | 619089.4 | 361468.1 | 55.544 | 0.0007 | 0.001036 | 1 | 0.0000 | NS | NS | NS | NS | NS | 4.1588 | 0.022496 | 0 |
| 283 | C05853 | HFB | FALSE | 2-Phenylethanol | 122.0728 | 1.063 | 0.2 | 530547.7 | 0 | | | Spectral Match | | 26017.23 | 75669.44 | 32851.53 | 86373.92 | NS | NS | NS | NS | NS | 1.6571 | 0.005631 | 0 | 0.949219 | 0.0000 | 22.946 | 1.84E-07 | 0.000929 |
| 284 | C05859 | H | FALSE | [Similar to: Methyl 3-[2-[(2,2-dichloroacetyl)amino]-4-(trifluoromethyl)phenoxy]thiophene-2-carboxylate; ΔMass: 168.8456 Da] | 258.1204 | 1.113 | 0.2 | 5268438 | 0 | | 60.6 | Spectral Match | | 35705.59 | 966764.4 | 216067.3 | 1154439 | 8.6842 | 0.0261 | 0 | 0.955 | 0.0008 | NS | NS | NS | NS | NS | 16.205 | 6.08E-06 | 0.00025 |
| 285 | C05913 | HFB | FALSE | (3Z)-Phytochromobilin | 584.2607 | 16.02 | 0.2 | 121016.6 | 0 | | | Spectral Mat | | 119.595 | 176669.79 | 665.8783 | 21920.76 | 53.674 | 0.000 | 0.001377 | 0.97 | 0.0000 | 21.841 | 3.18E-13 | 0.002725 | 1 | 0.000 | 6.893 | 0.002577 | 3.03E-05 |

| | | | | | | | | | | | | | | | | | | | | | | | | | | | | | | |
|---|---|---|---|---|---|---|---|---|---|---|---|---|---|---|---|---|---|---|---|---|---|---|---|---|---|---|---|---|---|---|
| | | | | | | | | | | | | ch | | | | | | | 0 | | | | | | | | 0 | 1 | | |
| 286 | C05922 | HB | FALSE | Formamidopyrimidine nucleoside triphosphate | 541.0026 | 10.041 | 0.2 | 486753.3 | 0 | | | Spectral Match | | 25339.08 | 22241.78 | 26189.77 | 21254.79 | 0.349996 | 0.0001 | 0 | 1 | 0.4132 | 0.53718 | 0.004905 | 0.000769 | 0.589844 | 0.4086 | 46.347 | 2.35E-11 | 0.002458 |
| 287 | C05924 | B | FALSE | Molybdopterin | 396.018 | 14.343 | 0.2 | 551351.3 | 0 | | | Spectral Match | | 108288.7 | 90034.72 | 51186.05 | 99746.89 | 0.30681 | 0.0000 | 0.001522 | 0.52 | 0.4991 | NS | NS | NS | NS | NS | 78.782 | 6.12E-15 | 0.004815 |
| 288 | C05932 | H | FALSE | N-Succinyl-L-glutamate 5-semialdehyde | 230.066 | 0.829 | 0.2 | 6198371 | 0 | | | Spectral Match | | 106831.1 | 965258.2 | 21912.15 | 1201095 | NS | NS | NS | NS | NS | 31.775 | 0.005336 | 0.000533 | 0.960938 | 0.0019 | 19.973 | 8.02E-07 | 0.000267 |
| 289 | C05944 | HFB | FALSE | Pantothenol | 205.1308 | 0.952 | 0.2 | 2169240 | 0 | | | Spectral Match | | 942930.9 | 77912.16 | 68997.89 | 80140.72 | 0.030079 | 0.0000 | 0 | 1 | 0.0000 | NS | NS | NS | NS | NS | 16.842 | 4.26E-06 | 0.000111 |
| 290 | C05947 | FB | FALSE | L-erythro-4-Hydroxyglutamate | 164.0556 | 0.856 | 0.2 | 972825.3 | 0 | | | Spectral Match | | 19260.8 | 135200.4 | 27332.56 | 162167.3 | NS | NS | NS | NS | NS | 3.5825 | 0.000065 | 0.000552 | 0.96875 | 0.0001 | 9.1981 | 0.000486 | 0 |
| 291 | C05966 | FB | FALSE | 15-Hydroperoxyeicosatetraenoate | 113.084 | 4.536 | 0.2 | 440074.1 | 0 | | | Spectral Match | | 30044.13 | 87202.36 | 119620 | 79097.96 | NS | NS | NS | NS | NS | 0.47037 | 0.000799 | 0.000154 | 0.683594 | 0.0774 | 6.6237 | 0.00316 | 0.000125 |
| 292 | C06002 | B | FALSE | (S)-Methylmalonatesemialdehyde | 102.032 | 0.963 | 0.2 | 1878785 | 0 | | 92.6 | Spectral Match | Superpathway of L-lysine degradation;Superpathway of L-threonine metabolism;Superpathway of L-arginine and L-ornithine degradation;Superpathway of branched chain amino acid | 91426.34 | 342295.4 | 252212.2 | 364816.2 | NS | NS | NS | NS | NS | NS | NS | NS | NS | NS | 11.951 | 7.79E-05 | 0.000462 |

| | | | | | | | | | | | | | | | | | | | | | | | | | | | | | | |
|---|---|---|---|---|---|---|---|---|---|---|---|---|---|---|---|---|---|---|---|---|---|---|---|---|---|---|---|---|---|---|
| | | | | | | | | | | | | | biosynthesis;Superpathway of L-isoleucine biosynthesis I;Superpathway of L-arginine, putrescine, and 4-aminobutanoate degradation;Other | | | | | | | | | | | | | | | | | |
| **293** | C06025 | H | FALSE | KDO2-lipid IVA | 923.4893 | 9.117 | 0.2 | 7980141 | 0 | | | Spectral Match | | 29906 | 61243.34 | 41703.51 | 66128.3 | NS | NS | NS | NS | NS | NS | NS | NS | NS | NS | 5.1589 | 0.009922 | 0 |
| **294** | C06044 | FB | FALSE | 2-(4-Hydroxyphenyl)ethanol (Tyrosol) | 138.0677 | 1.357 | 0.2 | 295432.4 | 0 | | 70.3 | Spectral Match | | 7561.565 | 33642.06 | 95244.66 | 18241.41 | NS | NS | NS | NS | NS | 0.13499 | 4.51E-08 | 8.00E-04 | 0.953125 | 0.0000 | 16.4667 | 5.25E-06 | 0.000728 |
| **295** | C06056 | H | FALSE | 4-Hydroxy-L-threonine | 136.0609 | 0.848 | 0.2 | 4072519 | 0 | | | Spectral Match | | 75782.61 | 982073.5 | 1041755 | 967153.1 | 4.9035 | 0.0000 | 0.000667 | 0.81 | 0.0000 | NS | NS | NS | NS | NS | 72.676 | 2.30E-14 | 2.00E-04 |
| **296** | C06123 | H | FALSE | 2-trans-Hexadecenal | 238.2289 | 15.193 | 0.2 | 239454.5 | 0 | | 63.6 | Spectral Match | | 1509.74 | 4857.84 | 10373.71 | 3478.871 | NS | NS | NS | NS | NS | 0.23326 | 8.85E-11 | 0.005325 | 0.984375 | 0.0000 | 5.1529 | 0.009997 | 0.000209 |
| **297** | C06165 | H | FALSE | Macarpine | 392.1149 | 19.77 | 0.2 | 9903320 | 0 | | | Spectral Match | | 1331343 | 1586334 | 1126148 | 1701381 | 0.4507 | 0.0011 | 0 | 0.855 | 0.7442 | NS | NS | NS | NS | NS | 18.307 | 1.92E-06 | 0.000683 |
| **298** | C06174 | H | FALSE | 7_8-Didehydro-4_5-epoxy-3-methoxy-17-methoxy-(5_)-morphinan- | 297.1354 | 7.149 | 0.2 | 520017.8 | 0 | | 82.1 | Spectral Match | | 69673.84 | 80168.47 | 81845.88 | 79749.12 | 0.45692 | 0.0000 | -0.00024 | 0.58 | 0.2810 | 0.6542 | 0.001216 | 0 | 0.5 | 0.8390 | 17.323 | 3.26E-06 | 0.001577 |

| | | | | | | | | | | | | | | | | | | | | | | | | | | | | | | |
|---|---|---|---|---|---|---|---|---|---|---|---|---|---|---|---|---|---|---|---|---|---|---|---|---|---|---|---|---|---|---|
| | | | | 6-one | | | | | | | | | | | | | | | | | | | | | | | | | | |
| 299 | C06178 | B | FALSE | 1-Methylpyrrolinium | 83.07384 | 0.712 | 0.2 | 290485.9 | 0 | | | Spectral Match | Superpathway of linamarin and lotaustralin biosynthesis | 28406.26 | 56135.65 | 56509.93 | 56042.09 | NS | NS | NS | NS | NS | NS | NS | NS | NS | NS | 4.4507 | 0.017656 | 0 |
| 300 | C06187 | B | FALSE | Arbutin 6-phosphate | 353.0626 | 8.624 | 0.2 | 4735374 | 0 | | | Spectral Match | | 14512.76 | 665269.1 | 48967.41 | 819344.5 | 16.011 | 0.0023 | 0 | 0.99 | 0.0001 | 10.682 | 0.000133 | 0.000977 | 0.945313 | 0.0000 | 5.6409 | 0.006762 | 0.000661 |
| 301 | C06188 | H | FALSE | Salicin 6-phosphate | 367.0786 | 8.797 | 0.2 | 1560126 | 0 | | | Spectral Match | | 1260.899 | 377261.4 | 539121.2 | 336796.5 | 125.73 | 0.0000 | 0.001983 | 1 | 0.0000 | NS | NS | NS | NS | NS | 47.297 | 1.75E-11 | 0.004747 |
| 302 | C06205 | H | FALSE | 1,2-Dihydronaphthalene-1,2-diol | 80.02645 | 0.984 | 0.2 | 1216461 | 0 | | | Spectral Match | | 23134.27 | 281589.8 | 300996.3 | 276738.2 | 4.6096 | 0.0000 | 0.000286 | 0.625 | 0.0000 | 0.56597 | 0.005867 | 0 | 0.511719 | 0.7796 | 36.774 | 5.89E-10 | 0.002325 |
| 303 | C06324 | H | FALSE | 1(2H)-Isoquinolinone | 145.0523 | 2.908 | 0.2 | 1709313 | 0 | | 69.1 | Spectral Match | | 17680.26 | 405180.2 | 593822.4 | 358019.7 | NS | NS | NS | NS | NS | 0.3117 | 0.027657 | 0 | 0.660156 | 0.1721 | 23.482 | 1.43E-07 | 0.000211 |
| 304 | C06326 | H | FALSE | (2S)-2-{[1-(R)-Carboxyethyl]amino}pentanoate | 189.0996 | 1.465 | 0.2 | 2960042 | 0 | | | Spectral Match | | 37041.34 | 557033.4 | 25629.73 | 689884.3 | NS | NS | NS | NS | NS | 17.456 | 8.35E-09 | 0.002354 | 1 | 0.0000 | 18.89 | 1.41E-06 | 0.000419 |
| 305 | C06368 | B | FALSE | 3-(Hydrohydroxyphosphoryl)pyruvate | 151.987 | 0.674 | 0.2 | 1006875 | 0 | | | Spectral Match | | 44919.57 | 139864.9 | 31096.58 | 167057 | NS | NS | NS | NS | NS | 4.6707 | 0.000568 | 0.000471 | 0.996094 | 0.0000 | 10.419 | 0.000212 | 0.000458 |
| 306 | C06397 | B | FALSE | ADP-D-glycero-beta-D-manno-heptose | 211.0813 | 2.338 | 0.2 | 493617.7 | 0 | | | Spectral Match | | 50805.91 | 119448.9 | 92576.59 | 126167 | NS | NS | NS | NS | NS | NS | NS | NS | NS | NS | 51.994 | 4.34E-12 | 0.000545 |
| 307 | C06507 | H | FALSE | Adenosyl cobyrinate hexaamide | 592.2682 | 7.918 | 0.2 | 2074649 | 0 | | | Spectral Mat | | 21398.19 | 25925.31 | 19196.86 | 27607.42 | 0.46552 | 0.000 | 0 | 0.68 | 0.2519 | NS | NS | NS | NS | NS | 4.896 | 0.012258 | 0 |

| | | | | | | | | | | | | | | | | | | | | | | | | | | | | | | |
|---|---|---|---|---|---|---|---|---|---|---|---|---|---|---|---|---|---|---|---|---|---|---|---|---|---|---|---|---|---|---|
| | | | | | | | | | | | | ch | | | | | | | 0 | | | | | | | | | 9 | | |
| 308 | C06508 | H | FALSE | Bis(4-ethylbenzylidene)sorbitol | 414.2023 | 12.101 | 0.2 | 3879799 | 0 | 80.1 | | Spectral Match | | 405890.8 | 834866.8 | 805438.9 | 842223.8 | NS | NS | NS | NS | NS | 0.65582 | 0.010461 | 0 | 0.511719 | 0.9092 | 16.859 | 4.22E-06 | 0.00085 |
| 309 | C06593 | H | FALSE | Caprolactam | 113.084 | 3.269 | 0.2 | 2080453 | 0 | 58 | | Spectral Match | | 60448.76 | 344975 | 304383 | 355123 | 2.1881 | 0.0000 | 0.000619 | 1 | 0.0000 | NS | NS | NS | NS | NS | 21.583 | 3.57E-07 | 0.000933 |
| 310 | C06658 | H | FALSE | Proclavaminicacid | 202.0947 | 0.991 | 0.2 | 768795.1 | 0 | | | Spectral Match | | 186802.7 | 130652.9 | 110989 | 135568.8 | 0.26894 | 0.0000 | 0.00242 | 1 | 0.0916 | NS | NS | NS | NS | NS | 10.427 | 0.000211 | 0.000254 |
| 311 | C06672 | FB | FALSE | Vanillate | 168.0394 | 1.3 | 0.2 | 264770 | 0 | | | Spectral Match | | 3234.623 | 8733.282 | 1383.936 | 10570.62 | NS | NS | NS | NS | NS | 5.4432 | 2.97E-11 | 0.004769 | 1 | 0.0000 | 9.9746 | 0.000285 | -0.00012 |
| 312 | C06725 | H | FALSE | 5_6-Dihydroxy-3-methyl-2-oxo-1_2-dihydroquinoline | 191.0577 | 1.55 | 0.2 | 387602.7 | 0 | | | Spectral Match | | 33206.34 | 88705.77 | 102486.5 | 85260.59 | NS | NS | NS | NS | NS | 0.50684 | 0.009817 | 0 | 0.554688 | 0.5000 | 39.806 | 2.01E-10 | 0.002781 |
| 313 | C06771 | H | FALSE | Triethanolamine | 149.1049 | 1.222 | 0.2 | 51511157 | 0 | 83.5 | | Spectral Match | | 1582604 | 2588910 | 1681589 | 2815741 | 0.60704 | 0.0049 | 0 | 0.74 | 0.0391 | NS | NS | NS | NS | NS | 5.3485 | 0.008526 | 0 |
| 314 | C06790 | H | FALSE | Trichloroethene | 130.9221 | 0.665 | 0.2 | 4306389 | 0 | | | Spectral Match | | 208057.3 | 471204.2 | 159135.4 | 549221.4 | NS | NS | NS | NS | NS | 2.6641 | 0.002731 | 0.000133 | 0.992188 | 0.0000 | 63.999 | 1.77E-13 | 0.000995 |
| 315 | C06848 | H | FALSE | Betamethasone | 392.2019 | 8.067 | 0.2 | 1065794 | 0 | | | Spectral Match | | 20820.65 | 31985.9 | 31948.63 | 31995.22 | 0.59158 | 0.0101 | 0 | 0.785 | 0.0962 | NS | NS | NS | NS | NS | 15.136 | 1.12E-05 | 0.000111 |
| 316 | C06850 | H | FALSE | Betanechol | 160.120 | 0.78 | 0.2 | 542480 | 0 | | 86.3 | Spectral | | 28599.5 | 95954.1 | 91417.2 | 97083.3 | NS | NS | NS | NS | NS | NS | NS | NS | NS | NS | 12.6 | 5.01E- | 0.00059 |

| | | | | | | | | | | | | | | | | | | | | | | | | | | | | | | |
|---|---|---|---|---|---|---|---|---|---|---|---|---|---|---|---|---|---|---|---|---|---|---|---|---|---|---|---|---|---|---|
| | | | | | 6 | 2 | | .7 | | | | Match | | 4 | 3 | 7 | 5 | | | | | | | | | | | 52 | 05 | |
| 317 | C06852 | H | FALSE | Bisoprolol | 325.224 | 9.318 | 0.2 | 8557069 | 0 | | | Spectral Match | | 58071.41 | 1653947 | 195822.1 | 2018479 | 10.977 | 0.0041 | -0.00011 | 0.74 | 0.0000 | 6.8604 | 0.006121 | 0 | 0.964844 | 0.0002 | 25.743 | 5.04E-08 | 0.000986 |
| 318 | C06943 | H | FALSE | Desipramine | 266.177 | 9.47 | 0.2 | 4557429 | 0 | | | Spectral Match | | 23533.94 | 644297.6 | 26826.43 | 798665.4 | NS | NS | NS | NS | NS | 17.796 | 0.000271 | 0.0001613 | 0.980469 | 0.0000 | 23.746 | 1.26E-07 | 0.0006069 |
| 319 | C06948 | H | FALSE | Diazepam | 284.0707 | 10.867 | 0.2 | 234811.8 | 0 | | | Spectral Match | | 713.9522 | 32387.69 | 971.9839 | 40241.62 | 16.386 | 0.0021 | 0 | 1 | 0.0001 | 27.411 | 2.66E-13 | 0.0003285 | 1 | 0.0000 | 20.22 | 7.07E-07 | 0.0000353 |
| 320 | C06971 | H | FALSE | Doxepin | 279.1613 | 8.566 | 0.2 | 76200.76 | 0 | | | Spectral Match | | 9939.779 | 9869.41 | 7435.785 | 10477.82 | 0.38593 | 0.0000 | 0.000375 | 1 | 0.8201 | NS | NS | NS | NS | NS | 25.842 | 4.82E-08 | 0.0001941 |
| 321 | C06980 | H | FALSE | Esmolol | 295.1772 | 8.134 | 0.2 | 506522.1 | 0 | | 74.6 | Spectral Match | | 249.0446 | 96164.13 | 5947.679 | 118718.2 | 142.07 | 0.0000 | 0.000711 | 0.525 | 0.0000 | 13.466 | 1.71E-07 | 0.0002757 | 1 | 0.0000 | 42.794 | 7.35E-11 | 0.0001617 |
| 322 | C07049 | H | FALSE | Imipramine | 280.1928 | 9.528 | 0.2 | 4.75E+08 | 0 | 90.3 | | Spectral Match | | 14668768 | 33162891 | 42293464 | 30880247 | NS | NS | NS | NS | NS | 0.49317 | 3.34E-08 | 0 | 0.992188 | 0.0000 | 3.9446 | 0.026921 | 0 |
| 323 | C07063 | H | FALSE | Labetalol | 328.1783 | 7.924 | 0.2 | 2583384 | 0 | | | Spectral Match | | 13264.38 | 76258.87 | 46259.49 | 83758.72 | 2.1398 | 0.0000 | 0 | 1 | 0.0000 | NS | NS | NS | NS | NS | 6.6928 | 0.002998 | 0 |
| 324 | C07073 | H | FALSE | Lidocaine | 234.1725 | 6.575 | 0.2 | 2798559 | 0 | 73 | | Spectral Match | | 63814.47 | 611704.7 | 643081.7 | 603860.4 | 3.8595 | 0.0001 | 0.001 | 1 | 0.0000 | 0.59304 | 0.047301 | 0.0000125 | 0.507813 | 0.8555 | 14.908 | 1.28E-05 | 0.0000525 |
| 325 | C07128 | H | FALSE | Meperidine | 247.156 | 6.25 | 0.2 | 884140 | 0 | | | Spectral | | 7918.88 | 160557. | 14165.0 | 197156 | NS | NS | NS | NS | NS | 7.4781 | 0.0044 | 0.0000276 | 0.9765 | 0.00 | 15.9 | 6.90E- | 0.000065 |

| | | | | | | | | | | | | | | | | | | | | | | | | | | | | | | |
|---|---|---|---|---|---|---|---|---|---|---|---|---|---|---|---|---|---|---|---|---|---|---|---|---|---|---|---|---|---|---|
| | | | | | 4 | 7 | | .1 | | | | Match | | 8 | 8 | 2 | | | | | | | | 29 | | 63 | 06 | 82 | 06 | |
| 326 | C07129 | H | FALSE | Terbutaline | 225.1357 | 7.624 | 0.2 | 591858.6 | 0 | | | Spectral Match | | 149333 | 112668.1 | 63783.86 | 124889.1 | NS | NS | NS | NS | NS | NS | NS | NS | NS | NS | 65.494 | 1.23E-13 | 0.004597 |
| 327 | C07151 | H | FALSE | Metformin | 129.1019 | 0.913 | 0.2 | 44328513 | 0 | | 85 | Spectral Match | | 774909.6 | 638796.6 | 419148.6 | 7880170 | NS | NS | NS | NS | NS | 10.928 | 7.45E-05 | 0.0000821 | 0.980469 | 0.0000 | 30.203 | 7.44E-09 | 0.0000776 |
| 328 | C07159 | FB | FALSE | 2-Oxo-delta3-4_5_5-trimethylcyclopentenyl acetate | 182.0939 | 7.281 | 0.2 | 533228 | 0 | | | Spectral Match | | 2031.938 | 7959.827 | 6627.845 | 8292.823 | 1.5636 | 0.0302 | 0.000105 | 1 | 0.0000 | NS | NS | NS | NS | NS | 8.1057 | 0.001055 | 4.00E-04 |
| 329 | C07189 | H | FALSE | 3'-hydroxyacetophenone; 3-ACETYLPHENOL | 136.0652 | 1.068 | 0.2 | 249180.8 | 0 | | | Spectral Match | Anaerobic aromatic compound degradation (Thauera aromatica);Other | 8963.862 | 43043.08 | 346510.8 | 451410.8 | 1.7466 | 0.0032 | 0 | 1 | 0.0000 | NS | NS | NS | NS | NS | 29.675 | 9.25E-09 | 0.001189 |
| 330 | C07196 | H | FALSE | Methylphenidate | 233.1409 | 7.436 | 0.2 | 141764.9 | 0 | | | Spectral Match | | 5957.974 | 9528.921 | 4991.563 | 10663.26 | 0.60224 | 0.0102 | -0.00013 | 1 | 0.0722 | NS | NS | NS | NS | NS | 12.836 | 4.46E-05 | 0.000125 |
| 331 | C07203 | H | FALSE | Metronidazole | 171.0639 | 2.626 | 0.2 | 1493324 | 0 | | | Spectral Match | | 693.704 | 223294.8 | 5548.988 | 27731.2 | 100.85 | 0.0005 | 0.000571 | 0.785 | 0.0000 | 29.188 | 0.002972 | 0.0000402 | 0.964844 | 0.0003 | 11.812 | 8.51E-05 | 0.00025 |
| 332 | C07204 | H | FALSE | Vincristine | 824.4036 | 0.937 | 0.2 | 499096.2 | 0 | | | Spectral Match | | 1647.986 | 18783.56 | 1962.385 | 22988.85 | 4.6628 | 0.0082 | -0.00014 | 1 | 0.0000 | 8.7191 | 1.20E-13 | 0.002919 | 1 | 0.0000 | 22.996 | 1.80E-07 | 0.000567 |
| 333 | C07207 | H | FALSE | Zalcitabine | 211.0948 | 2.299 | 0.2 | 897998.3 | 0 | | | Spectral Match | | 713004.9 | 123876.4 | 1599.027 | 154445.7 | NS | NS | NS | NS | NS | 53.964 | 8.06E-18 | 0.004493 | 1 | 0.0000 | 6.9266 | 0.002513 | 0 |

| | | | | | | | | | | | | | | | | | | | | | | | | | | | | | | |
|---|---|---|---|---|---|---|---|---|---|---|---|---|---|---|---|---|---|---|---|---|---|---|---|---|---|---|---|---|---|---|
| 334 | C07289 | H | FALSE | [FA(18:1)]9Z-Octadecen-12-ynoicacid | 278.2237 | 13.624 | 0.2 | 124910.6 | 0 | | | Spectral Match | Other | 5898.38 | 9433.77 | 8473.288 | 9673.891 | 0.62673 | 0.01449 | 0 | 1 | 0.0521 | NS | NS | NS | NS | NS | 7.11183 | 0.002177 | 0 |
| 335 | C07354 | B | FALSE | (7S_8S)-DiHODE | 312.229 | 11.126 | 0.2 | 256459.9 | 0 | | | Spectral Match | Other | 8112.801 | 10389.82 | 10705.47 | 10310.9 | 0.5103 | 0.00000 | 0 | 0.775 | 0.0000 | 0.64105 | 8.41E-06 | 0 | 0.605469 | 0.3822 | 36.344 | 6.89E-10 | 0.000898 |
| 336 | C07410 | H | FALSE | Pyridostigmine | 180.0893 | 2.101 | 0.2 | 461854.8 | 0 | | | Spectral Match | | 50120.08 | 85942.41 | 30088.48 | 99905.89 | 0.58421 | 0.04463 | 0 | 0.635 | 0.3961 | 1.9384 | 0.036979 | 0 | 0.851563 | 0.0026 | 5.6966 | 0.006472 | 0.000111 |
| 337 | C07475 | H | FALSE | Ethoxyquin | 217.146 | 6.662 | 0.2 | 670952.5 | 0 | | | Spectral Match | | 24721.33 | 97426.28 | 15847.69 | 117820.9 | NS | NS | NS | NS | NS | 4.3326 | 1.52E-05 | 0.001797 | 1 | 0.0000 | 14.435 | 1.69E-05 | 0.002455 |
| 338 | C07481 | H | FALSE | Cafeine | 194.0798 | 5.138 | 0.2 | 8725927 | 0 | | 87.5 | Spectral Match | Other | 22728.97 | 59481.85 | 20765.19 | 69161.02 | NS | NS | NS | NS | NS | 2.1387 | 6.32E-05 | 0.001699 | 1 | 0.0000 | 92.648 | 3.98E-16 | 0.000267 |
| 339 | C07490 | H | FALSE | [Similar to: Triethanolamine; ΔMass: 0.1724 Da] | 148.9328 | 0.674 | 0.046 | 2030649 | 0 | | 63.4 | Spectral Match | | 52352.54 | 139999.8 | 61634.64 | 159591.1 | NS | NS | NS | NS | NS | NS | NS | NS | NS | NS | 29.085 | 1.18E-08 | 0.000693 |
| 340 | C07525 | H | FALSE | Leu-Pro(Leucyl-Proline) | 228.1467 | 0.994 | 0.062 | 2064319 | 0 | | | Spectral Match | | 109844.8 | 378483.6 | 231548.3 | 415217.4 | 12.063 | 0.0003 | 0 | 0.975 | 0.0000 | NS | NS | NS | NS | NS | 34.839 | 1.20E-09 | 0.000369 |
| 341 | C07527 | H | FALSE | Benzocaine | 165.0784 | 7.028 | 0.2 | 1271644 | 0 | | | Spectral Match | Superpathway of chorismate metabolism;Superpathway of aromatic amino acid biosynthesis;Anaerobic aromatic compound | 7704.115 | 176640.4 | 7237.101 | 218991.2 | 6.9569 | 0.0442 | 0 | 1 | 0.0018 | 16.815 | 2.20E-08 | 0.005884 | 1 | 0.0000 | 27.595 | 2.23E-08 | 0.002975 |

| | | | | | | | | | | | | | | | | | | | | | | | | | | | | | | |
|---|---|---|---|---|---|---|---|---|---|---|---|---|---|---|---|---|---|---|---|---|---|---|---|---|---|---|---|---|---|---|
| | | | | | | | | | | | | | degradation (Thauera aromatica);Superpathway of indole-3-acetate conjugate biosynthesis;Superpathway of jasmonoyl-amino acid conjugates biosynthesis;Other | | | | | | | | | | | | | | | | | |
| 342 | C07537 | H | FALSE | Ethylmorphine | 313.1665 | 8.544 | 0.2 | 1000043 | 0 | | 91.1 | Spectral Match | | 19571.8 | 158336.9 | 32171.21 | 189878.3 | NS | NS | NS | NS | NS | 3.257 | 0.000633 | 0.000133 | 0.972656 | 0.0000 | 33.944 | 1.69E-09 | 0.003275 |
| 343 | C07545 | H | FALSE | α-Ergocryptine | 575.3124 | 8.586 | 0.2 | 2810876 | 0 | 50.5 | | Spectral Match | | 37949.43 | 19829.72 | 15154.18 | 20998.6 | 0.20712 | 0.0000 | 0.001817 | 0.87 | 0.0000 | NS | NS | NS | NS | NS | 8.5436 | 0.000771 | -7.35E-06 |
| 344 | C07585 | H | FALSE | N-Acetylisoniazid | 180.0766 | 10.138 | 0.2 | 226978.8 | 0 | | | Spectral Match | | 15180.53 | 37388.18 | 26260.22 | 40170.17 | NS | NS | NS | NS | NS | NS | NS | NS | NS | NS | 8.2761 | 0.000933 | -0.00022 |
| 345 | C07591 | H | FALSE | Phenacetin | 179.0941 | 0.389 | 0.2 | 1145799 | 0 | | | Spectral Match | | 91915.23 | 172570.8 | 107914.8 | 188734.8 | NS | NS | NS | NS | NS | NS | NS | NS | NS | NS | 5.5787 | 0.007102 | 0 |
| 346 | C07592 | H | FALSE | Colchicine | 399.1665 | 8.93 | 0.062 | 7434749 | 0 | | | Spectral Match | | 21430.52 | 1051967 | 59322.36 | 1300128 | 15.952 | 0.0071 | 0 | 1 | 0.0001 | 12.707 | 0.000172 | 0 | 0.988281 | 0.0000 | 17.681 | 2.69E-06 | 0 |
| 347 | C07644 | H | FALSE | 4-Ketocyclophosphamide | 138.0093 | 0.666 | 0.2 | 268434.3 | 0 | | | Spectral Match | | 28003.98 | 55676.4 | 28426.62 | 62488.85 | NS | NS | NS | NS | NS | NS | NS | NS | NS | NS | 5.4877 | 0.007632 | 0 |
| 3 | C07 | H | FA | Pirbuterol | 24 | 1. | 0. | 21 | 0 | | | Spe | | 74 | 44 | 63 | 39 | NS | N | NS | N | N | 0. | 0. | 0 | 0. | 0. | 1 | 4. | 0.00 |

| | | | | | | | | | | | | | | | | | | | | | | | | | | | | | | |
|---|---|---|---|---|---|---|---|---|---|---|---|---|---|---|---|---|---|---|---|---|---|---|---|---|---|---|---|---|---|---|
| 48 | 807 | | LSE | | 0.14464 | 112 | 2 | 14346 | | | | ctral Match | | 729.8 | 0804.7 | 6330 | 1923.4 | | S | | S | S | 38287 | 001726 | | 625 | 1763 | 2.918 | 24E-05 | 0886 |
| 349 | C07935 | H | FALSE | Orphenadrine | 269.1768 | 8.349 | 0.2 | 514580.6 | 0 | | | Spectral Match | | 366055.41 | 998023.1 | 481519.7 | 112714.9 | NS | NS | NS | NS | NS | NS | NS | NS | NS | NS | 18.229 | 2.00E-06 | 0.000503 |
| 350 | C08156 | H | FALSE | Testosterone Cypionate | 412.2959 | 14.749 | 0.2 | 1562025 | 0 | | | Spectral Match | | 9487.744 | 408666.9 | 176761.2 | 466643.3 | 15.265 | 0.0001 | 0.000353 | 0.97 | 0.0000 | NS | NS | NS | NS | NS | 4.9501 | 0.01174 | 0 |
| 351 | C08265 | HB | FALSE | Alliin | 177.0454 | 0.811 | 0.2 | 530531 | 0 | | | Spectral Match | Other | 133518.2 | 113586.5 | 152968.1 | 103741.1 | 0.34843 | 0.0007 | 0 | 0.98 | 0.4380 | 0.44457 | 0.001288 | 0 | 0.703125 | 0.1067 | 5.7831 | 0.006047 | 0.00025 |
| 352 | C08278 | H | FALSE | Suberic acid | 174.0872 | 7.373 | 0.109 | 575739.1 | 0 | | | Spectral Match | | 110460.7 | 116647.3 | 169002.9 | 103558.3 | 0.42156 | 0.0004 | 0 | 0.66 | 0.9816 | 0.41185 | 3.70E-05 | 0 | 0.683594 | 0.0499 | 37.934 | 3.88E-10 | 0.000651 |
| 353 | C08287 | H | FALSE | Hypoglycin A | 141.0786 | 0.993 | 0.2 | 1649256 | 0 | | 94.1 | Spectral Match | | 225339.8 | 352761.4 | 187820.8 | 393996.6 | 0.54827 | 0.0013 | 0 | 0.655 | 0.2373 | NS | NS | NS | NS | NS | 4.178 | 0.022139 | 0.000118 |
| 354 | C08288 | HB | FALSE | L-Indospicine | 173.1159 | 1.101 | 0.2 | 755970.9 | 0 | | | Spectral Match | | 127775.95 | 151244.3 | 146869.4 | 152338 | 4.5002 | 0.0019 | 0 | 0.675 | 0.0000 | NS | NS | NS | NS | NS | 90.348 | 6.11E-16 | 0.000901 |
| 355 | C08290 | H | FALSE | L-Lathyrine | 182.0801 | 0.998 | 0.2 | 53380.01 | 0 | | | Spectral Match | | 1156.88 | 5951.881 | 1056.312 | 7175.773 | NS | NS | NS | NS | NS | 4.5219 | 2.29E-13 | 0.004024 | 1 | 0.0000 | 20.929 | 4.94E-07 | 0.001664 |
| 356 | C08301 | H | FALSE | Cathinone | 149.0835 | 1.061 | 0.2 | 275757.7 | 0 | | | Spectral Match | | 14133.84 | 540966.1 | 418899.28 | 571445.94 | NS | NS | NS | NS | NS | NS | NS | NS | NS | NS | 31.878 | 3.78E-09 | 0.000948 |
| 3 | C08 | H | FA | 5-Methoxy- | 21 | 7. | 0. | 65 | 0 | | 67 | Spe | | 15 | 10 | 67 | 11 | 23 | 0. | 0. | 0. | 0. | NS | NS | NS | NS | N | 9. | 0. | - |

| | | | | | | | | | | | | | | | | | | | | | | | | | | | | | | |
|---|---|---|---|---|---|---|---|---|---|---|---|---|---|---|---|---|---|---|---|---|---|---|---|---|---|---|---|---|---|---|
| 57 | 309 | | LSE | N_N-dimethyltryptamine | 8.1413 | 418 | 2 | 6514.8 | | | .8 | ctral Match | | 83.368 | 4400.9 | 044.4 | 3740 | .373 | 0000 | 000756 | 99 | 0000 | | | | | S | 0909 | 000524 | 2.38E-05 |
| 358 | C08322 | B | FALSE | (9Z)-Tetradecenoicacid | 226.1923 | 12.204 | 0.2 | 1.18E+08 | 0 | | 73.7 | Spectral Match | | 1344186 | 2675131 | 3077528 | 2574532 | NS | NS | NS | NS | NS | 0.5619 | 3.27E-07 | 0.000767 | 0.824219 | 0.0009 | 3.9355 | 0.027129 | 0.000143 |
| 359 | C08334 | H | FALSE | Lotaustralin | 261.1201 | 0.977 | 0.2 | 604706.5 | 0 | | 74.8 | Spectral Match | Superpathway of linamarin and lotaustralin biosynthesis | 47696.83 | 157750.9 | 117717.4 | 167759.3 | NS | NS | NS | NS | NS | NS | NS | NS | NS | NS | 12.8331 | 4.48E-05 | 0.000421 |
| 360 | C08362 | H | FALSE | Palmitoleic acid | 254.2234 | 13.429 | 0.2 | 61900943 | 0 | | | Spectral Match | Other | 1058278 | 8867768.6 | 1344228 | 7724036 | 0.34864 | 0.00000 | 0.000477 | 1 | 0.0748 | 0.38663 | 4.68E-12 | 0.002663 | 1 | 0.00000 | 17.502 | 2.96E-06 | 0.00086 |
| 361 | C08439 | H | FALSE | Phalloidin | 788.3178 | 10.674 | 0.2 | 516742 | 0 | | | Spectral Match | | 17038.01 | 53543.11 | 54612.03 | 53275.88 | NS | NS | NS | NS | NS | 0.62907 | 0.00024 | 0.000669 | 0.507813 | 0.85155 | 15.128 | 1.13E-05 | 0 |
| 362 | C08497 | H | FALSE | [FA(6:1)]2-hexenal | 98.07311 | 1.007 | 0.2 | 937576.1 | 0 | | 83.8 | Spectral Match | Other | 85418.52 | 107485.2 | 97206.77 | 110054.8 | 0.484 | 0.0002 | 0 | 0.81 | 0.1971 | NS | NS | NS | NS | NS | 6.5274 | 0.003401 | -0.00013 |
| 363 | C08538 | H | FALSE | Betalamic acid | 211.0474 | 0.986 | 0.2 | 383463.6 | 0 | | | Spectral Match | Other | 1409.656 | 51915.7 | 1371.766 | 64551.68 | 12.512 | 0.0047 | 0 | 0.67 | 0.0002 | 29.279 | 1.39E-17 | 0.004489 | 1 | 0.0000 | 6.9948 | 0.002388 | 0.000389 |
| 364 | C08608 | H | FALSE | Norbixin | 380.197 | 13.175 | 0.2 | 3772911 | 0 | | 64.2 | Spectral Match | | 39505.47 | 70852.83 | 73842.47 | 70105.43 | NS | NS | NS | NS | NS | 0.62852 | 1.91E-06 | 0 | 0.640625 | 0.2258 | 4.0685 | 0.02426 | 0.000333 |
| 365 | C08869 | H | FALSE | Strophanthidin | 404.2191 | 8.9 | 0.2 | 2581029 | 0 | | 62 | Spectral Match | | 28757.81 | 26028.58 | 31346.5 | 24699.1 | 0.37512 | 0.00000 | 0 | 0.87 | 0.2299 | 0.53588 | 1.39E-05 | 0 | 0.851563 | 0.00002 | 31.8 | 3.90E-09 | 0.001049 |
| 3 | C09 | H | FA | 7- | 16 | 1 | 0. | 59 | 0 | 60 | | Spe | Other | 77 | 68 | 97 | 61 | 0. | 0. | 0 | 0. | 0. | 0. | 8. | 0.00 | 0. | 0. | 4 | 1. | 0.00 |

| | | | | | | | | | | | | | | | | | | | | | | | | | | | | | | |
|---|---|---|---|---|---|---|---|---|---|---|---|---|---|---|---|---|---|---|---|---|---|---|---|---|---|---|---|---|---|---|
| 66 | 315 | | LSE | Hydroxycoumarine | 2.0312 | 0.935 | 2 | 04873 | | .3 | | ctral Match | | 243.75 | 376.19 | 281.69 | 149.81 | 36951 | 0000 | | 685 | 1491 | 42079 | 45E-10 | 0286 | 996094 | 0000 | 0.594 | 54E-10 | 1597 |
| 367 | C09382 | H | FALSE | Costunolide | 232.1455 | 9.182 | 0.2 | 426785.9 | 0 | | | Spectral Match | | 3925.91 | 61710.75 | 2471.894 | 76520.46 | NS | NS | NS | NS | NS | 18.36 | 1.72E-14 | 0.003836 | 1 | 0.0000 | 7.0214 | 0.002341 | 0 |
| 368 | C09558 | H | FALSE | Laudanosine | 357.1927 | 7.873 | 0.2 | 1445942 | 0 | 69.8 | | Spectral Match | Superpathway of jasmonoyl-amino acid conjugates biosynthesis | 15399.57 | 74706.29 | 35131.96 | 84599.87 | NS | NS | NS | NS | NS | NS | NS | NS | NS | NS | 8.6981 | 0.000691 | 0.000235 |
| 369 | C09861 | H | FALSE | Geranylacetate | 196.1457 | 12.107 | 0.2 | 2620327 | 0 | | | Spectral Match | | 13132.16 | 7888.886 | 2240.883 | 9300.886 | 0.22851 | 0.0000 | 6.00E-04 | 0.8 | 0.0488 | 2.8308 | 3.19E-05 | 0.000429 | 1 | 0.0000 | 10.501 | 0.0002 | 0 |
| 370 | C10164 | FB | FALSE | Picolinic acid | 123.0338 | 0.432 | 0.2 | 1408044 | 0 | | 69.7 | Spectral Match | | 710488.8 | 622746.1 | 689053.9 | 606169.1 | NS | NS | NS | NS | NS | NS | NS | NS | NS | NS | 60.06 | 4.80E-13 | 0.002507 |
| 371 | C10172 | H | FALSE | DL-Stachydrine | 143.0941 | 20.579 | 0.2 | 1597620 | 0 | 83.7 | | Spectral Match | | 127742.8 | 148840.1 | 147637.3 | 149140.8 | 0.4561 | 0.0000 | 0 | 0.995 | 0.1431 | NS | NS | NS | NS | NS | 40.26 | 1.72E-10 | 0.001014 |
| 372 | C10503 | H | FALSE | (-)-Medicarpin | 270.0885 | 5.059 | 0.2 | 637975.1 | 0 | | 71.9 | Spectral Match | Other | 119327.4 | 154621.4 | 137658.6 | 158862.1 | 0.5016 | 0.0027 | 0.000133 | 0.81 | 0.3782 | NS | NS | NS | NS | NS | 33.106 | 2.34E-09 | 0.002501 |
| 373 | C10858 | H | FALSE | Ecgonine | 185.1046 | 1.883 | 0.2 | 289506.6 | 0 | | | Spectral Match | | 7684.211 | 5123.272 | 2259.811 | 5839.137 | 0.24815 | 0.0000 | 0.001948 | 0.63 | 0.0171 | 1.6928 | 4.31E-07 | 0.002742 | 1 | 0.0000 | 23.331 | 1.53E-07 | 0.005543 |
| 374 | C10929 | B | FALSE | Benalaxyl | 325.1665 | 7.85 | 0.2 | 1971309 | 0 | | 60.7 | Spectral Match | | 144728.1 | 184663.7 | 144531 | 194696.9 | 0.50105 | 0.0000 | 0.000419 | 0.91 | 0.0050 | NS | NS | NS | NS | NS | 5.9718 | 0.005218 | 0 |
| 3 | C10 | H | FA | Daminozide | 16 | 1. | 0. | 27 | 0 | | 79 | Spe | Other | 14 | 65 | 40 | 71 | NS | N | NS | N | N | NS | NS | NS | NS | N | 2 | 3. | 0.00 |

| | | | | | | | | | | | | | | | | | | | | | | | | | | | | | | |
|---|---|---|---|---|---|---|---|---|---|---|---|---|---|---|---|---|---|---|---|---|---|---|---|---|---|---|---|---|---|---|
| 75 | 996 | | LSE | | 0.0843 | 066 | 2 | 95731 | | | .8 | ctral Match | | 2257.3 | 4336.5 | 9174.9 | 5626.9 | | S | | S | S | | | | | S | 6.382 | 79E-08 | 1421 |
| 376 | C11005 | H | FALSE | Isoproturon | 206.1412 | 4.667 | 0.2 | 345146.5 | 0 | | 75.1 | Spectral Match | | 2881.432 | 50678.54 | 15474.49 | 59479.55 | 6.1156 | 0.0000 | 0.001167 | 0.86 | 0.0000 | 2.3435 | 0.022314 | 0 | 0.910156 | 0.0005 | 47.754 | 1.53E-11 | 0.001527 |
| 377 | C11071 | H | FALSE | Aminocarb | 208.1206 | 3.2 | 0.2 | 430642.6 | 0 | | 78.1 | Spectral Match | | 18388.1 | 102369 | 111592.6 | 100063.2 | NS | NS | NS | NS | NS | NS | NS | NS | NS | NS | 15.247 | 1.05E-05 | 0.000419 |
| 378 | C11073 | H | FALSE | Bufencarb | 221.1408 | 2.133 | 0.2 | 1203410 | 0 | | 61.4 | Spectral Match | | 15161.24 | 189381.8 | 168930.6 | 194494.7 | 4.7043 | 0.0000 | 0.001846 | 1 | 0.0000 | NS | NS | NS | NS | NS | 8.7426 | 0.000669 | 4.00E-04 |
| 379 | C11124 | H | FALSE | Spiroxamine | 297.2655 | 11.476 | 0.2 | 464548.5 | 0 | | 96 | Spectral Match | | 4083.605 | 9048.772 | 6549.786 | 9673.518 | NS | NS | NS | NS | NS | NS | NS | NS | NS | NS | 18.99 | 1.34E-06 | 0.003082 |
| 380 | C11134 | H | FALSE | Testosterone glucuronide | 464.2403 | 17.568 | 0.2 | 6560725 | 0 | | | Spectral Match | | 179690.4 | 1472671 | 1101814 | 1565385 | 3.211 | 0.0059 | 0 | 1 | 0.0000 | NS | NS | NS | NS | NS | 27.278 | 2.56E-08 | 0.002268 |
| 381 | C11156 | H | FALSE | Triadimefon | 293.0923 | 1.918 | 0.2 | 79791332 | 0 | | | Spectral Match | | 33495639 | 67904.66 | 46664.93 | 73214.59 | 0.000784 | 0.0000 | 0.000444 | 1 | 0.0000 | NS | NS | NS | NS | NS | 41.029 | 1.32E-10 | 0.000375 |
| 382 | C11173 | H | FALSE | SN-38 | 197.0767 | 0.88 | 0.2 | 349645.4 | 0 | | | Spectral Match | | 27770.3 | 74147.43 | 84506.52 | 71557.65 | NS | NS | NS | NS | NS | 0.51751 | 0.035921 | -0.00031 | 0.5 | 0.7193 | 12.031 | 7.40E-05 | 0.000721 |
| 383 | C11233 | B | FALSE | Dimethyl Phthalate | 194.0574 | 10.798 | 0.2 | 7.43E+08 | 0 | | 88.5 | Spectral Match | Other | 7776516 | 8758627 | 7277785 | 9128837 | 0.44578 | 0.0000 | 0.001239 | 0.89 | 0.2341 | NS | NS | NS | NS | NS | 6.6122 | 0.003188 | 0.000361 |
| 3 | C11 | H | FA | Bacteriochl | 30 | 1. | 0. | 59 | 0 | | | Spe | | 13 | 85 | 19 | 10 | NS | N | NS | N | N | 4. | 0. | 0 | 0. | 0. | 7 | 5. | 0.00 |

| | | | | | | | | | | | | | | | | | | | | | | | | | | | | | | |
|---|---|---|---|---|---|---|---|---|---|---|---|---|---|---|---|---|---|---|---|---|---|---|---|---|---|---|---|---|---|---|
| 84 | 242 | | LSE | orophyll a | 3.8521 | 712 | 2 | 9895.3 | | | | ctral Match | | 949.46 | 926.51 | 743.42 | 2472.3 | | S | | S | S | 9946 | 028193 | | 878906 | 0003 | 9.403 | 37E-15 | 5098 |
| 385 | C11284 | H | FALSE | 3-Indolebutyricacid | 203.0943 | 3.132 | 0.2 | 124906 | 0 | | | Spectral Match | | 16844.71 | 23387.1 | 17203.75 | 24932.94 | 0.53009 | 0.0074 | 0 | 0.685 | 0.1832 | NS | NS | NS | NS | NS | 95.157 | 2.52E-16 | 0.000672 |
| 386 | C11285 | FB | FALSE | Tridemorph | 297.30211 | 16.571 | 0.2 | 169085.6 | 0 | | | Spectral Match | | 2750.708 | 13573.55 | 14797.65 | 13267.52 | 1.9846 | 0.0005 | 0.000797 | 0.665 | 0.0000 | 0.60583 | 3.57E-05 | 0.000353 | 0.625 | 0.2433 | 49.919 | 7.96E-12 | 0.001443 |
| 387 | C11355 | B | FALSE | 4-Amino-4-deoxychorismate | 224.057 | 0.685 | 0.2 | 1216643 | 0 | | | Spectral Match | | 67488.74 | 172757.7 | 394082 | 1174426.6 | NS | NS | NS | NS | NS | 0.19109 | 4.30E-06 | 0.000025 | 0.824219 | 0.0038 | 76.827 | 9.27E-15 | 0.004943 |
| 388 | C11434 | H | FALSE | 2-C-Methyl-D-erythritol 4-phosphate | 109.0276 | 0.374 | 0.2 | 997253.5 | 0 | | | Spectral Match | | 297023.2 | 379715.4 | 363804.1 | 383693.2 | NS | NS | NS | NS | NS | NS | NS | NS | NS | NS | 38.814 | 2.84E-10 | 0.005492 |
| 389 | C11457 | FB | FALSE | 3-(3-Hydroxyphenyl)propanoic acid | 166.061 | 6.718 | 0.2 | 186846.8 | 0 | | | Spectral Match | | 9783.883 | 30286.25 | 33158.83 | 29568.11 | NS | NS | NS | NS | NS | 0.58567 | 0.001017 | 0 | 0.578125 | 0.4901 | 19.395 | 1.08E-06 | 0.001674 |
| 390 | C11480 | H | FALSE | 2-Methyl-6-oxohepta-2_4-dienal | 140.0833 | 1.063 | 0.2 | 4764614 | 0 | | 88.2 | Spectral Match | | 144870 | 1084198 | 263344.8 | 1289411 | NS | NS | NS | NS | NS | 3.144 | 0.047071 | 0.000308 | 0.851563 | 0.0029 | 13.02 | 3.98E-05 | 0.000786 |
| 391 | C11636 | H | FALSE | 7-Deoxyloganate | 360.1417 | 7.573 | 0.2 | 2165749 | 0 | | | Spectral Match | | 6436.621 | 311977.9 | 89051.21 | 367709.6 | 16.637 | 0.0000 | 0.002544 | 1 | 0.0000 | 2.5711 | 0.005201 | 0.000603 | 0.929688 | 0.0000 | 18.742 | 1.52E-06 | 0.000653 |
| 392 | C11770 | H | FALSE | Albuterol | 239.1514 | 8.342 | 0.2 | 818596.6 | 0 | | | Spectral Match | | 432.898 | 126910.3 | 2326.509 | 158056.2 | 109.04 | 0.0000 | 0.000502 | 1 | 0.0000 | 46.626 | 2.25E-10 | 0.003782 | 1 | 0.0000 | 5.8645 | 0.005673 | 0 |
| 3 | C11 | H | FA | N-Butyryl-L- | 17 | 5. | 0. | 58 | 0 | | | Spe | | 97 | 11 | 86 | 12 | 40 | 0. | 0. | 1 | 0. | NS | NS | NS | NS | N | 8. | 0. | 0 |

| | | | | | | | | | | | | | | | | | | | | | | | | | | | | | | |
|---|---|---|---|---|---|---|---|---|---|---|---|---|---|---|---|---|---|---|---|---|---|---|---|---|---|---|---|---|---|---|
| 93 | 837 | | LSE | homoserine lactone | 1.0891 | 735 | 2 | 32062 | | | | ctral Match | | 8.1054 | 96690 | 7581.5 | 78967 | 9.53 | 0000 | 000671 | | 0000 | | | | | S | 7426 | 000669 | |
| 394 | C11839 | B | FALSE | [FAoxo(6:0)]N-(3-oxo-hexanoyl)-homoserine lactone | 213.0993 | 8.865 | 0.2 | 266527.7 | 0 | | | Spectral Match | | 25195.42 | 36616.63 | 31451.5 | 37907.91 | 0.56836 | 0.0000 | 0 | 1 | 0.0005 | NS | NS | NS | NS | NS | 9.2468 | 0.00047 | 0.000511 |
| 395 | C11907 | FB | FALSE | 2-Phenylbenzimidazole-5-sulfonic acid | 274.0399 | 4.651 | 0.2 | 14227216 | 0 | 82.5 | | Spectral Match | | 767303.6 | 3335486 | 1434147 | 3810820 | NS | NS | NS | NS | NS | NS | NS | NS | NS | NS | 30.975 | 5.43E-09 | 0.003673 |
| 396 | C11918 | B | FALSE | 3-Hydroxy-2-methyl-4-pyrone | 126.0314 | 1.063 | 0.2 | 3152210 | 0 | | 79.8 | Spectral Match | | 166019.3 | 708495.2 | 430469.3 | 778001.6 | NS | NS | NS | NS | NS | NS | NS | NS | NS | NS | 49.952 | 7.88E-12 | 0.001665 |
| 397 | C11924 | B | FALSE | Perillic acid | 82.04226 | 0.986 | 0.2 | 1862435 | 0 | | | Spectral Match | | 39057.66 | 329095.6 | 246950.3 | 349631.9 | NS | NS | NS | NS | NS | NS | NS | NS | NS | NS | NS | NS | NS |
| 398 | C12026 | FB | FALSE | 3_5-Dihydroxy-phenylglycine | 183.0526 | 1.18 | 0.2 | 2142119 | 0 | | | Spectral Match | | 515773.3 | 411618.4 | 186496 | 467899 | 0.28178 | 0.0002 | 0 | 0.575 | 0.4836 | NS | NS | NS | NS | NS | NS | NS | NS |
| 399 | C12088 | FB | FALSE | 4-Hydroxycinnamoylmethane | 162.0676 | 13.075 | 0.2 | 469691.1 | 0 | | | Spectral Match | Superpathway of aromatic compound degradation via 2-oxopent-4-enoate;Other | 35202.53 | 32933.44 | 36393.44 | 32068.44 | 0.37582 | 0.0078 | 0 | 0.75 | 0.8693 | 0.57494 | 0.033126 | 0 | 0.566406 | 0.4942 | NS | NS | NS |
| 400 | C12109 | H | FALSE | AminoDHQ | 189.063 | 1.081 | 0.2 | 331548.3 | 0 | | | Spectral Match | Other | 7804.297 | 45884.62 | 18610.66 | 52703.12 | 2.107 | 0.0028 | 0.00025 | 0.555 | 0.0000 | 1.7642 | 0.002468 | 0.000857 | 0.972656 | 0.0000 | NS | NS | NS |
| 401 | C12110 | HF | FALSE | Gabaculine | 139.0629 | 1.08 | 0.2 | 1404040 | 0 | | 88.2 | Spectral Mat | | 56167.69 | 289720.9 | 125915.8 | 330672.1 | 1.7808 | 0.043 | 0 | 0.95 | 0.0001 | 1.5774 | 0.028741 | 0 | 0.863281 | 0.002 | NS | NS | NS |

| | | | | | | | | | | | | | | | | | | | | | | | | | | | | | | |
|---|---|---|---|---|---|---|---|---|---|---|---|---|---|---|---|---|---|---|---|---|---|---|---|---|---|---|---|---|---|---|
| | | | | | | | | | | | | ch | | | | | | | 3 | | | | | | | | 0 | | | |
| 402 | C12144 | HB | FALSE | 2-Amino-1,3,4-octadecanetriol | 317.2917 | 11.806 | 0.2 | 1075285 | 0 | 64.2 | | Spectral Match | | 38377.08 | 45648.45 | 46641.05 | 45400.3 | 0.48766 | 0.0001 | 0.000267 | 0.995 | 0.1185 | 0.64257 | 0.001877 | 0.000375 | 0.578125 | 0.6013 | NS | NS | NS |
| 403 | C12448 | H | FALSE | Ecgonine methyl ester | 199.1203 | 3.651 | 0.2 | 663348 0 | 0 | 57.1 | | Spectral Match | | 5006.911 | 5707.187 | 5005.65 | 5882.571 | 0.28821 | 0.0110 | 0.000921 | 0.53 | 0.2799 | 3.3056 | 3.93E-13 | 0 | 0.671875 | 0.0949 | NS | NS | NS |
| 404 | C12535 | B | FALSE | Tranexamic Acid | 157.1099 | 0.918 | 0.039 | 1168790 | 0 | | 79.9 | Spectral Match | | 145438.7 | 232434.4 | 123402 | 259692.5 | 0.60068 | 0.0317 | 0 | 0.95 | 0.4092 | NS | NS | NS | NS | NS | NS | NS | NS |
| 405 | C12835 | H | FALSE | 2,5-Dichloro-4-oxohex-2-enedioate | 225.9435 | 1.628 | 0.2 | 3122945 | 0 | | | Spectral Match | | 157773.87 | 131152.8 | 75775.54 | 144997.1 | 3.5718 | 0.0026 | 0 | 1 | 0.0000 | NS | NS | NS | NS | NS | NS | NS | NS |
| 406 | C12986 | H | FALSE | N2-Acetyl-L-aminoadipate | 203.0786 | 1.067 | 0.2 | 3441110 | 0 | | 72.4 | Spectral Match | | 864633.9 | 537494.3 | 228906.7 | 614641.1 | NS | NS | NS | NS | NS | NS | NS | NS | NS | NS | NS | NS | NS |
| 407 | C13235 | FB | FALSE | Atenolol | 266.1637 | 12.864 | 0.2 | 1033630 | 0 | | | Spectral Match | | 6758.388 | 9089.904 | 8285.648 | 9290.968 | 0.53988 | 0.0000 | 0.001887 | 0.64 | 0.0000 | NS | NS | NS | NS | NS | NS | NS | NS |
| 408 | C13635 | H | FALSE | 2'_4'-Dihydroxyacetophenone | 152.047 | 1.148 | 0.2 | 6008512 | 0 | | 67.3 | Spectral Match | Superpathway of aromatic compound degradation via 2-oxopent-4-enoate;Superpathway of aromatic compound degradation via 3-oxoadipate;Other | 43533.93 | 847373.9 | 202527.5 | 1008585 | 9.6118 | 0.0001 | 0.002144 | 1 | 0.0000 | 4.5856 | 0.003194 | 0.001479 | 1 | 0.0000 | NS | NS | NS |

| | | | | | | | | | | | | | | | | | | | | | | | | | | | | | | |
|---|---|---|---|---|---|---|---|---|---|---|---|---|---|---|---|---|---|---|---|---|---|---|---|---|---|---|---|---|---|---|
| 409 | C13688 | H | FALSE | delta-Guanidinovalericacid | 159.1002 | 0.846 | 0.2 | 2902877 | 0 | | | Spectral Match | | 39206.59 | 637875.9 | 250256.7 | 734780.7 | 5.568 | 0.0060 | 0.000333 | 0.985 | 0.0001 | NS | NS | NS | NS | NS | NS | NS | NS |
| 410 | C13747 | H | FALSE | Paraxanthine | 180.0641 | 2.563 | 0.2 | 1765280 | 0 | | 62.4 | Spectral Match | Superpathway of indole-3-acetate conjugate biosynthesis;Other | 18603.04 | 306948.7 | 225050.2 | 327423.3 | 5.6836 | 0.0022 | 0 | 0.93 | 0.0000 | NS | NS | NS | NS | NS | NS | NS | NS |
| 411 | C13800 | H | FALSE | 10-Nitrolinoleate | 325.2239 | 8.247 | 0.071 | 2073866 | 0 | 52.1 | | Spectral Match | | 133.7061 | 366454.9 | 4394.759 | 456969.9 | 1013.8 | 0.0000 | 0.000819 | 1 | 0.0000 | 71.477 | 3.47E-05 | 0.001943 | 1 | 0.0000 | NS | NS | NS |
| 412 | C13846 | H | FALSE | Stearamide | 283.2864 | 12.038 | 0.2 | 2482175 | 0 | 69.5 | | Spectral Match | | 16065.45 | 23449.03 | 30824.76 | 21605.1 | 0.60104 | 0.0035 | 0 | 1 | 0.0002 | 0.47633 | 2.62E-08 | 0.000986 | 0.953125 | 0.0000 | NS | NS | NS |
| 413 | C13914 | H | FALSE | N,N-Dimethylsphingosine | 327.3124 | 12.725 | 0.2 | 1100623 | 0 | 65.8 | | Spectral Match | | 12330.69 | 13993.11 | 13281.15 | 14171.1 | 0.45421 | 0.0000 | 0.000235 | 0.995 | 0.0996 | NS | NS | NS | NS | NS | NS | NS | NS |
| 414 | C13916 | H | FALSE | N-(Tetradecanoyl)-sphing-4-enine | 509.4785 | 19.95 | 0.2 | 3026026 | 0 | | | Spectral Match | | 141601.8 | 179546.5 | 148700.4 | 187258.1 | 0.4956 | 0.0000 | 0.000533 | 0.7 | 0.0258 | NS | NS | NS | NS | NS | NS | NS | NS |
| 415 | C14039 | H | FALSE | 1,1-Dichloroethylene | 95.952998 | 0.665 | 0.2 | 5273730 | 0 | | | Spectral Match | | 254210.4 | 726196.2 | 145815 | 871291.5 | NS | NS | NS | NS | NS | 5.0311 | 8.24E-05 | 0.00176 | 1 | 0.0000 | NS | NS | NS |
| 416 | C14175 | H | FALSE | Diethyl phthalate | 222.0884 | 10.938 | 0.2 | 1.33E+08 | 0 | 89 | | Spectral Match | | 1813110 | 4149143 | 3434557 | 4327790 | NS | NS | NS | NS | NS | NS | NS | NS | NS | NS | NS | NS | NS |
| 417 | C14214 | H | FALSE | Dibutyl Phthalate | 278.1508 | 0.34 | 0.2 | 709621.7 | 0 | | | Spectral Mat | Other | 176215.2 | 165630.3 | 164432.2 | 165929.8 | NS | NS | NS | NS | NS | NS | NS | NS | NS | NS | NS | NS | NS |

| | | | | | | | | | | | | | | | | | | | | | | | | | | | | | | |
|---|---|---|---|---|---|---|---|---|---|---|---|---|---|---|---|---|---|---|---|---|---|---|---|---|---|---|---|---|---|---|
| | | | | | | | | | | | | ch | | | | | | | | | | | | | | | | | | |
| 418 | C14235 | H | FALSE | Triphenyl phosphate | 326.0695 | 17.662 | 0.2 | 387159.5 | 0 | 82.4 | | Spectral Match | | 1759.608 | 1460.418 | 941.407 | 1589.496 | 0.31967 | 0.0000 | 0.003382 | 0.855 | 0.0553 | NS | NS | NS | NS | NS | NS | NS | NS |
| 419 | C14323 | H | FALSE | Butylate | 217.149 | 15.196 | 0.2 | 516038.1 | 0 | | 68.9 | Spectral Match | | 7429.247 | 19437.8 | 17697.71 | 19872.82 | NS | NS | NS | NS | NS | NS | NS | NS | NS | NS | NS | NS | NS |
| 420 | C14334 | H | FALSE | Propoxur | 209.1045 | 1.68 | 0.2 | 493246.6 | 0 | | | Spectral Match | | 13007.92 | 66617.46 | 14495.7 | 79647.9 | NS | NS | NS | NS | NS | 3.4619 | 4.13E-06 | 0.002 | 0.984375 | 0.0000 | NS | NS | NS |
| 421 | C14402 | H | FALSE | 1_2-Diaminobenzene | 108.0685 | 1.039 | 0.2 | 449469.3 | 0 | | 88.2 | Spectral Match | | 23173.97 | 86116.44 | 73957.31 | 89156.22 | NS | NS | NS | NS | NS | NS | NS | NS | NS | NS | NS | NS | NS |
| 422 | C14403 | H | FALSE | o-Toluidine | 107.0735 | 1.084 | 0.2 | 606238.9 | 0 | 75.6 | | Spectral Match | | 35478.2 | 102264.2 | 37990.67 | 118332.6 | NS | NS | NS | NS | NS | 1.9447 | 1.90E-07 | 0.000895 | 0.96875 | 0.0000 | NS | NS | NS |
| 423 | C14425 | FB | FALSE | Fenobucarb | 207.1254 | 2.055 | 0.2 | 1609225 | 0 | | 80.2 | Spectral Match | | 106121 | 245971.5 | 131776.4 | 274520.3 | NS | NS | NS | NS | NS | NS | NS | NS | NS | NS | NS | NS | NS |
| 424 | C14602 | H | FALSE | 3-Hydroxyaminophenol | 125.0475 | 1.094 | 0.2 | 429370.8 | 0 | | | Spectral Match | Other | 42279.21 | 87081.15 | 37405.83 | 99499.98 | NS | NS | NS | NS | NS | 1.6363 | 0.00063 | 0.000111 | 0.945313 | 0.0000 | NS | NS | NS |
| 425 | C14686 | H | FALSE | Dicyclohexylamine | 181.1826 | 7.703 | 0.2 | 1546433 | 0 | | | Spectral Match | | 183702.5 | 293575.2 | 311431.2 | 289111.2 | 0.64461 | 0.0109 | 0 | 0.845 | 0.0403 | 0.62312 | 0.002579 | 0 | 0.511719 | 0.6715 | NS | NS | NS |
| 426 | C14743 | H | FALSE | Isophorone | 138.1042 | 5.978 | 0.2 | 3963303 | 0 | 85.9 | 86.4 | Spectral Mat | Other | 11599.7 | 34263.55 | 32064.13 | 34813.4 | NS | NS | NS | NS | NS | NS | NS | NS | NS | NS | NS | NS | NS |

| | | | | | | | | | | | | | | | | | | | | | | | | | | | | | | |
|---|---|---|---|---|---|---|---|---|---|---|---|---|---|---|---|---|---|---|---|---|---|---|---|---|---|---|---|---|---|---|
| | | | | | | | | | | | | ch | | | | | | | | | | | | | | | | | | |
| 427 | C14832 | H | FALSE | [FA(18:0)]12R_13S-epoxy-9Z-octadecenoicacid | 296.2342 | 13.528 | 0.109 | 202074.1 | 0 | | | Spectral Match | | 8555.281 | 11585.05 | 18778.22 | 9786.758 | 0.57673 | 0.00061 | 0 | 0.81 | 0.1900 | 0.32831 | 5.61E-09 | 0.000143 | 0.90625 | 0.00000 | NS | NS | NS |
| 428 | C14853 | H | FALSE | Benzo[a]pyrene-7,8-dihydrodiol-9,10-oxide | 150.04 | 0.869 | 0.2 | 714548.6 | 0 | | | Spectral Match | | 18094.85 | 82495.03 | 81610.55 | 82716.15 | NS | NS | NS | NS | NS | NS | NS | NS | NS | NS | NS | NS | NS |
| 429 | C14857 | H | FALSE | 1,1-Dichloroethylene epoxide | 112.9554 | 1.306 | 0.2 | 431191.9 | 0 | | | Spectral Match | | 35122 | 31582.94 | 18396.69 | 34879.5 | 0.34805 | 0.00020 | 0.000143 | 0.69 | 0.6134 | NS | NS | NS | NS | NS | NS | NS | NS |
| 430 | C14874 | HFB | FALSE | Glutathione episulfonium ion | 166.0459 | 0.83 | 0.2 | 2962059 | 0 | | | Spectral Match | | 273215.5 | 678270.5 | 464283.4 | 731767.3 | NS | NS | NS | NS | NS | NS | NS | NS | NS | NS | NS | NS | NS |
| 431 | C15522 | B | FALSE | 4a-Hydroxytetrahydrobiopterin | 256.105 | 1.075 | 0.2 | 153669.4 | 0 | | | Spectral Match | | 17896 | 34637.76 | 28595.02 | 36148.45 | NS | NS | NS | NS | NS | NS | NS | NS | NS | NS | NS | NS | NS |
| 432 | C15532 | H | FALSE | N-Acetyl-L-citrulline | 216.099 | 7.49 | 0.2 | 15916720 | 0 | | | Spectral Match | | 3291667 | 2403000 | 2602678 | 2353080 | 0.28847 | 0.00000 | 0.000767 | 0.795 | 0.0253 | 0.57668 | 9.54E-06 | 0.0011 | 0.585938 | 0.3883 | NS | NS | NS |
| 433 | C15559 | B | FALSE | Gly-Pro(Glycylproline) | 172.0844 | 0.96 | 0.2 | 376677.4 | 0 | | 84 | Spectral Match | | 137827.6 | 69781.88 | 91307.13 | 64400.57 | 0.19468 | 0.00001 | 0.000854 | 0.565 | 0.0964 | 0.44969 | 0.004835 | -0.000014 | 0.617188 | 0.2846 | NS | NS | NS |
| 434 | C15584 | H | FALSE | Phenol | 94.04207 | 1.075 | 0.2 | 538190.7 | 0 | | | Spectral Match | Anaerobic aromatic compound degradation (Thauera aromatica) | 37686.54 | 89153.62 | 46233 | 99883.77 | NS | NS | NS | NS | NS | NS | NS | NS | NS | NS | NS | NS | NS |
| 435 | C15604 | HFB | FALSE | 3-Methylbut-2-enal | 84.0574 | 1.02 | 0.2 | 212956 | 0 | | | Spectral | | 80689.1 | 311916 | 64589.5 | 373747. | NS | NS | NS | NS | NS | 4.0272 | 2.27E- | 0.002051 | 1 | 0.00 | NS | NS | NS |

| | | | | | | | | | | | | | | | | | | | | | | | | | | | | | | |
|---|---|---|---|---|---|---|---|---|---|---|---|---|---|---|---|---|---|---|---|---|---|---|---|---|---|---|---|---|---|---|
| | | | | | 9 | 3 | | 1 | | | | Match | | 2 | | 7 | 6 | | | | | | | 05 | | | 00 | | | |
| 436 | C15606 | H | FALSE | 1,2-Dihydroxy-5-(methylthio)pent-1-en-3-one | 161.0272 | 0.688 | 0.2 | 6555881 | 0 | | | Spectral Match | | 341758.8 | 892008 | 2055151 | 601222.3 | NS | NS | NS | NS | NS | 0.18977 | 3.78E-08 | 0.000174 | 0.894531 | 0.0002 | NS | NS | NS |
| 437 | C15699 | H | FALSE | gamma-L-Glutamylputrescine | 217.1418 | 1.116 | 0.2 | 982099.7 | 0 | | | Spectral Match | Superpathway of L-arginine and L-ornithine degradation;Superpathway of L-arginine, putrescine, and 4-aminobutanoate degradation;Other | 2129.223 | 167609.7 | 5366.821 | 208170.5 | 24.645 | 0.0020 | 0.000526 | 0.71 | 0.0001 | 22.151 | 7.72E-06 | 0.003377 | 1 | 0.0000 | NS | NS | NS |
| 438 | C15700 | H | FALSE | gamma-Glutamyl-gamma-aminobutyraldehyde | 216.1103 | 1.012 | 0.04 | 2950666 | 0 | | 75 | Spectral Match | Superpathway of L-arginine and L-ornithine degradation;Superpathway of L-arginine, putrescine, and 4-aminobutanoate degradation;Other | 87371.2 | 692042 | 1031202 | 607252 | NS | NS | NS | NS | NS | NS | NS | NS | NS | NS | NS | NS | NS |
| 439 | C15767 | H | FALSE | gamma-Glutamyl-gamma-aminobutyraye | 232.1051 | 0.988 | 0.2 | 2841758 | 0 | | 93.1 | Spectral Match | Superpathway of L-arginine and L-ornithine degradation;Superpathway of L-arginine, putrescine, and 4-aminobutanoate | 131553 | 464776.1 | 112645.1 | 552808.8 | NS | NS | NS | NS | NS | 2.9006 | 0.024257 | 0 | 0.835938 | 0.0031 | NS | NS | NS |

| | | | | | | | | | | | | | | | | | | | | | | | | | | | | | | |
|---|---|---|---|---|---|---|---|---|---|---|---|---|---|---|---|---|---|---|---|---|---|---|---|---|---|---|---|---|---|---|
| | | | | | | | | | | | | | degradation;Other | | | | | | | | | | | | | | | | | |
| 440 | C15792 | H | FALSE | 3-Dehydroteasterone | 446.3382 | 16.0551 | 0.2 | 123377.5 | 0 | | | Spectral Match | Other | 9383.108 | 8768.561 | 6353.359 | 9372.361 | 0.36475 | 0.00000 | 0 | 1 | 0.5050 | NS | NS | NS | NS | NS | NS | NS | NS |
| 441 | C16309 | F | FALSE | [FAoxo(12:0)]12-oxo-10E-dodecenoic acid | 212.1406 | 7.604 | 0.2 | 178164.6 | 0 | | | Spectral Match | Other | 2130.404 | 27913.34 | 5172.663 | 33598.5 | 4.9955 | 0.00009 | 0 | 0.58 | 0.0000 | 4.4468 | 6.21E-06 | 0.000835 | 1 | 0.00000 | NS | NS | NS |
| 442 | C16328 | HFB | FALSE | trans-2-Enoyl-OPC8-CoA | 348.1111 | 13.113 | 0.2 | 456237.5 | 0 | | | Spectral Match | | 29273.23 | 29225.68 | 34301.3 | 27956.78 | 0.40408 | 0.00000 | 0.000836 | 0.555 | 0.8862 | 0.52791 | 2.82E-06 | 0.000593 | 0.664063 | 0.0744 | NS | NS | NS |
| 443 | C16334 | H | FALSE | 3-Oxo-OPC6-CoA | 513.6276 | 8.753 | 0.2 | 895115.7 | 0 | | | Spectral Match | | 22920.43 | 153245.9 | 68410.24 | 174454.8 | 2.4303 | 0.0268 | 0 | 1 | 0.0000 | NS | NS | NS | NS | NS | NS | NS | NS |
| 444 | C16359 | H | FALSE | 1-Methyluric acid | 183.0512 | 1.181 | 0.2 | 6182005 | 0 | | | Spectral Match | | 14140.7 | 1405088 | 1089119 | 1484080 | 35.882 | 0.0016 | 5.00E-04 | 1 | 0.0000 | NS | NS | NS | NS | NS | NS | NS | NS |
| 445 | C16365 | B | FALSE | 5-Acetylamino-6-formylamino-3-methyluracil | 112.0282 | 0.687 | 0.2 | 1200762 | 0 | | | Spectral Match | | 68305.37 | 175620.2 | 388345.8 | 122438.8 | NS | NS | NS | NS | NS | 0.2017 | 8.63E-06 | 0.001103 | 0.792969 | 0.0061 | NS | NS | NS |
| 446 | C16391 | B | FALSE | 2,4,6-Trinitrotoluene | 227.0177 | 1.071 | 0.2 | 2277712 | 0 | | | Spectral Match | | 102725.5 | 552686.8 | 707404.3 | 514007.4 | NS | NS | NS | NS | NS | NS | NS | NS | NS | NS | NS | NS | NS |
| 447 | C16399 | H | FALSE | 4-Ethylbenzaldehyde | 152.0834 | 5.732 | 0.2 | 531371.9 | 0 | 56.5 | 78.6 | Spectral Match | | 198322.34 | 378881.6 | 224983.7 | 417356.6 | NS | NS | NS | NS | NS | NS | NS | NS | NS | NS | NS | NS | NS |
| 4 | C16 | H | FA | 4-(N- | 20 | 1 | 0. | 23 | 0 | | | Spe | | 26 | 37 | 79 | 27 | 66 | 0. | 0. | 0. | 0. | 0. | 0. | 0 | 0. | 0. | N | NS | NS |

| | | | | | | | | | | | | | | | | | | | | | | | | | | | | | | |
|---|---|---|---|---|---|---|---|---|---|---|---|---|---|---|---|---|---|---|---|---|---|---|---|---|---|---|---|---|---|---|
| 48 | 453 | | LSE | Nitrosomethylamino)-1-(3-pyridyl)-1-butanone | 8.1081 | 0.444 | 2 | 20933 | | | | ctral Match | | 14.858 | 9538.2 | 2751.5 | 6234.8 | .842 | 0000 | 000706 | 99 | 0000 | 24312 | 013959 | | 640625 | 0869 | S | | |
| 449 | C16503 | H | FALSE | Methylfarnesoate | 250.1923 | 8.415 | 0.2 | 631668.5 | 0 | | | Spectral Match | | 468.0767 | 121237 | 813.561 | 151342.9 | 96.627 | 0.0010 | 0 | 1 | 0.0001 | 127.31 | 4.14E-11 | 0.001349 | 1 | 0.0000 | NS | NS | NS |
| 450 | C16526 | B | FALSE | [FA(20:0)]11Z-eicosenoicacid | 310.2858 | 19.25 | 0.2 | 1244241 | 0 | | 60.4 | Spectral Match | | 35647.52 | 122675.5 | 406330.3 | 51761.8 | NS | NS | NS | NS | NS | 0.082322 | 5.97E-09 | 0.002323 | 0.988281 | 0.0000 | NS | NS | NS |
| 451 | C16538 | H | FALSE | 1_5-Anhydro-D-mannitol | 164.068 | 1.028 | 0.2 | 15358894 | 0 | | | Spectral Match | Other | 99130.86 | 2618929 | 2428665 | 2666495 | 10.87 | 0.0003 | 0.000372 | 1 | 0.0000 | NS | NS | NS | NS | NS | NS | NS | NS |
| 452 | C16551 | H | FALSE | Alcophosphamide | 138.0102 | 0.681 | 0.2 | 7464535 | 0 | | 72.3 | Spectral Match | | 144458 | 1008630 | 2360886 | 670566.4 | NS | NS | NS | NS | NS | 0.18747 | 4.34E-08 | 0.00025 | 0.894531 | 0.0001 | NS | NS | NS |
| 453 | C16607 | H | FALSE | [Similar to: Amiloride; ΔMass: 57.9612 Da] | 171.0866 | 1.614 | 0.2 | 4103786 | 0 | | 83.3 | Spectral Match | | 46401.34 | 110981.3 | 452294.02 | 127403.1 | NS | NS | NS | NS | NS | NS | NS | NS | NS | NS | NS | NS | NS |
| 454 | C16612 | H | FALSE | Citalopram aldehyde | 294.0937 | 1.013 | 0.2 | 891316 | 0 | | | Spectral Match | | 311928.5 | 141068.2 | 175623.7 | 132429.3 | 0.17918 | 0.0001 | 0.00045 | 1 | 0.0458 | 0.46967 | 0.018455 | 0 | 0.640625 | 0.2496 | NS | NS | NS |
| 455 | C16615 | H | FALSE | 6-Methylthiopurine 5'-monophosphate ribonucleotide | 188.0124 | 9.325 | 0.2 | 39095221 | 0 | | | Spectral Match | | 1294932 | 5904605 | 13515096 | 4001982 | NS | NS | NS | NS | NS | NS | NS | NS | NS | NS | NS | NS | NS |
| 456 | C16618 | H | FALSE | [Similar to: Isoprenaline; ΔMass: | 191.016 | 0.86 | 0.2 | 106943 | 0 | | 85 | Spectral | | 10454.8 | 223701 | 138521. | 244996 | 7.5307 | 0.00 | 0.0005 | 0.765 | 0.0000 | NS | NS | NS | NS | NS | NS | NS | NS |

| | | | | | | | | | | | | | | | | | | | | | | | | | | | | | | |
|---|---|---|---|---|---|---|---|---|---|---|---|---|---|---|---|---|---|---|---|---|---|---|---|---|---|---|---|---|---|---|
| | | | | 20.1045 Da] | 4 | 1 | | 1 | | | | Match | | 8 | | 4 | | | 00 | 96 | | | | | | | | | | |
| 457 | C16633 | H | FALSE | [Similar to: Ethylene glycol tetraacetic acid (EGTA); ΔMass: 119.0912 Da] | 261.0519 | 20.75 | 0.2 | 1787632 | 0 | | 69.2 | Spectral Match | | 422007.5 | 313752 | 317194.7 | 312891.4 | 0.29886 | 0.0000 | 0 | 0.975 | 0.0570 | NS | NS | NS | NS | NS | NS | NS | NS |
| 458 | C16673 | H | FALSE | L-isoglutamine | 144.0894 | 1.191 | 0.2 | 857552.8 | 0 | | 64.1 | Spectral Match | | 20702.85 | 181315.9 | 65922.83 | 210164.2 | NS | NS | NS | NS | NS | NS | NS | NS | NS | NS | NS | NS | NS |
| 459 | C16715 | H | FALSE | 2-oxo-1_2-dihydroquinoline-4-carboxylate | 189.042 | 16.644 | 0.2 | 332363.4 | 0 | | 81.6 | Spectral Match | Other | 10776.97 | 29179.55 | 21984.15 | 30978.4 | NS | NS | NS | NS | NS | NS | NS | NS | NS | NS | NS | NS | NS |
| 460 | C16759 | H | FALSE | allylcysteine | 161.0506 | 1.271 | 0.2 | 1078737 | 0 | | | Spectral Match | Other | 10255.94 | 168861.5 | 86641.6 | 189416.5 | NS | NS | NS | NS | NS | NS | NS | NS | NS | NS | NS | NS | NS |
| 461 | C16843 | H | FALSE | 1-deoxynojirimycin | 163.0839 | 0.939 | 0.2 | 598184 | 0 | | | Spectral Match | Superpathway of linamarin and lotaustralin biosynthesis;Superpathway of butirocin biosynthesis;Other | 121461.9 | 142686.1 | 72154.85 | 160318.9 | 0.4146 | 0.0010 | 0.000235 | 0.67 | 0.8636 | NS | NS | NS | NS | NS | NS | NS | NS |
| 462 | C17241 | B | FALSE | 4-methylthiobutanaldoxime | 133.0558 | 2.318 | 0.2 | 253401 | 0 | | 75.9 | Spectral Match | | 10302.36 | 15006.8 | 14149.84 | 15221.04 | 0.57515 | 0.0058 | 0 | 0.525 | 0.1201 | NS | NS | NS | NS | NS | NS | NS | NS |
| 463 | C17355 | B | FALSE | [Similar to: Isoprenaline; ΔMass: 19.5635 Da] | 191.5574 | 10.945 | 0.2 | 5386740 | 0 | | 92 | Spectral Match | | 470119.3 | 373665.2 | 287640.6 | 395171.3 | 0.31254 | 0.0000 | 0.001976 | 0.83 | 0.0108 | NS | NS | NS | NS | NS | NS | NS | NS |

| | | | | | | | | | | | | | | | | | | | | | | | | | | | | | | |
|---|---|---|---|---|---|---|---|---|---|---|---|---|---|---|---|---|---|---|---|---|---|---|---|---|---|---|---|---|---|---|
| 464 | C17541 | B | FALSE | Undecaprenyl-diphospho-N-acetylmuramoyl-(N-acetylglucosamine)-L-alanyl-D-isoglutaminyl-L-lysyl-(glycyl)-D-alanyl-D-alanine | 964.5388 | 8.527 | 0.2 | 1164460 | 0 | | | Spectral Match | | 9594.842 | 37444.08 | 17001.54 | 42554.72 | NS | NS | NS | NS | NS | 1.682 | 0.008465 | 0.000275 | 1 | 0.0000 | NS | NS | NS |
| 465 | C17928 | H | FALSE | 2-methylhistamine | 125.0949 | 1.493 | 0.2 | 1608829 | 0 | | | Spectral Match | | 5733.456 | 278892.8 | 6970.422 | 346873.4 | NS | NS | NS | NS | NS | 27.892 | 4.83E-05 | 0.00161 | 1 | 0.0000 | NS | NS | NS |
| 466 | C17954 | H | FALSE | albaflavenone | 218.1663 | 13.616 | 0.2 | 513943.1 | 0 | | 78.3 | Spectral Match | | 19983.31 | 7336.245 | 5180.598 | 7875.156 | 0.14354 | 0.0000 | 0.001124 | 0.73 | 0.0000 | NS | NS | NS | NS | NS | NS | NS | NS |
| 467 | C18015 | H | FALSE | momilactoneA | 314.1872 | 10.307 | 0.2 | 90086.16 | 0 | | 92.9 | Spectral Match | | 8316.386 | 8235.23 | 2742.218 | 9608.484 | 0.37792 | 0.0000 | 0.000211 | 1 | 0.6199 | 2.4132 | 2.19E-05 | 0.001513 | 1 | 0.0000 | NS | NS | NS |
| 468 | C18043 | H | FALSE | Cholesterol sulfate | 465.3066 | 10.172 | 0.2 | 20517838 | 0 | | | Spectral Match | | 35583.88 | 4769261 | 3457076 | 5097307 | 49.451 | 0.0000 | 0.002472 | 0.51 | 0.0000 | NS | NS | NS | NS | NS | NS | NS | NS |
| 469 | C18059 | H | FALSE | Unsaturated CDP-archaeol | 1021.56 | 8.46 | 0.2 | 174397.5 | 0 | | | Spectral Match | Other | 5827.053 | 16064.06 | 16260.38 | 16014.98 | NS | NS | NS | NS | NS | NS | NS | NS | NS | NS | NS | NS | NS |
| 470 | C18132 | H | FALSE | 3''-hydroxy-geranylhydroquinone | 262.1559 | 7.644 | 0.2 | 171030.9 | 0 | | | Spectral Match | | 4702.484 | 28445.83 | 2778.474 | 34862.67 | NS | NS | NS | NS | NS | 7.1825 | 1.98E-08 | 0.00393 | 1 | 0.0000 | NS | NS | NS |
| 4 | C18 | H | FA | Desmediph | 30 | 8. | 0. | 14 | 0 | | | Spe | | 22 | 25 | 12 | 28 | 3. | 0. | 0. | 1 | 0. | NS | NS | NS | NS | N | N | NS | NS |

| | | | | | | | | | | | | | | | | | | | | | | | | | | | | | | |
|---|---|---|---|---|---|---|---|---|---|---|---|---|---|---|---|---|---|---|---|---|---|---|---|---|---|---|---|---|---|---|
| 71 | 417 | | LSE | am | 0.1096 | 666 | 2 | 16029 | | | | ctral Match | | 016.32 | 1201.9 | 1211.9 | 3699.4 | 8406 | 0194 | 000143 | | 0001 | | | | | S | S | | |
| 472 | C18525 | H | FALSE | Methoxyfenozide | 368.2118 | 10.016 | 0.2 | 455698.5 | 0 | | | Spectral Match | | 17065.53 | 8583.23 | 73757.85 | 88539.58 | NS | NS | NS | NS | NS | NS | NS | NS | NS | NS | NS | NS | NS |
| 473 | C18786 | H | FALSE | Dodemorph | 281.2708 | 15.187 | 0.2 | 280293.6 | 0 | | | Spectral Match | | 5417.748 | 7661.667 | 9435.597 | 7218.185 | 0.57091 | 0.0011 | 0 | 0.975 | 0.0004 | 0.50947 | 1.42E-07 | 0.00198 | 0.867188 | 0.0002 | NS | NS | NS |
| 474 | C18794 | B | FALSE | Monolinuron | 214.0502 | 3.447 | 0.2 | 157238.3 | 0 | | | Spectral Match | | 61235.66 | 36165.75 | 24527.79 | 39075.24 | 0.2334 | 0.0001 | 0 | 0.96 | 0.2753 | NS | NS | NS | NS | NS | NS | NS | NS |
| 475 | C18796 | H | FALSE | 2-Hydroxy-2-methylbutyronitrile | 99.068854 | 1.693 | 0.2 | 32059642 | 0 | | | Spectral Match | Superpathway of linamarin and lotaustralin biosynthesis | 1040180 | 6810907 | 1989673 | 8016216 | NS | NS | NS | NS | NS | NS | NS | NS | NS | NS | NS | NS | NS |
| 476 | C18868 | FB | FALSE | Napropamide | 271.1565 | 6.973 | 0.2 | 75173.72 | 0 | | | Spectral Match | | 5613.215 | 14076.42 | 21713.38 | 12167.18 | NS | NS | NS | NS | NS | 0.38116 | 0.000436 | -0.00032 | 0.679688 | 0.0506 | NS | NS | NS |
| 477 | C18885 | H | FALSE | Propamocarb | 188.1519 | 0.821 | 0.2 | 4583370 | 0 | | | Spectral Match | Biotin biosynthesis II;Other | 19706.39 | 721747.8 | 352194.4 | 814136.1 | 12.633 | 0.0016 | 0 | 1 | 0.0001 | NS | NS | NS | NS | NS | NS | NS | NS |
| 478 | C18894 | H | FALSE | Spinetoram 1 | 747.4955 | 0.838 | 0.2 | 75107.5 | 0 | | | Spectral Match | | 248.9721 | 1546.572 | 1278.027 | 1613.708 | 2.3874 | 0.0000 | 0 | 0.615 | 0.0000 | NS | NS | NS | NS | NS | NS | NS | NS |
| 479 | C19266 | H | FALSE | 4,4'-Bis(diethylamino)benzophenone | 324.2188 | 13.184 | 0.2 | 693909.7 | 0 | 74.5 | | Spectral Match | | 105155 | 111401 | 66501.74 | 122625.8 | 0.42209 | 0.0017 | 0.000105 | 0.975 | 0.9805 | NS | NS | NS | NS | NS | NS | NS | NS |
| 4 | C19 | H | FA | Sudan 1 | 24 | 9. | 0. | 79 | 0 | | | Spe | | 43 | 62 | 55 | 64 | 0. | 0. | 0. | 0. | 0. | NS | NS | NS | NS | N | N | NS | NS |

| | | | | | | | | | | | | | | | | | | | | | | | | | | | | | | |
|---|---|---|---|---|---|---|---|---|---|---|---|---|---|---|---|---|---|---|---|---|---|---|---|---|---|---|---|---|---|---|
| 80 | 525 | | LSE | | 8.094 | 944 | 2 | 7237.1 | | | | ctral Match | | 080.89 | 720.85 | 564.77 | 509.87 | 58154 | 0002 | 000235 | 53 | 0001 | | | | | S | S | | |
| 481 | C19539 | H | FALSE | [Similar to: 2,4,5-Trimethylaniline; ΔMass: 1.0684 Da] | 134.0364 | 14.687 | 0.2 | 722420.9 | 0 | | 65.5 | Spectral Match | | 13466.98 | 55321.73 | 30932.75 | 61418.97 | NS | NS | NS | NS | NS | NS | NS | NS | NS | NS | NS | NS | NS |
| 482 | C19566 | H | FALSE | 4-Hydroxy-4-(methylnitrosoamino)-1-(3-pyridinyl)-1-butanone | 222.088 | 10.319 | 0.2 | 1.25E+08 | 0 | 64.6 | 65.8 | Spectral Match | | 163460.6 | 1076908 | 470624.3 | 1228479 | 2.3786 | 0.0001 | 0.000941 | 0.97 | 0.0000 | 1.6404 | 0.002704 | 0 | 0.964844 | 0.0000 | NS | NS | NS |
| 483 | C19577 | H | FALSE | 1-(Methylnitrosoamino)-4-(3-pyridinyl)-1,4-butanediol | 225.1105 | 1.08 | 0.2 | 197042.5 | 0 | | | Spectral Match | | 21170.37 | 37623.72 | 16590.84 | 42881.94 | 0.61669 | 0.0240 | 0 | 1 | 0.1680 | NS | NS | NS | NS | NS | NS | NS | NS |
| 484 | C19580 | H | FALSE | alpha-[3-[(Hydroxymethyl)nitrosoamino]propyl]-3-pyridinemethanol | 226.1197 | 13.959 | 0.2 | 268982.9 | 0 | | | Spectral Match | | 12484.81 | 46304.21 | 54502.36 | 44254.68 | NS | NS | NS | NS | NS | NS | NS | NS | NS | NS | NS | NS | NS |
| 485 | C19781 | B | FALSE | Prolinamide | 114.0791 | 0.919 | 0.2 | 4100625 | 0 | 55.4 | | Spectral Match | | 186758.7 | 766251.4 | 455928.7 | 843832.1 | NS | NS | NS | NS | NS | NS | NS | NS | NS | NS | NS | NS | NS |
| 486 | C19787 | B | FALSE | 5'-S-Methyl-5'-thioinosine | 148.0302 | 1.013 | 0.2 | 592096.8 | 0 | | | Spectral Match | | 31651.65 | 53270.32 | 44007.42 | 55586.05 | NS | NS | NS | NS | NS | NS | NS | NS | NS | NS | NS | NS | NS |
| 487 | C19798 | B | FALSE | 4,4'-Diapolycopenedial | 427.2661 | 19.41 | 0.2 | 5006119 | 0 | | | Spectral Mat | | 1693737 | 1058856 | 1252672 | 1010402 | 0.23914 | 0.003 | 0.000333 | 0.66 | 0.2190 | NS | NS | NS | NS | NS | NS | NS | NS |

| | | | | | | | | | | | | | | | | | | | | | | | | | | | | | | |
|---|---|---|---|---|---|---|---|---|---|---|---|---|---|---|---|---|---|---|---|---|---|---|---|---|---|---|---|---|---|---|
| | | | | | | 6 | | | | | | ch | | | | | | | 2 | | | | | | | | | | | |
| 488 | C19884 | H | FALSE | Lipid IIA | 1536.926 | 13.603 | 0.2 | 3227083 | 0 | | | Spectral Match | | 588846.77 | 503977.67 | 528415.6 | 497866.7 | 0.34628 | 0.0000 | 0.000517 | 0.735 | 0.0955 | 0.6253 | 0.000287 | 0.001179 | 0.609375 | 0.4270 | NS | NS | NS |
| 489 | C19891 | F | FALSE | 1D-chiro-inositol | 180.0627 | 2.58 | 0.2 | 1158930 | 0 | | | Spectral Match | Superpathway of indole-3-acetate conjugate biosynthesis;Other | 6050.349 | 171302.8 | 131908.3 | 210830.9 | NS | NS | NS | NS | NS | 8.217 | 0.025073 | 0.000361 | 0.816406 | 0.0037 | NS | NS | NS |
| 490 | C20126 | H | FALSE | Methylone | 207.0888 | 1.306 | 0.2 | 219261.3 | 0 | | | Spectral Match | | 375569.08 | 442644.49 | 215119.83 | 499550.66 | 0.41133 | 0.0305 | 0 | 0.515 | 0.9835 | NS | NS | NS | NS | NS | NS | NS | NS |
| 491 | C20199 | HB | FALSE | 3-[(3aS,4S,7aS)-7a-Methyl-1,5-dioxo-octahydro-1H-inden-4-yl]propanoate | 238.1197 | 7.69 | 0.2 | 234785.1 | 0 | | | Spectral Match | Other | 472.8336 | 1447.208 | 2395.564 | 1210.119 | NS | NS | NS | NS | NS | 0.33487 | 1.70E-15 | 0.002451 | 1 | 0.0000 | NS | NS | NS |
| 492 | C20239 | B | FALSE | [Similar to: 2-[(2-Oxo-2-{4-[4-(trifluoromethyl)-2-pyrimidinyl]-1,4-diazepan-1-yl}ethyl)sulfanyl]acetic acid; ΔMass: 166.0186 Da] | 212.0788 | 1.054 | 0.2 | 3273628 | 0 | | 72.4 | Spectral Match | | 75612.61 | 489604.4 | 26316.73 | 605426.3 | NS | NS | NS | NS | NS | 13.412 | 2.16E-07 | 0.002913 | 1 | 0.0000 | NS | NS | NS |
| 493 | C20246 | H | FALSE | 2-[(2R,5Z)-2-Carboxy-4-methylthiazol-5(2H)- | 266.9974 | 6.866 | 0.2 | 5813016 | 0 | | | Spectral Match | Other | 1154793 | 836309.3 | 368537.3 | 953252.2 | 0.28494 | 0.0000 | 0.000444 | 0.78 | 0.0749 | 1.7876 | 0.003815 | 0 | 1 | 0.0000 | NS | NS | NS |

| | | | | | | | | | | | | | | | | | | | | | | | | | | | | | | |
|---|---|---|---|---|---|---|---|---|---|---|---|---|---|---|---|---|---|---|---|---|---|---|---|---|---|---|---|---|---|---|
| | | | | ylidene]ethyl phosphate | | | | | | | | | | | | | | | | | | | | | | | | | | |
| 494 | C20310 | H | FALSE | N-Hydroxy-L-isoleucine | 147.0892 | 0.914 | 0.117 | 1558812 | 0 | | | Spectral Match | Superpathway of linamarin and lotaustralin biosynthesis | 127418.7 | 345282.4 | 300893.9 | 356379.6 | NS | NS | NS | NS | NS | NS | NS | NS | NS | NS | NS | NS | NS |
| 495 | C20313 | H | FALSE | N-Hydroxy-L-valine | 133.0735 | 0.957 | 0.2 | 2662253 | 0 | | 93.9 | Spectral Match | Superpathway of linamarin and lotaustralin biosynthesis | 40509.41 | 378352.1 | 54435.6 | 459331.2 | NS | NS | NS | NS | NS | 4.9368 | 0.000412 | 0 | 0.960938 | 0.0000 | NS | NS | NS |
| 496 | C20314 | H | FALSE | N,N-Dihydroxy-L-valine | 149.0685 | 0.345 | 0.2 | 2352918 | 0 | | 64.7 | Spectral Match | Superpathway of linamarin and lotaustralin biosynthesis | 127365 | 122136.4 | 116359.9 | 123580.6 | NS | NS | NS | NS | NS | NS | NS | NS | NS | NS | NS | NS | NS |
| 497 | C20643 | B | FALSE | 2-Heptyl-4(1H)-quinolone | 243.1615 | 7.625 | 0.2 | 2256627 | 0 | | | Spectral Match | Other | 172397.3 | 409457.8 | 277497.3 | 442447.9 | NS | NS | NS | NS | NS | NS | NS | NS | NS | NS | NS | NS | NS |
| 498 | C20940 | B | FALSE | L-Dihydroanticapsin | 201.0994 | 1.038 | 0.2 | 1289681 | 0 | | | Spectral Match | | 192034.8 | 281971.6 | 146951.4 | 315726.7 | 0.53094 | 0.0132 | 0 | 0.66 | 0.3970 | NS | NS | NS | NS | NS | NS | NS | NS |
| 499 | C20941 | HB | FALSE | L-Anticapsin | 199.0838 | 1.071 | 0.2 | 42926376 | 0 | | | Spectral Match | | 413130 | 9482671 | 3736097 | 10919315 | 7.7004 | 0.000 | 0.000671 | 1 | 0.0000 | NS | NS | NS | NS | NS | NS | NS | NS |
| 500 | C21015 | H | FALSE | [Similar to: Propylthiouracil; ΔMass: 55.0058 Da] | 115.0456 | 11.807 | 0.2 | 787727.9 | 0 | | 71.8 | Spectral Match | | 175444.4 | 22736.46 | 19258.71 | 23605.9 | 0.51388 | 0.0000 | 0.000111 | 0.69 | 0.0020 | NS | NS | NS | NS | NS | NS | NS | NS |
| 501 | C21189 | H | FALSE | Symmetricdimethylarginine | 202.1424 | 0.884 | 0.2 | 861359.2 | 0 | | | Spectral Match | | 452744.4 | 159110.5 | 111632.3 | 170980 | NS | NS | NS | NS | NS | NS | NS | NS | NS | NS | NS | NS | NS |
| 5 | C21 | H | FA | [FAhydroxy | 18 | 1. | 0. | 21 | 0 | | | Spe | Superpathway | 40 | 32 | 19 | 35 | 0. | 0. | 0. | 0. | 0. | NS | NS | NS | NS | N | N | NS | NS |

| | | | | | | | | | | | | | | | | | | | | | | | | | | | | | | |
|---|---|---|---|---|---|---|---|---|---|---|---|---|---|---|---|---|---|---|---|---|---|---|---|---|---|---|---|---|---|---|
| 02 | 195 | | LSE | (4:0)]N-(3S-hydroxy-butanoyl)-homoserine lactone | 7.0838 | 082 | 105 | 12408 | | | | ctral Match | of L-lysine degradation | 7162.7 | 4769 | 8615.8 | 6307.3 | 29476 | 0000 | 000211 | 67 | 4381 | | | | | S | S | | |
| 503 | C21201 | B | FALSE | N-(3-Oxododecanoyl)homoserinelactone | 297.1929 | 8.133 | 0.2 | 4643381 | 0 | | 74.6 | Spectral Match | | 31825.55 | 949454.6 | 523395.64 | 1173719 | NS | NS | NS | NS | NS | 14.139 | 0.004919 | 0 | 0.960938 | 0.0006 | NS | NS | NS |
| 504 | C21559 | B | FALSE | N-[(2S)-2-Amino-2-carboxyethyl]-L-glutamate | 234.843 | 1.125 | 0.095 | 21730285 | 0 | | | Spectral Match | | 9389321 | 1062917 | 1218418 | 1024041 | 0.03933 | 0.0337 | 0 | 0.56 | 0.2882 | NS | NS | NS | NS | NS | NS | NS | NS |
| 505 | C21984 | B | FALSE | Deacyl-lipid IVA | 1178.656 | 9.013 | 0.2 | 2221257 | 0 | | | Spectral Match | | 37899.36 | 110906.3 | 111768.7 | 110690.7 | NS | NS | NS | NS | NS | 0.65873 | 0.001078 | -0.00025 | 0.507813 | 0.8966 | NS | NS | NS |
| 506 | C22072 | B | FALSE | [Similar to: (1S)-1,5-Anhydro-1-[2,4,6-trihydroxy-3-(4-hydroxybenzoyl)phenyl]-D-glucitol; ΔMass: 270.0759 Da] | 138.0297 | 8.309 | 0.2 | 24640933 | 0 | | 78 | Spectral Match | | 312683.8 | 617598 | 931077.3 | 539228.2 | NS | NS | NS | NS | NS | 0.37073 | 0.026619 | 0.000308 | 0.648438 | 0.4697 | NS | NS | NS |
| 507 | C22101 | F | FALSE | N5-Citryl-D-ornithine | 152.0453 | 8.106 | 0.2 | 389379.9 | 0 | | | Spectral Match | | 13128.53 | 69983.63 | 70295.52 | 69905.65 | NS | NS | NS | NS | NS | NS | NS | NS | NS | NS | NS | NS | NS |
| 508 | C22136 | H | FALSE | [ST(3:0)]cholest-5Z_7Z_24-trien-3beta-ol | 382.3217 | 19.148 | 0.2 | 1579939 | 0 | | | Spectral Match | Other | 611564.4 | 83555.18 | 59380.93 | 89598.75 | 0.048941 | 0.0000 | 0.001228 | 0.505 | 0.0000 | NS | NS | NS | NS | NS | NS | NS | NS |
| 509 | C22163 | H | FALSE | Roquefortine D | 391.201 | 12.7 | 0.2 | 344705 | 0 | | | Spectral | Other | 692.625 | 61095.7 | 10007.7 | 73867.7 | 28.885 | 0.00 | 0.0014 | 0.555 | 0.0000 | 4.3291 | 0.0006 | 0 | 0.9492 | 0.00 | NS | NS | NS |

| | | | | | | | | | | | | | | | | | | | | | | | | | | | | | | |
|---|---|---|---|---|---|---|---|---|---|---|---|---|---|---|---|---|---|---|---|---|---|---|---|---|---|---|---|---|---|---|
| | | | | | 6 | 98 | | .3 | | | | Match | | 4 | 4 | 6 | 3 | | 00 | 71 | | | | 85 | | 19 | 01 | | | |
| 510 | HMDB0000734 | H | FALSE | Indoleacrylicacid | 187.0628 | 2.614 | 0.2 | 10092105 | 0 | | | Spectral Match | | 1482077 | 2379980 | 2956731 | 2235792 | 0.59955 | 0.03006 | 0 | 1 | 0.5480 | 0.46637 | 0.009056 | 0 | 0.589844 | 0.3791 | NS | NS | NS |
| 511 | HMDB0000872 | H | FALSE | Tetradecanedioicacid | 258.1825 | 0.259 | 0.2 | 796083.1 | 0 | | 83.4 | Spectral Match | | 233469.8 | 234729.1 | 234764.5 | 234720.2 | NS | NS | NS | NS | NS | NS | NS | NS | NS | NS | NS | NS | NS |
| 512 | HMDB0001431 | H | FALSE | Pyridoxamine;2-Methyl-4-aminomethyl-5-hydroxymethyl-3-Pyridinol | 168.0894 | 1.076 | 0.2 | 311804 | 0 | | | Spectral Match | Other | 82623.52 | 75901.1 | 48004.79 | 82875.18 | 0.33829 | 0.0001 | 0.000111 | 0.635 | 0.8624 | NS | NS | NS | NS | NS | NS | NS | NS |
| 513 | HMDB0004824 | H | FALSE | N2,N2-Dimethylguanosine | 311.121 | 20.325 | 0.2 | 1660571 | 0 | | | Spectral Match | | 188065.8 | 285815.8 | 263881.9 | 291299.3 | 0.60288 | 0.0012 | 0.000107 | 0.515 | 0.0045 | NS | NS | NS | NS | NS | NS | NS | NS |
| 514 | HMDB0004949 | H | FALSE | N-Palmitoyl-D-erythro-sphingosine | 537.5096 | 17.86 | 0.2 | 3213028 | 0 | | | Spectral Match | | 840822.7 | 415051.3 | 35865.74 | 509847.7 | NS | NS | NS | NS | NS | 10.091 | 7.09E-13 | 0.003703 | 1 | 0.0000 | NS | NS | NS |
| 515 | HMDB0012273 | H | FALSE | Hexadecanamide;Palmiticamide | 255.2551 | 15.04 | 0.2 | 331385.1 | 0 | | | Spectral Match | | 34674.12 | 28358.62 | 42450.09 | 24835.75 | 0.3525 | 0.0000 | 0.000105 | 0.88 | 0.1985 | 0.39489 | 4.68E-07 | 0.000133 | 0.875 | 0.0001 | NS | NS | NS |
| 516 | HMDB0014010 | H | FALSE | 5-HydroxyOmeprazole | 361.108 | 8.779 | 0.2 | 777272.8 | 0 | | | Spectral Match | | 972.3651 | 127445.9 | 268879.3 | 92087.52 | 54.926 | 0.0000 | 0.00025 | 0.7 | 0.0000 | NS | NS | NS | NS | NS | NS | NS | NS |
| 517 | HMDB0014693 | H | FALSE | Xanthoxin;Methoxsalen | 250.156 | 7.974 | 0.2 | 739928.4 | 0 | | | Spectral Match | Other | 133166.4 | 133147.1 | 168195.8 | 162229 | 3.1818 | 0.0494 | 0 | 1 | 0.0006 | 5.6037 | 5.96E-07 | 0.003863 | 1 | 0.0000 | NS | NS | NS |

| | | | | | | | | | | | | | | | | | | | | | | | | | | | | | | |
|---|---|---|---|---|---|---|---|---|---|---|---|---|---|---|---|---|---|---|---|---|---|---|---|---|---|---|---|---|---|---|
| 518 | HMDB0028695 | H | FALSE | Ala-Pro(Alanyl-Proline) | 186.0998 | 0.939 | 0.068 | 2953545 | 0 | | | Spectral Match | | 33259.99 | 489797.7 | 311235.2 | 534438.4 | 5.1707 | 0.0011 | 0 | 0.96 | 0.0000 | NS | NS | NS | NS | NS | NS | NS | NS |
| 520 | HMDB0030283 | H | FALSE | Cinchonine; Hydrocinchonine | 296.1879 | 12.059 | 0.2 | 198195.2 | 0 | | | Spectral Match | | 13747.69 | 34998.98 | 24215.09 | 37694.95 | NS | NS | NS | NS | NS | NS | NS | NS | NS | NS | NS | NS | NS |
| 521 | HMDB003419 | H | FALSE | CucurbitacinA | 574.317 | 8.915 | 0.2 | 138617.9 | 0 | | | Spectral Match | | 743.876 | 7012.609 | 2811.454 | 8062.898 | 3.5195 | 0.0000 | 0.000917 | 0.975 | 0.0000 | 1.8963 | 2.67E-05 | 0.000725 | 1 | 0.0000 | NS | NS | NS |
| 522 | HMDB0059805 | H | FALSE | [Similar to: 5-Hydroxyindole; ΔMass: 27.9948 Da] | 105.058 | 3.94 | 0.2 | 2781025 | 0 | | 88.5 | Spectral Match | | 3161.782 | 3814.92 | 2251.621 | 4205.745 | 0.47045 | 0.0000 | 0.000267 | 1 | 0.3286 | NS | NS | NS | NS | NS | NS | NS | NS |
| 523 | HMDB0061822 | H | FALSE | PEG n6 | 282.1667 | 5.674 | 0.2 | 1300198 | 0 | 71.4 | | Spectral Match | | 12228.12 | 53453.8 | 24454.3 | 60703.68 | 1.6287 | 0.0209 | 0 | 0.675 | 0.0000 | 1.654 | 0.00326 | 0.000182 | 1 | 0.0000 | NS | NS | NS |
| 524 | HMDB0061835 | H | FALSE | PEG n7 | 326.1927 | 6.543 | 0.081 | 18977183 | 0 | 56.7 | | Spectral Match | | 369885.5 | 2621110 | 722268.2 | 3095820 | 2.5386 | 0.0079 | 0 | 1 | 0.0000 | 2.7489 | 1.94E-05 | 0.000462 | 0.996094 | 0.0000 | NS | NS | NS |
| 525 | HMDB00701 | H | FALSE | Hexanoylglycine | 173.1047 | 2.521 | 0.2 | 1990492 | 0 | | | Spectral Match | | 386582.8 | 497301 | 348398.4 | 534526.6 | NS | NS | NS | NS | NS | NS | NS | NS | NS | NS | NS | NS | NS |
| 526 | HMDB0094680 | H | FALSE | [Similar to: PEG n8; ΔMass: 0.0016 Da] | 370.2187 | 6.94 | 0.2 | 21715478 | 0 | | 82.4 | Spectral Match | | 317551 | 2995600 | 1037689 | 3485078 | 3.4183 | 0.0000 | 0.001417 | 0.99 | 0.0000 | 2.1575 | 0.000142 | 0.000987 | 0.992188 | 0.0000 | NS | NS | NS |
| 527 | HMDB0240294 | H | FALSE | 2-O-Methylascorbic acid | 190.0457 | 1.324 | 0.2 | 343998.2 | 0 | | | Spectral Mat | | 146720.8 | 9505.52 | 7476.404 | 10012.8 | 0.02332 | 0.000 | 0.001008 | 1 | 0.0000 | NS | NS | NS | NS | NS | NS | NS | NS |

| | | | | | | | | | | | | | | | | | | | | | | | | | | | | | | |
|---|---|---|---|---|---|---|---|---|---|---|---|---|---|---|---|---|---|---|---|---|---|---|---|---|---|---|---|---|---|---|
| | | | | | | | | | | | | ch | | | | | | | 0 | | | | | | | | | | | |
| 528 | HMDB0248559 | H | FALSE | ArachidoylEthanolamide | 355.3435 | 12.762 | 0.2 | 1211336 | 0 | | 68.8 | Spectral Match | | 150448.4 | 136059 | 64929.43 | 153841.3 | 0.33631 | 0.0000 | 0 | 0.945 | 0.5019 | NS | NS | NS | NS | NS | NS | NS | NS |
| 529 | HMDB0255195 | H | FALSE | Propylnorapomorphine | 295.1559 | 9.429 | 0.2 | 208964.5 | 0 | | | Spectral Match | | 25736.75 | 30379.54 | 20219.89 | 32919.46 | 0.50834 | 0.0006 | 0.000635 | 0.59 | 0.6392 | NS | NS | NS | NS | NS | NS | NS | NS |
| 530 | HMDB0259191 | H | FALSE | Triethyl phosphate | 182.0703 | 8.496 | 0.2 | 565958.2 | 0 | 51.4 | | Spectral Match | | 45712.12 | 52536.28 | 76683.63 | 46499.45 | 0.47512 | 0.0001 | 0 | 0.57 | 0.3037 | 0.40752 | 7.63E-12 | 0.002307 | 1 | 0.0000 | NS | NS | NS |
| 531 | HMDB05015 | H | FALSE | Gabapentin | 171.1253 | 1.05 | 0.2 | 3617778 | 0 | | | Spectral Match | | 153748.4 | 24311.03 | 12496.94 | 27264.55 | 0.005815 | 0.0000 | 0.002262 | 0.645 | 0.0000 | NS | NS | NS | NS | NS | NS | NS | NS |
| 532 | HMDB12273 | H | FALSE | Hexadecanamide | 255.2552 | 11.102 | 0.2 | 8717625 | 0 | 80.1 | | Spectral Match | | 378852.4 | 305613.2 | 404182.5 | 280970.9 | 0.33194 | 0.0000 | 0.000222 | 0.98 | 0.0423 | 0.4824 | 3.60E-07 | 0.002355 | 0.925781 | 0.0000 | NS | NS | NS |
| 533 | HMDB13078 | H | FALSE | Stearoyl ethanolamide | 327.3125 | 13.127 | 0.2 | 1848495 | 0 | 52.7 | | Spectral Match | | 10803.55 | 9724.911 | 13192.4 | 8858.039 | 0.3701 | 0.0000 | 0.001236 | 0.78 | 0.2020 | 0.45023 | 6.76E-10 | 0.001061 | 0.980469 | 0.0000 | NS | NS | NS |
| 534 | HMDB14494 | H | FALSE | Minoxidil | 209.127 | 6.638 | 0.2 | 2293349 | 0 | 69.7 | | Spectral Match | | 146273.2 | 403349.1 | 161481.9 | 463815.9 | NS | NS | NS | NS | NS | 1.9974 | 0.015381 | -0.00013 | 0.988281 | 0.0000 | NS | NS | NS |
| 535 | HMDB14515 | FB | FALSE | Meprobamate | 218.1273 | 0.867 | 0.2 | 118298.7 | 0 | | | Spectral Match | | 1732.949 | 13975.84 | 3992.254 | 16471.73 | 3.0012 | 0.0002 | 0 | 0.75 | 0.0000 | 2.7375 | 1.80E-07 | 0.000963 | 0.96875 | 0.0000 | NS | NS | NS |
| 536 | HMDB14581 | H | FALSE | Allopurinol | 136.0381 | 1.061 | 0.2 | 3310322 | 0 | | | Spectral Mat | Other | 42710.73 | 737327.1 | 318356.3 | 842069.8 | 5.8546 | 0.034 | 0 | 0.975 | 0.0006 | NS | NS | NS | NS | NS | NS | NS | NS |

| | | | | | | | | | | | | | | | | | | | | | | | | | | | | | | |
|---|---|---|---|---|---|---|---|---|---|---|---|---|---|---|---|---|---|---|---|---|---|---|---|---|---|---|---|---|---|---|
| | | | | | | | | | | | | ch | | | | | | | 1 | | | | | | | | | | | |
| 537 | HMDB15377 | H | FALSE | Isocarboxazide | 231.0999 | 1.573 | 0.2 | 490074.2 | 0 | | | Spectral Match | | 1278.761 | 73596.56 | 6946.355 | 90259.11 | 16.791 | 0.0002 | -0.000011 | 1 | 0.0000 | 7.1793 | 0.000505 | 0 | 0.945313 | 0.0001 | NS | NS | NS |
| 538 | HMDB15520 | H | FALSE | Oxprenolol | 265.1667 | 7.402 | 0.2 | 2274521 | 0 | | | Spectral Match | | 4191.819 | 376289.3 | 66043.4 | 453850.8 | 24.495 | 0.0000 | 0.001461 | 1 | 0.0000 | NS | NS | NS | NS | NS | NS | NS | NS |
| 539 | HMDB28760 | H | FALSE | Aspartyl-Phenylalanine | 280.1049 | 1.431 | 0.2 | 215667.1 | 0 | | | Spectral Match | | 1431.13 | 43180.57 | 72206.62 | 35924.05 | 11.539 | 0.0000 | 0.001011 | 1 | 0.0000 | 0.32464 | 0.013805 | 0 | 0.597656 | 0.2761 | NS | NS | NS |
| 540 | HMDB29846 | H | FALSE | Pseudocapsaicin | 293.1979 | 12.981 | 0.2 | 5592916 | 0 | | | Spectral Match | | 29114.35 | 52079.54 | 41857.78 | 54634.97 | NS | NS | NS | NS | NS | NS | NS | NS | NS | NS | NS | NS | NS |
| 541 | HMDB32142 | H | FALSE | 2,4-Dimethylbenzaldehyde | 134.0727 | 11.756 | 0.2 | 421715 | 0 | 53.8 | | Spectral Match | | 33510.99 | 74447.01 | 80724.2 | 72877.72 | NS | NS | NS | NS | NS | 0.5644 | 0.000515 | 0.000333 | 0.535156 | 0.6119 | NS | NS | NS |
| 542 | HMDB33843 | H | FALSE | N-Vinyl-2-pyrrolidone | 111.0685 | 1.253 | 0.2 | 1358235 | 0 | 80.4 | | Spectral Match | | 6773.527 | 17940.18 | 15948.9 | 18438 | NS | NS | NS | NS | NS | NS | NS | NS | NS | NS | NS | NS | NS |
| 543 | HMDB34738 | H | FALSE | Isotrichodermin | 292.1662 | 13.947 | 0.2 | 1415330 | 0 | | 83.5 | Spectral Match | Other | 19780.2 | 33202.76 | 28607.79 | 34351.5 | 0.66483 | 0.0016 | 0 | 1 | 0.0000 | NS | NS | NS | NS | NS | NS | NS | NS |
| 544 | HMDB38249 | H | FALSE | 1-Aminocyclohexanecarboxylic Acid | 143.0942 | 0.927 | 0.2 | 20635316 | 0 | | 92.9 | Spectral Match | | 1581505 | 4671604 | 5640961 | 4429264 | NS | NS | NS | NS | NS | NS | NS | NS | NS | NS | NS | NS | NS |
| 545 | HMDB38642 | H | FALSE | Fumitremorgin C | 379.1912 | 10.64 | 0.2 | 435192.7 | 0 | | | Spectral Mat | Other | 485.2776 | 82081.21 | 536.7893 | 102467.3 | 63.008 | 0.0015 | 0 | 1 | 0.0010 | 130.62 | 7.86E-08 | 0.001778 | 1 | 0.000 | NS | NS | NS |

| | | | | | | | | | | | | | | | | | | | | | | | | | | | | | | |
|---|---|---|---|---|---|---|---|---|---|---|---|---|---|---|---|---|---|---|---|---|---|---|---|---|---|---|---|---|---|---|
| | | | | | | 3 | | | | | | ch | | | | | | | 2 | | | | | | | | 0 | | | |
| 546 | HMDB41220 | H | FALSE | Dibutyl sebacate | 314.24448 | 0.31 | 0.2 | 4438177 | 0 | 92.2 | | Spectral Match | | 13664452 | 16289981 | 16250041 | 16299966 | NS | NS | NS | NS | NS | NS | NS | NS | NS | NS | NS | NS | NS |
| 547 | HMDB41925 | H | FALSE | Methandienone | 300.20082 | 11.416 | 0.2 | 166040.8 | 0 | | | Spectral Match | Other | 562.1957 | 28625.52 | 32732.82 | 27598.7 | 20.75 | 0.0000 | 0.000917 | 0.94 | 0.0000 | 0.55548 | 0.021882 | 0 | 0.574219 | 0.4703 | NS | NS | NS |
| 548 | HMDB41927 | H | FALSE | Methcathinone | 163.0994 | 3.943 | 0.2 | 876887.1 | 0 | | 78.6 | Spectral Match | | 3451.736 | 6348.502 | 4970.576 | 6692.984 | NS | NS | NS | NS | NS | NS | NS | NS | NS | NS | NS | NS | NS |

**Supplementary table 2- Total metaproteome identified in Acute liver failure Plasma samples in Healthy control , Survival ALF and Non-survival ALF**

| S.no. | Variable | Peptide | lca | superkingdom | phylum | subphylum | class | subclass | order | suborder | family | genus | speciesgroup | speciessubgroup | species | subspecies | strain | EC-names | GO (cellular component) | GO (molecular function) | GO (biological processes) | GO (cellular component) - name | GO (molecular function) - names | GO (biological process) - names | InterPro | InterPro-names | HEALTHY AVERAGE | SURVIVAL | NON-SURVIVAL | FC (ALF VS H) | pval (ALF VS H) | Mean Decrease Accuracy (Random forest: | AUC (ALF VS H) | FC (NS VS S) | pval (NS VS S) | Mean Decrease Accuracy (NSV | AUC (NS VS S) | Fc.value (H;S;NS | P value (H;S;NS) | Mean Decrease Accuracy (H |
|---|---|---|---|---|---|---|---|---|---|---|---|---|---|---|---|---|---|---|---|---|---|---|---|---|---|---|---|---|---|---|---|---|---|---|---|---|---|---|---|---|

| | | | | | | | | | | | | | | | | | | | | | | | | | | | | | | | | | | | | | | | | |
|---|---|---|---|---|---|---|---|---|---|---|---|---|---|---|---|---|---|---|---|---|---|---|---|---|---|---|---|---|---|---|---|---|---|---|---|---|---|---|---|---|
| | | | | | | | | | | | | | | | | | | | | | | **s** | | | | | | | | | | **(ALFVSH)** | | | | **SS)** | | | | **;S;NS)** |
| 129 | Variable 214 | LTPAHLHVLK | Streptomycescoelicolor | Bacteria | Actinobacteria | | Actinomycetia | | Streptomycetales | | Streptomycetaceae | Streptomyces | Streptomyces albidoflavus group | | Streptomyces coelicolor | | | 1.1.1.108(100%) | Carnitine 3-dehydrogenase (100%) | GO:0005737(100%) | cytoplasm(100%) | GO:0047728(100%);GO:0070403(67%);GO:0 | carnitine 3-dehydrogenase activity(100%);NAD+ binding(67%);oxidoreductase activity | acting on the CH-OH group of donors | NAD or NADP as acceptor(33%) | GO:0042413(100%);GO:0006631(67%) | 1.00E+00 | 1.00E+00 | 1.36E+06 | 90.988 | 0.0043083 | 0.00029412 | 0.9 | 54.342 | 0.013872 | 0.00082443 | 1 | | | |

| | | | | | | | | | | | | | | | | | | | | | | | | | | | | | | | | | | | | | | | | |
|---|---|---|---|---|---|---|---|---|---|---|---|---|---|---|---|---|---|---|---|---|---|---|---|---|---|---|---|---|---|---|---|---|---|---|---|---|---|---|---|---|
| | | | | | | | | | | | | | | | | | | | | | | 0166616(33%) | | | | | | | | | | | | | | | | | | |
| 203 | Variable 281 | TAAAAVAWASR | Bacillales | Bacteria | Firmicutes | | Bacilli | | Bacillales | | | | | | | | | 2.3.1.-(100%) | Transferring groups other than amino-acyl groups (100%) | | | GO:0031177(95%);GO:0008483(78%);GO | phosphopantetheine binding (95%);transaminase activity (78%);pyridoxal phosphate bindin | GO:0006633(77%);GO:0017000(32%) | fatty acid biosynthetic process (77%);antibiotic biosynthetic process | IPR000873(100%);IPR010071(99%);IPR | 1.00E+00 | 1.00E+00 | 1.11E+06 | 120.13 | 2.4691E-06 | 0.000057566 | 0.9 | 52.267 | 0.0019063 | 0.00069784 | 1 | 9.248 | 4.70E-04 | 0.000625 |

| | | | | | | | | | | | | | | | | | | | | | | | | | | | | | | | | | | | | | | | | |
|---|---|---|---|---|---|---|---|---|---|---|---|---|---|---|---|---|---|---|---|---|---|---|---|---|---|---|---|---|---|---|---|---|---|---|---|---|---|---|---|---|
| | | | | | | | | | | | | | | | | | | | | | | :0030170(78%) | g (78%) | | (32%) | 042099(99%) | | | | | | | | | | | | | | |
| 387 | Variable 447 | VVTLEVEGR | Gammaproteobacteria | Bacteria | Proteobacteria | | Gammaproteobacteria | | | | | | | | | | | 2.1.1.177(100%) | 23S rRNA (pseudouridine(1915)-N(3))-methyltransferase (100%) | GO:0005737(100%) | cytoplasm(100%) | GO:0070038(100%) | rRNA (pseudouridine-N3-)-methyltransferase activity (100%) | | | IPR003742(100%);IPR029026(100%); | 1.00E+00 | 1.00E+00 | 2.52E+06 | 133.65 | 0.00042097 | 0.00086806 | 0.9 | 105.28 | 0.00087904 | 0.00052013 | 1 | 5.7834 | 6.05E-03 | 0.00045386 |

| | | | | | | | | | | | | | | | | | | | | | | | | | | | | | | | | | | | | | |
|---|---|---|---|---|---|---|---|---|---|---|---|---|---|---|---|---|---|---|---|---|---|---|---|---|---|---|---|---|---|---|---|---|---|---|---|---|---|
| | | | | | | | | | | | | | | | | | | | | | | | | | | IPR029028(100%) | | | | | | | | | | | |
| 428 | Variable 484 | QVYGNANLYNK | Enterobacteriaceae | Bacteria | Proteobacteria | | Gammaproteobacteria | | Enterobacterales | | Enterobacteriaceae | | | | | | | | | GO:0005737(40%);GO:0031361(40 | cytoplasm(40%);integral component of thylakoid membran | GO:0005506(40%);GO:0009055(40 | iron ion binding(40%);electron transfer activity(40%);heme binding(40%) | GO:0015979(40%);GO:0035864(40%) | photosynthesis(40%);response to potassium ion(40%) | IPR018392(100%);IPR036779(100 | 1.00E+00 | 1.00E+00 | 1.52E+06 | 74.012 | 0.00085182 | 0.001 | 0.9 | 77.141 | 0.045234 | 0.00043536 | 1 |

| | | | | | | | | | | | | | | | | | | | | | | | | | | | | | | | | | | | | | | | | |
|---|---|---|---|---|---|---|---|---|---|---|---|---|---|---|---|---|---|---|---|---|---|---|---|---|---|---|---|---|---|---|---|---|---|---|---|---|---|---|---|---|
| | | | | | | | | | | | | | | | | | | | | %) | e(40%) | %);GO:0020037(40%) | | | | %);IPR007055(99%) | | | | | | | | | | | | | | |
| 498 | Variable 547 | GEFLDGNGEK | Mycoplasmamobile163K | Bacteria | Tenericutes | | Mollicutes | | Mycoplasmatales | | Mycoplasmataceae | Mycoplasma | | | Mycoplasmamobile | | Mycoplasmamobile163K | | | GO:0005840(100%) | ribosome(100%) | GO:0003729(100%);GO:0003735 | mRNA binding(100%);structural constituent of ribosome(100%);rRNA bin | GO:0006412(100%) | translation(100%) | IPR001351(100%);IPR004044(1 | 1.00E+00 | 1.00E+00 | 8.45E+05 | 49.409 | 0.000050401 | 0.000088235 | 0.9 | 45.547 | 0.021443 | 0.00072388 | 1 | 5.8067 | 5.94E-03 | 0.00052668 |

| | | | | | | | | | | | | | | | | | | | | | | | | | | | | | | | | | | | | | | | | |
|---|---|---|---|---|---|---|---|---|---|---|---|---|---|---|---|---|---|---|---|---|---|---|---|---|---|---|---|---|---|---|---|---|---|---|---|---|---|---|---|---|
| | | | | | | | | | | | | | | | | | | | | | | (100%);GO:0019843(100%) | ding(100%) | | | 00%);IPR004087(100%) | | | | | | | | | | | | | | |
| 1 | Variable 1 | FGTPDDFR | Bacteria | Bacteria | | | | | | | | | | | | | | 2.4.1.18(100%) | 1 | 4-alpha-glucan branching e | GO:0005737(0%);GO:0005829(0%);GO:00 | cytoplasm (0%);cytosol( | GO:0003844(100%);GO:0004553(97%);GO:010275 | 1 | 4-alpha-glucan branching enzyme activity(100 | hydrolyzing O-glycosyl com | 7.61E+05 | 1.79E+06 | 3.81E+05 | 0.12844 | 3.497E-08 | 0.00081645 | 0.73 | 0.0056218 | 0.00022439 | 0.0078274 | 0.88671875 | 259.16 | 2.35E-24 | 0.012504 |

| | | | | | | | | | | | | | | | | | | | | | | | | | | | | | | | | | | | | | | | | |
|---|---|---|---|---|---|---|---|---|---|---|---|---|---|---|---|---|---|---|---|---|---|---|---|---|---|---|---|---|---|---|---|---|---|---|---|---|---|---|---|---|
| | | | | | | | | | | | | | | | | | | | | nzyme (100%) | 05886 (0%) | 0%); plasma membrane (0%) | 2 (96%) | | %); hydrolase activity | pounds (97%); 1 | | | | | | | | | | | | | | |
| 10 | Variable 107 | AMGSLGMSSPGGVR | Mycobacterium tuberculosis complex | Bacteria | Actinobacteria | | Actinomycetia | | Corynebacteriales | | Mycobacteriaceae | Mycobacterium | Mycobacterium tuberculosis | | | | | | | GO:0005886 (100%); GO:00160 | plasma membrane (100%); integral component of m | | | GO:0006869 (11%); GO:0071555 (11%); GO:0046506 | lipid transport (11%); cell wall organization (11%); sul | IPR004869 (100%); IPR000731 | 1.00E+00 | 1.00E+00 | 2.94E+05 | 21.344 | 0.00083854 | 0.00016402 | 0.9 | 12.45 | 0.034845 | 0.00053477 | 1 | 5.3476 | 8.53E-03 | 0.0022893 |

| | | | | | | | | | | | | | | | | | | | | | | | | | | | | | | | | | | | | | | | | |
|---|---|---|---|---|---|---|---|---|---|---|---|---|---|---|---|---|---|---|---|---|---|---|---|---|---|---|---|---|---|---|---|---|---|---|---|---|---|---|---|---|
| | | | | | | | | | | | | | complex | | | | | | | 21(100%);GO:0005618(5%) | embrane(100%);cell wall(5%) | | | (5%) | folipid biosynthetic process(5%) | (74%);IPR004707(26%) | | | | | | | | | | | | | | |
| 12 | Variable 109 | TAPATVPLAR | Mycobacterium tuberculosis com | Bacteria | Actinobacteria | | Actinomycetia | | Corynebacteriales | | Mycobacteriaceae | Mycobacterium | Mycobacterium tubercul | | | | | 3.6.1.-(67%);3.6.1.55(3 | Inphosphorus-containing anhydrides(67%);8-oxo-dGTP dip | | | GO:0046872(94%);GO:001 | metal ion binding(94%);hydrolase activity(88%);8-oxo-7 | 8-dihydrodeoxyguanosine triphosphate pyrophosph | GO:0006260(12%);GO:0006281(12%) | DNA replication(12%);DNA repair | 1.00E+00 | 1.00E+00 | 1.06E+06 | 78.672 | 7.0114E-06 | 0.000078947 | 0.9 | 57.679 | 0.0034614 | 0.000088919 | 1 | 7.9347 | 1.19E-03 | 0.00037024 |

| | | | | | | | | | | | | | | | | | | | | | | | | | | | | | | | | | | | | | | | | |
|---|---|---|---|---|---|---|---|---|---|---|---|---|---|---|---|---|---|---|---|---|---|---|---|---|---|---|---|---|---|---|---|---|---|---|---|---|---|---|---|---|
| | | | plex | | | | | | | | | | osis complex | | | | | 3%) | hosphatase (33%) | | | 6787(88%);GO:0035539(12%) | | atase activity (12%) | | (12%) | | | | | | | | | | | | | | |
| 19 | Variable 115 | QWALAGLTPLR | Mycobacterium tuberculosis c | Bacteria | Actinobacteria | | Actinomycetia | | Corynebacteriales | | Mycobacteriaceae | Mycobacterium | Mycobacterium tuberc | | | | | 1.1.1.133(100%) | dTDP-4-dehydrorhamnose reductase (100%) | GO:0005829(6%) | cytosol (6%) | GO:0008831(100%);GO: | dTDP-4-dehydrorhamnose reductase activity (100%);meta | GO:0019305(100%);GO:0005975(6%);GO: | dTDP-rhamnose biosynthetic process | IPR029903(100%);IPR0 | 1.00E+00 | 1.00E+00 | 2.52E+05 | 20.091 | 1.0231E-06 | 0.0001794 | 0.9 | 11.716 | 0.0030259 | 0.0010458 | 1 | 10.772 | 1.67E-04 | 0.00052484 |

| | | | | | | | | | | | | | | | | | | | | | | | | | | |
|---|---|---|---|---|---|---|---|---|---|---|---|---|---|---|---|---|---|---|---|---|---|---|---|---|---|---|
| | | | omplex | | | | | | | | | | ulosis complex | | | | | | | | | 0046872(12%) | l ion binding (12%) | 00452226 (6%) | (100%);carbohydrate metabolic process (6%);extracellular polysaccharide biosynthetic proces | 36291(100%);IPR005913(94%) |

| | | | | | | | | | | | | | | | | | | | | | | | | | | | | | | | | | | | | | |
|---|---|---|---|---|---|---|---|---|---|---|---|---|---|---|---|---|---|---|---|---|---|---|---|---|---|---|---|---|---|---|---|---|---|---|---|---|---|
| | | | | | | | | | | | | | | | | | | | | | | | | | s(6%) | | | | | | | | | | | | |
| 20 | Variable 116 | GVAAVFQAAQLAGMGR | Mycobacterium tuberculosis complex | Bacteria | Actinobacteria | | Actinomycetia | | Corynebacteriales | | Mycobacteriaceae | Mycobacterium | Mycobacterium tuberculosis complex | | | | | 3.1.11.5(59%);3.6.4.12(41%) | Exodeoxyribonuclease V (59%); DNA helicase (41%) | GO:0005618(6%);GO:0005829(6%);GO:0005886( | cell wall(6%);cytosol(6%);plasma membrane(6%) | GO:0003677(100%);GO:0005524(100%);GO:0008 | DNA binding(100%);ATP binding(100%);exodeoxyribonuclease V activity(100%) | GO:0000724(59%);GO:0006281(41%);GO:0000725(6%) | double-strand break repair via homologous recombination(59%);DNA repair(41 | IPR002121(100%);IPR004586(100%);IPR027417 | 1.00E+00 | 1.00E+00 | 1.34E+06 | 34.5 | 0.022411 | 0.0001039 | 0.9 | 41.7 | 0.042103 | 0.00018669 | 1 |

| | | | | | | | | | | | | | | | | | | | | | | | | | | | | | | | | | | | | | |
|---|---|---|---|---|---|---|---|---|---|---|---|---|---|---|---|---|---|---|---|---|---|---|---|---|---|---|---|---|---|---|---|---|---|---|---|---|---|
| | | | | | | | | | | | | | | | | | | | | 6%) | | 854(100%) | | | %);recombinational repair(6%) | (100%) | | | | | | | | | | | |
| 33 | Variable 128 | ATEYPSAVAK | Mycobacterium tuberculosis complex | Bacteria | Actinobacteria | | Actinomycetia | | Corynebacteriales | | Mycobacteriaceae | Mycobacterium | Mycobacterium tuberculosis com | | | | | | | GO:0015934(100%);GO:0005618( | large ribosomal subunit(100%);cell wall(7%);cytosolic large rib | GO:0003735(100%);GO:0019843( | structural constituent of ribosome(100%);rRNA binding(100%) | GO:0006412(100%);GO:0042255(7%) | translation(100%);ribosome assembly(7%) | IPR001063(100%);IPR005727(10 | 1.00E+00 | 1.18E+05 | 2.16E+06 | NS | NS | NS | NS | 0.4839 | 0.002729 | 0.00032723 | 0.5117875 |

| | | | | | | | | | | | | | | | | | | | | | | | | | | | | | | | | | | | | | | | | |
|---|---|---|---|---|---|---|---|---|---|---|---|---|---|---|---|---|---|---|---|---|---|---|---|---|---|---|---|---|---|---|---|---|---|---|---|---|---|---|---|---|
| | | | | | | | | | | | | | plex | | | | | | | 7%);GO:0022625(7%) | osomal subunit(7%) | 100%) | | | | 0%);IPR018260(100%) | | | | | | | | | | | | | | |
| 69 | Variable 160 | QAALLQAFGSVR | Beutenbergia cavernae DSM 12333 | Bacteria | Actinobacteria | | Actinomycetia | | Micrococcales | | Beutenbergiaceae | Beutenbergia | | | Beutenbergia cavernae | | Beutenbergia cavernae DSM 12333 | | | GO:0005737(100%);GO:000938 | cytoplasm(100%);excinuclease repair complex(100% | GO:0003677(100%);GO:000938 | DNA binding(100%);excinuclease ABC activity(100%) | GO:0006289(100%);GO:0009432(100%) | nucleotide-excision repair(100%);SOS response(100 | IPR000305(100%);IPR001162( | 1.00E+00 | 1.00E+00 | 9.81E+04 | 9.9413 | 1.682E-07 | 0.0005397 | 0.9 | 4.7777 | 0.047017 | 0.0015496 | 1 | 13.993 | 2.20E-05 | 0.0018711 |

| | | | | | | | | | | | | | | | | | | | | | | | | | | | | | | | | | | | | | | | | |
|---|---|---|---|---|---|---|---|---|---|---|---|---|---|---|---|---|---|---|---|---|---|---|---|---|---|---|---|---|---|---|---|---|---|---|---|---|---|---|---|---|
| | | | | | | | | | | | | | | | | | | | | 0(100%) | ) | 1(100%) | | | %) | 100%);IPR001943(100%) | | | | | | | | | | | | | | |
| 80 | Variable 170 | FGLFYTDR | Tropherymawhipplei | Bacteria | Actinobacteria | | Actinomycetia | | Micrococcales | | Tropherymataceae | Tropheryma | | | Tropherymawhipplei | | | 6.1.1.3(100%) | Threonine--tRNA ligase (100%) | GO:0005737(100%) | cytoplasm(100%) | GO:0000049(100%);GO:0004 | tRNA binding(100%);threonine-tRNA ligase activity(100%);A | GO:0006435(100%) | threonyl-tRNA aminoacylation(100%) | IPR002314(100%);IPR002232 | 1.00E+00 | 3.88E+05 | 9.22E+04 | 305.71 | 1.573E-11 | 0.0038319 | 1 | 0.0067196 | 0.017369 | 0.0014924 | 0.5625 | 64.289 | 1.65E-13 | 0.0015392 |

| | | | | | | | | | | | | | | | | | | | | | | | | | | | | | | | | | | | | | | | | |
|---|---|---|---|---|---|---|---|---|---|---|---|---|---|---|---|---|---|---|---|---|---|---|---|---|---|---|---|---|---|---|---|---|---|---|---|---|---|---|---|---|
| | | | | | | | | | | | | | | | | | | | | | | 829(100%);GO:0005524(100%) | TP binding (100%) | | | 0(100%);IPR004154(100%) | | | | | | | | | | | | | | |
| 83 | Variable 173 | AAQESGDFER | Salinispora | Bacteria | Actinobacteria | | Actinomycetia | | Micromonosporales | | Micromonosporaceae | Salinispora | | | | | | | | GO:0005886(100%);GO | plasmamembrane(100%);integralco | | | | | IPR005372(100%) | 1.00E+00 | 1.42E+05 | 7.84E+04 | 133.11 | 3.4832E-09 | 0.0013312 | 1 | 0.014065 | 0.0060944 | 0.0043755 | 0.7421875 | 61.318 | 3.48E-13 | 0.001789 |

| | | | | | | | | | | | | | | | | | | | | | | | | | | | | | | | | | | | | | | | | |
|---|---|---|---|---|---|---|---|---|---|---|---|---|---|---|---|---|---|---|---|---|---|---|---|---|---|---|---|---|---|---|---|---|---|---|---|---|---|---|---|---|
| | | | | | | | | | | | | | | | | | | | | :0016021(100%) | mponent of membrane (100%) | | | | | | | | | | | | | | | | | | | |
| 93 | Variable 182 | GHQVVEASEAGEK | Streptomyces | Bacteria | Actinobacteria | | Actinomycetia | | Streptomycetales | | Streptomycetaceae | Streptomyces | | | | | | 1.14.13.170(100%) | Pentalenolactone D synthase (100%) | | | GO:0004497(67%);GO:0004499(17 | monooxygenase activity (67%);N | N-dimethylaniline monooxygenase activity (17%);oxidoreductase activity | acting on paired donors | with incorporation or reduction of molec | 1.00E+00 | 1.00E+00 | 7.67E+05 | 26.002 | 0.00015143 | 0 | 0.9 | 25.472 | 0.00066438 | 0.00037155 | 1 | 4.6913 | 1.45E-02 | 0.00046958 |

| | | | | | | | | | | | | | | | | | | | | | | | | | | | | | | | | | | | | | |
|---|---|---|---|---|---|---|---|---|---|---|---|---|---|---|---|---|---|---|---|---|---|---|---|---|---|---|---|---|---|---|---|---|---|---|---|---|---|
| | | | | | | | | | | | | | | | | | | | | | | %);GO:0016709(17%) | | | | ularoxygen | | | | | | | | | | | |
| 111 | Variable 199 | AQAFSLTTTFTPNGR | Streptomyces | Bacteria | Actinobacteria | | Actinomycetia | | Streptomycetales | | Streptomycetaceae | Streptomyces | | | | | | | | GO:0005886(100%);GO:0016021 | plasma membrane(100%);integral component of mem m | | | | | IPR005372(100%) | 1.00E+00 | 1.00E+00 | 1.01E+05 | 3.9479 | 0.0011088 | 0.00016581 | 0.9 | 3.8544 | 0.018187 | 0.00097573 | 1 |

| | | | | | | | | | | | | | | | | | | | | | | | | | | | | | | | | | | | | | |
|---|---|---|---|---|---|---|---|---|---|---|---|---|---|---|---|---|---|---|---|---|---|---|---|---|---|---|---|---|---|---|---|---|---|---|---|---|---|
| | | | | | | | | | | | | | | | | | | | | (97%) | brane (97%) | | | | | | | | | | | | | | | | |
| 112 | Variable 2 | SATQLDCGNSGSSMR | Candidatus Koribacter versatilis | Bacteria | Acidobacteria | | Acidobacteriia | | Acidobacteriales | | Acidobacteriaceae | Candidatus Koribacter | | | Candidatus Koribacter versatilis | | | 2.5.1.19 (100%) | 3-phosphoshikimate 1-carboxyvinyltransferase (100%) | GO:0005737 (100%) | cytoplasm (100%) | GO:0003866 (100%) | 3-phosphoshikimate 1-carboxyvinyltransferase activity (100%) | GO:0008652 (100%);GO:0009073 (100%);GO:0009423 (100%) | cellular amino acid biosynthetic process (100%);aromatic amino acid family bios | IPR001986 (100%);IPR006264 (100%);IPR013 | 1.00E+00 | 1.00E+00 | 5.62E+05 | 13.244 | 0.0241 | 0.000010684 | 0.9 | 16.927 | 0.03123 | 0.00031429 | 1 |

| | | | | | | | | | | | | | | | | | | | | | | | | | | | | | | | | | | | | | |
|---|---|---|---|---|---|---|---|---|---|---|---|---|---|---|---|---|---|---|---|---|---|---|---|---|---|---|---|---|---|---|---|---|---|---|---|---|---|
| | | | | | | | | | | | | | | | | | | | | | | | | | ynthetic process (100%);chorismate biosynthetic process (100%) | 792(100%) | | | | | | | | | | | |
| 114 | Variable 200 | VAEVTGQAAGK | Streptomyces | Bacteria | Actinobacteria | | Actinomycetia | | Streptomycetales | | Streptomycetaceae | Streptomyces | | | | | | | | GO:0000502(1 | proteasome complex (100 | GO:0005524(1 | ATP binding (100%);ATPase (100% | GO:0010498(98%);GO:00 | proteasomal protein cata | IPR003593(10 | 5.46E+05 | 1.14E+07 | 1.19E+07 | NS | NS | NS | NS | 0.027437 | 0.0010032 | 0.0032736 | 0.54296875 |

| | | | | | | | | | | | | | | | | | | | | | | | | | | | | | | | | | | | | | | | | |
|---|---|---|---|---|---|---|---|---|---|---|---|---|---|---|---|---|---|---|---|---|---|---|---|---|---|---|---|---|---|---|---|---|---|---|---|---|---|---|---|---|
| | | | | | | | | | | | | | | | | | | | | 00%) | %) | 00%);GO:0016887(100%);GO:0016853(2%) | );isomerase activity (2%) | 19941(98%) | bolic process (98%); modification-dependent protein catabolic process (98%) | 0%);IPR003959(100%);IPR022482(100%) | | | | | | | | | | | | | | |
| 122 | Variab | FWDFLVNGR | Streptom | Bacter | Actinob | | Actinomy | | Streptomyc | | Streptomyc | Strepto | | | Strepto | | | 2.3.1. | Neocarzinostatin | | | GO:001 | acyltransferase act | GO:0017000 | antibiotic bios | IPR001 | 1.00E+ | 1.07E+ | 1.18E+ | 44.323 | 0.0021 | 0.0012 | 1 | 0.0357 | 0.0058 | 0.0004 | 0.6054 | 15.034 | 1.19E- | 0.0000 |

| | | | | | | | | | | | | | | | | | | | | | | | | | | | | | | | | | | | | | | | | |
|---|---|---|---|---|---|---|---|---|---|---|---|---|---|---|---|---|---|---|---|---|---|---|---|---|---|---|---|---|---|---|---|---|---|---|---|---|---|---|---|---|
| | le 208 | | ycescarzinostaticus | ia | acteria | | cetia | | etales | | etaceae | myces | | | mycescarzinostaticus | | | 237(100%) | naphthoate synthase (100%) | | | 6746(100%);GO:0031177(100%) | ivity (100%);phosphopantetheine binding (100%) | (100%) | ynthetic process (100%) | 227(100%);IPR006162(100%);IPR009081(100%) | 00 | 06 | 06 | | 164 | 5 | | 13 | 52 | 4163 | 6875 | | 05 | 9375 |
| 132 | Vari | GLAADDFR | Strept | Bact | Actin | | Actino | | Streptos | | Nocardi | Strep | | | Strep | | | | | | | GO:0 | D-threo-ald | | | IPR0 | 1.00 | 1.00 | 9.20 | 0.38 | 3.88 | 0.000 | 0.9 | 12.2 | 0.00 | 0.00 | 1 | | | |

| | | | | | | | | | | | | | | | | | | | | | | | | | | | | | | | | | | | | | | | | |
|---|---|---|---|---|---|---|---|---|---|---|---|---|---|---|---|---|---|---|---|---|---|---|---|---|---|---|---|---|---|---|---|---|---|---|---|---|---|---|---|---|
| | able 217 | | omonosporaalba | eria | obacteria | | mycetia | | poranagiales | | opsaceae | tomonospora | | | tomonosporaalba | | | | | | | 047834(100%) | ose 1-dehydrogenase activity (100%) | | | 23210(100%);IPR036812(100%) | E+00 | E+00 | E+05 | 818 | 78E-07 | 9623 | | 23 | 37145 | 008235 | | | | |
| 143 | Variable 227 | GALDEMGK | CandidatusProtochlamy | Bacteria | Chlamydiae | | Chlamydiia | | Parachlamydiales | | Parachlamydiaceae | CandidatusProtochla | | | CandidatusProtochla | | | 6.1.1.3(100%) | Threonine--tRNA ligase (100%) | GO:0005737(100%) | cytoplasm(100%) | GO:0000049(100%);G | tRNA binding (100%);threonine-tRNA ligase | GO:0006435(100%) | threonyl-tRNA aminoacylation (100 | IPR002314(100%);IP | 1.00E+00 | 1.00E+00 | 5.10E+05 | 31.506 | 2.2363E-06 | 0.00023482 | 0.9 | 21.852 | 0.0056431 | 0.0012394 | 1 | 9.3698 | 4.32E-04 | 0.0017735 |

| | | | | | | | | | | | | | | | | | | | | | | | | | | | | | | | | | | | | | | | | |
|---|---|---|---|---|---|---|---|---|---|---|---|---|---|---|---|---|---|---|---|---|---|---|---|---|---|---|---|---|---|---|---|---|---|---|---|---|---|---|---|---|
| | | | diaamoebophila | | | | | | | | | mydia | | | mydiaamoebophila | | | | | | | O:0004829(100%);GO:0005524(100%) | activity(100%);ATP binding(100%) | | %) | R002320(100%);IPR004095(100%) | | | | | | | | | | | | | | |
| 145 | Variable 229 | FGSGLLEKPR | Chlorobiaceae | Bacteria | Chlorobi | | Chlorobia | | Chlorobiales | | Chlorobiaceae | | | | | | | 3.6.5.-(100%) | Acting on GTP; involved in cellular | GO:0005737(1 | cytoplasm(100%) | GO:0000287(1 | magnesium ion binding(100%);GTP | GO:0042254(100%) | ribosome biogenesis(100 | IPR006073(10 | 1.00E+00 | 1.00E+00 | 2.24E+05 | 15.664 | 0.00022128 | 0.00053644 | 0.9 | 11.455 | 0.04797 | 0.0008256 | 1 | 6.6277 | 3.15E-03 | 0.00017679 |

| | | | | | | | | | | | | | | | | | | | | | | | | | | | | | | | | | | | | | |
|---|---|---|---|---|---|---|---|---|---|---|---|---|---|---|---|---|---|---|---|---|---|---|---|---|---|---|---|---|---|---|---|---|---|---|---|---|---|
| | | | | | | | | | | | | | | | | | | | and subcellular movement (100%) | 00%) | | 00%);GO:0003924(100%);GO:0005525(100%) | ase activity (100%);GTP binding (100%) | | %) | 0%);IPR006074(100%);IPR006169(100%) | | | | | | | | | | | |
| 150 | Variabl | SNACGDETAMLAWN | Chlorobiu | Bacteri | Chlorob | | Chlorobia | | Chlorobiales | | Chlorobiaceae | Chlorob | | | Chlorob | | Chlorob | 2.7.1.1 | Tetraacyldisaccharide | | | GO:0005 | ATP binding (100% | GO:0009245 (1 | lipid A biosynth | IPR0037 | 1.00E+0 | 1.00E+0 | 1.34E+0 | 3.5635 | 0.01389 | 0 | 0.9 | 4.591 | 0.02911 | 0.00012 | 1 |

| | | | | | | | | | | | | | | | | | | | | | | | | | | | | | | | | | | | | | | | | |
|---|---|---|---|---|---|---|---|---|---|---|---|---|---|---|---|---|---|---|---|---|---|---|---|---|---|---|---|---|---|---|---|---|---|---|---|---|---|---|---|---|
| | e 233 | NR | mphaeobacteroides DSM 266 | a | i | | | | | | | ium | | | ium phaeobacteroides | | ium phaeobacteroides DSM 266 | 30(100%) | 4'-kinase(100%) | | | 524(100%);GO:0009029(100%) | );tetraacyldisaccharide 4'-kinase activity(100%) | 00%) | etic process(100%) | 58(100%);IPR027417(100%) | 0 | 0 | 5 | | | | | | | 973 | | | | |
| 153 | Variable 236 | AGAWFSYENDK | Chloroherpetonthalassium A | Bacteria | Chlorobi | | Chlorobia | | Chlorobiales | | Chlorobiaceae | Chloroherpeton | | | Chloroherpetonthalassi | | Chloroherpetonthalassi | | | GO:0005737(100%) | cytoplasm(100%) | GO:0003684(100%);GO | damaged DNA binding(100%);single-strande | GO:0006281(100%);GO:0006310(100%) | DNA repair(100%);DNA recombina | IPR003593(100%);IPR | 1.00E+00 | 1.99E+05 | 2.43E+05 | NS | NS | NS | NS | 0.032046 | 0.012725 | 0.0013144 | 0.6171875 | 21.789 | 3.23E-07 | 0.0019453 |

| | | | | | | | | | | | | | | | | | | | | | | | | | | | | | | | | | | | | | | | | |
|---|---|---|---|---|---|---|---|---|---|---|---|---|---|---|---|---|---|---|---|---|---|---|---|---|---|---|---|---|---|---|---|---|---|---|---|---|---|---|---|---|
| | | | TCC35110 | | | | | | | | | | | | um | | um ATCC35110 | | | | | :0003697(100%);GO:0005524(100%) | dDNA binding(100%);ATP binding(100%) | ;GO:0009432(100%) | tion(100%);SOS response(100%) | 013765(100%);IPR020584(100%) | | | | | | | | | | | | | | |
| 163 | Variable 245 | VGALSSPLR | Cyanobacteria | Bacteria | Cyanobacteria | | | | | | | | | | | | | 3.6.4.12(100% | DNA helicase (100%) | | | GO:0003677(10 | DNA binding(100%);ATP binding | GO:0006281(100%);GO:0006 | DNA repair(100%);DNA | IPR003593(100 | 1.00E+00 | 2.91E+07 | 4.69E+07 | 6.1467 | 3.334E-10 | 0.0010409 | 1 | 0.043857 | 0.003452 | 0.005007 | 0.66015625 | 5.8833 | 5.59E-03 | 0.00022165 |

| | | | | | | | | | | | | | | | | | | | | | | | | | | | | | | | | | | | | | |
|---|---|---|---|---|---|---|---|---|---|---|---|---|---|---|---|---|---|---|---|---|---|---|---|---|---|---|---|---|---|---|---|---|---|---|---|---|---|
| | | | | | | | | | | | | | | | | | | ) | | | | 0%);GO:0000524(100%);GO:0009378(100%) | (100%);four-way junction helicase activity (100%) | 310(100%);GO:0009432(100%) | recombination (100%);SOS response (100%) | %);IPR004605(100%);IPR008823(100%) | | | | | | | | | | | |
| 165 | Variable | FADDEFADAFAR | Leptolyngb | Bacteria | Cyanobac | | | | Synechococcale | | Leptolyngbyacea | Leptolyn | | | Leptolyn | | | 3.4.11.5 | Prolyl aminopeptidase (10 | GO:00057 | cytoplasm (100% | GO:00041 | aminopeptidase activit | | | IPR00007 | 1.00E+00 | 1.00E+00 | 8.59E+04 | 3.7974 | 0.000238 | 0.0001683 | 0.9 | 3.4448 | 0.021239 | 0.001013 | 1 |

| | | | | | | | | | | | | | | | | | | | | | | | | | | | | | | | | | | | | | | | | |
|---|---|---|---|---|---|---|---|---|---|---|---|---|---|---|---|---|---|---|---|---|---|---|---|---|---|---|---|---|---|---|---|---|---|---|---|---|---|---|---|---|
| | 247 | | yaboryana | | teria | | | | s | | e | gbya | | | gbyaboryana | | | (100%) | 0%) | 37(100%) | ) | 77(100%) | y(100%) | | | 3(100%);IPR0024100(100%);IPR0059444(100%) | | | | | 42 | | | | | 5 | | | | |
| 170 | Variab | CAGSSAQLMAK | Prochlo | Bacter | Cyanob | | | | Synechococ | | Prochloracea | Prochl | | | Prochl | Prochl | Prochl | | | GO:001 | largeribosom | GO:000 | structural const | GO:0006412 | translation(1 | IPR002 | 1.00E+ | 1.00E+ | 2.63E+ | 16.469 | 0.0000 | 0.0002 | 0.9 | 11.139 | 0.0100 | 0.0003 | 1 | 6.7584 | 2.85E- | 0.0005 |

| | | | | | | | | | | | | | | | | | | | | | | | | | | | | | | | | | | | | | | | | |
|---|---|---|---|---|---|---|---|---|---|---|---|---|---|---|---|---|---|---|---|---|---|---|---|---|---|---|---|---|---|---|---|---|---|---|---|---|---|---|---|---|
| | le 251 | | rococcus marinus subsp. marinus str. CCMP1375 | ia | acteria | | | | cales | | e | orococcus | | | orococcus marinus | orococcus marinus subsp. marinus | orococcus marinus subsp. marinus str. CCMP1375 | | | 5934(100%) | al subunit(100%) | 3735(100%);GO:0016740(100%);GO:0019843(100%) | ituent of ribosome(100%);transferase activity(100%);rRNA binding(100%) | (100%) | 00%) | 171(100%);IPR005880(100%);IPR008991(100%) | 00 | 00 | 05 | | 18766 | 7778 | | | 68 | 9302 | | | 03 | 0559 |
| 1 | V | QQ | S | B | C | | | | S | | S | S | | | S | | | 4 | Ph | | | G | ma | G | tri | I | 1 | 1 | 2 | N | N | N | N | 5 | 0 | 0 | 1 | | | |

| | | | | | | | | | | | | | | | | | | | | | | | | | | | | | | | | | | | | |
|---|---|---|---|---|---|---|---|---|---|---|---|---|---|---|---|---|---|---|---|---|---|---|---|---|---|---|---|---|---|---|---|---|---|---|---|---|
| 81 | ariable 261 | TETTGLMR | ynechococcus elongatus | acteria | yanobacteria | | | | ynechococcales | | ynechococcaceae | ynechococcus | | | ynechococcus elongatus | | | .1.1.31(100%) | osphoenolpyruvate carboxylase(100%) | | | O:0000287(100%);GO:0008964(100%) | gnesium ion binding(100%);phosphoenolpyruvate carboxylase activity(100%) | O:0006099(100%);GO:0006107(100%);GO:0015977(100%) | carboxylic acid cycle(100%);oxaloacetate metabolic process(100%);carbon fixation(100% | PR015813(100%);IPR018129(100%);IPR021135(100% | .00E+00 | .00E+00 | .14E+05 | S | S | S | S | .0401 | .039536 | .0018908 |

| | | | | | | | | | | | | | | | | | | | | | | | | | | | | | | | | | | | | | | | | |
|---|---|---|---|---|---|---|---|---|---|---|---|---|---|---|---|---|---|---|---|---|---|---|---|---|---|---|---|---|---|---|---|---|---|---|---|---|---|---|---|---|
| | | | | | | | | | | | | | | | | | | | | | | | | | ) | ) | | | | | | | | | | | | | | |
| 188 | Variable 268 | LRPEPGEGEPGVVAR | Thermus | Bacteria | Deinococcus-Thermus | | Deinococci | | Thermales | | Thermaceae | Thermus | | | | | | 2.1.2.9(100%) | Methionyl-tRNA formyltransferase (100%) | | | GO:0004479(100%) | methionyl-tRNA formyltransferase activity (100%) | | | IPR002376(100%);IPR005793(100%);IPR005794(10 | 1.00E+00 | 1.09E+05 | 2.46E+05 | NS | NS | NS | NS | 0.062095 | 0.0003467 | 0.001069 | 0.6367875 | 19.846 | 8.56E-07 | 0.00085965 |

| | | | | | | | | | | | | | | | | | | | | | | | | | | | | | | | | | | | | | |
|---|---|---|---|---|---|---|---|---|---|---|---|---|---|---|---|---|---|---|---|---|---|---|---|---|---|---|---|---|---|---|---|---|---|---|---|---|---|
| | | | | | | | | | | | | | | | | | | | | | | | | | | 0%) | | | | | | | | | | | |
| 189 | Variable 269 | QDPLVVLEK | Dictyoglomus thermophilum | Bacteria | Dictyoglomi | | Dictyoglomia | | Dictyoglomales | | Dictyoglomaceae | Dictyoglomus | | | Dictyoglomus thermophilum | | | | | GO:0015935(100%) | small ribosomal subunit(100%) | GO:0000049(100%);GO:0003735(100%);GO:00198 | tRNA binding(100%);structural constituent of ribosome(100%);rRNA binding(100%) | GO:0006412(100%) | translation(100%) | IPR000235(100%);IPR005717(100%);IPR020606( | 1.00E+00 | 1.00E+00 | 1.09E+06 | NS | NS | NS | NS | 18.872 | 0.040695 | 0.0001623 | 1 |

| | | | | | | | | | | | | | | | | | | | | | | | | | | | | | | | | | | | | | | | | |
|---|---|---|---|---|---|---|---|---|---|---|---|---|---|---|---|---|---|---|---|---|---|---|---|---|---|---|---|---|---|---|---|---|---|---|---|---|---|---|---|---|
| | | | | | | | | | | | | | | | | | | | | | | 43(100%) | | | | 100%) | | | | | | | | | | | | | | |
| 197 | Variable 276 | QPNGAFSWR | Bacilli | Bacteria | Firmicutes | | Bacilli | | | | | | | | | | | | | GO:0005886(100%);GO:0016021(100%) | plasma membrane(100%);integral component of membrane(100%) | GO:0022857(100%) | transmembrane transporter activity(100%) | | | IPR011701(100%);IPR036259(100%);IPR00 | 1.00E+00 | 1.00E+00 | 8.21E+05 | 94.161 | 4.7868E-07 | 0.00028729 | 0.9 | 43.493 | 0.00204 | 0.00062636 | 1 | 12.107 | 7.05E-05 | 0.004478 |

| | | | | | | | | | | | | | | | | | | | | | | | | | | | | | | | | | | | | | |
|---|---|---|---|---|---|---|---|---|---|---|---|---|---|---|---|---|---|---|---|---|---|---|---|---|---|---|---|---|---|---|---|---|---|---|---|---|---|
| | | | | | | | | | | | | | | | | | | | | | | | | | | 4751(97%) | | | | | | | | | | | |
| 198 | Variable 277 | GSDYYAK | Bacilli | Bacteria | Firmicutes | | Bacilli | | | | | | | | | | | | | GO:0045261(60%);GO:0005886(53%);GO:0 | proton-transporting ATP synthase complex | catalytic core F(1)(60%);plasma membrane(53 | GO:0046933(60%);GO:0005524(53%);GO:0016740(7%) | proton-transporting ATP synthase activity | rotational mechanism(60%);ATP binding(53%);transferase activ | GO:0042777(53%);GO:0009847(33%) | 1.00E+00 | 1.00E+00 | 1.03E+06 | 27.045 | 0.001802 | 0.0017647 | 0.9 | 46.209 | 0.008547 | 0.002989 | 1 |

| | | | | | | | | | | | | | | | | | | | | | | | | | | | | | | | | | | | | | |
|---|---|---|---|---|---|---|---|---|---|---|---|---|---|---|---|---|---|---|---|---|---|---|---|---|---|---|---|---|---|---|---|---|---|---|---|---|---|
| | | | | | | | | | | | | | | | | | | | | 016020(33%) | | %); membrane(33%) | | | ity(7%) | | | | | | | | | | | | |
| 206 | Variable 284 | LENADALYEK | Bacillales | Bacteria | Firmicutes | | Bacilli | | Bacillales | | | | | | | | | | | | | GO:0005525(100%) | GTP binding(100%) | GO:0032297(2%) | negative regulation of DNA-dependent DNA replication initiatio | IPR006073(100%);IPR019988(100% | 1.00E+00 | 1.00E+00 | 9.75E+05 | NS | NS | NS | NS | 20.817 | 0.00056431 | 0.00037618 | 1 |

| | | | | | | | | | | | | | | | | | | | | | | | | | | | | | | | | | | | | | | | | |
|---|---|---|---|---|---|---|---|---|---|---|---|---|---|---|---|---|---|---|---|---|---|---|---|---|---|---|---|---|---|---|---|---|---|---|---|---|---|---|---|---|
| | | | | | | | | | | | | | | | | | | | | | | | | | n (2%) | );IPR027417(100%) | | | | | | | | | | | | | | |
| 207 | Variable 285 | EPSGTQEEAR | Bacillaceae | Bacteria | Firmicutes | | Bacilli | | Bacillales | | Bacillaceae | | | | | | | 4.2.2.2(100%) | Pectate lyase (100%) | GO:0005576(93%) | extracellular region (93%) | GO:0030570(51%);GO:0016829(4 | pectate lyase activity (51%);lyase activity (49%);metal ion binding (2%) | GO:0000272(91%);GO:0045490(2%) | polysaccharide catabolic process (91%);pectin catabolic pr | IPR002022(100%);IPR011050(10 | 1.00E+00 | 1.00E+00 | 4.95E+04 | 0.069387 | 9.0975E-17 | 0.0030225 | 0.9 | 0.63708 | 0.00081286 | 0.0019317 | 1 | 144.17 | 1.55E-19 | 0.017887 |

| | | | | | | | | | | | | | | | | | | | | | | | | | | | | | | | | | | | | | |
|---|---|---|---|---|---|---|---|---|---|---|---|---|---|---|---|---|---|---|---|---|---|---|---|---|---|---|---|---|---|---|---|---|---|---|---|---|---|
| | | | | | | | | | | | | | | | | | | | | | | 9%);GO:0046872(2%) | | | ocess(2%) | 0%);IPR012334(100%) | | | | | | | | | | | |
| 215 | Variable 292 | GYEAMVELMR | Bacillus | Bacteria | Firmicutes | | Bacilli | | Bacillales | | Bacillaceae | Bacillus | | | | | | 2.3.1.204(100%) | Octanoyl-[GcvH]:protein N-octanoyltransferase(100%) | | | GO:0016415(100%);GO:00168 7 | octanoyltransferase activity(100%);ligase activity(15%) | GO:0009107(100%);GO:0009249(100%) | lipoate biosynthetic process(100%);protein lipoylatio | IPR004143(100%);IPR024897( | 1.00E+00 | 1.00E+00 | 1.29E+06 | 0.57132 | 2.9872E-06 | 0.00055529 | 0.9 | 23.142 | 0.0080637 | 0.000086445 | 1 |

| | | | | | | | | | | | | | | | | | | | | | | | | | | | | | | | | | | | | | |
|---|---|---|---|---|---|---|---|---|---|---|---|---|---|---|---|---|---|---|---|---|---|---|---|---|---|---|---|---|---|---|---|---|---|---|---|---|---|
| | | | | | | | | | | | | | | | | | | | | | | 4(15%) | | | n(100%) | 100%) | | | | | | | | | | | |
| 217 | Variable 294 | GLSPVSTEQSPPNPMK | Bacillus | Bacteria | Firmicutes | | Bacilli | | Bacillales | | Bacillaceae | Bacillus | | | | | | 7.1.2.2(100%) | H(+)-transporting two-sector ATPase (100%) | GO:0005737(100%);GO:0030257(100%);GO:00 | cytoplasm(100%);type III protein secretion system complex(100%);proton-transpo | catalytic core F(1)(3%) | GO:0005524(100%);GO:0016887(100%);GO:0046933(100%) | ATP binding(100%);ATPase(100%);proton-transporting ATP synthase activity | rotational mechanism(100%) | GO:0030254(100%);GO:0044780(100%);GO:00719 | 1.00E+00 | 1.00E+00 | 1.89E+05 | 4.5255 | 0.014043 | 0.000060662 | 0.9 | 6.0899 | 0.029043 | 0.00035607 | 1 |

| | | | | | | | | | | | | | | | | | | | | | | | | | | | | | | | | | | | | | |
|---|---|---|---|---|---|---|---|---|---|---|---|---|---|---|---|---|---|---|---|---|---|---|---|---|---|---|---|---|---|---|---|---|---|---|---|---|---|
| | | | | | | | | | | | | | | | | | | | | 45261(3%) | rtingATP synthase complex | | | | | 73(97%) | | | | | | | | | | | |
| 231 | Variable 306 | LGLSDFFPR | Bacillus | Bacteria | Firmicutes | | Bacilli | | Bacillales | | Bacillaceae | Bacillus | | | | | | 3.1.-.-(100%) | Acting on ester bonds (100%) | | | GO:0005524(100%);GO:0008408(100 | ATP binding (100%);3'-5' exonuclease activity (100%);helicase activity (96%) | GO:0006260 (93%) | DNA replication (93%) | IPR001650(100%);IPR006054(100%); | 1.00E+00 | 1.00E+00 | 1.68E+05 | 5.4035 | 0.0038918 | 0 | 0.9 | 5.9739 | 0.0025914 | 0.0013876 | 1 |

| | | | | | | | | | | | | | | | | | | | | | | | | | | | | | | | | | | | | | | | | |
|---|---|---|---|---|---|---|---|---|---|---|---|---|---|---|---|---|---|---|---|---|---|---|---|---|---|---|---|---|---|---|---|---|---|---|---|---|---|---|---|---|
| | | | | | | | | | | | | | | | | | | | | | | %);GO:0004386(96%) | | | | IPR006310(100%) | | | | | | | | | | | | | | |
| 251 | Variable 324 | QNSEAVALLQK | Bacillus | Bacteria | Firmicutes | | Bacilli | | Bacillales | | Bacillaceae | Bacillus | | | | | | | | | | | | | | | 1.00E+00 | 5.09E+05 | 4.33E+04 | NS | NS | NS | NS | 0.002271 | 0.0014019 | 0.0045066 | 0.71484375 | 96.008 | 2.1583E-16 | 0.0012117 |
| 253 | Variable 326 | FDDFDLDVVK | Bacillus | Bacteria | Firmicutes | | Bacilli | | Bacillales | | Bacillaceae | Bacillus | | | | | | | | GO:0005576(100 | extracellular region (100%) | GO:0005102(100 | signaling receptor binding (100%) | GO:0019835(100%);GO:004274 | cytolysis(100%); defense resp | IPR006079(100% | 1.00E+00 | 3.78E+05 | 7.52E+05 | NS | NS | NS | NS | 0.052725 | 0.0059969 | 0.00074455 | 0.65625 | 6.8699 | 0.0026228 | 0.00039315 |

| | | | | | | | | | | | | | | | | | | | | | | | | | | | | | | | | | | | | | | | | |
|---|---|---|---|---|---|---|---|---|---|---|---|---|---|---|---|---|---|---|---|---|---|---|---|---|---|---|---|---|---|---|---|---|---|---|---|---|---|---|---|---|
| | | | | | | | | | | | | | | | | | | | | %) | | %) | | 2 (100%) | onsetobacterium (100%) | ) | | | | | | | | | | | | | | |
| 255 | Variable 328 | DMSPSSVK | Bacillus | Bacteria | Firmicutes | | Bacilli | | Bacillales | | Bacillaceae | Bacillus | | | | | | | | GO:0005886(100%);GO:0016021(100% | plasma membrane(100%);integral component of membrane (100 | | | GO:0030435 (100%) | sporulation resulting in formation of a cellular spore (100%) | IPR029046(100%);IPR025377(6%) | 1.00E+00 | 3.91E+05 | 3.52E+05 | 162.43 | 1.096E-07 | 0.00044402 | 1 | 0.022522 | 0.00086054 | 0.001125 | 0.64453125 | 18.114 | 2.1259E-06 | 0.00044732 |

| | | | | | | | | | | | | | | | | | | | | | | | | | | | | | | | | | | | | | | | | |
|---|---|---|---|---|---|---|---|---|---|---|---|---|---|---|---|---|---|---|---|---|---|---|---|---|---|---|---|---|---|---|---|---|---|---|---|---|---|---|---|---|
| | | | | | | | | | | | | | | | | | | | | ) | %) | | | | | | | | | | | | | | | | | | | |
| 262 | Variable 334 | MDVMSKPLGDEK | Bacillus subtilis group | Bacteria | Firmicutes | | Bacilli | | Bacillales | | Bacillaceae | Bacillus | Bacillus subtilis group | | | | | | | | | GO:0005524(100%);GO:0003677(87%);GO:0016887 | ATP binding(100%);DNA binding(87%);ATPase(87%) | | | IPR003439(100%);IPR003593(100%);IPR017871(1 | 1.00E+00 | 1.00E+00 | 2.58E+05 | 44.917 | 1.1998E-07 | 0.00077877 | 0.9 | 12.406 | 0.0079064 | 0.00082969 | 1 | 14.802 | 1.36E-05 | 0.0041994 |

| | | | | | | | | | | | | | | | | | | | | | | | | | | | | | | | | | | | | | | | | |
|---|---|---|---|---|---|---|---|---|---|---|---|---|---|---|---|---|---|---|---|---|---|---|---|---|---|---|---|---|---|---|---|---|---|---|---|---|---|---|---|---|
| | | | | | | | | | | | | | | | | | | | | | | (87%) | | | | 00%) | | | | | | | | | | | | | | |
| 285 | Variable 355 | QWMDVMGDNPSGFK | Paenibacillusbarcinonensis | Bacteria | Firmicutes | | Bacilli | | Bacillales | | Paenibacillaceae | Paenibacillus | | | Paenibacillusbarcinonensis | | | | | GO:0005886(100%) | plasmamembrane(100%) | GO:0044183(100%) | proteinfoldingchaperone(100%) | GO:0006457(100%) | proteinfolding(100%) | IPR005532(100%);IPR016187(100%);IPR04209 | 1.00E+00 | 1.00E+00 | 3.67E+04 | 0.13548 | 2.9227E-14 | 0.0025357 | 0.9 | 0.65209 | 0.0007971 | 0.0038479 | 1 | 70.489 | 3.7828E-14 | 0.014771 |

| | | | | | | | | | | | | | | | | | | | | | | | | | | | | | | | | | | | | | |
|---|---|---|---|---|---|---|---|---|---|---|---|---|---|---|---|---|---|---|---|---|---|---|---|---|---|---|---|---|---|---|---|---|---|---|---|---|---|
| | | | | | | | | | | | | | | | | | | | | | | | | | | 5(100%) | | | | | | | | | | | |
| 291 | Variable 360 | ESGNLDEMK | Lactobacillaceae | Bacteria | Firmicutes | | Bacilli | | Lactobacillales | | Lactobacillaceae | | | | | | | | | | | GO:0005524(100%);GO:0016887(100%);GO:0 | ATP binding(100%);ATPase(100%);unfolded protein binding(100%) | GO:0006457(100%) | protein folding(100%) | IPR012725(100%);IPR013126(100%);IPR018 | 1.00E+00 | 1.00E+00 | 9.80E+05 | NS | NS | NS | NS | 18.561 | 0.001948 | 0.001159 | 1 |

| | | | | | | | | | | | | | | | | | | | | | | | | | | | | | | | | | | | | | |
|---|---|---|---|---|---|---|---|---|---|---|---|---|---|---|---|---|---|---|---|---|---|---|---|---|---|---|---|---|---|---|---|---|---|---|---|---|---|
| | | | | | | | | | | | | | | | | | | | | | | 0510082(100%) | | | | 181(100%) | | | | | | | | | | | |
| 299 | Variable 368 | LFQQLLVFPK | Streptococcus equi subsp. zooepidemicus | Bacteria | Firmicutes | | Bacilli | | Lactobacillales | | Streptococcaceae | Streptococcus | Streptococcus dysgalactiae group | | Streptococcus equi | Streptococcus equi subsp. zooepide | | 4.2.3.4(100%) | 3-dehydroquinate synthase (100%) | GO:0005737(100%) | cytoplasm(100%) | GO:0000166(100%);GO:0003856(100% | nucleotide binding(100%);3-dehydroquinate synthase activity(100%);metal ion bindin | GO:0008652(100%);GO:0009073(100%);GO:0009423(100%) | cellular amino acid biosynthetic process(100%);aromatic amino acid | IPR016037(100%);IPR030960(100%);I | 1.00E+00 | 1.00E+00 | 1.05E+06 | 23.251 | 0.0092791 | 0 | 0.9 | 35.826 | 0.012894 | 0.001448 | 1 |

| | | | | | | | | | | | | | | | | | | | | | | | | | | | | | | | | | | | | | |
|---|---|---|---|---|---|---|---|---|---|---|---|---|---|---|---|---|---|---|---|---|---|---|---|---|---|---|---|---|---|---|---|---|---|---|---|---|---|
| | | | | | | | | | | | | | | | | micus | | | | | | );GO:0046872(100%) | g(100%) | | family biosynthetic process(100%);chorismate biosynthetic process(100%) | PR030963(100%) | | | | | | | | | | | |
| 308 | Variabl | VPMMEVSSR | Lachnocl | Bacteri | Firmicut | | Clostridia | | Eubacteriale | | Lachnospirace | Lachnocl | | | Lachnocl | | | 2.7.7.1 | Nicotinate-nucleotid | | | GO:0004 | nicotinate-nucleoti | GO:0009435(1 | NAD biosynth | IPR0048 | 1.00E+0 | 1.00E+0 | 2.30E+0 | 7.4038 | 0.00270 | 0 | 0.9 | 7.8382 | 0.02549 | 0.00036 | 1 |

| | | | | | | | | | | | | | | | | | | | | | | | | | | | | | | | | | | | | | | | | |
|---|---|---|---|---|---|---|---|---|---|---|---|---|---|---|---|---|---|---|---|---|---|---|---|---|---|---|---|---|---|---|---|---|---|---|---|---|---|---|---|---|
| | e 376 | | ostridium phytofermentans | a | es | | | | s | | ae | ostridium | | | ostridium phytofermentans | | | 8(100%) | e adenylyltransferase(100%) | | | 515(100%);GO:0005524(100%) | de adenylyltransferase activity(100%);ATP binding(100%) | 00%) | etic process(100%) | 21(100%);IPR005248(100%);IPR014729(100%) | 0 | 0 | 5 | | 41 | | | | 1 | 05 | | | | |
| 317 | Varia | DESQNSVNR | Thermo | Bacte | Nitros | | Thermo | | Thermo | | Thermod | Thermo | | | Thermo | | Thermo | 2.3.1 | Phosphate acy | GO:00 | cytoplasm | GO:00 | phosphate:acyl | GO:0006663 | fatty acid bi | IPR00 | 1.00E | 1.14E | 1.33E | 16.46 | 0.000 | 0 | 1 | 0.030 | 0.000 | 0.002 | 0.539 | 33.61 | 1.919 | 0.000 |

| | | | | | | | | | | | | | | | | | | | | | | | | | | | | | | | | | | | | | | | | |
|---|---|---|---|---|---|---|---|---|---|---|---|---|---|---|---|---|---|---|---|---|---|---|---|---|---|---|---|---|---|---|---|---|---|---|---|---|---|---|---|---|
| | ble 384 | | desulfovibrioyellowstonii DSM 11347 | ria | pirae | | desulfovibrionia | | desulfovibrionales | | esulfovibrionaceae | desulfovibrio | | | desulfovibrioyellowstonii | | desulfovibrioyellowstonii DSM 11347 | .274(100%) | ltransferase(100%) | 057337(100%) | (100%) | 43811(100%) | -[acyl carrier protein] acyltransferase activity(100%) | 3(100%);GO:0008654(100%) | osynthetic process(100%); phospholipid biosynthetic process(100%) | 3664(100%);IPR012281(100%) | +00 | +05 | +05 | 5 | 76619 | | | 852 | 97073 | 6362 | 0625 | 4 | 5E-09 | 68036 |
| 320 | Variable 3 | VAEVFHSVAK | Proteobacteria | Bacteria | Proteobact | | | | | | | | | | | | | 2.1.1.163( | Demethylmenaquinone methy | 4-benzoqui | | | GO:0043333(96%);GO: | 2-octaprenyl-6-methoxy-1 | 4-benzoquinone methy | GO:0032259( | 1.00E+00 | 1.00E+00 | 4.13E+05 | 15.408 | 0.0004551 | 0.0001943 | 0.9 | 15.435 | 0.0072658 | 0.0011958 | 1 | | | |

| | | | | | | | | | | | | | | | | | | | | | | | | | | |
|---|---|---|---|---|---|---|---|---|---|---|---|---|---|---|---|---|---|---|---|---|---|---|---|---|---|---|
| | 87 | | | | eria | | | | | | | | | | | | | 100%);2.1.1.201(100%) | ltransferase(100%);2-methoxy-6-polyprenyl-1 | nolmethylase(100%) | | | 0043770(96%);GO:0102027(96%) | | lase activity(96%);demethylmenaquinone methyltransferase activity(96%);S-adenosylmethi | 100%);GO:0009060(99%);GO:0009234(99%) |

| | | | | | | | | | | | | | | | | | | | | | | | | | | | | | | | | | | | | | |
|---|---|---|---|---|---|---|---|---|---|---|---|---|---|---|---|---|---|---|---|---|---|---|---|---|---|---|---|---|---|---|---|---|---|---|---|---|---|
| | | | | | | | | | | | | | | | | | | | | | | | | | onine: 2-demethylquinol-8 methyltransferase activity (96%) | | | | | | | | | | | | |
| 321 | Variable 388 | MAGHYGNTK | Proteobacteria | Bacteria | Proteobacteria | | | | | | | | | | | | | | | GO:0005840(100 | ribosome (100%) | GO:0003735(100 | structural constituent of ribosome (100% | GO:0006412 (100%) | translation (100%) | IPR000597(100% | 1.00E+00 | 1.00E+00 | 6.21E+05 | 0.42674 | 1.052E-07 | 0.0010699 | 0.9 | 9.8822 | 0.0001708 | 0.0022511 | 1 |

| | | | | | | | | | | | | | | | | | | | | | | | | | | | | | | | | | | | | | | | | |
|---|---|---|---|---|---|---|---|---|---|---|---|---|---|---|---|---|---|---|---|---|---|---|---|---|---|---|---|---|---|---|---|---|---|---|---|---|---|---|---|---|
| | | | | | | | | | | | | | | | | | | | | %) | | %);GO:0019843(1%) | );rRNA binding (1%) | | | );IPR009000(100%);IPR019927(100%) | | | | | | | | | | | | | | |
| 334 | Variable 4 | QYVGPPLADGVK | CandidatusSoliba | Bacteria | Acidobacteria | | Acidobacteriia | | Bryobacterales | | Solibacteraceae | CandidatusSoli | | | CandidatusSoli | | | 6.1.1.20(100 | Phenylalanine--tRNA ligase (100%) | GO:0005737(1 | cytoplasm (100%) | GO:0000049(1 | tRNA binding (100%);magnesiu | GO:0006432 (100%) | phenylalanyl-tRNA ami | IPR002547(10 | 1.00E+00 | 1.00E+00 | 1.88E+06 | 139.54 | 6.4739E-06 | 0.00029059 | 0.9 | 77.776 | 0.00048219 | 0.00086989 | 1 | 7.8761 | 1.25E-03 | 0.00024084 |

| | | | | | | | | | | | | | | | | | | | | | | | | | | | | | | | | | | | | | |
|---|---|---|---|---|---|---|---|---|---|---|---|---|---|---|---|---|---|---|---|---|---|---|---|---|---|---|---|---|---|---|---|---|---|---|---|---|---|
| | | | cterusitatus | | | | | | | | | bacter | | | bacterusitatus | | | %) | | 00%) | | 00%);GO:0000287(100%);GO:0000482 6(100%) | mion binding (100%);phenylalanine-tRNA ligase activity (100%) | | noacylation (100%) | 0%);IPR004532(100%);IPR005121(100%) | | | | | | | | | | | |
| 337 | Variabl | ANVVPGEGGSGGK | Caulobact | Bacteri | Proteob | | Alphaprote | | Caulobacteral | | Caulobacterac | Cauloba | | | | | | 3.4.16. | Serine-type D-Ala | GO:0016 | integral compone | GO:0008 | penicillin binding | GO:0000917(1 | division septu | IPR0014 | 1.00E+0 | 1.00E+0 | 1.72E+0 | 4.742 | 0.00592 | 0 | 0.9 | 6.0205 | 0.01818 | 0 | 1 |

| | | | | | | | | | | | | | | | | | | | | | | | | | | | | | | | | | | | |
|---|---|---|---|---|---|---|---|---|---|---|---|---|---|---|---|---|---|---|---|---|---|---|---|---|---|---|---|---|---|---|---|---|---|---|---|
| | e 401 | | er | a | acteria | | obacteria | | es | | eae | cter | | | | | | 4(100%) | -D-Ala carboxypeptidase(100%) | 021(100%);GO:0005886(13%) | nt of membrane(100%);plasma membrane(13%) | 658(100%);GO:0009002(13%) | (100%);serine-type D-Ala-D-Ala carboxypeptidase activity(13%) | 3%);GO:0008360(13%);GO:0009252(13%) | massembly(13%);regulation of cell shape(13%);peptidoglycan biosynthetic process(13% | 60(100%);IPR005311(100%);IPR012338(100%) | 0 | 0 | 5 | | 81 | | | | 7 |

| | | | | | | | | | | | | | | | | | | | | | | | | | | | | | | | | | | | | | |
|---|---|---|---|---|---|---|---|---|---|---|---|---|---|---|---|---|---|---|---|---|---|---|---|---|---|---|---|---|---|---|---|---|---|---|---|---|---|
| | | | | | | | | | | | | | | | | | | | | | | | | | ) | | | | | | | | | | | | |
| 338 | Variable 402 | QLPETVVNR | Bradyrhizobiaceae | Bacteria | Proteobacteria | | Alphaproteobacteria | | Hyphomicrobiales | | Bradyrhizobiaceae | | | | | | | | | GO:0032300(100%) | mismatch repair complex(100%) | GO:0005524(100%);GO:0016887(100%);GO:0030983 | ATP binding(100%);ATPase(100%);mismatched DNA binding(100%) | GO:0006298(100%) | mismatch repair(100%) | IPR002099(100%);IPR013507(100%);IPR014721(10 | 1.00E+00 | 1.00E+00 | 9.90E+05 | 0.60126 | 2.9209E-06 | 0.00039314 | 0.9 | 15.233 | 0.0015072 | 0.00069849 | 1 |

| | | | | | | | | | | | | | | | | | | | | | | | | | | | | | | | | | | | | | | | | |
|---|---|---|---|---|---|---|---|---|---|---|---|---|---|---|---|---|---|---|---|---|---|---|---|---|---|---|---|---|---|---|---|---|---|---|---|---|---|---|---|---|
| | | | | | | | | | | | | | | | | | | | | | | (100%) | | | | 0%) | | | | | | | | | | | | | | |
| 339 | Variable 403 | VAEADASR | Bradyrhizobium | Bacteria | Proteobacteria | | Alphaproteobacteria | | Hyphomicrobiales | | Bradyrhizobiaceae | Bradyrhizobium | | | | | | 2.7.7.18 (100%) | Nicotinate-nucleotide adenylyltransferase (100%) | | | GO:0004515(100%);GO:0005524(100%) | nicotinate-nucleotide adenylyltransferase activity (100%);ATP binding (100%) | GO:0009435 (100%) | NAD biosynthetic process (100%) | IPR004821(100%);IPR005248(100%);IPR0147 | 2.35E+06 | 1.00E+00 | 1.08E+06 | 0.017767 | 2.4685E-15 | 0.0037978 | 0.785 | 30.977 | 0.019053 | 0 | 1 | 20.612 | 5.7945E-07 | 0.00029708 |

| | | | | | | | | | | | | | | | | | | | | | | | | | | | | | | | | | | | | | | | | |
|---|---|---|---|---|---|---|---|---|---|---|---|---|---|---|---|---|---|---|---|---|---|---|---|---|---|---|---|---|---|---|---|---|---|---|---|---|---|---|---|---|
| | | | | | | | | | | | | | | | | | | | | | | | | | | 29(100%) | | | | | | | | | | | | | | |
| 341 | Variable 405 | TLGGNCGACHQAYR | Rhizobiaceae | Bacteria | Proteobacteria | | Alphaproteobacteria | | Hyphomicrobiales | | Rhizobiaceae | | | | | | | | | GO:0042597(100%) | periplasmic space (100%) | GO:0005506(100%);GO:0009055(100%);GO: | iron ion binding (100%);electron transfer activity (100%);heme binding (100%) | | | IPR002321(100%);IPR010980(100%);IPR01 | 1.00E+00 | 1.00E+00 | 1.95E+05 | 27.018 | 5.2406E-07 | 0.00016096 | 0.9 | 10.151 | 0.029755 | 0.00079446 | 1 | 11.901 | 8.04E-05 | 0.0060988 |

| | | | | | | | | | | | | | | | | | | | | | | | | | | | | | | | | | | | | | | | | |
|---|---|---|---|---|---|---|---|---|---|---|---|---|---|---|---|---|---|---|---|---|---|---|---|---|---|---|---|---|---|---|---|---|---|---|---|---|---|---|---|---|
| | | | | | | | | | | | | | | | | | | | | | | 0020037(100%) | | | | 2127(100%) | | | | | | | | | | | | | | |
| 342 | Variable 406 | CPPGDSAVTGLSGGER | Rhizobiaceae | Bacteria | Proteobacteria | | Alphaproteobacteria | | Hyphomicrobiales | | Rhizobiaceae | | | | | | | 3.6.1.-(100%) | Inphosphorus-containing anhydrides(100%) | GO:0005737(96%) | cytoplasm(96%) | GO:0005524(100%);GO:0000049(98% | ATP binding(100%);tRNA binding(98%);ATPase(98%) | GO:0045900(100%);GO:0006412(93%) | negative regulation of translational elongation(100%);translation | IPR003439(100%);IPR022374(100%); | 1.00E+00 | 1.00E+00 | 4.11E+05 | 34.191 | 0.000022923 | 0.00015257 | 0.9 | 17.32 | 0.012725 | 0.00024485 | 1 | 6.5264 | 3.40E-03 | 0.00069886 |

| | | | | | | | | | | | | | | | | | | | | | | | | | | | | | | | | | | | | | | | | |
|---|---|---|---|---|---|---|---|---|---|---|---|---|---|---|---|---|---|---|---|---|---|---|---|---|---|---|---|---|---|---|---|---|---|---|---|---|---|---|---|---|
| | | | | | | | | | | | | | | | | | | | | | | );GO:0016887(98%) | | | (93%) | IPR027417(100%) | | | | | | | | | | | | | | |
| 344 | Variable 408 | TTLEFPATGDDHLAR | Rhizobiaceae | Bacteria | Proteobacteria | | Alphaproteobacteria | | Hyphomicrobiales | | Rhizobiaceae | | | | | | | | | GO:0032300(98%) | mismatch repair complex(98%) | GO:0005524(98%);GO:0016887(9 | ATP binding(98%);ATPase(98%);mismatched DNA binding(98%) | GO:0006298(100%) | mismatch repair(100%) | IPR002099(100%);IPR013507(10 | 1.00E+00 | 4.87E+04 | 6.68E+04 | 23.25 | 0.002124 | 0.0001 | 1 | 0.034983 | 0.00034767 | 0.0034775 | 0.75390625 | 61.638 | 3.2053E-13 | 0.00091348 |

| | | | | | | | | | | | | | | | | | | | | | | | | | | | | | | | | | | | | | | | | |
|---|---|---|---|---|---|---|---|---|---|---|---|---|---|---|---|---|---|---|---|---|---|---|---|---|---|---|---|---|---|---|---|---|---|---|---|---|---|---|---|---|
| | | | | | | | | | | | | | | | | | | | | | | 8%);GO:0030983(98%) | | | | 0%);IPR014721(100%) | | | | | | | | | | | | | | |
| 346 | Variable 41 | EDALLVEVNPLVR | Actinomycetia | Bacteria | Actinobacteria | | Actinomycetia | | | | | | | | | | | 6.2.1.5(100%) | Succinate--CoA ligase (ADP-forming) (100%) | GO:0005829(0%);GO:0042709( | cytosol(0%);succinate-CoA ligase complex(0%) | GO:0004775(100%);GO:000552 | succinate-CoA ligase (ADP-forming) activity (100%);ATP binding (99 | GO:0006099(100%);GO:0006104(0%) | tricarboxylic acid cycle (100%);succinyl-CoA metab | IPR005809(100%);IPR005811( | 1.00E+00 | 1.70E+05 | 2.45E+05 | NS | NS | NS | NS | 0.037274 | 0.00082024 | 0.002335 | 0.75390625 | 26.332 | 3.8759E-08 | 0.0008939 |

| | | | | | | | | | | | | | | | | | | | | | | | | | | | | | | | | | | | | | |
|---|---|---|---|---|---|---|---|---|---|---|---|---|---|---|---|---|---|---|---|---|---|---|---|---|---|---|---|---|---|---|---|---|---|---|---|---|---|
| | | | | | | | | | | | | | | | | | | | | 0%) | | 4(99%);GO:0000287(99%) | %);magnesium ion binding (99%) | | olic process (0%) | 100%);IPR016102(100%) | | | | | | | | | | | |
| 355 | Variable 418 | CAAVAAEK | Cereibacter sphaeroides | Bacteria | Proteobacteria | | Alphaproteobacteria | | Rhodobacterales | | Rhodobacteraceae | Cereibacter | | | Cereibacter sphaeroides | | | 4.2.1.10(100%) | 3-dehydroquinate dehydratase (100%) | | | GO:0003855(100%) | 3-dehydroquinate dehydratase activity (100%) | GO:0008652(100%);GO:0009073(100%);GO:0009 | cellular amino acid biosynthetic process (100%);a | IPR001874(100%);IPR0185 | 1.00E+00 | 1.00E+00 | 9.46E+05 | NS | NS | NS | NS | 15.171 | 0.00049783 | 0.00015296 | 1 |

| | | | | | | | | | | | | | | | | | | | | | | | | | | |
|---|---|---|---|---|---|---|---|---|---|---|---|---|---|---|---|---|---|---|---|---|---|---|---|---|---|---|
| | | | | | | | | | | | | | | | | | | | | | | | | 423 (100%) | romatic amino acid family biosynthetic process (100%);chorismate biosynthetic process (100 | 09 (100%); IPR036441 (100%) |

| | | | | | | | | | | | | | | | | | | | | | | | | | | | | | | | | | | | | | | | | |
|---|---|---|---|---|---|---|---|---|---|---|---|---|---|---|---|---|---|---|---|---|---|---|---|---|---|---|---|---|---|---|---|---|---|---|---|---|---|---|---|---|
| | | | | | | | | | | | | | | | | | | | | | | | | %) | | | | | | | | | | | | | | | | |
| 363 | Variable 425 | VMLCEQCGAR | Burkholderiales | Bacteria | Proteobacteria | | Betaproteobacteria | | Burkholderiales | | | | | | | | | | | | | GO:0003677(100%) | DNA binding(100%) | GO:0006310(100%);GO:0032196(100%) | DNA recombination(100%);transposition(100%) | IPR029261(100%);IPR002560(94%);IPR032877(94 | 1.00E+00 | 2.41E+05 | 6.52E+04 | 0.43847 | 3.6624E-10 | 0.0081699 | 1 | 0.0075342 | 0.0023704 | 0.003018 | 0.8125 | 102.81 | 6.5905E-17 | 0.011258 |

| | | | | | | | | | | | | | | | | | | | | | | | | | | | | | | | | | | | | | | | | |
|---|---|---|---|---|---|---|---|---|---|---|---|---|---|---|---|---|---|---|---|---|---|---|---|---|---|---|---|---|---|---|---|---|---|---|---|---|---|---|---|---|
| | | | | | | | | | | | | | | | | | | | | | | | | | | %) | | | | | | | | | | | | | | |
| 376 | Variable 437 | HDDMPVSVR | Neisseria | Bacteria | Proteobacteria | | Betaproteobacteria | | Neisseriales | | Neisseriaceae | Neisseria | | | | | | | | GO:0009279(100%);GO:0046930(99%) | cell outer membrane(100%);pore complex(99%) | GO:0015288(100%) | porin activity(100%) | GO:0034220(100%) | ion transmembrane transport(100%) | IPR001702(100%);IPR002299(100%);IPR023614(1 | 1.00E+00 | 1.00E+00 | 3.72E+04 | 0.076459 | 9.0975E-17 | 0.0034777 | 0.9 | 0.48904 | 1.9244E-07 | 0.0038723 | 1 | 228.44 | 2.694E-23 | 0.018275 |

| | | | | | | | | | | | | | | | | | | | | | | | | | | | | | | | | | | | | | | | | |
|---|---|---|---|---|---|---|---|---|---|---|---|---|---|---|---|---|---|---|---|---|---|---|---|---|---|---|---|---|---|---|---|---|---|---|---|---|---|---|---|---|
| | | | | | | | | | | | | | | | | | | | | | | | | | | 00%) | | | | | | | | | | | | | | |
| 386 | Variable 446 | YNLYLGGNK | Gammaproteobacteria | Bacteria | Proteobacteria | | Gammaproteobacteria | | | | | | | | | | | 1.8.1.2(100%) | Assimilatory sulfite reductase (NADPH) (100%) | GO:0009337(100%) | sulfite reductase complex (NADPH) (100%) | GO:0004783(100%);GO:0020037(100%);GO:0046 | sulfite reductase (NADPH) activity (100%);heme binding (100%);metal ion binding (100%) | GO:0019344(100%);GO:0000103(92%);GO:0070814(92%) | cysteine biosynthetic process(100%);sulfate assimilation(92%);hydrogen sulfide bi | IPR005117(100%);IPR006066(100%);IPR006067 | 1.00E+00 | 1.00E+00 | 5.91E+05 | 25.785 | 0.00008803 | 0.000090278 | 0.9 | 22.002 | 0.0013755 | 0.00035303 | 1 | 5.136 | 1.01E-02 | 0.00021218 |

| | | | | | | | | | | | | | | | | | | | | | | | | | | | | | | | | | | | | | | | | |
|---|---|---|---|---|---|---|---|---|---|---|---|---|---|---|---|---|---|---|---|---|---|---|---|---|---|---|---|---|---|---|---|---|---|---|---|---|---|---|---|---|
| | | | | | | | | | | | | | | | | | | | | | | 872(100%) | | | osynthetic process (92%) | (100%) | | | | | | | | | | | | | | |
| 390 | Variable 45 | GFTGDLDSVR | Actinomycetia | Bacteria | Actinobacteria | | Actinomycetia | | | | | | | | | | | | | GO:0005737(1%);GO:0005886(1%);GO | cytoplasm(1%);plasma membrane(1%);excinuclease repair co | GO:0005524(100%);GO:0005525(98% | ATP binding(100%);GTP binding(98%);kinase activity(2%) | GO:0006289(1%);GO:0009432(1%) | nucleotide-excision repair(1%);SOS response(1%) | IPR005337(100%);IPR027417(100%); | 1.00E+00 | 4.83E+05 | 2.53E+05 | 0.21195 | 3.516E-12 | 0.001259 | 1 | 0.01458 | 0.00036081 | 0.0039645 | 0.625 | 40.324 | 1.6839E-10 | 0.013486 |

| | | | | | | | | | | | | | | | | | | | | | | | | | | | | | | | | | | | | | |
|---|---|---|---|---|---|---|---|---|---|---|---|---|---|---|---|---|---|---|---|---|---|---|---|---|---|---|---|---|---|---|---|---|---|---|---|---|---|
| | | | | | | | | | | | | | | | | | | | | :0009380(1%) | mplex(1%) | );GO:0016301(2%) | | | | IPR000305(1%) | | | | | | | | | | | |
| 395 | Variable 454 | YSQGNVSAVGVTYDGHTALTR | Gammaproteobacteria | Bacteria | Proteobacteria | | Gammaproteobacteria | | | | | | | | | | | 3.4.21.50(100%) | Lysylendopeptidase (100%) | GO:0005576(4%);GO:0005615(4%) | extracellular region(4%);extracellular space(4%) | GO:0008233(75%);GO:0016787(17 | peptidase activity(75%);hydrolase activity(17%);serine-type peptidase act | GO:0009405(8%);GO:0006508(4%);GO:0006554(4%) | obsolete pathogenesis(8%);proteolysis(4%);lysine catabolic | IPR009003(100%);IPR043504(100 | 1.00E+00 | 6.17E+05 | 3.03E+06 | 6.8621 | 7.1188E-06 | 0.0013941 | 1 | 0.14527 | 0.034845 | 0.0010892 | 0.87109375 |

| | | | | | | | | | | | | | | | | | | | | | | | | | | | | | | | | | | | | | | | | |
|---|---|---|---|---|---|---|---|---|---|---|---|---|---|---|---|---|---|---|---|---|---|---|---|---|---|---|---|---|---|---|---|---|---|---|---|---|---|---|---|---|
| | | | | | | | | | | | | | | | | | | | | | | %);GO:0008236(8%) | ivity (8%) | | process (4%) | %) | | | | | | | | | | | | | | |
| 396 | Variable 455 | HLVLASGNAGK | Gammaproteobacteria | Bacteria | Proteobacteria | | Gammaproteobacteria | | | | | | | | | | | 3.6.1.66(100%) | XTP/dITP diphosphatase (100%) | GO:0005737(3%);GO:0005829(3% | cytoplasm(3%);cytosol(3%) | GO:0000166(97%);GO:0017111(9 | nucleotide binding(97%);nucleoside-triphosphatase activity(97%);dITP diphos | GO:0009146(100%);GO:0009117(97%);GO:0009143(3%) | purine nucleoside triphosphate catabolic process(100%);nu | IPR002637(100%);IPR020922(10 | 1.00E+00 | 1.00E+00 | 5.50E+05 | 16.761 | 0.0016074 | 0.0026087 | 0.9 | 20.805 | 0.001312 | 0.0030214 | 1 | 4.6326 | 1.52E-02 | 0.0043056 |

| | | | | | | | | | | | | | | | | | | | | | | | | | | | | | | | | | | | | | | | | |
|---|---|---|---|---|---|---|---|---|---|---|---|---|---|---|---|---|---|---|---|---|---|---|---|---|---|---|---|---|---|---|---|---|---|---|---|---|---|---|---|---|
| | | | | | | | | | | | | | | | | | | | | ) | | 7%);GO:0035870(97%) | phatase activity (97%) | | cleotide metabolic process (97%); nucleoside triphosphate catabolic process (3%) | 0%); IPR029001 (100%) | | | | | | | | | | | | | | |
| 403 | Vari | QLLDLAVT | Nitro | Bact | Prot | | Gamm | | Chrom | | Chrom | Nitro | | | Nitro | | | | | | | GO:0 | damaged | GO:0006 | mismat | IPR0 | 1.00 | 1.00 | 4.71 | 37.2 | 0.00 | -0.00 | 0.9 | 20.6 | 0.00 | 0.00 | 1 | 6.93 | 2.51 | 0.00 |

| | | | | | | | | | | | | | | | | | | | | | | | | | | | | | | | | | | | | | | | | |
|---|---|---|---|---|---|---|---|---|---|---|---|---|---|---|---|---|---|---|---|---|---|---|---|---|---|---|---|---|---|---|---|---|---|---|---|---|---|---|---|---|
| | able 461 | H | sococcusoceani | eria | eobacteria | | aproteobacteria | | atiales | | atiaceae | sococcus | | | sococcusoceani | | | | | | | 0036884(100%);GO:0005524(100%);GO:0030983(100% | DNA binding(100%);ATP binding(100%);mismatched DNA binding(100%) | 298(100%) | chrepair(100%) | 00432(100%);IPR005748(100%);IPR007695(100%) | E+00 | E+00 | E+05 | 09 | 0016586 | 0040545 | | 05 | 66656 | 043626 | | 01 | E-03 | 074345 |

| | | | | | | | | | | | | | | | | | | | | | | | | | | | | | | | | | | | | | |
|---|---|---|---|---|---|---|---|---|---|---|---|---|---|---|---|---|---|---|---|---|---|---|---|---|---|---|---|---|---|---|---|---|---|---|---|---|---|
| | | | | | | | | | | | | | | | | | | | | | | ) | | | | | | | | | | | | | | | |
| 408 | Variable 466 | ADLVQQVSR | Enterobacterales | Bacteria | Proteobacteria | | Gammaproteobacteria | | Enterobacterales | | | | | | | | | | | | | GO:0003677(100%) | DNA binding(100%) | | | IPR003477(100%) | 1.00E+00 | 1.00E+00 | 3.09E+05 | NS | NS | NS | NS | 9.634 | 0.047436 | 0.00015385 | 1 |
| 411 | Variable 469 | AASVLVELAK | Enterobacterales | Bacteria | Proteobacteria | | Gammaproteobacteria | | Enterobacterales | | | | | | | | | | | GO:0005829(0%);GO:0005737(0 | cytosol(0%);cytoplasm(0%) | GO:0003700(100%);GO:0003677 | DNA-binding transcription factor activity(100%);DNA binding(5%);ba | GO:0070689(0%);GO:0006355(0%);GO:0045893(0%) | L-threonine catabolic process to propionate(0%);regulati | DNA-templated(0%);positive regulatio | 1.00E+00 | 1.00E+00 | 4.46E+06 | NS | NS | NS | NS | 84.601 | 0.00017088 | 0.00033004 | 1 |

| | | | | | | | | | | | | | | | | | | | | | | | | | | | | | | | | | | | | | | | | |
|---|---|---|---|---|---|---|---|---|---|---|---|---|---|---|---|---|---|---|---|---|---|---|---|---|---|---|---|---|---|---|---|---|---|---|---|---|---|---|---|---|
| | | | | | | | | | | | | | | | | | | | | %) | | (5%);GO:0000986(0%) | cterial-type cis-regulatory region sequence-specific DNA binding (0%) | | on of transcription | n of transcription | | | | | | | | | | | | | | |
| 413 | Variable 470 | AYELFLSEK | Enterobacterales | Bacteria | Proteobacteria | | Gammaproteobacteria | | Enterobacterales | | | | | | | | | 2.3.1.245(100%); | 3-hydroxy-5-phosphonooxypentane-2 | 4-dionethiolase( | GO:0005737(98%) | cytoplasm (98%) | GO:0043332(98%);GO:0016747(98 | fructose-bisphosphate aldolase activity(9 | transferring groups other than a | | 1.00E+00 | 4.43E+05 | 5.51E+05 | 0.59345 | 4.2426E-06 | 0.0014757 | 1 | 0.032039 | 0.015138 | 0.0096771 | 0.58203125 | 14.705 | 0.00014431 | 0.0019397 |

| | | | | | | | | | | | | | | | | | | | | | | | | | | | | | | | | | | | | | |
|---|---|---|---|---|---|---|---|---|---|---|---|---|---|---|---|---|---|---|---|---|---|---|---|---|---|---|---|---|---|---|---|---|---|---|---|---|---|
| | | | | | | | | | | | | | | | | | | 4.1.2.-(0%) | | 100%);Aldehyde-lyases(0%) | | | %);GO:0016829 (1%) | 8%);acyltransferase activity | mino-acyl groups (98%);lyase activity (1%) | | | | | | | | | | | | |
| 415 | Variable 472 | SHGAAHGSWK | Enterobacteriaceae | Bacteria | Proteobacteria | | Gammaproteobacteria | | Enterobacterales | | Enterobacteriaceae | | | | | | | | | GO:0016021(91%);GO:00 | integral component of membrane (91%);plas | | | GO:0006935 (1%);GO:0097588 (1%) | chemotaxis (1%);archaeal or bacterial-type pe | IPR025713(100%);IPR03 | 1.00E+00 | 1.00E+00 | 2.89E+06 | NS | NS | NS | NS | 63.402 | 0.000083694 | 0.00027276 | 1 |

| | | | | | | | | | | | | | | | | | | | | | | | | | | | | | | | | | | | | | | | | |
|---|---|---|---|---|---|---|---|---|---|---|---|---|---|---|---|---|---|---|---|---|---|---|---|---|---|---|---|---|---|---|---|---|---|---|---|---|---|---|---|---|
| | | | | | | | | | | | | | | | | | | | | 05886(90%);GO:0016020(4%) | mamembrane(90%);membrane(4%) | | | | flagellum-dependent cell motility(1%) | 6737(95%);IPR006665(93%) | | | | | | | | | | | | | | |
| 416 | Variable 473 | LTAENGVMSSR | Enterobacteriaceae | Bacteria | Proteobacteria | | Gammaproteobacteria | | Enterobacterales | | Enterobacteriaceae | | | | | | | 2.7.13.3(100%);2.7.3 | Histidine kinase(100%);Phosphotransferases with a nitroge | GO:0005887(95%);GO:0 | integral component of plasma membrane | GO:0000155(97%);GO:0 | phosphorelay sensor kinase activity(97%);ATP binding | GO:0006355(96%);GO:0046777(95%) | regulation of transcription | DNA-templated(96%);prot | 1.00E+00 | 1.00E+00 | 2.67E+05 | 51.391 | 3.8533E-08 | 0.000056896 | 0.9 | 12.357 | 0.0075165 | 0.000047089 | 1 | 17.225 | 3.45E-06 | 0.0061836 |

| | | | | | | | | | | | | | | | | | | | | | | | | | | | | | | | | | | | | | |
|---|---|---|---|---|---|---|---|---|---|---|---|---|---|---|---|---|---|---|---|---|---|---|---|---|---|---|---|---|---|---|---|---|---|---|---|---|---|
| | | | | | | | | | | | | | | | | | | .-(2%) | nous group as acceptor (2%) | 0160021(4%);GO:0005886(1%) | (95%);integral component of membrane (4%);plasma membrane (1%) | 005524(96%);GO:0004673(2%) | (96%);protein histidine kinase activity (2%) | | | ein autophosphorylation (95%) | | | | | | | | | | | |
| 417 | Variable | GAVVLVNFDTDQR | Enterobacteria | Bacteria | Proteobac | | Gammaproteo | | Enterobacterales | | Enterobacteriacea | | | | | | | | | GO:0009927 | cell outer membran | GO:0015477 | fimbrial usher porin activit | GO:0009297 (100%) | pilus assembly (100 | IPR000015 | 1.00E+00 | 1.00E+00 | 2.20E+05 | 6.5701 | 0.0020781 | 0.00005 | 0.9 | 8.3302 | 0.011536 | 0.0002166 | 1 |

| | | | | | | | | | | | | | | | | | | | | | | | | | | | | | | | | | | | | | |
|---|---|---|---|---|---|---|---|---|---|---|---|---|---|---|---|---|---|---|---|---|---|---|---|---|---|---|---|---|---|---|---|---|---|---|---|---|---|
| | 474 | | ceae | | teria | | bacteria | | | | e | | | | | | | | | 9(93%);GO:0016021(93%);GO:0016020(7%) | e(93%);integral component of membrane(93%);membrane(7%) | 3(100%) | y(100%) | ;GO:0006974(0%) | %);cellular response to DNA damage stimulus(0%) | (100%);IPR042186(99%);IPR025949(99%) | | | | | | | | | | 7 | |
| 425 | Variable | LNQGTPLLNAVSK | Enterobacter | Bacteria | Proteoba | | Gammaprote | | Enterobacteral | | Enterobacteriac | | | | | | | 7.6.2.14 | ABC-type aliphatic sul | GO:00058 | plasmamembr | GO:00055 | ATP binding(100%);A | GO:0015074(0%) | DNA integration | IPR02741 | 1.00E+00 | 1.00E+00 | 6.95E+05 | 30.936 | 0.008370 | 0.000520 | 0.9 | 28.461 | 0.038985 | 0.000138 | 1 |

| | | | | | | | | | | | | | | | | | | | | | | | | | | | | | | | | | | | | | |
|---|---|---|---|---|---|---|---|---|---|---|---|---|---|---|---|---|---|---|---|---|---|---|---|---|---|---|---|---|---|---|---|---|---|---|---|---|---|
| | 481 | | iaceae | | cteria | | obacteria | | es | | eae | | | | | | | (95%);3.6.3.-(5%) | fonate transporter(95%);Acting on acid anhydrides; catalyzing transmembrane movement of substances(5%) | 86(89%) | ane(89%) | 24(100%);GO:0016887(89%);GO:0008509(87%) | TPase(89%);anion transmembrane transporter activity(87%) | | (0%) | 7(100%);IPR003439(99%);IPR003593(95%) | | | | | 9 | 83 | | | | 1 | |
| 436 | Varia | ANDFNAWLNQ | Salmon | Bacte | Prote | | Gamma | | Enteroba | | Enterobac | Salmon | | | Salmon | | | 2.4.2 | Lipid IV(A) 4- | GO:00 | integral com | GO:01 | 4-amino-4-de | GO:000649 | protein O- | IPR00 | 1.00E | 1.00E | 1.69E | 7.288 | 0.000 | -0.000 | 0.9 | 7.334 | 0.021 | 0.000 | 1 |

| | | | | | | | | | | | | | | | | | | | | | | | | | | | | | | | | | | | | |
|---|---|---|---|---|---|---|---|---|---|---|---|---|---|---|---|---|---|---|---|---|---|---|---|---|---|---|---|---|---|---|---|---|---|---|---|---|
| | ble 491 | HR | ellaenterica | ria | obacteria | | proteobacteria | | cterales | | teriaceae | ella | | | ellaenterica | | | .43(100%) | amino-4-deoxy-L-arabinosyltransferase(100%) | 16021(94%);GO:0005886(89%) | ponent of membrane(94%);plasma membrane(89%) | 03015(100%);GO:0000030(94%) | oxy-L-arabinose transferase activity(100%);mannosyltransferase activity(94%) | 3(94%);GO:0009245(89%);GO:0009103(72%) | linked glycosylation(94%);lipid A biosynthetic process(89%);lipopolysaccharide biosyntheti | 3342(100%);IPR022839(94%) | +00 | +00 | +05 | 9 | 89611 | 014286 | | | 239 | 17292 |

| | | | | | | | | | | | | | | | | | | | | | | | | | | | | | | | | | | | | | | | | |
|---|---|---|---|---|---|---|---|---|---|---|---|---|---|---|---|---|---|---|---|---|---|---|---|---|---|---|---|---|---|---|---|---|---|---|---|---|---|---|---|---|
| | | | | | | | | | | | | | | | | | | | | | | | | | c process (72%) | | | | | | | | | | | | | | | |
| 437 | Variable 492 | FVTAWLGEDGMVR | Salmonella enterica | Bacteria | Proteobacteria | | Gammaproteobacteria | | Enterobacterales | | Enterobacteriaceae | Salmonella | | | Salmonella enterica | | | | | GO:0042597(100%) | periplasmic space (100%) | GO:0004867(100%) | serine-type endopeptidase inhibitor activity (100%) | | | IPR005658(100%);IPR023084(100%);IPR0 | 1.00E+00 | 1.00E+00 | 1.05E+05 | 14.895 | 2.1861E-07 | 0.00055254 | 0.9 | 4.9257 | 0.034845 | 0.00074912 | 1 | 13.491 | 2.98E-05 | 0.0058613 |

| | | | | | | | | | | | | | | | | | | | | | | | | | | | | | | | | | | | | | |
|---|---|---|---|---|---|---|---|---|---|---|---|---|---|---|---|---|---|---|---|---|---|---|---|---|---|---|---|---|---|---|---|---|---|---|---|---|---|
| | | | | | | | | | | | | | | | | | | | | | | | | | | 27438(100%) | | | | | | | | | | | |
| 462 | Variable 514 | GAGELLLTSVDFEGK | Campylobacter | Bacteria | Proteobacteria | delta/epsilon subdivisions | Epsilonproteobacteria | | Campylobacterales | | Campylobacteraceae | Campylobacter | | | | | | 4.3.2.10(92%);4.1.3.-(8%) | Imidazoleglycerol-phosphate synthase (92%); Oxo-acid-lyases (8%) | GO:0005737(1%) | cytoplasm(1%) | GO:0000107(98%);GO:0016829(73%);GO | imidazoleglycerol-phosphate synthase activity (98%);lyase activity (73%);oxo-acid-lyase act | GO:0000105(100%) | histidine biosynthetic process(100%) | IPR006062(100%);IPR011060(100%);IP | 1.00E+00 | 1.00E+00 | 2.29E+05 | 7.7597 | 0.0047197 | 0.000064286 | 0.9 | 9.8118 | 0.021239 | 0.00012698 | 1 |

| | | | | | | | | | | | | | | | | | | | | | | | | | | | | | | | | | | | | | |
|---|---|---|---|---|---|---|---|---|---|---|---|---|---|---|---|---|---|---|---|---|---|---|---|---|---|---|---|---|---|---|---|---|---|---|---|---|---|
| | | | | | | | | | | | | | | | | | | | | | | :0016833(26%) | ivity(26%) | | | R0137785(100%) | | | | | | | | | | | |
| 474 | Variable 525 | ALALLSVAYK | CandidatusPhytoplasma | Bacteria | Tenericutes | | Mollicutes | | Acholeplasmatales | | Acholeplasmataceae | CandidatusPhytoplasma | | | | | | | | GO:0005840(100%) | ribosome(100%) | GO:0003735(100%);GO:0019843(100 | structural constituent of ribosome(100%);rRNA binding(100%) | GO:0006412(100%) | translation(100%) | IPR002583(100%);IPR036510(100%) | 1.00E+00 | 1.00E+00 | 7.28E+05 | 33.29 | 0.0025382 | 0.0028775 | 0.9 | 32.296 | 0.02768 | 0.0050663 | 1 |

| | | | | | | | | | | | | | | | | | | | | | | | | | | | | | | | | | | | | | |
|---|---|---|---|---|---|---|---|---|---|---|---|---|---|---|---|---|---|---|---|---|---|---|---|---|---|---|---|---|---|---|---|---|---|---|---|---|---|
| | | | | | | | | | | | | | | | | | | | | | | %) | | | | | | | | | | | | | | | |
| 488 | Variable 538 | SSVVNHDFFK | Mycoplasma genitalium G37 | Bacteria | Tenericutes | | Mollicutes | | Mycoplasmatales | | Mycoplasmataceae | Mycoplasma | | | Mycoplasma genitalium | | Mycoplasma genitalium G37 | 6.1.1.20 (100%) | Phenylalanine--tRNA ligase (100%) | GO:0009328 (100%) | phenylalanine-tRNA ligase complex (100%) | GO:0000049(100%);GO:0000287(100%);GO:000482 | tRNA binding (100%);magnesium ion binding (100%);phenylalanine-tRNA ligase activity (100%) | GO:0006432 (100%) | phenylalanyl-tRNA aminoacylation (100%) | IPR002547(100%);IPR004532(100%);IPR005146(1 | 1.00E+00 | 1.00E+00 | 4.94E+05 | 12.206 | 0.0051223 | 0.00010633 | 0.9 | 16.651 | 0.017369 | 0.00031311 | 1 |

| | | | | | | | | | | | | | | | | | | | | | | | | | | | | | | | | | | | | | |
|---|---|---|---|---|---|---|---|---|---|---|---|---|---|---|---|---|---|---|---|---|---|---|---|---|---|---|---|---|---|---|---|---|---|---|---|---|---|
| | | | | | | | | | | | | | | | | | | | | | | 6(100%) | | | | 00%) | | | | | | | | | | | |
| 493 | Variable 542 | FNNLEELLER | MycoplasmagenitaliumG37 | Bacteria | Tenericutes | | Mollicutes | | Mycoplasmatales | | Mycoplasmataceae | Mycoplasma | | | Mycoplasmagenitalium | | Mycoplasmagenitalium G37 | | | | | GO:0005524(100%);GO:0016887(100%) | ATP binding(100%);ATPase(100%) | | | IPR003439(100%);IPR003593(100%);IPR017 | 1.00E+00 | 1.00E+00 | 8.05E+05 | 34.19 | 0.002961 | 0.000513 | 0.9 | 29.047 | 0.01515 | 0 | 1 |

| | | | | | | | | | | | | | | | | | | | | | | | | | | | | | | | | | | | | | | | | |
|---|---|---|---|---|---|---|---|---|---|---|---|---|---|---|---|---|---|---|---|---|---|---|---|---|---|---|---|---|---|---|---|---|---|---|---|---|---|---|---|---|
| | | | | | | | | | | | | | | | | | | | | | | | | | | 871(100%) | | | | | | | | | | | | | | |
| 510 | Variable 558 | LQEVVNHAYTPFSK | Mycoplasmapneumoniae | Bacteria | Tenericutes | | Mollicutes | | Mycoplasmatales | | Mycoplasmataceae | Mycoplasma | | | Mycoplasmapneumoniae | | | 3.5.4.5(100%) | Cytidine deaminase (100%) | | | GO:0004126(100%);GO:0008270(100%);GO | cytidine deaminase activity (100%);zinc ion binding (100%);deoxycytidine deaminase activity (10 | | | IPR002125(100%);IPR006262(100%);IPR0 | 1.00E+00 | 1.00E+00 | 2.07E+05 | 16.596 | 5.6962E-06 | 0.00072875 | 0.9 | 10.433 | 0.035433 | 0.00061682 | 1 | 8.318 | 9.05E-04 | 0.00078695 |

| | | | | | | | | | | | | | | | | | | | | | | | | | | | | | | | | | | | | | | | | |
|---|---|---|---|---|---|---|---|---|---|---|---|---|---|---|---|---|---|---|---|---|---|---|---|---|---|---|---|---|---|---|---|---|---|---|---|---|---|---|---|---|
| | | | | | | | | | | | | | | | | | | | | | | :0047844(100%) | 0%) | | | 16192(100%) | | | | | | | | | | | | | | |
| 512 | Variable 56 | GVEVGDGFAEAR | Corynebacterium | Bacteria | Actinobacteria | | Actinomycetia | | Corynebacteriales | | Corynebacteriaceae | Corynebacterium | | | | | | 4.2.3.5(100%) | Chorismate synthase (100%) | | | GO:0004107(100%) | chorismate synthase activity (100%) | GO:0008652(100%);GO:0009073(100%);GO:0009423(100%) | cellular amino acid biosynthetic process(100%);aromatic amino | IPR000453(100%);IPR020541(100% | 1.00E+00 | 1.00E+00 | 9.75E+04 | 0.19415 | 6.971E-13 | 0.002602 | 0.9 | 1.8496 | 0.047017 | 0.0017334 | 1 | 37.108 | 5.2194E-10 | 0.0011893 |

| | | | | | | | | | | | | | | | | | | | | | | | | | | | | | | | | | | | | | | | | |
|---|---|---|---|---|---|---|---|---|---|---|---|---|---|---|---|---|---|---|---|---|---|---|---|---|---|---|---|---|---|---|---|---|---|---|---|---|---|---|---|---|
| | | | | | | | | | | | | | | | | | | | | | | | | | acid family biosynthetic process (100%);chorismate biosynthetic process (100%) | );IPR035904(100%) | | | | | | | | | | | | | | |
| 517 | Varia | SLTDSDYNTGN | Mycopl | Bacte | Teneri | | Mollicut | | Mycoplas | | Mycoplas | Mycop | | | Mycop | | | | | GO:00 | integral com | | | | | | 1.00E | 2.57E | 5.92E | NS | NS | NS | NS | 0.006 | 0.001 | 0.004 | 0.601 | 42.80 | 7.323 | 0.001 |

| | | | | | | | | | | | | | | | | | | | | | | | | | | | | | | | | | | | | | | | | |
|---|---|---|---|---|---|---|---|---|---|---|---|---|---|---|---|---|---|---|---|---|---|---|---|---|---|---|---|---|---|---|---|---|---|---|---|---|---|---|---|---|
| | ble 564 | GQK | asmapneumoniae | ria | cutes | | es | | matales | | mataceae | lasma | | | lasmapneumoniae | | | | | 16021(100%);GO:0005886(33%) | ponent of membrane(100%);plasma membrane(33%) | | | | | | +00 | +05 | +04 | | | | | 8548 | 1432 | 3746 | 5625 | 4 | 8E-11 | 0231 |
| 527 | Variable 573 | VEGDLKPMYANFGLLPPLK | Thermotoga | Bacteria | Thermotogae | | Thermotogae | | Thermotogales | | Thermotogaceae | Thermotoga | | | | | | 2.1.1.74(100%) | Methylenetetrahydrofolate--tRNA-(uracil(54)- | GO:0005737(91%);G | cytoplasm(91%);cytosol(9%) | GO:0047151(100%); | methylenetetrahydrofolate-tRNA-(uracil- | 10-methylenetetrahydrofolate-dependent | GO:0002098(9%);GO:0008033(9 | tRNA wobble uridine mo | 1.00E+00 | 1.00E+00 | 2.86E+06 | 121.64 | 0.00052751 | 0.00018824 | 0.9 | 140.03 | 0.0034891 | 0.00046935 | 1 | 5.5997 | 6.99E-03 | 0.00029375 |

| | | | | | | | | | | | | | | | | | | | | | | | | | | | | | | | | | | | | | | | | |
|---|---|---|---|---|---|---|---|---|---|---|---|---|---|---|---|---|---|---|---|---|---|---|---|---|---|---|---|---|---|---|---|---|---|---|---|---|---|---|---|---|
| | | | | | | | | | | | | | | | | | | | C(5))-methyltransferas(NAD(P)H-oxidizing)(100%) | O:0005829(9%) | | GO:0050660(100%);GO:0030698(91%) | 5-)-methyltransferase(FADH2-oxidizing)activity(100%);flavinadeninedinucleotidebinding(100%);5 | tRNA(m5U54)methyltransferaseactivity(91%) | %);GO:0030488(9%) | dification(9%);tRNAprocessing(9%);tRNAmethylation(9%) | | | | | | | | | | | | | | |
| 52 | Va | YGAG | Th | Ba | Th | | Th | | The | | The | Th | | | Th | | Th | 2. | DNA- | GO | cyto | GO | DN | GO: | DN | IP | 1. | 1. | 9. | 0. | 3. | 0.0 | 0. | 2. | 0. | 0. | 1 | 15 | 9. | 0. |

| | | | | | | | | | | | | | | | | | | | | | | | | | | | | | | | | | | | | | | | | |
|---|---|---|---|---|---|---|---|---|---|---|---|---|---|---|---|---|---|---|---|---|---|---|---|---|---|---|---|---|---|---|---|---|---|---|---|---|---|---|---|---|
| 8 | riable 574 | YDLPNK | ermotogamaritimaMSB8 | cteria | ermotogae | | ermotogae | | rmotogales | | rmotogaceae | ermotoga | | | ermotogamaritima | | ermotogamaritimaMSB8 | 7.7(100%) | directed DNA polymerase (100%) | :0005737(100%) | plasm(100%) | :0003677(100%);GO:0003887(100%);GO:0008408(10 | A binding (100%);DNA-directed DNA polymerase activity (100%);3'-5' exonuclease activity (100%) | 0006261(100%) | A-dependent DNA replication (100%) | R003141(100%);IPR004013(100%);IPR006054(100%) | 00E+00 | 00E+00 | 92E+04 | 49934 | 388E-09 | 015156 | 9 | 6567 | 017518 | 00033127 | | .382 | 7254E-06 | 0020351 |

| | | | | | | | | | | | | | | | | | | | | | | | | | | | | | | | | | | | | | |
|---|---|---|---|---|---|---|---|---|---|---|---|---|---|---|---|---|---|---|---|---|---|---|---|---|---|---|---|---|---|---|---|---|---|---|---|---|---|
| | | | | | | | | | | | | | | | | | | | | | | 0%) | | | | | | | | | | | | | | | |
| 536 | Variable 6 | EEGATGRPR | Actinomycetia | Bacteria | Actinobacteria | | Actinomycetia | | | | | | | | | | | 4.1.2.-(100%) | Aldehyde-lyases (100%) | | | GO:0016832(100%) | aldehyde-lyase activity (100%) | GO:0005975 (93%) | carbohydrate metabolic process (93%) | IPR005593(100%);IPR009014(100%);IPR018969( | 1.00E+00 | 1.00E+00 | 2.06E+05 | 5.8492 | 0.0047524 | 0.000038462 | 0.9 | 7.8453 | 0.021239 | 0.000025797 | 1 |

| | | | | | | | | | | | | | | | | | | | | | | | | | | | | | | | | | | | | | | | | |
|---|---|---|---|---|---|---|---|---|---|---|---|---|---|---|---|---|---|---|---|---|---|---|---|---|---|---|---|---|---|---|---|---|---|---|---|---|---|---|---|---|
| | | | | | | | | | | | | | | | | | | | | | | | | | | 100%) | | | | | | | | | | | | | | |
| 544 | Variable 67 | DFVDAQQFSK | Mycobacteriaceae | Bacteria | Actinobacteria | | Actinomycetia | | Corynebacteriales | | Mycobacteriaceae | | | | | | | 2.8.1.1(100%) | Thiosulfate sulfurtransferase (100%) | GO:0005618(1%);GO:0005829(1%);GO:0005886 | cell wall (1%);cytosol (1%);plasma membrane (1%) | GO:0004792(97%);GO:0005506(2%);GO:0016670 | thiosulfate sulfurtransferase activity (97%);iron ion binding (2%);oxidoreductase activity | acting on paired donors | with incorporation or reduction of molecular oxygen (2%) | | 1.00E+00 | 2.75E+05 | 5.55E+04 | 21.026 | 0.0027144 | 0.000025 | 1 | 0.0051874 | 0.00066033 | 0.0062269 | 0.72265625 | 78.923 | 5.938E-15 | 0.0004383 |

| | | | | | | | | | | | | | | | | | | | | | | | | | | | | | | | | | | | | | |
|---|---|---|---|---|---|---|---|---|---|---|---|---|---|---|---|---|---|---|---|---|---|---|---|---|---|---|---|---|---|---|---|---|---|---|---|---|---|
| | | | | | | | | | | | | | | | | | | | | (1%) | | 5(2%) | | | | | | | | | | | | | | | |
| 546 | Variable 69 | GPDVDAGGSQHNR | Mycobacterium | Bacteria | Actinobacteria | | Actinomycetia | | Corynebacteriales | | Mycobacteriaceae | Mycobacterium | | | | | | 2.7.11.1(100%) | Non-specific serine/threonine protein kinase(100%) | GO:0016021(97%);GO:0005886(90%) | integral component of membrane(97%);plasma membrane(90%) | GO:0005524(100%);GO:0106310(90%);GO:0106 | ATP binding(100%);protein serine kinase activity(90%);protein threonine kinase activity(90%) | | | IPR005543(100%);IPR000719(97%);IPR011009 | 1.00E+00 | 1.00E+00 | 3.60E+05 | 8.4397 | 0.0056032 | 0 | 0.9 | 11.632 | 0.0077516 | 0.00011487 | 1 |

| | | | | | | | | | | | | | | | | | | | | | | | | | | | | | | | | | | | | | | | | |
|---|---|---|---|---|---|---|---|---|---|---|---|---|---|---|---|---|---|---|---|---|---|---|---|---|---|---|---|---|---|---|---|---|---|---|---|---|---|---|---|---|
| | | | | | | | | | | | | | | | | | | | | | | 311(90%) | | | | (97%) | | | | | | | | | | | | | | |
| 547 | Variable 7 | HFDLVGFDPR | Actinomycetia | Bacteria | Actinobacteria | | Actinomycetia | | | | | | | | | | | 3.4.-.-(56%);3.1.1.-(44%);3.4.14.-(28%) | Acting on peptide bonds (peptidases) (56%); Carboxylic ester hydrolases (44%); Dipeptidyl-peptidases and | GO:0005576(2%);GO:0005618(1%);GO:0005 | extracellular region (2%);cell wall (1%);plasma membrane (1%) | GO:0016787(84%);GO:0008233(16%) | hydrolase activity (84%);peptidase activity (16%) | | | IPR000073(100%);IPR029058(100%);IPR01 | 1.00E+00 | 1.00E+00 | 1.31E+05 | 27.44 | 1.1439E-09 | 0.0069649 | 0.9 | 6.429 | 0.0013228 | 0.0017125 | 1 | 30.444 | 6.74E-09 | 0.0071094 |

| | | | | | | | | | | | | | | | | | | | | | | | | | | | | | | | | | | | | | |
|---|---|---|---|---|---|---|---|---|---|---|---|---|---|---|---|---|---|---|---|---|---|---|---|---|---|---|---|---|---|---|---|---|---|---|---|---|---|
| | | | | | | | | | | | | | | | | | | | tripeptidyl-peptidases (28%) | 886(1%) | | | | | | 3595(19%) | | | | | | | | | | | |
| 548 | Variable 70 | ALTLLMPALR | Mycobacterium | Bacteria | Actinobacteria | | Actinomycetia | | Corynebacteriales | | Mycobacteriaceae | Mycobacterium | | | | | | 3.6.1.66(100%) | XTP/dITP diphosphatase (100%) | | | GO:0000166(100%);GO:0017111(100%);G | nucleotide binding (100%);nucleoside-triphosphatase activity (100%);dITP diphosphatase act | GO:0009117(100%);GO:0009146(100%) | nucleotide metabolic process (100%);purine nucleoside triphosphate ca | IPR002637(100%);IPR020922(100%);IPR | 1.00E+00 | 1.00E+00 | 2.24E+05 | 10.934 | 0.0034662 | 0 | 0.9 | 8.585 | 0.04211 | 0.000033333 | 1 |

| | | | | | | | | | | | | | | | | | | | | | | | | | | | | | | | | | | | | | | | | |
|---|---|---|---|---|---|---|---|---|---|---|---|---|---|---|---|---|---|---|---|---|---|---|---|---|---|---|---|---|---|---|---|---|---|---|---|---|---|---|---|---|
| | | | | | | | | | | | | | | | | | | | | | | O:0035870(100%) | ivity (100%) | | tabolic process (100%) | 0290001(100%) | | | | | | | | | | | | | | |
| 549 | Variable 71 | TVTAVLPHAK | Mycobacterium | Bacteria | Actinobacteria | | Actinomycetia | | Corynebacteriales | | Mycobacteriaceae | Mycobacterium | | | | | | 1.2.-.-(100%);1.-.-.-(43%) | Acting on the aldehyde or oxo group of donors (100%);Oxidoreductases (43%) | GO:0005829(4%);GO:0005886(4%) | cytosol (4%);plasma membrane (4%) | GO:0016705(100%);GO:0004497(4 | oxidoreductase activity | acting on paired donors | within corporation or reduction of molecular oxygen (100%);m | GO:0006629(19%) | 1.00E+00 | 1.00E+00 | 7.66E+05 | 116.75 | 2.6251E-07 | 0.00021685 | 0.9 | 44.35 | 0.015138 | 0.0010365 | 1 | 12.865 | 4.38E-05 | 0.0035895 |

| | | | | | | | | | | | | | | | | | | | | | | | | | | | | | | | | | | | | | |
|---|---|---|---|---|---|---|---|---|---|---|---|---|---|---|---|---|---|---|---|---|---|---|---|---|---|---|---|---|---|---|---|---|---|---|---|---|---|
| | | | | | | | | | | | | | | | | | | | | | | %) | | | onooxygenase activity(4%) | | | | | | | | | | | | |
| 551 | Variable 73 | APDHFLVGR | Mycobacterium | Bacteria | Actinobacteria | | Actinomycetia | | Corynebacteriales | | Mycobacteriaceae | Mycobacterium | | | | | | | | GO:0005618(100%);GO:0005886(100% | cell wall(100%);plasma membrane(100%) | | | | | IPR016709(100%);IPR029069(100%);I | 1.00E+00 | 1.00E+00 | 6.10E+05 | 13.204 | 0.02708 | 0.000035714 | 0.9 | 18.478 | 0.034296 | 0.000033333 | 1 |

| | | | | | | | | | | | | | | | | | | | | | | | | | | | | | | | | | | | | | | | | |
|---|---|---|---|---|---|---|---|---|---|---|---|---|---|---|---|---|---|---|---|---|---|---|---|---|---|---|---|---|---|---|---|---|---|---|---|---|---|---|---|---|
| | | | | | | | | | | | | | | | | | | | | ) | | | | | | PR039569(100%) | | | | | | | | | | | | | | |
| 558 | Variable 8 | ATDLMSER | Actinomycetia | Bacteria | Actinobacteria | | Actinomycetia | | | | | | | | | | | 6.3.4.4(100%) | Adenylosuccinate synthase (100%) | GO:0005737(93%) | cytoplasm(93%) | GO:0004019(100%);GO:0005525(10 | adenylosuccinate synthase activity (100%);GTP binding (100%);magnesium ion bindin | GO:0044208(93%);GO:0006164(7%) | 'de novo' AMP biosynthetic process (93%);purine nucleotid | IPR001114(100%);IPR027417(100% | 1.00E+00 | 1.00E+00 | 7.97E+05 | 100.54 | 0.000016918 | 0 | 0.9 | 36.491 | 0.0097842 | 0.00043452 | 1 | 6.8396 | 2.68E-03 | 0.00098601 |

| | | | | | | | | | | | | | | | | | | | | | | | | | | | | | | | | | | | | | | | | |
|---|---|---|---|---|---|---|---|---|---|---|---|---|---|---|---|---|---|---|---|---|---|---|---|---|---|---|---|---|---|---|---|---|---|---|---|---|---|---|---|---|
| | | | | | | | | | | | | | | | | | | | | | | 0%);GO:0000287(93%) | g(93%) | | ebiosynthetic process(7%) | );IPR042109(100%) | | | | | | | | | | | | | | |
| 560 | Variable 81 | VGHPDAAR | Mycobacterium | Bacteria | Actinobacteria | | Actinomycetia | | Corynebacteriales | | Mycobacteriaceae | Mycobacterium | | | | | | 2.4.1.227(100%) | Undecaprenyldiphospho-muramoylpentapeptide beta-N-acetylglucosaminyl | GO:0005886(96%);GO:0005887 | plasma membrane(96%);integral component of plas | GO:0050511(99%);GO:0051991 | undecaprenyldiphospho-muramoylpentapeptide beta-N-acetylglucosaminy | 6-diaminopimelyl-D-alanyl-D-alanine-diphosphoundecaprenol | GO:0005975(99%);GO:0030259(99%);GO:000836 | carbohydrate metabolic process(99% | 1.00E+00 | 8.49E+04 | 3.78E+04 | NS | NS | NS | NS | 0.011602 | 0.00017464 | 0.004465 | 0.7734375 | 93.612 | 3.3323E-16 | 0.0005032 |

| | | | | | | | | | | | | | | | | | | | | | | | | | | | | | | | | | | | | | |
|---|---|---|---|---|---|---|---|---|---|---|---|---|---|---|---|---|---|---|---|---|---|---|---|---|---|---|---|---|---|---|---|---|---|---|---|---|---|
| | | | | | | | | | | | | | | | | | | | transferase (100%) | (1%); GO:0032153 (1%) | mamembrane (1%);cell division site (1%) | (95%); GO:0005524 (1%) | ltransferase activity (99%);UDP-N-acetyl-D-glucosamine:N-acetylmuramoyl-L-alanyl-D-glutamyl-meso-2 | 4-beta-N-acetylglucosaminlytransferase activity (95%);ATP binding (1%) | 0 (98%) | ); lipid glycosylation (99%); regulation of cell shape (98%) | | | | | | | | | | | |
| 568 | Vari | TLGSQVGK | Myco | Bact | Actin | | Actino | | Coryne | | Mycobac | Myco | | | Myco | | | | | GO:0 | plasma | GO:0 | P-P-bond- | GO:0006 | protein | IPR0 | 1.00 | 1.00 | 1.88 | 4.65 | 0.00 | 0.000 | 0.9 | 6.17 | 0.00 | 0.00 | 1 |

| | | | | | | | | | | | | | | | | | | | | | | | | | | | | | | | | | | | | |
|---|---|---|---|---|---|---|---|---|---|---|---|---|---|---|---|---|---|---|---|---|---|---|---|---|---|---|---|---|---|---|---|---|---|---|---|---|
| | able 89 | | bacteriumleprae | eria | obacteria | | mycetia | | bacteriales | | teriaceae | bacterium | | | bacterium leprae | | | | | 0058886(100%);GO:0016021(100%) | membrane(100%);integral component of membrane(100%) | 015450(100%) | hydrolysis-driven protein transmembrane transporter activity(100%) | 605(100%);GO:0043952(100%);GO:0065002(100%) | targeting(100%);protein transport by the Sec complex(100%);intracellular protein transmem | 05665(100%);IPR022645(100%);IPR022646(100%) | E+00 | E+00 | E+05 | 66 | 23745 | 069048 | | 62 | 45827 | 08295 |

| | | | | | | | | | | | | | | | | | | | | | | | | | | | | | | | | | | | | | | | | |
|---|---|---|---|---|---|---|---|---|---|---|---|---|---|---|---|---|---|---|---|---|---|---|---|---|---|---|---|---|---|---|---|---|---|---|---|---|---|---|---|---|
| | | | | | | | | | | | | | | | | | | | | | | | | | brane transport(100%) | | | | | | | | | | | | | | | |
| 569 | Variable 9 | HAQWEAER | Actinomycetia | Bacteria | Actinobacteria | | Actinomycetia | | | | | | | | | | | | | GO:0005737(98%) | cytoplasm(98%) | GO:0005524(100%);GO:0016887(98%);G | ATP binding(100%);ATPase(98%);peptidase activity(21%) | GO:0009408(94%);GO:0042026(94%) | response to heat(94%);protein refolding(94%) | IPR027417(100%);IPR003593(99%);IPR | 1.00E+00 | 1.00E+00 | 1.29E+05 | 9.675 | 2.107E-06 | 0.00028142 | 0.9 | 5.7759 | 0.01403 | 0.0005387 | 1 | 9.8546 | 3.10E-04 | 0.0025428 |

| | | | | | | | | | | | | | | | | | | | | | | | | | | | | | | | | | | | | | | | | |
|---|---|---|---|---|---|---|---|---|---|---|---|---|---|---|---|---|---|---|---|---|---|---|---|---|---|---|---|---|---|---|---|---|---|---|---|---|---|---|---|---|
| | | | | | | | | | | | | | | | | | | | | | | O:0008233(21%) | | | | 003959(99%) | | | | | | | | | | | | | | |
| 570 | Variable 90 | SRPVFEDASAR | Mycobacterium leprae | Bacteria | Actinobacteria | | Actinomycetia | | Corynebacteriales | | Mycobacteriaceae | Mycobacterium | | | Mycobacterium leprae | | | | | | | | | | | | 1.00E+00 | 1.00E+00 | 2.65E+04 | 0.26276 | 1.5094E-12 | 0.0033605 | 0.9 | 0.62839 | 0.000081286 | 0.0037591 | 1 | 53.923 | 2.5103E-12 | 0.012371 |
| 572 | Variable 92 | GDSNVDWLHR | Mycobacterium aviu | Bacteria | Actinobacteria | | Actinomycetia | | Corynebacteriales | | Mycobacteriaceae | Mycobacterium | Mycobacterium | | | | | 2.1.1.45(100%) | Thymidylate synthase (100%) | GO:0005737(10 | cytoplasm(100%) | GO:0004799(10 | thymidylate synthase activity (100%) | GO:0006231(100%);GO:0006 | dTMP biosynthetic proces | IPR000398(100 | 1.00E+00 | 1.00E+00 | 2.99E+05 | 7.6806 | 0.018975 | 0.00019141 | 0.9 | 9.5015 | 0.034845 | 0.00017077 | 1 | | | |

| | | | | | | | | | | | | | | | | | | | | | | | | | | | | | | | | | | | | | | | | |
|---|---|---|---|---|---|---|---|---|---|---|---|---|---|---|---|---|---|---|---|---|---|---|---|---|---|---|---|---|---|---|---|---|---|---|---|---|---|---|---|---|
| | | | mcomplex(MAC) | | | | | | | | | avium complex(MAC) | | | | | | ) | | 0%) | | 0%) | | 235(100%);GO:0032259(100%) | s(100%);dTTP biosynthetic process(100%);methylation(100%) | %);IPR020940(100%);IPR023451(100%) | | | | | | | | | | | | | | |
| 577 | Variable 9 | LLGAKPGAR | Mycobacterium | Bacteria | Actinobacte | | Actinomycetia | | Corynebacteriales | | Mycobacteriaceae | Mycobacteriu | Mycobacteri | | | | | 3.1.11.5(1 | Exodeoxyribonuclease V(10 | GO:0009338 | exodeoxyribonuclease | GO:0008854 | exodeoxyribonuclease V act | GO:0000724(81%);G | double-strand break re | IPR027417( | 1.00E+00 | 1.00E+00 | 2.93E+05 | 14.928 | 0.0010493 | 0.0011275 | 0.9 | 11.525 | 0.0051813 | 0.00062151 | 1 | 5.0548 | 1.08E-02 | 0.0004065 |

| | | | | | | | | | | | | | | | | | | | | | | | | | | |
|---|---|---|---|---|---|---|---|---|---|---|---|---|---|---|---|---|---|---|---|---|---|---|---|---|---|---|
| | 7 | | tuberculosis complex | | ria | | | | | | | m | um tuberculosis complex | | | | | 00%) | 0%) | (94%);GO:0005618(6%);GO:0005829(6%) | V complex(94%);cell wall(6%);cytosol(6%) | (100%);GO:0005524(88%);GO:0003677(81%) | ivity(100%);ATP binding(88%);DNA binding(81%) | O:0006310(19%);GO:0006302(13%) | pair via homologous recombination(81%);DNA recombination(19%);double-strand brea | 100%);IPR006344(94%);IPR027785(88%) |

| | | | | | | | | | | | | | | | | | | | | | | | | | | | | | | | | | | | | | | | | |
|---|---|---|---|---|---|---|---|---|---|---|---|---|---|---|---|---|---|---|---|---|---|---|---|---|---|---|---|---|---|---|---|---|---|---|---|---|---|---|---|---|
| | | | | | | | | | | | | | | | | | | | | | | | | | krepair (13%) | | | | | | | | | | | | | | | |
| 578 | Variable 98 | FGEAGFQSVVDK | Mycobacterium tuberculosis complex | Bacteria | Actinobacteria | | Actinomycetia | | Corynebacteriales | | Mycobacteriaceae | Mycobacterium | Mycobacterium tuberculosis complex | | | | | | | GO:0005886(100%) | plasma membrane (100%) | | | | | IPR012312(100%) | 1.00E+00 | 1.00E+00 | 1.48E+05 | 8.0053 | 0.00015379 | 0.00008844 | 0.9 | 6.0058 | 0.034845 | 0.00052254 | 1 | 4.736 | 1.40E-02 | 0.00027092 |
| 2 | Vari | AQSAADDA | Actino | Bact | Actin | | Actino | | | | | | | | | | | | | GO:0 | integralco | GO:0 | transmemb | | | IPR0 | 1.00 | 1.00 | 8.67 | 0.35 | 4.85 | 0.001 | 0.9 | NS | NS | NS | NS | 17.3 | 3.25 | 0.00 |

| | | | | | | | | | | | | | | | | | | | | | | | | | | | | | | | | | | | | | | | | |
|---|---|---|---|---|---|---|---|---|---|---|---|---|---|---|---|---|---|---|---|---|---|---|---|---|---|---|---|---|---|---|---|---|---|---|---|---|---|---|---|---|
| | able 10 | R | mycetia | eria | obacteria | | mycetia | | | | | | | | | | | | | 016021(64%);GO:0005737(4%) | mponent of membrane (64%);cytoplasm (4%) | 022857(60%);GO:0005524(36%);GO:0016887(36%) | rane transporter activity (60%);ATP binding (36%);ATPase (36%) | | | 11701(60%);IPR036259(60%);IPR000641(36%) | E+00 | E+00 | E+04 | 383 | 34E-10 | 4314 | | | | | | 32 | E-06 | 098583 |
| 3 | Va | VPYS | My | Ba | Ac | | Act | | Cor | | Myc | My | My | | | | | | | | | | | | | | 1. | 1. | 5. | 11 | 3. | 0.0 | 0. | NS | NS | NS | NS | 9. | 5. | 0. |

| | | | | | | | | | | | | | | | | | | | | | | | | | | | | | | | | | | | | | | | | |
|---|---|---|---|---|---|---|---|---|---|---|---|---|---|---|---|---|---|---|---|---|---|---|---|---|---|---|---|---|---|---|---|---|---|---|---|---|---|---|---|---|
| | riable 100 | AESLVR | cobacterium tuberculosis complex | cteria | tinobacteria | | inomycetia | | ynebacteriales | | obacteriaceae | cobacterium | cobacterium tuberculosis complex | | | | | | | | | | | | | | 00E+00 | 00E+00 | 70E+04 | .133 | 6717E-06 | 0027987 | 9 | | | | | 0454 | 41E-04 | 00073623 |
| 4 | Variable 101 | MNVSAESGAPR | Mycobacterium tuberc | Bacteria | Actinobacteria | | Actinomycetia | | Corynebacteriales | | Mycobacteriaceae | Mycobacterium | Mycobacterium t | | | | | 2.3.1.-(83%);2.3 | Transferring groups other than amino-acy | | | GO:0016747(93% | acyltransferase activity | transferring groups other than amino | GO:0006633(100%);GO:0009 | fatty acid biosynthetic | 1.00E+00 | 1.00E+00 | 3.93E+04 | NS | NS | NS | NS | NS | NS | NS | NS | | | |

| | | | | | | | | | | | | | | | | | | | | | | | | | | |
|---|---|---|---|---|---|---|---|---|---|---|---|---|---|---|---|---|---|---|---|---|---|---|---|---|---|---|
| | | | ulosis complex | | | | | | | | | | uberculosis complex | | | | | .1.74 (17%) | l groups (83%); Chalcone synthase (17%) | | | ); GO:0016210 (7%) | | -acyl groups (93%);naringenin-chalcone synthase activity (7%) | 715 (7%); GO:0030639 (7%) | process (100%); chalcone biosynthetic process (7%); polyketid |

| | | | | | | | | | | | | | | | | | | | | | | | | | | | | | | | | | | | | | |
|---|---|---|---|---|---|---|---|---|---|---|---|---|---|---|---|---|---|---|---|---|---|---|---|---|---|---|---|---|---|---|---|---|---|---|---|---|---|
| | | | | | | | | | | | | | | | | | | | | | | | | | | ebiosynthetic process(7%) | | | | | | | | | | | |
| 5 | Variable 102 | EVNAEQFADR | Mycobacterium tuberculosis complex | Bacteria | Actinobacteria | | Actinomycetia | | Corynebacteriales | | Mycobacteriaceae | Mycobacterium | Mycobacterium tuberculosis | | | | | 3.6.3.31(40%);7.3.2.5(40% | Transferred entry: 7.6.2.11(40%);ABC-type molybdate transporter(40 | GO:0016020(73%);GO:0005588 | membrane(73%);plasmamembrane(13%) | GO:0005524(100%);GO:00168 | ATP binding(100%);ATPase(93%);molybdenum ion transmembrane | GO:0015689(67%);GO:0055085(7%) | molybdate ion transport(67%);transmembrane tra | IPR003439(100%);IPR027417 | 1.00E+00 | 1.00E+00 | 2.61E+05 | 1.5321 | 0.01805 | 0.00028595 | 0.9 | NS | NS | NS | NS |

| | | | | | | | | | | | | | | | | | | | | | | | | | | | | | | | | | | | | | |
|---|---|---|---|---|---|---|---|---|---|---|---|---|---|---|---|---|---|---|---|---|---|---|---|---|---|---|---|---|---|---|---|---|---|---|---|---|---|
| | | | | | | | | | | | | | complex | | | | | );3.6.3.30(20%) | %);Transferred entry:7.2.2.7(20%) | 6(13%) | | 87(93%);GO:0042888(87%) | transporter activity (87%) | | nsport(7%) | (100%);IPR003593(93%) | | | | | | | | | | | |
| 6 | Variable 103 | QCVTAAEHR | Mycobacterium tuberculosis com | Bacteria | Actinobacteria | | Actinomycetia | | Corynebacteriales | | Mycobacteriaceae | Mycobacterium | Mycobacterium tubercul | | | | | | | GO:0005576(100%);GO:00 | extracellular region(100%);cell wall(100%);plasm | | | GO:0009405(100%) | obsolete pathogenesis(100%) | IPR000084(100%) | 1.00E+00 | 1.00E+00 | 9.69E+04 | NS | NS | NS | NS | NS | NS | NS | NS |

| | | | | | | | | | | | | | | | | | | | | | | | | | | | | | | | | | | | | | |
|---|---|---|---|---|---|---|---|---|---|---|---|---|---|---|---|---|---|---|---|---|---|---|---|---|---|---|---|---|---|---|---|---|---|---|---|---|---|
| | | | plex | | | | | | | | | | osis complex | | | | | | | 05618(100%);GO:0005886(100%) | a membrane(100%) | | | | | | | | | | | | | | | | |
| 7 | Variable 104 | AGDGDAVLGLLR | Mycobacterium tuberculo | Bacteria | Actinobacteria | | Actinomycetia | | Corynebacteriales | | Mycobacteriaceae | Mycobacterium | Mycobacterium tub | | | | | 3.1.11.5(100%) | Exodeoxyribonuclease V (100%) | GO:0009338(86%);G | exodeoxyribonuclease V complex (8 | GO:0008854(100%); | exodeoxyribonuclease V activity (100%);ATP | GO:0000724(67%);GO:0006310(2 | double-strand break repair via hom | IPR027417(100%);I | 1.00E+00 | 1.00E+00 | 1.13E+05 | 8.6226 | 0.00057258 | 0.00016721 | 0.9 | NS | NS | NS | NS |

| | | | | | | | | | | | | | | | | | | | | | | | | | | |
|---|---|---|---|---|---|---|---|---|---|---|---|---|---|---|---|---|---|---|---|---|---|---|---|---|---|---|
| | | | sis complex | | | | | | | | | | erculosis complex | | | | | | | O:0005618(5%);GO:0005829(5%) | 6%);cell wall(5%);cytosol(5%) | GO:0005524(81%);GO:0003677(67%) | binding(81%);DNA binding(67%) | 4%);GO:0006302(19%) | ologous recombination(67%);DNA recombination(24%);double-strand break repair(19 | PR006344(86%);IPR027785(86%) |

| | | | | | | | | | | | | | | | | | | | | | | | | | | | | | | | | | | | | | |
|---|---|---|---|---|---|---|---|---|---|---|---|---|---|---|---|---|---|---|---|---|---|---|---|---|---|---|---|---|---|---|---|---|---|---|---|---|---|
| | | | | | | | | | | | | | | | | | | | | | | | | | %) | | | | | | | | | | | | |
| 8 | Variable 105 | GAAQFDAGESVTLAQTGLR | Mycobacterium tuberculosis complex | Bacteria | Actinobacteria | | Actinomycetia | | Corynebacteriales | | Mycobacteriaceae | Mycobacterium | Mycobacterium tuberculosis complex | | | | | 2.3.1.41(75%);2.3.1.292(25%) | Beta-ketoacyl-[acyl-carrier-protein] synthase I (75%);(Phenol)carboxyphthiodiolenone synthase (25%) | GO:0034081(11%);GO:0005829(5%);GO:0005886(5 | polyketide synthase complex(11%);cytosol(5%);plasma membrane (5%) | GO:0016491(100%);GO:0031177(95%);GO:0004315 | oxidoreductase activity(100%);phosphopantetheine binding(95%);3-oxoacyl-[acyl-carrier-protein] synthase activity | GO:0006633(68%);GO:0071766(11%);GO:0097040(11%) | fatty acid biosynthetic process(68%);Actinobacterium-type cell wall biogenesis(11% | IPR011032(100%);IPR013154(100%);IPR013968(1 | 1.00E+00 | 1.00E+00 | 1.23E+05 | NS | NS | NS | NS | NS | NS | NS | NS |

| | | | | | | | | | | | | | | | | | | | | | | | | | | | | | | | | | | | | | |
|---|---|---|---|---|---|---|---|---|---|---|---|---|---|---|---|---|---|---|---|---|---|---|---|---|---|---|---|---|---|---|---|---|---|---|---|---|---|
| | | | | | | | | | | | | | | | | | | | | %) | | (84%) | (84%) | | ); phthiocerol biosynthetic process (11%) | 00%) | | | | | | | | | | | |
| 9 | Variable 106 | APSQADGR | Mycobacterium tuberculosis complex | Bacteria | Actinobacteria | | Actinomycetia | | Corynebacteriales | | Mycobacteriaceae | Mycobacterium | Mycobacterium tuberculosi | | | | | 4.6.1.1(100%) | Adenylate cyclase (100%) | GO:0016021(100%);GO:0005 | integral component of membrane (100%);plasmam | GO:0004016(27%);GO:00168 | adenylate cyclase activity (27%);lyase activity (9%) | GO:0035556(95%);GO:0009190(91%);GO:000617 | intracellular signal transduction (95%);cyclic nucleotid | IPR003660(100%);IPR00105 | 1.00E+00 | 1.00E+00 | 6.06E+04 | 3.4599 | 0.0090362 | 0 | 0.9 | NS | NS | NS | NS |

| | | | | | | | | | | | | | | | | | | | | | | | | | | | | | | | | | | | | | |
|---|---|---|---|---|---|---|---|---|---|---|---|---|---|---|---|---|---|---|---|---|---|---|---|---|---|---|---|---|---|---|---|---|---|---|---|---|---|
| | | | x | | | | | | | | | | scomplex | | | | | | | 886(14%);GO:0005576(5%) | embrane(14%);extracellular region(5%) | 29(9%) | | 1(5%) | ebiosynthetic process(91%);cAMP biosynthetic process(5%) | 4(96%);IPR029787(96%) | | | | | | | | | | | |
| 11 | Variable 108 | ELPEMDPSGAGK | Mycobacterium tub | Bacteria | Actinobacteria | | Actinomycetia | | Corynebacteriales | | Mycobacteriaceae | Mycobacterium | Mycobacterium | | | | | 2.3.1.-(100%) | Transferring groups other than amino | GO:0005618(4 | cell wall(4%);cytosol(4% | GO:0016491(9 | oxidoreductase activity(96%);phosp | GO:0006633(70%);GO:00 | fatty acid biosynthetic proc | IPR014043(10 | 1.00E+00 | 1.00E+00 | 2.41E+05 | NS | NS | NS | NS | NS | NS | NS | NS |

| | | | | | | | | | | | | | | | | | | | | | | | | | | | | | | | | | | | | | |
|---|---|---|---|---|---|---|---|---|---|---|---|---|---|---|---|---|---|---|---|---|---|---|---|---|---|---|---|---|---|---|---|---|---|---|---|---|---|
| | | | erculosis complex | | | | | | | | | | tuberculosis complex | | | | | | -acyl groups (100%) | %);GO:0005829(4%);GO:0005886(4%) | );plasmamembrane(4%) | 6%);GO:0031177(87%);GO:0004315(70%) | hopantetheine binding (87%);3-oxoacyl-[acyl-carrier-protein] synthase activity (70%) | 08152(13%);GO:0006631(9%) | ess(70%);metabolic process(13%);fatty acid metabolic process(9%) | 0%);IPR016035(100%);IPR020807(100%) | | | | | | | | | | | |
| 13 | Variable | EMGVVAHNLGNR | Actinomycet | Bacteria | Actinobac | | Actinomyceti | | | | | | | | | | | | | GO:00057 | cytoplasm(100% | GO:00161 | nickel cation binding (10 | GO:0006457(94 | protein folding (9 | IPR00402 | 1.00E+00 | 1.00E+00 | 6.75E+04 | NS | NS | NS | NS | NS | NS | NS | NS |

| | | | | | | | | | | | | | | | | | | | | | | | | | | | | | | | | | | | | | |
|---|---|---|---|---|---|---|---|---|---|---|---|---|---|---|---|---|---|---|---|---|---|---|---|---|---|---|---|---|---|---|---|---|---|---|---|---|---|
| | 11 | | ia | | teria | | a | | | | | | | | | | | | | 37(100%) | ) | 51(100%);GO:0051082(100%) | 0%);unfolded protein binding (100%) | %);GO:0019627 (94%);GO:0065003 (94%) | 4%); urea metabolic process (94%); protein-containing complex assembly (94%) | 9(100%);IPR012406(100%);IPR036118(100%) | | | | | | | | | | | |
| 14 | Var | SLGYLDV | Myc | Bac | Acti | | Actin | | Cory | | Mycob | Myc | Myc | | | | | | | GO: | integral | | | | | | 1.0 | 1.0 | 5.0 | NS | NS | NS | NS | NS | NS | NS | NS |

| | | | | | | | | | | | | | | | | | | | | | | | | | | | | | | | | | | | | | |
|---|---|---|---|---|---|---|---|---|---|---|---|---|---|---|---|---|---|---|---|---|---|---|---|---|---|---|---|---|---|---|---|---|---|---|---|---|---|
| | iable 110 | SADGR | obacteriumtuberculosiscomplex | teria | nobacteria | | omycetia | | nebacteriales | | acteriaceae | obacterium | obacterium tuberculosiscomplex | | | | | | | 0016021(100%);GO:0005886(16%);GO:0005576(5%) | component of membrane(100%);plasma membrane(16%);extracellular region(5%) | | | | | | 0E+00 | 0E+00 | 7E+04 | | | | | | | | |
| 1 | V | VG | M | B | A | | A | | C | | M | M | M | | | | | 1 | Ox | G | ce | G | oxi | G | re | I | 1 | 1 | 4 | N | N | N | N | N | N | N | N |

| | | | | | | | | | | | | | | | | | | | | | | | | | | | | | | | | | | | | | |
|---|---|---|---|---|---|---|---|---|---|---|---|---|---|---|---|---|---|---|---|---|---|---|---|---|---|---|---|---|---|---|---|---|---|---|---|---|---|
| 5 | ariable 111 | LAPEMEEPPPATPR | ycobacterium tuberculosis complex | acteria | ctinobacteria | | ctinomycetia | | orynebacteriales | | ycobacteriaceae | ycobacterium | ycobacterium tuberculosis complex | | | | | .-.-.-(100%) | idoreductases(100%) | O:0005618(5%);GO:0005829(5%);GO:0005886(5%) | ll wall(5%);cytosol(5%);plasma membrane(5%) | O:0016491(100%) | doreductase activity(100%) | O:0001666(5%);GO:0052572(5%) | sponse to hypoxia(5%);response to host immune response(5%) | PR004415(100%);IPR029479(11%) | .00E+00 | .00E+00 | .76E+04 | S | S | S | S | S | S | S | S |
| 16 | Va | YQPPQ | My | Ba | Ac | | Act | | Cor | | Myc | My | My | | | | | 6. | Methi | GO | cyto | GO | methi | GO: | met | IP | 1. | 1. | 4. | NS | NS | NS | NS | NS | NS | NS | NS |

| | | | | | | | | | | | | | | | | | | | | | | | | | | | | | | | | | | | | | |
|---|---|---|---|---|---|---|---|---|---|---|---|---|---|---|---|---|---|---|---|---|---|---|---|---|---|---|---|---|---|---|---|---|---|---|---|---|---|
| | riable 112 | PPEGK | cobacterium tuberculosis complex | cteria | tinobacteria | | inomycetia | | ynebacteriales | | obacteriaceae | cobacterium | cobacterium tuberculosis complex | | | | | 1.1.10(100%) | onine--tRNA ligase(100%) | :0005737(100%);GO:0005886(6%) | plasm(100%);plasmamembrane(6%) | :0004825(100%);GO:0005524(100%) | onine-tRNA ligase activity(100%);ATP binding(100%) | 0006431(94%);GO:0006418(6%) | hionyl-tRNA aminoacylation(94%);tRNA aminoacylation for protein translation(6%) | R0090800(100%);IPR014729(94%);IPR015413(94%) | 00E+00 | 00E+00 | 37E+04 | | | | | | | | |
| 1 | V | QG | M | B | A | | A | | C | | M | M | M | | | | | 2 | Tra | G | ex | G | tra | G | pe | I | 1 | 1 | 1 | N | N | N | N | N | N | N | N |

| | | | | | | | | | | | | | | | | | | | | | | | | | | | | | | | | | | | | | |
|---|---|---|---|---|---|---|---|---|---|---|---|---|---|---|---|---|---|---|---|---|---|---|---|---|---|---|---|---|---|---|---|---|---|---|---|---|---|
| 7 | ariable 113 | VPVLDGTDAEGVAR | ycobacterium tuberculosis complex | acteria | ctinobacteria | | ctinomycetia | | orynebacteriales | | ycobacteriaceae | ycobacterium | ycobacterium tuberculosis complex | | | | | .2.1.1(100%) | nsketolase (100%) | O:0005576(5%);GO:0005618(5%);GO:0005829(5%) | tracellular region(5%);cell wall(5%);cytosol(5%) | O:0004802(90%);GO:0046872(90%);GO:0016740(10% | nsketolase activity(90%);metal ion binding(90%);transferase activity(10%) | O:0006098(5%) | ntose-phosphate shunt(5%) | PR009014(100%);IPR033247(100%);IPR005475(95%) | .00E+00 | .00E+00 | .03E+05 | S | S | S | S | S | S | S | S |

| | | | | | | | | | | | | | | | | | | | | | | | | | | | | | | | | | | | | | |
|---|---|---|---|---|---|---|---|---|---|---|---|---|---|---|---|---|---|---|---|---|---|---|---|---|---|---|---|---|---|---|---|---|---|---|---|---|---|
| | | | | | | | | | | | | | | | | | | | | | | ) | | | | | | | | | | | | | | | |
| 18 | Variable 114 | SGEGSPDDLVVR | Mycobacterium tuberculosis complex | Bacteria | Actinobacteria | | Actinomycetia | | Corynebacteriales | | Mycobacteriaceae | Mycobacterium | Mycobacterium tuberculosis complex | | | | | | | | | | | | | IPR007367(100%);IPR009057(100%);IPR017277(94 | 1.00E+00 | 1.00E+00 | 9.52E+04 | 1.7332 | 0.0039863 | 0.009619 | 0.9 | NS | NS | NS | NS |

| | | | | | | | | | | | | | | | | | | | | | | | | | | | | | | | | | | | | | |
|---|---|---|---|---|---|---|---|---|---|---|---|---|---|---|---|---|---|---|---|---|---|---|---|---|---|---|---|---|---|---|---|---|---|---|---|---|---|
| | | | | | | | | | | | | | | | | | | | | | | | | | | %) | | | | | | | | | | | |
| 21 | Variable 117 | VPDGFQLPHPPR | Mycobacterium tuberculosis complex | Bacteria | Actinobacteria | | Actinomycetia | | Corynebacteriales | | Mycobacteriaceae | Mycobacterium | Mycobacterium tuberculosis complex | | | | | 1.2.1.2(63%);1.-.-.-(38%) | Transferred entry: 1.17.1.9 (63%); Oxidoreductases (38%) | GO:0005886(6%) | plasma membrane (6%) | GO:0008863(100%);GO:0030151(100%);GO:005153 | formate dehydrogenase (NAD+) activity (100%);molybdenum ion binding (100%);4 iron | 4 sulfur cluster binding (100%) | | | 1.00E+00 | 1.00E+00 | 3.82E+04 | NS | NS | NS | NS | NS | NS | NS | NS |

| | | | | | | | | | | | | | | | | | | | | | | | | | | | | | | | | | | | | | | | | |
|---|---|---|---|---|---|---|---|---|---|---|---|---|---|---|---|---|---|---|---|---|---|---|---|---|---|---|---|---|---|---|---|---|---|---|---|---|---|---|---|---|
| | | | | | | | | | | | | | | | | | | | | | | 9(100%) | | | | | | | | | | | | | | | | | | |
| 22 | Variable 118 | SQQSAYASHPALLDACFQSVLVHPEVQK | Mycobacterium tuberculosis complex | Bacteria | Actinobacteria | | Actinomycetia | | Corynebacteriales | | Mycobacteriaceae | Mycobacterium | Mycobacterium tuberculosis complex | | | | | 2.3.1.-(75%);2.7.1.-(63%);2.3.1.252(25%) | Transferring groups other than amino-acyl groups (75%); Phosphotransferases with an alcohol group as acc | GO:0005886(6%);GO:0034081(6%) | plasma membrane (6%);polyketide synthase complex (6%) | GO:0016491(100%);GO:0031177(82%);GO:00 | oxidoreductase activity (100%);phosphopantetheine binding (82%);3-oxoacyl-[acyl-carrier-protein] syn | GO:0006633(76%);GO:0008152(6%);GO:1902321(6%) | fatty acid biosynthetic process (76%); metabolic process (6%); methyl-bran | IPR011032(100%);IPR013154(100%);IPR014 | 1.00E+00 | 1.00E+00 | 2.32E+04 | 2.3784 | 0.00065199 | 0.00014028 | 0.9 | NS | NS | NS | NS | 6.0165 | 5.04E-03 | 0 |

| | | | | | | | | | | | | | | | | | | | | | | | | | | | | | | | | | | | | | | | | |
|---|---|---|---|---|---|---|---|---|---|---|---|---|---|---|---|---|---|---|---|---|---|---|---|---|---|---|---|---|---|---|---|---|---|---|---|---|---|---|---|---|
| | | | | | | | | | | | | | | | | | | | eptor (63%); Mycolipanoate synthase (25%) | | | 0043315 (76%) | thase activity (76%) | | ched fatty acid biosynthetic process (6%) | 043 (100%) | | | | | | | | | | | | | | |
| 23 | Variable 119 | WGLAALADGVVDESNAR | Mycobacterium tuberculosis complex | Bacteria | Actinobacteria | | Actinomycetia | | Corynebacteriales | | Mycobacteriaceae | Mycobacterium | Mycobacterium tuberculosis | | | | | 4.2.1.51 (100%) | Prephenate dehydratase (100%) | GO:0005737 (6%) | cytoplasm (6%) | GO:0004664 (100%); GO:00041 | prephenate dehydratase activity (100%);chorismate mutase activit | GO:0009094 (89%);GO:0033585 (6%) | L-phenylalanine biosynthetic process (89%); L-ph | IPR002912 (100%); IPR001086 | 1.00E+00 | 1.00E+00 | 8.35E+04 | 16.288 | 9.216E-09 | 0.00046595 | 0.9 | NS | NS | NS | NS | 21.976 | 2.94E-07 | 0.0053771 |

| | | | | | | | | | | | | | | | | | | | | | | | | | | | | | | | | | | | | | |
|---|---|---|---|---|---|---|---|---|---|---|---|---|---|---|---|---|---|---|---|---|---|---|---|---|---|---|---|---|---|---|---|---|---|---|---|---|---|
| | | | | | | | | | | | | | complex | | | | | | | | | 06(67%);GO:0042803(22%) | y(67%);protein homodimerization activity(22%) | | enylalanine biosynthetic process from chorismate via phenylpyruvate(6%) | (89%);IPR018528(83%) | | | | | | | | | | | |
| 24 | Variab | MASRPSAAK | Actinomy | Bacter | Actinob | | Actinomy | | | | | | | | | | | | | GO:001 | integral compo | GO:000 | ATP binding(93 | GO:0051301 | cell division | IPR002 | 1.00E+ | 1.00E+ | 5.91E+ | NS | NS | NS | NS | NS | NS | NS | NS |

| | | | | | | | | | | | | | | | | | | | | | | | | | | | | | | | | | | | | | | | | |
|---|---|---|---|---|---|---|---|---|---|---|---|---|---|---|---|---|---|---|---|---|---|---|---|---|---|---|---|---|---|---|---|---|---|---|---|---|---|---|---|---|
| | le 12 | | cetia | ia | acteria | | cetia | | | | | | | | | | | | | 6021(94%);GO:0005886(3%) | nent of membrane(94%);plasma membrane(3%) | 5524(93%);GO:0003677(87%) | %);DNA binding(87%) | (43%);GO:0007049(1%);GO:0007059(1%) | (43%);cell cycle(1%);chromosome segregation(1%) | 543(100%);IPR003593(100%);IPR018541(100%) | 00 | 00 | 04 | | | | | | | | | | | |
| 25 | Vari | GLDGATGG | Myco | Bact | Actin | | Actino | | Coryne | | Mycobac | Myco | Myco | | | | | | | GO:0 | host cell | | | GO:0009 | obsolete | IPR0 | 1.00 | 1.00 | 2.21 | 39.7 | 0.00 | 0.000 | 0.9 | NS | NS | NS | NS | 6.27 | 4.12 | 0.00 |

| | | | | | | | | | | | | | | | | | | | | | | | | | | | | | | | | | | | | | | | | |
|---|---|---|---|---|---|---|---|---|---|---|---|---|---|---|---|---|---|---|---|---|---|---|---|---|---|---|---|---|---|---|---|---|---|---|---|---|---|---|---|---|
| | able 120 | K | bacterium tuberculosis complex | eria | obacteria | | mycetia | | bacteriales | | teriaceae | bacterium | bacterium tuberculosis complex | | | | | | | 044165(100%) | endoplasmic reticulum(100%) | | | 405(100%) | pathogenesis(100%) | 00084(100%) | E+00 | E+00 | E+05 | 25 | 00807 | 24013 | | | | | | 68 | E-03 | 070925 |
| 26 | Variable 121 | NLRPHLAGASSGVFNAVR | Mycobacterium tuberculo | Bacteria | Actinobacteria | | Actinomycetia | | Corynebacteriales | | Mycobacteriaceae | Mycobacterium | Mycobacterium tub | | | | | | | GO:0016021(100%); | integral component of membrane (1 | GO:0022857(90%) | transmembrane transporter activity (90%) | | | IPR036259(100%);I | 1.00E+00 | 1.00E+00 | 5.49E+04 | NS | NS | NS | NS | NS | NS | NS | NS | | | |

| | | | | | | | | | | | | | | | | | | | | | | | | | | | | | | | | | | | | | |
|---|---|---|---|---|---|---|---|---|---|---|---|---|---|---|---|---|---|---|---|---|---|---|---|---|---|---|---|---|---|---|---|---|---|---|---|---|---|
| | | | siscomplex | | | | | | | | | | erculosiscomplex | | | | | | | GO:0005886(75%);GO:0016020(5%) | 00%);plasmamembrane(75%);membrane(5%) | | | | | PR011701(90%);IPR004638(75%) | | | | | | | | | | | |
| 27 | Variable 122 | GVDLVALHAWSDMGPLDFPR | Mycobacteriumtuberc | Bacteria | Actinobacteria | | Actinomycetia | | Corynebacteriales | | Mycobacteriaceae | Mycobacterium | Mycobacterium t | | | | | | | GO:0005618(25% | cellwall(25%);plasmamem | GO:0005524(75% | ATP binding(75%) | GO:0071500(25%) | cellular response to nitrosative str | IPR006015(100% | 1.00E+00 | 1.00E+00 | 4.06E+04 | NS | NS | NS | NS | NS | NS | NS | NS |

| | | | | | | | | | | | | | | | | | | | | | | | | | | | | | | | | | | | | | |
|---|---|---|---|---|---|---|---|---|---|---|---|---|---|---|---|---|---|---|---|---|---|---|---|---|---|---|---|---|---|---|---|---|---|---|---|---|---|
| | | | ulosis complex | | | | | | | | | | uberculosis complex | | | | | | | );GO:0005886(25%) | brane (25%) | ) | | | ess (25%) | );IPR006016(100%) | | | | | | | | | | | |
| 28 | Variable 123 | APAATVPAAAAAAATR | Mycobacterium tuberculosis complex | Bacteria | Actinobacteria | | Actinomyceta | | Corynebacteriales | | Mycobacteriaceae | Mycobacterium | Mycobacterium tuberculosisco | | | | | | | GO:0016021(100%);GO:0005886 | integral component of membrane (100%);plasma membr | | | | | IPR000030(100%);IPR038332(1 | 1.00E+00 | 1.00E+00 | 5.24E+04 | NS | NS | NS | NS | NS | NS | NS | NS |

| | | | | | | | | | | | | | | | | | | | | | | | | | | | | | | | | | | | | | |
|---|---|---|---|---|---|---|---|---|---|---|---|---|---|---|---|---|---|---|---|---|---|---|---|---|---|---|---|---|---|---|---|---|---|---|---|---|---|
| | | | | | | | | | | | | | mplex | | | | | | | (15%) | ane (15%) | | | | | 00%);IPR0043641(93%) | | | | | | | | | | | |
| 29 | Variable 124 | QSVDLTPALYQELR | Mycobacterium tuberculosis complex | Bacteria | Actinobacteria | | Actinomyceta | | Corynebacteriales | | Mycobacteriaceae | Mycobacterium | Mycobacterium tuberculosis c | | | | | 2.4.2.-(100%) | Pentosyltransferases (100%) | | | GO:0016779(100%) | nucleotidyltransferase activity (100%) | | | IPR014914(100%) | 1.00E+00 | 1.00E+00 | 3.60E+04 | 3.6185 | 0.002349 | 0.00004359 | 0.9 | NS | NS | NS | NS |

| | | | | | | | | | | | | | | | | | | | | | | | | | | | | | | | | | | | | | | | | |
|---|---|---|---|---|---|---|---|---|---|---|---|---|---|---|---|---|---|---|---|---|---|---|---|---|---|---|---|---|---|---|---|---|---|---|---|---|---|---|---|---|
| | | | | | | | | | | | | | omplex | | | | | | | | | | | | | | | | | | | | | | | | | | | |
| 30 | Variable 125 | DVLATTGLDR | Mycobacterium tuberculosis complex | Bacteria | Actinobacteria | | Actinomycetia | | Corynebacteriales | | Mycobacteriaceae | Mycobacterium | Mycobacterium tuberculosis complex | | | | | 3.-.-.-(100%);3.2.1.-(50%);3.1.3.-(25%) | Hydrolases(100%);Glycosidases | i.e. enzymes hydrolyzing O- and S-glycosylco | | | GO:0016787(88%);GO:0016798(13%) | hydrolase activity(88%);hydrolase activity | acting on glycosyl bonds(13%) | GO:0008152(13%) | 1.00E+00 | 1.00E+00 | 2.19E+05 | 0.1745 | 1.3481E-11 | 0.0025733 | 0.9 | NS | NS | NS | NS | 18.018 | 2.24E-06 | 0.0005753 |

| | | | | | | | | | | | | | | | | | | | | | | | | | | | | | | | | | | | | | |
|---|---|---|---|---|---|---|---|---|---|---|---|---|---|---|---|---|---|---|---|---|---|---|---|---|---|---|---|---|---|---|---|---|---|---|---|---|---|
| | | | | | | | | | | | | | | | | | | | | mpounds(50%);Phosphoric monoester hydrolases(25%) | | | | | | | | | | | | | | | | | |
| 3 | V | QG | M | B | A | | A | | C | | M | M | M | | | | | 2 | Bet | G | po | G | oxi | G | fat | I | 1 | 4 | 1 | N | N | N | N | N | N | N | N |

| | | | | | | | | | | | | | | | | | | | | | | | | | | | | | | | | | | | | | |
|---|---|---|---|---|---|---|---|---|---|---|---|---|---|---|---|---|---|---|---|---|---|---|---|---|---|---|---|---|---|---|---|---|---|---|---|---|---|
| 1 | ariable 126 | PSSNSALVVPVR | ycobacterium tuberculosis complex | acteria | ctinobacteria | | ctinomycetia | | orynebacteriales | | ycobacteriaceae | ycobacterium | ycobacterium tuberculosis complex | | | | | .3.1.41(71%);2.3.1.292(29%) | a-ketoacyl-[acyl-carrier-protein] synthase I (71%);(Phenol)carboxyphthiodiolenone synthase (29%) | O:0034081(11%);GO:0005829(6%);GO:0005886(6%) | lyketide synthase complex(11%);cytosol(6%);plasma membrane(6%) | O:0016491(100%);GO:0031177(94%);GO:0004315(83 | doreductase activity (100%);phosphopantetheine binding (94%);3-oxoacyl-[acyl-carrier-protein] synthase activity (83%) | O:0006633(72%);GO:0071766(11%);GO:0097040(11%) | tyacid biosynthetic process(72%);Actinobacterium-type cell wall biogenesis(11%);phthi | PR020807(100%);IPR036291(100%);IPR042104(100% | .00E+00 | .68E+04 | .00E+00 | S | S | S | S | S | S | S | S |

| | | | | | | | | | | | | | | | | | | | | | | | | | | | | | | | | | | | | | |
|---|---|---|---|---|---|---|---|---|---|---|---|---|---|---|---|---|---|---|---|---|---|---|---|---|---|---|---|---|---|---|---|---|---|---|---|---|---|
| | | | | | | | | | | | | | | | | | | | | | | %) | | | ocerol biosynthetic process (11%) | ) | | | | | | | | | | | |
| 32 | Variable 127 | HSGATTASVR | Mycobacterium tuberculosis complex | Bacteria | Actinobacteria | | Actinomycetia | | Corynebacteriales | | Mycobacteriaceae | Mycobacterium | Mycobacterium tuberculosis co | | | | | 2.7.-.-(91%);2.7.13.3(73%) | Transferring phosphorus-containing groups (91%); Histidine kinase (73%) | GO:0016021(71%);GO:0005886( | integral component of membrane (71%);plasma membr | GO:0016301(57%);GO:0004673( | kinase activity (57%);protein histidine kinase activity (38%) | GO:0000160 (5%) | phosphorelay signal transduction system (5%) | IPR036890(100%);IPR003594(9 | 1.00E+00 | 5.37E+05 | 1.00E+00 | NS | NS | NS | NS | NS | NS | NS | NS |

| | | | | | | | | | | | | | | | | | | | | | | | | | | | | | | | | | | | | | |
|---|---|---|---|---|---|---|---|---|---|---|---|---|---|---|---|---|---|---|---|---|---|---|---|---|---|---|---|---|---|---|---|---|---|---|---|---|---|
| | | | | | | | | | | | | | mplex | | | | | | | 5%) | ane(5%) | 38%) | | | | 0%);IPR005467(67%) | | | | | | | | | | | |
| 34 | Variable 129 | AAAAASAYASR | Mycobacterium tuberculosis complex | Bacteria | Actinobacteria | | Actinomycetia | | Corynebacteriales | | Mycobacteriaceae | Mycobacterium | Mycobacterium tuberculosis co | | | | | | | GO:0016021(100%);GO:0005886 | integral component of membrane(100%);plasma membr | GO:0016209(91%);GO:0016491( | antioxidant activity(91%);oxidoreductase activity(91%) | GO:0017004(100%) | cytochrome complex assembly(100%) | IPR003834(100%);IPR036249(1 | 1.00E+00 | 1.64E+05 | 1.00E+00 | NS | NS | NS | NS | NS | NS | NS | NS |

| | | | | | | | | | | | | | | | | | | | | | | | | | | | | | | | | | | | | | |
|---|---|---|---|---|---|---|---|---|---|---|---|---|---|---|---|---|---|---|---|---|---|---|---|---|---|---|---|---|---|---|---|---|---|---|---|---|---|
| | | | | | | | | | | | | | mplex | | | | | | | (91%) | ane(91%) | 91%) | | | | 00%);IPR000866(91%) | | | | | | | | | | | |
| 35 | Variable 13 | YLEELAGTYHR | Actinomycetia | Bacteria | Actinobacteria | | Actinomyceta | | | | | | | | | | | 6.1.1.19(100%) | Arginine--tRNA ligase(100%) | GO:0005737(98%) | cytoplasm(98%) | GO:0004814(100%);GO:00552 | arginine-tRNA ligase activity(100%);ATP binding(100%) | GO:0006420(100%) | arginyl-tRNA aminoacylation(100%) | IPR008909(100%);IPR009080( | 1.00E+00 | 1.00E+00 | 6.36E+04 | NS | NS | NS | NS | NS | NS | NS | NS |

| | | | | | | | | | | | | | | | | | | | | | | | | | | | | | | | | | | | | | |
|---|---|---|---|---|---|---|---|---|---|---|---|---|---|---|---|---|---|---|---|---|---|---|---|---|---|---|---|---|---|---|---|---|---|---|---|---|---|
| | | | | | | | | | | | | | | | | | | | | | | 4(100%) | | | | 100%);IPR001278(99%) | | | | | | | | | | | |
| 36 | Variable 130 | LGPCPAQWPVADDCR | Mycobacterium tuberculosis complex | Bacteria | Actinobacteria | | Actinomycetia | | Corynebacteriales | | Mycobacteriaceae | Mycobacterium | Mycobacterium tuberculosis | | | | | 5.3.3.2(100%) | Isopentenyl-diphosphate Delta-isomerase (100%) | GO:0005737(92%) | cytoplasm(92%) | GO:0004452(100%);GO:00167 | isopentenyl-diphosphate delta-isomerase activity (100%);hydrolase | GO:0008299(92%);GO:0050992(92%) | isoprenoid biosynthetic process (92%);dimethylal | IPR015797(100%);IPR00086 | 1.00E+00 | 9.92E+04 | 1.00E+00 | NS | NS | NS | NS | NS | NS | NS | NS |

| | | | | | | | | | | | | | | | | | | | | | | | | | | | | | | | | | | | | | |
|---|---|---|---|---|---|---|---|---|---|---|---|---|---|---|---|---|---|---|---|---|---|---|---|---|---|---|---|---|---|---|---|---|---|---|---|---|---|
| | | | | | | | | | | | | | complex | | | | | | | | | 87(100%);GO:0046872(92%) | activity(100%);metal ion binding(92%) | | lyldiphosphate biosynthetic process(92%) | (92%);IPR011876(92%) | | | | | | | | | | | |
| 37 | Variable 131 | LLADGPASR | Mycobacterium tuberculosis co | Bacteria | Actinobacteria | | Actinomycetia | | Corynebacteriales | | Mycobacteriaceae | Mycobacterium | Mycobacterium tubercu | | | | | 6.5.1.2(100%) | DNA ligase (NAD(+))(100%) | GO:0005618(5%);GO:000 | cell wall(5%);cytosol(5%);plasma membr | GO:0003911(89%);GO:00 | DNA ligase (NAD+) activity(89%);metal ion bindin | GO:0006260(68%);GO:0006281(63%);GO: | DNA replication(68%);DNA repair(63 | IPR001357(100%);IPR03 | 1.00E+00 | 4.59E+05 | 1.00E+00 | NS | NS | NS | NS | NS | NS | NS | NS |

| | | | | | | | | | | | | | | | | | | | | | | | | | | | | | | | | | | | | | |
|---|---|---|---|---|---|---|---|---|---|---|---|---|---|---|---|---|---|---|---|---|---|---|---|---|---|---|---|---|---|---|---|---|---|---|---|---|---|
| | | | mplex | | | | | | | | | | losis complex | | | | | | | 5829(5%);GO:0005886(5%) | ane(5%) | 46872(63%);GO:0016874(11%) | g(63%);ligase activity(11%) | 0006266(5%) | %);DNA ligation(5%) | 6420(100%);IPR001679(68%) | | | | | | | | | | | |
| 38 | Variable 132 | MTAFGVEPYGQPK | Mycobacterium tuberculosis | Bacteria | Actinobacteria | | Actinomycetia | | Corynebacteriales | | Mycobacteriaceae | Mycobacterium | Mycobacterium tuber | | | | | 3.8.1.5(100%) | Haloalkane dehalogenase(100%) | GO:0005618(5%);GO:0 | cell wall(5%);plasma membrane(5% | GO:0018786(100%);GO | haloalkane dehalogenase activity(100%);hydrolase | GO:0042206(5%) | halogenated hydrocarbon catabolic proc | IPR000073(100%);IPR | 1.00E+00 | 9.74E+04 | 1.00E+00 | NS | NS | NS | NS | NS | NS | NS | NS |

| | | | | | | | | | | | | | | | | | | | | | | | | | | | | | | | | | | | | | |
|---|---|---|---|---|---|---|---|---|---|---|---|---|---|---|---|---|---|---|---|---|---|---|---|---|---|---|---|---|---|---|---|---|---|---|---|---|---|
| | | | complex | | | | | | | | | | culosis complex | | | | | | | 0058886(5%) | ) | :0016787(5%) | activity (5%) | | ess (5%) | 0290058(100%);IPR000639(95%) | | | | | | | | | | | |
| 39 | Variable 133 | LFPTGFDEQR | Mycobacterium tuberculosi | Bacteria | Actinobacteria | | Actinomycetia | | Corynebacteriales | | Mycobacteriaceae | Mycobacterium | Mycobacterium tube | | | | | 3.1.11.5(100%) | Exodeoxyribonuclease V (100%) | GO:0009338(100%);G | exodeoxyribonuclease V complex (100 | GO:0008854(100%);G | exodeoxyribonuclease V activity (100%);DNA | GO:0000724(93%);GO:0006310(13 | double-strand break repair via homol | IPR006344(100%);IP | 5.03E+06 | 1.00E+00 | 1.00E+00 | NS | NS | NS | NS | NS | NS | NS | NS |

| | | | | | | | | | | | | | | | | | | | | | | | | | | |
|---|---|---|---|---|---|---|---|---|---|---|---|---|---|---|---|---|---|---|---|---|---|---|---|---|---|---|
| | | | s complex | | | | | | | | | | rculosis complex | | | | | | | O:0005618(7%);GO:0005829(7%) | %);cell wall(7%);cytosol(7%) | O:0003677(93%);GO:0005524(93%) | binding(93%);ATP binding(93%) | %);GO:0006302(7%) | ogous recombination(93%);DNA recombination(13%);double-strand break repair(7%) | R027417(100%);IPR027785(93%) |

| | | | | | | | | | | | | | | | | | | | | | | | | | | | | | | | | | | | | | |
|---|---|---|---|---|---|---|---|---|---|---|---|---|---|---|---|---|---|---|---|---|---|---|---|---|---|---|---|---|---|---|---|---|---|---|---|---|---|
| 40 | Variable 134 | YPPDGVSADSVR | Mycobacterium tuberculosis complex | Bacteria | Actinobacteria | | Actinomycetia | | Corynebacteriales | | Mycobacteriaceae | Mycobacterium | Mycobacterium tuberculosis complex | | | | | 2.7.7.42(100%);2.7.7.89(67%) | [Glutamine synthetase] adenylyltransferase(100%);[Glutamine synthetase]-adenylyl-L-tyrosine phosphorylase(67%) | GO:0005618(5%);GO:0005829(5%);GO:0005886(5%) | cell wall(5%);cytosol(5%);plasma membrane(5%) | GO:0005524(100%);GO:0008882(100%);GO:0016874( | ATP binding(100%);[glutamate-ammonia-ligase] adenylyltransferase activity(100%);ligase activity(70%) | GO:0000820(60%);GO:0060359(5%) | regulation of glutamine family amino acid metabolic process(60%);response to ammonium | IPR005190(100%);IPR013546(100%);IPR023057(100 | 4.65E+05 | 1.00E+00 | 1.00E+00 | NS | NS | NS | NS | NS | NS | NS | NS |

| | | | | | | | | | | | | | | | | | | | | | | | | | | | | | | | | | | | | | |
|---|---|---|---|---|---|---|---|---|---|---|---|---|---|---|---|---|---|---|---|---|---|---|---|---|---|---|---|---|---|---|---|---|---|---|---|---|---|
| | | | | | | | | | | | | | | | | | | | | | | 70%) | | | ion(5%) | %) | | | | | | | | | | | |
| 41 | Variable 135 | CCDEATPSAGPR | Mycobacterium tuberculosis | Bacteria | Actinobacteria | | Actinomycetia | | Corynebacteriales | | Mycobacteriaceae | Mycobacterium | Mycobacterium tuberculosis complex | | Mycobacterium tuberculosis | | | 2.8.4.3(100%) | tRNA-2-methylthio-N(6)-dimethylallyladenosine synthase(100%) | GO:0005737(75%);GO:0005829(25%) | cytoplasm(75%);cytosol(25%) | GO:0035596(100%);GO:0046872(100%);GO:005 | methylthiotransferase activity(100%);metal ion binding(100%);4iron | 4 sulfur cluster binding(100%) | GO:0006400(75%);GO:0035600(25%) | tRNA modification(75%);tRNA methylthiolation(25%) | 1.00E+00 | 1.00E+00 | 1.60E+05 | 9.7989 | 0.0031978 | 0.0003512 9 | 0.9 | NS | NS | NS | NS |

| | | | | | | | | | | | | | | | | | | | | | | | | | | | | | | | | | | | | | |
|---|---|---|---|---|---|---|---|---|---|---|---|---|---|---|---|---|---|---|---|---|---|---|---|---|---|---|---|---|---|---|---|---|---|---|---|---|---|
| | | | | | | | | | | | | | | | | | | | | | | 1539(100%) | | | | | | | | | | | | | | | |
| 42 | Variable 136 | RPVPGGYSGMHAASR | Mycobacteroides abscessus | Bacteria | Actinobacteria | | Actinomycetia | | Corynebacteriales | | Mycobacteriaceae | Mycobacteroides | | | Mycobacteroides abscessus | | | | | GO:0005886(100%);GO:0016021(92%);GO | plasma membrane(100%);integral component of membrane(92%);p | coupling factor F(o)(92%) | GO:0046933(100%);GO:0016787(8%) | proton-transporting ATP synthase activity | rotational mechanism(100%);hydrolase activity(8%) | | 1.00E+00 | 1.00E+00 | 8.78E+04 | NS | NS | NS | NS | NS | NS | NS | NS |

| | | | | | | | | | | | | | | | | | | | | | | | | | | | | | | | | | | | | | |
|---|---|---|---|---|---|---|---|---|---|---|---|---|---|---|---|---|---|---|---|---|---|---|---|---|---|---|---|---|---|---|---|---|---|---|---|---|---|
| | | | | | | | | | | | | | | | | | | | | :0045263(92%) | roton-transporting ATP synthase complex | | | | | | | | | | | | | | | | |
| 43 | Variable 137 | LAPSLPAPQR | Mycobacteroidesabscessus | Bacteria | Actinobacteria | | Actinomycetia | | Corynebacteriales | | Mycobacteriaceae | Mycobacteroides | | | Mycobacteroidesabscessus | | | 2.7.7.105(100%) | -1 | GO:0016021(15%) | integral component of membrane(15%) | GO:0043814(100%);GO:00055 | phospholactate guanylyltransferase activity(100%);GTP binding(85%) | GO:0052645(100%) | F420-0 metabolic process(100%) | IPR002835(100%);IPR029044 | 1.00E+00 | 1.00E+00 | 2.71E+05 | NS | NS | NS | NS | NS | NS | NS | NS |

| | | | | | | | | | | | | | | | | | | | | | | | | | | | | | | | | | | | | | |
|---|---|---|---|---|---|---|---|---|---|---|---|---|---|---|---|---|---|---|---|---|---|---|---|---|---|---|---|---|---|---|---|---|---|---|---|---|---|
| | | | | | | | | | | | | | | | | | | | | | | 25(85%) | | | | (100%) | | | | | | | | | | | |
| 44 | Variable 138 | LASGNTLLLSK | Mycolicibacterium | Bacteria | Actinobacteria | | Actinomycetia | | Corynebacteriales | | Mycobacteriaceae | Mycolicibacterium | | | | | | 2.7.13.3(100%);3.1.3.-(50%) | Histidine kinase(100%);Phosphoric monoesterhydrolases(50%) | GO:0005886(100%);GO:0016021(100%) | plasma membrane(100%);integral component of membrane(100%) | GO:0000155(100%);GO:0004721(50%);GO:00 | phosphorelay sensor kinase activity(100%);phosphoprotein phosphatase activity(50%);ATP binding(50%) | GO:0009405(50%) | obsolete pathogenesis(50%) | IPR003594(100%);IPR003660(100%);IPR003 | 1.00E+00 | 1.00E+00 | 3.06E+05 | NS | NS | NS | NS | NS | NS | NS | NS |

| | | | | | | | | | | | | | | | | | | | | | | | | | | | | | | | | | | | | | |
|---|---|---|---|---|---|---|---|---|---|---|---|---|---|---|---|---|---|---|---|---|---|---|---|---|---|---|---|---|---|---|---|---|---|---|---|---|---|
| | | | | | | | | | | | | | | | | | | | | | | 05524(50%) | | | | 661(100%) | | | | | | | | | | | |
| 45 | Variable 139 | NQNPAGEGERP | Mycolicibacterium gilvum | Bacteria | Actinobacteria | | Actinomycetia | | Corynebacteriales | | Mycobacteriaceae | Mycolicibacterium | | | Mycolicibacterium gilvum | | | 2.7.7.18(100%) | Nicotinate-nucleotide adenylyltransferase (100%) | | | GO:0004515(100%);GO:0005524(100%) | nicotinate-nucleotide adenylyltransferase activity (100%);ATP binding (100%) | GO:0009435 (100%) | NAD biosynthetic process (100%) | IPR004821(100%);IPR005248(100%);IPR | 1.00E+00 | 1.00E+00 | 3.14E+04 | NS | NS | NS | NS | NS | NS | NS | NS |

| | | | | | | | | | | | | | | | | | | | | | | | | | | | | | | | | | | | | | | | | |
|---|---|---|---|---|---|---|---|---|---|---|---|---|---|---|---|---|---|---|---|---|---|---|---|---|---|---|---|---|---|---|---|---|---|---|---|---|---|---|---|---|
| | | | | | | | | | | | | | | | | | | | | | | | | | | 014729(100%) | | | | | | | | | | | | | | |
| 46 | Variable 14 | GNSVAGVAALDR | Actinomycetia | Bacteria | Actinobacteria | | Actinomycetia | | | | | | | | | | | 3.5.1.2(100%) | Glutaminase(100%) | | | GO:0004359(100%) | glutaminase activity(100%) | GO:0006541(97%);GO:0006537(0%);GO:0006543(0%) | glutamine metabolic process(97%);glutamate biosynthetic pr | IPR012338(100%);IPR015868(100%) | 1.00E+00 | 1.00E+00 | 8.94E+04 | NS | NS | NS | NS | NS | NS | NS | NS | 4.9408 | 1.18E-02 | 0.00029959 |

| | | | | | | | | | | | | | | | | | | | | | | | | | | | | | | | | | | | | | |
|---|---|---|---|---|---|---|---|---|---|---|---|---|---|---|---|---|---|---|---|---|---|---|---|---|---|---|---|---|---|---|---|---|---|---|---|---|---|
| | | | | | | | | | | | | | | | | | | | | | | | | | ocess(0%);glutaminecatabolicprocess(0%) | | | | | | | | | | | | |
| 47 | Variable 140 | LAEADAMHTK | Mycolicibacterium gilvum | Bacteria | Actinobacteria | | Actinomycetia | | Corynebacteriales | | Mycobacteriaceae | Mycolicibacterium | | | Mycolicibacterium gilvum | | | | | GO:0005886(100%);GO:00 | plasmamembrane(100%);integralcomponent | couplingfactorF(o)(100% | GO:0046933(100%) | proton-transportingATPsynthaseactivity | rotationalmechanism(100%) | | 1.00E+00 | 1.38E+05 | 1.00E+00 | NS | NS | NS | NS | NS | NS | NS | NS |

| | | | | | | | | | | | | | | | | | | | | | | | | | | | | | | | | | | | | | | | | |
|---|---|---|---|---|---|---|---|---|---|---|---|---|---|---|---|---|---|---|---|---|---|---|---|---|---|---|---|---|---|---|---|---|---|---|---|---|---|---|---|---|
| | | | | | | | | | | | | | | | | | | | | 16021(100%);GO:0045263(100%) | nt of membrane (100%);proton-transporting ATP synthase complex | ) | | | | | | | | | | | | | | | | | | |
| 48 | Variable 141 | LATPDQGFTSSQAAAR | Mycolicibacteriums | Bacteria | Actinobacteria | | Actinomycetia | | Corynebacteriales | | Mycobacteriaceae | Mycolicibacteriu | | | Mycolicibacteriu | | | 1.-.-.-(100%);1. | Oxidoreductases (100%);2 | 5-didehydrog | | | GO:0047834(100%);GO:0050 | D-threo-aldose 1-dehydrogen | 5-didehydrogluconate redu | | 1.00E+00 | 1.00E+00 | 6.86E+05 | 0.3468 | 2.3375E-09 | 0.00043481 | 0.9 | NS | NS | NS | NS | 9.6395 | 3.59E-04 | 0.00038022 |

| | | | | | | | | | | | | | | | | | | | | | | | | | |
|---|---|---|---|---|---|---|---|---|---|---|---|---|---|---|---|---|---|---|---|---|---|---|---|---|---|
| | | | megmatis | | | | | | | | | m | | | m smegmatis | | | 1.1.274 (50%) | | luconate reductase (2-dehydro-D-gluconate-forming) | | | 580 (25%) | ase activity (100%);2 | ctase activity (25%) |

| | | | | | | | | | | | | | | | | | | | | | | | | | | | | | | | | | | | | | |
|---|---|---|---|---|---|---|---|---|---|---|---|---|---|---|---|---|---|---|---|---|---|---|---|---|---|---|---|---|---|---|---|---|---|---|---|---|---|
| | | | | | | | | | | | | | | | | | | | | (50%) | | | | | | | | | | | | | | | | | |
| 49 | Variable 142 | FGALDALVR | Mycolicibacterium smegmatis | Bacteria | Actinobacteria | | Actinomycetia | | Corynebacteriales | | Mycobacteriaceae | Mycolicibacterium | | | Mycolicibacterium smegmatis | | | 6.5.1.1(100%) | DNA ligase (ATP) (100%) | | | GO:0003677(100%);GO:0003910(100%);GO:000 | DNA binding(100%);DNA ligase (ATP) activity (100%);ATP binding (100%) | GO:0006260(100%);GO:0006281(100%);GO:0006310(100%) | DNA replication(100%);DNA repair(100%);DNA recombination(100%) | IPR009777(100%);IPR012308(100%);IPR012300 | 4.96E+05 | 2.03E+06 | 1.08E+07 | NS | NS | NS | NS | NS | NS | NS | NS |

| | | | | | | | | | | | | | | | | | | | | | | | | | | | | | | | | | | | | | |
|---|---|---|---|---|---|---|---|---|---|---|---|---|---|---|---|---|---|---|---|---|---|---|---|---|---|---|---|---|---|---|---|---|---|---|---|---|---|
| | | | | | | | | | | | | | | | | | | | | | | 55524(100%) | | | | 9(100%) | | | | | | | | | | | |
| 50 | Variable 143 | WEAVMMNNYGTPPLSLVSGEGAVVTDADGR | Mycolicibacterium smegmatis | Bacteria | Actinobacteria | | Actinomycetia | | Corynebacteriales | | Mycobacteriaceae | Mycolicibacterium | | | Mycolicibacterium smegmatis | | | 2.6.1.11(100%) | Acetylornithine transaminase (100%) | GO:0005737(100%) | cytoplasm(100%) | GO:0003992(100%);GO:0030170(100%) | N2-acetyl-L-ornithine:2-oxoglutarate 5-aminotransferase activity (100%);pyridoxal phosphat | GO:0006526(100%) | arginine biosynthetic process (100%) | IPR004636(100%);IPR005814(100%);IPR | 1.00E+00 | 6.60E+05 | 1.00E+00 | NS | NS | NS | NS | NS | NS | NS | NS |

| | | | | | | | | | | | | | | | | | | | | | | | | | | | | | | | | | | | | | |
|---|---|---|---|---|---|---|---|---|---|---|---|---|---|---|---|---|---|---|---|---|---|---|---|---|---|---|---|---|---|---|---|---|---|---|---|---|---|
| | | | | | | | | | | | | | | | | | | | | | | | e binding (100%) | | | 015421 (100%) | | | | | | | | | | | |
| 51 | Variable 144 | AVALVPEER | Nocardiaceae | Bacteria | Actinobacteria | | Actinomycetia | | Corynebacteriales | | Nocardiaceae | | | | | | | 2.7.1.39 (100%) | Homoserine kinase (100%) | GO:0005737 (100%) | cytoplasm (100%) | GO:0004413 (100%);GO:0005524 (100% | homoserine kinase activity (100%);ATP binding (100%) | GO:0009088 (100%) | threonine biosynthetic process (100%) | IPR000870 (100%);IPR006204 (100%);I | 1.00E+00 | 3.85E+05 | 1.00E+00 | NS | NS | NS | NS | NS | NS | NS | NS |

| | | | | | | | | | | | | | | | | | | | | | | | | | | | | | | | | | | | | | |
|---|---|---|---|---|---|---|---|---|---|---|---|---|---|---|---|---|---|---|---|---|---|---|---|---|---|---|---|---|---|---|---|---|---|---|---|---|---|
| | | | | | | | | | | | | | | | | | | | | | | ) | | | | PR014721(100%) | | | | | | | | | | | |
| 52 | Variable 145 | DAALCLGTLK | Nocardia | Bacteria | Actinobacteria | | Actinomycetia | | Corynebacteriales | | Nocardiaceae | Nocardia | | | | | | | | | | | | | | IPR003774(100%) | 1.00E+00 | 1.00E+00 | 4.01E+04 | NS | NS | NS | NS | NS | NS | NS | NS |
| 53 | Variable 146 | SAVDVDAMR | Nocardia farcinica | Bacteria | Actinobacteria | | Actinomycetia | | Corynebacteriales | | Nocardiaceae | Nocardia | | | Nocardia farcinica | | | 5.4.99.12(100% | tRNA pseudouridine(38-40) synthase (100% | | | GO:0003723(100 | RNA binding (100%);tRNA pseudouridi | GO:0031119 (100%) | tRNA pseudouridine synthesis (1 | IPR001406(100% | 1.00E+00 | 1.00E+00 | 4.00E+04 | NS | NS | NS | NS | NS | NS | NS | NS |

| | | | | | | | | | | | | | | | | | | | | | | | | | | | | | | | | | | | | | |
|---|---|---|---|---|---|---|---|---|---|---|---|---|---|---|---|---|---|---|---|---|---|---|---|---|---|---|---|---|---|---|---|---|---|---|---|---|---|
| | | | | | | | | | | | | | | | | | | ) | ) | | | %);GO:0106029(100%) | ne synthase activity (100%) | | 00%) | );IPR020095(100%);IPR020097(100%) | | | | | | | | | | | |
| 54 | Variable 147 | AHAANAALLQPLGK | Nocardia farcinica | Bacteria | Actinobacteria | | Actinomycetia | | Corynebacteriales | | Nocardiaceae | Nocardia | | | Nocardia farcinica | | | 1.2.1.10(100 | Acetaldehyde dehydrogenase (acetylati | | | GO:0008774(1 | acetaldehyde dehydrogenase (ac | GO:0019439 (100%) | aromatic compound cata | IPR000534(10 | 1.00E+00 | 1.68E+05 | 1.00E+00 | NS | NS | NS | NS | NS | NS | NS | NS |

| | | | | | | | | | | | | | | | | | | | | | | | | | | | | | | | | | | | | | |
|---|---|---|---|---|---|---|---|---|---|---|---|---|---|---|---|---|---|---|---|---|---|---|---|---|---|---|---|---|---|---|---|---|---|---|---|---|---|
| | | | | | | | | | | | | | | | | | | %) | ng)(100%) | | | 00%);GO:0051287(100%) | etylating) activity (100%);NAD binding (100%) | | bolic process (100%) | 0%);IPR003361(100%);IPR015426(100%) | | | | | | | | | | | |
| 55 | Variable 1 | GLPHEARPLR | Nocardiafarcinic | Bacteria | Actinobacte | | Actinomycetia | | Corynebacteriales | | Nocardiaceae | Nocardia | | | Nocardiafarcin | | | 4.2.1.109( | Methylthioribulose 1-phosphat | GO:0005737 | cytoplasm(100%) | GO:0008270 | zinc ion binding (100%);S- | GO:0019284 (100%);G | L-methionine salvag | IPR000649( | 1.00E+00 | 5.90E+04 | 1.00E+00 | NS | NS | NS | NS | NS | NS | NS | NS |

| | | | | | | | | | | | | | | | | | | | | | | | | | | |
|---|---|---|---|---|---|---|---|---|---|---|---|---|---|---|---|---|---|---|---|---|---|---|---|---|---|---|
| | 48 | | a | | ria | | | | | | | | | | ica | | | 100%);5.3.1.23(100%) | e dehydratase (100%);S-methyl-5-thioribose-1-phosphate isomerase (100%) | (100%) | | (100%);GO:0046523(100%);GO:0046570(100%) | methyl-5-thioribose-1-phosphate isomerase activity (100%);methylthioribulose 1-phosphate dehydratase activity (10 | O:0019509(100%) | e from S-adenosylmethionine (100%);L-methionine salvage from methylthioadenosine | 100%);IPR001303(100%);IPR005251(100%) |

| | | | | | | | | | | | | | | | | | | | | | | | | | | | | | | | | | | | | | |
|---|---|---|---|---|---|---|---|---|---|---|---|---|---|---|---|---|---|---|---|---|---|---|---|---|---|---|---|---|---|---|---|---|---|---|---|---|---|
| | | | | | | | | | | | | | | | | | | | | | | | 0%) | | (100%) | | | | | | | | | | | | |
| 56 | Variable 149 | AVVAEELASLR | Rhodococcus | Bacteria | Actinobacteria | | Actinomycetia | | Corynebacteriales | | Nocardiaceae | Rhodococcus | | | | | | | | | | GO:0003824(100%);GO:0015420(100%) | catalytic activity (100%);ABC-type vitamin B12 transporter activity (100%) | GO:0006541(100%);GO:0009236(100%) | glutamine metabolic process (100%);cobalamin biosynthetic process (100%) | IPR002586(100%);IPR004459(100%);IPR011698 | 1.00E+00 | 1.22E+05 | 1.00E+00 | NS | NS | NS | NS | NS | NS | NS | NS |

| | | | | | | | | | | | | | | | | | | | | | | | | | | | | | | | | | | | | | |
|---|---|---|---|---|---|---|---|---|---|---|---|---|---|---|---|---|---|---|---|---|---|---|---|---|---|---|---|---|---|---|---|---|---|---|---|---|---|
| | | | | | | | | | | | | | | | | | | | | | | | | | | (100%) | | | | | | | | | | | |
| 57 | Variable 15 | GEVDLDTDDEK | Actinomycetia | Bacteria | Actinobacteria | | Actinomycetia | | | | | | | | | | | | | GO:0005737(100%) | cytoplasm(100%) | GO:0005524(100%);GO:0016887(100%);GO:00 | ATP binding(100%);ATPase(100%);unfolded protein binding(100%) | GO:0006457(100%) | protein folding(100%) | IPR001404(100%);IPR003594(100%);IPR0198 | 1.00E+00 | 1.00E+00 | 6.81E+04 | NS | NS | NS | NS | NS | NS | NS | NS |

| | | | | | | | | | | | | | | | | | | | | | | | | | | | | | | | | | | | | | | | | |
|---|---|---|---|---|---|---|---|---|---|---|---|---|---|---|---|---|---|---|---|---|---|---|---|---|---|---|---|---|---|---|---|---|---|---|---|---|---|---|---|---|
| | | | | | | | | | | | | | | | | | | | | | | 51082(100%) | | | | 05(100%) | | | | | | | | | | | | | | |
| 58 | Variable 150 | AGPTGQLMNSYLGR | Rhodococcus opacus | Bacteria | Actinobacteria | | Actinomycetia | | Corynebacteriales | | Nocardiaceae | Rhodococcus | | | Rhodococcus opacus | | | 1.13.11.-(100%) | With incorporation of two atoms of oxygen (100%) | | | GO:0008199(100%);GO:0016702(50%);G | ferric iron binding (100%);oxidoreductase activity | acting on single donors with incorporation of molecular oxygen | incorporation of two atoms of oxygen (50%);chlorocatechol 1 | 2-dioxygenase activity (50%) | 1.00E+00 | 1.00E+00 | 5.27E+04 | 0.27101 | 1.0459E-11 | 0.0026585 | 0.9 | NS | NS | NS | NS | 33.834 | 1.76E-09 | 0.0056748 |

| | | | | | | | | | | | | | | | | | | | | | | | | | | | | | | | | | | | | | |
|---|---|---|---|---|---|---|---|---|---|---|---|---|---|---|---|---|---|---|---|---|---|---|---|---|---|---|---|---|---|---|---|---|---|---|---|---|---|
| | | | | | | | | | | | | | | | | | | | | | | O:0018575(50%) | | | | | | | | | | | | | | | |
| 59 | Variable 151 | MAEDLHLPSR | Frankia | Bacteria | Actinobacteria | | Actinomycetia | | Frankiales | | Frankiaceae | Frankia | | | | | | 6.3.2.1(100%) | Pantoate--beta-alanine ligase (AMP-forming)(100%) | GO:0005737(100%) | cytoplasm(100%) | GO:0004592(100%);GO:0005524(10 | pantoate-beta-alanine ligase activity(100%);ATP binding(100%) | GO:0015940(100%) | pantothenate biosynthetic process(100%) | IPR003721(100%);IPR014729(100% | 1.00E+00 | 1.00E+00 | 7.45E+05 | NS | NS | NS | NS | NS | NS | NS | NS |

| | | | | | | | | | | | | | | | | | | | | | | | | | | | | | | | | | | | | | |
|---|---|---|---|---|---|---|---|---|---|---|---|---|---|---|---|---|---|---|---|---|---|---|---|---|---|---|---|---|---|---|---|---|---|---|---|---|---|
| | | | | | | | | | | | | | | | | | | | | | | 0%) | | | | );IPR0421776(100%) | | | | | | | | | | | |
| 60 | Variable 152 | EDDLFAGLVP | FrankiaalniACN14a | Bacteria | Actinobacteria | | Actinomycetia | | Frankiales | | Frankiaceae | Frankia | | | Frankiaalni | | FrankiaalniACN14a | 3.5.1.103(100%) | N-acetyl-1-D-myo-inositol-2-amino-2-deoxy-alpha-D-glucopyranoside deacetylase | | | GO:0008270(100%);GO:0035595( | zinc ion binding(100%);N-acetylglucosaminylinositol deacetylase activity (10 | GO:0010125(100%) | mycothiol biosynthetic process (100%) | IPR003737(100%);IPR017810(10 | 1.00E+00 | 2.21E+06 | 1.00E+00 | NS | NS | NS | NS | NS | NS | NS | NS |

| | | | | | | | | | | | | | | | | | | | | | | | | | | | | | | | | | | | | | |
|---|---|---|---|---|---|---|---|---|---|---|---|---|---|---|---|---|---|---|---|---|---|---|---|---|---|---|---|---|---|---|---|---|---|---|---|---|---|
| | | | | | | | | | | | | | | | | | | | (100%) | | | 100%) | 0%) | | | 0%);IPR024078(100%) | | | | | | | | | | | |
| 61 | Variable 153 | GVLALGGAVLDAENR | Frankiacasuarinae | Bacteria | Actinobacteria | | Actinomycetia | | Frankiales | | Frankiaceae | Frankia | | | Frankiacasuarinae | | | 2.7.1.71(100%) | Shikimate kinase (100%) | GO:0005737(100%) | cytoplasm(100%) | GO:0000287(100%);GO:000476 | magnesium ion binding (100%);shikimate kinase activity (100%);ATP bindin | GO:0008652(100%);GO:0009073(100%);GO:0009423(1 | cellular amino acid biosynthetic process(100%);aromati | IPR000623(100%);IPR023000( | 1.00E+00 | 1.00E+00 | 4.81E+05 | NS | NS | NS | NS | NS | NS | NS | NS |

| | | | | | | | | | | | | | | | | | | | | | | | | | | | | | | | | | | | | | |
|---|---|---|---|---|---|---|---|---|---|---|---|---|---|---|---|---|---|---|---|---|---|---|---|---|---|---|---|---|---|---|---|---|---|---|---|---|---|
| | | | | | | | | | | | | | | | | | | | | | | 5(100%);GO:0005524(100%) | g(100%) | 00%) | c amino acid family biosynthetic process(100%);chorismate biosynthetic process(100%) | 100%);IPR027417(100%) | | | | | | | | | | | |
| 6 | V | QA | F | B | A | | A | | Fr | | Fr | F | | | F | | | 3 | Act | G | cy | G | me | G | rR | I | 1 | 1 | 1 | N | N | N | N | N | N | N | N |

| | | | | | | | | | | | | | | | | | | | | | | | | | | | | | | | | | | | | | |
|---|---|---|---|---|---|---|---|---|---|---|---|---|---|---|---|---|---|---|---|---|---|---|---|---|---|---|---|---|---|---|---|---|---|---|---|---|---|
| 2 | ariable 154 | EQAGHSFDR | rankiacasuarinae | acteria | ctinobacteria | | ctinomycetia | | ankiales | | ankiaceae | rankia | | | rankiacasuarinae | | | .1.-.-(100%) | ingonesterbonds(100%) | O:0005737(100%) | toplasm(100%) | O:0004222(100%);GO:0004521(100%);GO:0008270(1 | talloendopeptidase activity(100%);endoribonuclease activity(100%);zinc ion binding(100%) | O:0006364(100%) | NA processing(100%) | PR002036(100%);IPR020549(100%);IPR023091(100% | .00E+00 | .00E+00 | .79E+05 | S | S | S | S | S | S | S | S |

| | | | | | | | | | | | | | | | | | | | | | | | | | | | | | | | | | | | | | | | | |
|---|---|---|---|---|---|---|---|---|---|---|---|---|---|---|---|---|---|---|---|---|---|---|---|---|---|---|---|---|---|---|---|---|---|---|---|---|---|---|---|---|
| | | | | | | | | | | | | | | | | | | | | | | 00%) | | | | ) | | | | | | | | | | | | | | |
| 63 | Variable 155 | MDLSATLNSLVSK | Frankiacasuarinae | Bacteria | Actinobacteria | | Actinomycetia | | Frankiales | | Frankiaceae | Frankia | | | Frankiacasuarinae | | | 2.8.1.6(100%) | Biotin synthase (100%) | | | GO:0004076(100%);GO:0005506(100%);GO:0051 | biotin synthase activity (100%);iron ion binding (100%);2 iron | 2 sulfur cluster binding (100%) | GO:0009102(100%) | biotin biosynthetic process(100%) | 1.00E+00 | 1.00E+00 | 1.92E+05 | 27.44 | 1.895E-06 | 0.00043431 | 0.9 | NS | NS | NS | NS | 9.7646 | 3.29E-04 | 0.0071566 |

| | | | | | | | | | | | | | | | | | | | | | | | | | | | | | | | | | | | | | | | | |
|---|---|---|---|---|---|---|---|---|---|---|---|---|---|---|---|---|---|---|---|---|---|---|---|---|---|---|---|---|---|---|---|---|---|---|---|---|---|---|---|---|
| | | | | | | | | | | | | | | | | | | | | | | 537(100%) | | | | | | | | | | | | | | | | | | |
| 64 | Variable 156 | ALVDNAGDFEK | Frankiacasuarinae | Bacteria | Actinobacteria | | Actinomycetia | | Frankiales | | Frankiaceae | Frankia | | | Frankiacasuarinae | | | | | GO:0005737(100%) | cytoplasm(100%) | GO:0003746(100%) | translation elongation factor activity(100%) | | | IPR001816(100%);IPR009060(100%);IPR0 | 1.00E+00 | 1.00E+00 | 4.81E+04 | 0.42075 | 5.8296E-09 | 0.0016606 | 0.9 | NS | NS | NS | NS | 12.863 | 4.39E-05 | 0.0003351 |

| | | | | | | | | | | | | | | | | | | | | | | | | | | | | | | | | | | | | | |
|---|---|---|---|---|---|---|---|---|---|---|---|---|---|---|---|---|---|---|---|---|---|---|---|---|---|---|---|---|---|---|---|---|---|---|---|---|---|
| | | | | | | | | | | | | | | | | | | | | | | | | | | 14039(100%) | | | | | | | | | | | |
| 65 | Variable 157 | GLDPADAELTAVVAAYLGGSLR | Kineococcus radiotolerans SRS30216=ATCCB | Bacteria | Actinobacteria | | Actinomycetia | | Kineosporiales | | Kineosporiaceae | Kineococcus | | | Kineococcus radiotolerans | | Kineococcus radiotolerans SRS30216=AT | 6.5.1.1(100%) | DNA ligase (ATP) (100%) | | | GO:0003677(100%);GO:0003910(100%); | DNA binding(100%);DNA ligase (ATP) activity(100%);ATP binding(100%) | GO:0006260(100%);GO:0006281(100%);GO:0006310(100%) | DNA replication(100%);DNA repair(100%);DNA recombination | IPR012308(100%);IPR012309(100%);IP | 1.00E+00 | 1.00E+00 | 3.59E+05 | NS | NS | NS | NS | NS | NS | NS | NS |

| | | | | | | | | | | | | | | | | | | | | | | | | | | | | | | | | | | | | | |
|---|---|---|---|---|---|---|---|---|---|---|---|---|---|---|---|---|---|---|---|---|---|---|---|---|---|---|---|---|---|---|---|---|---|---|---|---|---|
| | | | AA-149 | | | | | | | | | | | | | | CCBAA-149 | | | | | GO:0005524(100%) | | | (100%) | R0123100(100%) | | | | | | | | | | | |
| 66 | Variable 158 | ASPETVQLAR | Kineococcus radiotolerans SRS30216= | Bacteria | Actinobacteria | | Actinomycetia | | Kineosporiales | | Kineosporiaceae | Kineococcus | | | Kineococcus radiotolerans | | Kineococcus radiotolerans SRS302 | | | GO:0005886(100%);GO:0045261( | plasma membrane (100%);proton-transporting ATP synt | catalytic core F(1)(100%) | GO:0046933(100%) | proton-transporting ATP synthase activity | rotational mechanism(100%) | | 1.00E+00 | 2.28E+05 | 1.00E+00 | NS | NS | NS | NS | NS | NS | NS | NS |

| | | | | | | | | | | | | | | | | | | | | | | | | | | | | | | | | | | | | | | | | |
|---|---|---|---|---|---|---|---|---|---|---|---|---|---|---|---|---|---|---|---|---|---|---|---|---|---|---|---|---|---|---|---|---|---|---|---|---|---|---|---|---|
| | | | ATCC BAA-149 | | | | | | | | | | | | | | 16 = ATCC BAA-149 | | | 100%) | hase complex | | | | | | | | | | | | | | | | | | | |
| 67 | Variable 159 | HAGGAGPVQVGDTWSR | Beutenbergia cavernae DSM 12333 | Bacteria | Actinobacteria | | Actinomycetia | | Micrococcales | | Beutenbergiaceae | Beutenbergia | | | Beutenbergia cavernae | | Beutenbergia cavernae DSM 12333 | 5.1.1.1 (100%) | Alanine racemase (100%) | | | GO:0008784(100%);GO:0030170(10 | alanine racemase activity(100%);pyridoxal phosphate binding (100%) | GO:0030632 (100%) | D-alanine biosynthetic process (100%) | IPR000821(100%);IPR001608(100% | 1.00E+00 | 1.00E+00 | 3.35E+04 | 0.5486 | 1.306E-07 | 0.0013514 | 0.9 | NS | NS | NS | NS | 7.9025 | 1.22E-03 | 0.0086593 |

| | | | | | | | | | | | | | | | | | | | | | | | | | | | | | | | | | | | | | |
|---|---|---|---|---|---|---|---|---|---|---|---|---|---|---|---|---|---|---|---|---|---|---|---|---|---|---|---|---|---|---|---|---|---|---|---|---|---|
| | | | | | | | | | | | | | | | | | | | | | | 0%) | | | | );IPR009006(100%) | | | | | | | | | | | |
| 68 | Variable 16 | EFVMPTHCPECGTELAPAK | Actinomycetia | Bacteria | Actinobacteria | | Actinomycetia | | | | | | | | | | | 6.5.1.2(100%) | DNA ligase (NAD(+)) (100%) | | | GO:0003911(100%);GO:0046872( | DNA ligase (NAD+) activity (100%);metal ion binding (100%);dioxygena | GO:0006260(100%);GO:0006281(100%) | DNA replication (100%); DNA repair (100%) | IPR001357(100%);IPR001679(10 | 1.00E+00 | 1.00E+00 | 5.89E+04 | 2.8392 | 0.02724 1 | 0 | 0.9 | NS | NS | NS | NS |

| | | | | | | | | | | | | | | | | | | | | | | | | | | | | | | | | | | | | | |
|---|---|---|---|---|---|---|---|---|---|---|---|---|---|---|---|---|---|---|---|---|---|---|---|---|---|---|---|---|---|---|---|---|---|---|---|---|---|
| | | | | | | | | | | | | | | | | | | | | | | 100%);GO:0051213(3%) | se activity (3%) | | | 0%);IPR004149(100%) | | | | | | | | | | | |
| 70 | Variable 161 | FLAEAPGMLEAFK | Clavibactermichiganensis | Bacteria | Actinobacteria | | Actinomycetia | | Micrococcales | | Microbacteriaceae | Clavibacter | | | Clavibactermichiganensis | | | 6.1.1.4(100%) | Leucine--tRNA ligase (100%) | GO:0005737(89%) | cytoplasm(89%) | GO:0004823(100%);GO:00055 | leucine-tRNA ligase activity (100%);ATP binding (100%);aminoacy | GO:0006429(100%) | leucyl-tRNA aminoacylation (100%) | IPR002300(100%);IPR002302 | 1.00E+00 | 1.00E+00 | 3.40E+04 | NS | NS | NS | NS | NS | NS | NS | NS |

| | | | | | | | | | | | | | | | | | | | | | | | | | | | | | | | | | | | | | |
|---|---|---|---|---|---|---|---|---|---|---|---|---|---|---|---|---|---|---|---|---|---|---|---|---|---|---|---|---|---|---|---|---|---|---|---|---|---|
| | | | | | | | | | | | | | | | | | | | | | | 24(100%);GO:0002161(95%) | l-tRNA editing activity(95%) | | | (100%);IPR009080(100%) | | | | | | | | | | | |
| 71 | Variable 162 | VATGVPATPVEVAAVDALLGVALGSPAA | Clavibacter michiganensis | Bacteria | Actinobacteria | | Actinomycetia | | Micrococcales | | Microbacteriaceae | Clavibacter | | | Clavibacter michiganensis | | | 6.3.1.13(100%) | L-cysteine:1D-myo-inositol 2-amino-2-deoxy-alpha-D-glu | | | GO:0005524(100%);GO:0 | ATP binding(100%);zinc ion binding(88%);cysteine-glucos | GO:0010125(88%) | mycothiol biosynthetic process(88%) | IPR014729(100%);IPR02 | 1.00E+00 | 1.00E+00 | 2.25E+05 | 14.442 | 0.0051247 | 0.0001806 | 0.9 | NS | NS | NS | NS |

| | | | | | | | | | | | | | | | | | | | | | | | | | | | | | | | | | | | | | | | | |
|---|---|---|---|---|---|---|---|---|---|---|---|---|---|---|---|---|---|---|---|---|---|---|---|---|---|---|---|---|---|---|---|---|---|---|---|---|---|---|---|---|
| | | | | | | | | | | | | | | | | | | | copyranosidligase (100%) | | | 0082700(88%);GO:0035446(88%) | aminylinositol ligase activity (88%) | | | 49009(100%);IPR032678(100%) | | | | | | | | | | | | | | |
| 72 | Variable 163 | VAAGYDGDLAALSR | Leifsoniaxyli | Bacteria | Actinobacteria | | Actinomycetia | | Micrococcales | | Microbacteriaceae | Leifsonia | | | Leifsoniaxyli | | | 6.1.1.19(100%) | Arginine--tRNA ligase (100%) | GO:0005737(100%) | cytoplasm (100%) | GO:0004814(100%);G | arginine-tRNA ligase activity (100%);ATP bindin | GO:0006420 (100%) | arginyl-tRNA aminoacylation (100% | IPR001278(100%);IP | 1.00E+00 | 1.00E+00 | 7.90E+04 | 0.58126 | 1.0251E-06 | 0.00224 | 0.9 | NS | NS | NS | NS | 6.5618 | 3.31E-03 | 0.0005556 |

| | | | | | | | | | | | | | | | | | | | | | | | | | | | | | | | | | | | | | |
|---|---|---|---|---|---|---|---|---|---|---|---|---|---|---|---|---|---|---|---|---|---|---|---|---|---|---|---|---|---|---|---|---|---|---|---|---|---|
| | | | | | | | | | | | | | | | | | | | | | | O:0005524(100%) | g(100%) | | ) | R001412(100%);IPR005148(100%) | | | | | | | | | | | |
| 73 | Variable 164 | WVDFFVNK | Micrococcaceae | Bacteria | Actinobacteria | | Actinomycetia | | Micrococcales | | Micrococcaceae | | | | | | | 6.1.1.7(100%) | Alanine--tRNA ligase (100%) | GO:0005737(100% | cytoplasm(100%) | GO:0000049(100% | tRNA binding(100%);alanine-tRNA lig | GO:0006419(100%) | alanyl-tRNA aminoacylation(1 | IPR002318(88%);I | 1.00E+00 | 1.00E+00 | 7.44E+04 | NS | NS | NS | NS | NS | NS | NS | NS |

| | | | | | | | | | | | | | | | | | | | | | | | | | | | | | | | | | | | | | |
|---|---|---|---|---|---|---|---|---|---|---|---|---|---|---|---|---|---|---|---|---|---|---|---|---|---|---|---|---|---|---|---|---|---|---|---|---|---|
| | | | | | | | | | | | | | | | | | | | | ) | | );GO:0004813(100%);GO:0005524(100%) | ase activity (100%);ATP binding (100%) | | 00%) | PR003156(88%);IPR009000(88%) | | | | | | | | | | | |
| 74 | Variable 1 | DEVLGALAGQFEGR | Micrococcac | Bacteria | Actinobacte | | Actinomycetia | | Micrococcales | | Micrococcaceae | | | | | | | 2.7.7.8(10 | Polyribonucleotide nucleotid | GO:0005737 | cytoplasm (100%) | GO:0000287 | magnesium ion binding (100% | GO:0006396(100%);G | RNA processing (100 | IPR001247( | 1.00E+00 | 1.00E+00 | 2.55E+04 | NS | NS | NS | NS | NS | NS | NS | NS |

| | | | | | | | | | | | | | | | | | | | | | | | | | | | | | | | | | | | | | | | | |
|---|---|---|---|---|---|---|---|---|---|---|---|---|---|---|---|---|---|---|---|---|---|---|---|---|---|---|---|---|---|---|---|---|---|---|---|---|---|---|---|---|
| | 65 | | eae | | ria | | | | | | | | | | | | | 0%) | yltransferase(100%) | (100%) | | (100%);GO:0003723(100%);GO:0004654(100%) | );RNA binding(100%);polyribonucleotide nucleotidyltransferase activity(100%) | O:0006402(100%) | %);mRNA catabolic process(100%) | 100%);IPR003029(100%);IPR004087(100%) | | | | | | | | | | | | | | |
| 75 | Varia | QYSAAWSGGM | Pseud | Bacte | Actino | | Actinom | | Micrococc | | Micrococc | Pseud | | | Pseud | | Pseud | 2.7.7 | Nucleotidyltra | | | GO:00 | magnesium ion | | | IPR00 | 1.00E | 1.00E | 3.85E | 4.494 | 6.473 | 0.0001 | 0.9 | NS | NS | NS | NS | 8.185 | 9.96E | 0.000 |

| | | | | | | | | | | | | | | | | | | | | | | | | | | | | | | | | | | | | | | | | |
|---|---|---|---|---|---|---|---|---|---|---|---|---|---|---|---|---|---|---|---|---|---|---|---|---|---|---|---|---|---|---|---|---|---|---|---|---|---|---|---|---|
| | ble 166 | ATK | arthrobacterchlorophenolicusA6 | ria | bacteria | | ycetia | | ales | | aceae | arthrobacter | | | arthrobacterchlorophenolicus | | arthrobacterchlorophenolicusA6 | .-(100%);2.7.7.0(100%) | nsferases(100%);(100%) | | | 00287(100%);GO:0005524(100%);GO:0016779(100%) | binding(100%);ATP binding(100%);nucleotidyltransferase activity(100%) | | | 3846(100%) | +00 | +00 | +04 | 3 | 9E-06 | 7876 | | | | | | 6 | -04 | 50466 |

| | | | | | | | | | | | | | | | | | | | | | | | | | | | | | | | | | | | | | |
|---|---|---|---|---|---|---|---|---|---|---|---|---|---|---|---|---|---|---|---|---|---|---|---|---|---|---|---|---|---|---|---|---|---|---|---|---|---|
| 76 | Variable 167 | FLADLEANSK | Renibacterium salmoninarum ATCC 33209 | Bacteria | Actinobacteria | | Actinomycetia | | Micrococcales | | Micrococcaceae | Renibacterium | | | Renibacterium salmoninarum | | Renibacterium salmoninarum ATCC 33209 | | | GO:0005886(100%);GO:0016021(100%);GO:0045263( | plasma membrane(100%);integral component of membrane(100%);proton-transporting AT | coupling factor F(o)(100%) | GO:0046933(100%) | proton-transporting ATP synthase activity | rotational mechanism(100%) | | 1.00E+00 | 1.00E+00 | 3.46E+04 | NS | NS | NS | NS | NS | NS | NS | NS |

| | | | | | | | | | | | | | | | | | | | | | | | | | | | | | | | | | | | | | |
|---|---|---|---|---|---|---|---|---|---|---|---|---|---|---|---|---|---|---|---|---|---|---|---|---|---|---|---|---|---|---|---|---|---|---|---|---|---|
| | | | | | | | | | | | | | | | | | | | | 100%) | P synthase complex | | | | | | | | | | | | | | | | |
| 77 | Variable 168 | QFWSHMVDK | Renibacterium salmoninarum ATCC 33209 | Bacteria | Actinobacteria | | Actinomycetia | | Micrococcales | | Micrococcaceae | Renibacterium | | | Renibacterium salmoninarum | | Renibacterium salmoninarum ATCC 33209 | 1.1.5.4 (100%) | Malate dehydrogenase (quinone) (100%) | | | GO:0008924(100%);GO:0052589(100%) | malate dehydrogenase (quinone) activity (100%);malate dehydrogenase (menaquinone) activity | GO:0006099 (100%) | tricarboxylic acid cycle (100%) | IPR006231(100%);IPR036188(100%) | 1.00E+00 | 5.09E+04 | 1.00E+00 | NS | NS | NS | NS | NS | NS | NS | NS |

| | | | | | | | | | | | | | | | | | | | | | | | | | | | | | | | | | | | | | |
|---|---|---|---|---|---|---|---|---|---|---|---|---|---|---|---|---|---|---|---|---|---|---|---|---|---|---|---|---|---|---|---|---|---|---|---|---|---|
| | | | | | | | | | | | | | | | | | | | | | | | (100%) | | | | | | | | | | | | | | |
| 78 | Variable 169 | FFYGSVGPDAEAAR | Sinomonas | Bacteria | Actinobacteria | | Actinomycetia | | Micrococcales | | Micrococcaceae | Sinomonas | | | | | | | | | | GO:0016740(100%) | transferase activity (100%) | | | IPR006222(100%);IPR028896(100%);IPR029043( | 1.00E+00 | 1.00E+00 | 8.04E+04 | 2.6099 | 0.045445 | 0.000029412 | 0.9 | NS | NS | NS | NS |

| | | | | | | | | | | | | | | | | | | | | | | | | | | | | | | | | | | | | | | | | |
|---|---|---|---|---|---|---|---|---|---|---|---|---|---|---|---|---|---|---|---|---|---|---|---|---|---|---|---|---|---|---|---|---|---|---|---|---|---|---|---|---|
| | | | | | | | | | | | | | | | | | | | | | | | | | | 100%) | | | | | | | | | | | | | | |
| 79 | Variable 17 | YWMHNGWVTMSGEK | Actinomycetia | Bacteria | Actinobacteria | | Actinomycetia | | | | | | | | | | | 6.1.1.16(100%) | Cysteine--tRNA ligase (100%) | GO:0005737(100%) | cytoplasm (100%) | GO:0004817(100%);GO:0005524(100%);GO:000 | cysteine-tRNA ligase activity (100%);ATP binding (100%);zinc ion binding (100%) | GO:0006423 (100%) | cysteinyl-tRNA aminoacylation (100%) | IPR009080(100%);IPR014729(100%);IPR015273 | 1.00E+00 | 1.00E+00 | 6.88E+04 | 8.2759 | 0.000031993 | 0.0004506 | 0.9 | NS | NS | NS | NS | 6.2665 | 4.15E-03 | 0.00072153 |

| | | | | | | | | | | | | | | | | | | | | | | | | | | | | | | | | | | | | | |
|---|---|---|---|---|---|---|---|---|---|---|---|---|---|---|---|---|---|---|---|---|---|---|---|---|---|---|---|---|---|---|---|---|---|---|---|---|---|
| | | | | | | | | | | | | | | | | | | | | | | 8270(100%) | | | | 3(100%) | | | | | | | | | | | |
| 81 | Variable 171 | GAVMSVPGHDTR | Tropherymawhipplei | Bacteria | Actinobacteria | | Actinomycetia | | Micrococcales | | Tropherymataceae | Tropheryma | | | Tropherymawhipplei | | | 6.1.1.4(100%) | Leucine--tRNA ligase(100%) | GO:0005737(100%) | cytoplasm(100%) | GO:0002161(100%);GO:0004823(100%);G | aminoacyl-tRNA editing activity(100%);leucine-tRNA ligase activity(100%);ATP binding | GO:0006429(100%) | leucyl-tRNA aminoacylation(100%) | IPR001412(100%);IPR002300(100%);IPR | 2.48E+05 | 1.00E+00 | 1.00E+00 | NS | NS | NS | NS | NS | NS | NS | NS |

| | | | | | | | | | | | | | | | | | | | | | | | | | | | | | | | | | | | | | |
|---|---|---|---|---|---|---|---|---|---|---|---|---|---|---|---|---|---|---|---|---|---|---|---|---|---|---|---|---|---|---|---|---|---|---|---|---|---|
| | | | | | | | | | | | | | | | | | | | | | | O:0005524(100%) | (100%) | | | 0023002(100%) | | | | | | | | | | | |
| 82 | Variable 172 | DGVVVVNDLTVTSLR | Tropherymawhippleistr.Twist | Bacteria | Actinobacteria | | Actinomycetia | | Micrococcales | | Tropherymataceae | Tropheryma | | | Tropherymawhipplei | | Tropherymawhipplei str. Twist | | | GO:0005737(100%) | cytoplasm(100%) | GO:0003743(100%);GO:0003924(1 | translation initiation factor activity(100%);GTPase activity(100%);GTP binding(10 | | | IPR000178(100%);IPR000795(100 | 1.00E+00 | 1.00E+00 | 3.08E+04 | NS | NS | NS | NS | NS | NS | NS | NS |

| | | | | | | | | | | | | | | | | | | | | | | | | | | | | | | | | | | | | | |
|---|---|---|---|---|---|---|---|---|---|---|---|---|---|---|---|---|---|---|---|---|---|---|---|---|---|---|---|---|---|---|---|---|---|---|---|---|---|
| | | | | | | | | | | | | | | | | | | | | | | 00%);GO:0005525(100%) | 0%) | | | %);IPR005225(100%) | | | | | | | | | | | |
| 84 | Variable 174 | WADALAAVTLDADAPR | Nakamurella multipartita DSM 442 | Bacteria | Actinobacteria | | Actinomycetia | | Nakamurellales | | Nakamurellaceae | Nakamurella | | | Nakamurella multipartita | | Nakamurella multipartita DSM 44 | 2.4.1.250(100%) | D-inositol-3-phosphate glycosyltransferase (100%) | | | GO:0000287(100%);GO:0008 | magnesium ion binding (100%);acetylglucosaminyltransferase activit | GO:0010125 (100%) | mycothiol biosynthetic process (100%) | IPR001296(100%);IPR01781 | 1.00E+00 | 1.00E+05 | 1.00E+00 | NS | NS | NS | NS | NS | NS | NS | NS |

| | | | | | | | | | | | | | | | | | | | | | | | | | | | | | | | | | | | | | | | | |
|---|---|---|---|---|---|---|---|---|---|---|---|---|---|---|---|---|---|---|---|---|---|---|---|---|---|---|---|---|---|---|---|---|---|---|---|---|---|---|---|---|
| | | | 33 | | | | | | | | | | | | | | 233 | | | | | 375(100%);GO:0102710(100%) | y(100%);D-inositol-3-phosphate glycosyltransferase activity(100%) | | | 4(100%);IPR028098(100%) | | | | | | | | | | | | | | |
| 85 | Variable 175 | GEVPSNEGMLPTR | Kribbella flavida DSM 178 | Bacteria | Actinobacteria | | Actinomycetia | | Propionibacteriales | | Kribbellaceae | Kribbella | | | Kribbella flavida | | Kribbella flavida DSM 17 | 3.2.1.205(100%) | Isomaltose glucohydrolase(100%) | GO:0005737(100%) | cytoplasm(100%) | GO:0016798(100%) | hydrolase activity | acting on glycosyl bonds(100%) | GO:0000272(100%) | polysaccharide catabolic p | 1.00E+00 | 1.00E+00 | 2.29E+04 | 3.1953 | 0.00046529 | 0.00086601 | 0.9 | NS | NS | NS | NS | 6.1697 | 4.48E-03 | 0.0017895 |

| | | | | | | | | | | | | | | | | | | | | | | | | | | | | | | | | | | | | | |
|---|---|---|---|---|---|---|---|---|---|---|---|---|---|---|---|---|---|---|---|---|---|---|---|---|---|---|---|---|---|---|---|---|---|---|---|---|---|
| | | | 36 | | | | | | | | | | | | | | 836 | | | | | | | | | rocess(100%) | | | | | | | | | | | |
| 86 | Variable 176 | LASPLLDQALR | Kribbellaflavida DSM 17836 | Bacteria | Actinobacteria | | Actinomycetia | | Propionibacteriales | | Kribbellaceae | Kribbella | | | Kribbella flavida | | Kribbella flavida DSM 17836 | | | GO:0000502(100%) | proteasome complex(100%) | GO:0005524(100%);GO:0016887(100%) | ATP binding(100%);ATPase(100%) | GO:0010498(100%);GO:0019941(100%) | proteasomal protein catabolic process(100%);modification-dependen | IPR003593(100%);IPR003959(100%);IP | 1.00E+00 | 1.00E+00 | 1.48E+06 | 7.6745 | 0.00051974 | 0.00068995 | 0.9 | NS | NS | NS | NS |

| | | | | | | | | | | | | | | | | | | | | | | | | | | | | | | | | | | | | | |
|---|---|---|---|---|---|---|---|---|---|---|---|---|---|---|---|---|---|---|---|---|---|---|---|---|---|---|---|---|---|---|---|---|---|---|---|---|---|
| | | | | | | | | | | | | | | | | | | | | | | | | | t protein catabolic process (100%) | R003960 (100%) | | | | | | | | | | | |
| 87 | Variable 177 | GASVVPCDFSLGMLR | Nocardioides | Bacteria | Actinobacteria | | Actinomycetia | | Propionibacteriales | | Nocardioidaceae | Nocardioides | | | | | | 2.1.1.163 (100%) | Demethylmenaquinone methyltransferase (100%) | | | GO:0043770 (100%);GO:0102027 | demethylmenaquinone methyltransferase activity (100%);S-adenosylmethi | GO:0009234 (100%);GO:0032259 (100%) | menaquinone biosynthetic process (100%); methylation | IPR004033 (100%);IPR023576 (1 | 1.00E+00 | 2.18E+05 | 1.00E+00 | NS | NS | NS | NS | NS | NS | NS | NS |

| | | | | | | | | | | | | | | | | | | | | | | | | | | |
|---|---|---|---|---|---|---|---|---|---|---|---|---|---|---|---|---|---|---|---|---|---|---|---|---|---|---|
| | | | | | | | | | | | | | | | | | | | | | | (100%);GO:0102094(100%) | onine:2-demethylquinol-8methyltransferase activity (100%);S-adenosylmethionine:2-demethylmenaquinolmethyltransferase | | (100%) | 00%);IPR029063(100%) |

| | | | | | | | | | | | | | | | | | | | | | | | | | | | | | | | | | | | | | |
|---|---|---|---|---|---|---|---|---|---|---|---|---|---|---|---|---|---|---|---|---|---|---|---|---|---|---|---|---|---|---|---|---|---|---|---|---|---|
| | | | | | | | | | | | | | | | | | | | | | | | activity (100%) | | | | | | | | | | | | | | |
| 88 | Variable 178 | AGSGVGTGGMTTK | Propionibacteriaceae | Bacteria | Actinobacteria | | Actinomycetia | | Propionibacteriales | | Propionibacteriaceae | | | | | | | 2.7.2.11(100%) | Glutamate 5-kinase (100%) | GO:0005737(95%) | cytoplasm(95%) | GO:0004349(100%);GO:0005524(95%);GO:000 | glutamate 5-kinase activity (100%);ATP binding (95%);RNA binding (90%) | GO:0055129 (85%);GO:0006561 (15%) | L-proline biosynthetic process (85%); proline biosynthetic process (15%) | IPR001048(100%);IPR036393(100%);IPR0010 | 1.00E+00 | 1.00E+00 | 8.94E+04 | 4.058 | 0.0013198 | 0.00013333 | 0.9 | NS | NS | NS | NS |

| | | | | | | | | | | | | | | | | | | | | | | | | | | | | | | | | | | | | | | | | |
|---|---|---|---|---|---|---|---|---|---|---|---|---|---|---|---|---|---|---|---|---|---|---|---|---|---|---|---|---|---|---|---|---|---|---|---|---|---|---|---|---|
| | | | | | | | | | | | | | | | | | | | | | | 3723 (90%) | | | | 57 (95%) | | | | | | | | | | | | | | |
| 89 | Variable 179 | ASTTALQLASGQLGER | Tessaracoccus | Bacteria | Actinobacteria | | Actinomycetia | | Propionibacteriales | | Propionibacteriaceae | Tessaracoccus | | | | | | 2.7.13.3 (100%) | Histidine kinase (100%) | GO:0016021 (100%) | integral component of membrane (100%) | GO:0000155 (100%) | phosphorelay sensor kinase activity (100%) | | | IPR003594 (100%); IPR003660 (100%); IPR0 | 1.00E+00 | 1.00E+00 | 9.73E+04 | 26.667 | 7.8051E-07 | 0.00052416 | 0.9 | NS | NS | NS | NS | 11.229 | 1.24E-04 | 0.001442 |

| | | | | | | | | | | | | | | | | | | | | | | | | | | | | | | | | | | | | | | | | |
|---|---|---|---|---|---|---|---|---|---|---|---|---|---|---|---|---|---|---|---|---|---|---|---|---|---|---|---|---|---|---|---|---|---|---|---|---|---|---|---|---|
| | | | | | | | | | | | | | | | | | | | | | | | | | | 03661(100%) | | | | | | | | | | | | | | |
| 90 | Variable 18 | TLSETVADLMR | Actinomycetia | Bacteria | Actinobacteria | | Actinomycetia | | | | | | | | | | | | | | | | | GO:0006355(100%) | regulation of transcription | DNA-templated(100%) | 1.00E+00 | 1.00E+00 | 2.78E+04 | 3.5138 | 2.2363E-06 | 0.00017193 | 0.9 | NS | NS | NS | NS | 9.707 | 3.42E-04 | 0.00051499 |
| 91 | Variable 180 | QNLPEGAA | Saccharopolysporaer | Bacteria | Actinobacteria | | Actinomycetia | | Pseudonocardiales | | Pseudonocardiaceae | Saccharopolyspora | | | Saccharopolyspora | | Saccharopolyspora | | | GO:0005840(100% | ribosome(100%) | GO:0003735(100% | structural constituent of ribosome(100%);r | GO:0000027(100%);GO:0006412 | ribosomal large subunit assembly | IPR005813(100%); | 1.00E+00 | 1.00E+00 | 3.72E+04 | 7.7157 | 1.5671E-09 | 0.00039636 | 0.9 | NS | NS | NS | NS | 30.858 | 5.69E-09 | 0.0039165 |

| | | | | | | | | | | | | | | | | | | | | | | | | | | | | | | | | | | | | | |
|---|---|---|---|---|---|---|---|---|---|---|---|---|---|---|---|---|---|---|---|---|---|---|---|---|---|---|---|---|---|---|---|---|---|---|---|---|---|
| | | | ythraeaNRRL2338 | | | | | | | | | | | | erythraea | | erythraeaNRRL2338 | | | ) | | );GO:0019843(100%) | RNA binding (100%) | (100%) | (100%);translation (100%) | IPR035566(100%) | | | | | | | | | | | |
| 92 | Variable 181 | SSSPGFYPR | Saccharothrix mutabilis subsp. capreol | Bacteria | Actinobacteria | | Actinomycetia | | Pseudonocardiales | | Pseudonocardiaceae | Saccharothrix | | | Saccharothrix mutabilis | Saccharothrix mutabilis subsp | | 3.4.17.18(100%) | Carboxypeptidase T (100%) | | | GO:0004181(100%);GO:0008270 | metallocarboxypeptidase activity (100%);zinc ion binding (100%) | | | IPR000834(100%);IPR033810(1 | 1.00E+00 | 1.00E+00 | 2.89E+04 | NS | NS | NS | NS | NS | NS | NS | NS |

| | | | | | | | | | | | | | | | | | | | | | | | | | | | | | | | | | | | | | | | | |
|---|---|---|---|---|---|---|---|---|---|---|---|---|---|---|---|---|---|---|---|---|---|---|---|---|---|---|---|---|---|---|---|---|---|---|---|---|---|---|---|---|
| | | | us | | | | | | | | | | | | | .capreolus | | | | | | (100%) | | | | 00%) | | | | | | | | | | | | | | |
| 94 | Variable 183 | DGAPEVHVAPVEGGPAK | Streptomyces | Bacteria | Actinobacteria | | Actinomycetia | | Streptomycetales | | Streptomycetaceae | Streptomyces | | | | | | 3.4.21.-(100%) | Serine endopeptidases (100%) | GO:0005737(100%) | cytoplasm (100%) | GO:0008236(100%) | serine-type peptidase activity (100%) | GO:0006508 (7%) | proteolysis (7%) | IPR005151(100%);IPR011659(100%);IPR | 1.00E+00 | 1.00E+00 | 1.04E+05 | 20.935 | 1.549E-08 | 0.00036501 | 0.9 | NS | NS | NS | NS | 19.996 | 7.92E-07 | 0.0051456 |

| | | | | | | | | | | | | | | | | | | | | | | | | | | | | | | | | | | | | | |
|---|---|---|---|---|---|---|---|---|---|---|---|---|---|---|---|---|---|---|---|---|---|---|---|---|---|---|---|---|---|---|---|---|---|---|---|---|---|
| | | | | | | | | | | | | | | | | | | | | | | | | | | 0123933(100%) | | | | | | | | | | | |
| 95 | Variable 184 | MLTSGMER | Streptomyces | Bacteria | Actinobacteria | | Actinomycetia | | Streptomycetales | | Streptomycetaceae | Streptomyces | | | | | | | | GO:0005840(97%);GO:0022627(3%) | ribosome(97%);cytosolic small ribosomal subunit(3%) | GO:0003729(97%);GO:0019843(97%);G | mRNA binding(97%);rRNA binding(97%);structural constituent of ribosome(94%) | GO:0006412(100%) | translation(100%) | IPR004044(100%);IPR004087(100%);I | 1.00E+00 | 1.00E+00 | 1.73E+05 | 5.678 | 0.0326 | 0.001333 | 0.9 | NS | NS | NS | NS |

| | | | | | | | | | | | | | | | | | | | | | | | | | | | | | | | | | | | | | |
|---|---|---|---|---|---|---|---|---|---|---|---|---|---|---|---|---|---|---|---|---|---|---|---|---|---|---|---|---|---|---|---|---|---|---|---|---|---|
| | | | | | | | | | | | | | | | | | | | | | | O:0003735(94%) | | | | PR005704(100%) | | | | | | | | | | | |
| 96 | Variable 185 | GTAVMAALTR | Streptomyces | Bacteria | Actinobacteria | | Actinomycetia | | Streptomycetales | | Streptomycetaceae | Streptomyces | | | | | | | | GO:0005886(100%) | plasmamembrane(100%) | | | | | IPR029046(100%) | 1.00E+00 | 1.00E+00 | 3.39E+05 | NS | NS | NS | NS | NS | NS | NS | NS |
| 97 | Variable 186 | AGSAPAPDATATALDGAR | Streptomyces | Bacteria | Actinobacteria | | Actinomycetia | | Streptomycetales | | Streptomycetaceae | Streptomyces | | | | | | 1.17.1.8(100% | 4-hydroxy-tetrahydrodipicolinate reduct | GO:0005737(97 | cytoplasm(97%);cytosol(3 | GO:0008839(10 | 4-hydroxy-tetrahydrodipicolinate red | acting on CH or CH2 groups | NAD or NADP as accept | GO:0009089(100 | 1.00E+00 | 1.00E+00 | 6.47E+04 | NS | NS | NS | NS | NS | NS | NS | NS |

| | | | | | | | | | | | | | | | | | | | | | | | | | | | | | | | | | | | | | | | | |
|---|---|---|---|---|---|---|---|---|---|---|---|---|---|---|---|---|---|---|---|---|---|---|---|---|---|---|---|---|---|---|---|---|---|---|---|---|---|---|---|---|
| | | | | | | | | | | | | | | | | | | ) | ase(100%) | %);GO:0005829(3%) | %) | 0%);GO:0016726(100%);GO:0050661(100%) | uctase(100%);oxidoreductase activity | | or(100%);NADP binding(100%) | %);GO:0019877(100%) | | | | | | | | | | | | | | |
| 98 | Variable | ATHDATAALMLR | Streptomyc | Bacteria | Actinobac | | Actinomyceti | | Streptomycetal | | Streptomycetac | Streptomy | | | | | | 7.5.2.7( | ABC-type D-ribose tra | GO:00058 | plasmamembr | GO:00055 | ATP binding(100%);A | | | IPR00343 | 1.00E+00 | 1.00E+00 | 3.93E+04 | 0.29563 | 1.2944E- | 0.001581 | 0.9 | NS | NS | NS | NS | 36.856 | 5.72E-10 | 0.004962 |

| | | | | | | | | | | | | | | | | | | | | | | | | | | | | | | | | | | | | | | |
|---|---|---|---|---|---|---|---|---|---|---|---|---|---|---|---|---|---|---|---|---|---|---|---|---|---|---|---|---|---|---|---|---|---|---|---|---|---|---|
| | 187 | | es | | teria | | a | | es | | eae | ces | | | | | | 100%) | nsporter (100%) | 86(91%) | ane (91%) | 24(100%);GO:0015611(91%);GO:0016887(82%) | BC-type D-ribose transporter activity (91%);ATPase (82%) | | | 9(100%);IPR003593(100%);IPR027417(100%) | | | | | 11 | | | | | | | 5 |
| 99 | Varia | AADHTAYLAW | Strepto | Bacte | Actino | | Actinom | | Streptom | | Streptom | Strept | | | | | | 2.1.1 | 5-methyltetrah | | | GO:00 | 5-methyltetrah | GO:000908 | methionin | IPR00 | 1.00E | 1.00E | 1.30E | NS | NS | NS | NS | NS | NS | NS | NS | |

| | | | | | | | | | | | | | | | | | | | | | | | | | | | | | | | | | | | | | | | | |
|---|---|---|---|---|---|---|---|---|---|---|---|---|---|---|---|---|---|---|---|---|---|---|---|---|---|---|---|---|---|---|---|---|---|---|---|---|---|---|---|---|
| | ble 188 | ATEAFR | myces | ria | bacteria | | ycetia | | ycetales | | ycetaceae | omyces | | | | | | .14(100%) | ydropteroyltriglutamate--homocysteine S-methyltransferase(100%) | | | 03871(100%);GO:0008270(67%) | ydropteroyltriglutamate-homocysteine S-methyltransferase activity(100%);zinc ion binding(67%) | 6(100%);GO:0032259(100%) | ebiosynthetic process(100%); methylation(100%) | 2629(100%);IPR006276(100%);IPR013215(100%) | +00 | +00 | +05 | | | | | | | | | | | |
| 100 | Var | GGMAEV | Stre | Bac | Acti | | Actin | | Strept | | Strept | Stre | | | | | | 2.7 | Non-spe | GO: | integral | GO: | ATP bin | | | IPR | 1.0 | 1.0 | 3.7 | 0.2 | 1.1 | 0.00 | 0.9 | NS | NS | NS | NS | 32. | 3.1 | 0.0 |

| | | | | | | | | | | | | | | | | | | | | | | | | | | | | | | | | | | | | | | | | |
|---|---|---|---|---|---|---|---|---|---|---|---|---|---|---|---|---|---|---|---|---|---|---|---|---|---|---|---|---|---|---|---|---|---|---|---|---|---|---|---|---|
| | iable 189 | YHAHDTR | ptomyces | teria | nobacteria | | omycetia | | omycetales | | omycetaceae | ptomyces | | | | | | .11.1(100%) | cific serine/threonine protein kinase (100%) | 0016021(90%) | component of membrane (90%) | 0005524(100%);GO:0106310(90%);GO:0106311(90%) | ding (100%);protein serine kinase activity (90%);protein threonine kinase activity (90%) | | | 000719(100%);IPR011009(100%);IPR005543(98%) | 0E+00 | 0E+00 | 9E+04 | 7073 | 253E-11 | 14294 | | | | | | 342 | 5E-09 | 010183 |

| | | | | | | | | | | | | | | | | | | | | | | | | | | | | | | | | | | | | | |
|---|---|---|---|---|---|---|---|---|---|---|---|---|---|---|---|---|---|---|---|---|---|---|---|---|---|---|---|---|---|---|---|---|---|---|---|---|---|
| 101 | Variable 19 | WPMDDAAR | Actinomycetia | Bacteria | Actinobacteria | | Actinomycetia | | | | | | | | | | | 3.5.1.-(100%) | In linear amides (100%) | | | GO:0016787(100%) | hydrolase activity (100%) | | | IPR002589(100%);IPR043472(100%) | 1.00E+00 | 1.00E+00 | 1.56E+05 | 7.4747 | 0.034861 | 0.00025476 | 0.9 | NS | NS | NS | NS |
| 102 | Variable 190 | VGDTVTSQHK | Streptomyces | Bacteria | Actinobacteria | | Actinomycetia | | Streptomycetales | | Streptomycetaceae | Streptomyces | | | | | | 3.6.5.0(100%) | -1 | GO:0005886(10 | plasma membrane (100% | GO:0003746(10 | translation elongation factor activity | GO:0045727 (100%) | positive regulation of translat | IPR000640(100 | 1.00E+00 | 1.00E+00 | 1.42E+05 | NS | NS | NS | NS | NS | NS | NS | NS |

| | | | | | | | | | | | | | | | | | | | | | | | | | | | | | | | | | | | | | |
|---|---|---|---|---|---|---|---|---|---|---|---|---|---|---|---|---|---|---|---|---|---|---|---|---|---|---|---|---|---|---|---|---|---|---|---|---|---|
| | | | | | | | | | | | | | | | | | | | | 0%) | ) | 0%);GO:0003924(100%);GO:0005525(100%) | (100%);GTPase activity(100%);GTP binding(100%) | | ion(100%) | %);IPR000795(100%);IPR005225(100%) | | | | | | | | | | | |
| 103 | Variable | SGAVDTAQAR | Streptomyc | Bacteria | Actinobac | | Actinomyceti | | Streptomycetal | | Streptomycetac | Streptomy | | | | | | 3.5.99.6 | Glucosamine-6-phosp | GO:0057 | cytoplasm(17% | GO:0043 | glucosamine-6-phosphat | GO:0005975(83 | carbohydrate met | IPR00454 | 1.00E+00 | 1.00E+00 | 4.54E+04 | NS | NS | NS | NS | NS | NS | NS | NS |

| | | | | | | | | | | | | | | | | | | | | | | | | | | |
|---|---|---|---|---|---|---|---|---|---|---|---|---|---|---|---|---|---|---|---|---|---|---|---|---|---|---|
| | 191 | | es | | teria | | a | | es | | eae | ces | | | | | | (100%) | hate deaminase (100%) | 37(17%) | ) | 42(100%);GO:0042802(17%) | e deaminase activity (100%);identical protein binding (17%) | %);GO:0006044(83%);GO:0019262(83%) | abolic processes (83%); N-acetylglucosamine metabolic processes (83%); N-acetylneuramina | 7(100%);IPR006148(100%);IPR018321(100%) |

| | | | | | | | | | | | | | | | | | | | | | | | | | | | | | | | | | | | | | | | | |
|---|---|---|---|---|---|---|---|---|---|---|---|---|---|---|---|---|---|---|---|---|---|---|---|---|---|---|---|---|---|---|---|---|---|---|---|---|---|---|---|---|
| | | | | | | | | | | | | | | | | | | | | | | | | | tecatabolic process (83%) | | | | | | | | | | | | | | | |
| 104 | Variable 192 | ANLFEGVQVYK | Streptomyces | Bacteria | Actinobacteria | | Actinomycetia | | Streptomycetales | | Streptomycetaceae | Streptomyces | | | | | | 3.2.1.55 (100%) | Non-reducing end alpha-L-arabinofuranosidase (100%) | GO:0005576 (100%) | extracellular region (100%) | GO:0046556 (100%);GO:0030246 (6%) | alpha-L-arabinofuranosidase activity (100%);carbohydrate binding (6%) | GO:0046373 (100%);GO:0045493 (4%) | L-arabinose metabolic process (100%); xylan catabolic process | IPR007772 (100%);IPR005193 (100%); | 1.00E+00 | 1.00E+00 | 6.95E+04 | 8.078 | 0.00020347 | 0.00041052 | 0.9 | NS | NS | NS | NS | 6.9039 | 2.56E-03 | 0.00015694 |

| | | | | | | | | | | | | | | | | | | | | | | | | | | | | | | | | | | | | | |
|---|---|---|---|---|---|---|---|---|---|---|---|---|---|---|---|---|---|---|---|---|---|---|---|---|---|---|---|---|---|---|---|---|---|---|---|---|---|
| | | | | | | | | | | | | | | | | | | | | | | | | | (4%) | IPR023296(100%) | | | | | | | | | | | |
| 105 | Variable 193 | ALGSALLAASGK | Streptomyces | Bacteria | Actinobacteria | | Actinomycetia | | Streptomycetales | | Streptomycetaceae | Streptomyces | | | | | | 2.3.1.189(100%) | Mycothiol synthase(100%) | | | GO:0035447(100%);GO:0008080(9 | mycothiol synthase activity(100%);N-acetyltransferase activity(98%) | GO:0010125(100%) | mycothiol biosynthetic process(100%) | IPR000182(100%);IPR016181(100 | 1.00E+00 | 1.00E+00 | 1.23E+05 | NS | NS | NS | NS | NS | NS | NS | NS |

| | | | | | | | | | | | | | | | | | | | | | | | | | | | | | | | | | | | | | | | | |
|---|---|---|---|---|---|---|---|---|---|---|---|---|---|---|---|---|---|---|---|---|---|---|---|---|---|---|---|---|---|---|---|---|---|---|---|---|---|---|---|---|
| | | | | | | | | | | | | | | | | | | | | | | 8%) | | | | %);IPR017813(100%) | | | | | | | | | | | | | | |
| 106 | Variable 194 | YVDTAELNK | Streptomyces | Bacteria | Actinobacteria | | Actinomycetia | | Streptomycetales | | Streptomycetaceae | Streptomyces | | | | | | 2.3.1.8(100%) | Phosphate acetyltransferase(100%) | GO:0005737(80%) | cytoplasm(80%) | GO:0008959(90%);GO:0016407( | phosphate acetyltransferase activity(90%);acetyltransferase activity(10%) | GO:0006085(80%) | acetyl-CoA biosynthetic process(80%) | IPR002505(100%);IPR004614(1 | 1.00E+00 | 1.00E+00 | 2.91E+04 | NS | NS | NS | NS | NS | NS | NS | NS | 6.2418 | 4.23E-03 | 0.0021631 |

| | | | | | | | | | | | | | | | | | | | | | | | | | | | | | | | | | | | | | |
|---|---|---|---|---|---|---|---|---|---|---|---|---|---|---|---|---|---|---|---|---|---|---|---|---|---|---|---|---|---|---|---|---|---|---|---|---|---|
| | | | | | | | | | | | | | | | | | | | | | | 10%) | | | | 00%);IPR010766(100%) | | | | | | | | | | | |
| 107 | Variable 195 | AGEPGETGALGPLR | Streptomyces | Bacteria | Actinobacteria | | Actinomycetia | | Streptomycetales | | Streptomycetaceae | Streptomyces | | | | | | 2.3.1.180(100%) | Beta-ketoacyl-[acyl-carrier-protein] synthase III(100%) | GO:0005737(100%) | cytoplasm(100%) | GO:0004315(100%);GO:00338 | 3-oxoacyl-[acyl-carrier-protein] synthase activity(100%);beta-ket | GO:0006633(100%);GO:0006631(13%) | fatty acid biosynthetic process(100%);fatty acid met | IPR004655(100%);IPR013747 | 1.00E+00 | 1.00E+00 | 5.82E+04 | NS | NS | NS | NS | NS | NS | NS | NS |

| | | | | | | | | | | | | | | | | | | | | | | | | | | | | | | | | | | | | | | | | |
|---|---|---|---|---|---|---|---|---|---|---|---|---|---|---|---|---|---|---|---|---|---|---|---|---|---|---|---|---|---|---|---|---|---|---|---|---|---|---|---|---|
| | | | | | | | | | | | | | | | | | | | | | | 18(100%) | oacyl-acyl-carrier-protein synthase III activity (100%) | | abolic process (13%) | (100%);IPR013751(100%) | | | | | | | | | | | | | | |
| 108 | Variable 196 | EAGFPDLDGATR | Streptomyces | Bacteria | Actinobacteria | | Actinomycetia | | Streptomycetales | | Streptomycetaceae | Streptomyces | | | | | | 4.3.1.30(100%);1.1.99.-( | dTDP-4-amino-4 | 6-dideoxy-D-glucoseamm | | | GO:0016841(100%);GO:0046872(100%);GO:0051539 | ammonia-lyase activity (100%);metal ion binding (100%) | 4 sulfur cluster binding (100%) | GO:0033068(100%) | 1.00E+00 | 1.00E+00 | 4.85E+04 | 4.4477 | 0.00020333 | 0.00013575 | 0.9 | NS | NS | NS | NS | 4.6048 | 1.56E-02 | 0.00019375 |

| | | | | | | | | | | | | | | | | | | | | | | | | | | | | | | | | | | | | | |
|---|---|---|---|---|---|---|---|---|---|---|---|---|---|---|---|---|---|---|---|---|---|---|---|---|---|---|---|---|---|---|---|---|---|---|---|---|---|
| | | | | | | | | | | | | | | | | | | 50%) | | onia-lyase(100%);Withotheracceptors(50%) | | | (100%) | ;4 iron | | | | | | | | | | | | | |
| 109 | Variable | MHLLVATAVSVER | Streptomyc | Bacteria | Actinobac | | Actinomyceti | | Streptomycetal | | Streptomycetac | Streptomy | | | | | | 3.2.2.26 | Futalosine hydrolase | GO:0058 | cytosol(13%) | GO:00167 | hydrolase activity | hydrolyzing N-gly | GO:0009234(100 | menaquino | 1.00E+00 | 1.00E+00 | 6.94E+04 | NS | NS | NS | NS | NS | NS | NS | NS |

| | | | | | | | | | | | | | | | | | | | | | | | | | | |
|---|---|---|---|---|---|---|---|---|---|---|---|---|---|---|---|---|---|---|---|---|---|---|---|---|---|---|
| | 197 | | es | | teria | | a | | es | | eae | ces | | | | | | (100%) | (100%) | 29(13%) | | 99(88%);GO:0008782(13%);GO:0008930(13%) | | cosyl compounds (88%);adenosylhomocysteine nucleosidase activity (13%);methylthioadenosine nucleosidase | %);GO:0009116(88%);GO:0019284(13%) | nebiosynthetic process(100%);nucleoside metabolic process( |

| | | | | | | | | | | | | | | | | | | | | | | | | | | |
|---|---|---|---|---|---|---|---|---|---|---|---|---|---|---|---|---|---|---|---|---|---|---|---|---|---|---|
| | | | | | | | | | | | | | | | | | | | | | | | | activity (13%) | | 88%); L-methionine salvage from S-adenosylmethionine (13%) |

| | | | | | | | | | | | | | | | | | | | | | | | | | | | | | | | | | | | | | |
|---|---|---|---|---|---|---|---|---|---|---|---|---|---|---|---|---|---|---|---|---|---|---|---|---|---|---|---|---|---|---|---|---|---|---|---|---|---|
| 110 | Variable 198 | ELTADLLTEAK | Streptomyces | Bacteria | Actinobacteria | | Actinomycetia | | Streptomycetales | | Streptomycetaceae | Streptomyces | | | | | | 6.3.5.5(100%) | Carbamoyl-phosphate synthase (glutamine-hydrolyzing) (100%) | GO:0005737(6%) | cytoplasm(6%) | GO:0004088(100%);GO:0005524(100%);GO:0046872( | carbamoyl-phosphate synthase (glutamine-hydrolyzing) activity (100%);ATP binding (100%);metal ion binding (88%) | GO:0006526(100%);GO:0044205(88%);GO:0006221(12%) | arginine biosynthetic process (100%);'de novo' UMP biosynthetic process (88%); pyrimidi | IPR005479(100%);IPR005480(100%);IPR005483(100 | 1.00E+00 | 1.00E+00 | 2.04E+05 | 7.3602 | 0.019719 | 0 | 0.9 | NS | NS | NS | NS |

| | | | | | | | | | | | | | | | | | | | | | | | | | | | | | | | | | | | | | | | | |
|---|---|---|---|---|---|---|---|---|---|---|---|---|---|---|---|---|---|---|---|---|---|---|---|---|---|---|---|---|---|---|---|---|---|---|---|---|---|---|---|---|
| | | | | | | | | | | | | | | | | | | | | | | 88%) | | | ne nucleotide biosynthetic process(12%) | %) | | | | | | | | | | | | | | |
| 113 | Variable 20 | WVAQTDFSNDEAVGYLADR | Actinomycetia | Bacteria | Actinobacteria | | Actinomyceta | | | | | | | | | | | 3.6.5.-(100%) | Acting on GTP; involved in cellular and subcellular movement(100% | GO:0005737(80%) | cytoplasm(80%) | GO:0000287(100%);GO:0003 | magnesium ion binding(100%);GTPase activity(100%);GTP binding | GO:0042254(80%) | ribosome biogenesis(80%) | IPR006073(100%);IPR00616 | 1.00E+00 | 1.00E+00 | 2.70E+04 | 4.2638 | 1.5344E-08 | 0.0002532 | 0.9 | NS | NS | NS | NS | 21.294 | 4.12E-07 | 0.0019276 |

| | | | | | | | | | | | | | | | | | | | | | | | | | | | | | | | | | | | | | |
|---|---|---|---|---|---|---|---|---|---|---|---|---|---|---|---|---|---|---|---|---|---|---|---|---|---|---|---|---|---|---|---|---|---|---|---|---|---|
| | | | | | | | | | | | | | | | | | | | ) | | | 924(100%);GO:0005525(100%) | (100%) | | | 9(100%);IPR014100(100%) | | | | | | | | | | | |
| 115 | Variable 201 | GLFTGLPGAK | Streptomyces | Bacteria | Actinobacteria | | Actinomycetia | | Streptomycetales | | Streptomycetaceae | Streptomyces | | | | | | 2.1.1.235(100%) | dTDP-3-amino-3 | 6-dideoxy-alpha-D-gl | N-dimethyltransferase(100%) | | | GO:0008168(50%);GO:0008757(50%) | methyltransferase activity(50%);S-ad | GO:0032259(100%);GO:0 | 1.00E+00 | 1.73E+07 | 1.00E+00 | NS | NS | NS | NS | NS | NS | NS | NS |

| | | | | | | | | | | | | | | | | | | | | | | | | | | |
|---|---|---|---|---|---|---|---|---|---|---|---|---|---|---|---|---|---|---|---|---|---|---|---|---|---|---|
| | | | | | | | | | | | | | | | | | | | | ucopyranoseN | | | | ;GO:0042803(50%) | enosylmethionine-dependent methyltransferase activity(50%); protein homodimerization ac | 0170000(50%) |

| | | | | | | | | | | | | | | | | | | | | | | | | | | | | | | | | | | | | | |
|---|---|---|---|---|---|---|---|---|---|---|---|---|---|---|---|---|---|---|---|---|---|---|---|---|---|---|---|---|---|---|---|---|---|---|---|---|---|
| | | | | | | | | | | | | | | | | | | | | | | | | | tivity(50%) | | | | | | | | | | | | |
| 116 | Variable 202 | TLSLALTGK | Streptomyces | Bacteria | Actinobacteria | | Actinomycetia | | Streptomycetales | | Streptomycetaceae | Streptomyces | | | | | | 6.3.2.4(82%);6.1.2.1(9%);6.3.2.-(9%) | D-alanine--D-alanine ligase(82%);D-alanine--(R)-lactate ligase(9%);Acid--amino-acid ligases (peptide | GO:0005737(90%);GO:0005829(10%);GO:0005 | cytoplasm(90%);cytosol(10%);plasma membrane(10%) | GO:0005524(100%);GO:0008716(100%);GO:00 | ATP binding(100%);D-alanine-D-alanine ligase activity(100%);metal ion binding(100%) | GO:0008360(100%);GO:0009252(100%);GO:0071555(100%) | regulation of cell shape(100%);peptidoglycan biosynthetic process(100%);cell w | IPR000291(100%);IPR005905(100%);IPR0110 | 1.00E+00 | 4.68E+05 | 1.00E+00 | NS | NS | NS | NS | NS | NS | NS | NS |

| | | | | | | | | | | | | | | | | | | | | | | | | | | | | | | | | | | | | | |
|---|---|---|---|---|---|---|---|---|---|---|---|---|---|---|---|---|---|---|---|---|---|---|---|---|---|---|---|---|---|---|---|---|---|---|---|---|---|
| | | | | | | | | | | | | | | | | | | | synthases)(9%) | 886(10%) | | 46872(100%) | | | all organization(100%) | 95(100%) | | | | | | | | | | | |
| 117 | Variable 203 | GEPVVTER | Streptomyces | Bacteria | Actinobacteria | | Actinomycetia | | Streptomycetales | | Streptomycetaceae | Streptomyces | | | | | | 3.5.2.2(81%);3.5.2.-(19%) | Dihydropyrimidinase(81%); In cyclic amides(19%) | GO:0005737(94%) | cytoplasm(94%) | GO:0016810(58%);GO:0004157(39%);GO | hydrolase activity | acting on carbon-nitrogen (but not peptide) bonds(58%);dihydropyrimidinase act | acting on carbon-nitrogen (but not peptide) bonds | in cyclic amides(3%) | 1.00E+00 | 1.73E+05 | 1.00E+00 | NS | NS | NS | NS | NS | NS | NS | NS |

| | | | | | | | | | | | | | | | | | | | | | | | | | | | | | | | | | | | | | |
|---|---|---|---|---|---|---|---|---|---|---|---|---|---|---|---|---|---|---|---|---|---|---|---|---|---|---|---|---|---|---|---|---|---|---|---|---|---|
| | | | | | | | | | | | | | | | | | | | | | | :0016812(3%) | | ivity(39%);hydrolase activity | | | | | | | | | | | | | |
| 118 | Variable 204 | SWDGTVANGVAATGVK | Streptomyces | Bacteria | Actinobacteria | | Actinomycetia | | Streptomycetales | | Streptomycetaceae | Streptomyces | | | | | | 3.4.14.11(100%) | Xaa-Prodipeptidyl-peptidase(100%) | | | GO:0008239(100%);GO:0004177(13%); | dipeptidyl-peptidase activity(100%);aminopeptidase activity(13%);serine-type pe | | | IPR000383(100%);IPR008979(100%);I | 1.00E+00 | 3.78E+05 | 1.00E+00 | NS | NS | NS | NS | NS | NS | NS | NS |

| | | | | | | | | | | | | | | | | | | | | | | | | | | | | | | | | | | | | | | | | |
|---|---|---|---|---|---|---|---|---|---|---|---|---|---|---|---|---|---|---|---|---|---|---|---|---|---|---|---|---|---|---|---|---|---|---|---|---|---|---|---|---|
| | | | | | | | | | | | | | | | | | | | | | | GO:0008236(13%) | ptidase activity (13%) | | | PR013736(100%) | | | | | | | | | | | | | | |
| 119 | Variable 205 | NAHSATTWSGQYVGGAEAR | Streptomyces | Bacteria | Actinobacteria | | Actinomycetia | | Streptomycetales | | Streptomycetaceae | Streptomyces | | | | | | | | GO:0005576(100%) | extracellular region (100%) | GO:0009374(100%) | biotin binding (100%) | | | IPR005468(100%);IPR005469(100 | 1.00E+00 | 1.00E+00 | 6.15E+04 | 0.41888 | 2.255E-09 | 0.0016884 | 0.9 | NS | NS | NS | NS | 15.713 | 8.04E-06 | 0.0098109 |

| | | | | | | | | | | | | | | | | | | | | | | | | | | | | | | | | | | | | | |
|---|---|---|---|---|---|---|---|---|---|---|---|---|---|---|---|---|---|---|---|---|---|---|---|---|---|---|---|---|---|---|---|---|---|---|---|---|---|
| | | | | | | | | | | | | | | | | | | | | | | | | | | %); IPR017889 (100%) | | | | | | | | | | | |
| 120 | Variable 206 | STLVGHDTFTK | Streptomyces | Bacteria | Actinobacteria | | Actinomycetia | | Streptomycetales | | Streptomycetaceae | Streptomyces | | | | | | | | GO:0005576 (100%) | extracellular region (100%) | GO:0009374 (100%) | biotin binding (100%) | | | IPR005468 (100%); IPR005469 (1 | 2.85E+05 | 1.00E+00 | 1.00E+00 | NS | NS | NS | NS | NS | NS | NS | NS |

| | | | | | | | | | | | | | | | | | | | | | | | | | | | | | | | | | | | | | |
|---|---|---|---|---|---|---|---|---|---|---|---|---|---|---|---|---|---|---|---|---|---|---|---|---|---|---|---|---|---|---|---|---|---|---|---|---|---|
| | | | | | | | | | | | | | | | | | | | | | | | | | | 00%);IPR017889(100%) | | | | | | | | | | | |
| 121 | Variable 207 | LLGTQVSGTAVDLVAGTVR | Streptomyces | Bacteria | Actinobacteria | | Actinomycetia | | Streptomycetales | | Streptomycetaceae | Streptomyces | | | | | | | | GO:0005886(100%);GO:00452 | plasma membrane (100%);proton-transporting AT | catalytic core F(1) (100%) | GO:0046933 (100%) | proton-transporting ATP synthase activity | rotational mechanism (100%) | GO:0015986(11%) | 2.13E+05 | 1.00E+00 | 1.00E+00 | NS | NS | NS | NS | NS | NS | NS | NS |

| | | | | | | | | | | | | | | | | | | | | | | | | | | | | | | | | | | | | | |
|---|---|---|---|---|---|---|---|---|---|---|---|---|---|---|---|---|---|---|---|---|---|---|---|---|---|---|---|---|---|---|---|---|---|---|---|---|---|
| | | | | | | | | | | | | | | | | | | | | 61(100%) | P synthase complex | | | | | | | | | | | | | | | | |
| 123 | Variable 209 | FMVANTSHLAR | Streptomyces davaonensis JCM 4913 | Bacteria | Actinobacteria | | Actinomycetia | | Streptomycetales | | Streptomycetaceae | Streptomyces | | | Streptomyces davaonensis | | Streptomyces davaonensis JCM 4913 | 2.6.1.114(100%) | 8-demethyl-8-aminoriboflavin-5'-phosphate synthase (100%) | | | GO:0008483(100%);GO:0016491(100%) | transaminase activity (100%);oxidoreductase activity (100%) | GO:0016999(100%);GO:0017000(100%) | antibiotic metabolic process (100%);antibiotic biosynthetic process (100% | IPR005025(100%);IPR029039(100%) | 1.05E+06 | 1.00E+00 | 1.00E+00 | NS | NS | NS | NS | NS | NS | NS | NS |

| | | | | | | | | | | | | | | | | | | | | | | | | | | | | | | | | | | | | | | | | |
|---|---|---|---|---|---|---|---|---|---|---|---|---|---|---|---|---|---|---|---|---|---|---|---|---|---|---|---|---|---|---|---|---|---|---|---|---|---|---|---|---|
| | | | | | | | | | | | | | | | | | | | | | | | | | ) | | | | | | | | | | | | | | | |
| 124 | Variable 21 | LVAVVEDNSR | Actinomycetia | Bacteria | Actinobacteria | | Actinomycetia | | | | | | | | | | | 2.2.1.7(100%) | 1-deoxy-D-xylulose-5-phosphate synthase(100%) | GO:0005829(0%) | cytosol(0%) | GO:0008661(100%);GO:0000287(97%);GO:0030976( | 1-deoxy-D-xylulose-5-phosphate synthase activity(100%);magnesium ion binding(97%);thiamine pyrophosphate bindin | GO:0009228(100%);GO:0016114(100%);GO:0052865(97%) | thiamine biosynthetic process(100%);terpenoid biosynthetic process(100%);1-deoxy-D-xy | IPR009014(100%);IPR033248(100%);IPR005475(10 | 1.00E+00 | 1.00E+00 | 6.86E+04 | 0.63426 | 7.4657E-07 | 0.00071696 | 0.9 | NS | NS | NS | NS | 6.6971 | 2.99E-03 | 0.00034884 |

| | | | | | | | | | | | | | | | | | | | | | | | | | | | | | | | | | | | | | |
|---|---|---|---|---|---|---|---|---|---|---|---|---|---|---|---|---|---|---|---|---|---|---|---|---|---|---|---|---|---|---|---|---|---|---|---|---|---|
| | | | | | | | | | | | | | | | | | | | | | | 97%) | g (97%) | | lulose 5-phosphate biosynthetic process (97%) | 0%) | | | | | | | | | | | |
| 125 | Variable 210 | AQPLTSVER | Streptomyces lasalocidi | Bacteria | Actinobacteria | | Actinomycetia | | Streptomycetales | | Streptomycetaceae | Streptomyces | | | Streptomyces lasalocidi | | | 1.14.13.-(100%) | With NADH or NADPH as one donor | and incorporation of one | | | GO:0071949 (100%);GO:0004497 (20%) | FAD binding (100%);monooxygenase activity (20 | GO:0017000 (20%) | antibiotic biosynthetic process | 1.00E+00 | 1.00E+00 | 1.08E+05 | NS | NS | NS | NS | NS | NS | NS | NS |

| | | | | | | | | | | | | | | | | | | | | | | | | | | | | | | | | | | | | | |
|---|---|---|---|---|---|---|---|---|---|---|---|---|---|---|---|---|---|---|---|---|---|---|---|---|---|---|---|---|---|---|---|---|---|---|---|---|---|
| | | | | | | | | | | | | | | | | | | | | atom of oxygen (100%) | | | | %) | | (20%) | | | | | | | | | | | |
| 126 | Variable 211 | LVGMPPTDR | Streptomyces nogalater | Bacteria | Actinobacteria | | Actinomycetia | | Streptomycetales | | Streptomycetaceae | Streptomyces | | | Streptomyces nogalater | | | 5.5.1.26 (100%) | Nogalonic acid methyl ester cyclase (100%) | | | | | GO:0030638 (100%) | polyketide metabolic process (100%) | IPR009959 (100%); IPR032710 ( | 1.00E+00 | 1.25E+05 | 1.00E+00 | NS | NS | NS | NS | NS | NS | NS | NS |

| | | | | | | | | | | | | | | | | | | | | | | | | | | | | | | | | | | | | | |
|---|---|---|---|---|---|---|---|---|---|---|---|---|---|---|---|---|---|---|---|---|---|---|---|---|---|---|---|---|---|---|---|---|---|---|---|---|---|
| | | | | | | | | | | | | | | | | | | | | | | | | | | 100%) | | | | | | | | | | | |
| 127 | Variable 212 | TQDWGSCFEGK | Streptomyces olivaceoviridis | Bacteria | Actinobacteria | | Actinomycetia | | Streptomycetales | | Streptomycetaceae | Streptomyces | | | Streptomyces olivaceoviridis | | | 3.2.1.14(100%) | Chitinase(100%) | | | GO:0004568(100%) | chitinase activity(100%) | GO:0000272(100%);GO:0006032(100%) | polysaccharide catabolic process(100%);chitin catabolic process(100%) | | 1.00E+00 | 1.00E+00 | 2.55E+05 | NS | NS | NS | NS | NS | NS | NS | NS |
| 128 | Varia | SAAPHWLAPPLD | Strepto | Bacte | Actino | | Actinom | | Streptom | | Streptom | Strept | Strept | | Strept | | | 6.3.3 | Dethiobiotin syn | GO:00 | cytosol (5 | GO:00 | magnesium ion | GO:000910 | biotin bios | IPR00 | 1.00E | 1.00E | 4.23E | NS | NS | NS | NS | NS | NS | NS | NS |

| | | | | | | | | | | | | | | | | | | | | | | | | | | | | | |
|---|---|---|---|---|---|---|---|---|---|---|---|---|---|---|---|---|---|---|---|---|---|---|---|---|---|---|---|---|---|
| | ble 213 | GTWDADAFGDR | mycescoelicolor | ria | bacteria | | ycetia | | ycetales | | ycetaceae | omyces | omycesalbidoflavusgroup | | omycescoelicolor | | | .3(100%) | thase(100%) | 05829(50%) | 0%) | 00287(100%);GO:0004141(100%);GO:0005524(100%) | binding(100%);dethiobiotin synthase activity(100%);ATP binding(100%) | 2(100%) | ynthetic process(100%) | 4472(100%);IPR027417(100%) | +00 | +00 | +04 |

| | | | | | | | | | | | | | | | | | | | | | | | | | | | | | | | | | | | | | |
|---|---|---|---|---|---|---|---|---|---|---|---|---|---|---|---|---|---|---|---|---|---|---|---|---|---|---|---|---|---|---|---|---|---|---|---|---|---|
| 130 | Variable 215 | FHLAGDGSGELVAR | Streptomyces griseus | Bacteria | Actinobacteria | | Actinomycetia | | Streptomycetales | | Streptomycetaceae | Streptomyces | Streptomyces griseus group | Streptomyces griseus subgroup | Streptomyces griseus | | | | | GO:0005737 (100%) | cytoplasm (100%) | | | GO:0042274 (100%) | ribosomal small subunit biogenesis (100%) | IPR003728(100%);IPR028989(100%);IPR028998(100 | 1.65E+06 | 1.00E+00 | 1.00E+00 | NS | NS | NS | NS | NS | NS | NS | NS |

| | | | | | | | | | | | | | | | | | | | | | | | | | | | | | | | | | | | | | |
|---|---|---|---|---|---|---|---|---|---|---|---|---|---|---|---|---|---|---|---|---|---|---|---|---|---|---|---|---|---|---|---|---|---|---|---|---|---|
| | | | | | | | | | | | | | | | | | | | | | | | | | | %) | | | | | | | | | | | |
| 131 | Variable 216 | DAAPDAGFPGFAPFVR | Streptomyces cinnamonensis | Bacteria | Actinobacteria | | Actinomycetia | | Streptomycetales | | Streptomycetaceae | Streptomyces | Streptomyces virginiae group | | Streptomyces cinnamonensis | | | 5.4.99.2(100%) | Methylmalonyl-CoA mutase(100%) | | | GO:0004494(100%);GO:0031419(100%) | methylmalonyl-CoA mutase activity(100%);cobalamin binding(100%) | GO:0019652(50%) | lactate fermentation to propionate and acetate(50%) | IPR006099(100%);IPR016176(100%);IPR004608(5 | 1.00E+00 | 4.94E+04 | 1.00E+00 | NS | NS | NS | NS | NS | NS | NS | NS |

| | | | | | | | | | | | | | | | | | | | | | | | | | | | | | | | | | | | | | | | | |
|---|---|---|---|---|---|---|---|---|---|---|---|---|---|---|---|---|---|---|---|---|---|---|---|---|---|---|---|---|---|---|---|---|---|---|---|---|---|---|---|---|
| | | | | | | | | | | | | | | | | | | | | | | | | | | 0%) | | | | | | | | | | | | | | |
| 133 | Variable 218 | ASGTCADTVTAHGPVPGATVV | ThermobifidafuscaYX | Bacteria | Actinobacteria | | Actinomycetia | | Streptosporangiales | | Nocardiopsaceae | Thermobifida | | | Thermobifidafusca | | ThermobifidafuscaYX | 2.7.1.148(100%) | 4-(cytidine 5'-diphospho)-2-C-methyl-D-erythritol kinase(100%) | | | GO:0005524(100%);GO:0050515(100%) | ATP binding(100%);4-(cytidine 5'-diphospho)-2-C-methyl-D-erythritol kinase activity(100%) | GO:0016114(100%);GO:0019288(100%) | terpenoid biosynthetic process(100%);isopentenyl diphosphate biosynthetic process | methylerythritol 4-phosphate pathway(100%) | 1.00E+00 | 1.00E+00 | 3.28E+05 | NS | NS | NS | NS | NS | NS | NS | NS | 8.81746 | 6.35E-044 | 0.000540230 |
| 1 | V | QT | A | B | A | | A | | A | | A | A | | | | | | | | G | lar | G | str | G | tra | I | 1 | 1 | 1 | N | N | N | N | N | N | N | N | 6 | 4 | 0 |

| | | | | | | | | | | | | | | | | | | | | | | | | | | | | | | | | | | | | | | | | |
|---|---|---|---|---|---|---|---|---|---|---|---|---|---|---|---|---|---|---|---|---|---|---|---|---|---|---|---|---|---|---|---|---|---|---|---|---|---|---|---|---|
| 34 | ariable 219 | YLNVADNSGAK | quifex | acteria | quificae | | quificae | | quificales | | quificaceae | quifex | | | | | | | | O:0015934(75%);GO:0022625(25%) | geribosomal subunit(75%);cytosolic large ribosomal subunit(25%) | O:0003735(100%);GO:0019843(75%);GO:0070180(25 | uctural constituent of ribosome(100%);rRNA binding(75%);large ribosomal subunit rRNA binding(25%) | O:0006412(100%) | nslation(100%) | PR002218(100%);IPR005745(100%);IPR019972(100% | .00E+00 | .00E+00 | .08E+05 | S | S | S | S | S | S | S | S | .3105 | .02E-03 | .000410021 |

| | | | | | | | | | | | | | | | | | | | | | | | | | | | | | | | | | | | | | |
|---|---|---|---|---|---|---|---|---|---|---|---|---|---|---|---|---|---|---|---|---|---|---|---|---|---|---|---|---|---|---|---|---|---|---|---|---|---|
| | | | | | | | | | | | | | | | | | | | | | | %) | | | | ) | | | | | | | | | | | |
| 135 | Variable 22 | FAPVPLYSTYHDCWR | Actinomycetia | Bacteria | Actinobacteria | | Actinomycetia | | | | | | | | | | | 3.7.1.3(100%) | Kynureninase(100%) | GO:0005737(97%) | cytoplasm(97%) | GO:0030170(100%);GO:0030429(100%);GO:0061998 | pyridoxal phosphate binding(100%);kynureninase activity(100%);3-hydroxykynureninase activity(97%) | GO:0006569(100%);GO:0019805(100%);GO:0034354(100%) | tryptophan catabolic process(100%);quinolinate biosynthetic process(100%);'de novo' N | IPR000192(100%);IPR010111(100%);IPR015421(1 | 1.00E+00 | 1.00E+00 | 3.42E+05 | NS | NS | NS | NS | NS | NS | NS | NS |

| | | | | | | | | | | | | | | | | | | | | | | | | | | | | | | | | | | | | | | | | |
|---|---|---|---|---|---|---|---|---|---|---|---|---|---|---|---|---|---|---|---|---|---|---|---|---|---|---|---|---|---|---|---|---|---|---|---|---|---|---|---|---|
| | | | | | | | | | | | | | | | | | | | | | | 1 (97%) | | | AD biosynthetic process from tryptophan (100%) | 00%) | | | | | | | | | | | | | | |
| 136 | Variable 220 | VTVDLGSGK | Bacteroidia | Bacteria | Bacteroidetes | | Bacteroidia | | | | | | | | | | | | | | | GO:0046556 (100%) | alpha-L-arabinofuranosidase activity (100%) | GO:0046373 (100%) | L-arabinose metabolic process (100%) | IPR010720 (100%); IPR0 | 1.00E+00 | 1.00E+00 | 7.36E+04 | 0.2316 | 5.663E-12 | 0.0027739 | 0.9 | NS | NS | NS | NS | 32.843 | 2.59E-09 | 0.0048401 |

| | | | | | | | | | | | | | | | | | | | | | | | | | | | | | | | | | | | | | |
|---|---|---|---|---|---|---|---|---|---|---|---|---|---|---|---|---|---|---|---|---|---|---|---|---|---|---|---|---|---|---|---|---|---|---|---|---|---|
| | | | | | | | | | | | | | | | | | | | | | | | | | | 13780(100%);IPR017853(100%) | | | | | | | | | | | |
| 137 | Variable 221 | EQVGLSYDSHAR | Bacteroides | Bacteria | Bacteroidetes | | Bacteroidia | | Bacteroidales | | Bacteroidaceae | Bacteroides | | | | | | | | GO:0005737(100%) | cytoplasm(100%) | GO:0003697(100%);G | single-stranded DNA binding(100%);ATP | GO:0006260(100%);GO:0009432(100 | DNA replication(100%);SOS resp | IPR001238(100%);IP | 1.00E+00 | 3.71E+04 | 1.00E+00 | NS | NS | NS | NS | NS | NS | NS | NS |

| | | | | | | | | | | | | | | | | | | | | | | | | | | | | | | | | | | | | | |
|---|---|---|---|---|---|---|---|---|---|---|---|---|---|---|---|---|---|---|---|---|---|---|---|---|---|---|---|---|---|---|---|---|---|---|---|---|---|
| | | | | | | | | | | | | | | | | | | | | | | O:0005524(100%) | binding(100%) | %);GO:0006281(95%) | onse(100%);DNA repair(95%) | R0274177(100%);IPR0421774(100%) | | | | | | | | | | | |
| 138 | Variable 222 | TFAASHTVSETQK | Bacteroidesthetaiotaom | Bacteria | Bacteroidetes | | Bacteroidia | | Bacteroidales | | Bacteroidaceae | Bacteroides | | | Bacteroidesthetaiota | | | 6.3.2.1(100%) | Pantoate--beta-alanine ligase (AMP-for | GO:0005737(90%); | cytoplasm(90%);cytosol(10% | GO:0004592(100% | pantoate-beta-alanine ligase activity(10 | GO:0015940(100%) | pantothenate biosynthetic proces | IPR003721(100%); | 1.00E+00 | 1.00E+00 | 3.80E+04 | NS | NS | NS | NS | NS | NS | NS | NS |

| | | | | | | | | | | | | | | | | | | | | | | | | | | | | | | | | | | | | | |
|---|---|---|---|---|---|---|---|---|---|---|---|---|---|---|---|---|---|---|---|---|---|---|---|---|---|---|---|---|---|---|---|---|---|---|---|---|---|
| | | | icron | | | | | | | | | | | | omicron | | | | ming)(100%) | GO:0005829(10%) | ) | );GO:0005524(100%) | 0%);ATP binding(100%) | | s(100%) | IPR014729(100%);IPR042176(100%) | | | | | | | | | | | |
| 139 | Variable 223 | APFGGGNQAGR | Prevotellaruminicola | Bacteria | Bacteroidetes | | Bacteroidia | | Bacteroidales | | Prevotellaceae | Prevotella | | | Prevotellaruminic | | | 3.1.1.-(100%);3 | Carboxylic ester hydrolases(100%);F | | | GO:0016787(80 | hydrolase activity(80%);feruloyl est | GO:0045493(20%) | xylan catabolic processes(20 | IPR000801(100 | 1.00E+00 | 1.00E+00 | 1.67E+05 | 7.9198 | 0.043543 | 0.00015074 | 0.9 | NS | NS | NS | NS |

| | | | | | | | | | | | | | | | | | | | | | | | | | | | | | | | | | | | | | |
|---|---|---|---|---|---|---|---|---|---|---|---|---|---|---|---|---|---|---|---|---|---|---|---|---|---|---|---|---|---|---|---|---|---|---|---|---|---|
| | | | | | | | | | | | | | | | ola | | | .1.1.73(100%) | eruloyl esterase (100%) | | | %);GO:0030600(20%) | erase activity (20%) | | %) | %);IPR005181(100%);IPR013783(100%) | | | | | | | | | | | |
| 140 | Variable 22 | QQLFDLEQDMHDLTK | Chlamydiamurid | Bacteria | Chlamydiae | | Chlamydia | | Chlamydiales | | Chlamydiaceae | Chlamydia | | | Chlamydiamurid | | | | | | | | | | | IPR035336(1 | 1.00E+00 | 1.00E+00 | 1.84E+05 | NS | NS | NS | NS | NS | NS | NS | NS |

| | | | | | | | | | | | | | | | | | | | | | | | | | | | | | | | | | | | | | |
|---|---|---|---|---|---|---|---|---|---|---|---|---|---|---|---|---|---|---|---|---|---|---|---|---|---|---|---|---|---|---|---|---|---|---|---|---|---|
| | 4 | | arum | | | | | | | | | | | | arum | | | | | | | | | | | 00%) | | | | | | | | | | | |
| 141 | Variable 225 | AFHLNDSMFPLGANK | Chlamydiapneumoniae | Bacteria | Chlamydiae | | Chlamydia | | Chlamydiales | | Chlamydiaceae | Chlamydia | | | Chlamydiapneumoniae | | | 3.1.21.2(100%) | Deoxyribonuclease IV (100%) | | | GO:0003677(100%);GO:0008270(100%);GO:0008 | DNA binding (100%);zinc ion binding (100%);deoxyribonuclease IV (phage-T4-induced) activity (100%) | GO:0006281 (100%) | DNA repair (100%) | IPR001719(100%);IPR013022(100%);IPR018246 | 1.00E+00 | 1.00E+00 | 4.90E+05 | 21.008 | 0.017885 | 0 | 0.9 | NS | NS | NS | NS |

| | | | | | | | | | | | | | | | | | | | | | | | | | | | | | | | | | | | | | |
|---|---|---|---|---|---|---|---|---|---|---|---|---|---|---|---|---|---|---|---|---|---|---|---|---|---|---|---|---|---|---|---|---|---|---|---|---|---|
| | | | | | | | | | | | | | | | | | | | | | | 833(100%) | | | | (100%) | | | | | | | | | | | |
| 142 | Variable 226 | NVVVENPVPDGYAHSSGQR | Chlamydiatrachomatis | Bacteria | Chlamydiae | | Chlamydiia | | Chlamydiales | | Chlamydiaceae | Chlamydia | | | Chlamydiatrachomatis | | | | | GO:0042597(70%) | periplasmic space(70%) | GO:0005201(100%) | extracellular matrix structural constituent(100%) | GO:0008360(70%) | regulation of cell shape(70%) | IPR001434(100%);IPR003506(100%) | 1.00E+00 | 1.00E+00 | 9.59E+04 | NS | NS | NS | NS | NS | NS | NS | NS |
| 144 | Vari | AGMTVCAV | Candi | Bact | Chla | | Chlam | | Parachl | | Parachla | Cand | | | Cand | | | | | GO:0 | cytoplas | GO:0 | ATP bindin | GO:0042 | protein | IPR0 | 1.00 | 1.00 | 3.81 | 4.57 | 0.00 | 0.000 | 0.9 | NS | NS | NS | NS |

| | | | | | | | | | | | | | | | | | | | | | | | | | | | | | | | | |
|---|---|---|---|---|---|---|---|---|---|---|---|---|---|---|---|---|---|---|---|---|---|---|---|---|---|---|---|---|---|---|---|---|
| | able 228 | K | datusProtochlamydiaamoebophila | eria | mydiae | | ydia | | amydiales | | mydiaceae | idatusProtochlamydia | | | idatusProtochlamydiaamoebophila | | | | | 0057377(100%) | m(100%) | 0055244(100%);GO:0016887(100%);GO:0051082(100% | g(100%);ATPase(100%);unfolded protein binding(100%) | 026(100%) | refolding(100%) | 01844(100%);IPR002423(100%);IPR027409(100%) | E+00 | E+00 | E+04 | 99 | 27762 | 10238 |

| | | | | | | | | | | | | | | | | | | | | | | | | | | | | | | | | | | | | | |
|---|---|---|---|---|---|---|---|---|---|---|---|---|---|---|---|---|---|---|---|---|---|---|---|---|---|---|---|---|---|---|---|---|---|---|---|---|---|
| | | | | | | | | | | | | | | | | | | | | | | ) | | | | | | | | | | | | | | | |
| 146 | Variable 23 | HTWGEVHER | Actinomycetia | Bacteria | Actinobacteria | | Actinomycetia | | | | | | | | | | | 6.2.1.-(85%);2.3.1.86(8%);6.2.1.20(4%) | Acid--thiol ligases(85%);Fatty-acyl-CoA synthase system(8%);Long-chain-fatty-acid--[acyl-carrier-protein] ligase(4 | GO:0005618(0%);GO:0005886(0%) | cell wall(0%);plasma membrane(0%) | GO:0016874(86%);GO:0004321(1%);GO:0004467(0% | ligase activity(86%);fatty-acyl-CoA synthase activity(1%);long-chain fatty acid-CoA ligase activity(0%) | GO:0008610(97%);GO:0006633(0%) | lipid biosynthetic process(97%);fatty acid biosynthetic process(0%) | IPR042099(100%);IPR000873(100%);IPR040097(97 | 1.00E+00 | 1.00E+00 | 5.75E+04 | NS | NS | NS | NS | NS | NS | NS | NS |

| | | | | | | | | | | | | | | | | | | | | | | | | | | | | | | | | | | | | | |
|---|---|---|---|---|---|---|---|---|---|---|---|---|---|---|---|---|---|---|---|---|---|---|---|---|---|---|---|---|---|---|---|---|---|---|---|---|---|
| | | | | | | | | | | | | | | | | | | | %) | | | ) | | | | %) | | | | | | | | | | | |
| 147 | Variable 230 | CAFCVVPFTR | Chlorobiaceae | Bacteria | Chlorobi | | Chlorobia | | Chlorobiales | | Chlorobiaceae | | | | | | | 2.8.4.3(100%) | tRNA-2-methylthio-N(6)-dimethylallyladenosine synthase(100%) | GO:0005737(100%) | cytoplasm(100%) | GO:0035596(100%);GO:0046872(100%);GO:005153 | methylthiotransferase activity(100%);metal ion binding(100%);4 iron | 4 sulfur cluster binding(100%) | GO:0006400(100%) | tRNA modification(100%) | 1.00E+00 | 1.00E+00 | 5.29E+04 | NS | NS | NS | NS | NS | NS | NS | NS |

| | | | | | | | | | | | | | | | | | | | | | | | | | | | | | | | | | | | | | |
|---|---|---|---|---|---|---|---|---|---|---|---|---|---|---|---|---|---|---|---|---|---|---|---|---|---|---|---|---|---|---|---|---|---|---|---|---|---|
| | | | | | | | | | | | | | | | | | | | | | | 9(100%) | | | | | | | | | | | | | | | |
| 148 | Variable 231 | ALGLGRPNHK | Chlorobaculum | Bacteria | Chlorobi | | Chlorobia | | Chlorobiales | | Chlorobiaceae | Chlorobaculum | | | | | | | | GO:0015934(100%) | large ribosomal subunit (100%) | GO:0003735(100%) | structural constituent of ribosome (100%) | GO:0006412(100%) | translation (100%) | IPR005996(100%);IPR016082(100%);IPR036 | 1.00E+00 | 8.68E+05 | 1.00E+00 | NS | NS | NS | NS | NS | NS | NS | NS |

| | | | | | | | | | | | | | | | | | | | | | | | | | | | | | | | | | | | | | |
|---|---|---|---|---|---|---|---|---|---|---|---|---|---|---|---|---|---|---|---|---|---|---|---|---|---|---|---|---|---|---|---|---|---|---|---|---|---|
| | | | | | | | | | | | | | | | | | | | | | | | | | | 919(100%) | | | | | | | | | | | |
| 149 | Variable 232 | APALDLTLPGR | Chlorobiumlimicola | Bacteria | Chlorobi | | Chlorobia | | Chlorobiales | | Chlorobiaceae | Chlorobium | | | Chlorobium limicola | | | 6.1.1.20(100%) | Phenylalanine--tRNA ligase (100%) | GO:0005737(100%) | cytoplasm(100%) | GO:0000049(100%);GO:0000287(100%);GO | tRNA binding(100%);magnesium ion binding(100%);phenylalanine-tRNA ligase activity(10 | GO:0006432(100%) | phenylalanyl-tRNA aminoacylation(100%) | IPR002319(100%);IPR004188(100%);IPR0 | 1.00E+00 | 1.00E+00 | 4.41E+04 | NS | NS | NS | NS | NS | NS | NS | NS |

| | | | | | | | | | | | | | | | | | | | | | | | | | | | | | | | | | | | | | |
|---|---|---|---|---|---|---|---|---|---|---|---|---|---|---|---|---|---|---|---|---|---|---|---|---|---|---|---|---|---|---|---|---|---|---|---|---|---|
| | | | | | | | | | | | | | | | | | | | | | | :0004826(100%) | 0%) | | | 04529(100%) | | | | | | | | | | | |
| 151 | Variable 234 | HLSGGRPLK | Chlorobiumphaeovibrioides | Bacteria | Chlorobi | | Chlorobia | | Chlorobiales | | Chlorobiaceae | Chlorobium | | | Chlorobium phaeovibrioides | | | 3.6.1.27(100%) | Undecaprenyl-diphosphate phosphatase(100%) | GO:0005886(100%);GO:0016021(10 | plasma membrane(100%);integral component of membrane | GO:0050380(100%) | undecaprenyl-diphosphatase activity(100%) | GO:0008360(100%);GO:0009252(100%);GO:0016311(100%) | regulation of cell shape(100%); peptidoglycan biosynthetic proc | IPR003824(100%) | 1.00E+00 | 1.52E+06 | 1.00E+00 | NS | NS | NS | NS | NS | NS | NS | NS |

| | | | | | | | | | | | | | | | | | | | | | | | | | | | | | | | | | | | | | |
|---|---|---|---|---|---|---|---|---|---|---|---|---|---|---|---|---|---|---|---|---|---|---|---|---|---|---|---|---|---|---|---|---|---|---|---|---|---|
| | | | | | | | | | | | | | | | | | | | | 0%) | (100%) | | | | es s (100%); dephosphorylation (100%) | | | | | | | | | | | | |
| 152 | Variable 235 | DEELEGVEFDAYAR | Chloroherpetonthalassium ATCC35111 | Bacteria | Chlorobi | | Chlorobia | | Chlorobiales | | Chlorobiaceae | Chloroherpeton | | | Chloroherpetonthalassium | | Chloroherpetonthalassium ATCC | 3.4.21.53(100%) | EndopeptidaseLa(100%) | GO:0005737(100%) | cytoplasm(100%) | GO:0004176(100%);GO:000425 | ATP-dependentpeptidase activity(100%);serine-type endopeptidas | GO:0006515(100%);GO:0034605(100%) | protein quality control for misfolded or incompletel | IPR003111(100%);IPR003593( | 1.00E+00 | 7.08E+04 | 1.00E+00 | NS | NS | NS | NS | NS | NS | NS | NS |

| | | | | | | | | | | | | | | | | | | | | | | | | | | | | | | | | | | | | | |
|---|---|---|---|---|---|---|---|---|---|---|---|---|---|---|---|---|---|---|---|---|---|---|---|---|---|---|---|---|---|---|---|---|---|---|---|---|---|
| | | | 0 | | | | | | | | | | | | | | 35110 | | | | | 2(100%);GO:0005524(100%) | e activity(100%);ATP binding(100%) | | y synthesized proteins(100%);cellular response to heat(100%) | 100%);IPR003959(100%) | | | | | | | | | | | |
| 154 | Variable 237 | DAYMLQAVDEGTLK | PelodictyonluteolumD | Bacteria | Chlorobi | | Chlorobia | | Chlorobiales | | Chlorobiaceae | Pelodictyon | | | Pelodictyonluteol | | Pelodictyonluteol | 2.1.1.182(100% | 16S rRNA(adenine(1518)-N(6)/adenin | GO:0005737(100 | cytoplasm(100%) | GO:0003723(100 | RNA binding(100%);16S rRNA(adeni | | | IPR001737(100% | 1.00E+00 | 1.00E+00 | 2.66E+04 | NS | NS | NS | NS | NS | NS | NS | NS |

| | | | | | | | | | | | | | | | | | | | | | | | | | | | | | | | | | | | | | | | | |
|---|---|---|---|---|---|---|---|---|---|---|---|---|---|---|---|---|---|---|---|---|---|---|---|---|---|---|---|---|---|---|---|---|---|---|---|---|---|---|---|---|
| | | | SM 273 | | | | | | | | | | | | um | | um DSM 273 | ) | e(1519)-N(6))-dimethyltransferase(100%) | %) | | %);GO:0052908(100%) | ne(1518)-N(6)/adenine(1519)-N(6))-dimethyltransferase activity(100%) | | | );IPR011530(100%);IPR020596(100%) | | | | | | | | | | | | | | |
| 155 | Variable 238 | AAEQDAPK | Pelodictyonphae | Bacteria | Chlorobi | | Chlorobia | | Chlorobiales | | Chlorobiaceae | Pelodictyon | | | Pelodictyonpha | | Pelodictyonpha | 2.7.7.8(100% | Polyribonucleotide nucleotidyltrans | GO:0005737(1 | cytoplasm(100%) | GO:0000287(1 | magnesium ion binding(100%);RN | GO:0006396(100%);GO:00 | RNA processing(100%); | IPR001247(10 | 1.00E+00 | 1.00E+00 | 3.35E+04 | 9.6572 | 1.491E-08 | 0.00079672 | 0.9 | NS | NS | NS | NS | 20.283 | 6.85E-07 | 0.0053735 |

| | | | | | | | | | | | | | | | | | | | | | | | | | | | | | | | | | | | | | |
|---|---|---|---|---|---|---|---|---|---|---|---|---|---|---|---|---|---|---|---|---|---|---|---|---|---|---|---|---|---|---|---|---|---|---|---|---|---|
| | | | oclathratiformeBU-1 | | | | | | | | | | | | eoclathratiforme | | eoclathratiformeBU-1 | ) | ferase(100%) | 00%) | | 00%);GO:0003723(100%);GO:0004654(100%) | A binding (100%);polyribonucleotide nucleotidyltransferase activity (100%) | 06402(100%) | mRNA catabolic process (100%) | 0%);IPR003029(100%);IPR004087(100%) | | | | | | | | | | | |
| 156 | Variabl | AAGPSSTTDDLYR | Prosthec | Bacteri | Chlorob | | Chlorobia | | Chlorobiales | | Chlorobiaceae | Prosthec | | | | | | | | GO:0032 | mismatch repa | GO:0005 | ATP binding (100% | GO:0006298(1 | mismatch repa | IPR0020 | 1.00E+0 | 1.00E+0 | 5.24E+0 | NS | NS | NS | NS | NS | NS | NS | NS |

| | | | | | | | | | | | | | | | | | | | | | | | | | | | | | | | | | | | | | | | | |
|---|---|---|---|---|---|---|---|---|---|---|---|---|---|---|---|---|---|---|---|---|---|---|---|---|---|---|---|---|---|---|---|---|---|---|---|---|---|---|---|---|
| | e 239 | | ochloris | a | i | | | | | | | ochloris | | | | | | | | 300(100%) | ircomplex(100%) | 524(100%);GO:0016887(100%);GO:0030983(100%) | );ATPase(100%);mismatched DNA binding(100%) | 00%) | ir(100%) | 99(100%);IPR013507(100%);IPR014721(100%) | 0 | 0 | 4 | | | | | | | | | | | |
| 15 | Va | VF VT | Act | Ba | Ac | | Act | | | | | | | | | | | 6. | Propio | GO | cell | GO | ligase | GO: | fatty | s | 1. | 1. | 3. | NS | NS | NS | NS | NS | NS | NS | NS | 4. | 1. | 0. |

| | | | | | | | | | | | | | | | | | | | | | | | | | | | | | | | | | | | | | | |
|---|---|---|---|---|---|---|---|---|---|---|---|---|---|---|---|---|---|---|---|---|---|---|---|---|---|---|---|---|---|---|---|---|---|---|---|---|---|---|
| 7 | riable 24 | GPDVVR | inomycetia | cteria | tinobacteria | | inomycetia | | | | | | | | | | | 4.1.3(96%);2.1.3.-(3%);2.1.3.15(3%) | nyl-CoA carboxylase (96%); Carboxy-and carbamoyltransferases (3%); Acetyl-CoA carboxytransferase (3%) | :0005618(0%);GO:0009317(0%) | wall(0%);acetyl-CoA carboxylase complex(0%) | :0016874(86%);GO:0016740(19%);GO:0004658(14%) | activity(86%);transferase activity(19%);propionyl-CoA carboxylase activity(14%) | 00193367(0%) | acid elongation | aturated fatty acid(0%) | 00E+00 | 00E+00 | 55E+05 | | | | | | | | | 4423 | 78E-02 | 00011905 |

| | | | | | | | | | | | | | | | | | | | | | | | | | | | | | | | | | | | | | | | | |
|---|---|---|---|---|---|---|---|---|---|---|---|---|---|---|---|---|---|---|---|---|---|---|---|---|---|---|---|---|---|---|---|---|---|---|---|---|---|---|---|---|
| 158 | Variable 240 | EAGCSACLGMNDDK | Chloroflexales | Bacteria | Chloroflexi | | Chloroflexia | | Chloroflexales | | | | | | | | | 4.2.1.33(100%) | 3-isopropylmalate dehydratase (100%) | | | GO:0003861(100%);GO:0046872(100%);GO:0051539( | 3-isopropylmalate dehydratase activity (100%);metal ion binding (100%);4 iron | 4 sulfur cluster binding (100%) | GO:0009082(100%);GO:0009098(100%) | branched-chain amino acid biosynthetic process (100%);leucin | 1.00E+00 | 1.00E+00 | 1.97E+05 | NS | NS | NS | NS | NS | NS | NS | NS | 5.384 | 8.29E-03 | 0.00029871 |

| | | | | | | | | | | | | | | | | | | | | | | | | | | | | | | | | | | | | | |
|---|---|---|---|---|---|---|---|---|---|---|---|---|---|---|---|---|---|---|---|---|---|---|---|---|---|---|---|---|---|---|---|---|---|---|---|---|---|
| | | | | | | | | | | | | | | | | | | | | | | 100%) | | | | ebiosynthetic process(100%) | | | | | | | | | | | |
| 159 | Variable 241 | VLVEGQGQTDDGR | Roseiflexus castenholzii DSM | Bacteria | Chloroflexi | | Chloroflexia | | Chloroflexales | Roseiflexineae | Roseiflexaceae | Roseiflexus | | | Roseiflexus castenholzii | | Roseiflexus castenholzii DSM | 2.8.4.4(100%) | [Ribosomal protein S12] (aspartate(89)-C(3))-methylthiotrans | GO:0005737(100%) | cytoplasm(100%) | GO:0046872(100%);GO:005 | metal ion binding(100%);4iron | 4 sulfur cluster binding(100%);protein methylthiotransferas | GO:0006400(100%);GO:0018339(100%) | tRNA modification(100%);peptid | 1.00E+00 | 1.00E+00 | 5.59E+04 | NS | NS | NS | NS | NS | NS | NS | NS |

| | | | | | | | | | | | | | | | | | | | | | | | | | | |
|---|---|---|---|---|---|---|---|---|---|---|---|---|---|---|---|---|---|---|---|---|---|---|---|---|---|---|
| | | | 13941 | | | | | | | | | | | | | | 13941 | | ferase(100%) | | | 1539(100%);GO:0103039(100%) | | e activity(100%) | | yl-L-beta-methylthioaspartic acid biosynthetic process from peptid |

| | | | | | | | | | | | | | | | | | | | | | | | | | | | | | | | | | | | | | |
|---|---|---|---|---|---|---|---|---|---|---|---|---|---|---|---|---|---|---|---|---|---|---|---|---|---|---|---|---|---|---|---|---|---|---|---|---|---|
| | | | | | | | | | | | | | | | | | | | | | | | | | | yl-aspartic acid (100%) | | | | | | | | | | | |
| 160 | Variable 242 | YGPEAAEAYR | Thermomicrobium roseum DSM 5159 | Bacteria | Chloroflexi | | Thermomicrobia | | Thermomicrobiales | | Thermomicrobiaceae | Thermomicrobium | | | Thermomicrobium roseum | | Thermomicrobium roseum DSM 5 | 7.1.1.-(100%) | Hydron translocation or charge separation linked to oxidoreductase reactions | GO:0005886(100%) | plasma membrane (100%) | GO:0005506(100%);GO:0008137 | iron ion binding (100%);NADH dehydrogenase (ubiquinone) activity (10 | | | IPR006137(100%);IPR006138(1 | 1.00E+00 | 1.00E+00 | 3.56E+04 | NS | NS | NS | NS | NS | NS | NS | NS |

| | | | | | | | | | | | | | | | | | | | | | | | | | | | | | | | | | | | | | | | | |
|---|---|---|---|---|---|---|---|---|---|---|---|---|---|---|---|---|---|---|---|---|---|---|---|---|---|---|---|---|---|---|---|---|---|---|---|---|---|---|---|---|
| | | | | | | | | | | | | | | | | | 159 | | (100%) | | | (100%);GO:0048038(100%) | 0%);quinone binding(100%) | | | 00%) | | | | | | | | | | | | | | |
| 161 | Variable 243 | NDSMQSFR | Cyanobacteria | Bacteria | Cyanobacteria | | | | | | | | | | | | | | | GO:0005737(100%) | cytoplasm(100%) | GO:0003688(100%);GO:00 | DNA replication origin binding(100%);ATP binding(100% | GO:0006270(100%);GO:0006275(100%) | DNA replication initiation(100%);regulation of | IPR001957(100%);IPR003 | 1.00E+00 | 1.00E+00 | 2.33E+05 | 0.16843 | 7.6367E-13 | 0.0018745 | 0.9 | NS | NS | NS | NS | 30.647 | 6.20E-09 | 0.0019321 |

| | | | | | | | | | | | | | | | | | | | | | | | | | | | | | | | | | | | | | |
|---|---|---|---|---|---|---|---|---|---|---|---|---|---|---|---|---|---|---|---|---|---|---|---|---|---|---|---|---|---|---|---|---|---|---|---|---|---|
| | | | | | | | | | | | | | | | | | | | | | | 05524(100%) | ) | | DNA replication(100%) | 593(100%);IPR010921(100%) | | | | | | | | | | | |
| 162 | Variable 244 | LTGFTPLLR | Cyanobacteria | Bacteria | Cyanobacteria | | | | | | | | | | | | | | | GO:0005737(100%) | cytoplasm(100%) | GO:0051082(100%) | unfolded protein binding(100%) | GO:0006457(100%) | protein folding(100%) | IPR000397(100%);IPR0 | 1.00E+00 | 1.00E+00 | 7.80E+04 | NS | NS | NS | NS | NS | NS | NS | NS |

| | | | | | | | | | | | | | | | | | | | | | | | | | | | | | | | | | | | | | | | | |
|---|---|---|---|---|---|---|---|---|---|---|---|---|---|---|---|---|---|---|---|---|---|---|---|---|---|---|---|---|---|---|---|---|---|---|---|---|---|---|---|---|
| | | | | | | | | | | | | | | | | | | | | | | | | | | 16153(100%);IPR016154(100%) | | | | | | | | | | | | | | |
| 164 | Variable 246 | MVYQPAAGAR | Nostocales | Bacteria | Cyanobacteria | | | | Nostocales | | | | | | | | | | | GO:0005737(100%) | cytoplasm(100%) | GO:0016757(74%);GO | glycosyltransferase activity(74%);aminoacyl-tR | GO:0000105(100%) | histidine biosynthetic process(100% | IPR004516(100%);IP | 1.00E+00 | 1.00E+00 | 4.64E+04 | 0.43599 | 3.5256E-09 | 0.00065083 | 0.9 | NS | NS | NS | NS | 13.977 | 2.22E-05 | 0.00046905 |

| | | | | | | | | | | | | | | | | | | | | | | | | | | | | | | | | | | | | | |
|---|---|---|---|---|---|---|---|---|---|---|---|---|---|---|---|---|---|---|---|---|---|---|---|---|---|---|---|---|---|---|---|---|---|---|---|---|---|
| | | | | | | | | | | | | | | | | | | | | | | :0004812(24%);GO:0016874(6%) | NA ligase activity (24%);ligase activity (6%) | | ) | R004517(100%);IPR041715(100%) | | | | | | | | | | | |
| 166 | Variable 248 | EYEVLTSAEHK | Synechocystis | Bacteria | Cyanobacteria | | | | Synechococcales | | Merismopediaceae | Synechocystis | | | | | | | | GO:0031676(100% | plasma membrane-derived thyl | GO:0005524(100% | ATP binding (100%);helicase activity (50 | GO:0006605(100%);GO:0017038 | protein targeting (100%); protei | IPR000185(100%); | 1.00E+00 | 1.00E+00 | 1.08E+05 | NS | NS | NS | NS | NS | NS | NS | NS |

| | | | | | | | | | | | | | | | | | | | | | | | | | | |
|---|---|---|---|---|---|---|---|---|---|---|---|---|---|---|---|---|---|---|---|---|---|---|---|---|---|---|
| | | | | | | | | | | | | | | | | | | | | );GO:0005829(50%);GO:0005887(50%) | akoid membrane(100%);cytosol(50%);integral component of plasma membrane(50%) | );GO:0004386(50%);GO:0015462(50%) | %);ABC-type protein transporter activity(50%) | (100%);GO:0065002(100%) | n import(100%);intracellular protein transmembrane transport(100%) | IPR011115(100%);IPR011116(100%) |

| | | | | | | | | | | | | | | | | | | | | | | | | | | | | | | | | | | | | | |
|---|---|---|---|---|---|---|---|---|---|---|---|---|---|---|---|---|---|---|---|---|---|---|---|---|---|---|---|---|---|---|---|---|---|---|---|---|---|
| 167 | Variable 249 | ADPQQDFLSVDGK | Synechocystis | Bacteria | Cyanobacteria | | | | Synechococcales | | Merismopediaceae | Synechocystis | | | | | | 5.4.99.-(100%) | Transferring other groups (100%) | | | GO:0003723(100%);GO:0009982(100%) | RNA binding (100%);pseudouridine synthase activity (100%) | GO:0001522 (100%) | pseudouridine synthesis (100%) | IPR000748(100%);IPR002942(100%);IPR006145(100 | 1.00E+00 | 3.77E+05 | 1.00E+00 | NS | NS | NS | NS | NS | NS | NS | NS |

| | | | | | | | | | | | | | | | | | | | | | | | | | | | | | | | | | | | | | |
|---|---|---|---|---|---|---|---|---|---|---|---|---|---|---|---|---|---|---|---|---|---|---|---|---|---|---|---|---|---|---|---|---|---|---|---|---|---|
| | | | | | | | | | | | | | | | | | | | | | | | | | | %) | | | | | | | | | | | |
| 168 | Variable 25 | MDWPTGTR | Actinomycetia | Bacteria | Actinobacteria | | Actinomycetia | | | | | | | | | | | 2.1.1.-(100%) | Methyltransferases(100%) | | | GO:0008168(100%) | methyltransferase activity(100%) | GO:0032259(100%) | methylation(100%) | IPR007213(100%);IPR011610(100%);IPR029063(1 | 1.00E+00 | 1.00E+00 | 5.07E+04 | NS | NS | NS | NS | NS | NS | NS | NS |

| | | | | | | | | | | | | | | | | | | | | | | | | | | | | | | | | | | | | | |
|---|---|---|---|---|---|---|---|---|---|---|---|---|---|---|---|---|---|---|---|---|---|---|---|---|---|---|---|---|---|---|---|---|---|---|---|---|---|
| | | | | | | | | | | | | | | | | | | | | | | | | | | 00%) | | | | | | | | | | | |
| 169 | Variable 250 | AKPGDQCFYDNK | Prochlorococcus marinus str. MIT 9211 | Bacteria | Cyanobacteria | | | | Synechococcales | | Prochloraceae | Prochlorococcus | | | Prochlorococcus marinus | | Prochlorococcus marinus str. MIT 9211 | 6.1.1.15 (100%) | Proline--tRNA ligase (100%) | GO:0005737 (100%) | cytoplasm (100%) | GO:0002161(100%);GO:0004827(100%);GO:0005 | aminoacyl-tRNA editing activity (100%);proline-tRNA ligase activity (100%);ATP binding (100%) | GO:0006433 (100%) | prolyl-tRNA aminoacylation (100%) | IPR002314(100%);IPR002316(100%);IPR004154 | 1.00E+00 | 1.00E+00 | 6.56E+04 | NS | NS | NS | NS | NS | NS | NS | NS |

| | | | | | | | | | | | | | | | | | | | | | | | | | | | | | | | | | | | | | |
|---|---|---|---|---|---|---|---|---|---|---|---|---|---|---|---|---|---|---|---|---|---|---|---|---|---|---|---|---|---|---|---|---|---|---|---|---|---|
| | | | | | | | | | | | | | | | | | | | | | | 524(100%) | | | | (100%) | | | | | | | | | | | |
| 171 | Variable 252 | SGGGLGELLVLEPR | Synechococcaceae | Bacteria | Cyanobacteria | | | | Synechococcales | | Synechococcaceae | | | | | | | 2.4.99.17(100%) | S-adenosylmethionine:tRNA ribosyltransferase-isomerase (100%) | GO:0005737(100%) | cytoplasm(100%) | GO:0016853(100%);GO:0016740(94%);GO: | isomerase activity(100%);transferase activity(94%);glycosyltransferase activity(6%) | GO:0008616(100%) | queuosine biosynthetic process(100%) | IPR003699(100%);IPR036100(100%);IPR0 | 1.00E+00 | 1.00E+00 | 6.44E+04 | NS | NS | NS | NS | NS | NS | NS | NS |

| | | | | | | | | | | | | | | | | | | | | | | | | | | | | | | | | | | | | | |
|---|---|---|---|---|---|---|---|---|---|---|---|---|---|---|---|---|---|---|---|---|---|---|---|---|---|---|---|---|---|---|---|---|---|---|---|---|---|
| | | | | | | | | | | | | | | | | | | | | | | 0016757(6%) | | | | 42118(100%) | | | | | | | | | | | |
| 172 | Variable 253 | LLVVPPAER | Synechococcaceae | Bacteria | Cyanobacteria | | | | Synechococcales | | Synechococcaceae | | | | | | | 2.4.99.17(100%) | S-adenosylmethionine:tRNA ribosyltransferase-isomerase(100%) | GO:0005737(100%) | cytoplasm(100%) | GO:0016740(100%);GO:0016853(100%) | transferase activity(100%);isomerase activity(100%) | GO:0008616(100%) | queuosine biosynthetic process(100%) | IPR003699(100%);IPR036100(100%);IP | 1.00E+00 | 1.96E+05 | 1.00E+00 | NS | NS | NS | NS | NS | NS | NS | NS |

| | | | | | | | | | | | | | | | | | | | | | | | | | | | | | | | | | | | | | |
|---|---|---|---|---|---|---|---|---|---|---|---|---|---|---|---|---|---|---|---|---|---|---|---|---|---|---|---|---|---|---|---|---|---|---|---|---|---|
| | | | | | | | | | | | | | | | | | | | | | | | | | | R0421118(100%) | | | | | | | | | | | |
| 173 | Variable 254 | EQPDMVTLVPER | Synechococcaceae | Bacteria | Cyanobacteria | | | | Synechococcales | | Synechococcaceae | | | | | | | 2.6.99.2(100%) | Pyridoxine 5'-phosphate synthase (100%) | GO:0005737(100%) | cytoplasm (100%) | GO:0033856(60%);GO:0016769(40%) | pyridoxine 5'-phosphate synthase activity (60%);transferase activity | transferring nitrogenous groups (40%) | GO:0008615(100%) | pyridoxine biosynthetic process (100%) | 1.00E+00 | 1.09E+05 | 1.00E+00 | NS | NS | NS | NS | NS | NS | NS | NS |

| | | | | | | | | | | | | | | | | | | | | | | | | | | | | | | | | | | | | | |
|---|---|---|---|---|---|---|---|---|---|---|---|---|---|---|---|---|---|---|---|---|---|---|---|---|---|---|---|---|---|---|---|---|---|---|---|---|---|
| 174 | Variable 255 | HGTVWDWQNELR | Synechococcus | Bacteria | Cyanobacteria | | | | Synechococcales | | Synechococcaceae | Synechococcus | | | | | | 2.4.99.17(100%) | S-adenosylmethionine:tRNA ribosyltransferase-isomerase (100%) | GO:0005737(100%) | cytoplasm(100%) | GO:0016740(100%);GO:0016853(100%) | transferase activity(100%);isomerase activity(100%) | GO:0008616(100%) | queuosine biosynthetic process(100%) | IPR003699(100%);IPR036100(100%);IPR042118(100 | 1.00E+00 | 1.00E+00 | 5.90E+04 | 3.521 | 0.0013173 | 0.00013844 | 0.9 | NS | NS | NS | NS |

| | | | | | | | | | | | | | | | | | | | | | | | | | | | | | | | | | | | | | | | | |
|---|---|---|---|---|---|---|---|---|---|---|---|---|---|---|---|---|---|---|---|---|---|---|---|---|---|---|---|---|---|---|---|---|---|---|---|---|---|---|---|---|
| | | | | | | | | | | | | | | | | | | | | | | | | | | %) | | | | | | | | | | | | | | |
| 175 | Variable 256 | CGYVNPDQVTFEYLK | Synechococcus | Bacteria | Cyanobacteria | | | | Synechococcales | | Synechococcaceae | Synechococcus | | | | | | 4.2.1.33(100%) | 3-isopropylmalate dehydratase (100%) | | | GO:0003861(100%);GO:0046872(67%);GO:0051539 | 3-isopropylmalate dehydratase activity (100%);metal ion binding (67%);4 iron | 4 sulfur cluster binding (67%) | GO:0009098 (100%); GO:0009082 (67%) | leucine biosynthetic process (100%); branched-chain amino | 1.00E+00 | 1.00E+00 | 3.03E+04 | NS | NS | NS | NS | NS | NS | NS | NS | 4.8585 | 1.26E-02 | 0.000652 |

| | | | | | | | | | | | | | | | | | | | | | | | | | | | | | | | | | | | | | |
|---|---|---|---|---|---|---|---|---|---|---|---|---|---|---|---|---|---|---|---|---|---|---|---|---|---|---|---|---|---|---|---|---|---|---|---|---|---|
| | | | | | | | | | | | | | | | | | | | | | | (67%) | | | | acid biosynthetic process (67%) | | | | | | | | | | | |
| 176 | Variable 257 | MVLDLGLAVR | Synechococcus | Bacteria | Cyanobacteria | | | | Synechococcales | | Synechococcaceae | Synechococcus | | | | | | 2.7.4.25(100%);6.3.2.1 | (d)CMP kinase (100%);Pantoate--beta-alanine ligase (AMP- | GO:0005737(100%) | cytoplasm(100%) | GO:0004592(100%);GO:00 | pantoate-beta-alanine ligase activity (100%);ATP binding | GO:0006220(100%);GO:0015940(100%) | pyrimidine nucleotide metabolic process (100 | IPR003136(100%);IPR003 | 1.00E+00 | 1.00E+00 | 1.52E+05 | NS | NS | NS | NS | NS | NS | NS | NS |

| | | | | | | | | | | | | | | | | | | | | | | | | | | | | | | | | | | | | | |
|---|---|---|---|---|---|---|---|---|---|---|---|---|---|---|---|---|---|---|---|---|---|---|---|---|---|---|---|---|---|---|---|---|---|---|---|---|---|
| | | | | | | | | | | | | | | | | | | (100%) | forming)(100%) | | | 05524(100%);GO:0036430(75%) | (100%);CMP kinase activity (75%) | | %); pantothenate biosynthetic process (100%) | 721(100%);IPR011994(100%) | | | | | | | | | | | |
| 177 | Variable 258 | LDPPEEQSSLLHPGR | Synechococcus | Bacteria | Cyanobacteria | | | | Synechococcales | | Synechococcaceae | Synechococcus | | | | | | 6.1.1.20(100%) | Phenylalanine--tRNA ligase (100%) | GO:0005737(100%) | cytoplasm(100%) | GO:0000049(100%);G | tRNA binding (100%);magnesium ion binding | GO:0006432(100%) | phenylalanyl-tRNA aminoacylation | IPR002547(100%);IP | 1.00E+00 | 1.00E+00 | 5.00E+04 | NS | NS | NS | NS | NS | NS | NS | NS |

| | | | | | | | | | | | | | | | | | | | | | | | | | | | | | | | | | | | | | | | | |
|---|---|---|---|---|---|---|---|---|---|---|---|---|---|---|---|---|---|---|---|---|---|---|---|---|---|---|---|---|---|---|---|---|---|---|---|---|---|---|---|---|
| | | | | | | | | | | | | | | | | | | | | | | O:0000287(100%);GO:0004826(100%) | (100%);phenylalanine-tRNA ligase activity(100%) | | (100%) | R004532(100%);IPR005121(100%) | | | | | | | | | | | | | | |
| 178 | Variable 259 | LDGFDLGNSPMSVVPDTVK | Synechococcu | Bacteria | Cyanobacteria | | | | Synechococcales | | Synechococcaceae | Synechococcu | | | | | | 3.1.3.71(100 | 2-phosphosulfolactate phosphatase | | | GO:0000287(1 | magnesium ion binding(100%);2- | | | IPR005238(10 | 1.00E+00 | 1.00E+00 | 3.35E+04 | 4.0076 | 9.2623E-06 | 0.00023595 | 0.9 | NS | NS | NS | NS | 8.123 | 1.04E-03 | 0.00062398 |

| | | | | | | | | | | | | | | | | | | | | | | | | | | | | | | | | | | | | | | | | |
|---|---|---|---|---|---|---|---|---|---|---|---|---|---|---|---|---|---|---|---|---|---|---|---|---|---|---|---|---|---|---|---|---|---|---|---|---|---|---|---|---|
| | | | s | | | | | | | | | s | | | | | | %) | (100%) | | | 00%);GO:0050532(100%) | phosphosulfolactate phosphatase activity (100%) | | | 0%);IPR036702(100%) | | | | | | | | | | | | | | |
| 179 | Variable 26 | GLATGLGDEGGFAPNLGSNR | Actinomycetia | Bacteria | Actinobacteria | | Actinomycetia | | | | | | | | | | | 4.2.1.11(100%) | Phosphopyruvate hydratase (100%) | GO:0009986(99%);GO:00000 | cell surface (99%);phosphopyruvate hydratase com | GO:0000287(100%);GO:0004 | magnesium ion binding (100%);phosphopyruvate hydratase activit | GO:0006096(100%);GO:0032889(0%) | glycolytic process (100%);regulation of vacuole fu | non-autophagic (0%) | 1.00E+00 | 1.00E+00 | 1.93E+05 | 0.49483 | 4.5478E-08 | 0.0008971 | 0.9 | NS | NS | NS | NS | 9.6717 | 3.51E-04 | 0.00035347 |

| | | | | | | | | | | | | | | | | | | | | | | | | | | | | | | | | | | | | | |
|---|---|---|---|---|---|---|---|---|---|---|---|---|---|---|---|---|---|---|---|---|---|---|---|---|---|---|---|---|---|---|---|---|---|---|---|---|---|
| | | | | | | | | | | | | | | | | | | | | 15(98%);GO:0005576(98%) | plex(98%);extracellular region(98%) | 634(100%) | y(100%) | | sion | | | | | | | | | | | | |
| 180 | Variable 260 | TAAGETLQQGR | Synechococcus | Bacteria | Cyanobacteria | | | | Synechococcales | | Synechococcaceae | Synechococcus | | | | | | | | GO:0005737(100%) | cytoplasm(100%) | GO:0016151(100%) | nickel cation binding(100%) | GO:0006807(45%) | nitrogen compound metabolic process(45% | IPR002669(100%) | 1.00E+00 | 1.47E+05 | 1.00E+00 | NS | NS | NS | NS | NS | NS | NS | NS |

| | | | | | | | | | | | | | | | | | | | | | | | | | | | | | | | | | | | | | |
|---|---|---|---|---|---|---|---|---|---|---|---|---|---|---|---|---|---|---|---|---|---|---|---|---|---|---|---|---|---|---|---|---|---|---|---|---|---|
| | | | | | | | | | | | | | | | | | | | | | | | | | ) | | | | | | | | | | | | |
| 182 | Variable 262 | GMGESEFK | Rippkaeaorientalis PCC 8801 | Bacteria | Cyanobacteria | | | Oscillatoriophycideae | Chroococcales | | Aphanothecaceae | Rippkaea | | | Rippkaea orientalis | | Rippkaea orientalis PCC 8801 | 2.1.2.1 (100%) | Glycine hydroxymethyltransferase (100%) | GO:0005737 (100%) | cytoplasm (100%) | GO:0004372 (100%);GO:0030170 (100%) | glycine hydroxymethyltransferase activity (100%);pyridoxal phosphate binding (100%) | GO:0019264 (100%);GO:0035999 (100%) | glycine biosynthetic process from serine (100%);tetrahydrofolate interconversion (100%) | IPR001085 (100%);IPR015421 (100%);IPR015422 (10 | 1.00E+00 | 1.00E+00 | 6.15E+04 | NS | NS | NS | NS | NS | NS | NS | NS |

| | | | | | | | | | | | | | | | | | | | | | | | | | | | | | | | | | | | | | |
|---|---|---|---|---|---|---|---|---|---|---|---|---|---|---|---|---|---|---|---|---|---|---|---|---|---|---|---|---|---|---|---|---|---|---|---|---|---|
| | | | | | | | | | | | | | | | | | | | | | | | | | | 0%) | | | | | | | | | | | |
| 183 | Variable 263 | CPYCSAPDSK | Deinococcus | Bacteria | Deinococcus-Thermus | | Deinococci | | Deinococcales | | Deinococcaceae | Deinococcus | | | | | | | | | | GO:0005524(100%);GO:0008270(100%);GO:00036 | ATP binding(100%);zinc ion binding(100%);DNA binding(96%) | GO:0045892(100%) | negative regulation of transcription | DNA-templated(100%) | 1.00E+00 | 1.00E+00 | 2.58E+05 | NS | NS | NS | NS | NS | NS | NS | NS |

| | | | | | | | | | | | | | | | | | | | | | | | | | | | | | | | | | | | | | | | | |
|---|---|---|---|---|---|---|---|---|---|---|---|---|---|---|---|---|---|---|---|---|---|---|---|---|---|---|---|---|---|---|---|---|---|---|---|---|---|---|---|---|
| | | | | | | | | | | | | | | | | | | | | | | 77(96%) | | | | | | | | | | | | | | | | | | |
| 184 | Variable 264 | LGSGAPVPNVQLSALAVK | Deinococcus radiodurans R1 | Bacteria | Deinococcus-Thermus | | Deinococci | | Deinococcales | | Deinococcaceae | Deinococcus | | | Deinococcus radiodurans | | Deinococcus radiodurans R1 | | | | | GO:0004866(100%) | endopeptidase inhibitor activity (100%) | | | IPR001599(100%);IPR002890(100%);IPR011 | 1.00E+00 | 1.00E+00 | 7.75E+04 | 0.26307 | 1.294E-11 | 0.001449 | 0.9 | NS | NS | NS | NS | 25.642 | 5.28E-08 | 0.001256 |

| | | | | | | | | | | | | | | | | | | | | | | | | | | | | | | | | | | | | | |
|---|---|---|---|---|---|---|---|---|---|---|---|---|---|---|---|---|---|---|---|---|---|---|---|---|---|---|---|---|---|---|---|---|---|---|---|---|---|
| | | | | | | | | | | | | | | | | | | | | | | | | | | 625(100%) | | | | | | | | | | | |
| 185 | Variable 265 | ANLSPSATPCTPWK | Deinococcus radiodurans R1 | Bacteria | Deinococcus-Thermus | | Deinococci | | Deinococcales | | Deinococcaceae | Deinococcus | | | Deinococcus radiodurans | | Deinococcus radiodurans R1 | 1.1.1.35(100%) | 3-hydroxyacyl-CoA dehydrogenase (100%) | | | GO:0003857(100%);GO:0070403(100%) | 3-hydroxyacyl-CoA dehydrogenase activity (100%);NAD+ binding (100%) | GO:0006631 (100%) | fatty acid metabolic process (100%) | IPR006108(100%);IPR006176(100%);IPR0 | 1.00E+00 | 1.00E+00 | 5.80E+04 | NS | NS | NS | NS | NS | NS | NS | NS |

| | | | | | | | | | | | | | | | | | | | | | | | | | | | | | | | | | | | | | |
|---|---|---|---|---|---|---|---|---|---|---|---|---|---|---|---|---|---|---|---|---|---|---|---|---|---|---|---|---|---|---|---|---|---|---|---|---|---|
| | | | | | | | | | | | | | | | | | | | | | | | | | | 08927 (100%) | | | | | | | | | | | |
| 186 | Variable 266 | VGPPPAALETK | Deinococcus radiodurans R1 | Bacteria | Deinococcus-Thermus | | Deinococci | | Deinococcales | | Deinococcaceae | Deinococcus | | | Deinococcus radiodurans | | Deinococcus radiodurans R1 | 2.1.1.33 (100%) | tRNA (guanine(46)-N(7))-methyltransferase (100%) | GO:0043527 (100%) | tRNA methyltransferase complex (100%) | GO:0008176 (100%) | tRNA (guanine-N7-)-methyltransferase activity (100%) | GO:0030488 (100%);GO:0036265 (100%) | tRNA methylation (100%);RNA (guanine-N7)-methylation (100 | IPR003358 (100%);IPR029063 (100%) | 1.00E+00 | 1.00E+00 | 1.03E+05 | NS | NS | NS | NS | NS | NS | NS | NS |

| | | | | | | | | | | | | | | | | | | | | | | | | | | | | | | | | | | | | | |
|---|---|---|---|---|---|---|---|---|---|---|---|---|---|---|---|---|---|---|---|---|---|---|---|---|---|---|---|---|---|---|---|---|---|---|---|---|---|
| | | | | | | | | | | | | | | | | | | | | | | | | | %) | | | | | | | | | | | | |
| 187 | Variable 267 | LGLDVDVVPPGASLR | Thermus | Bacteria | Deinococcus-Thermus | | Deinococci | | Thermales | | Thermaceae | Thermus | | | | | | 3.2.1.23(100%) | Beta-galactosidase(100%) | GO:0009341(100%) | beta-galactosidase complex(100%) | GO:0004565(100%);GO:0046872(100%) | beta-galactosidase activity(100%);metal ion binding(100%) | GO:0006012(100%) | galactose metabolic process(100%) | IPR003476(100%);IPR013529(100%);IPR013738(1 | 1.00E+00 | 1.00E+00 | 9.27E+04 | 6.1699 | 0.007364 | 0.00025 | 0.9 | NS | NS | NS | NS |

| | | | | | | | | | | | | | | | | | | | | | | | | | | | | | | | | | | | | | |
|---|---|---|---|---|---|---|---|---|---|---|---|---|---|---|---|---|---|---|---|---|---|---|---|---|---|---|---|---|---|---|---|---|---|---|---|---|---|
| | | | | | | | | | | | | | | | | | | | | | | | | | | 00%) | | | | | | | | | | | |
| 190 | Variable 27 | ETASYVLESGSAR | Actinomycetia | Bacteria | Actinobacteria | | Actinomycetia | | | | | | | | | | | 1.13.11.27(100%) | 4-hydroxyphenylpyruvate dioxygenase (100%) | | | GO:0003868(100%);GO:0046872(96%) | 4-hydroxyphenylpyruvate dioxygenase activity (100%);metal ion binding (96%) | GO:0009072(95%);GO:0006559(1%);GO:0006572(1%) | aromatic amino acid family metabolic process (95%);L-phenylalanine catabolic pr | IPR004360(100%);IPR005956(100%);IPR029068 | 1.00E+00 | 1.00E+00 | 3.47E+04 | NS | NS | NS | NS | NS | NS | NS | NS |

| | | | | | | | | | | | | | | | | | | | | | | | | | | | | | | | | | | | | | |
|---|---|---|---|---|---|---|---|---|---|---|---|---|---|---|---|---|---|---|---|---|---|---|---|---|---|---|---|---|---|---|---|---|---|---|---|---|---|
| | | | | | | | | | | | | | | | | | | | | | | | | | oces s (1%);tyrosine catabolic process (1%) | (100%) | | | | | | | | | | | |
| 191 | Variable 270 | LEVETVLVK | Dictyoglomus turgidum DSM 672 | Bacteria | Dictyoglomi | | Dictyoglomia | | Dictyoglomales | | Dictyoglomaceae | Dictyoglomus | | | Dictyoglomus turgidum | | Dictyoglomus turgidum DS | 3.5.1.2(100%);4.3.3.6(1 | Glutaminase (100%);Pyridoxal 5'-phosphate synthase (glutami | GO:0005829(100%);GO:190 | cytosol (100%);glutaminase complex (100%) | GO:0004359(100%);GO:003 | glutaminase activity (100%);pyridoxal 5'-phosphate synthase | GO:0006543(100%);GO:0008614(100%);GO:0042 | glutamine catabolic process (100%);pyridoxine | IPR002161(100%);IPR0211 | 1.00E+00 | 2.68E+05 | 1.00E+00 | NS | NS | NS | NS | NS | NS | NS | NS |

| | | | | | | | | | | | | | | | | | | | | | | | | | | | | | | | | | | | | | |
|---|---|---|---|---|---|---|---|---|---|---|---|---|---|---|---|---|---|---|---|---|---|---|---|---|---|---|---|---|---|---|---|---|---|---|---|---|---|
| | | | 4 | | | | | | | | | | | | | | M 6724 | 00%) | nehydrolyzing)(100%) | 3600(100%) | | 6381(100%) | (glutaminehydrolysing)activity(100%) | 823(100%) | metabolicprocess(100%);pyridoxalphosphatebiosyntheticprocess(100%) | 96(100%);IPR029062(100%) | | | | | | | | | | | |
| 192 | Variab | ANEELCDFLSDFFDV | Candidat | Bacter | Elusim | | Endomic | | Endomic | | Endomicro | Endomic | | | Candid | | | | | | | | | | | IPR003 | 1.00E+ | 1.00E+ | 6.77E+ | NS | NS | NS | NS | NS | NS | NS | NS |

| | | | | | | | | | | | | | | | | | | | | | | | | | | | | | | | | | | | | | | | | |
|---|---|---|---|---|---|---|---|---|---|---|---|---|---|---|---|---|---|---|---|---|---|---|---|---|---|---|---|---|---|---|---|---|---|---|---|---|---|---|---|---|
| | le 271 | K | usEndomicrobiumtrichonymphae | ia | icrobia | | robia | | robiales | | biaceae | robium | | | atusEndomicrobium trichonymphae | | | | | | | | | | | 746(100%);IPR036591(100%) | 00 | 00 | 04 | | | | | | | | | | | |
| 193 | Variable 272 | NLNNDDEQPDVTK | CandidatusEndomicro | Bacteria | Elusimicrobia | | Endomicrobia | | Endomicrobiales | | Endomicrobiaceae | Endomicrobium | | | CandidatusEndomicr | | | 2.7.7.6(100%) | DNA-directed RNA polymerase (100%) | | | GO:0000287(100%); | magnesium ion binding(100%);DNA binding | GO:0006351(100%) | transcription | DNA-templated(100% | 1.00E+00 | 1.00E+00 | 2.71E+04 | 3.4607 | 1.189E-06 | 0.00045429 | 0.9 | NS | NS | NS | NS | 10.815 | 1.63E-04 | 0.0017573 |

| | | | | | | | | | | | | | | | | | | | | | | | | | | | | | | | | | | | | | |
|---|---|---|---|---|---|---|---|---|---|---|---|---|---|---|---|---|---|---|---|---|---|---|---|---|---|---|---|---|---|---|---|---|---|---|---|---|---|
| | | | bium trichonymphae | | | | | | | | | | | | obium trichonymphae | | | | | | | GO:0003677(100%);GO:0003899(100%) | (100%);DNA-directed 5'-3' RNA polymerase activity(100%) | | | ) | | | | | | | | | | | |
| 194 | Variable 27 | EYSTDTATK | Firmicutes | Bacteria | Firmicutes | | | | | | | | | | | | | 5.2.1.8(100 | Peptidylprolyl isomerase (100% | GO:0005886( | plasmamembrane(9 | GO:0003755( | peptidyl-prolyl cis-trans iso | GO:0006457(92%) | protein folding(92%) | IPR002297(9 | 1.00E+00 | 1.00E+00 | 4.60E+04 | 3.2902 | 0.001037 | 0.00031326 | 0.9 | NS | NS | NS | NS |

| | | | | | | | | | | | | | | | | | | | | | | | | | | | | | | | | | | | | | | | | |
|---|---|---|---|---|---|---|---|---|---|---|---|---|---|---|---|---|---|---|---|---|---|---|---|---|---|---|---|---|---|---|---|---|---|---|---|---|---|---|---|---|
| | 3 | | | | | | | | | | | | | | | | | %) | ) | 94%);GO:0016021(8%) | 4%);integral component of membrane (8%) | 100%) | merase activity (100%) | | | 8%);IPR023059(98%);IPR027304(96%) | | | | | | | | | | | | | | |
| 195 | Variable 274 | GGEFHAVMGPNGTGK | Firmicutes | Bacteria | Firmicutes | | | | | | | | | | | | | 3.6.1.3(100% | Adenosinetriphosphatase(100%) | | | GO:0005524(1 | ATP binding(100%);ATPase(94%) | | | IPR003439(10 | 1.00E+00 | 1.00E+00 | 3.27E+04 | 3.5194 | 2.2363E-06 | 0.00014854 | 0.9 | NS | NS | NS | NS | 10.017 | 2.77E-04 | 0.00097072 |

| | | | | | | | | | | | | | | | | | | | | | | | | | | | | | | | | | | | | | |
|---|---|---|---|---|---|---|---|---|---|---|---|---|---|---|---|---|---|---|---|---|---|---|---|---|---|---|---|---|---|---|---|---|---|---|---|---|---|
| | | | | | | | | | | | | | | | | | | ) | | | | 00%);GO:0016887(94%) | | | | 0%);IPR010230(100%);IPR027417(100%) | | | | | | | | | | | |
| 196 | Variable 2 | ELMAEVSK | Firmicutes | Bacteria | Firmicutes | | | | | | | | | | | | | 3.6.5.-(57% | Acting on GTP; involved in cell | GO:0005737 | cytoplasm(67%);int | GO:0000287 | magnesium ion binding(67%) | GO:0042254(67%) | ribosome biogenesis | IPR006073( | 1.00E+00 | 3.87E+05 | 1.55E+06 | NS | NS | NS | NS | NS | NS | NS | NS |

| | | | | | | | | | | | | | | | | | | | | | | | | | | |
|---|---|---|---|---|---|---|---|---|---|---|---|---|---|---|---|---|---|---|---|---|---|---|---|---|---|---|
| | 75 | | | | | | | | | | | | | | | | | );2.4.1.129(43%);3.4.16.4(43%) | ular and subcellular movement (57%);Peptidoglycan glycosyltransferase (43%);Serine-type D-Ala-D-Ala carboxypeptidase (43%) | (67%);GO:0016021(33%) | egral component of membrane (33%) | (67%);GO:0003924(67%);GO:0005525(67%) | ;GTPase activity (67%);GTP binding (67%) | | (67%) | 50%);IPR006074(50%);IPR006169(50%) |

| | | | | | | | | | | | | | | | | | | | | | | | | | | | | | | | | | | | | | |
|---|---|---|---|---|---|---|---|---|---|---|---|---|---|---|---|---|---|---|---|---|---|---|---|---|---|---|---|---|---|---|---|---|---|---|---|---|---|
| 199 | Variable 278 | GYAYESAGDVYYR | Bacilli | Bacteria | Firmicutes | | Bacilli | | | | | | | | | | | 6.1.1.16(100%) | Cysteine--tRNA ligase (100%) | GO:0005737(99%) | cytoplasm(99%) | GO:0004817(100%);GO:0005524(100%);GO:0008270( | cysteine-tRNA ligase activity (100%);ATP binding (100%);zinc ion binding (99%) | GO:0006423 (100%) | cysteinyl-tRNA aminoacylation (100%) | IPR014729(100%);IPR015803(100%);IPR024909(100 | 1.00E+00 | 1.00E+00 | 2.20E+05 | NS | NS | NS | NS | NS | NS | NS | NS |

| | | | | | | | | | | | | | | | | | | | | | | | | | | | | | | | | | | | | | |
|---|---|---|---|---|---|---|---|---|---|---|---|---|---|---|---|---|---|---|---|---|---|---|---|---|---|---|---|---|---|---|---|---|---|---|---|---|---|
| | | | | | | | | | | | | | | | | | | | | | | 99%) | | | | %) | | | | | | | | | | | |
| 200 | Variable 279 | LLQQTNSFK | Bacilli | Bacteria | Firmicutes | | Bacilli | | | | | | | | | | | 3.4.24.13(94%);3.4.24.-(6%) | IgA-specific metalloendopeptidase (94%); Metalloendopeptidases (6%) | GO:0005576(100%);GO:0005618(100%);GO:0016 | extracellular region(100%);cell wall(100%);integral component of membrane(96%) | GO:0004222(100%);GO:0008270(100%) | metalloendopeptidase activity (100%);zinc ion binding (100%) | GO:0009405 (1%) | obsolete pathogenesis (1%) | IPR011505(100%);IPR008006(96%);IPR011098( | 1.00E+00 | 1.00E+00 | 3.45E+04 | NS | NS | NS | NS | NS | NS | NS | NS |

| | | | | | | | | | | | | | | | | | | | | | | | | | | | | | | | | | | | | | |
|---|---|---|---|---|---|---|---|---|---|---|---|---|---|---|---|---|---|---|---|---|---|---|---|---|---|---|---|---|---|---|---|---|---|---|---|---|---|
| | | | | | | | | | | | | | | | | | | | | 021(96%) | | | | | | 84%) | | | | | | | | | | | |
| 201 | Variable 28 | AYLPFDER | Actinomycetia | Bacteria | Actinobacteria | | Actinomycetia | | | | | | | | | | | 2.3.1.-(100%) | Transferring groups other than amino-acyl groups (100%) | | | GO:0016746(94%);GO:0004315(6%) | acyltransferase activity (94%);3-oxoacyl-[acyl-carrier-protein] synthase activity (6%) | GO:0006633(6%);GO:0017000(6%) | fatty acid biosynthetic process (6%);antibiotic biosynthetic process (6%) | IPR000794(100%);IPR014030(100%);IPR01 | 1.00E+00 | 1.00E+00 | 1.30E+05 | NS | NS | NS | NS | NS | NS | NS | NS |

| | | | | | | | | | | | | | | | | | | | | | | | | | | | | | | | | | | | | | |
|---|---|---|---|---|---|---|---|---|---|---|---|---|---|---|---|---|---|---|---|---|---|---|---|---|---|---|---|---|---|---|---|---|---|---|---|---|---|
| | | | | | | | | | | | | | | | | | | | | | | | | | | 40031(100%) | | | | | | | | | | | |
| 202 | Variable 280 | AQASVDEDK | Bacillales | Bacteria | Firmicutes | | Bacilli | | Bacillales | | | | | | | | | | | GO:0048500(100%) | signal recognition particle (100%) | GO:0003924(100%);GO:0005525(100%);G | GTPase activity (100%);GTP binding (100%);7S RNA binding (98%) | GO:0006614(98%);GO:0006612(2%) | SRP-dependent cotranslational protein targeting to membrane (98% | IPR000897(100%);IPR003593(100%);IPR | 1.00E+00 | 1.00E+00 | 6.90E+04 | NS | NS | NS | NS | NS | NS | NS | NS |

| | | | | | | | | | | | | | | | | | | | | | | | | | | | | | | | | | | | | | |
|---|---|---|---|---|---|---|---|---|---|---|---|---|---|---|---|---|---|---|---|---|---|---|---|---|---|---|---|---|---|---|---|---|---|---|---|---|---|
| | | | | | | | | | | | | | | | | | | | | | | O:0008312(98%) | | | ); protein targeting to membrane (2%) | 004125(100%) | | | | | | | | | | | |
| 204 | Variable 282 | DTCAVVTR | Bacillales | Bacteria | Firmicutes | | Bacilli | | Bacillales | | | | | | | | | 3.4.13.9(100%) | Xaa-Prodipeptidase(100%) | | | GO:0102009(50%) | proline dipeptidase activity (50%) | GO:0006446(50%) | regulation of translational initiation (50%) | IPR001498(100%);IPR020568 | 1.00E+00 | 1.00E+00 | 1.06E+05 | 6.0572 | 0.0029188 | 0.000077381 | 0.9 | NS | NS | NS | NS |

| | | | | | | | | | | | | | | | | | | | | | | | | | | | | | | | | | | | | | |
|---|---|---|---|---|---|---|---|---|---|---|---|---|---|---|---|---|---|---|---|---|---|---|---|---|---|---|---|---|---|---|---|---|---|---|---|---|---|
| | | | | | | | | | | | | | | | | | | | | | | | | | | (100%);IPR023582(100%) | | | | | | | | | | | |
| 205 | Variable 283 | MADQGEVK | Bacillales | Bacteria | Firmicutes | | Bacilli | | Bacillales | | | | | | | | | 2.3.1.-(100%) | Transferring groups other than amino-acyl groups (100%) | | | GO:0031177(100%);GO:000 | phosphopantetheine binding(100%);catalytic activity(64%);3- | GO:0017000(100%);GO:0006633(27%) | antibiotic biosynthetic process(100%);fatty acid | IPR000873(100%);IPR0012 | 1.00E+00 | 1.00E+00 | 2.51E+04 | NS | NS | NS | NS | NS | NS | NS | NS |

| | | | | | | | | | | | | | | | | | | | | | | | | | | | | | | | | | | | | | | | | |
|---|---|---|---|---|---|---|---|---|---|---|---|---|---|---|---|---|---|---|---|---|---|---|---|---|---|---|---|---|---|---|---|---|---|---|---|---|---|---|---|---|
| | | | | | | | | | | | | | | | | | | | | | | 3824(64%);GO:0004315(27%) | oxoacyl-[acyl-carrier-protein] synthase activity(27%) | | biosynthetic process(27%) | 42(100%);IPR009081(100%) | | | | | | | | | | | | | | |
| 208 | Variable 286 | NDLQLQTALK | Bacillaceae | Bacteria | Firmicutes | | Bacilli | | Bacillales | | Bacillaceae | | | | | | | 3.4.21.102(100%) | C-terminal processing peptidase(100%) | GO:0030288(1%) | outer membrane-bounded periplasmic spac | GO:0008236(81%);GO:0 | serine-type peptidase activity(81%);serine-type | GO:0006508(1%);GO:0006518(1%);GO:00 | proteolysis(1%);peptide metabolic process | IPR002477(100%);IPR0 | 1.00E+00 | 1.00E+00 | 2.65E+04 | 6.304 | 3.4832E-09 | 0.00019596 | 0.9 | NS | NS | NS | NS | 26.795 | 3.16E-08 | 0.0055814 |

| | | | | | | | | | | | | | | | | | | | | | | | | | | | | | | | | | | | | | |
|---|---|---|---|---|---|---|---|---|---|---|---|---|---|---|---|---|---|---|---|---|---|---|---|---|---|---|---|---|---|---|---|---|---|---|---|---|---|
| | | | | | | | | | | | | | | | | | | | | | e(1%) | 004252(17%);GO:0008233(4%) | endopeptidase activity(17%);peptidase activity(4%) | 07165(1%) | s(1%);signal transduction(1%) | 36365(100%);IPR036366(100%) | | | | | | | | | | | |
| 209 | Variable 287 | EEQFTPEHMR | Bacillaceae | Bacteria | Firmicutes | | Bacilli | | Bacillales | | Bacillaceae | | | | | | | | | GO:0005737(100%);G | cytoplasm(100%);plasma membran | GO:0005524(100%);G | ATP binding(100%);metal ion binding(100%);h | GO:0006605(100%);GO:0017038(100 | protein targeting(100%);protein im | IPR000185(100%);IP | 1.00E+00 | 1.00E+00 | 3.19E+04 | 4.4218 | 0.0011088 | 0.000033333 | 0.9 | NS | NS | NS | NS |

| | | | | | | | | | | | | | | | | | | | | | | | | | | | | | | | | | | | | | |
|---|---|---|---|---|---|---|---|---|---|---|---|---|---|---|---|---|---|---|---|---|---|---|---|---|---|---|---|---|---|---|---|---|---|---|---|---|---|
| | | | | | | | | | | | | | | | | | | | | O:0005886(100%) | e (100%) | O:0046872(100%);GO:0004386(8%) | elicase activity (8%) | %);GO:0065002(83%) | port (100%);intracellular protein transmembrane transport (83%) | R004027(100%);IPR011116(100%) | | | | | | | | | | | |
| 210 | Variable 288 | NSAGGTGFEQVK | Bacillaceae | Bacteria | Firmicutes | | Bacilli | | Bacillales | | Bacillaceae | | | | | | | 4.3.2.1(100% | Argininosuccinate lyase (100%) | GO:0005737(1 | cytoplasm (100%) | GO:0004056(1 | argininosuccinate lyase activity (10 | GO:0042450(100%) | arginine biosynthetic proc | IPR000362(10 | 1.00E+00 | 1.00E+00 | 5.16E+05 | 16.624 | 0.0358 | 0.00029412 | 0.9 | NS | NS | NS | NS |

| | | | | | | | | | | | | | | | | | | | | | | | | | | | | | | | | | | | | | |
|---|---|---|---|---|---|---|---|---|---|---|---|---|---|---|---|---|---|---|---|---|---|---|---|---|---|---|---|---|---|---|---|---|---|---|---|---|---|
| | | | | | | | | | | | | | | | | | | ) | | 00%) | | 00%) | 0%) | | ess via ornithine (100%) | 0%);IPR008948(100%);IPR009049(100%) | | | | | | | | | | | |
| 211 | Variable 2 | LSEHLEELGFELVR | Bacillaceae | Bacteria | Firmicutes | | Bacilli | | Bacillales | | Bacillaceae | | | | | | | | | GO:0005737 | cytoplasm(99%) | GO:0050660 | flavin adenine dinucleotide bin | GO:0002098(95%);G | tRNA wobble uridine | IPR002218( | 1.00E+00 | 1.00E+00 | 3.75E+06 | NS | NS | NS | NS | NS | NS | NS | NS |

| | | | | | | | | | | | | | | | | | | | | | | | | | | | | | | | | | | | | | |
|---|---|---|---|---|---|---|---|---|---|---|---|---|---|---|---|---|---|---|---|---|---|---|---|---|---|---|---|---|---|---|---|---|---|---|---|---|---|
| | 89 | | | | | | | | | | | | | | | | | | | (99%) | | (100%) | ding(100%) | O:0008033(5%) | modification(95%);tRNA processing(5%) | 100%);IPR004416(100%);IPR020595(100%) | | | | | | | | | | | |
| 212 | Variable | DLQFELAAR | Actinomycet | Bacteria | Actinobac | | Actinomyceti | | | | | | | | | | | 3.1.25.-( | Site-specific endodeox | GO:0093 | excinucleasepa | GO:0036 | DNA binding(10 0% | GO:0006289(100 | nucleotide-excisi | IPR00194 | 1.00E+00 | 1.00E+00 | 1.11E+05 | NS | NS | NS | NS | NS | NS | NS | NS |

| | | | | | | | | | | | | | | | | | | | | | | | | | | | | | | | | | | | | | |
|---|---|---|---|---|---|---|---|---|---|---|---|---|---|---|---|---|---|---|---|---|---|---|---|---|---|---|---|---|---|---|---|---|---|---|---|---|---|
| | 29 | | ia | | teria | | a | | | | | | | | | | | 100%) | yribonucleases specific for altered bases (100%) | 80(99%);GO:0005737(97%);GO:0005886(0%) | ircomplex(99%);cytoplasm(97%);plasma membrane(0%) | 77(100%);GO:0005524(100%);GO:0016887(98%) | );ATP binding (100%);ATPase (98%) | %);GO:0009432(96%);GO:0051409(0%) | on repair (100%);SOS response (96%);response to nitrosative stress (0%) | 3(100%);IPR036876(100%);IPR004807(99%) | | | | | | | | | | | |
| 213 | Vari | LGGDAAFSG | Bacill | Bact | Firmi | | Bacilli | | Bacillal | | Bacillac | | | | | | | 2.7. | Fructokinase | | | GO:0 | kinase activit | alcohol | | | 1.00 | 4.50 | 1.00 | NS | NS | NS | NS | NS | NS | NS | NS |

| | | | | | | | | | | | | | | | | | | | | | | | | | | | | | | | | | | | | | | | | |
|---|---|---|---|---|---|---|---|---|---|---|---|---|---|---|---|---|---|---|---|---|---|---|---|---|---|---|---|---|---|---|---|---|---|---|---|---|---|---|---|---|
| | able 290 | K | aceae | eria | cutes | | | | es | | eae | | | | | | | 1.4(67%);2.7.1.-(33%) | (67%);Phosphotransferases with an alcohol group as acceptor (33%) | | | 0163001(97%);GO:0016773(88%);GO:0000865(3%) | y (97%);phosphotransferase activity | group as acceptor (88%);fructokinase activity (3%) | | | E+00 | E+05 | E+00 | | | | | | | | | | | |
| 214 | Var | CYTYTM | Alkal | Bac | Firm | | Bacill | | Bacill | | Bacill | Alk | | | Alk | | | 3.2 | Hydrolyz | | | GO: | DNA | GO:00 | base- | IPR | 1.0 | 1.0 | 4.2 | 2.7 | 0.0 | 0.00 | 0.9 | NS | NS | NS | NS | 5.4 | 0.0 | 0.0 |

| | | | | | | | | | | | | | | | | | | | | | | | | | | | | | | | | | | | | | | | | |
|---|---|---|---|---|---|---|---|---|---|---|---|---|---|---|---|---|---|---|---|---|---|---|---|---|---|---|---|---|---|---|---|---|---|---|---|---|---|---|---|---|
| | iable 291 | HTHCLLNVVCEQK | ihalobacillusclausii | teria | icutes | | i | | ales | | aceae | alihalobacillus | | | alihalobacillusclausii | | | .2.-(100%) | ingN-glycosylcompounds(100%) | | | 0003677(100%);GO:0003905(100%) | binding(100%);alkylbaseDNAN-glycosylaseactivity(100%) | 06284(100%) | excisionrepair(100%) | 003180(100%);IPR011034(100%);IPR036995(100%) | 0E+00 | 0E+00 | 9E+04 | 124 | 008803 | 0029412 | | | | | | 128 | 081004 | 0026738 |
| 2 | V | TA | B | B | F | | B | | B | | B | B | | | | | | 5 | Pe | G | cy | G | pe | G | pr | I | 1 | 1 | 3 | 0 | 1 | 0. | 0 | N | N | N | N | 1 | 1 | 0 |

| | | | | | | | | | | | | | | | | | | | | | | | | | | | | | | | | | | | | | | | | |
|---|---|---|---|---|---|---|---|---|---|---|---|---|---|---|---|---|---|---|---|---|---|---|---|---|---|---|---|---|---|---|---|---|---|---|---|---|---|---|---|---|
| 16 | ariable 293 | LDDAFK | acillus | acteria | irmicutes | | acilli | | acillales | | acillaceae | acillus | | | | | | .2.1.8(100%) | ptidylprolyl isomerase (100%) | O:0005737(95%) | toplasm(95%) | O:0003755(100%);GO:0043022(2%);GO:0044183(2%) | ptidyl-prolyl cis-trans isomerase activity (100%);ribosome binding (2%);protein folding chaperone (2%) | O:0015031(100%);GO:0006457(98%);GO:0007049(95%) | otein transport(100%); protein folding(98%);cell cycle(95%) | PR001179(100%);IPR005215(100%);IPR008880(100% | .00E+00 | .00E+00 | .69E+05 | .14952 | .4528E-12 | 0019305 | .9 | S | S | S | S | 8.505 | .7259E-06 | .0084231 |

| | | | | | | | | | | | | | | | | | | | | | | | | | | | | | | | | | | | | | |
|---|---|---|---|---|---|---|---|---|---|---|---|---|---|---|---|---|---|---|---|---|---|---|---|---|---|---|---|---|---|---|---|---|---|---|---|---|---|
| | | | | | | | | | | | | | | | | | | | | | | | | | ) | | | | | | | | | | | | |
| 218 | Variable 295 | EQPETASYR | Bacillus | Bacteria | Firmicutes | | Bacilli | | Bacillales | | Bacillaceae | Bacillus | | | | | | | | | | GO:0003677(100%);GO:0046983(100%);GO:0003700 | DNA binding(100%);protein dimerization activity(100%);DNA-binding transcription factor activity(4%) | GO:0006355(100%) | regulation of transcription | DNA-templated(100%) | 1.00E+00 | 1.00E+00 | 1.09E+05 | NS | NS | NS | NS | NS | NS | NS | NS |

| | | | | | | | | | | | | | | | | | | | | | | | | | | | | | | | | | | | | | |
|---|---|---|---|---|---|---|---|---|---|---|---|---|---|---|---|---|---|---|---|---|---|---|---|---|---|---|---|---|---|---|---|---|---|---|---|---|---|
| | | | | | | | | | | | | | | | | | | | | | | (4%) | | | | | | | | | | | | | | | |
| 219 | Variable 296 | HLMETGLSNK | Bacillus | Bacteria | Firmicutes | | Bacilli | | Bacillales | | Bacillaceae | Bacillus | | | | | | | | | | GO:0003677(100%) | DNA binding(100%) | GO:0006310(91%);GO:0015074(91%);GO:0044826(9%) | DNA recombination(91%);DNA integration(91%);viral genome integration into | IPR002104(100%);IPR004107(100%);IPR010998 | 1.00E+00 | 1.00E+00 | 4.91E+04 | NS | NS | NS | NS | NS | NS | NS | NS |

| | | | | | | | | | | | | | | | | | | | | | | | | | | | | | | | | | | | | | |
|---|---|---|---|---|---|---|---|---|---|---|---|---|---|---|---|---|---|---|---|---|---|---|---|---|---|---|---|---|---|---|---|---|---|---|---|---|---|
| | | | | | | | | | | | | | | | | | | | | | | | | | host DNA (9%) | (100%) | | | | | | | | | | | |
| 220 | Variable 297 | AAAGQVNGK | Bacillus | Bacteria | Firmicutes | | Bacilli | | Bacillales | | Bacillaceae | Bacillus | | | | | | 6.1.1.3(100%) | Threonine--tRNA ligase (100%) | GO:0005737(97%);GO:0005829(3%) | cytoplasm (97%);cytosol (3%) | GO:0004829(100%);GO:0005524(100%);GO: | threonine-tRNA ligase activity (100%);ATP binding (100%);tRNA binding (97%) | GO:0006435 (100%) | threonyl-tRNA aminoacylation (100%) | IPR002314(100%);IPR002320(100%);IPR00 | 1.00E+00 | 1.00E+00 | 5.53E+04 | NS | NS | NS | NS | NS | NS | NS | NS |

| | | | | | | | | | | | | | | | | | | | | | | | | | | | | | | | | | | | | | |
|---|---|---|---|---|---|---|---|---|---|---|---|---|---|---|---|---|---|---|---|---|---|---|---|---|---|---|---|---|---|---|---|---|---|---|---|---|---|
| | | | | | | | | | | | | | | | | | | | | | | 000000049(97%) | | | | 4095(100%) | | | | | | | | | | | |
| 221 | Variable 298 | TALNNGLTYLK | Bacillus | Bacteria | Firmicutes | | Bacilli | | Bacillales | | Bacillaceae | Bacillus | | | | | | 3.2.1.96(100%) | Mannosyl-glycoprotein endo-beta-N-acetylglucosaminidase (100%) | GO:0005576(93%);GO:0005618(3%) | extracellular region (93%);cell wall (3%) | GO:0004040(100%);GO:0042834(97%); | amidase activity (100%);peptidoglycan binding (97%);mannosyl-glycoprotein endo-beta- | GO:0008152(10%);GO:0071555(3%) | metabolic process (10%);cell wall organization (3%) | IPR002901(100%);IPR036446(100%);I | 1.00E+00 | 1.00E+00 | 4.46E+04 | 2.7225 | 0.001461 | 0.00032722 | 0.9 | NS | NS | NS | NS |

| | | | | | | | | | | | | | | | | | | | | | | | | | | | | | | | | | | | | | |
|---|---|---|---|---|---|---|---|---|---|---|---|---|---|---|---|---|---|---|---|---|---|---|---|---|---|---|---|---|---|---|---|---|---|---|---|---|---|
| | | | | | | | | | | | | | | | | | | | | | | GO:0033925(14%) | N-acetylglucosaminidase activity (14%) | | | PR007730(97%) | | | | | | | | | | | |
| 222 | Variable 299 | MSNGLDVYVLPK | Bacillus | Bacteria | Firmicutes | | Bacilli | | Bacillales | | Bacillaceae | Bacillus | | | | | | 3.4.24.-(100%) | Metalloendopeptidases (100%) | | | GO:0046872(100%);GO:0008233(1 | metal ion binding (100%);peptidase activity (19%);metallopeptidase activity (4 | | | IPR007863(100%);IPR011249(100 | 1.00E+00 | 1.00E+00 | 6.28E+04 | NS | NS | NS | NS | NS | NS | NS | NS |

| | | | | | | | | | | | | | | | | | | | | | | | | | | | | | | | | | | | | | | | | |
|---|---|---|---|---|---|---|---|---|---|---|---|---|---|---|---|---|---|---|---|---|---|---|---|---|---|---|---|---|---|---|---|---|---|---|---|---|---|---|---|---|
| | | | | | | | | | | | | | | | | | | | | | | 9%);GO:0008237(4%) | %) | | | %);IPR011765(100%) | | | | | | | | | | | | | | |
| 223 | Variable 3 | VTPGVLFFAFPGSK | CandidatusSolibacterusitatus | Bacteria | Acidobacteria | | Acidobacteriia | | Bryobacterales | | Solibacteraceae | CandidatusSolibacter | | | CandidatusSolibacterusitatus | | | 6.3.2.13(100%) | UDP-N-acetylmuramoyl-L-alanyl-D-glutamate--2 | 6-diaminopimelate ligase(100%) | GO:0005737(100%) | cytoplasm (100%) | GO:0000287(100%);GO:0005524(100%);GO:0008765(100%) | magnesium ion binding (100%);ATP binding (100%);UDP-N-acetylm | 6-diaminopimelate ligase activity (100%) | GO:0070049(100%);GO:0008360(1 | 1.00E+00 | 1.00E+00 | 5.26E+04 | 5.8172 | 0.00010409 | 0.00026283 | 0.9 | NS | NS | NS | NS | 7.7376 | 0.0013777 | 0.0014836 |

| | | | | | | | | | | | | | | | | | | | | | | | | | | | | | | | | | | | | | |
|---|---|---|---|---|---|---|---|---|---|---|---|---|---|---|---|---|---|---|---|---|---|---|---|---|---|---|---|---|---|---|---|---|---|---|---|---|---|
| | | | | | | | | | | | | | | | | | | | | | | | | uramoylalanyl-D-glutamate-2 | | 00%);GO:0009252(100%) | | | | | | | | | | | |
| 224 | Variable 30 | AVVTYCTGGVR | Actinomycetia | Bacteria | Actinobacteria | | Actinomycetia | | | | | | | | | | | 1.14.-.-(100%) | Acting on paired donors | with incorporation or reduction | | | GO:0016705(100%);GO:0016740(25%) | oxidoreductase activity | acting on paired donors | with incorporation or reduction of | 1.00E+00 | 1.00E+00 | 3.37E+04 | NS | NS | NS | NS | NS | NS | NS | NS |

| | | | | | | | | | | | | | | | | | | | | | | | | | | |
|---|---|---|---|---|---|---|---|---|---|---|---|---|---|---|---|---|---|---|---|---|---|---|---|---|---|---|
| | | | | | | | | | | | | | | | | | | | | of molecular oxygen. The oxygen incorporated need not be | | | | | | molecular oxygen (100%); transferase activity (25%) |

| | | | | | | | | | | | | | | | | | | | | | | | | | | | | | | | | | | | | | |
|---|---|---|---|---|---|---|---|---|---|---|---|---|---|---|---|---|---|---|---|---|---|---|---|---|---|---|---|---|---|---|---|---|---|---|---|---|---|
| | | | | | | | | | | | | | | | | | | | | derived from O(2)(100%) | | | | | | | | | | | | | | | | | |
| 225 | Variable 300 | YLESQHTCK | Bacillus | Bacteria | Firmicutes | | Bacilli | | Bacillales | | Bacillaceae | Bacillus | | | | | | 2.3.1.-(100%) | Transferring groups other than amino-acyl groups(100%) | GO:0005737(7%) | cytoplasm(7%) | GO:0016746(73%);GO:0016747 | acyltransferase activity(73%);transferase activity(27%) | GO:0017000(7%) | antibiotic biosynthetic process(7%) | IPR001227(100%);IPR014004 | 1.00E+00 | 1.00E+00 | 5.01E+04 | NS | NS | NS | NS | NS | NS | NS | NS |

| | | | | | | | | | | | | | | | | | | | | | | | | | | | | | | | | | | | | | |
|---|---|---|---|---|---|---|---|---|---|---|---|---|---|---|---|---|---|---|---|---|---|---|---|---|---|---|---|---|---|---|---|---|---|---|---|---|---|
| | | | | | | | | | | | | | | | | | | | | | | 40(27%) | | | | 3(100%);IPR016035(100%) | | | | | | | | | | | |
| 226 | Variable 301 | YPACMLFVPSR | Bacillus | Bacteria | Firmicutes | | Bacilli | | Bacillales | | Bacillaceae | Bacillus | | | | | | 3.5.3.9(100%) | Allantoate deiminase (100%) | GO:0005737(4%) | cytoplasm(4%) | GO:0046872(88%);GO:004 | metal ion binding (88%);allantoate deiminase activity (54%) | acting on carbon-nitrogen (but not peptide) bonds | in linear amidines (46%) | GO:0000256(4%);GO:000614 | 1.00E+00 | 1.00E+00 | 2.67E+04 | NS | NS | NS | NS | NS | NS | NS | NS |

| | | | | | | | | | | | | | | | | | | | | | | | | | | | | | | | | | | | | | |
|---|---|---|---|---|---|---|---|---|---|---|---|---|---|---|---|---|---|---|---|---|---|---|---|---|---|---|---|---|---|---|---|---|---|---|---|---|---|
| | | | | | | | | | | | | | | | | | | | | | | 7652(54%);GO:0016813(46%) | ;hydrolase activity | | | 4(4%) | | | | | | | | | | | |
| 227 | Variable 302 | VHTVFHYFLT | Bacillus | Bacteria | Firmicutes | | Bacilli | | Bacillales | | Bacillaceae | Bacillus | | | | | | 2.3.1.-(100%) | Transferring groups other than amino-acyl groups (100%) | | | GO:0008080(77%);GO:0 | N-acetyltransferase activity (77%);transferase activity | GO:0006048 (9%) | UDP-N-acetylglucosamine biosynthetic | IPR000182(100%);IPR0 | 1.00E+00 | 1.00E+00 | 1.58E+05 | 7.7529 | 0.0041522 | 0.00015754 | 0.9 | NS | NS | NS | NS |

| | | | | | | | | | | | | | | | | | | | | | | | | | | | | | | | | | | | | | |
|---|---|---|---|---|---|---|---|---|---|---|---|---|---|---|---|---|---|---|---|---|---|---|---|---|---|---|---|---|---|---|---|---|---|---|---|---|---|
| | | | | | | | | | | | | | | | | | | | ) | | | 0167400(14%);GO:0004343(9%) | (14%);glucosamine 6-phosphate N-acetyltransferase activity (9%) | | process (9%) | 16181(100%);IPR039143(9%) | | | | | | | | | | | |
| 228 | Variable 303 | GAFTLTDGK | Bacillus | Bacteria | Firmicutes | | Bacilli | | Bacillales | | Bacillaceae | Bacillus | | | | | | | | GO:0016021(100%) | integral component of membrane (100% | | | | | IPR020534(100%) | 1.00E+00 | 1.00E+00 | 4.89E+04 | 3.2058 | 0.0090238 | 0.0001333 | 0.9 | NS | NS | NS | NS |

| | | | | | | | | | | | | | | | | | | | | | | | | | | | | | | | | | | | | | | | | |
|---|---|---|---|---|---|---|---|---|---|---|---|---|---|---|---|---|---|---|---|---|---|---|---|---|---|---|---|---|---|---|---|---|---|---|---|---|---|---|---|---|
| | | | | | | | | | | | | | | | | | | | | | ) | | | | | | | | | | | | | | | | | | | |
| 229 | Variable 304 | MADLGGSEK | Bacillus | Bacteria | Firmicutes | | Bacilli | | Bacillales | | Bacillaceae | Bacillus | | | | | | | | | | GO:0005524(100%) | ATP binding (100%) | | | IPR003439(100%);IPR003593(100%);IPR027417(10 | 1.00E+00 | 1.00E+00 | 3.66E+04 | 4.7036 | 9.0757E-07 | 0.00028194 | 0.9 | NS | NS | NS | NS | 11.394 | 0.001136 | 0.0019521 |

| | | | | | | | | | | | | | | | | | | | | | | | | | | | | | | | | | | | | | | | | |
|---|---|---|---|---|---|---|---|---|---|---|---|---|---|---|---|---|---|---|---|---|---|---|---|---|---|---|---|---|---|---|---|---|---|---|---|---|---|---|---|---|
| | | | | | | | | | | | | | | | | | | | | | | | | | | 0%) | | | | | | | | | | | | | | |
| 230 | Variable 305 | TWYEAAEK | Bacillus | Bacteria | Firmicutes | | Bacilli | | Bacillales | | Bacillaceae | Bacillus | | | | | | 7.6.2.-(100%) | Linked to the hydrolysis of a nucleoside triphosphate (100%) | GO:0016021(96%);GO:0005886(4%) | integral component of membrane(96%);plasma membrane(4%) | GO:0005524(100%);GO:0016887(96%);GO:004262 | ATP binding(100%);ATPase(96%);ATPase-coupled transmembrane transporter activity(96%) | GO:0046677(4%);GO:0055085(4%) | response to antibiotic(4%);transmembrane transport(4%) | IPR003439(100%);IPR003593(100%);IPR011527( | 1.00E+00 | 1.00E+00 | 3.09E+04 | 0.27697 | 2.0824E-11 | 0.0012357 | 0.9 | NS | NS | NS | NS | 31.735 | 4.005E-09 | 0.0021027 |

| | | | | | | | | | | | | | | | | | | | | | | | | | | | | | | | | | | | | | | | | |
|---|---|---|---|---|---|---|---|---|---|---|---|---|---|---|---|---|---|---|---|---|---|---|---|---|---|---|---|---|---|---|---|---|---|---|---|---|---|---|---|---|
| | | | | | | | | | | | | | | | | | | | | | | 6(96%) | | | | 100%) | | | | | | | | | | | | | | |
| 232 | Variable 307 | DNGYMNK | Bacillus | Bacteria | Firmicutes | | Bacilli | | Bacillales | | Bacillaceae | Bacillus | | | | | | | | GO:0005886(100%) | plasmamembrane (100%) | | | GO:0008643 (100%) | carbohydrate transport (100%) | IPR006059(100%) | 1.00E+00 | 1.00E+00 | 4.66E+04 | 0.6513 | 1.2465E-07 | 0.00078159 | 0.9 | NS | NS | NS | NS | 8.8388 | 0.00062546 | 0.00099091 |
| 233 | Variable 308 | QQYEQLVEELK | Bacillus | Bacteria | Firmicutes | | Bacilli | | Bacillales | | Bacillaceae | Bacillus | | | | | | 2.7.13.3(100%);3.1.3.-( | Histidine kinase (100%);Phosphoric monoester hydrolases (100% | GO:0005737(100%);GO:00 | cytoplasm (100%);integral component of mem | GO:0000155(100%);GO:00 | phosphorelay sensor kinase activity (100%);phosphoprotein ph | | | IPR003594(100%);IPR008 | 1.00E+00 | 1.00E+00 | 1.17E+05 | NS | NS | NS | NS | NS | NS | NS | NS | | | |

| | | | | | | | | | | | | | | | | | | | | | | | | | | | | | | | | | | | | | |
|---|---|---|---|---|---|---|---|---|---|---|---|---|---|---|---|---|---|---|---|---|---|---|---|---|---|---|---|---|---|---|---|---|---|---|---|---|---|
| | | | | | | | | | | | | | | | | | | 100%) | ) | 16021(96%) | brane(96%) | 04721(100%);GO:0005524(100%) | osphatase activity(100%);ATP binding(100%) | | | 595(100%);IPR011712(100%) | | | | | | | | | | | |
| 234 | Variable 309 | ENLQLADHMAK | Bacillus | Bacteria | Firmicutes | | Bacilli | | Bacillales | | Bacillaceae | Bacillus | | | | | | | | GO:0005737(3%) | cytoplasm(3%) | GO:0003677(3%);GO | DNA binding(3%);DNA/DNA annea | GO:0009294(97%);GO:0030420(3 | DNA mediated transformation(97 | IPR003488(100%) | 1.00E+00 | 1.00E+00 | 8.02E+04 | NS | NS | NS | NS | NS | NS | NS | NS |

| | | | | | | | | | | | | | | | | | | | | | | | | | | | | | | | | | | | | | |
|---|---|---|---|---|---|---|---|---|---|---|---|---|---|---|---|---|---|---|---|---|---|---|---|---|---|---|---|---|---|---|---|---|---|---|---|---|---|
| | | | | | | | | | | | | | | | | | | | | | | :1990814(3%) | ling activity (3%) | %) | %);establishment of competence for transformation (3%) | | | | | | | | | | | | |
| 235 | Variable 31 | EVLLTVLDR | Actinomycetia | Bacteria | Actinobacteria | | Actinomycetia | | | | | | | | | | | | | GO:0005886(100% | plasmamembrane(100%);cyt | GO:0005524(100% | ATP binding(100%);helicase activity(22 | GO:0006605(100%);GO:0017038 | protein targeting(100%); protei | IPR000185(100%); | 1.00E+00 | 1.00E+00 | 3.75E+04 | NS | NS | NS | NS | NS | NS | NS | NS |

| | | | | | | | | | | | | | | | | | | | | | | | | | | | | | | | | | | | | | |
|---|---|---|---|---|---|---|---|---|---|---|---|---|---|---|---|---|---|---|---|---|---|---|---|---|---|---|---|---|---|---|---|---|---|---|---|---|---|
| | | | | | | | | | | | | | | | | | | | | );GO:0005737(78%) | oplasm(78%) | );GO:0004386(22%) | %) | (100%);GO:0065002(78%) | n import(100%);intracellular protein transmembrane transport(78%) | IPR011116(100%);IPR014018(100%) | | | | | | | | | | | |
| 236 | Variable 3 | EANSTEGK | Bacillus | Bacteria | Firmicutes | | Bacilli | | Bacillales | | Bacillaceae | Bacillus | | | | | | | | GO:0005576 | extracellular region(100 | GO:0090729 | toxin activity(100%) | GO:0001906(100%);G | cell killing(100%);ob | IPR023888( | 1.00E+00 | 1.00E+00 | 7.29E+04 | NS | NS | NS | NS | NS | NS | NS | NS |

| | | | | | | | | | | | | | | | | | | | | | | | | | | | | | | | | | | | | | |
|---|---|---|---|---|---|---|---|---|---|---|---|---|---|---|---|---|---|---|---|---|---|---|---|---|---|---|---|---|---|---|---|---|---|---|---|---|---|
| | 10 | | | | | | | | | | | | | | | | | | | (100%) | %) | (100%) | | O:0009405(100%);GO:0019835(100%) | solete pathogenesis(100%);cytolysis(100%) | 100%) | | | | | | | | | | | |
| 237 | Variable 311 | AVVEVTCVADR | Bacillus | Bacteria | Firmicutes | | Bacilli | | Bacillales | | Bacillaceae | Bacillus | | | | | | 3.2.1.22(86%);3.2.1.-(14 | Alpha-galactosidase(86%);Glycosidases | i.e. enzymes hydrolyzing O- | GO:0005737(3%) | cytoplasm (3%) | GO:0016616(100%);GO:0046872(100%);GO:0004553 | oxidoreductase activity | acting on the CH-OH group of donors | NAD or NADP as acceptor (10 | 1.00E+00 | 1.00E+00 | 1.00E+05 | NS | NS | NS | NS | NS | NS | NS | NS |

| | | | | | | | | | | | | | | | | | | | | | | | | | | | | | | | | | | | | | |
|---|---|---|---|---|---|---|---|---|---|---|---|---|---|---|---|---|---|---|---|---|---|---|---|---|---|---|---|---|---|---|---|---|---|---|---|---|---|
| | | | | | | | | | | | | | | | | | | %) | | andS-glycosylcompounds(14%) | | | (83%) | | | 0%);metalionbinding(100%);hydrolaseactivity | | | | | | | | | | | |
| 238 | Variable 31 | WPHAAPEGK | Bacillus | Bacteria | Firmicutes | | Bacilli | | Bacillales | | Bacillaceae | Bacillus | | | | | | 1.3.3.15(10 | Coproporphyrinogen III oxidase | GO:0005737( | cytoplasm(100%);integ | GO:0004729( | oxygen-dependent protoporphyr | GO:0006783(100%) | heme biosynthetic proc | IPR002937(1 | 1.00E+00 | 1.00E+00 | 5.89E+04 | 5.3486 | 0.0669 | 0.0111 | 0.9 | NS | NS | NS | NS |

| | | | | | | | | | | | | | | | | | | | | | | | | | | | | | | | | | | | | | |
|---|---|---|---|---|---|---|---|---|---|---|---|---|---|---|---|---|---|---|---|---|---|---|---|---|---|---|---|---|---|---|---|---|---|---|---|---|---|
| | 2 | | | | | | | | | | | | | | | | | 0%) | (coproporphyrin-forming)(100%) | 100%);GO:0016021(78%);GO:0005886(2%) | ral component of membrane(78%);plasma membrane(2%) | 100%);GO:0016491(2%) | inogen oxidase activity(100%);oxidoreductase activity(2%) | | es(100%) | 00%);IPR004572(100%);IPR036188(100%) | | | | | | | | | | | |
| 239 | Variable | KPQNGSLTLK | Bacillus | Bacteria | Firmicutes | | Bacilli | | Bacillales | | Bacillaceae | Bacillus | | | | | | | | GO:0005888 | plasmamembran | GO:0003677 | DNA binding(50%);A | GO:0030420(50%) | establishment of co | IPR001482 | 1.00E+00 | 1.00E+00 | 5.43E+04 | NS | NS | NS | NS | NS | NS | NS | NS |

| | | | | | | | | | | | | | | | | | | | | | | | | | | | | | | | | | | | | | |
|---|---|---|---|---|---|---|---|---|---|---|---|---|---|---|---|---|---|---|---|---|---|---|---|---|---|---|---|---|---|---|---|---|---|---|---|---|---|
| | 313 | | | | | | | | | | | | | | | | | | | 6(50%) | e(50%) | 7(50%);GO:0005524(50%) | TP binding(50%) | | mpetence for transformation(50%) | (100%);IPR003593(100%);IPR027417(100%) | | | | | | | | | | | |
| 240 | Variabl | NASAGGSLDAER | Bacillus | Bacteri | Firmicut | | Bacilli | | Bacillales | | Bacillaceae | Bacillus | | | | | | | | | | GO:0031 | phosphopantetheine | GO:0017000(8 | antibiotic biosyn | IPR0008 | 1.00E+0 | 1.00E+0 | 7.71E+0 | 4.5361 | 0.0047 | 0.000238 | 0.9 | NS | NS | NS | NS |

| | | | | | | | | | | | | | | | | | | | | | | | | | | | | | | | | | | | | | | | | |
|---|---|---|---|---|---|---|---|---|---|---|---|---|---|---|---|---|---|---|---|---|---|---|---|---|---|---|---|---|---|---|---|---|---|---|---|---|---|---|---|---|
| | e 314 | | | a | es | | | | | | | | | | | | | | | | | 177(100%);GO:0003824(91%);GO:0016874(9%) | binding(100%);catalytic activity(91%);ligase activity(9%) | 3%);GO:0030435(4%) | thetic process(83%);sporulation resulting in formation of a cellular spore(4%) | 73(100%);IPR001242(100%);IPR006162(100%) | 0 | 0 | 4 | | 49 | 64 | | | | | | | | |
| 241 | Varia | HQPAADLK | Bacillu | Bacte | Firmic | | Bacilli | | Bacillales | | Bacillacea | Bacillu | | | | | | | | | | GO:00 | ATP binding | GO:000635 | regulation | DNA- | 1.00E | 1.00E | 4.51E | 7.016 | 5.409 | 0.0003 | 0.9 | NS | NS | NS | NS | 11.94 | 0.000 | 0.001 |

| | | | | | | | | | | | | | | | | | | | | | | | | | | | | | | | | | | | | | | | | |
|---|---|---|---|---|---|---|---|---|---|---|---|---|---|---|---|---|---|---|---|---|---|---|---|---|---|---|---|---|---|---|---|---|---|---|---|---|---|---|---|---|
| | ble 315 | | s | ria | utes | | | | | | e | s | | | | | | | | | | 05524(100%);GO:0008134(100%);GO:0043565(100%) | (100%);transcription factor binding(100%);sequence-specific DNA binding(100%) | 5(100%);GO:0000160(3%);GO:0006525(3%) | of transcription | templated(100%);phosphorelay signal transduction system(3% | +00 | +00 | +04 | 9 | 3E-07 | 5915 | | | | | | 1 | 078386 | 3212 |

| | | | | | | | | | | | | | | | | | | | | | | | | | | | | | | | | | | | | | |
|---|---|---|---|---|---|---|---|---|---|---|---|---|---|---|---|---|---|---|---|---|---|---|---|---|---|---|---|---|---|---|---|---|---|---|---|---|---|
| | | | | | | | | | | | | | | | | | | | | | | | | | | ); arginine metabolic process (3%) | | | | | | | | | | | |
| 242 | Variable 316 | SLEMAVGLLAVLK | Bacillus | Bacteria | Firmicutes | | Bacilli | | Bacillales | | Bacillaceae | Bacillus | | | | | | | | | | GO:0031177(100%);GO:00 | phosphopantetheine binding (100%);catalytic activity (97%) | GO:0009058 (100%) | biosynthetic process (100%) | IPR000873(100%);IPR042 | 1.00E+00 | 1.00E+00 | 1.43E+05 | 2.0687 | 0.00060329 | 0.00036552 | 0.9 | NS | NS | NS | NS |

| | | | | | | | | | | | | | | | | | | | | | | | | | | | | | | | | | | | | | |
|---|---|---|---|---|---|---|---|---|---|---|---|---|---|---|---|---|---|---|---|---|---|---|---|---|---|---|---|---|---|---|---|---|---|---|---|---|---|
| | | | | | | | | | | | | | | | | | | | | | | 03824(97%);GO:0016874(3%) | ;ligase activity (3%) | | | 099(100%);IPR001031(97%) | | | | | | | | | | | |
| 243 | Variable 317 | VTGSGNNGR | Bacillus | Bacteria | Firmicutes | | Bacilli | | Bacillales | | Bacillaceae | Bacillus | | | | | | 2.3.1.-(98%);2.3.1.12 | Transferring groups other than amino-acyl groups (98%); | | | GO:0016746(95%);GO:0 | acyltransferase activity (95%);dihydrolipoyllysine-res | GO:0006096(2%) | glycolytic process (2%) | IPR000089(100%);IPR0 | 1.00E+00 | 1.63E+05 | 1.00E+00 | NS | NS | NS | NS | NS | NS | NS | NS |

| | | | | | | | | | | | | | | | | | | | | | | | | | | | | | | | | | | | | | |
|---|---|---|---|---|---|---|---|---|---|---|---|---|---|---|---|---|---|---|---|---|---|---|---|---|---|---|---|---|---|---|---|---|---|---|---|---|---|
| | | | | | | | | | | | | | | | | | | (2%) | Dihydrolipoyllysine-residue acetyltransferase (2%) | | | 004742(5%) | idue acetyltransferase activity (5%) | | | 010078(100%); IPR004167(100%) | | | | | | | | | | | |
| 244 | Variable 318 | QDGDDLSFPR | Bacillus | Bacteria | Firmicutes | | Bacilli | | Bacillales | | Bacillaceae | Bacillus | | | | | | 2.4.-.-(100%) | Glycosyltransferases (100%) | | | GO:0016740(97%);GO | transferase activity (97%);glycosyltransferase act | GO:0000271 (3%) | polysaccharide biosynthetic process | IPR001173(100%);IP | 1.00E+00 | 2.75E+05 | 1.00E+00 | NS | NS | NS | NS | NS | NS | NS | NS |

| | | | | | | | | | | | | | | | | | | | | | | | | | | | | | | | | | | | | | |
|---|---|---|---|---|---|---|---|---|---|---|---|---|---|---|---|---|---|---|---|---|---|---|---|---|---|---|---|---|---|---|---|---|---|---|---|---|---|
| | | | | | | | | | | | | | | | | | | | | | | :0016757(3%) | ivity (3%) | | (3%) | R029044(100%) | | | | | | | | | | | |
| 245 | Variable 319 | VEPSNTFSTESQASQAVAK | Bacillus | Bacteria | Firmicutes | | Bacilli | | Bacillales | | Bacillaceae | Bacillus | | | | | | 3.2.1.96(100%) | Mannosyl-glycoprotein endo-beta-N-acetylglucosaminidase (100%) | GO:0005576(89%);GO:0005618(5%) | extracellular region (89%);cell wall (5%) | GO:0004040(100%);GO:0042834(100 | amidase activity (100%);peptidoglycan binding (100%);mannosyl-glycoprotein endo-bet | GO:0008152(5%);GO:0071555(5%) | metabolic process (5%);cell wall organization (5%) | IPR002901(100%);IPR003646(100%); | 1.00E+00 | 6.14E+05 | 1.00E+00 | NS | NS | NS | NS | NS | NS | NS | NS |

| | | | | | | | | | | | | | | | | | | | | | | | | | | | | | | | | | | | | | |
|---|---|---|---|---|---|---|---|---|---|---|---|---|---|---|---|---|---|---|---|---|---|---|---|---|---|---|---|---|---|---|---|---|---|---|---|---|---|
| | | | | | | | | | | | | | | | | | | | | | | %);GO:0033925(11%) | a-N-acetylglucosaminidase activity(11%) | | | IPR007730(100%) | | | | | | | | | | | |
| 246 | Variable 32 | GSTASAPGER | Actinomycetia | Bacteria | Actinobacteria | | Actinomycetia | | | | | | | | | | | | | | | GO:0003677(99%);GO:0000976( | DNA binding(99%);transcription cis-regulatory region binding(1%);D | GO:0006355(99%) | regulation of transcription | DNA-templated(99%) | 1.00E+00 | 1.00E+00 | 2.36E+04 | NS | NS | NS | NS | NS | NS | NS | NS |

| | | | | | | | | | | | | | | | | | | | | | | | | | | | | | | | | | | | | | |
|---|---|---|---|---|---|---|---|---|---|---|---|---|---|---|---|---|---|---|---|---|---|---|---|---|---|---|---|---|---|---|---|---|---|---|---|---|---|
| | | | | | | | | | | | | | | | | | | | | | | 1%);GO:0003700(1%) | NA-binding transcription factor activity (1%) | | | | | | | | | | | | | | |
| 247 | Variable 320 | ECWASFFSGR | Bacillus | Bacteria | Firmicutes | | Bacilli | | Bacillales | | Bacillaceae | Bacillus | | | | | | 2.7.9.2(100%) | Pyruvate | water dikinase(100%) | | | GO:0005524(95%);GO:0016301(92%);GO:0016740(5%) | ATP binding (95%);kinase activity (92%);transferase activity (5 | | | 1.00E+00 | 2.02E+05 | 1.00E+00 | NS | NS | NS | NS | NS | NS | NS | NS |

| | | | | | | | | | | | | | | | | | | | | | | | | | | | | | | | | | | | | | |
|---|---|---|---|---|---|---|---|---|---|---|---|---|---|---|---|---|---|---|---|---|---|---|---|---|---|---|---|---|---|---|---|---|---|---|---|---|---|
| | | | | | | | | | | | | | | | | | | | | | | | | %) | | | | | | | | | | | | | |
| 248 | Variable 321 | QVFLTGHPEYDTNTLLQEYER | Bacillus | Bacteria | Firmicutes | | Bacilli | | Bacillales | | Bacillaceae | Bacillus | | | | | | 2.3.1.31(97%);2.3.1.-(3%) | Homoserine O-acetyltransferase (97%);Transferring groups other than amino-acyl groups (3%) | GO:0005737(100%) | cytoplasm(100%) | GO:0008899(100%);GO:0004414(89%) | homoserine O-succinyltransferase activity (100%);homoserine O-acetyltransferase activity (89%) | GO:0019281(86%);GO:0009086(14%) | L-methionine biosynthetic process from homoserine via O-succinyl-L-homoserine and cy | IPR029062(100%);IPR033752(100%);IPR005697(97 | 1.00E+00 | 4.26E+04 | 1.00E+00 | NS | NS | NS | NS | NS | NS | NS | NS |

| | | | | | | | | | | | | | | | | | | | | | | | | | | | | | | | | | | | | | |
|---|---|---|---|---|---|---|---|---|---|---|---|---|---|---|---|---|---|---|---|---|---|---|---|---|---|---|---|---|---|---|---|---|---|---|---|---|---|
| | | | | | | | | | | | | | | | | | | | | | | | | | stathionine(86%); methionine biosynthetic process(14%) | %) | | | | | | | | | | | |
| 249 | Variable 322 | QLNYQNPYQPMTSDGVYYDK | Bacillus | Bacteria | Firmicutes | | Bacilli | | Bacillales | | Bacillaceae | Bacillus | | | | | | | | GO:0005829(10 | cytosol(100%) | | | GO:0044781(100%) | bacterial-type flagellum organi | | 1.00E+00 | 7.58E+04 | 1.00E+00 | NS | NS | NS | NS | NS | NS | NS | NS |

| | | | | | | | | | | | | | | | | | | | | | | | | | | | | | | | | | | | | | |
|---|---|---|---|---|---|---|---|---|---|---|---|---|---|---|---|---|---|---|---|---|---|---|---|---|---|---|---|---|---|---|---|---|---|---|---|---|---|
| | | | | | | | | | | | | | | | | | | | | 0%) | | | | | zation (100%) | | | | | | | | | | | | |
| 250 | Variable 323 | LAVVAVNFTSAEK | Bacillus | Bacteria | Firmicutes | | Bacilli | | Bacillales | | Bacillaceae | Bacillus | | | | | | | | GO:0016021(86%);GO:0005886(5%) | integral component of membrane (86%);plasma membrane (5%) | GO:0016209(100%);GO:0016491(100%) | antioxidant activity (100%);oxidoreductase activity (100%) | | | IPR000866(100%);IPR013766(100%);IPR036 | 1.00E+00 | 1.65E+05 | 1.00E+00 | NS | NS | NS | NS | NS | NS | NS | NS |

| | | | | | | | | | | | | | | | | | | | | | | | | | | | | | | | | | | | | | |
|---|---|---|---|---|---|---|---|---|---|---|---|---|---|---|---|---|---|---|---|---|---|---|---|---|---|---|---|---|---|---|---|---|---|---|---|---|---|
| | | | | | | | | | | | | | | | | | | | | | | | | | | 249(100%) | | | | | | | | | | | |
| 252 | Variable 325 | LTEHMDDPADVQTR | Bacillus | Bacteria | Firmicutes | | Bacilli | | Bacillales | | Bacillaceae | Bacillus | | | | | | | | GO:0016021(100%);GO:0005886(4%) | integral component of membrane (100%);plasma membrane (4%) | | | GO:0009405(100%);GO:0030435(4%) | obsolete pathogenesis (100%);sporulation resulting in formation of a cellular | IPR025940(100%) | 1.00E+00 | 1.37E+06 | 1.00E+00 | NS | NS | NS | NS | NS | NS | NS | NS |

| | | | | | | | | | | | | | | | | | | | | | | | | | | | | | | | | | | | | | |
|---|---|---|---|---|---|---|---|---|---|---|---|---|---|---|---|---|---|---|---|---|---|---|---|---|---|---|---|---|---|---|---|---|---|---|---|---|---|
| | | | | | | | | | | | | | | | | | | | | | | | | | spore(4%) | | | | | | | | | | | | |
| 254 | Variable 327 | TFFGAGSK | Bacillus | Bacteria | Firmicutes | | Bacilli | | Bacillales | | Bacillaceae | Bacillus | | | | | | 3.6.3.34(100%) | Transferred entry: 7.2.2.16(100%) | GO:0005886(97%);GO:0016021(97%);GO:0005 | plasma membrane(97%);integral component of membrane(97%);integral co | GO:0022857(100%);GO:0016787(3%) | transmembrane transporter activity(100%);hydrolase activity(3%) | GO:0033214(3%) | siderophore-dependent iron import into cell(3%) | IPR000522(100%);IPR037294(100%) | 1.00E+00 | 4.30E+04 | 1.00E+00 | NS | NS | NS | NS | NS | NS | NS | NS |

| | | | | | | | | | | | | | | | | | | | | | | | | | | | | | | | | | | | | | |
|---|---|---|---|---|---|---|---|---|---|---|---|---|---|---|---|---|---|---|---|---|---|---|---|---|---|---|---|---|---|---|---|---|---|---|---|---|---|
| | | | | | | | | | | | | | | | | | | | | 887(3%) | mponent of plasma membrane (3%) | | | | | | | | | | | | | | | | |
| 256 | Variable 329 | QEPVTFEK | Bacillus | Bacteria | Firmicutes | | Bacilli | | Bacillales | | Bacillaceae | Bacillus | | | | | | | | GO:0005737(5%) | cytoplasm(5%) | GO:0031177(100%);GO:000431 | phosphopantetheine binding(100%);3-oxoacyl-[acyl-carrier-protein] | GO:0006633(95%);GO:0017000(5%) | fatty acid biosynthetic process(95%);antibiotic biosyn | IPR009081(100%);IPR013968( | 1.00E+00 | 8.39E+04 | 1.00E+00 | NS | NS | NS | NS | NS | NS | NS | NS |

| | | | | | | | | | | | | | | | | | | | | | | | | | | | | | | | | | | | | | | | | |
|---|---|---|---|---|---|---|---|---|---|---|---|---|---|---|---|---|---|---|---|---|---|---|---|---|---|---|---|---|---|---|---|---|---|---|---|---|---|---|---|---|
| | | | | | | | | | | | | | | | | | | | | | | 5(95%);GO:0016746(5%) | synthase activity (95%);acyltransferase activity (5%) | | thetic process (5%) | 100%);IPR014030(100%) | | | | | | | | | | | | | | |
| 257 | Variable 33 | MHTVVVGTSGVTASDVLAVAR | Actinomycetia | Bacteria | Actinobacteria | | Actinomycetia | | | | | | | | | | | 4.3.1.3(100%) | Histidine ammonia-lyase (100%) | GO:0005737(100%) | cytoplasm (100%) | GO:0004397(100%);GO:0016 | histidine ammonia-lyase activity (100%);ammonia-lyase activit | GO:0019556(97%);GO:0019557(97%);GO:000654 | histidine catabolic process to glutamate and for | IPR001106(100%);IPR00592 | 1.00E+00 | 1.00E+00 | 1.75E+05 | 0.50158 | 1.0494E-07 | 0.00050658 | 0.9 | NS | NS | NS | NS | 8.6773 | 0.00070098 | 0.000013614 |

| | | | | | | | | | | | | | | | | | | | | | | | | | | |
|---|---|---|---|---|---|---|---|---|---|---|---|---|---|---|---|---|---|---|---|---|---|---|---|---|---|---|
| | | | | | | | | | | | | | | | | | | | | | | 841 (1%) | y (1%) | 8 (4%) | mamide (97%); histidine catabolic process to glutamate and formate (97%); histidine ca | 1 (100%); IPR008948 (100%) |

| | | | | | | | | | | | | | | | | | | | | | | | | | | | | | | | | | | | | | |
|---|---|---|---|---|---|---|---|---|---|---|---|---|---|---|---|---|---|---|---|---|---|---|---|---|---|---|---|---|---|---|---|---|---|---|---|---|---|
| | | | | | | | | | | | | | | | | | | | | | | | | | tabolic process (4%) | | | | | | | | | | | | |
| 258 | Variable 330 | TAMNADYSWGK | Bacillus | Bacteria | Firmicutes | | Bacilli | | Bacillales | | Bacillaceae | Bacillus | | | | | | 2.4.1.21(100%) | Starch synthase (100%) | | | GO:0009011(100%);GO:0033201(100%);G | starch synthase activity (100%);alpha-1 | 4-glucan synthase activity (100%);glycogen (starch) synthase activity (90%) | GO:0005978(100%) | glycogen biosynthetic process (100%) | 1.00E+00 | 1.03E+05 | 1.00E+00 | NS | NS | NS | NS | NS | NS | NS | NS |

| | | | | | | | | | | | | | | | | | | | | | | | | | | | | | | | | | | | | | |
|---|---|---|---|---|---|---|---|---|---|---|---|---|---|---|---|---|---|---|---|---|---|---|---|---|---|---|---|---|---|---|---|---|---|---|---|---|---|
| | | | | | | | | | | | | | | | | | | | | | | O:0004373(90%) | | | | | | | | | | | | | | | |
| 259 | Variable 331 | EGAPFYQVLK | Bacillus | Bacteria | Firmicutes | | Bacilli | | Bacillales | | Bacillaceae | Bacillus | | | | | | | | GO:0005886(100%);GO:0016021(10 | plasma membrane(100%);integral component of membrane | | | GO:0009306(92%);GO:0030420(4%) | protein secretion(92%);establishment of competence for tra | IPR003004(100%);IPR018076(100% | 1.00E+00 | 1.80E+05 | 1.00E+00 | NS | NS | NS | NS | NS | NS | NS | NS |

| | | | | | | | | | | | | | | | | | | | | | | | | | | | | | | | | | | | | | |
|---|---|---|---|---|---|---|---|---|---|---|---|---|---|---|---|---|---|---|---|---|---|---|---|---|---|---|---|---|---|---|---|---|---|---|---|---|---|
| | | | | | | | | | | | | | | | | | | | | 0%) | (100%) | | | | nsformation (4%) | );IPR0042094(100%) | | | | | | | | | | | |
| 260 | Variable 332 | GTYVEDQNGK | Bacillus | Bacteria | Firmicutes | | Bacilli | | Bacillales | | Bacillaceae | Bacillus | | | | | | 2.6.1.11(100%) | Acetylornithine transaminase (100%) | GO:0005737(100%) | cytoplasm(100%) | GO:0003992(100%);GO:0030170( | N2-acetyl-L-ornithine:2-oxoglutarate 5-aminotransferase activity (100%);p | GO:0006526(100%) | arginine biosynthetic process (100%) | IPR005814(100%);IPR015421(10 | 8.23E+05 | 1.00E+00 | 1.00E+00 | NS | NS | NS | NS | NS | NS | NS | NS |

| | | | | | | | | | | | | | | | | | | | | | | | | | | | | | | | | | | | | | |
|---|---|---|---|---|---|---|---|---|---|---|---|---|---|---|---|---|---|---|---|---|---|---|---|---|---|---|---|---|---|---|---|---|---|---|---|---|---|
| | | | | | | | | | | | | | | | | | | | | | | 96%);GO:0042802(4%) | yridoxal phosphate binding(96%);identical protein binding(4%) | | | 0%);IPR015422(100%) | | | | | | | | | | | |
| 261 | Variable 333 | FFGCSNHPNCNK | Bacillus cereus group | Bacteria | Firmicutes | | Bacilli | | Bacillales | | Bacillaceae | Bacillus | Bacillus cereus group | | | | | 5.6.2.1(100%) | DNA topoisomerase (100%) | | | GO:0003677(100%);GO: | DNA binding(100%);DNA topoisomerase type I (si | ATP-independent) activity (100%);metal ion | GO:0006265(100%);GO:0009405(25% | DNA topological change (10 | 1.00E+00 | 1.16E+05 | 1.00E+00 | NS | NS | NS | NS | NS | NS | NS | NS |

| | | | | | | | | | | | | | | | | | | | | | | | | | | | | | | | | | | | | | | | | |
|---|---|---|---|---|---|---|---|---|---|---|---|---|---|---|---|---|---|---|---|---|---|---|---|---|---|---|---|---|---|---|---|---|---|---|---|---|---|---|---|---|
| | | | | | | | | | | | | | | | | | | | | | | 0003917(100%);GO:0046872(25%) | ngle strand cut | binding (25%) | ) | 0%); obsolete pathogenesis (25%) | | | | | | | | | | | | | | |
| 263 | Variable 335 | QLGFMCGAELVR | Bacillus subtilis grou | Bacteria | Firmicutes | | Bacilli | | Bacillales | | Bacillaceae | Bacillus | Bacillus subtilis gr | | | | | 2.6.1.62(95%);2. | Adenosylmethionine-8-amino-7-oxononanoa | GO:0005737(95%) | cytoplasm(95%) | GO:0030170(100% | pyridoxal phosphate binding (100%);adenos | GO:0009102(95%) | biotin biosynthetic process (95 | IPR005814(100%); | 1.00E+00 | 1.00E+00 | 4.60E+04 | 6.936 | 3.6717E-06 | 0.00014726 | 0.9 | NS | NS | NS | NS | 8.8002 | 0.00064271 | 0.000888 |

| | | | | | | | | | | | | | | | | | | | | | | | | | | | | | | | | | | | | | |
|---|---|---|---|---|---|---|---|---|---|---|---|---|---|---|---|---|---|---|---|---|---|---|---|---|---|---|---|---|---|---|---|---|---|---|---|---|---|
| | | | p | | | | | | | | | | oup | | | | | 6.1.105(5%) | tetransaminase (95%); Lysine--8-amino-7-oxononanoatetransaminase (5%) | | | );GO:0004015(95%);GO:0008483(5%) | ylmethionine-8-amino-7-oxononanoate transaminase activity (95%);transaminase activity (5%) | | %) | IPR015421(100%);IPR015422(100%) | | | | | | | | | | | |
| 264 | Variable | EQSLYEELQSAQMK | Bacillussu | Bacteria | Firmicute | | Bacilli | | Bacillales | | Bacillaceae | Bacillus | Bacillus | | | | | | | | | GO:0045 | exonuclease activity | GO:0006260 (9%) | DNA replication | IPR0241 | 1.00E+00 | 1.00E+00 | 2.63E+05 | 30.02 | 0.001769 | 0.0000804 | 0.9 | NS | NS | NS | NS |

| | | | | | | | | | | | | | | | | | | | | | | | | | | | | | | | | | | | | | |
|---|---|---|---|---|---|---|---|---|---|---|---|---|---|---|---|---|---|---|---|---|---|---|---|---|---|---|---|---|---|---|---|---|---|---|---|---|---|
| | 336 | | btilisgroup | | s | | | | | | | | ubtilisgroup | | | | | | | | | 27(100%);GO:0004519(9%);GO:0005524(9%) | (100%);endonuclease activity (9%);ATP binding (9%) | ;GO:0006310 (9%) | (9%); DNA recombination (9%) | 7(100%);IPR038729(100%) | | | | | 6 | 09 | | | | | |
| 265 | Variabl | EVANLDDSEFSGK | Bacillus | Bacteri | Firmicut | | Bacilli | | Bacillales | | Bacillaceae | Bacillus | Bacillus | | Bacillus | | | 3.1.-.-(1 | Acting on ester bonds | | | GO:0004 | nuclease activity (10 | | | IPR0068 | 1.00E+0 | 1.00E+0 | 7.20E+0 | NS | NS | NS | NS | NS | NS | NS | NS |

| | | | | | | | | | | | | | | | | | | | | | | | | | | | | | | | | | | | | | |
|---|---|---|---|---|---|---|---|---|---|---|---|---|---|---|---|---|---|---|---|---|---|---|---|---|---|---|---|---|---|---|---|---|---|---|---|---|---|
| | e 337 | | ubtilis | a | es | | | | | | | | subtilis group | | ubtilis | | | 00%) | (100%) | | | 518(100%) | 0%) | | | 29(100%);IPR026834(100%) | 0 | 0 | 4 | | | | | | | | |
| 266 | Variable 338 | EDAAEETAAPGK | Bacillus subtilis | Bacteria | Firmicutes | | Bacilli | | Bacillales | | Bacillaceae | Bacillus | Bacillus subtilis group | | Bacillus subtilis | | | 2.7.1.199(100%) | Protein-N(pi)-phosphohistidine--D-glucose phosphotransfe | GO:0005886(93%);GO:00 | plasma membrane(93%);integral compo | GO:0008982(100%);GO:0 | protein-N(PI)-phosphohistidine-sugar phosphotransferas | GO:0009401(100%) | phosphoenolpyruvate-dependent sugar phos | IPR001127(100%);IPR00 | 1.00E+00 | 1.00E+00 | 3.75E+04 | NS | NS | NS | NS | NS | NS | NS | NS |

| | | | | | | | | | | | | | | | | | | | | | | | | | | | | | | | | | | | | | |
|---|---|---|---|---|---|---|---|---|---|---|---|---|---|---|---|---|---|---|---|---|---|---|---|---|---|---|---|---|---|---|---|---|---|---|---|---|---|
| | | | | | | | | | | | | | | | | | | | rase (100%) | 16021 (93%);GO:0005887 (7%) | nent of membrane (93%);integral component of plasma membrane (7%) | 0163001 (100%);GO:0005355 (93%) | e activity (100%);kinase activity (100%);glucose transmembrane transporter activity (93%) | | photransferase system (100%) | 1996 (100%);IPR003352 (100%) | | | | | | | | | | | |
| 267 | Variable | FHYGGSYK | Bacillussu | Bacteria | Firmicute | | Bacilli | | Bacillales | | Bacillaceae | Bacillus | Bacilluss | | Bacillussu | | | | | | | GO:00167 | hydrolase activity (10 | | | IPR00132 | 1.00E+00 | 1.00E+00 | 6.14E+05 | 30.319 | 0.031963 | 0.000333 | 0.9 | NS | NS | NS | NS |

| | | | | | | | | | | | | | | | | | | | | | | | | | | | | | | | | | | | | | | | | |
|---|---|---|---|---|---|---|---|---|---|---|---|---|---|---|---|---|---|---|---|---|---|---|---|---|---|---|---|---|---|---|---|---|---|---|---|---|---|---|---|---|
| | 339 | | btilis | | s | | | | | | | | ubtilis group | | btilis | | | | | | | 87(100%) | 0%) | | | 2(100%);IPR004843(100%);IPR011401(100%) | | | | | | 33 | | | | | | | | |
| 268 | Variab | EYAEAQPLAGAR | Actinomy | Bacter | Actinob | | Actinomy | | | | | | | | | | | 3.3.1. | Adenosylhomocys | GO:000 | cytoplasm(9 | GO:000 | adenosylhomocyst | GO:0006730 | one-carbonom | IPR000 | 1.00E+ | 1.00E+ | 2.82E+ | 4.1603 | 6.3574 | 0.00019 | 0.9 | NS | NS | NS | NS | 8.3771 | 0.008 | 0.005 |

| | | | | | | | | | | | | | | | | | | | | | | | | | | | | | | | | | | | | | | | | |
|---|---|---|---|---|---|---|---|---|---|---|---|---|---|---|---|---|---|---|---|---|---|---|---|---|---|---|---|---|---|---|---|---|---|---|---|---|---|---|---|---|
| | le 34 | | cetia | ia | acteria | | cetia | | | | | | | | | | | 1(100%) | teinase(100%) | 5737(96%);GO:0005829(0%) | 6%);cytosol(0%) | 4013(100%) | einase activity(100%) | (98%);GO:0033353(0%) | etabolic process(98%);S-adenosylmethionine cycle(0%) | 0433(100%);IPR0042172(100%);IPR015878(98%) | 00 | 00 | 04 | | E-06 | 496 | | | | | | | 6777 | 3438 |
| 269 | Varia | NDSTGSDTNK | Bacillu | Bacte | Firmic | | Bacilli | | Bacillales | | Bacillacea | Bacillu | Bacill | | Bacillu | | | 2.4.1 | Peptidoglycan | GO:00 | plasmam | GO:00 | penicillin bin | GO:000836 | regulation | IPR00 | 1.00E | 1.00E | 2.96E | 0.368 | 9.046 | 0.0024 | 0.9 | NS | NS | NS | NS | 23.42 | 1.467 | 0.006 |

| | | | | | | | | | | | | | | | | | | | | | | | | | | | | | | | | | | | | | | | | |
|---|---|---|---|---|---|---|---|---|---|---|---|---|---|---|---|---|---|---|---|---|---|---|---|---|---|---|---|---|---|---|---|---|---|---|---|---|---|---|---|---|
| | ble 340 | | ssubtilis | ria | utes | | | | | | e | s | ussubtilisgroup | | ssubtilis | | | .129(100%);3.4.16.4(100%) | glycosyltransferase(100%);Serine-type D-Ala-D-Ala carboxypeptidase(100%) | 05886(8%);GO:0016021(8%) | embrane(8%);integral component of membrane(8%) | 08658(100%);GO:0008955(100%);GO:0009002(100%) | ding(100%);peptidoglycan glycosyltransferase activity(100%);serine-type D-Ala-D-Ala carboxypeptidase activity(100% | 0(8%);GO:0009252(8%);GO:0046677(8%) | of cell shape(8%);peptidoglycan biosynthetic process(8%);response to antibiotic(8%) | 1264(100%);IPR001460(100%);IPR003961(100%) | +00 | +00 | +04 | 33 | 4E-11 | 102 | | | | | | 5 | 1E-07 | 1506 |

| | | | | | | | | | | | | | | | | | | | | | | | | | | | | | | | | | | | | | |
|---|---|---|---|---|---|---|---|---|---|---|---|---|---|---|---|---|---|---|---|---|---|---|---|---|---|---|---|---|---|---|---|---|---|---|---|---|---|
| | | | | | | | | | | | | | | | | | | | | | | | ) | | | | | | | | | | | | | | |
| 270 | Variable 341 | LLLAEMQVLK | Bacillus subtilis | Bacteria | Firmicutes | | Bacilli | | Bacillales | | Bacillaceae | Bacillus | Bacillus subtilis group | | Bacillus subtilis | | | 2.1.1.37 (100%) | DNA (cytosine-5-)-methyltransferase (100%) | | | GO:0003886 (100%) | DNA (cytosine-5-)-methyltransferase activity (100%) | GO:0009307 (100%) | DNA restriction-modification system (100%) | IPR001525 (100%); IPR018117 (100%); IPR029063 (10 | 1.00E+00 | 1.00E+00 | 2.09E+05 | NS | NS | NS | NS | NS | NS | NS | NS |

| | | | | | | | | | | | | | | | | | | | | | | | | | | | | | | | | | | | | | | | | |
|---|---|---|---|---|---|---|---|---|---|---|---|---|---|---|---|---|---|---|---|---|---|---|---|---|---|---|---|---|---|---|---|---|---|---|---|---|---|---|---|---|
| | | | | | | | | | | | | | | | | | | | | | | | | | | 0%) | | | | | | | | | | | | | | |
| 271 | Variable 342 | GVPLDDMK | Bacillus subtilis | Bacteria | Firmicutes | | Bacilli | | Bacillales | | Bacillaceae | Bacillus | Bacillus subtilis group | | Bacillus subtilis | | | 3.4.14.13 (100%) | Gamma-D-glutamyl-L-lysine dipeptidyl-peptidase (100%) | | | GO:0016787 (88%);GO:0008234 (13%) | hydrolase activity (88%);cysteine-type peptidase activity (13%) | GO:0016998 (13%);GO:0071555 (13%) | cell wall macromolecule catabolic process (13%);cell wall organization (13%) | IPR000064 (100%);IPR038765 (100%) | 1.00E+00 | 1.00E+00 | 3.81E+04 | 0.14238 | 9.6104E-15 | 0.0022668 | 0.9 | NS | NS | NS | NS | 104.15 | 5.2519E-17 | 0.005817 |
| 272 | Var | FGEVGLTL | Baci | Bac | Firm | | Bacill | | Bacill | | Bacill | Baci | Bac | | Baci | | | 3.1 | Type II | | | GO: | type II site | GO:00 | DNA | | 1.0 | 1.0 | 2.4 | NS | NS | NS | NS | NS | NS | NS | NS | | | |

| | | | | | | | | | | | | | | | | | | | | | | | | | | | | | | | | | | | | | | | | |
|---|---|---|---|---|---|---|---|---|---|---|---|---|---|---|---|---|---|---|---|---|---|---|---|---|---|---|---|---|---|---|---|---|---|---|---|---|---|---|---|---|
| | iable 343 | R | llussubtilis | teria | icutes | | i | | ales | | aceae | llus | illussubtilisgroup | | llussubtilis | | | .21.4(100%) | site-specific deoxyribonuclease (100%) | | | 0009036(100%) | -specific deoxyribonuclease activity (100%) | 09307(100%) | restriction-modification system (100%) | | 0E+00 | 0E+00 | 8E+04 | | | | | | | | | | | |
| 273 | Variable 344 | HPSLSAPAQSR | Bacillussubtilis | Bacteria | Firmicutes | | Bacilli | | Bacillales | | Bacillaceae | Bacillus | Bacillussubtilisgroup | | Bacillussubtilis | | | | | GO:0005886(100%);GO:00160 | plasmamembrane(100%);integral component of m | GO:0022857(100%) | transmembrane transporter activity (100%) | | | IPR001958(100%);IPR011701 | 1.00E+00 | 1.00E+00 | 2.85E+04 | 0.29713 | 1.049E-11 | 0.0019002 | 0.9 | NS | NS | NS | NS | 36.085 | 7.58E-10 | 0.0061438 |

| | | | | | | | | | | | | | | | | | | | | | | | | | | | | | | | | | | | | | | | | |
|---|---|---|---|---|---|---|---|---|---|---|---|---|---|---|---|---|---|---|---|---|---|---|---|---|---|---|---|---|---|---|---|---|---|---|---|---|---|---|---|---|
| | | | | | | | | | | | | | | | | | | | | 21(100%) | embrane(100%) | | | | | (100%); IPR020846(100%) | | | | | | | | | | | | | | |
| 274 | Variable 345 | YFLSAEMQK | Bacillus subtilis | Bacteria | Firmicutes | | Bacilli | | Bacillales | | Bacillaceae | Bacillus | Bacillus subtilis group | | Bacillus subtilis | | | 3.1.21.4(100%) | Type II site-specific deoxyribonuclease (100%) | | | GO:0009036(100%) | type II site-specific deoxyribonuclease activity (100%) | GO:0009307(100%) | DNA restriction-modification system (100%) | | 1.00E+00 | 1.00E+00 | 2.89E+04 | NS | NS | NS | NS | NS | NS | NS | NS | 4.9958 | 0.01314 | 0.0043179 |

| | | | | | | | | | | | | | | | | | | | | | | | | | | | | | | | | | | | | | |
|---|---|---|---|---|---|---|---|---|---|---|---|---|---|---|---|---|---|---|---|---|---|---|---|---|---|---|---|---|---|---|---|---|---|---|---|---|---|
| 275 | Variable 346 | ESVMSFTLSK | Bacillus subtilis | Bacteria | Firmicutes | | Bacilli | | Bacillales | | Bacillaceae | Bacillus | Bacillus subtilis group | | Bacillus subtilis | | | | | GO:0016021(100%);GO:0005886(25%) | integral component of membrane (100%);plasma membrane (25%) | GO:0003677(25%) | DNA binding (25%) | | | IPR011646(100%);IPR027417(100%) | 1.00E+00 | 1.71E+05 | 1.00E+00 | NS | NS | NS | NS | NS | NS | NS | NS |
| 276 | Variable 34 | SFEDQDSAR | Bacillus subtilis | Bacteria | Firmicutes | | Bacilli | | Bacillales | | Bacillaceae | Bacillus | Bacillus subti | | Bacillus subtili | | | | | | | GO:0004527( | exonuclease activity (100%) | | | IPR011990(1 | 1.00E+00 | 8.12E+05 | 1.00E+00 | NS | NS | NS | NS | NS | NS | NS | NS |

| | | | | | | | | | | | | | | | | | | | | | | | | | | | | | | | | | | | | | |
|---|---|---|---|---|---|---|---|---|---|---|---|---|---|---|---|---|---|---|---|---|---|---|---|---|---|---|---|---|---|---|---|---|---|---|---|---|---|
| | 7 | | | | | | | | | | | | lisgroup | | s | | | | | | | 100%) | | | | 00%); IPR012337 (100%); IPR038720 (100%) | | | | | | | | | | | |
| 277 | Variable | ETSADSVK | Bacillus subt | Bacteria | Firmicutes | | Bacilli | | Bacillales | | Bacillaceae | Bacillus | Bacillus su | | Bacillus sub | | | | | GO:0000557 | extracellular region (1 | | | GO:0030435 (100%) | sporulation resulting | IPR007253 | 1.00E+00 | 8.60E+04 | 1.00E+00 | NS | NS | NS | NS | NS | NS | NS | NS |

| | | | | | | | | | | | | | | | | | | | | | | | | | | | | | | | | | | | | | |
|---|---|---|---|---|---|---|---|---|---|---|---|---|---|---|---|---|---|---|---|---|---|---|---|---|---|---|---|---|---|---|---|---|---|---|---|---|---|
| | 348 | | ilis | | | | | | | | | | btilis group | | tilis | | | | | 6(11%);GO:0005618(11%);GO:0005886(11%) | 1%);cell wall(11%);plasma membrane(11%) | | | ;GO:0071555(11%) | information of a cellular spore(100%);cell wall organization(11%) | (100%);IPR013486(100%);IPR013693(100%) | | | | | | | | | | | |
| 278 | Variabl | NNGDTPNNPK | Bacillus | Bacteri | Firmicut | | Bacilli | | Bacillales | | Bacillaceae | Bacillus | Bacillus | | Bacillus | | | 3.1.3.1 | Alkaline phosphatase | | | GO:0004 | alkaline phosphatase | GO:0016311(6 | dephosphorylati | IPR0019 | 1.00E+0 | 4.83E+0 | 1.00E+0 | NS | NS | NS | NS | NS | NS | NS | NS |

| | | | | | | | | | | | | | | | | | | | | | | | | | | | | | | | | | | | | | |
|---|---|---|---|---|---|---|---|---|---|---|---|---|---|---|---|---|---|---|---|---|---|---|---|---|---|---|---|---|---|---|---|---|---|---|---|---|---|
| | e 349 | | ubtilis | a | es | | | | | | | | subtilis group | | ubtilis | | | (100%) | (100%) | | | 035(50%);GO:0016791(50%);GO:0046872(6%) | activity (50%);phosphatase activity (50%);metal ion binding (6%) | %) | on (6%) | 52(100%);IPR017850(100%);IPR018299(100%) | 0 | 4 | 0 | | | | | | | | |
| 279 | Varia | SEGKPEAALPK | Actinom | Bacte | Actino | | Actinom | | | | | | | | | | | | | GO:00 | cytoplasm | GO:00 | translation elo | | | IPR00 | 1.00E | 1.00E | 1.27E | NS | NS | NS | NS | NS | NS | NS | NS |

| | | | | | | | | | | | | | | | | | | | | | | | | | | | | | | | | | | | | | |
|---|---|---|---|---|---|---|---|---|---|---|---|---|---|---|---|---|---|---|---|---|---|---|---|---|---|---|---|---|---|---|---|---|---|---|---|---|---|
| | ble 35 | | ycetia | ria | bacteria | | ycetia | | | | | | | | | | | | | 05737(100%) | (100%) | 03746(100%) | ngation factor activity (100%) | | | 1816(100%);IPR0090600(100%);IPR0140039(100%) | +00 | +00 | +06 | | | | | | | | |
| 280 | Var | QNVADLS | Geo | Bac | Firm | | Bacill | | Bacill | | Bacill | Geo | | | | | | | | GO: | cytopl | GO: | isomeras | | | IPR | 1.0 | 1.0 | 4.3 | 4.2 | 0.0 | 0.00 | 0.9 | NS | NS | NS | NS |

| | | | | | | | | | | | | | | | | | | | | | | | | | | | | | | | | | | | | | |
|---|---|---|---|---|---|---|---|---|---|---|---|---|---|---|---|---|---|---|---|---|---|---|---|---|---|---|---|---|---|---|---|---|---|---|---|---|---|
| | iable 350 | VLAECAVK | bacillus | teria | icutes | | i | | ales | | aceae | bacillus | | | | | | | | 0005737(14%) | asm(14%) | 0016853(86%) | e activity(86%) | | | 036249(100%) | 0E+00 | 0E+00 | 3E+04 | 523 | 018942 | 01534 | | | | | |
| 281 | Variable 351 | EMADFFXATVANGADPK | Geobacillus stearothermophilus | Bacteria | Firmicutes | | Bacilli | | Bacillales | | Bacillaceae | Geobacillus | | | Geobacillus stearothermophilus | | | 6.3.5.-(100%) | Carbon--nitrogen ligases with glutamine as amido-N-donor(100%) | | | GO:0005524(100%);GO:0050566(100% | ATP binding(100%);asparaginyl-tRNA synthase(glutamine-hydrolyzing) activity(10 | GO:0006412(100%) | translation(100%) | IPR003789(100%);IPR004413(100%);I | 1.00E+00 | 1.00E+00 | 2.74E+04 | NS | NS | NS | NS | NS | NS | NS | NS |

| | | | | | | | | | | | | | | | | | | | | | | | | | | | | | | | | | | | | | |
|---|---|---|---|---|---|---|---|---|---|---|---|---|---|---|---|---|---|---|---|---|---|---|---|---|---|---|---|---|---|---|---|---|---|---|---|---|---|
| | | | | | | | | | | | | | | | | | | | | | | ); GO:0050567(100%) | 0%);glutaminyl-tRNA synthase (glutamine-hydrolyzing) activity (100%) | | | PR006075(100%) | | | | | | | | | | | |
| 282 | Variable 352 | HYMPGLDGLR | Lysinibacillus | Bacteria | Firmicutes | | Bacilli | | Bacillales | | Bacillaceae | Lysinibacillus | | | | | | | | GO:0016021(100%) | integral component of membrane (1 | GO:0016747(100%) | acyltransferase activity | transferring groups other than amino-ac | | | 1.00E+00 | 5.95E+05 | 1.00E+00 | NS | NS | NS | NS | NS | NS | NS | NS |

| | | | | | | | | | | | | | | | | | | | | | | | | | | | | | | | | | | | | | |
|---|---|---|---|---|---|---|---|---|---|---|---|---|---|---|---|---|---|---|---|---|---|---|---|---|---|---|---|---|---|---|---|---|---|---|---|---|---|
| | | | | | | | | | | | | | | | | | | | | | 00%) | | | ylgroups(100%) | | | | | | | | | | | | | |
| 283 | Variable 353 | FYAVEEEDAK | Listeria | Bacteria | Firmicutes | | Bacilli | | Bacillales | | Listeriaceae | Listeria | | | | | | | | GO:0005737(100%) | cytoplasm(100%) | | | GO:0000917(100%);GO:0043093(100%) | division septum assembly(100%);FtsZ-dependent cytokinesis(100%) | IPR007561(100%);IPR023052(100%);IPR038 | 1.00E+00 | 1.00E+00 | 8.79E+04 | 2.1387 | 0.012821 | 0.00077456 | 0.9 | NS | NS | NS | NS |

| | | | | | | | | | | | | | | | | | | | | | | | | | | | | | | | | | | | | | |
|---|---|---|---|---|---|---|---|---|---|---|---|---|---|---|---|---|---|---|---|---|---|---|---|---|---|---|---|---|---|---|---|---|---|---|---|---|---|
| | | | | | | | | | | | | | | | | | | | | | | | | | | 594(100%) | | | | | | | | | | | |
| 284 | Variable 354 | NAPDEQDGMFK | Listeria | Bacteria | Firmicutes | | Bacilli | | Bacillales | | Listeriaceae | Listeria | | | | | | 6.3.5.-(100%) | Carbon--nitrogen ligases with glutamine as amido-N-donor(100%) | | | GO:0050566(100%);GO:0050567(100%);GO | asparaginyl-tRNA synthase (glutamine-hydrolyzing) activity(100%);glutaminyl-tRNA syntha | GO:0006412(100%);GO:0006450(100%);GO:0070681(5%) | translation(100%);regulation of translational fidelity(100%);glutaminyl- | IPR003837(100%);IPR036113(100%) | 2.41E+06 | 1.00E+00 | 1.00E+00 | NS | NS | NS | NS | NS | NS | NS | NS |

| | | | | | | | | | | | | | | | | | | | | | | | | | | | | | | | | | | | | | |
|---|---|---|---|---|---|---|---|---|---|---|---|---|---|---|---|---|---|---|---|---|---|---|---|---|---|---|---|---|---|---|---|---|---|---|---|---|---|
| | | | | | | | | | | | | | | | | | | | | | | :0005524(95%) | se(glutamine-hydrolyzing) activity(100%);ATP binding(95%) | | tRNA Gln biosynthesis via transamidation(5%) | | | | | | | | | | | | |
| 286 | Variable 356 | QLDEVTQLATEK | Jeotgalibacaporci | Bacteria | Firmicutes | | Bacilli | | Lactobacillales | | Carnobacteriaceae | Jeotgalibaca | | | Jeotgalibacaporci | | | | | | | GO:0016491(100%) | oxidoreductase activity(100%) | | | IPR000683(100%);IPR03 | 1.00E+00 | 1.00E+00 | 1.11E+05 | NS | NS | NS | NS | NS | NS | NS | NS |

| | | | | | | | | | | | | | | | | | | | | | | | | | | | | | | | | | | | | | |
|---|---|---|---|---|---|---|---|---|---|---|---|---|---|---|---|---|---|---|---|---|---|---|---|---|---|---|---|---|---|---|---|---|---|---|---|---|---|
| | | | | | | | | | | | | | | | | | | | | | | | | | | 6291(100%) | | | | | | | | | | | |
| 287 | Variable 357 | NGQNGMSK | Lactobacillaceae | Bacteria | Firmicutes | | Bacilli | | Lactobacillales | | Lactobacillaceae | | | | | | | 3.6.5.-(100%) | Acting on GTP; involved in cellular and subcellular movement (100%) | GO:0005737(100%) | cytoplasm(100%) | GO:0000287(100%);GO:0003924(100%);G | magnesium ion binding(100%);GTPase activity(100%);GTP binding(100%) | GO:0042254(100%) | ribosome biogenesis(100%) | IPR006073(100%);IPR006169(100%);IPR | 1.00E+00 | 1.00E+00 | 3.82E+04 | NS | NS | NS | NS | NS | NS | NS | NS |

| | | | | | | | | | | | | | | | | | | | | | | | | | | | | | | | | | | | | | |
|---|---|---|---|---|---|---|---|---|---|---|---|---|---|---|---|---|---|---|---|---|---|---|---|---|---|---|---|---|---|---|---|---|---|---|---|---|---|
| | | | | | | | | | | | | | | | | | | | | | | O:0005525(100%) | | | | 014100(100%) | | | | | | | | | | | |
| 288 | Variable 358 | LFNDDQQ | Lactobacillaceae | Bacteria | Firmicutes | | Bacilli | | Lactobacillales | | Lactobacillaceae | | | | | | | | | GO:0005840(100%) | ribosome(100%) | GO:0003735(100%);GO:0019843(1 | structural constituent of ribosome(100%);rRNA binding(100%) | GO:0006412(100%) | translation(100%) | IPR012677(100%);IPR012678(100 | 1.00E+00 | 1.00E+00 | 8.44E+04 | NS | NS | NS | NS | NS | NS | NS | NS |

| | | | | | | | | | | | | | | | | | | | | | | | | | | | | | | | | | | | | | |
|---|---|---|---|---|---|---|---|---|---|---|---|---|---|---|---|---|---|---|---|---|---|---|---|---|---|---|---|---|---|---|---|---|---|---|---|---|---|
| | | | | | | | | | | | | | | | | | | | | | | 00%) | | | | %);IPR013025(100%) | | | | | | | | | | | |
| 289 | Variable 359 | LNELKPATGSR | Lactobacillaceae | Bacteria | Firmicutes | | Bacilli | | Lactobacillales | | Lactobacillaceae | | | | | | | | | GO:0015934(100%) | large ribosomal subunit (100%) | GO:0003735(100%);GO:0019843 | structural constituent of ribosome (100%);rRNA binding (100%) | GO:0006412 (100%) | translation (100%) | IPR001196(100%);IPR005749(1 | 1.00E+00 | 6.48E+04 | 1.00E+00 | NS | NS | NS | NS | NS | NS | NS | NS |

| | | | | | | | | | | | | | | | | | | | | | | | | | | | | | | | | | | | | | |
|---|---|---|---|---|---|---|---|---|---|---|---|---|---|---|---|---|---|---|---|---|---|---|---|---|---|---|---|---|---|---|---|---|---|---|---|---|---|
| | | | | | | | | | | | | | | | | | | | | | | (100%) | | | | 00%);IPR0211131(100%) | | | | | | | | | | | |
| 290 | Variable 36 | TQVCGVMAGR | Actinomycetia | Bacteria | Actinobacteria | | Actinomycetia | | | | | | | | | | | 2.6.1.36(100%) | L-lysine 6-transaminase (100%) | | | GO:0030170(100%);GO:00454 | pyridoxal phosphate binding (100%);L-lysine 6-transaminase activit | GO:0017000(97%);GO:0009450(1%) | antibiotic biosynthetic process (97%); gamma-ami | IPR005814(100%);IPR015421 | 1.00E+00 | 1.00E+00 | 4.70E+04 | NS | NS | NS | NS | NS | NS | NS | NS |

| | | | | | | | | | | | | | | | | | | | | | | | | | | | | | | | | | | | | | |
|---|---|---|---|---|---|---|---|---|---|---|---|---|---|---|---|---|---|---|---|---|---|---|---|---|---|---|---|---|---|---|---|---|---|---|---|---|---|
| | | | | | | | | | | | | | | | | | | | | | | 84(98%);GO:0008483(2%) | y(98%);transaminase activity(2%) | | nobutyric acid catabolic process(1%) | (100%);IPR015424(100%) | | | | | | | | | | | |
| 292 | Variable 361 | AADLVSEVEAQEAEAAVKPK | Lactobacillus delbrueckii | Bacteria | Firmicutes | | Bacilli | | Lactobacillales | | Lactobacillaceae | Lactobacillus | | | Lactobacillus delbrueckii | | | 3.6.5.-(100%) | Acting on GTP; involved in cellular and subcellular movement(10 | GO:0005737(100%) | cytoplasm(100%) | GO:0000287(100%);GO:000 | magnesium ion binding(100%);GTPase activity(100%);GTP bindin | GO:0042254(100%) | ribosome biogenesis(100%) | IPR006073(100%);IPR0060 | 1.00E+00 | 1.00E+00 | 1.28E+05 | 1.901 | 0.0041522 | 0.00089542 | 0.9 | NS | NS | NS | NS |

| | | | | | | | | | | | | | | | | | | | | | | | | | | | | | | | | | | | | | |
|---|---|---|---|---|---|---|---|---|---|---|---|---|---|---|---|---|---|---|---|---|---|---|---|---|---|---|---|---|---|---|---|---|---|---|---|---|---|
| | | | | | | | | | | | | | | | | | | | 0%) | | | 3924(100%);GO:0005525(100%) | g(100%) | | | 74(100%);IPR006169(100%) | | | | | | | | | | | |
| 293 | Variable 362 | VAVQTMDALGMTNYGR | Pediococcuspentosac | Bacteria | Firmicutes | | Bacilli | | Lactobacillales | | Lactobacillaceae | Pediococcus | | | Pediococcuspentos | | | 6.3.2.4(100%) | D-alanine--D-alanine ligase(100%) | GO:0005737(100%) | cytoplasm(100%) | GO:0005524(100%);G | ATP binding(100%);D-alanine-D-alanine lig | GO:0008360(100%);GO:0009252(100 | regulation of cell shape(100%); pepti | IPR000291(100%);IP | 1.47E+05 | 1.00E+00 | 1.00E+00 | NS | NS | NS | NS | NS | NS | NS | NS |

| | | | | | | | | | | | | | | | | | | | | | | | | | | | | | | | | | | | | | |
|---|---|---|---|---|---|---|---|---|---|---|---|---|---|---|---|---|---|---|---|---|---|---|---|---|---|---|---|---|---|---|---|---|---|---|---|---|---|
| | | | eus | | | | | | | | | | | | aceus | | | | | | | O:0008716(100%);GO:0046872(100%) | ase activity(100%);metal ion binding(100%) | %);GO:0071555(100%) | doglycan biosynthetic process(100%);cell wall organization(100%) | R0059005(100%);IPR011095(100%) | | | | | | | | | | | |
| 294 | Variable 363 | YLPLVEV | Streptococcus | Bacteria | Firmicutes | | Bacilli | | Lactobacillales | | Streptococcaceae | Streptococcus | | | | | | | | | | GO:0005524(1 | ATP binding(100%);proton-transp | rotational mechanism(93%) | GO:0042777(93%);GO: | plasmamembrane | 1.00E+00 | 1.00E+00 | 6.33E+04 | 3.1776 | 0.04545 | 0.00005824 | 0.9 | NS | NS | NS | NS |

| | | | | | | | | | | | | | | | | | | | | | | | | | |
|---|---|---|---|---|---|---|---|---|---|---|---|---|---|---|---|---|---|---|---|---|---|---|---|---|---|
| | | | | | | | | | | | | | | | | | | | | | 00%);GO:0046933(93%);GO:0016787(43%) | orting ATP synthase activity | ;hydrolase activity (43%) | 0006811 (7%) | ATP synthesis coupled proton transport (93%); ion transport ( |

| | | | | | | | | | | | | | | | | | | | | | | | | | | | | | | | | | | | | | | | |
|---|---|---|---|---|---|---|---|---|---|---|---|---|---|---|---|---|---|---|---|---|---|---|---|---|---|---|---|---|---|---|---|---|---|---|---|---|---|---|---|
| | | | | | | | | | | | | | | | | | | | | | | | | | 7%) | | | | | | | | | | | | | | |
| 295 | Variable 364 | SGLDTQADVLVDGK | Streptococcus | Bacteria | Firmicutes | | Bacilli | | Lactobacillales | | Streptococcaceae | Streptococcus | | | | | 3.5.2.3(100%) | Dihydroorotase (100%) | | | GO:0004151(100%);GO:0008270(93%);GO:004687 | dihydroorotase activity (100%);zinc ion binding (93%);metal ion binding (7%) | GO:0044205(93%);GO:0006221(2%) | 'de novo' UMP biosynthetic process(93%);pyrimidine nucleotide biosynthetic proc | IPR002195(100%);IPR006680(100%);IPR011059( | 1.00E+00 | 1.00E+00 | 3.37E+04 | 0.2144 | 5.1943E-13 | 0.0019461 | 0.9 | NS | NS | NS | NS | 61.384 | 3.4193E-13 | 0.0066573 |

| | | | | | | | | | | | | | | | | | | | | | | | | | | | | | | | | | | | | | |
|---|---|---|---|---|---|---|---|---|---|---|---|---|---|---|---|---|---|---|---|---|---|---|---|---|---|---|---|---|---|---|---|---|---|---|---|---|---|
| | | | | | | | | | | | | | | | | | | | | | | 2(7%) | | | ess(2%) | 100%) | | | | | | | | | | | |
| 296 | Variable 365 | AVVVTTGTALR | Streptococcus | Bacteria | Firmicutes | | Bacilli | | Lactobacillales | | Streptococcaceae | Streptococcus | | | | | | | | GO:0005737(99%) | cytoplasm(99%) | GO:0050660(100%) | flavin adenine dinucleotide binding(100%) | GO:0002098(97%);GO:0008033(3%) | tRNA wobble uridine modification(97%);tRNA processing(3%) | IPR002218(100%);IPR004416(100%);IPR02690 | 1.00E+00 | 3.77E+05 | 1.00E+00 | NS | NS | NS | NS | NS | NS | NS | NS |

| | | | | | | | | | | | | | | | | | | | | | | | | | | | | | | | | | | | | | |
|---|---|---|---|---|---|---|---|---|---|---|---|---|---|---|---|---|---|---|---|---|---|---|---|---|---|---|---|---|---|---|---|---|---|---|---|---|---|
| | | | | | | | | | | | | | | | | | | | | | | | | | | 4(100%) | | | | | | | | | | | |
| 297 | Variable 366 | LTSSQSAGK | Streptococcus mutans | Bacteria | Firmicutes | | Bacilli | | Lactobacillales | | Streptococcaceae | Streptococcus | | | Streptococcus mutans | | | | | GO:0005618(100%);GO:0005576(83%);GO:00 | cell wall(100%);extracellular region(83%);integral component of membrane(5 | GO:0005518(83%) | collagen binding(83%) | GO:0007155(83%) | cell adhesion(83%) | IPR008456(100%);IPR008966(100%);IPR011 | 1.00E+00 | 1.00E+00 | 2.04E+05 | NS | NS | NS | NS | NS | NS | NS | NS |

| | | | | | | | | | | | | | | | | | | | | | | | | | | | | | | | | | | | | | |
|---|---|---|---|---|---|---|---|---|---|---|---|---|---|---|---|---|---|---|---|---|---|---|---|---|---|---|---|---|---|---|---|---|---|---|---|---|---|
| | | | | | | | | | | | | | | | | | | | | 16021(50%) | 0%) | | | | | 252(100%) | | | | | | | | | | | |
| 298 | Variable 367 | EXPQWVR | Streptococcus pneumoniae G54 | Bacteria | Firmicutes | | Bacilli | | Lactobacillales | | Streptococcaceae | Streptococcus | | | Streptococcus pneumoniae | | Streptococcus pneumoniae G54 | 1.14.-.-(100%) | Acting on paired donors | with incorporation or reduction of molecular | | | GO:0016705 (100%) | oxidoreductase activity | acting on paired donors | with incorporation or reduction of molecular oxy | 1.00E+00 | 1.00E+00 | 2.75E+04 | NS | NS | NS | NS | NS | NS | NS | NS |

oxygen. The oxygen incorporated need not be derived from

gen (100%)

| | | | | | | | | | | | | | | | | | | | | | | | | | | | | | | | | | | | | | |
|---|---|---|---|---|---|---|---|---|---|---|---|---|---|---|---|---|---|---|---|---|---|---|---|---|---|---|---|---|---|---|---|---|---|---|---|---|---|
| | | | | | | | | | | | | | | | | | | | | O(2)(100%) | | | | | | | | | | | | | | | | | |
| 300 | Variable 369 | MGLTLTQK | Clostridia | Bacteria | Firmicutes | | Clostridia | | | | | | | | | | | 4.2.1.33(100%) | 3-isopropylmalatedehydratase(100%) | | | GO:0003861(100%);GO:0046872(100%); | 3-isopropylmalate dehydratase activity(100%);metal ion binding(100%);4 iron | 4 sulfur cluster binding(100%) | GO:0009082(100%);GO:0009098(100%) | branched-chain amino acid biosynthetic process | 1.00E+00 | 1.00E+00 | 5.04E+04 | NS | NS | NS | NS | NS | NS | NS | NS |

| | | | | | | | | | | | | | | | | | | | | | | | | | | | | | | | | | | | | | |
|---|---|---|---|---|---|---|---|---|---|---|---|---|---|---|---|---|---|---|---|---|---|---|---|---|---|---|---|---|---|---|---|---|---|---|---|---|---|
| | | | | | | | | | | | | | | | | | | | | | | GO:0051539(100%) | | | | ss(100%);leucine biosynthetic process(100%) | | | | | | | | | | | |
| 301 | Variable 37 | AYNNASPASGR | Actinomycetia | Bacteria | Actinobacteri | | Actinomycetia | | | | | | | | | | | 3.6.4.-(100% | Acting on ATP; involved in cellula | GO:0005618( | cell wall(0%);plasmam | GO:0003723( | RNA binding(100%);ATP bindin | GO:0006353(100%);GO: | DNA-templated transcri | termination(10 | 1.00E+00 | 1.00E+00 | 4.31E+04 | NS | NS | NS | NS | NS | NS | NS | NS |

| | | | | | | | | | | | | | | | | | | | | | | | | | | | | | | | | | | | | | | | | |
|---|---|---|---|---|---|---|---|---|---|---|---|---|---|---|---|---|---|---|---|---|---|---|---|---|---|---|---|---|---|---|---|---|---|---|---|---|---|---|---|---|
| | | | | | a | | | | | | | | | | | | | ) | r and subcellular movement(100%) | 0%);GO:0005886(0%) | embrane(0%) | 100%);GO:0005524(100%);GO:0004386(99%) | g(100%);helicase activity(99%) | 0006363(0%) | ption | 0%);termination of RNA polymerase I transcription(0%) | | | | | | | | | | | | | | |
| 302 | Variab | ELEDTEVELEEVEEE | Alkaliphilu | Bacter | Firmicu | | Clostridia | | Eubacteria | | Clostridiacea | Alkalip | | | Alkalip | | Alkalip | 3.6.5.- | Acting on GTP; inv | GO:000 | cytoplasm(1 | GO:000 | magnesium ion bin | GO:0042254 | ribosome bi | IPR005 | 1.00E+ | 1.00E+ | 5.13E+ | 6.51 | 6.935 | 0.0026 | 0.9 | NS | NS | NS | NS | 8.1869 | 0.009 | 0.009 |

| | | | | | | | | | | | | | | | | | | | | | | | | | | | | | | | | | | | | | | | |
|---|---|---|---|---|---|---|---|---|---|---|---|---|---|---|---|---|---|---|---|---|---|---|---|---|---|---|---|---|---|---|---|---|---|---|---|---|---|---|---|
| | le 370 | K | smetalliredigensQYMF | ia | tes | | | | les | | e | hilus | | | hilusmetalliredigens | | hilusmetalliredigensQYMF | (100%) | olved in cellular and subcellular movement (100%) | 5737(100%) | 00%) | 0287(100%);GO:0003924(100%);GO:0005525(100%) | ding(100%);GTPase activity(100%);GTP binding(100%) | (100%) | ogenesis(100%) | 225(100%);IPR006073(100%);IPR006074(100%) | 00 | 00 | 04 | | E-06 | 771 | | | | | | 9466 | 008 |
| 3 | V | SD | A | B | F | | C | | E | | Cl | A | | | A | | A | 3 | Tri | G | cy | G | me | G | pe | I | 1 | 1 | 9 | N | N | N | N | N | N | N | N | | |

| | | | | | | | | | | | | | | | | | | | | | | | | | | | | | | | | | | | | | |
|---|---|---|---|---|---|---|---|---|---|---|---|---|---|---|---|---|---|---|---|---|---|---|---|---|---|---|---|---|---|---|---|---|---|---|---|---|---|
| 03 | ariable 371 | HESESFPSTK | lkaliphilusoremlandiiOhILAs | acteria | irmicutes | | lostridia | | ubacteriales | | ostridiaceae | lkaliphilus | | | lkaliphilusoremlandii | | lkaliphilusoremlandiiOhILAs | .4.11.4(100%) | peptide aminopeptidase(100%) | O:0005737(100%) | toplasm(100%) | O:0008237(100%);GO:0008270(100%);GO:0045148(1 | talloptidase activity(100%);zinc ion binding(100%);tripeptide aminopeptidase activity(100%) | O:0043171(100%) | ptide catabolic process(100%) | PR001261(100%);IPR002933(100%);IPR010161(100% | .00E+00 | .00E+00 | .18E+04 | S | S | S | S | S | S | S | S |

| | | | | | | | | | | | | | | | | | | | | | | | | | | | | | | | | | | | | | |
|---|---|---|---|---|---|---|---|---|---|---|---|---|---|---|---|---|---|---|---|---|---|---|---|---|---|---|---|---|---|---|---|---|---|---|---|---|---|
| | | | | | | | | | | | | | | | | | | | | | | 00%) | | | | ) | | | | | | | | | | | |
| 304 | Variable 372 | FAEACYQTDGK | Clostridium | Bacteria | Firmicutes | | Clostridia | | Eubacteriales | | Clostridiaceae | Clostridium | | | | | | | | | | GO:0005524(100%);GO:0005525(100%) | ATP binding(100%);GTP binding(100%) | | | IPR005337(100%);IPR027417(100%) | 1.00E+00 | 1.00E+00 | 3.84E+04 | NS | NS | NS | NS | NS | NS | NS | NS |
| 305 | Variabl | SEYCVAVTGEVK | Clostridium | Bacteri | Firmicut | | Clostridia | | Eubacteriale | | Clostridiaceae | Clostridi | | | | | | 6.1.1.1 | Aspartate--tRNA lig | GO:0005 | cytoplasm (100 | GO:0003 | nucleic acid bindin | GO:0006422 (1 | aspartyl-tRN | IPR0023 | 1.00E+0 | 5.15E+0 | 1.00E+0 | NS | NS | NS | NS | NS | NS | NS | NS |

| | | | | | | | | | | | | | | | | | | | | | | | | | | | | | | | | | | | | | |
|---|---|---|---|---|---|---|---|---|---|---|---|---|---|---|---|---|---|---|---|---|---|---|---|---|---|---|---|---|---|---|---|---|---|---|---|---|---|
| | e 373 | | | a | es | | | | s | | | um | | | | | | 2(100%) | ase(100%) | 737(100%) | %) | 676(100%);GO:0004815(100%);GO:0005524(100%) | g(100%);aspartate-tRNA ligase activity(100%);ATP binding(100%) | 00%) | A aminoacylation(100%) | 12(100%);IPR004115(100%);IPR004364(100%) | 0 | 4 | 0 | | | | | | | | |
| 30 | Va | ASQY | Clo | Ba | Fir | | Clo | | Eu | | Clos | Cl | | | | | | 2. | Starch | | | GO | starch | 4-glu | | | 4. | 1. | 1. | NS | NS | NS | NS | NS | NS | NS | NS |

| | | | | | | | | | | | | | | | | | | | | | | | | | | | | | | | | | | | | | | | | |
|---|---|---|---|---|---|---|---|---|---|---|---|---|---|---|---|---|---|---|---|---|---|---|---|---|---|---|---|---|---|---|---|---|---|---|---|---|---|---|---|---|
| 6 | riable 374 | SWQK | stridium | cteria | micutes | | stridia | | bacteriales | | tridiaceae | ostridium | | | | | | 4.1.21(100%) | synthase (100%) | | | :0009011(100%);GO:0033201(100%) | synthase activity (100%);alpha-1 | can synthase activity (100%) | | | 00E+05 | 00E+00 | 00E+00 | | | | | | | | | | | |
| 307 | Variable 375 | EYSVDSDSLHQQLK | Clostridium botulinum | Bacteria | Firmicutes | | Clostridia | | Eubacteriales | | Clostridiaceae | Clostridium | | | Clostridium bot | | | | | | | GO:0080146(10 | L-cysteine desulfhydrase activity (10 | GO:0019450 (100%) | L-cysteine catabolic process | IPR005130(100 | 1.00E+00 | 1.00E+00 | 5.77E+04 | 0.48387 | 1.2597E-07 | 0.00034662 | 0.9 | NS | NS | NS | NS | 8.9157 | 0.0059254 | 0.00042176 |

| | | | | | | | | | | | | | | | | | | | | | | | | | | | | | | | | | | | | | | | | |
|---|---|---|---|---|---|---|---|---|---|---|---|---|---|---|---|---|---|---|---|---|---|---|---|---|---|---|---|---|---|---|---|---|---|---|---|---|---|---|---|---|
| | | | | | | | | | | | | | | | ulinum | | | | | | | 0%) | 0%) | | to pyruvate (100%) | %);IPR021144(100%) | | | | | | | | | | | | | | |
| 309 | Variable 377 | LEMAQLQLK | Desulfitobacterium | Bacteria | Firmicutes | | Clostridia | | Eubacteriales | | Peptococcaceae | Desulfitobacterium | | | | | | | | GO:0005887(88%);GO:0005886( | integral component of plasma membrane (88%);plasma | GO:0015103(88%) | inorganic anion transmembrane transporter activity (88%) | | | IPR003691(100%) | 1.00E+00 | 1.00E+00 | 7.19E+04 | NS | NS | NS | NS | NS | NS | NS | NS | 5.9679 | 0.005234 | 0.0068542 |

| | | | | | | | | | | | | | | | | | | | | | | | | | | | | | | | | | | | | | |
|---|---|---|---|---|---|---|---|---|---|---|---|---|---|---|---|---|---|---|---|---|---|---|---|---|---|---|---|---|---|---|---|---|---|---|---|---|---|
| | | | | | | | | | | | | | | | | | | | | 13%);GO:0016021(13%) | membrane(13%);integral component of membrane(13%) | | | | | | | | | | | | | | | | |
| 310 | Variable 378 | EAGVEVGEEEDR | Desulfofundulusth | Bacteria | Firmicutes | | Clostridia | | Eubacteriales | | Peptococcaceae | Desulfofundulus | | | Desulfofundulus | | | 2.5.1.7(100%) | UDP-N-acetylglucosamine1-carboxyvin | GO:0005737(100 | cytoplasm(100%) | GO:0008760(100 | UDP-N-acetylglucosamine1-carboxyvin | GO:0007049(100%);GO:000836 | cell cycle(100%);regulation of | IPR001986(100% | 1.00E+00 | 1.00E+00 | 4.67E+04 | NS | NS | NS | NS | NS | NS | NS | NS |

| | | | | | | | | | | | | | | | | | | | | | | | | | | | | | | | | | | | | | |
|---|---|---|---|---|---|---|---|---|---|---|---|---|---|---|---|---|---|---|---|---|---|---|---|---|---|---|---|---|---|---|---|---|---|---|---|---|---|
| | | | ermocisternus | | | | | | | | | | | | thermocisternus | | | | yltransferase(100%) | %) | | %) | yltransferase activity(100%) | 0(100%);GO:0009252(100%) | cell shape(100%); peptidoglycan biosynthetic process(100%) | );IPR013792(100%);IPR036968(100%) | | | | | | | | | | | |
| 311 | Variable 379 | ADLDLRPALK | Peptostreptococc | Bacteria | Firmicutes | | Clostridia | | Eubacteriales | | Peptostreptococcaceae | | | | | | | | | GO:0005840(1 | ribosome(100%) | GO:0003735(1 | structural constituent of ribosome | GO:0006412(100%) | translation(100%) | IPR000754(10 | 1.00E+00 | 1.00E+00 | 5.05E+04 | NS | NS | NS | NS | NS | NS | NS | NS |

| | | | | | | | | | | | | | | | | | | | | | | | | | | | | | | | | | | | | | | | | |
|---|---|---|---|---|---|---|---|---|---|---|---|---|---|---|---|---|---|---|---|---|---|---|---|---|---|---|---|---|---|---|---|---|---|---|---|---|---|---|---|---|
| | | | aceae | | | | | | | | | | | | | | | | | 00%) | | 00%) | (100%) | | | 0%);IPR014721(100%);IPR020568(100%) | | | | | | | | | | | | | | |
| 312 | Variable 3 | LTGFAALHTR | Actinomycetia | Bacteria | Actinobacte | | Actinomycetia | | | | | | | | | | | 2.7.-.-(100% | Transferring phosphorus-contai | | | GO:0016772 | transferase activity | transferring phosphorus- | GO:0000271(98%) | polysaccharid | 1.00E+00 | 1.00E+00 | 1.72E+05 | 51.254 | 0.000540 | 0.000714 | 0.9 | NS | NS | NS | NS | 5.7534 | 0.006895 | 0.0015 |

| | | | | | | | | | | | | | | | | | | | | | | | | | | | | | | | | | | | | | |
|---|---|---|---|---|---|---|---|---|---|---|---|---|---|---|---|---|---|---|---|---|---|---|---|---|---|---|---|---|---|---|---|---|---|---|---|---|---|
| | 8 | | | | ria | | | | | | | | | | | | | ) | ning groups (100%) | | | (98%); GO:0016740 (2%) | | containing groups (98%);transferase activity (2%) | | ebiosynthetic process (98%) | | | | | 1 | | | | | | |
| 313 | Variable 380 | EDLYQFFDR | Symbiobacterium thermophilum IAM | Bacteria | Firmicutes | | Clostridia | | Eubacteriales | | Symbiobacteriaceae | Symbiobacterium | | | Symbiobacterium thermophilu | | Symbiobacterium thermophilu | | | GO:0005737 (100%) | cytoplasm (100%) | GO:0016757 (50%) | glycosyltransferase activity (50%) | GO:0000105 (100%) | histidine biosynthetic process (100%) | IPR004516 (100%); IPR00451 | 1.00E+00 | 1.00E+00 | 3.56E+04 | NS | NS | NS | NS | NS | NS | NS | NS |

| | | | | | | | | | | | | | | | | | | | | | | | | | | | | | | | | | | | | | | | | |
|---|---|---|---|---|---|---|---|---|---|---|---|---|---|---|---|---|---|---|---|---|---|---|---|---|---|---|---|---|---|---|---|---|---|---|---|---|---|---|---|---|
| | | | 14863 | | | | | | | | | | | | m | | m IAM 14863 | | | | | | | | | 7(100%);IPR006195(100%) | | | | | | | | | | | | | | |
| 314 | Variable 381 | CGHEWEPEQGGK | Moorellathermoacetica | Bacteria | Firmicutes | | Clostridia | | Thermoanaerobacterales | | Thermoanaerobacteraceae | Moorella | | | Moorellathermoacetica | | | | | | | GO:0003677(100%) | DNA binding(100%) | | | IPR002852(100%) | 1.00E+00 | 1.00E+00 | 1.10E+05 | 0.07923 | 5.1747E-15 | 0.002287 | 0.9 | NS | NS | NS | NS | 47.82 | 1.4946E-11 | 0.016388 |
| 315 | Varia | DEGLEEGESND | Thermo | Bacte | Firmic | | Clostridi | | Thermoa | | Thermoa | Thermo | | | | | | | | | | GO:00 | RNA binding | GO:000635 | DNA-tem | termin | 1.00E | 1.00E | 5.39E | NS | NS | NS | NS | NS | NS | NS | NS | | | |

| | | | | | | | | | | | | | | | | | | | | | | | | | | | | | | | | | | | | | |
|---|---|---|---|---|---|---|---|---|---|---|---|---|---|---|---|---|---|---|---|---|---|---|---|---|---|---|---|---|---|---|---|---|---|---|---|---|---|
| | ble 382 | N | anaerobacter | ria | utes | | a | | naerobacterales | | naerobacteraceae | anaerobacter | | | | | | | | | | 03723(100%) | (100%) | 3(100%);GO:0006355(100%);GO:0031564(100%) | plated transcription | ation(100%);regulation of transcription | +00 | +00 | +04 | | | | | | | | |
| 316 | Variable 383 | TYVSTLLYK | Fusobacterium | Bacteria | Fusobacteria | | Fusobacteriia | | Fusobacteriales | | Fusobacteriaceae | Fusobacterium | | | | | | 6.3.3.3(100%) | Dethiobiotin synthase (100%) | GO:0005737(92%);GO | cytoplasm(92%);cytosol(2%) | GO:0002887(100%);G | magnesium ion binding(100%);dethiobiotin syntha | GO:0009102(100%) | biotin biosynthetic process(100%) | IPR004472(100%);IP | 1.34E+05 | 1.00E+00 | 1.00E+00 | NS | NS | NS | NS | NS | NS | NS | NS |

| | | | | | | | | | | | | | | | | | | | | | | | | | | | | | | | | | | | | | |
|---|---|---|---|---|---|---|---|---|---|---|---|---|---|---|---|---|---|---|---|---|---|---|---|---|---|---|---|---|---|---|---|---|---|---|---|---|---|
| | | | | | | | | | | | | | | | | | | | | :0005829(2%) | | O:0004141(100%);GO:0005524(100%) | se activity(100%);ATP binding(100%) | | | R027417(100%) | | | | | | | | | | | |
| 318 | Variable 385 | KPLRPTDEEVQANPR | Rhodopirellula | Bacteria | Planctomycet | | Planctomycetia | | Pirellulales | | Pirellulaceae | Rhodopirellula | | | | | | 2.1.1.199(10 | 16S rRNA(cytosine(1402)-N(4)) | GO:0005737(8 | cytoplasm(89%) | GO:0071424(8 | rRNA(cytosine-N4-)-methyltra | GO:0070475(89%);GO:00 | rRNA base methylation(8 | IPR002903(10 | 1.00E+00 | 7.77E+04 | 1.00E+00 | NS | NS | NS | NS | NS | NS | NS | NS |

| | | | | | | | | | | | | | | | | | | | | | | | | | | | | | | | | | | | | | |
|---|---|---|---|---|---|---|---|---|---|---|---|---|---|---|---|---|---|---|---|---|---|---|---|---|---|---|---|---|---|---|---|---|---|---|---|---|---|
| | | | | | es | | | | | | | | | | | | | 0%) | -methyltransferase (100%) | 9%) | | 9%);GO:0008168(11%) | nsferase activity (89%);methyltransferase activity (11%) | 06364(11%);GO:0032259(11%) | 9%);rRNA processing(11%);methylation(11%) | 0%);IPR023397(100%);IPR029063(100%) | | | | | | | | | | | |
| 319 | Variable 3 | GHHCSDK | Rhodopirellula | Bacteria | Planctomyc | | Planctomycetia | | Pirellulales | | Pirellulaceae | Rhodopirellu | | | | | | 4.2.1.33(1 | 3-isopropylmalate dehydratase | | | GO:0003861 | 3-isopropylmalate dehydra | 4 sulfur cluster binding (1 | GO:0009082(100%); | branched-ch | 1.00E+00 | 6.05E+04 | 1.00E+00 | NS | NS | NS | NS | NS | NS | NS | NS |

| | | | | | | | | | | | | | | | | | | | | | | | | | | |
|---|---|---|---|---|---|---|---|---|---|---|---|---|---|---|---|---|---|---|---|---|---|---|---|---|---|---|
| | 86 | | | | etes | | | | | | | la | | | | | | 00%) | (100%) | | | (100%);GO:0046872(100%);GO:0051539(100%) | tase activity(100%);metal ion binding(100%);4iron | 00%) | GO:0009098(100%) | ainaminoacidbiosynthetic process(100%);leucine biosynthetic p |

| | | | | | | | | | | | | | | | | | | | | | | | | | | | | | | | | | | | | | |
|---|---|---|---|---|---|---|---|---|---|---|---|---|---|---|---|---|---|---|---|---|---|---|---|---|---|---|---|---|---|---|---|---|---|---|---|---|---|
| | | | | | | | | | | | | | | | | | | | | | | | | | | | rocess(100%) | | | | | | | | | | |
| 322 | Variable 389 | LLLLYGGSVK | Proteobacteria | Bacteria | Proteobacteria | | | | | | | | | | | | | 5.3.1.1(100%) | Triose-phosphate isomerase(100%) | GO:0005737(94%) | cytoplasm(94%) | GO:0004807(100%) | triose-phosphate isomerase activity(100%) | GO:0006096(100%);GO:0006094(94%) | glycolytic process(100%); gluconeogenesis(94%) | IPR000652(100%);IPR013785(100%);IP | 1.00E+00 | 1.00E+00 | 5.37E+04 | NS | NS | NS | NS | NS | NS | NS | NS |

| | | | | | | | | | | | | | | | | | | | | | | | | | | | | | | | | | | | | | | | | |
|---|---|---|---|---|---|---|---|---|---|---|---|---|---|---|---|---|---|---|---|---|---|---|---|---|---|---|---|---|---|---|---|---|---|---|---|---|---|---|---|---|
| | | | | | | | | | | | | | | | | | | | | | | | | | | R0208861(100%) | | | | | | | | | | | | | | |
| 323 | Variable 39 | FLGGYLTAR | Actinomycetia | Bacteria | Actinobacteria | | Actinomycetia | | | | | | | | | | | 5.1.3.2(100%) | UDP-glucose 4-epimerase (100%) | GO:0005576(0%);GO:0005829(0%);GO | extracellular region (0%);cytosol (0%);plasma membrane (0%) | GO:0003824(91%);GO:0003978(9%) | catalytic activity (91%);UDP-glucose 4-epimerase activity (9%) | | | IPR036291(100%);IPR001509(99%);I | 1.00E+00 | 1.00E+00 | 4.18E+04 | 8.4731 | 1.8995E-06 | 0.00014663 | 0.9 | NS | NS | NS | NS | 9.876 | 0.00030518 | 0.00015459 |

| | | | | | | | | | | | | | | | | | | | | | | | | | | | | | | | | | | | | | |
|---|---|---|---|---|---|---|---|---|---|---|---|---|---|---|---|---|---|---|---|---|---|---|---|---|---|---|---|---|---|---|---|---|---|---|---|---|---|
| | | | | | | | | | | | | | | | | | | | | :0005886(0%) | | | | | | PR0088886(1%) | | | | | | | | | | | |
| 324 | Variable 390 | GGENPFEGK | Proteobacteria | Bacteria | Proteobacteria | | | | | | | | | | | | | | | | | GO:0005525(100%) | GTP binding(100%) | GO:0042254(99%) | ribosome biogenesis(99%) | IPR015946(100%);IPR027417(100%);I | 1.00E+00 | 1.00E+00 | 5.66E+04 | NS | NS | NS | NS | NS | NS | NS | NS |

| | | | | | | | | | | | | | | | | | | | | | | | | | | | | | | | | | | | | | | | | |
|---|---|---|---|---|---|---|---|---|---|---|---|---|---|---|---|---|---|---|---|---|---|---|---|---|---|---|---|---|---|---|---|---|---|---|---|---|---|---|---|---|
| | | | | | | | | | | | | | | | | | | | | | | | | | | PR032859(100%) | | | | | | | | | | | | | | |
| 325 | Variable 391 | QRPELAWPADLYLEGSDQHR | Proteobacteria | Bacteria | Proteobacteria | | | | | | | | | | | | | 6.1.1.5(100%) | Isoleucine--tRNA ligase(100%) | GO:0005737(96%) | cytoplasm(96%) | GO:0004822(100%);GO:0005524(10 | isoleucine-tRNA ligase activity(100%);ATP binding(100%);tRNA binding(97%) | GO:0006428(99%);GO:0006418(1%) | isoleucyl-tRNA aminoacylation(99%);tRNA aminoacylation | IPR002300(100%);IPR014729(100% | 1.00E+00 | 1.00E+00 | 1.28E+05 | 10.969 | 0.0018932 | 0.0042757 | 0.9 | NS | NS | NS | NS | 4.664 | 0.014818 | 0.0017619 |

| | | | | | | | | | | | | | | | | | | | | | | | | | | | | | | | | | | | | | |
|---|---|---|---|---|---|---|---|---|---|---|---|---|---|---|---|---|---|---|---|---|---|---|---|---|---|---|---|---|---|---|---|---|---|---|---|---|---|
| | | | | | | | | | | | | | | | | | | | | | | 0%);GO:0000049(97%) | | | for protein translation (1%) | );IPR002301(99%) | | | | | | | | | | | |
| 326 | Variable 392 | TPVDLDGLR | Proteobacteria | Bacteria | Proteobacteria | | | | | | | | | | | | | 3.1.-.-(100%) | Acting on ester bonds (100%) | | | GO:0004519(100%);GO:000367 | endonuclease activity (100%);DNA binding (13%) | GO:0043571 (13%);GO:0051607 (13%) | maintenance of CRISPR repeat elements (13%);de | IPR010152(100%) | 1.00E+00 | 9.93E+04 | 1.00E+00 | NS | NS | NS | NS | NS | NS | NS | NS |

| | | | | | | | | | | | | | | | | | | | | | | | | | | | | | | | | | | | | | |
|---|---|---|---|---|---|---|---|---|---|---|---|---|---|---|---|---|---|---|---|---|---|---|---|---|---|---|---|---|---|---|---|---|---|---|---|---|---|
| | | | | | | | | | | | | | | | | | | | | | | 7 (13%) | | | fense response to virus (13%) | | | | | | | | | | | | |
| 327 | Variable 393 | VSQLDEAR | Proteobacteria | Bacteria | Proteobacteria | | | | | | | | | | | | | 1.17.7.3 (100%) | (E)-4-hydroxy-3-methylbut-2-enyl-diphosphate synthase (flavodoxin) (100%) | GO:0005737 (6%) | cytoplasm (6%) | GO:0005506 (88%);GO:0046429 (88 | iron ion binding (88%);4-hydroxy-3-methylbut-2-en-1-yl diphosphate synthase | 4 sulfur cluster binding (88%) | GO:0016114 (88%);GO:0019288 (82%);GO:0006260 (6% | terpenoid biosynthetic process (88%); iso | 1.00E+00 | 7.69E+04 | 1.00E+00 | NS | NS | NS | NS | NS | NS | NS | NS |

| | | | | | | | | | | | | | | | | | | | | | | | | | | | | | | | | | | | | | |
|---|---|---|---|---|---|---|---|---|---|---|---|---|---|---|---|---|---|---|---|---|---|---|---|---|---|---|---|---|---|---|---|---|---|---|---|---|---|
| | | | | | | | | | | | | | | | | | | | | | | %);GO:0051539(88%) | activity(88%);4iron | | ) | pentenyldiphosphate biosynthetic process | | | | | | | | | | | |
| 328 | Variable 394 | CAALVVHAK | Proteobacteria | Bacteria | Proteobacteria | | | | | | | | | | | | | 5.1.1.1(100%) | Alanine racemase(100%) | | | GO:0003677(100 | DNA binding(100%);alanine racemase | GO:0006355(100%);GO:000601 | regulation of transcription | DNA-templated(10 | 1.00E+00 | 6.34E+05 | 2.22E+06 | 2.9138 | 0.02786 | 0.00042326 | 1 | NS | NS | NS | NS |

| | | | | | | | | | | | | | | | | | | | | | | | | | | | | | | | | | | | | | |
|---|---|---|---|---|---|---|---|---|---|---|---|---|---|---|---|---|---|---|---|---|---|---|---|---|---|---|---|---|---|---|---|---|---|---|---|---|---|
| | | | | | | | | | | | | | | | | | | | | | | %);GO:0008784(0%);GO:0000976(0%) | activity (0%);transcription cis-regulatory region binding (0%) | 2 (0%);GO:0006351 (0%) | | 0%);galactose metabolic process (0%);transcription | | | | | | | | | | | |
| 329 | Variable | VSVPAAFR | Proteobact | Bacteria | Proteoba | | | | | | | | | | | | | | | GO:0057 | cytoplasm (100% | GO:0037 | DNA-binding transc | GO:0051301 (28 | cell division (28 | IPR00344 | 1.00E+00 | 2.42E+05 | 1.00E+00 | NS | NS | NS | NS | NS | NS | NS | NS |

| | | | | | | | | | | | | | | | | | | | | | | | | | | | | | | | | | | | | | |
|---|---|---|---|---|---|---|---|---|---|---|---|---|---|---|---|---|---|---|---|---|---|---|---|---|---|---|---|---|---|---|---|---|---|---|---|---|---|
| | 395 | | eria | | cteria | | | | | | | | | | | | | | | 37(100%);GO:0009295(96%) | );nucleoid(96%) | 00(100%);GO:0003677(99%) | ription factor activity(100%);DNA binding(99%) | %) | %) | 4(100%);IPR035642(100%);IPR035644(100%) | | | | | | | | | | | |
| 330 | Variab | LTALYAQK | Proteob | Bacter | Proteo | | | | | | | | | | | | | 2.7.4. | (d)CMP kinase (10 | GO:000 | cytoplasm (9 | GO:000 | ATP binding (98 | GO:0006220 | pyrimidine nu | IPR003 | 1.00E+ | 1.47E+ | 1.00E+ | NS | NS | NS | NS | NS | NS | NS | NS |

| | | | | | | | | | | | | | | | | | | | | | | | | | | | | | |
|---|---|---|---|---|---|---|---|---|---|---|---|---|---|---|---|---|---|---|---|---|---|---|---|---|---|---|---|---|---|
| | le 396 | | acteria | ia | bacteria | | | | | | | | | | | | | 25(100%) | 0%) | 5737(97%);GO:0005829(2%) | 7%);cytosol(2%) | 5524(98%);GO:0036430(95%);GO:0036431(95%) | %);CMP kinase activity(95%);dCMP kinase activity(95%) | (98%);GO:0006281(2%);GO:0015949(2%) | cleotide metabolic process(98%);DNA repair(2%);nucleobase-containing small molec | 136(98%);IPR011994(98%);IPR027417(98%) | 00 | 05 | 00 |

| | | | | | | | | | | | | | | | | | | | | | | | | | | | | | | | | | | | | | |
|---|---|---|---|---|---|---|---|---|---|---|---|---|---|---|---|---|---|---|---|---|---|---|---|---|---|---|---|---|---|---|---|---|---|---|---|---|---|
| | | | | | | | | | | | | | | | | | | | | | | | | | ule interconversion (2%) | | | | | | | | | | | | |
| 331 | Variable 397 | AAEQVAATR | Proteobacteria | Bacteria | Proteobacteria | | | | | | | | | | | | | 3.1.26.3(100%) | Ribonuclease III (100%) | GO:0005737(98%) | cytoplasm(98%) | GO:0004525(100%);GO:0019843(97%); | ribonuclease III activity(100%);rRNA binding(97%);metal ion binding(97%) | GO:0006397(99%);GO:0006364(98%);GO:0016075(98%) | mRNA processing(99%);rRNA processing(98%);rRNA catabo | IPR000999(99%);IPR011907(99%);IPR | 1.00E+00 | 1.01E+05 | 1.00E+00 | NS | NS | NS | NS | NS | NS | NS | NS |

| | | | | | | | | | | | | | | | | | | | | | | | | | | | | | | | | | | | | | |
|---|---|---|---|---|---|---|---|---|---|---|---|---|---|---|---|---|---|---|---|---|---|---|---|---|---|---|---|---|---|---|---|---|---|---|---|---|---|
| | | | | | | | | | | | | | | | | | | | | | | GO:0046872(97%) | | | lic process(98%) | 0147200(99%) | | | | | | | | | | | |
| 332 | Variable 398 | TAQSGLQEK | Alphaproteobacteria | Bacteria | Proteobacteria | | Alphaproteobacteria | | | | | | | | | | | 3.6.1.31(100%) | Phosphoribosyl-ATP diphosphatase (100%) | GO:0005737(97%) | cytoplasm(97%) | GO:0000463 6(100%);GO:0005524(9 | phosphoribosyl-ATP diphosphatase activity(100%);ATP binding(97%) | GO:0000105(100%) | histidine biosynthetic process(100%) | IPR008179(100%);IPR021130(100 | 1.00E+00 | 1.00E+00 | 2.55E+05 | 2.5216 | 0.0039216 | 0.001016 | 0.9 | NS | NS | NS | NS |

| | | | | | | | | | | | | | | | | | | | | | | | | | | | | | | | | | | | | | | | | |
|---|---|---|---|---|---|---|---|---|---|---|---|---|---|---|---|---|---|---|---|---|---|---|---|---|---|---|---|---|---|---|---|---|---|---|---|---|---|---|---|---|
| | | | | | | | | | | | | | | | | | | | | | | 7%) | | | | %) | | | | | | | | | | | | | | |
| 333 | Variable 399 | FVASGTVTSDGDMDLASR | Alphaproteobacteria | Bacteria | Proteobacteria | | Alphaproteobacteria | | | | | | | | | | | 3.1.22.4(100%) | Crossover junction endodeoxyribonuclease (100%) | | | GO:0000287(100%);GO:0008821(100%);GO:00036 | magnesium ion binding(100%);crossover junction endodeoxyribonuclease activity(100%);nucleic acid binding(97 | GO:0006281(100%);GO:0006310(100%) | DNA repair(100%);DNA recombination(100%) | IPR002176(100%);IPR012337(100%);IPR036397( | 1.00E+00 | 1.00E+00 | 1.04E+05 | 0.5022 | 6.3212E-08 | 0.00092037 | 0.9 | NS | NS | NS | NS | 10.095 | 0.002631 | 0.0013262 |

| | | | | | | | | | | | | | | | | | | | | | | | | | | | | | | | | | | | | | |
|---|---|---|---|---|---|---|---|---|---|---|---|---|---|---|---|---|---|---|---|---|---|---|---|---|---|---|---|---|---|---|---|---|---|---|---|---|---|
| | | | | | | | | | | | | | | | | | | | | | | 76(97%) | %) | | | 100%) | | | | | | | | | | | |
| 335 | Variable 40 | DLVDGTPKPVLEK | Actinomycetia | Bacteria | Actinobacteria | | Actinomycetia | | | | | | | | | | | | | GO:0005840(100%) | ribosome(100%) | GO:0003735(98%) | structural constituent of ribosome(98%) | GO:0006412(100%) | translation(100%) | IPR013823(100%);IPR014719(100%);IPR000 | 1.00E+00 | 1.33E+05 | 1.00E+00 | NS | NS | NS | NS | NS | NS | NS | NS |

| | | | | | | | | | | | | | | | | | | | | | | | | | | | | | | | | | | | | | |
|---|---|---|---|---|---|---|---|---|---|---|---|---|---|---|---|---|---|---|---|---|---|---|---|---|---|---|---|---|---|---|---|---|---|---|---|---|---|
| | | | | | | | | | | | | | | | | | | | | | | | | | | 206(100%) | | | | | | | | | | | |
| 336 | Variable 400 | MSVDLATVK | Alphaproteobacteria | Bacteria | Proteobacteria | | Alphaproteobacteria | | | | | | | | | | | 6.3.5.-(100%);6.3.5.6(0%);6.3.5.7(0%) | Carbon--nitrogen ligases with glutamine as amido-N-donor(100%);Asparaginyl-tRNA synthase(gl | | | GO:0050567(100%);GO:0050566(98%);GO: | glutaminyl-tRNA synthase(glutamine-hydrolyzing) activity(100%);asparaginyl-tRNA syn | GO:0006412(100%);GO:0006450(98%) | translation(100%);regulation of translational fidelity(98%) | IPR003837(100%);IPR036113(100%) | 1.00E+00 | 1.15E+05 | 1.00E+00 | NS | NS | NS | NS | NS | NS | NS | NS |

| | | | | | | | | | | | | | | | | | | | | | | | | | | | | | | | | | | | | | |
|---|---|---|---|---|---|---|---|---|---|---|---|---|---|---|---|---|---|---|---|---|---|---|---|---|---|---|---|---|---|---|---|---|---|---|---|---|---|
| | | | | | | | | | | | | | | | | | | | utamine-hydrolyzing)(0%);Glutaminyl-tRNA synthase(glutamine-hydrolyzing)(0%) | | | 0005524(97%) | thase(glutamine-hydrolyzing) activity(98%);ATP binding(97%) | | | | | | | | | | | | | | |
| 340 | Variable 404 | DAPVEAEEACATAR | Phylobacteriacea | Bacteria | Proteobacteri | | Alphaproteobacteri | | Hyphomicrobiales | | Phyllobacteriaceae | | | | | | | 5.4.99.25(10 | tRNA pseudouridine(55) synthase(10 | | | GO:0003723(9 | RNA binding(98%);tRNA pseud | GO:0031119(94%);GO:00 | tRNA pseudouridine synthe | IPR015240(99 | 1.00E+00 | 1.00E+00 | 2.94E+05 | 9.0159 | 0.016491 | 0 | 0.9 | NS | NS | NS | NS |

| | | | | | | | | | | | | | | | | | | | | | | | | | | | | | | | | | | | | | | | | |
|---|---|---|---|---|---|---|---|---|---|---|---|---|---|---|---|---|---|---|---|---|---|---|---|---|---|---|---|---|---|---|---|---|---|---|---|---|---|---|---|---|
| | | | e | | a | | a | | | | | | | | | | | 0%) | 0%) | | | 8%);GO:01006029(93%);GO:0009982(7%) | ouridine synthase activity (93%);pseudouridine synthase activity (7%) | 01522(6%);GO:0006396(1%) | sis(94%);pseudouridine synthesis(6%);RNA processing(1%) | %);IPR020103(96%);IPR032819(96%) | | | | | | | | | | | | | | |
| 343 | Variable 40 | DPAAMAGAFAK | Rhizobiaceae | Bacteria | Proteobacte | | Alphaproteobacte | | Hyphomicrobiales | | Rhizobiaceae | | | | | | | 2.8.1.10(10 | Thiazole synthase (100%) | GO:0005737( | cytoplasm (100%) | GO:1990107( | thiazole synthase activity (75%) | GO:0009229(75%);GO: | thiamine diphosphate bi | IPR008867(1 | 1.00E+00 | 1.00E+00 | 1.49E+05 | 1.5327 | 0.0001896 | 0.00073188 | 0.9 | NS | NS | NS | NS | 6.4655 | 0.0035649 | 0.0042979 |

| | | | | | | | | | | | | | | | | | | | | | | | | | | | | | | | | | | | | | |
|---|---|---|---|---|---|---|---|---|---|---|---|---|---|---|---|---|---|---|---|---|---|---|---|---|---|---|---|---|---|---|---|---|---|---|---|---|---|
| | 7 | | | | ria | | ria | | | | | | | | | | | 0%) | | 100%) | | 75%);GO:0016783(25%) | ;sulfurtransferase activity (25%) | 0009228(25%) | osynthetic process (75%);thiamine biosynthetic process (25%) | 00%);IPR013785(100%);IPR033983(100%) | | | | | | | | | | | |
| 345 | Variable | VTAGHCSLAR | Rhizobiacea | Bacteria | Proteobac | | Alphaproteob | | Hyphomicrobi | | Rhizobiaceae | | | | | | | 3.5.4.1(9 | Cytosine deaminase (94%); | | | GO:001681 | hydrolase activity | acting on carbon-nitro | | | 5.37E+05 | 1.00E+00 | 1.00E+00 | NS | NS | NS | NS | NS | NS | NS | NS |

| | | | | | | | | | | | | | | | | | | | | | | | | |
|---|---|---|---|---|---|---|---|---|---|---|---|---|---|---|---|---|---|---|---|---|---|---|---|---|
| | 409 | | e | | teria | | acteria | | ales | | | | | | | | | 4%);3.5.4.11(6%) | Pterin deaminase (6%) | | | 0(57%);GO:0004131(41%);GO:0102480(41%) | | gen (but not peptide) bonds (57%);cytosine deaminase activity (41%);5-fluorocytosine deaminase activity (41 |

| | | | | | | | | | | | | | | | | | | | | | | | | | | | | | | | | | | | | | |
|---|---|---|---|---|---|---|---|---|---|---|---|---|---|---|---|---|---|---|---|---|---|---|---|---|---|---|---|---|---|---|---|---|---|---|---|---|---|
| | | | | | | | | | | | | | | | | | | | | | | | | %) | | | | | | | | | | | | | |
| 347 | Variable 410 | AGVALGGQAK | Agrobacterium | Bacteria | Proteobacteria | | Alphaproteobacteria | | Hyphomicrobiales | | Rhizobiaceae | Agrobacterium | | | | | | 3.2.1.21(100%) | Beta-glucosidase(100%) | GO:0005737(25%) | cytoplasm(25%) | GO:0008422(100%);GO:0102483(100%) | beta-glucosidase activity(100%);scopolin beta-glucosidase activity(100%) | GO:0005975(100%) | carbohydrate metabolic process(100%) | IPR001764(100%);IPR002772(100%);IPR008979(10 | 1.00E+00 | 1.00E+00 | 1.80E+05 | 10.12 | 0.002349 | 0.00016275 | 0.9 | NS | NS | NS | NS |

| | | | | | | | | | | | | | | | | | | | | | | | | | | | | | | | | | | | | | |
|---|---|---|---|---|---|---|---|---|---|---|---|---|---|---|---|---|---|---|---|---|---|---|---|---|---|---|---|---|---|---|---|---|---|---|---|---|---|
| | | | | | | | | | | | | | | | | | | | | | | | | | | 0%) | | | | | | | | | | | |
| 348 | Variable 411 | SFCFTDWTLNLR | Agrobacterium | Bacteria | Proteobacteria | | Alphaproteobacteria | | Hyphomicrobiales | | Rhizobiaceae | Agrobacterium | | | | | | | | GO:0005737(14%) | cytoplasm(14%) | GO:0003677(100%);GO:0004519(7%) | DNA binding(100%);endonuclease activity(7%) | GO:0000160(100%);GO:0006355(100%) | phosphorelay signal transduction system(100%);regulation of transcription | DNA-templated(100%) | 1.00E+00 | 1.00E+00 | 3.13E+04 | NS | NS | NS | NS | NS | NS | NS | NS |
| 349 | Variable | VDHEVLV | Agrobacter | Bacteria | Proteoba | | Alphaproteo | | Hyphomicro | | Rhizobiaceae | Agrobacter | | | | | | 6.3.4.21 | Nicotinate phosphoribo | | | GO:0045 | nicotinate-nucleotide | GO:0009435(100 | NAD biosyntheti | IPR006400 | 1.00E+00 | 1.00E+00 | 3.00E+04 | NS | NS | NS | NS | NS | NS | NS | NS |

| | | | | | | | | | | | | | | | | | | | | | | | | | | | | | | | | | | | | | | | | |
|---|---|---|---|---|---|---|---|---|---|---|---|---|---|---|---|---|---|---|---|---|---|---|---|---|---|---|---|---|---|---|---|---|---|---|---|---|---|---|---|---|
| | 412 | | ium | | cteria | | bacteria | | biales | | | ium | | | | | | (100%) | syltransferase (100%) | | | 14(100%);GO:0004516(100%) | diphosphorylase (carboxylating) activity (100%);nicotinate phosphoribosyltransferase activity (100%) | %) | cprocess (100%) | 6(100%);IPR007229(100%);IPR036068(100%) | | | | | | | | | | | | | | |
| 350 | Variab | LMDPESVVAMAAT | Agroba | Bacter | Proteo | | Alphaprot | | Hyphom | | Rhizobiaceae | Agroba | | | | | | 2.8.1. | Biotin synthase (10 | | | GO:000 | biotin synthase act | 2 sulfur cluster | GO:0009102 | biotin bios | 1.00E+ | 1.00E+ | 6.73E+ | 13.451 | 7.215 | 0.0040 | 0.9 | NS | NS | NS | NS | 1.54 | 0.001 | 0.028 |

| | | | | | | | | | | | | | | | | | | | | | | | | | | | | | | | | | | | | | | | |
|---|---|---|---|---|---|---|---|---|---|---|---|---|---|---|---|---|---|---|---|---|---|---|---|---|---|---|---|---|---|---|---|---|---|---|---|---|---|---|---|
| | le 413 | AR | cterium | ia | bacteria | | eobacteria | | icrobiales | | e | cterium | | | | | | 6(100%) | 0%) | | | 4076(100%);GO:0005506(100%);GO:0051537(100%) | ivity(100%);iron ion binding(100%);2iron | binding(100%) | (100%) | ynthetic process(100%) | 00 | 00 | 04 | | E-07 | 327 | | | | | | 0138 | 358 |
| 3 | V | DLS | A | B | P | | A | | H | | R | A | | | | | | 6 | Thr | G | cy | G | tR | G | th | I | 1 | 3 | 1 | N | N | N | N | N | N | N | N | | |

| | | | | | | | | | | | | | | | | | | | | | | | | | | | | | | | | | | | | | |
|---|---|---|---|---|---|---|---|---|---|---|---|---|---|---|---|---|---|---|---|---|---|---|---|---|---|---|---|---|---|---|---|---|---|---|---|---|---|
| 51 | ariable 414 | EPVADGK | grobacterium | acteria | roteobacteria | | lphaproteobacteria | | yphomicrobiales | | hizobiaceae | grobacterium | | | | | | .1.1.3(100%) | eonine--tRNA ligase(100%) | O:0005737(100%) | toplasm(100%) | O:0000049(100%);GO:0004829(100%);GO:0005524(1 | NA binding(100%);threonine-tRNA ligase activity(100%);ATP binding(100%) | O:0006435(100%) | reonyl-tRNA aminoacylation(100%) | PR002314(100%);IPR002320(100%);IPR004095(100% | .00E+00 | .51E+05 | .00E+00 | S | S | S | S | S | S | S | S |

| | | | | | | | | | | | | | | | | | | | | | | | | | | | | | | | | | | | | | | | | |
|---|---|---|---|---|---|---|---|---|---|---|---|---|---|---|---|---|---|---|---|---|---|---|---|---|---|---|---|---|---|---|---|---|---|---|---|---|---|---|---|---|
| | | | | | | | | | | | | | | | | | | | | | | 00%) | | | | ) | | | | | | | | | | | | | | |
| 352 | Variable 415 | AMSGFLAEK | Agrobacterium tumefaciens complex | Bacteria | Proteobacteria | | Alphaproteobacteria | | Hyphomicrobiales | | Rhizobiaceae | Agrobacterium | Agrobacterium tumefaciens complex | | | | | | | | | GO:0003677(100%);GO:0003700(100%) | DNA binding(100%);DNA-binding transcription factor activity(100%) | GO:0019619(100%);GO:0045893(100%) | 3 | 4-dihydroxybenzoate catabolic process(100%);positi | 1.00E+00 | 1.00E+00 | 2.29E+04 | 2.3121 | 0.00001813 | 0.0002249 | 0.9 | NS | NS | NS | NS | 7.5124 | 0.0016252 | 0.00088076 |

| | | | | | | | | | | | | | | | | | | | | | | | | | | | | | | | | | | | | | | | | |
|---|---|---|---|---|---|---|---|---|---|---|---|---|---|---|---|---|---|---|---|---|---|---|---|---|---|---|---|---|---|---|---|---|---|---|---|---|---|---|---|---|
| | | | | | | | | | | | | | | | | | | | | | | | | | | veregulation of transcription | | | | | | | | | | | | | | |
| 353 | Variable 416 | AFQPVTDHDGR | Agrobacterium tumefaciens complex | Bacteria | Proteobacteria | | Alphaproteobacteria | | Hyphomicrobiales | | Rhizobiaceae | Agrobacterium | Agrobacterium tumefaciens | | | | | 1.-.-.-(100%) | Oxidoreductases (100%) | | | GO:0016491(100%) | oxidoreductase activity (100%) | GO:0019469 (17%) | octopine catabolic process (17%) | IPR007419(100%); IPR017224 | 1.00E+00 | 1.00E+00 | 6.52E+04 | 8.3609 | 0.0013978 | 0.0014706 | 0.9 | NS | NS | NS | NS | 5.0352 | 0.01096 | 0.0012941 |

| | | | | | | | | | | | | | | | | | | | | | | | | | | | | | | | | | | | | | |
|---|---|---|---|---|---|---|---|---|---|---|---|---|---|---|---|---|---|---|---|---|---|---|---|---|---|---|---|---|---|---|---|---|---|---|---|---|---|
| | | | | | | | | | | | | | complex | | | | | | | | | | | | | (100%);IPR023753(100%) | | | | | | | | | | | |
| 354 | Variable 417 | HDEEDYTHK | Agrobacterium tumefaciens | Bacteria | Proteobacteria | | Alphaproteobacteria | | Hyphomicrobiales | | Rhizobiaceae | Agrobacterium | Agrobacterium tumefacie | | Agrobacterium tumefaciens | | | | | GO:0005737(100%) | cytoplasm(100%) | GO:0003746(100%);GO:000 | translation elongation factor activity (100%);GTPase activity (10 | | | IPR000640(100%);IPR007 | 1.00E+00 | 1.00E+00 | 1.20E+05 | NS | NS | NS | NS | NS | NS | NS | NS |

| | | | | | | | | | | | | | | | | | | | | | | | | | | | | | | | | | | | | | |
|---|---|---|---|---|---|---|---|---|---|---|---|---|---|---|---|---|---|---|---|---|---|---|---|---|---|---|---|---|---|---|---|---|---|---|---|---|---|
| | | | | | | | | | | | | | nscomplex | | | | | | | | | 3924(100%);GO:0005525(100%) | 0%);GTP binding(100%) | | | 95(100%);IPR004161(100%) | | | | | | | | | | | |
| 356 | Variable 419 | AVLAMYGAQPANQAAFPR | Acidiphilium | Bacteria | Proteobacteria | | Alphaproteobacteria | | Rhodospirillales | | Acetobacteraceae | Acidiphilium | | | | | | 3.4.24.-(100%) | Metalloendopeptidases(100%) | GO:0005886(100%);G | plasma membrane(100%);integral | GO:0004222(100%);G | metalloendopeptidase activity(100%);zinc ion bin | | | IPR001915(100%);IP | 1.00E+00 | 1.00E+00 | 5.65E+04 | NS | NS | NS | NS | NS | NS | NS | NS |

| | | | | | | | | | | | | | | | | | | | | | | | | | | | | | | | | | | | | | |
|---|---|---|---|---|---|---|---|---|---|---|---|---|---|---|---|---|---|---|---|---|---|---|---|---|---|---|---|---|---|---|---|---|---|---|---|---|---|
| | | | | | | | | | | | | | | | | | | | | O:0016021(100%) | component of membrane (100%) | O:0008270(100%) | ding (100%) | | | R022919(100%) | | | | | | | | | | | |
| 357 | Variable 42 | DQQAAADR | Actinomycetia | Bacteria | Actinobacteria | | Actinomycetia | | | | | | | | | | | | | GO:0005737(96%);GO:0005618(1 | cytoplasm(96%);cell wall(1%);integral component of mem | GO:0000774(99%);GO:0042803(9 | adenyl-nucleotide exchange factor activity(99%);protein homodimerizati | GO:0006457(99%);GO:0009267(1%) | protein folding(99%);cellular response to starvation(1%) | IPR000740(99%);IPR013805(99% | 1.00E+00 | 5.49E+05 | 1.00E+00 | NS | NS | NS | NS | NS | NS | NS | NS |

| | | | | | | | | | | | | | | | | | | | | | | | | | | | | | | | | | | | | | |
|---|---|---|---|---|---|---|---|---|---|---|---|---|---|---|---|---|---|---|---|---|---|---|---|---|---|---|---|---|---|---|---|---|---|---|---|---|---|
| | | | | | | | | | | | | | | | | | | | | %);GO:0016021(1%) | brane(1%) | 9%);GO:0051087(99%) | on activity(99%);chaperone binding(99%) | | | );IPR009012(99%) | | | | | | | | | | | |
| 358 | Variable 420 | ELGFPAANFSEK | Neorickettsia | Bacteria | Proteobacteria | | Alphaproteobacteria | | Rickettsiales | | Anaplasmataceae | Neorickettsia | | | | | | | | | | GO:0016793(100%) | triphosphoric monoester hydrolase activity(100%) | | | IPR003607(100%);IPR006261( | 1.00E+00 | 1.00E+00 | 1.09E+05 | 4.345 | 0.017563 | 0.00011324 | 0.9 | NS | NS | NS | NS |

| | | | | | | | | | | | | | | | | | | | | | | | | | | | | | | | | | | | | | |
|---|---|---|---|---|---|---|---|---|---|---|---|---|---|---|---|---|---|---|---|---|---|---|---|---|---|---|---|---|---|---|---|---|---|---|---|---|---|
| | | | | | | | | | | | | | | | | | | | | | | | | | | 100%);IPR006674(100%) | | | | | | | | | | | |
| 359 | Variable 421 | GDSYECGFDK | Neorickettsia | Bacteria | Proteobacteria | | Alphaproteobacteria | | Rickettsiales | | Anaplasmataceae | Neorickettsia | | | | | | 7.1.1.-(100%) | Hydron translocation or charge separation linked to oxidoreductase | GO:0005886(100%);GO:0016 | plasma membrane(100%);integral component of | GO:0008137(100%);GO:0048 | NADH dehydrogenase (ubiquinone) activity (100%);quinone | | | IPR000440(100%);IPR038443 | 1.00E+00 | 1.00E+00 | 3.99E+04 | NS | NS | NS | NS | NS | NS | NS | NS |

| | | | | | | | | | | | | | | | | | | | | | | | | | | | | | | | | | | | | | | | | |
|---|---|---|---|---|---|---|---|---|---|---|---|---|---|---|---|---|---|---|---|---|---|---|---|---|---|---|---|---|---|---|---|---|---|---|---|---|---|---|---|---|
| | | | | | | | | | | | | | | | | | | | reactions (100%) | 021 (100%) | membrane (100%) | 038 (100%) | binding (100%) | | | 0 (100%); IPR023043 (80%) | | | | | | | | | | | | | | |
| 360 | Variable 422 | FMQQDEAPVMPSVCADHASALK | Zymomonas mobilis subsp. mobi | Bacteria | Proteobacteria | | Alphaproteobacteria | | Sphingomonadales | | Zymomonadaceae | Zymomonas | | | Zymomonas mobilis | Zymomonas mobilis subsp. m | Zymomonas mobilis subsp. mob | 3.1.3.1 (100%) | Alkaline phosphatase (100%) | | | GO:0004035 (100%); GO:004 | alkaline phosphatase activity (100%);metal ion binding (100%) | | | IPR002591 (100%); IPR0178 | 1.00E+00 | 1.00E+00 | 5.16E+04 | 4.1055 | 0.00001409 | 0.00003125 | 0.9 | NS | NS | NS | NS | 7.4469 | 0.0017056 | 0.00028741 |

| | | | | | | | | | | | | | | | | | | | | | | | | | | | | | | | | | | | | | |
|---|---|---|---|---|---|---|---|---|---|---|---|---|---|---|---|---|---|---|---|---|---|---|---|---|---|---|---|---|---|---|---|---|---|---|---|---|---|
| | | | lisZM4=ATCC31821 | | | | | | | | | | | | | obilis | ilisZM4=ATCC31821 | | | | | 6872(100%) | | | | 50(100%);IPR026263(100%) | | | | | | | | | | | |
| 361 | Variable 423 | LFTVERPDAVR | Betaproteobacteria | Bacteria | Proteobacteria | | Betaproteobacteria | | | | | | | | | | | 6.1.1.18(100%) | Glutamine--tRNA ligase (100%) | GO:0005737(99%);GO:00 | cytoplasm(99%);cytosol(1%) | GO:0004819(100%);GO:0 | glutamine-tRNA ligase activity (100%);ATP binding (10 | GO:0006424(98%);GO:0006425(89%);GO: | glutamyl-tRNA aminoacylation (98%); | IPR011035(100%);IPR02 | 1.00E+00 | 3.78E+06 | 1.00E+00 | NS | NS | NS | NS | NS | NS | NS | NS |

| | | | | | | | | | | | | | | | | | | | | | | | | | | |
|---|---|---|---|---|---|---|---|---|---|---|---|---|---|---|---|---|---|---|---|---|---|---|---|---|---|---|
| | | | | | | | | | | | | | | | | | | | | 05829(1%) | | 005524(100%) | 0%) | 0006418(2%) | glutaminyl-tRNA aminoacylation (89%);tRNA aminoacylation for protein translation (2% | 0056(100%);IPR020059(100%) |

| | | | | | | | | | | | | | | | | | | | | | | | | | | | | | | | | | | | | | |
|---|---|---|---|---|---|---|---|---|---|---|---|---|---|---|---|---|---|---|---|---|---|---|---|---|---|---|---|---|---|---|---|---|---|---|---|---|---|
| | | | | | | | | | | | | | | | | | | | | | | | | | ) | | | | | | | | | | | | |
| 362 | Variable 424 | CPFCDVGHGRPDPLDAQEPENLAR | Burkholderiales | Bacteria | Proteobacteria | | Betaproteobacteria | | Burkholderiales | | | | | | | | | 2.8.1.8(100%) | Lipoyl synthase(100%) | GO:0005737(100%) | cytoplasm(100%) | GO:0016992(100%);GO:0046872(100%);GO:0051539 | lipoate synthase activity(100%);metal ion binding(100%);4 iron | 4 sulfur cluster binding(100%) | GO:0009249(100%) | protein lipoylation(100%) | 1.00E+00 | 1.00E+00 | 2.68E+04 | NS | NS | NS | NS | NS | NS | NS | NS |

| | | | | | | | | | | | | | | | | | | | | | | | | | | | | | | | | | | | | | |
|---|---|---|---|---|---|---|---|---|---|---|---|---|---|---|---|---|---|---|---|---|---|---|---|---|---|---|---|---|---|---|---|---|---|---|---|---|---|
| | | | | | | | | | | | | | | | | | | | | | | (100%) | | | | | | | | | | | | | | | |
| 364 | Variable 426 | GPAPVAAALPAR | Bordetella avium 197N | Bacteria | Proteobacteria | | Betaproteobacteria | | Burkholderiales | | Alcaligenaceae | Bordetella | | | Bordetella avium | | Bordetella avium 197N | 3.1.1.61(100%);3.5.1.44(100%) | Protein-glutamate methylesterase (100%);Protein-glutamine glutaminase (100%) | GO:0005737(100%) | cytoplasm (100%) | GO:0000156(100%);GO:0008984(100%);GO:00 | phosphorelay response regulator activity (100%);protein-glutamate methylesterase activity (100%);protein | GO:0006482(100%);GO:0006935(100%);GO:0018277(100%) | protein demethylation (100%);chemotaxis (100%);protein deamination (100% | IPR000673(100%);IPR001789(100%);IPR0082 | 1.00E+00 | 1.00E+00 | 1.15E+05 | NS | NS | NS | NS | NS | NS | NS | NS |

| | | | | | | | | | | | | | | | | | | | | | | | | | | | | | | | | | | | | | | | | |
|---|---|---|---|---|---|---|---|---|---|---|---|---|---|---|---|---|---|---|---|---|---|---|---|---|---|---|---|---|---|---|---|---|---|---|---|---|---|---|---|---|
| | | | | | | | | | | | | | | | | | | | | | | 50568(100%) | -glutamine glutaminase activity (100%) | | ) | 48(100%) | | | | | | | | | | | | | | |
| 365 | Variable 427 | VTPATCFTPETVK | Ralstonia | Bacteria | Proteobacteria | | Betaproteobacteria | | Burkholderiales | | Burkholderiaceae | Ralstonia | | | | | | 3.2.1.15(100%) | Endo-polygalacturonase (100%) | GO:0005576(6%) | extracellular region (6%) | GO:0004650(100%) | polygalacturonase activity (100%) | GO:0005975(100%);GO:0071555(6%) | carbohydrate metabolic process (100%);cell wall organizatio | IPR000743(100%);IPR011050(10 | 1.00E+00 | 1.00E+00 | 1.23E+05 | 6.8838 | 0.00027388 | 0.00029412 | 0.9 | NS | NS | NS | NS | 6.4781 | 0.0035309 | 0.0005963 |

| | | | | | | | | | | | | | | | | | | | | | | | | | | | | | | | | | | | | | |
|---|---|---|---|---|---|---|---|---|---|---|---|---|---|---|---|---|---|---|---|---|---|---|---|---|---|---|---|---|---|---|---|---|---|---|---|---|---|
| | | | | | | | | | | | | | | | | | | | | | | | | | n (6%) | 0%); IPR012334 (100%) | | | | | | | | | | | |
| 366 | Variable 428 | GEPNEGNGPQPFER | Ralstonia | Bacteria | Proteobacteria | | Betaproteobacteria | | Burkholderiales | | Burkholderiaceae | Ralstonia | | | | | | | | | | GO:0004866 (100%) | endopeptidase inhibitor activity (100%) | | | IPR001599 (100%); IPR002890 ( | 1.00E+00 | 1.00E+00 | 4.43E+04 | 1.9695 | 0.027457 | 0 | 0.9 | NS | NS | NS | NS |

| | | | | | | | | | | | | | | | | | | | | | | | | | | | | | | | | | | | | | |
|---|---|---|---|---|---|---|---|---|---|---|---|---|---|---|---|---|---|---|---|---|---|---|---|---|---|---|---|---|---|---|---|---|---|---|---|---|---|
| | | | | | | | | | | | | | | | | | | | | | | | | | | 100%);IPR008930(100%) | | | | | | | | | | | |
| 367 | Variable 429 | LQFGQGEDFDK | Neisseriaceae | Bacteria | Proteobacteria | | Betaproteobacteria | | Neisseriales | | Neisseriaceae | | | | | | | 6.3.2.1(100%) | Pantoate--beta-alanine ligase (AMP-forming) (100%) | GO:0005737(97%);GO:00058 | cytoplasm(97%);cytosol(1%) | GO:0004592(100%);GO:0005 | pantoate-beta-alanine ligase activity (100%);ATP binding (96%) | GO:0015940 (100%) | pantothenate biosynthetic process (100%) | IPR003721(100%);IPR01472 | 1.00E+00 | 4.83E+04 | 1.00E+00 | NS | NS | NS | NS | NS | NS | NS | NS |

| | | | | | | | | | | | | | | | | | | | | | | | | | | | | | | | | | | | | | |
|---|---|---|---|---|---|---|---|---|---|---|---|---|---|---|---|---|---|---|---|---|---|---|---|---|---|---|---|---|---|---|---|---|---|---|---|---|---|
| | | | | | | | | | | | | | | | | | | | | 29(1%) | | 524(96%) | | | | 9(100%);IPR004821(99%) | | | | | | | | | | | |
| 368 | Variable 43 | LDKPGDFVGK | Actinomycetia | Bacteria | Actinobacteria | | Actinomycetia | | | | | | | | | | | 2.1.2.10(100%) | Aminomethyltransferase (100%) | | | GO:0004047(100%);GO:000 | aminomethyltransferase activity (100%);transaminase activity (10 | GO:0019464(87%);GO:0032259(77%);GO:0006 | glycine decarboxylation via glycine cleavage sy | IPR006222(100%);IPR0062 | 1.00E+00 | 2.37E+05 | 1.00E+00 | NS | NS | NS | NS | NS | NS | NS | NS |

| | | | | | | | | | | | | | | | | | | | | | | | | | | | | | | | | | | | | | |
|---|---|---|---|---|---|---|---|---|---|---|---|---|---|---|---|---|---|---|---|---|---|---|---|---|---|---|---|---|---|---|---|---|---|---|---|---|---|
| | | | | | | | | | | | | | | | | | | | | | | 8483(100%) | 0%) | 546(13%) | stem(87%); methylation(77%); glycine catabolic process(13%) | 23(100%); IPR013977(100%) | | | | | | | | | | | |
| 369 | Variable 4 | DAGLAVFHER | Neisseria | Bacteria | Proteobact | | Betaproteobacte | | Neisseriales | | Neisseriaceae | Neisseria | | | | | | | | GO:0009279 | cell outer membrane | | | GO:0009405(3%) | obsolete pathogenesis | IPR007655( | 1.00E+00 | 1.00E+00 | 1.62E+05 | 6.5369 | 0.007394 | 0.000125 | 0.9 | NS | NS | NS | NS |

| | | | | | | | | | | | | | | | | | | | | | | | | | | | | | | | | | | | | | |
|---|---|---|---|---|---|---|---|---|---|---|---|---|---|---|---|---|---|---|---|---|---|---|---|---|---|---|---|---|---|---|---|---|---|---|---|---|---|
| | 30 | | | | eria | | ria | | | | | | | | | | | | | (100%);GO:0016021(100%) | (100%);integral component of membrane(100%) | | | | (3%) | 100%);IPR011990(100%);IPR013026(97%) | | | | | | | | | | | |
| 370 | Variable | AEAAGHVVK | Neisseria | Bacteria | Proteobac | | Betaproteobac | | Neisseriales | | Neisseriaceae | Neisseria | | | | | | 1.1.1.86( | Ketol-acid reductoisomera | GO:000582 | cytosol(3%) | GO:00028 | magnesium ion binding(10 | GO:0009097(100%) | isoleucine biosynth | IPR000506 | 1.00E+00 | 1.00E+00 | 4.97E+05 | NS | NS | NS | NS | NS | NS | NS | NS |

| | | | | | | | | | | | | | | | | | | | | | | | | | | | | | | | | | | | | | |
|---|---|---|---|---|---|---|---|---|---|---|---|---|---|---|---|---|---|---|---|---|---|---|---|---|---|---|---|---|---|---|---|---|---|---|---|---|---|
| | 431 | | | | teria | | teria | | | | | | | | | | | 100%) | se(NADP(+))(100%) | 9(3%) | | 7(100%);GO:0004455(100%);GO:0050661(97%) | 0%);ketol-acid reductoisomerase activity(100%);NADP binding(97%) | ;GO:0009099(100%) | etic process(100%);valine biosynthetic process(100%) | (100%);IPR008927(100%);IPR013023(100%) | | | | | | | | | | | |
| 371 | Varia | ENGVLVNSGD | Neisseria | Bacte | Prote | | Betaprot | | Neisseria | | Neisseria | Neisser | | | | | | 6.1.1 | Leucine--tR | GO:00 | cytoplasm | GO:00 | leucine-tRN | GO:000642 | leucyl-tR | IPR00 | 1.00E | 1.00E | 1.99E | 2.097 | 0.013 | 0.006 | 0.9 | NS | NS | NS | NS |

| | | | | | | | | | | | | | | | | | | | | | | | | | | | | | | | | | | | | | |
|---|---|---|---|---|---|---|---|---|---|---|---|---|---|---|---|---|---|---|---|---|---|---|---|---|---|---|---|---|---|---|---|---|---|---|---|---|---|
| | ble 432 | LDGLDFQTAFDAVAAK | | ria | obacteria | | eobacteria | | les | | ceae | ia | | | | | | .4(100%) | NA ligase (100%) | 0573 37(97%);GO:0005829(3%) | (97%);cytosol(3%) | 048233(100%);GO:0005524(100%);GO:0002161(90%) | A ligase activity (100%);ATP binding (100%);aminoacyl-tRNA editing activity (90%) | 9(100%) | NA aminoacylation (100%) | 1412(100%);IPR002300(100%);IPR002302(100%) | +00 | +00 | +05 | 3 | 89 | 0895 | | | | | |
| 3 | V | TTL | N | B | P | | B | | N | | N | N | | | | | | | | G | cy | G | sin | | G | D | 1 | 1 | 3 | N | N | N | N | N | N | N | N |

| | | | | | | | | | | | | | | | | | | | | | | | | | | | | | | | | | | | | | |
|---|---|---|---|---|---|---|---|---|---|---|---|---|---|---|---|---|---|---|---|---|---|---|---|---|---|---|---|---|---|---|---|---|---|---|---|---|---|
| 72 | ariable 433 | CLEAVAQCQK | eisseria | acteria | roteobacteria | | etaproteobacteria | | eisseriales | | eisseriaceae | eisseria | | | | | | | | O:0005737(100%) | toplasm(100%) | O:0003697(100%);GO:0005524(100%);GO:0008094(1 | gle-stranded DNA binding(100%);ATP binding(100%);ATPase | acting on DNA(100%) | O:0006281(100%);GO:0006310(100%);GO:0009432(96%) | NA repair(100%);DNA recombination(100%);SOS response(96 | .00E+00 | .00E+00 | .44E+04 | S | S | S | S | S | S | S | S |

| | | | | | | | | | | | | | | | | | | | | | | | | | | | | | | | | | | | | | |
|---|---|---|---|---|---|---|---|---|---|---|---|---|---|---|---|---|---|---|---|---|---|---|---|---|---|---|---|---|---|---|---|---|---|---|---|---|---|
| | | | | | | | | | | | | | | | | | | | | | | 00%) | | | | %) | | | | | | | | | | | |
| 373 | Variable 434 | GVAQELGGR | Neisseria | Bacteria | Proteobacteria | | Betaproteobacteria | | Neisseriales | | Neisseriaceae | Neisseria | | | | | | 6.3.2.3(100%) | Glutathione synthase (100%) | GO:0005737(5%) | cytoplasm(5%) | GO:0004363(100%);GO:0005524(100%);GO:0046 | glutathione synthase activity(100%);ATP binding(100%);metal ion binding(100%) | | | IPR004215(100%);IPR004218(100%);IPR006284 | 1.00E+00 | 1.00E+00 | 2.93E+04 | NS | NS | NS | NS | NS | NS | NS | NS |

| | | | | | | | | | | | | | | | | | | | | | | | | | | | | | | | | | | | | | |
|---|---|---|---|---|---|---|---|---|---|---|---|---|---|---|---|---|---|---|---|---|---|---|---|---|---|---|---|---|---|---|---|---|---|---|---|---|---|
| | | | | | | | | | | | | | | | | | | | | | | 872(100%) | | | | (100%) | | | | | | | | | | | |
| 374 | Variable 435 | WVTPSTQDLFGK | Neisseria | Bacteria | Proteobacteria | | Betaproteobacteria | | Neisseriales | | Neisseriaceae | Neisseria | | | | | | 4.99.1.1(97%);4.99.1.9(3%) | Protoporphyrin ferrochelatase (97%); Coproporphyrin ferrochelatase (3%) | GO:0005737(97%) | cytoplasm(97%) | GO:0004325(100%);GO:0046872(94%) | ferrochelatase activity (100%);metal ion binding (94%) | GO:0006783(100%) | heme biosynthetic process (100%) | IPR001015(100%);IPR033644(100%);IPR0 | 1.00E+00 | 1.00E+00 | 4.61E+04 | NS | NS | NS | NS | NS | NS | NS | NS |

| | | | | | | | | | | | | | | | | | | | | | | | | | | | | | | | | | | | | | |
|---|---|---|---|---|---|---|---|---|---|---|---|---|---|---|---|---|---|---|---|---|---|---|---|---|---|---|---|---|---|---|---|---|---|---|---|---|---|
| | | | | | | | | | | | | | | | | | | | | | | | | | | 19772(97%) | | | | | | | | | | | |
| 375 | Variable 436 | FAVEQELVAASSR | Neisseria | Bacteria | Proteobacteria | | Betaproteobacteria | | Neisseriales | | Neisseriaceae | Neisseria | | | | | | 6.4.1.2(100%) | Acetyl-CoA carboxylase(100%) | GO:0009279(86%) | cell outer membrane(86%) | GO:0003989(14%) | acetyl-CoA carboxylase activity(14%) | GO:0007155(86%);GO:0009405(86%) | cell adhesion(86%); obsolete pathogenesis(86%) | | 1.00E+00 | 1.00E+00 | 4.68E+04 | 2.5678 | 0.034996 | 0.00016776 | 0.9 | NS | NS | NS | NS |
| 377 | Variable | EGAASLDLCAVATGR | Neisseria | Bacteria | Proteobac | | Betaproteobac | | Neisseriales | | Neisseriaceae | Neisseria | | | | | | 3.1.3.25( | Inositol-phosphate phosp | GO:000573 | cytoplasm(7%) | GO:000893 | inositol monophosphate | GO:0046854(100%) | phosphatidylinositol | IPR000760 | 1.00E+00 | 9.28E+04 | 1.00E+00 | NS | NS | NS | NS | NS | NS | NS | NS |

| | | | | | | | | | | | | | | | | | | | | | | | | | | |
|---|---|---|---|---|---|---|---|---|---|---|---|---|---|---|---|---|---|---|---|---|---|---|---|---|---|---|
| | 438 | | | | teria | | teria | | | | | | | | | | | 100%) | hatase(100%) | 7(7%) | | 4(100%);GO:0046872(100%);GO:0052832(100%) | 1-phosphatase activity(100%);metal ion binding(100%);inositol monophosphate 3-phosphatase activity(100%) | ;GO:0046855(100%);GO:0031564(7%) | phosphate biosynthetic process(100%);inositol phosphate dephosphorylation(100%);transcripti | (100%);IPR020550(100%);IPR020583(100%) |

| | | | | | | | | | | | | | | | | | | | | | | | | | | | | | | | | | | | | | |
|---|---|---|---|---|---|---|---|---|---|---|---|---|---|---|---|---|---|---|---|---|---|---|---|---|---|---|---|---|---|---|---|---|---|---|---|---|---|
| | | | | | | | | | | | | | | | | | | | | | | | | | onantitermination (7%) | | | | | | | | | | | | |
| 378 | Variable 439 | AAPFPAALLANQK | Neisseriameningitidis | Bacteria | Proteobacteria | | Betaproteobacteria | | Neisseriales | | Neisseriaceae | Neisseria | | | Neisseriameningitidis | | | 1.8.1.2(100%) | Assimilatory sulfite reductase (NADPH) (100%) | GO:0005829(11%) | cytosol (11%) | GO:0004783(100%);GO:0010181(100%); | sulfite reductase (NADPH) activity (100%);FMN binding (100%);flavin adenine dinucleotide | GO:0000103(100%);GO:0019344(100%);GO:0070814(100%) | sulfate assimilation (100%);cysteine biosynthetic process (100%);hydr | IPR001094(100%);IPR001433(100%);IP | 1.00E+00 | 1.00E+00 | 4.24E+04 | NS | NS | NS | NS | NS | NS | NS | NS |

| | | | | | | | | | | | | | | | | | | | | | | | | | | | | | | | | | | | | | |
|---|---|---|---|---|---|---|---|---|---|---|---|---|---|---|---|---|---|---|---|---|---|---|---|---|---|---|---|---|---|---|---|---|---|---|---|---|---|
| | | | | | | | | | | | | | | | | | | | | | | GO:0050660(100%) | binding(100%) | | ogensulfide biosynthetic process(100%) | R001709(100%) | | | | | | | | | | | |
| 379 | Variable 44 | FFDQLGDYAVSQGAK | Actinomycetia | Bacteria | Actinobacteria | | Actinomycetia | | | | | | | | | | | 6.1.1.12(100%) | Aspartate--tRNA ligase(100%) | GO:0005737(100%) | cytoplasm(100%) | GO:0005524(100%);GO:00048 | ATP binding(100%);aspartate-tRNA ligase activity(98%);nucleic | GO:0006422(98%);GO:0006418(2%) | aspartyl-tRNA aminoacylation(98%);tRNA ami | IPR002312(100%);IPR004115 | 1.00E+00 | 1.20E+05 | 1.00E+00 | NS | NS | NS | NS | NS | NS | NS | NS |

| | | | | | | | | | | | | | | | | | | | | | | | | | | | | | | | | | | | | | | | | |
|---|---|---|---|---|---|---|---|---|---|---|---|---|---|---|---|---|---|---|---|---|---|---|---|---|---|---|---|---|---|---|---|---|---|---|---|---|---|---|---|---|
| | | | | | | | | | | | | | | | | | | | | | | 15(98%);GO:0003676(93%) | acid binding (93%) | | noacylation for protein translation (2%) | (100%);IPR004364(100%) | | | | | | | | | | | | | | |
| 380 | Variable 440 | FQTALSHAFHQLLTHR | Neisseriameningitidis | Bacteria | Proteobacteria | | Betaproteobacteria | | Neisseriales | | Neisseriaceae | Neisseria | | | Neisseriameningitidis | | | 3.1.11.6(100%) | Exodeoxyribonuclease VII (100%) | GO:0005737(100%);GO:00 | cytoplasm (100%);exodeoxyribonuclease VII co | GO:0003676(100%);GO:00 | nucleic acid binding (100%);exodeoxyribonuclease VII act | GO:0006308 (100%) | DNA catabolic process (100%) | IPR003753(100%);IPR020 | 1.00E+00 | 1.00E+00 | 3.87E+04 | 0.5124 | 3.757E-08 | 0.00043542 | 0.9 | NS | NS | NS | NS | 9.7085 | 0.003421 | 0.008302 |

| | | | | | | | | | | | | | | | | | | | | | | | | | | | | | | | | | | | | | |
|---|---|---|---|---|---|---|---|---|---|---|---|---|---|---|---|---|---|---|---|---|---|---|---|---|---|---|---|---|---|---|---|---|---|---|---|---|---|
| | | | | | | | | | | | | | | | | | | | | 09318(100%) | mplex(100%) | 08855(100%) | ivity(100%) | | | 579(100%);IPR025824(100%) | | | | | | | | | | | |
| 381 | Variable 441 | LLAEENAALALPAYELVLK | Neisseria meningitidis | Bacteria | Proteobacteria | | Betaproteobacteria | | Neisseriales | | Neisseriaceae | Neisseria | | | Neisseria meningitidis | | | 6.1.1.14(100%) | Glycine--tRNA ligase(100%) | GO:0005737(100%) | cytoplasm(100%) | GO:0004820(100%);GO: | glycine-tRNA ligase activity(100%);ATP binding(10 | GO:0006426(100%) | glycyl-tRNA aminoacylation(100%) | IPR002310(100%);IPR0 | 1.00E+00 | 1.00E+00 | 3.64E+04 | NS | NS | NS | NS | NS | NS | NS | NS |

| | | | | | | | | | | | | | | | | | | | | | | | | | | | | | | | | | | | | | |
|---|---|---|---|---|---|---|---|---|---|---|---|---|---|---|---|---|---|---|---|---|---|---|---|---|---|---|---|---|---|---|---|---|---|---|---|---|---|
| | | | | | | | | | | | | | | | | | | | | | | 0005524(100%) | 0%) | | | 06194(100%) | | | | | | | | | | | |
| 382 | Variable 442 | MTDTAENQTQNNWQAGHPR | Neisseriameningitidis | Bacteria | Proteobacteria | | Betaproteobacteria | | Neisseriales | | Neisseriaceae | Neisseria | | | Neisseriameningitidis | | | 2.1.1.33(100%) | tRNA(guanine(46)-N(7))-methyltransferase(100%) | GO:0043527(25%) | tRNAmethyltransferase complex(25%) | GO:0008176(100%) | tRNA(guanine-N7-)-methyltransferase activity(100%) | GO:0030488(25%);GO:0036265(25%) | tRNA methylation(25%);RNA(guanine-N7)-methylation | IPR003358(100%);IPR029063(100%) | 1.00E+00 | 1.00E+00 | 4.84E+04 | 3.7743 | 0.0018115 | 0.000076316 | 0.9 | NS | NS | NS | NS |

| | | | | | | | | | | | | | | | | | | | | | | | | | | | | | | | | | | | | | |
|---|---|---|---|---|---|---|---|---|---|---|---|---|---|---|---|---|---|---|---|---|---|---|---|---|---|---|---|---|---|---|---|---|---|---|---|---|---|
| | | | | | | | | | | | | | | | | | | | | | | | | | (25%) | | | | | | | | | | | | |
| 383 | Variable 443 | FVATAGVGLR | Neisseriameningitidis | Bacteria | Proteobacteria | | Betaproteobacteria | | Neisseriales | | Neisseriaceae | Neisseria | | | Neisseriameningitidis | | | | | GO:0009279(100%);GO:0046930(100%) | cell outer membrane(100%);pore complex(100%) | GO:0015288(100%) | porin activity(100%) | GO:0034220(100%) | ion transmembrane transport(100%) | IPR001702(100%);IPR002299(100%);IPR013793 | 1.00E+00 | 6.45E+04 | 1.00E+00 | NS | NS | NS | NS | NS | NS | NS | NS |

| | | | | | | | | | | | | | | | | | | | | | | | | | | | | | | | | | | | | | |
|---|---|---|---|---|---|---|---|---|---|---|---|---|---|---|---|---|---|---|---|---|---|---|---|---|---|---|---|---|---|---|---|---|---|---|---|---|---|
| | | | | | | | | | | | | | | | | | | | | | | | | | | (100%) | | | | | | | | | | | |
| 384 | Variable 444 | LNYSTNTVGYR | Neisseriameningitidis | Bacteria | Proteobacteria | | Betaproteobacteria | | Neisseriales | | Neisseriaceae | Neisseria | | | Neisseriameningitidis | | | 2.7.4.14(50%);6.1.1.22(50%) | UMP/CMP kinase(50%);Asparagine--tRNA ligase(50%) | GO:0009279(93%);GO:0016021(3%);GO:00312 | cell outer membrane(93%);integral component of membrane(3%);intrinsic com | GO:0004127(3%);GO:0004816(3%);GO:001534 | cytidylate kinase activity(3%);asparagine-tRNA ligase activity(3%);siderophore uptake transmembran | GO:0044718(3%) | siderophore transmembrane transport(3%) | IPR039426(100%);IPR000531(97%);IPR01091 | 1.00E+00 | 4.85E+04 | 1.00E+00 | NS | NS | NS | NS | NS | NS | NS | NS |

| | | | | | | | | | | | | | | | | | | | | | | | | | | | | | | | | | | | | | |
|---|---|---|---|---|---|---|---|---|---|---|---|---|---|---|---|---|---|---|---|---|---|---|---|---|---|---|---|---|---|---|---|---|---|---|---|---|---|
| | | | | | | | | | | | | | | | | | | | | 30(3%) | ponent of cell outer membrane(3%) | 4(3%) | e transporter activity (3%) | | | 7(97%) | | | | | | | | | | | |
| 385 | Variable 445 | WLGAEGHAFTPYPGR | Neisseriameningitidis | Bacteria | Proteobacteria | | Betaproteobacteria | | Neisseriales | | Neisseriaceae | Neisseria | | | Neisseriameningitidis | | | 2.7.1.170(100%) | Anhydro-N-acetylmuramic acid kinase(100%) | | | GO:0005524(100%);GO:0016301 | ATP binding(100%);kinase activity(100%);phosphotransferase activity | alcohol group as acceptor(100%) | GO:0005975(100%);GO:0006040(100%);GO:0009254 | carbohydrate metabolic process(100% | 1.47E+07 | 1.00E+00 | 1.00E+00 | NS | NS | NS | NS | NS | NS | NS | NS |

| | | | | | | | | | | | | | | | | | | | | | | | | | | |
|---|---|---|---|---|---|---|---|---|---|---|---|---|---|---|---|---|---|---|---|---|---|---|---|---|---|---|
| | | | | | | | | | | | | | | | | | | | | | | (100%);GO:0016773(100%) | | | (100%) | );aminosugarmetabolicprocess(100%);peptidoglycanturnover |

| | | | | | | | | | | | | | | | | | | | | | | | | | | | | | | | | | | | | | |
|---|---|---|---|---|---|---|---|---|---|---|---|---|---|---|---|---|---|---|---|---|---|---|---|---|---|---|---|---|---|---|---|---|---|---|---|---|---|
| | | | | | | | | | | | | | | | | | | | | | | | | | | (100%) | | | | | | | | | | | |
| 388 | Variable 448 | CTLFVSGCVHQCK | Gammaproteobacteria | Bacteria | Proteobacteria | | Gammaproteobacteria | | | | | | | | | | | 1.97.1.-(100%) | Other oxidoreductases (100%) | GO:0005737(2%) | cytoplasm (2%) | GO:0043365(100%);GO:0046872(100%);GO:00 | [formate-C-acetyltransferase]-activating enzyme activity (100%);metal ion binding (100%);4 iron | 4 sulfur cluster binding (100%) | GO:0009265 (2%) | 2'-deoxyribonucleotide biosynthetic process (2%) | 1.00E+00 | 1.00E+00 | 1.36E+05 | NS | NS | NS | NS | NS | NS | NS | NS |

| | | | | | | | | | | | | | | | | | | | | | | | | | | | | | | | | | | | | | |
|---|---|---|---|---|---|---|---|---|---|---|---|---|---|---|---|---|---|---|---|---|---|---|---|---|---|---|---|---|---|---|---|---|---|---|---|---|---|
| | | | | | | | | | | | | | | | | | | | | | | 515539(100%) | | | | | | | | | | | | | | | |
| 389 | Variable 449 | TPTSDEMYFTFK | Gammaproteobacteria | Bacteria | Proteobacteria | | Gammaproteobacteria | | | | | | | | | | | | | GO:0009279(79%);GO:0016021(79%);GO | cell outer membrane(79%);integral component of membrane(79% | GO:0022857(93%);GO:0015344(7%) | transmembrane transporter activity(93%);siderophore uptake transmembrane transporter | GO:0044718(7%) | siderophore transmembrane transport(7%) | IPR000531(100%);IPR039426(100%);IP | 1.00E+00 | 1.00E+00 | 4.17E+04 | NS | NS | NS | NS | NS | NS | NS | NS |

| | | | | | | | | | | | | | | | | | | | | | | | | | | | | | | | | | | | | | | | | |
|---|---|---|---|---|---|---|---|---|---|---|---|---|---|---|---|---|---|---|---|---|---|---|---|---|---|---|---|---|---|---|---|---|---|---|---|---|---|---|---|---|
| | | | | | | | | | | | | | | | | | | | | :0019867(14%) | );outer membrane(14%) | | activity(7%) | | | R0109498(88%) | | | | | | | | | | | | | | |
| 391 | Variable 450 | DDHESGTDR | Gammaproteobacteria | Bacteria | Proteobacteria | | Gammaproteobacteria | | | | | | | | | | | 2.7.7.38(100%) | 3-deoxy-manno-octulosonate cytidylyltransferase (100%) | GO:0005737(94%) | cytoplasm(94%) | GO:0008690(100%) | 3-deoxy-manno-octulosonate cytidylyltransferase activity (100%) | GO:0009103(100%);GO:0033468(94%) | lipopolysaccharide biosynthetic process(100%);CMP-keto-3-de | IPR003329(100%);IPR004528(100%); | 1.00E+00 | 1.00E+00 | 1.12E+05 | NS | NS | NS | NS | NS | NS | NS | NS | 7.1136 | 0.0021845 | 0.00059904 |

| | | | | | | | | | | | | | | | | | | | | | | | | | | | | | | | | | | | | | |
|---|---|---|---|---|---|---|---|---|---|---|---|---|---|---|---|---|---|---|---|---|---|---|---|---|---|---|---|---|---|---|---|---|---|---|---|---|---|
| | | | | | | | | | | | | | | | | | | | | | | | | | oxy-D-manno-octulosonic acid biosynthetic process (94%) | IPR029044(100%) | | | | | | | | | | | |
| 392 | Variable 451 | ALTQVGLVLRPFLPEQVR | Gammaproteobacteria | Bacteria | Proteobacteria | | Gammaproteobacteria | | | | | | | | | | | 1.1.99.14(97%);1 | Glycolate dehydrogenase (97%); Glycerol-3-ph | GO:0005886(0%) | plasma membrane (0%) | GO:0051536(96%); | iron-sulfur cluster binding (96%);metal | GO:0046296(0%) | glycolate catabolic process (0% | IPR012257(100%); | 1.00E+00 | 1.00E+00 | 1.49E+05 | 10.129 | 0.00060329 | 0.00027014 | 0.9 | NS | NS | NS | NS |

| | | | | | | | | | | | | | | | | | | | | | | | | | | | | | | | | | | | | | | | | |
|---|---|---|---|---|---|---|---|---|---|---|---|---|---|---|---|---|---|---|---|---|---|---|---|---|---|---|---|---|---|---|---|---|---|---|---|---|---|---|---|---|
| | | | | | | | | | | | | | | | | | | .1.5.3(3%);1.1.3.15(1%) | osphate dehydrogenase (3%);(S)-2-hydroxy-acid oxidase (1%) | | | GO:0046872(94%);GO:0019154(40%) | ion binding(94%);glycolate dehydrogenase activity(40%) | | ) | IPR004017(99%);IPR017896(94%) | | | | | | | | | | | | | | |
| 393 | Variable 452 | VTDCGGVLHCFTEDR | Gammaproteobact | Bacteria | Proteobacteria | | Gammaproteobacteri | | | | | | | | | | | 3.1.21.-(89%);3 | Endodeoxyribonucleases producing 5'-ph | GO:0005829(0% | cytosol(0%);DNA polym | GO:0016888(88 | endodeoxyribonuclease activity | producing 5'-phosphomonoeste | GO:0006260(0%) | DNA replication(0%) | 1.00E+00 | 1.00E+00 | 1.30E+05 | 19.244 | 5.0797E-07 | 0.0004065 | 0.9 | NS | NS | NS | NS | 12.048 | 0.0007327 | 0.0023 |

| | | | | | | | | | | | | | | | | | | | | | | | | | | | | | | | | | | | | | |
|---|---|---|---|---|---|---|---|---|---|---|---|---|---|---|---|---|---|---|---|---|---|---|---|---|---|---|---|---|---|---|---|---|---|---|---|---|---|
| | | | eria | | | | a | | | | | | | | | | | .1.-.-(11%);2.7.7.7(2%) | osphomonoesters (89%);Acting on ester bonds (11%);DNA-directed DNA polymerase (2%) | );GO:0009360(0%) | erase III complex (0%) | %);GO:0004536(10%);GO:0016787(2%) | | rs (88%);deoxyribonuclease activity (10%);hydrolase activity (2%) | | | | | | | | | | | | | |
| 394 | Variable 453 | VDWPSAVVEGGSSGSGLLTVAGDGS | Gammaproteoba | Bacteria | Proteobacteri | | Gammaproteobacte | | | | | | | | | | | 3.4.21.50(10 | Lysylendopeptidase (100%) | GO:0005576(4 | extracellular region (4%);extr | GO:0008233(7 | peptidase activity (75%);hydrolas | GO:0009405 (8%);GO:0006 | obsolete pathogenesis (8% | IPR009003(10 | 5.05E+05 | 1.00E+00 | 1.00E+00 | NS | NS | NS | NS | NS | NS | NS | NS |

| | | | | | | | | | | | | | | | | | | | | | | | | | | | | | | | | | | | | | | | | |
|---|---|---|---|---|---|---|---|---|---|---|---|---|---|---|---|---|---|---|---|---|---|---|---|---|---|---|---|---|---|---|---|---|---|---|---|---|---|---|---|---|
| | | YQLR | cteria | | a | | ria | | | | | | | | | | | 0%) | | %);GO:0005615(4%) | acellular space (4%) | 5%);GO:0016787(17%);GO:0008236(8%) | e activity (17%);serine-type peptidase activity (8%) | 508(4%);GO:0006554(4%) | ); proteolysis (4%);lysine catabolic process (4%) | 0%); IPR043504(100%) | | | | | | | | | | | | | | |
| 397 | Variable 45 | NLPQVAGVSK | Pseudoalterom | Bacteria | Proteobacte | | Gammaproteobac | | Alteromonadales | | Pseudoalteromonadaceae | Pseudoalterom | | | | | | 6.1.1.20(10 | Phenylalanine--tRNA ligase (100% | GO:0005737( | cytoplasm (100%) | GO:0000049( | tRNA binding (100%);magnes | GO:0006432(100%) | phenylalanyl-tRNA | IPR002547(1 | 1.00E+00 | 1.00E+00 | 4.51E+04 | 6.9641 | 4.763E-08 | 0.0001765 | 0.9 | NS | NS | NS | NS | 16.955 | 3.9996E-06 | 0.0050712 |

| | | | | | | | | | | | | | | | | | | | | | | | | | | | | | | | | | | | | | |
|---|---|---|---|---|---|---|---|---|---|---|---|---|---|---|---|---|---|---|---|---|---|---|---|---|---|---|---|---|---|---|---|---|---|---|---|---|---|
| | 6 | | onas | | ria | | teria | | | | e | onas | | | | | | 0%) | ) | 100%) | | 100%);GO:0000287(100%);GO:0004826(100%) | ium ion binding (100%);phenylalanine-tRNA ligase activity (100%) | | minoacylation (100%) | 00%);IPR004532(100%);IPR005121(100%) | | | | | | | | | | | |
| 398 | Variab | VAGPAYTALAK | Pseudo | Bacter | Proteo | | Gammapr | | Alteromo | | Pseudoaltero | Pseudo | | | | | | 3.2.1. | Beta-N-acetylhex | | | GO:000 | hydrolase activit | hydrolyzing | GO:0005975 | carbohy | 1.00E+ | 4.19E+ | 1.00E+ | NS | NS | NS | NS | NS | NS | NS | NS |

| | | | | | | | | | | | | | | | | | | | | | | | | | | | | | |
|---|---|---|---|---|---|---|---|---|---|---|---|---|---|---|---|---|---|---|---|---|---|---|---|---|---|---|---|---|---|
| | le 457 | | alteromonas | ia | bacteria | | oteobacteria | | nadales | | monadaceae | alteromonas | | | | | | 52(100%) | osaminidase(100%) | | | 4553(71%);GO:0004563(29%);GO:0102148(29%) | y | O-glycosyl compounds(71%);beta-N-acetylhexosaminidase activity(29%);N-acetyl-beta-D-galactosaminida | (100%) | drate metabolic process(100%) | 00 | 05 | 00 |

| | | | | | | | | | | | | | | | | | | | | | | | | | | | | | | | | | | | | | |
|---|---|---|---|---|---|---|---|---|---|---|---|---|---|---|---|---|---|---|---|---|---|---|---|---|---|---|---|---|---|---|---|---|---|---|---|---|---|
| | | | | | | | | | | | | | | | | | | | | | | | | se activity (29%) | | | | | | | | | | | | | |
| 399 | Variable 458 | NDDDFDDDSADR | CelvibriojaponicusUeda107 | Bacteria | Proteobacteria | | Gammaproteobacteria | | Cellvibrionales | | Cellvibrionaceae | Cellvibrio | | | Cellvibriojaponicus | | CellvibriojaponicusUeda107 | | | GO:0005737(100%) | cytoplasm(100%) | GO:0003743(100%);GO:0003924(100%);GO:0 | translation initiation factor activity(100%);GTPase activity(100%);GTP binding(100%) | | | IPR000178(100%);IPR000795(100%);IPR005 | 1.00E+00 | 9.47E+04 | 1.00E+00 | NS | NS | NS | NS | NS | NS | NS | NS |

| | | | | | | | | | | | | | | | | | | | | | | | | | | | | | | | | | | | | | |
|---|---|---|---|---|---|---|---|---|---|---|---|---|---|---|---|---|---|---|---|---|---|---|---|---|---|---|---|---|---|---|---|---|---|---|---|---|---|
| | | | | | | | | | | | | | | | | | | | | | | 0055525(100%) | | | | 225(100%) | | | | | | | | | | | |
| 400 | Variable 459 | GESAYNDDLAPVVAEALEAQGLAVEDQGAK | Saccharophagus degradans 2-40 | Bacteria | Proteobacteria | | Gammaproteobacteria | | Cellvibrionales | | Cellvibrionaceae | Saccharophagus | | | Saccharophagus degradans | | Saccharophagus degradans 2-40 | 6.1.1.19(100%) | Arginine--tRNA ligase(100%) | GO:0005737(100%) | cytoplasm(100%) | GO:0004814(100%);GO:0005524(100% | arginine-tRNA ligase activity(100%);ATP binding(100%) | GO:0006420(100%) | arginyl-tRNA aminoacylation(100%) | IPR001278(100%);IPR001412(100%);I | 1.00E+00 | 1.00E+00 | 7.77E+04 | 3.9607 | 0.02892 | 0.0019211 | 0.9 | NS | NS | NS | NS |

| | | | | | | | | | | | | | | | | | | | | | | | | | | | | | | | | | | | | | |
|---|---|---|---|---|---|---|---|---|---|---|---|---|---|---|---|---|---|---|---|---|---|---|---|---|---|---|---|---|---|---|---|---|---|---|---|---|---|
| | | | | | | | | | | | | | | | | | | | | | | ) | | | | PR005148(100%) | | | | | | | | | | | |
| 401 | Variable 46 | VLLAGHLDTVPVAGNLPSR | Actinomycetia | Bacteria | Actinobacteria | | Actinomycetia | | | | | | | | | | | 3.5.1.18(98%);3.-.-.-(2%) | Succinyl-diaminopimelate desuccinylase (98%);Hydrolases (2%) | | | GO:0009014(100%);GO:0046872(1% | succinyl-diaminopimelate desuccinylase activity (100%);metal ion binding (1%) | GO:0009089 (100%);GO:0019877 (1%);GO:0006526 (1%) | lysine biosynthetic process via diaminopimelate (100%);di | IPR002933(100%);IPR010174(100% | 1.00E+00 | 8.53E+04 | 1.00E+00 | NS | NS | NS | NS | NS | NS | NS | NS |

| | | | | | | | | | | | | | | | | | | | | | | | | | | | | | | | | | | | | | | | | |
|---|---|---|---|---|---|---|---|---|---|---|---|---|---|---|---|---|---|---|---|---|---|---|---|---|---|---|---|---|---|---|---|---|---|---|---|---|---|---|---|---|
| | | | | | | | | | | | | | | | | | | | | | | );GO:0008777(1%) | ;acetylornithine deacetylase activity (1%) | | aminopimelate biosynthetic process (1%);arginine biosynthetic process (1%) | );IPR011650(100%) | | | | | | | | | | | | | | |
| 402 | Variab | DVTTGDTLCAESNK | Sacchar | Bacter | Proteo | | Gammapr | | Celvibrio | | Cellvibriona | Sacchar | | | Sacchar | | Sacchar | | | GO:000 | cytoplasm (1 | GO:000 | translation elong | | | IPR000 | 1.00E+ | 1.00E+ | 5.24E+ | 0.4434 | 1.491E | 0.0010 | 0.9 | NS | NS | NS | NS | 11.699 | 0.000 | 0.005 |

| | | | | | | | | | | | | | | | | | | | | | | | | | | | | | | | | | | | | | | | | |
|---|---|---|---|---|---|---|---|---|---|---|---|---|---|---|---|---|---|---|---|---|---|---|---|---|---|---|---|---|---|---|---|---|---|---|---|---|---|---|---|---|
| | le 460 | | ophagusdegradans2-40 | ia | bacteria | | oteobacteria | | nales | | ceae | ophagus | | | ophagusdegradans | | ophagusdegradans2-40 | | | 5737(100%) | 00%) | 3746(100%);GO:0003924(100%);GO:0005525(100%) | ation factor activity (100%);GTPase activity (100%);GTP binding (100%) | | | 640(100%);IPR000795(100%);IPR004161(100%) | 00 | 00 | 04 | 6 | -08 | 2 | | | | | | | 91495 | 0876 |
| 4 | V | AY | E | B | P | | G | | E | | | | | | | | | 5 | Ph | 6 | | | G | ma | | G | 1 | 1 | 2 | 0 | 3 | 0. | 0 | N | N | N | N | 8 | 0 | 0 |

| | | | | | | | | | | | | | | | | | | | | | | | | | | | | | | | | | | | | | | | | |
|---|---|---|---|---|---|---|---|---|---|---|---|---|---|---|---|---|---|---|---|---|---|---|---|---|---|---|---|---|---|---|---|---|---|---|---|---|---|---|---|---|
| 04 | ariable 462 | GEFLKPGK | nterobacterales | acteria | roteobacteria | | ammaproteobacteria | | nterobacterales | | | | | | | | | .4.2.8(79%);5.4.2.2(24%);2.7.7.13(1%) | osphomannomutase(79%);Phosphoglucomutase(alpha-D-glucose-1 | -bisphosphate-dependent)(24%);Mannose-1-phosphateg | | | O:0000287(92%);GO:0004615(54%);GO:0016868(33%) | gnesium ion binding(92%);phosphomannomutase activity(54%);intramolecular transferase activity | phosphotransferases(33%) | O:0005975(96%);GO:0009298(2%);GO:0000271(1%) | .00E+00 | .00E+00 | .92E+05 | .4197 | .4703E-08 | 00085995 | .9 | S | S | S | S | .6464 | .00071645 | .00030989 |

| | | | | | | | | | | | | | | | | | | | | | | | | | | | | | | | | | | | | | |
|---|---|---|---|---|---|---|---|---|---|---|---|---|---|---|---|---|---|---|---|---|---|---|---|---|---|---|---|---|---|---|---|---|---|---|---|---|---|
| | | | | | | | | | | | | | | | | | | | | uanylyltransferase(1%) | | | | | | | | | | | | | | | | | |
| 405 | Variable 463 | FHLMDFGTR | Enterobacterales | Bacteria | Proteobacteria | | Gammaproteobacteria | | Enterobacterales | | | | | | | | | 6.3.4.21(100%);2.4.2.11(0% | Nicotinate phosphoribosyltransferase (100%);Transferred entry: 6.3.4.21 | GO:0005829(0%) | cytosol(0%) | GO:0004516(100%);GO:000451 | nicotinate phosphoribosyltransferase activity (100%);nicotinate-nucleoti | GO:0009435(100%);GO:0034355(0%);GO:0008218(0%) | NAD biosynthetic process (100%);NAD salvage (0 | IPR007229(100%);IPR036068( | 1.00E+00 | 1.00E+00 | 6.75E+04 | NS | NS | NS | NS | NS | NS | NS | NS |

| | | | | | | | | | | | | | | | | | | | | | | | | | | | | | | | | | | | | | |
|---|---|---|---|---|---|---|---|---|---|---|---|---|---|---|---|---|---|---|---|---|---|---|---|---|---|---|---|---|---|---|---|---|---|---|---|---|---|
| | | | | | | | | | | | | | | | | | | ) | (0%) | | | 4(99%);GO:0016757(1%) | de diphosphorylase (carboxylating) activity (99%);glycosyltransferase activity (1%) | | %);bioluminescence (0%) | 99%);IPR041525(99%) | | | | | | | | | | | |
| 406 | 464 Variable | QHFEESSLR | Enterobacterales | Bacteria | Proteobacteria | | Gammaproteobacteria | | Enterobacterales | | | | | | | | | 3.1.21.3(100%) | Type I site-specific deoxyribonuclease | GO:0005829(0%) | cytosol (0%) | GO:0009035(98% | type I site-specific deoxyribonuclease | GO:0006304(97%);GO:000930 | DNA modification (97%);D | IPR013670(100% | 1.00E+00 | 1.00E+00 | 1.78E+05 | NS | NS | NS | NS | NS | NS | NS | NS |

| | | | | | | | | | | | | | | | | | | | | | | | | | | | | | | | | | | | | | | | | |
|---|---|---|---|---|---|---|---|---|---|---|---|---|---|---|---|---|---|---|---|---|---|---|---|---|---|---|---|---|---|---|---|---|---|---|---|---|---|---|---|---|
| | | | | | | | | | | | | | | | | | | | (100%) | | | );GO:0003677(97%);GO:0005524(95%) | activity(98%);DNA binding(97%);ATP binding(95%) | 7(0%);GO:0016032(0%) | NA restriction-modification system(0%);viral process(0%) | );IPR001650(99%);IPR027417(99%) | | | | | | | | | | | | | | |
| 407 | Variable 465 | YLQLHPEDTEK | Enterobacterales | Bacteria | Proteobacteri | | Gammaproteobacte | | Enterobacterales | | | | | | | | | | | | | | | | | IPR041256(10 | 1.00E+00 | 1.00E+00 | 2.01E+05 | 1.8468 | 0.000008845 | 0.0023692 | 0.9 | NS | NS | NS | NS | 4.7421 | 0.013903 | 0.0014234 |

| | | | | | | | | | | | | | | | | | | | | | | | | | | | | | | | | | | | | | |
|---|---|---|---|---|---|---|---|---|---|---|---|---|---|---|---|---|---|---|---|---|---|---|---|---|---|---|---|---|---|---|---|---|---|---|---|---|---|
| | | | | | a | | ria | | | | | | | | | | | | | | | | | | | 0%) | | | | | | | | | | | |
| 409 | Variable 467 | CDGKPCVK | Enterobacterales | Bacteria | Proteobacteria | | Gammaproteobacteria | | Enterobacterales | | | | | | | | | | | GO:0016021(100%) | integral component of membrane (100%) | GO:0000155(46%) | phosphorelay sensor kinase activity (46%) | | | IPR003661(100%) | 1.00E+00 | 1.00E+00 | 6.08E+04 | NS | NS | NS | NS | NS | NS | NS | NS |
| 410 | Variable 468 | RPPQLSGGQLQR | Enterobacterales | Bacteria | Proteobacteria | | Gammaproteobacteria | | Enterobacterales | | | | | | | | | 7.2.2.11(93%);3.6.3.24 | ABC-type Ni(2+) transporter (93%);Transferred entry: 7.2.2.11 | GO:0005886(98%);GO:001 | plasma membrane (98%);membrane (0 | GO:0005524(100%);GO:00 | ATP binding (100%);ABC-type nickel transporter activity | GO:0006974(0%);GO:0010045(0%);GO:0035544 | cellular response to DNA damage stimulus (0% | IPR003439(100%);IPR027 | 1.00E+00 | 1.00E+00 | 5.24E+04 | NS | NS | NS | NS | NS | NS | NS | NS |

| | | | | | | | | | | | | | | | | | | | | | | | | | | | | | | | | | | | | | |
|---|---|---|---|---|---|---|---|---|---|---|---|---|---|---|---|---|---|---|---|---|---|---|---|---|---|---|---|---|---|---|---|---|---|---|---|---|---|
| | | | | | | | | | | | | | | | | | | (5%);3.6.3.-(3%) | (5%); Acting on acid anhydrides; catalyzing transmembrane movement of substances (3%) | 6020(0%) | %) | 15413(98%);GO:0016151(95%) | (98%);nickel cation binding (95%) | 4(0%) | );response to nickel cation (0%); nickel cation transmembrane transport (0%) | 417(100%);IPR003593(98%) | | | | | | | | | | | |
| 412 | Variabl | EAVVTGLR | Actinomyc | Bacteri | Actinoba | | Actinomyce | | | | | | | | | | | | | GO:0005 | plasmam | GO:0005 | ATP binding (3%) | GO:0009058(3 | biosynthetic pr | IPR0048 | 1.00E+0 | 1.00E+0 | 6.37E+0 | 3.0454 | 0.0094 | 0.00714 | 0.9 | NS | NS | NS | NS |

| | | | | | | | | | | | | | | | | | | | | | | | | | | | | | | | | | | | | | |
|---|---|---|---|---|---|---|---|---|---|---|---|---|---|---|---|---|---|---|---|---|---|---|---|---|---|---|---|---|---|---|---|---|---|---|---|---|---|
| | e 47 | | etia | a | cteria | | tia | | | | | | | | | | | | | 886(97%);GO:0016021(97%) | brane(97%);integral component of membrane(97%) | 524(3%);GO:0016874(3%) | ;ligase activity(3%) | %) | ocess(3%) | 69(97%);IPR000731(95%);IPR027463(29%) | 0 | 0 | 5 | | 144 | 1 | | | | | |
| 414 | Variable | AEMSEYLFDK | Enterobacter | Bacteria | Proteoba | | Gammaprote | | Enterobacteral | | | | | | | | | | | GO:00058 | cytosol(0%);nuc | GO:00036 | DNA binding(100% | GO:0006310(100 | DNA recombina | DNA-templ | 1.00E+00 | 2.17E+05 | 1.00E+00 | NS | NS | NS | NS | NS | NS | NS | NS |

| | | | | | | | | | | | | | | | | | | | | | | | | | | | | | | | | | | | | | | | | |
|---|---|---|---|---|---|---|---|---|---|---|---|---|---|---|---|---|---|---|---|---|---|---|---|---|---|---|---|---|---|---|---|---|---|---|---|---|---|---|---|---|
| | 471 | | ales | | cteria | | obacteria | | es | | | | | | | | | | | 29(0%);GO:0009295(0%);GO:0032993(0%) | leoid(0%);protein-DNA complex(0%) | 77(100%);GO:0000976(0%);GO:0001216(0%) | );transcription cis-regulatory region binding(0%);DNA-binding transcription activator activity(0%) | %);GO:0006417(100%);GO:0006355(99%) | tion(100%);regulation of translation(100%);regulation of transcription | ated(99%) | | | | | | | | | | | | | | |
| 418 | Variabl | HGTLAAVAPLPR | Enterobact | Bacteri | Proteob | | Gammaprot | | Enterobacter | | Enterobacteri | | | | | | | 1.8.-.-(3 | Acting on a sulfur gro | GO:0016 | integral compone | GO:0051 | 4 iron | 4 sulfur cluster bin | GO:0019645(1 | anaerobic | 1.00E+0 | 1.00E+0 | 1.88E+0 | NS | NS | NS | NS | NS | NS | NS | NS | 6.1206 | 0.00464 | 0.00014 |

| | | | | | | | | | | | | | | | | | | | | | | | | | | | | | | | | | | | | | | | | |
|---|---|---|---|---|---|---|---|---|---|---|---|---|---|---|---|---|---|---|---|---|---|---|---|---|---|---|---|---|---|---|---|---|---|---|---|---|---|---|---|---|
| | e 475 | | eriaceae | a | acteria | | eobacteria | | ales | | aceae | | | | | | | 8%);1.8.5.3(38%);1.9.7.1.9(38%) | up of donors (38%);Respiratory dimethylsulfoxide reductase (38%);Selenate reductase (38%) | 021(1%) | nt of membrane (1%) | 539(97%);GO:0046872(96%);GO:0016491(3%) | | ding (97%);metal ion binding (96%);oxidoreductase activity (3%) | %) | electron transport chain (1%) | 0 | 0 | 5 | | | | | | | | | | 87 | 454 |
| 419 | Variab | HNVGATPR | Enteroba | Bacter | Proteo | | Gammapr | | Enterobact | | Enterobacte | | | | | | | 2.4.1.- | Hexosyltransferase | GO:001 | integral compo | GO:001 | transferase activit | GO:0017000 | antibiotic bios | IPR010 | 1.00E+ | 1.00E+ | 1.14E+ | 0.3327 | 1.7539 | 0.00178 | 0.9 | NS | NS | NS | NS | 20.313 | 6.7405 | 0.0008 |

| | | | | | | | | | | | | | | | | | | | | | | | | | | | | | | | | | | | | | | | |
|---|---|---|---|---|---|---|---|---|---|---|---|---|---|---|---|---|---|---|---|---|---|---|---|---|---|---|---|---|---|---|---|---|---|---|---|---|---|---|---|
| | le 476 | | cteriaceae | ia | bacteria | | oteobacteria | | erales | | riaceae | | | | | | | (75%);2.4.1.17(17%);2.4.1.278(8%) | s (75%); Glucuronosyltransferase (17%); 3-alpha-mycarosylerythronolide B desosaminyltransferase (8%) | 6021(21%) | nent of membrane (21%) | 6740(79%);GO:0016757(17%);GO:0015020(4%) | y (79%);glycosyltransferase activity (17%);glucuronosyltransferase activity (4%) | (100%) | ynthetic process (100%) | 610(100%) | 00 | 00 | 05 | | E-10 | 13 | | | | | | E-07 | 3542 |
| 420 | Varia | AGNYNSAEPK | Enterob | Bacte | Prote | | Gamma | | Enteroba | | Enterobac | | | | | | | | | GO:00 | cell outerm | GO:00 | fimbrial usher | GO:000929 | pilusassem | IPR00 | 1.00E | 1.00E | 4.99E | 3.607 | 0.002 | 0 | 0.9 | NS | NS | NS | NS | | |

| | | | | | | | | | | | | | | | | | | | | | | | | | | | | | | | | | | | | | |
|---|---|---|---|---|---|---|---|---|---|---|---|---|---|---|---|---|---|---|---|---|---|---|---|---|---|---|---|---|---|---|---|---|---|---|---|---|---|
| | ble 477 | | acteriaceae | ria | obacteria | | proteobacteria | | cterales | | teriaceae | | | | | | | | | 09279(100%);GO:0016021(100%) | embrane(100%);integral component of membrane(100%) | 15473(100%) | porin activity(100%) | 7(100%) | bly(100%) | 0015(100%);IPR025885(99%);IPR037224(98%) | +00 | +00 | +04 | 8 | 5789 | | | | | | |
| 421 | Varia | DLATMTPDQA | Enterob | Bacte | Prote | | Gamma | | Enteroba | | Enterobac | | | | | | | 1.1.1 | 1-deoxy-D-xyl | | | GO:00 | 1-deoxy-D- | GO:001928 | isopenteny | methyl | 1.00E | 1.00E | 4.49E | 3.446 | 0.000 | 0.0000 | 0.9 | NS | NS | NS | NS |

| | | | | | | | | | | | | | | | | | | | | | | | | | | | | | | | | |
|---|---|---|---|---|---|---|---|---|---|---|---|---|---|---|---|---|---|---|---|---|---|---|---|---|---|---|---|---|---|---|---|---|
| | ble 478 | CR | acteriaceae | ria | obacteria | | proteobacteria | | cterales | | teriaceae | | | | | | | .267(100%);2.7.4.-(1%) | ulose-5-phosphate reductoisomerase(100%);Phosphotransferases with a phosphate group as acceptor(1%) | | | 30604(100%);GO:0046872(98%);GO:0070402(97%) | xylulose-5-phosphate reductoisomerase activity(100%);metal ion binding(98%);NADPH binding(97%) | 8(97%);GO:0016114(96%);GO:0008299(2%) | l diphosphate biosynthetic process | erythritol 4-phosphate pathway(97%);terpenoid biosynthetic p | +00 | +00 | +04 | 7 | 57695 | 62745 |

| | | | | | | | | | | | | | | | | | | | | | | | | | | | | | | | | | | | | | |
|---|---|---|---|---|---|---|---|---|---|---|---|---|---|---|---|---|---|---|---|---|---|---|---|---|---|---|---|---|---|---|---|---|---|---|---|---|---|
| | | | | | | | | | | | | | | | | | | | | | | | | | | rocess (96%); isoprenoid biosynthetic process (2%) | | | | | | | | | | | |
| 422 | Variable | LSAGSPGAALALFQGDNWQ | Enterobacter | Bacteria | Proteoba | | Gammaprote | | Enterobacteral | | Enterobacteriac | | | | | | | 2.7.7.7( | DNA-directed DNA pol | GO:00093 | DNA polymeras | GO:00038 | DNA-directed DN | GO:0006260 (100 | DNA replication | IPR008892 | 1.00E+00 | 1.00E+00 | 7.36E+04 | NS | NS | NS | NS | NS | NS | NS | NS |

| | | | | | | | | | | | | | | | | | | | | | | | | | | | | | | | | | | | | | |
|---|---|---|---|---|---|---|---|---|---|---|---|---|---|---|---|---|---|---|---|---|---|---|---|---|---|---|---|---|---|---|---|---|---|---|---|---|---|
| | 479 | AR | iaceae | | cteria | | obacteria | | es | | eae | | | | | | | 100%) | ymerase (100%) | 60(100%) | e III complex (100%) | 87(100%);GO:0003677(100%);GO:0008408(95%) | A polymerase activity (100%);DNA binding (100%);3'-5' exonuclease activity (95%) | %);GO:00062261 (0%) | (100%); DNA-dependent DNA replication (0%) | 1(100%);IPR015199(100%);IPR027417(98%) | | | | | | | | | | | |
| 423 | Vari | MQAAFFPTSR | Acidot | Bact | Actin | | Actino | | Acidot | | Acidoth | Acido | | | Acido | | Acido | 2.1. | Aspartate car | | | GO:0 | aspartate car | GO:0006 | 'de novo | IPR0 | 1.00 | 1.00 | 5.20 | 3.77 | 0.00 | 0.000 | 0.9 | NS | NS | NS | NS |

| | | | | | | | | | | | | | | | | | | | | | | | | | | | | | | | | |
|---|---|---|---|---|---|---|---|---|---|---|---|---|---|---|---|---|---|---|---|---|---|---|---|---|---|---|---|---|---|---|---|---|
| | able 48 | | hermuscellulolyticus11B | eria | obacteria | | mycetia | | hermales | | ermaceae | thermus | | | thermuscellulolyticus | | thermuscellulolyticus11B | 3.2(100%) | bamoyltransferase(100%) | | | 0040700(100%);GO:0016597(100%) | bamoyltransferase activity(100%);amino acid binding(100%) | 207(100%);GO:0006520(100%);GO:0044205(100%) | 'pyrimidine nucleobase biosynthetic process(100%);cellular amino acid metabolic process(1 | 02082(100%);IPR006130(100%);IPR006131(100%) | E+00 | E+00 | E+04 | 74 | 1801 | 16667 |

| | | | | | | | | | | | | | | | | | | | | | | | | | | | | | | | | | | | | | |
|---|---|---|---|---|---|---|---|---|---|---|---|---|---|---|---|---|---|---|---|---|---|---|---|---|---|---|---|---|---|---|---|---|---|---|---|---|---|
| | | | | | | | | | | | | | | | | | | | | | | | | | 00%);'de novo' UMP biosynthetic process (100%) | | | | | | | | | | | | |
| 424 | Variable 480 | EPLATLWR | Enterobacteriaceae | Bacteria | Proteobacteria | | Gammaproteobacteria | | Enterobacterales | | Enterobacteriaceae | | | | | | | 2.7.1.-(100%) | Phosphotransferases with an alcohol group as acceptor | | | GO:0016301(100%);GO: | kinase activity (100%);ATP binding (94%) | GO:0009244(94%);GO:0009103(2%);GO: | lipopolysaccharide core region biosynthetic | IPR011009(100%);IPR0 | 1.00E+00 | 1.00E+00 | 4.87E+05 | 19.417 | 0.026871 | 0 | 0.9 | NS | NS | NS | NS |

| | | | | | | | | | | | | | | | | | | | | | | | | | | |
|---|---|---|---|---|---|---|---|---|---|---|---|---|---|---|---|---|---|---|---|---|---|---|---|---|---|---|
| | | | | | | | | | | | | | | | | | | | (100%) | | | 0005524 (94%) | | 0046835 (1%) | process (94%);lipopolysaccharide biosynthetic process (2%);carbohydrate phosphorylation (1 | 17172 (98%) |

| | | | | | | | | | | | | | | | | | | | | | | | | | | | | | | | | | | | | | |
|---|---|---|---|---|---|---|---|---|---|---|---|---|---|---|---|---|---|---|---|---|---|---|---|---|---|---|---|---|---|---|---|---|---|---|---|---|---|
| | | | | | | | | | | | | | | | | | | | | | | | | | %) | | | | | | | | | | | | |
| 426 | Variable 482 | QHQAAELVEEQ | Enterobacteriaceae | Bacteria | Proteobacteria | | Gammaproteobacteria | | Enterobacterales | | Enterobacteriaceae | | | | | | | | | GO:0005886(96%);GO:0016021(96%);GO:0005887( | plasma membrane(96%);integral component of membrane(96%);integral component | GO:0022857(96%);GO:0015125(4%);GO:0015385(4 | transmembrane transporter activity(96%);bile acid transmembrane transporter activity(4%);sodium:proton antipo | GO:0015721(4%);GO:0030641(4%);GO:0036376(4%) | bile acid and bile salt transport(4%);regulation of cellular pH(4%);sodium ion export across | IPR011701(100%);IPR036259(100%);IPR005829(9 | 1.00E+00 | 1.00E+00 | 1.31E+05 | 4.0739 | 0.0097716 | 0.00018466 | 0.9 | NS | NS | NS | NS |

| | | | | | | | | | | | | | | | | | | | | | | | | | | | | | | | | | | | | | |
|---|---|---|---|---|---|---|---|---|---|---|---|---|---|---|---|---|---|---|---|---|---|---|---|---|---|---|---|---|---|---|---|---|---|---|---|---|---|
| | | | | | | | | | | | | | | | | | | | | 4%) | of plasma membrane (4%) | %) | rter activity (4%) | | plasma membrane (4%) | 6%) | | | | | | | | | | | |
| 427 | Variable 483 | EGNFEADK | Enterobacteriaceae | Bacteria | Proteobacteria | | Gammaproteobacteria | | Enterobacterales | | Enterobacteriaceae | | | | | | | 2.7.-.-(78%);3.1.6.-(29%) | Transferring phosphorus-containing groups (78%); Sulfuric ester hydrolases (29%) | GO:0005886(97%);GO:0016021(97% | plasma membrane (97%);integral component of membrane | GO:0008484(97%);GO:0016772(97% | sulfuric ester hydrolase activity (97%);transferase activity | transferring phosphorus-containing groups (97%);transferase activity (3 | GO:0009244(0%) | lipopolysaccharide core region biosynthetic | 1.00E+00 | 1.00E+00 | 3.84E+04 | NS | NS | NS | NS | NS | NS | NS | NS |

| | | | | | | | | | | | | | | | | | | | | | | | | | | | | | | | | | | | | | |
|---|---|---|---|---|---|---|---|---|---|---|---|---|---|---|---|---|---|---|---|---|---|---|---|---|---|---|---|---|---|---|---|---|---|---|---|---|---|
| | | | | | | | | | | | | | | | | | | | | ) | (97%) | );GO:0016740(3%) | | %) | | process(0%) | | | | | | | | | | | |
| 429 | Variable 485 | ANAPSGYVR | Enterobacteriaceae | Bacteria | Proteobacteria | | Gammaproteobacteria | | Enterobacterales | | Enterobacteriaceae | | | | | | | 2.4.1.129(100%);2.4.2.-(0%) | Peptidoglycan glycosyltransferase(100%);Pentosyltransferases(0%) | GO:0005886(100%);GO:0009274(9 | plasma membrane(100%);peptidoglycan-based cell wall(9 | GO:0008955(98%);GO:0016763(98 | peptidoglycan glycosyltransferase activity(98%);pentosyltransferase activity(98%) | GO:0009252(100%);GO:0008360(96%);GO:0071555(96%) | peptidoglycan biosynthetic process(100%);regulation of cell sh | IPR023346(100%);IPR001264(100 | 1.00E+00 | 1.00E+00 | 8.54E+04 | NS | NS | NS | NS | NS | NS | NS | NS |

| | | | | | | | | | | | | | | | | | | | | | | | | | | | | | | | | | | | | | |
|---|---|---|---|---|---|---|---|---|---|---|---|---|---|---|---|---|---|---|---|---|---|---|---|---|---|---|---|---|---|---|---|---|---|---|---|---|---|
| | | | | | | | | | | | | | | | | | | | | 8%);GO:0016021(98%) | 8%);integral component of membrane(98%) | %);GO:0016757(0%) | ;glycosyltransferase activity(0%) | | ape(96%);cell wall organization(96%) | %);IPR011812(100%) | | | | | | | | | | | |
| 430 | Variable 486 | ESAEVFDTK | Enterobacteriaceae | Bacteria | Proteobacteria | | Gammaproteobacteria | | Enterobacterales | | Enterobacteriaceae | | | | | | | 6.1.1.9(100%) | Valine--tRNA ligase(100%) | GO:0005737(98%);GO:000 | cytoplasm(98%);cytosol(0%) | GO:0004832(100%);GO:00 | valine-tRNA ligase activity(100%);ATP binding(100%);a | GO:0006438(100%);GO:0006418(0%) | valyl-tRNA aminoacylation(100%);tRNA a | IPR002300(100%);IPR009 | 1.00E+00 | 1.00E+00 | 4.83E+05 | 22.111 | 0.0069144 | 0.00015466 | 0.9 | NS | NS | NS | NS |

| | | | | | | | | | | | | | | | | | | | | | | | | | | | | | | | | | | | | | |
|---|---|---|---|---|---|---|---|---|---|---|---|---|---|---|---|---|---|---|---|---|---|---|---|---|---|---|---|---|---|---|---|---|---|---|---|---|---|
| | | | | | | | | | | | | | | | | | | | | 5829(0%) | | 05524(100%);GO:0002161(100%) | minoacyl-tRNA editing activity (100%) | | minoacylation for protein translation (0%) | 008(100%);IPR002303(100%) | | | | | | | | | | | |
| 431 | Variable 487 | HMDGYEFNDGDEDTPMWSR | Enterobacteriaceae | Bacteria | Proteobacteria | | Gammaproteobacteria | | Enterobacterales | | Enterobacteriaceae | | | | | | | | | GO:0009279(96%);G | cell outer membrane (96%);integ | GO:0015473(100%); | fimbrial usher porin activity (100%);heparin bin | GO:0009297 (100%);GO:0043711 (0 | pilus assembly (100%); pilus organi | IPR000015(100%);I | 1.00E+00 | 1.35E+05 | 1.00E+00 | NS | NS | NS | NS | NS | NS | NS | NS |

| | | | | | | | | | | | | | | | | | | | | | | | | | | | | | | | | | | | | | |
|---|---|---|---|---|---|---|---|---|---|---|---|---|---|---|---|---|---|---|---|---|---|---|---|---|---|---|---|---|---|---|---|---|---|---|---|---|---|
| | | | | | | | | | | | | | | | | | | | | O:0016021(96%);GO:0016020(4%) | ral component of membrane (96%); membrane (4%) | GO:0008201(0%) | ding (0%) | %) | zation (0%) | PR042186(93%);IPR018030(93%) | | | | | | | | | | | |
| 432 | Variable 488 | HTTDGVVK | Enterobacteriaceae | Bacteria | Proteobacteria | | Gammaproteobacteria | | Enterobacterales | | Enterobacteriaceae | | | | | | | | | GO:0042597(100%) | periplasmic space (100%) | | | GO:0008643 (100%) | carbohydrate transport (100%) | IPR010794(100%) | 1.00E+00 | 2.36E+05 | 1.00E+00 | NS | NS | NS | NS | NS | NS | NS | NS |

| | | | | | | | | | | | | | | | | | | | | | | | | | | | | | | | | | | | | | |
|---|---|---|---|---|---|---|---|---|---|---|---|---|---|---|---|---|---|---|---|---|---|---|---|---|---|---|---|---|---|---|---|---|---|---|---|---|---|
| | | | | | | | | | | | | | | | | | | | | ) | | | | | | | | | | | | | | | | | |
| 433 | Variable 489 | NFASLSLPGLSEER | Enterobacteriaceae | Bacteria | Proteobacteria | | Gammaproteobacteria | | Enterobacterales | | Enterobacteriaceae | | | | | | | 3.6.1.11(100%) | Exopolyphosphatase(100%) | GO:0005886(93%) | plasmamembrane(93%) | GO:0004309(100%);GO:0016462(0%);GO:0042802(0 | exopolyphosphatase activity(100%);pyrophosphatase activity(0%);identical protein binding(0%) | GO:0006793(93%);GO:0006798(0%) | phosphorus metabolic process(93%);polyphosphate catabolic process(0%) | IPR003695(100%);IPR043129(100%);IPR030673(93 | 1.00E+00 | 1.21E+05 | 1.00E+00 | NS | NS | NS | NS | NS | NS | NS | NS |

| | | | | | | | | | | | | | | | | | | | | | | | | | | | | | | | | | | | | | |
|---|---|---|---|---|---|---|---|---|---|---|---|---|---|---|---|---|---|---|---|---|---|---|---|---|---|---|---|---|---|---|---|---|---|---|---|---|---|
| | | | | | | | | | | | | | | | | | | | | | | %) | | | | %) | | | | | | | | | | | |
| 434 | Variable 49 | QGEVVMFK | Acidothermus cellulolyticus 11B | Bacteria | Actinobacteria | | Actinomycetia | | Acidothermales | | Acidothermaceae | Acidothermus | | | Acidothermus cellulolyticus | | Acidothermus cellulolyticus 11B | | | GO:0000502(100%) | proteasome complex(100%) | GO:0005524(100%);GO:0016887(100%) | ATP binding(100%);ATPase(100%) | GO:0010498(100%);GO:0019941(100%) | proteasomal protein catabolic process(100%); modification-dependent protein catabolic | IPR003593(100%);IPR003959(100%);IPR003960(1 | 1.00E+00 | 1.00E+00 | 3.23E+04 | 2.0684 | 0.0445 | 0.000095651 | 0.9 | NS | NS | NS | NS |

| | | | | | | | | | | | | | | | | | | | | | | | | | | | | | | | | | | | | | |
|---|---|---|---|---|---|---|---|---|---|---|---|---|---|---|---|---|---|---|---|---|---|---|---|---|---|---|---|---|---|---|---|---|---|---|---|---|---|
| | | | | | | | | | | | | | | | | | | | | | | | | | process (100%) | 00%) | | | | | | | | | | | |
| 435 | Variable 490 | AQAMSAAVEQAQTPWTLDPVAVGALDYR | Escherichia coli | Bacteria | Proteobacteria | | Gammaproteobacteria | | Enterobacterales | | Enterobacteriaceae | Escherichia | | | Escherichia coli | | | 2.7.1.50(100%) | Hydroxyethylthiazole kinase (100%) | | | GO:0000287(100%);GO:0004417(100%);GO: | magnesium ion binding (100%);hydroxyethylthiazole kinase activity (100%);ATP binding (100%) | GO:0009228(100%);GO:0009229(100%) | thiamine biosynthetic process (100%);thiamine diphosphate biosynthetic pr | IPR000417(100%);IPR029056(100%) | 1.00E+00 | 1.00E+00 | 2.88E+04 | NS | NS | NS | NS | NS | NS | NS | NS |

| | | | | | | | | | | | | | | | | | | | | | | | | | | | | | | | | | | | | | | | | |
|---|---|---|---|---|---|---|---|---|---|---|---|---|---|---|---|---|---|---|---|---|---|---|---|---|---|---|---|---|---|---|---|---|---|---|---|---|---|---|---|---|
| | | | | | | | | | | | | | | | | | | | | | | 0005524(100%) | | | ocess(100%) | | | | | | | | | | | | | | | |
| 438 | Variable 493 | QQETLLNEFATSDAR | Salmonella enterica | Bacteria | Proteobacteria | | Gammaproteobacteria | | Enterobacterales | | Enterobacteriaceae | Salmonella | | | Salmonella enterica | | | 2.1.1.176(100%) | 16S rRNA (cytosine(967)-C(5))-methyltransferase (100%) | GO:0005737(99%);GO:0005829(0%) | cytoplasm(99%);cytosol(0%) | GO:0003723(100%);GO:0016434(95% | RNA binding(100%);rRNA (cytosine) methyltransferase activity(95%);rRNA methy | GO:0006355(100%);GO:0032259(1%);GO:0001510(0%) | regulation of transcription | DNA-templated(100%);methylation(1%);RN | 1.00E+00 | 1.00E+00 | 4.10E+04 | 0.5158 | 4.8013E-08 | 0.00081708 | 0.9 | NS | NS | NS | NS | 8.6287 | 0.00072551 | 0.00010719 |

| | | | | | | | | | | | | | | | | | | | | | | | | | | | | | | | | | | | | | | | | |
|---|---|---|---|---|---|---|---|---|---|---|---|---|---|---|---|---|---|---|---|---|---|---|---|---|---|---|---|---|---|---|---|---|---|---|---|---|---|---|---|---|
| | | | | | | | | | | | | | | | | | | | | | | );GO:0008649(4%) | ltransferase activity (4%) | | | Amethylation (0%) | | | | | | | | | | | | | | |
| 439 | Variable 494 | SVYASATSPLTGFLEVSA | Salmonella enterica | Bacteria | Proteobacteria | | Gammaproteobacteria | | Enterobacterales | | Enterobacteriaceae | Salmonella | | | Salmonella enterica | | | 2.8.1.10 (100%) | Thiazole synthase (100%) | GO:0005737 (93%) | cytoplasm (93%) | GO:1990107 (100%) | thiazole synthase activity (100%) | GO:0009229 (100%) | thiamine diphosphate biosynthetic process (100%) | IPR008867 (100%); IPR013785 (100 | 1.00E+00 | 1.00E+00 | 5.14E+04 | 0.43674 | 2.6945E-09 | 0.0012043 | 0.9 | NS | NS | NS | NS | 13.715 | 0.00026044 | 0.00027743 |

| | | | | | | | | | | | | | | | | | | | | | | | | | | | | | | | | | | | | | |
|---|---|---|---|---|---|---|---|---|---|---|---|---|---|---|---|---|---|---|---|---|---|---|---|---|---|---|---|---|---|---|---|---|---|---|---|---|---|
| | | | | | | | | | | | | | | | | | | | | | | | | | | %);IPR033983(100%) | | | | | | | | | | | |
| 440 | Variable 495 | TETEFFHR | Salmonella enterica | Bacteria | Proteobacteria | | Gammaproteobacteria | | Enterobacterales | | Enterobacteriaceae | Salmonella | | | Salmonella enterica | | | 7.-.-.-(99%);3.6.3.-(1%) | Translocases(99%);Acting on acid anhydrides;catalyzing transmembrane mo | GO:0005886(98%);GO:0043190( | plasma membrane(98%);ATP-binding cassette(A | GO:0005524(100%);GO:0016887 | ATP binding(100%);ATPase(79%);cobalt ion transmembrane transporter | GO:0009236(78%);GO:0006824(20%) | cobalamin biosynthetic process(78%);cobalt ion transpo | IPR027417(100%);IPR003439(9 | 1.00E+00 | 6.50E+04 | 1.00E+00 | NS | NS | NS | NS | NS | NS | NS | NS |

| | | | | | | | | | | | | | | | | | | | | | | | | | | | | | | | | | | | | | |
|---|---|---|---|---|---|---|---|---|---|---|---|---|---|---|---|---|---|---|---|---|---|---|---|---|---|---|---|---|---|---|---|---|---|---|---|---|---|
| | | | | | | | | | | | | | | | | | | | vement of substances (1%) | 0%) | BC) transporter complex (0%) | (79%); GO:0015087(78%) | activity (78%) | | rt (20%) | 9%); IPR005876(98%) | | | | | | | | | | | |
| 441 | Variable 496 | VSAAEGDYGEESEH | Salmonella enterica | Bacteria | Proteobacteria | | Gammaproteobacteria | | Enterobacterales | | Enterobacteriaceae | Salmonella | | | Salmonella enterica | | | | | GO:0005737(97%);GO:00058 | cytoplasm (97%);cytosol (1%) | GO:0008237(100%);GO:0016 | metallopeptidase activity (100%);transferase activity (1%);amino | GO:0051090 (1%) | regulation of DNA-binding transcription factor activ | IPR024079(100%);IPR010138 | 1.00E+00 | 7.10E+04 | 1.00E+00 | NS | NS | NS | NS | NS | NS | NS | NS |

| | | | | | | | | | | | | | | | | | | | | | | | | | | | | | | | | | | | | | |
|---|---|---|---|---|---|---|---|---|---|---|---|---|---|---|---|---|---|---|---|---|---|---|---|---|---|---|---|---|---|---|---|---|---|---|---|---|---|
| | | | | | | | | | | | | | | | | | | | | 29(1%) | | 740(1%);GO:0004177(1%) | peptidase activity (1%) | | ity (1%) | 4(99%);IPR042252(99%) | | | | | | | | | | | |
| 442 | Variable 497 | QNSVNVNGTLK | Salmonella enterica | Bacteria | Proteobacteria | | Gammaproteobacteria | | Enterobacterales | | Enterobacteriaceae | Salmonella | | | Salmonella enterica | | | | | GO:0005887(100%);GO:000 | integral component of plasma membrane (100% | | | GO:0009306(100%);GO:0009405(0%) | protein secretion (100%); obsolete pathogenesis | IPR007452(100%);IPR0001 | 1.00E+00 | 1.17E+06 | 1.00E+00 | NS | NS | NS | NS | NS | NS | NS | NS |

| | | | | | | | | | | | | | | | | | | | | | | | | | | | | | | | | | | | | | |
|---|---|---|---|---|---|---|---|---|---|---|---|---|---|---|---|---|---|---|---|---|---|---|---|---|---|---|---|---|---|---|---|---|---|---|---|---|---|
| | | | | | | | | | | | | | | | | | | | | 9279(0%);GO:0097347(0%) | );cell outer membrane(0%);TAM protein secretion complex(0%) | | | | (0%) | 84(0%);IPR010827(0%) | | | | | | | | | | | |
| 443 | Variable 498 | TPAYSSLAAGELNGLVAR | Salmonella enterica | Bacteria | Proteobacteri | | Gammaproteobacte | | Enterobacterales | | Enterobacteriaceae | Salmonella | | | Salmonella enteric | | | 2.4.2.52(100 | Triphosphoribosyl-dephospho-Co | | | GO:0005524(1 | ATP binding(100%);triphosphoribo | GO:0016310(100%);GO:00 | phosphorylation(100%);pr | IPR002736(10 | 1.00E+00 | 1.37E+06 | 1.00E+00 | NS | NS | NS | NS | NS | NS | NS | NS |

| | | | | | | | | | | | | | | | | | | | | | | | | | | | | | | | | | | | | | |
|---|---|---|---|---|---|---|---|---|---|---|---|---|---|---|---|---|---|---|---|---|---|---|---|---|---|---|---|---|---|---|---|---|---|---|---|---|---|
| | | | | | a | | ria | | | | | | | | a | | | %) | A synthase (100%) | | | 00%);GO:0046917(100%);GO:0016757(79%) | syl-dephospho-CoA synthase activity (100%);glycosyltransferase activity (79%) | 51191 (1%) | osthetic group biosynthetic process (1%) | 0%);IPR017551(98%);IPR005551(0%) | | | | | | | | | | | |
| 444 | Variable | AEALAMGFTHAACGPCV | Salmonellae | Bacteria | Proteoba | | Gammaprote | | Enterobacteral | | Enterobacteriac | Salmonella | | | Salmonellae | Salmonell | | 2.8.1.8( | Lipoyl synthase (100%) | GO:00057 | cytoplasm (100% | GO:00169 | lipoate synthase activit | 4 sulfur cluster bindin | GO:0009249 (100 | proteinlipo | 1.00E+00 | 2.18E+05 | 1.00E+00 | NS | NS | NS | NS | NS | NS | NS | NS |

| | | | | | | | | | | | | | | | | | | | | | | | | | | | | | | | | | | | | | |
|---|---|---|---|---|---|---|---|---|---|---|---|---|---|---|---|---|---|---|---|---|---|---|---|---|---|---|---|---|---|---|---|---|---|---|---|---|---|
| | 499 | R | ntericasubsp.enterica | | cteria | | obacteria | | es | | eae | | | | nterica | aentericasubsp.enterica | | 100%) | | 37(100%) | ) | 92(100%);GO:0046872(100%);GO:0051539(100%) | y(100%);metalionbinding(100%);4iron | g(100%) | %) | ylation(100%) | | | | | | | | | | | |
| 445 | Var | DVTGVPP | Actin | Bac | Acti | | | | | | | | | | | | | 7.6 | ABC-typ | GO: | ATP- | GO: | ATPbin | | | IPR | 2.0 | 1.0 | 1.0 | NS | NS | NS | NS | NS | NS | NS | NS |

| | | | | | | | | | | | | | | | | | | | | | | | | | | | | | |
|---|---|---|---|---|---|---|---|---|---|---|---|---|---|---|---|---|---|---|---|---|---|---|---|---|---|---|---|---|---|
| | iable 5 | NR | obacteria | teria | nobacteria | | | | | | | | | | | | | .2.11(100%) | e polyamine transporter (100%) | 0043190(88%);GO:0005886(13%) | binding cassette (ABC) transporter complex (88%);plasma membrane (13%) | 0005524(100%);GO:0015594(88%);GO:0016887(88%) | ding (100%);ABC-type putrescine transporter activity (88%);ATPase (88%) | | | 003439(100%);IPR003593(100%);IPR005893(100%) | 7E+06 | 0E+00 | 0E+00 |

| | | | | | | | | | | | | | | | | | | | | | | | | | | | | | | | | | | | | | |
|---|---|---|---|---|---|---|---|---|---|---|---|---|---|---|---|---|---|---|---|---|---|---|---|---|---|---|---|---|---|---|---|---|---|---|---|---|---|
| 446 | Variable 50 | VFAKPEVHGHR | Bifidobacterium | Bacteria | Actinobacteria | | Actinomycetia | | Bifidobacteriales | | Bifidobacteriaceae | Bifidobacterium | | | | | | 2.1.2.-(100%) | Hydroxymethyl- | formyl-and related transferases (100%) | | | GO:0005524 (100%);GO:0000287 (94%);GO:0004644 (94%) | ATP binding (100%);magnesium ion binding (94%);phosphoribosylglycinamide formyltransferase activity (94%) | GO:0006189 (94%); GO:0006164 (6%) | 'de novo' IMP biosynthetic process (94%); purine nucleotide bios | 1.00E+00 | 1.00E+00 | 7.55E+04 | NS | NS | NS | NS | NS | NS | NS | NS |

| | | | | | | | | | | | | | | | | | | | | | | | | | | | | | | | | | | | | | |
|---|---|---|---|---|---|---|---|---|---|---|---|---|---|---|---|---|---|---|---|---|---|---|---|---|---|---|---|---|---|---|---|---|---|---|---|---|---|
| | | | | | | | | | | | | | | | | | | | | | | | | | | ynthetic process (6%) | | | | | | | | | | | |
| 447 | Variable 500 | MGDFAEPR | Salmonella enterica subsp. enterica serovar P | Bacteria | Proteobacteria | | Gammaproteobacteria | | Enterobacterales | | Enterobacteriaceae | Salmonella | | | Salmonella enterica | Salmonella enterica subsp. enter | Salmonella enterica subsp. enterica serov | | | | | GO:0003684(100%);GO:0005524(1 | damaged DNA binding(100%);ATP binding(100%);mismatched DNA bin | GO:0006298(100%) | mismatch repair(100%) | IPR000432(100%);IPR005748(100 | 1.00E+00 | 1.00E+00 | 7.96E+04 | NS | NS | NS | NS | NS | NS | NS | NS |

| | | | | | | | | | | | | | | | | | | | | | | | | | | | | | | | | | | | | | |
|---|---|---|---|---|---|---|---|---|---|---|---|---|---|---|---|---|---|---|---|---|---|---|---|---|---|---|---|---|---|---|---|---|---|---|---|---|---|
| | | | aratyphiC str.RKS4594 | | | | | | | | | | | | | ica | arParatyphiC str.RKS4594 | | | | | 00%);GO:0030983(100%) | ding(100%) | | | %);IPR007695(100%) | | | | | | | | | | | |
| 448 | Variable 501 | FSGGCNGCSMVQK | Buchnera aphidicola str.Bp(Baizo | Bacteria | Proteobacteria | | Gammaproteobacteria | | Enterobacterales | | Erwiniaceae | Buchnera | | | Buchnera aphidicola | | Buchnera aphidicola str.Bp(Baiz | | | | | GO:0005506(100%);GO:0051 | iron ion binding(100%);4 iron | 4 sulfur cluster binding(100%) | GO:0016226(100%);GO:0051604(100%) | iron-sulfur cluster assembly(100 | 1.00E+00 | 1.00E+00 | 9.33E+04 | 5.3922 | 0.02103 | -0.000035714 | 0.9 | NS | NS | NS | NS |

| | | | | | | | | | | | | | | | | | | | | | | | | | | | | | | | | | | | | | |
|---|---|---|---|---|---|---|---|---|---|---|---|---|---|---|---|---|---|---|---|---|---|---|---|---|---|---|---|---|---|---|---|---|---|---|---|---|---|
| | | | ngiapistaciae) | | | | | | | | | | | | | | ongiapistaciae) | | | | | 539(100%) | | | | %); protein maturation (100%) | | | | | | | | | | | |
| 449 | Variable 502 | SVVQGEVLVK | Yersinia pseudotuberculosis compl | Bacteria | Proteobacteria | | Gammaproteobacteria | | Enterobacterales | | Yersiniaceae | Yersinia | Yersinia pseudotuberculosi | | | | | | | GO:0005887(100%);GO:0033 | integral component of plasma membrane (100%); | GO:0008320(100%) | protein transmembrane transporter activity (100%) | GO:0043953 (100%) | protein transport by the Tat complex (100%) | IPR018448(100%) | 1.00E+00 | 1.00E+00 | 4.70E+04 | NS | NS | NS | NS | NS | NS | NS | NS |

| | | | | | | | | | | | | | | | | | | | | | | | | | | | | | | | | | | | | | |
|---|---|---|---|---|---|---|---|---|---|---|---|---|---|---|---|---|---|---|---|---|---|---|---|---|---|---|---|---|---|---|---|---|---|---|---|---|---|
| | | | ex | | | | | | | | | | scomplex | | | | | | | 281(100%) | TAT protein transport complex(100%) | | | | | | | | | | | | | | | | |
| 450 | Variable 503 | TFVAETVK | [Haemophilus] ducreyi 35000HP | Bacteria | Proteobacteria | | Gammaproteobacteria | | Pasteurellales | | Pasteurellaceae | Haemophilus | | | [Haemophilus] ducreyi | | [Haemophilus] ducreyi 35000HP | | | GO:0005840(100%) | ribosome(100%) | GO:0003735(100%) | structural constituent of ribosome(100%) | GO:0006412(100%) | translation(100%) | IPR005822(100%);IPR005823 | 1.00E+00 | 1.00E+00 | 4.50E+04 | 2.4867 | 0.0038738 | 0.00011949 | 0.9 | NS | NS | NS | NS |

| | | | | | | | | | | | | | | | | | | | | | | | | | | | | | | | | | | | | | |
|---|---|---|---|---|---|---|---|---|---|---|---|---|---|---|---|---|---|---|---|---|---|---|---|---|---|---|---|---|---|---|---|---|---|---|---|---|---|
| | | | | | | | | | | | | | | | | | | | | | | | | | | (100%);IPR023563(100%) | | | | | | | | | | | |
| 451 | Variable 504 | AYDNATAAVAK | Pseudomonas | Bacteria | Proteobacteria | | Gammaproteobacteria | | Pseudomonadales | | Pseudomonadaceae | Pseudomonas | | | | | | 3.4.21.50(100%) | Lysylendopeptidase (100%) | GO:0005576(4%);GO:0005576 | extracellular region(4%);extracellular space (4%) | GO:0008233(78%);GO:0016 | peptidase activity(78%);hydrolase activity(13%);serine-typ | GO:0009405(9%);GO:0006508(4%);GO:0006554 | obsolete pathogenesis(9%);proteolysis(4%);lys | IPR009003(100%);IPR0435 | 1.00E+00 | 1.00E+00 | 6.40E+05 | 1.7359 | 0.01426 | 0 | 0.9 | NS | NS | NS | NS |

| | | | | | | | | | | | | | | | | | | | | | | | | | | | | | | | | | | | | | | | | |
|---|---|---|---|---|---|---|---|---|---|---|---|---|---|---|---|---|---|---|---|---|---|---|---|---|---|---|---|---|---|---|---|---|---|---|---|---|---|---|---|---|
| | | | | | | | | | | | | | | | | | | | | 15(4%) | | 787(13%);GO:0008236(9%) | epeptidase activity (9%) | (4%) | ine catabolic process (4%) | 04(100%) | | | | | | | | | | | | | | |
| 452 | Variable 505 | EAHWGNYPCR | Pseudomonas | Bacteria | Proteobacteria | | Gammaproteobacteria | | Pseudomonadales | | Pseudomonadaceae | Pseudomonas | | | | | | | | | | GO:0003676(100%);GO:00 | nucleic acid binding (100%);nuclease activity (88%);endo | | | IPR003509(100%);IPR011 | 1.00E+00 | 1.00E+00 | 2.87E+04 | 0.50962 | 1.2661E-08 | 0.0013622 | 0.9 | NS | NS | NS | NS | 10.472 | 0.0002042 | 0.0026916 |

| | | | | | | | | | | | | | | | | | | | | | | | | | | | | | | | | | | | | | |
|---|---|---|---|---|---|---|---|---|---|---|---|---|---|---|---|---|---|---|---|---|---|---|---|---|---|---|---|---|---|---|---|---|---|---|---|---|---|
| | | | | | | | | | | | | | | | | | | | | | | 04518(88%);GO:0004519(12%) | nuclease activity (12%) | | | 335(100%);IPR011856(100%) | | | | | | | | | | | |
| 453 | Variable 506 | NDSMSQLGDR | Vibrio | Bacteria | Proteobacteria | | Gammaproteobacteria | | Vibrionales | | Vibrionaceae | Vibrio | | | | | | | | GO:0005737(100%);GO | cytoplasm(100%);plasma membrane | | | | | IPR007451(100%);IPR | 1.00E+00 | 1.00E+00 | 6.56E+04 | NS | NS | NS | NS | NS | NS | NS | NS |

| | | | | | | | | | | | | | | | | | | | | | | | | | | | | | | | | | | | | | | | | |
|---|---|---|---|---|---|---|---|---|---|---|---|---|---|---|---|---|---|---|---|---|---|---|---|---|---|---|---|---|---|---|---|---|---|---|---|---|---|---|---|---|
| | | | | | | | | | | | | | | | | | | | | :0005886(100%) | (100%) | | | | | 0359332(100%) | | | | | | | | | | | | | | |
| 454 | Variable 507 | SEQTESVLANPNPAED | Vibrio | Bacteria | Proteobacteria | | Gammaproteobacteria | | Vibrionales | | Vibrionaceae | Vibrio | | | | | | 5.4.99.18(100%) | 5-(carboxyamino)imidazole ribonucleotide mutase (100%) | | | GO:0034023(100%) | 5-(carboxyamino)imidazole ribonucleotide mutase activity (100%) | GO:0006189(100%) | 'de novo' IMP biosynthetic process (100%) | IPR000031(100%);IPR024694(100% | 1.00E+00 | 1.00E+00 | 4.06E+04 | 0.61929 | 1.7817E-08 | 0.0012426 | 0.9 | NS | NS | NS | NS | 9.7858 | 0.00032453 | 0.0010032 |

| | | | | | | | | | | | | | | | | | | | | | | | | | | | | | | | | | | | | | | | | |
|---|---|---|---|---|---|---|---|---|---|---|---|---|---|---|---|---|---|---|---|---|---|---|---|---|---|---|---|---|---|---|---|---|---|---|---|---|---|---|---|---|
| | | | | | | | | | | | | | | | | | | | | | | | | | | );IPR033747(100%) | | | | | | | | | | | | | | |
| 455 | Variable 508 | MHCPFCQHSDTR | Xanthomonadaceae | Bacteria | Proteobacteria | | Gammaproteobacteria | | Xanthomonadales | | Xanthomonadaceae | | | | | | | | | | | GO:0005524(100%);GO:0008270( | ATP binding(100%);zinc ion binding(100%);DNA binding(82%) | GO:0045892(100%) | negative regulation of transcription | DNA-templated(100%) | 1.00E+00 | 1.00E+00 | 2.92E+04 | 0.30695 | 6.0036E-12 | 0.0028215 | 0.9 | NS | NS | NS | NS | 37.303 | 4.8649E-10 | 0.0056456 |

| | | | | | | | | | | | | | | | | | | | | | | | | | | | | | | | | | | | | | | | | |
|---|---|---|---|---|---|---|---|---|---|---|---|---|---|---|---|---|---|---|---|---|---|---|---|---|---|---|---|---|---|---|---|---|---|---|---|---|---|---|---|---|
| | | | | | | | | | | | | | | | | | | | | | | 100%);GO:0003677(82%) | | | | | | | | | | | | | | | | | | |
| 456 | Variable 509 | GLPPALGLVK | Bdellovibriobacteriovorus | Bacteria | Proteobacteria | | Oligoflexia | | Bdellovibrionales | | Bdellovibrionaceae | Bdellovibrio | | | Bdellovibriobacteriovorus | | | 3.5.4.2(100%) | Adenine deaminase (100%) | | | GO:0000034(100%) | adenine deaminase activity (100%) | GO:0006146 (100%) | adenine catabolic process (100%) | IPR006679(100%);IPR006668 | 1.00E+00 | 1.00E+00 | 1.11E+05 | 0.1918 | 1.3481E-14 | 0.002553 | 0.9 | NS | NS | NS | NS | 41.639 | 1.0786E-10 | 0.004818 |

| | | | | | | | | | | | | | | | | | | | | | | | | | | | | | | | | | | | | | |
|---|---|---|---|---|---|---|---|---|---|---|---|---|---|---|---|---|---|---|---|---|---|---|---|---|---|---|---|---|---|---|---|---|---|---|---|---|---|
| | | | | | | | | | | | | | | | | | | | | | | | | | | 0(100%);IPR011059(100%) | | | | | | | | | | | |
| 457 | Variable 51 | FFAHQFAESLAK | Bifidobacterium longum | Bacteria | Actinobacteria | | Actinomycetia | | Bifidobacteriales | | Bifidobacteriaceae | Bifidobacterium | | | Bifidobacterium longum | | | 6.3.2.8(100%) | UDP-N-acetylmuramate--L-alanine ligase(100%) | GO:0005737(100%) | cytoplasm(100%) | GO:0005524(100%);GO:00 | ATP binding(100%);UDP-N-acetylmuramate-L-alanine lig | GO:0009252(100%);GO:0007049(93%);GO:00 | peptidoglycan biosynthetic process(100%);cell | IPR000713(100%);IPR004 | 1.00E+00 | 1.00E+00 | 1.12E+05 | NS | NS | NS | NS | NS | NS | NS | NS |

| | | | | | | | | | | | | | | | | | | | | | | | | | | | | | | | | | | | | | |
|---|---|---|---|---|---|---|---|---|---|---|---|---|---|---|---|---|---|---|---|---|---|---|---|---|---|---|---|---|---|---|---|---|---|---|---|---|---|
| | | | | | | | | | | | | | | | | | | | | | | 08763(100%) | ase activity (100%) | 08360(93%) | cycle (93%);regulation of cell shape (93%) | 101(100%);IPR005758(100%) | | | | | | | | | | | |
| 458 | Variable 510 | AVMTSDACR | Desulfatibacillum aliphaticivo | Bacteria | Proteobacteria | delta/epsilon subdi | Deltaproteobacteria | | Desulfobacterales | | Desulfobacteraceae | Desulfatibacillum | | | Desulfatibacillum aliphat | | | 2.7.7.38(100%) | 3-deoxy-manno-octulosonate cytidylyltransferase (100% | GO:0005737(100%) | cytoplasm(100%) | GO:0008690(100%) | 3-deoxy-manno-octulosonate cytidylyltransferase act | GO:0009103(100%);GO:0033468(100%) | lipopolysaccharide biosynthetic process (1 | IPR003329(100%);IPR0 | 1.00E+00 | 5.25E+04 | 1.00E+00 | NS | NS | NS | NS | NS | NS | NS | NS |

| | | | | | | | | | | | | | | | | | | | | | | | | | | | | | | | | | | | | | | | | |
|---|---|---|---|---|---|---|---|---|---|---|---|---|---|---|---|---|---|---|---|---|---|---|---|---|---|---|---|---|---|---|---|---|---|---|---|---|---|---|---|---|
| | | | rans | | | visions | | | | | | | | | icivorans | | | | ) | | | | ivity (100%) | | 00%); CMP-keto-3-deoxy-D-manno-octulosonic acid biosynthetic process (100%) | 04528(100%); IPR0290044(100%) | | | | | | | | | | | | | | |
| 459 | Vari | GVCDGENG | Desul | Bact | Prot | delt | Deltapr | | Desulfo | | Desulfo | Desu | | | | | | 6.3. | tRNA(Ile)- | GO:0 | cytoplas | GO:0 | ATP bindin | forming | GO:0006 | tRNA | 1.00 | 1.00 | 1.10 | 0.30 | 3.75 | 0.001 | 0.9 | NS | NS | NS | NS | 18.1 | 2.07 | 0.00 |

| | | | | | | | | | | | | | | | | | | | | | | | | | | | | | | | | | | | | | | | | |
|---|---|---|---|---|---|---|---|---|---|---|---|---|---|---|---|---|---|---|---|---|---|---|---|---|---|---|---|---|---|---|---|---|---|---|---|---|---|---|---|---|
| | able 511 | EGVR | fovibrio | eria | eobacteria | a/epsilon subdivisions | oteobacteria | | vibrionales | | vibrionaceae | lfovibrio | | | | | | 4.19(100%) | lysidine synthetase (100%) | 0057337(100%) | m(100%) | 0005524(100%);GO:0016879(100%) | g(100%);ligase activity | carbon-nitrogen bonds (100%) | 400(100%) | modification (100%) | E+00 | E+00 | E+05 | 921 | 22E-10 | 4046 | | | | | | 56 | 87E-06 | 081026 |
| 460 | Variable 512 | AEGDLEAAWAYAMSGTR | Anaeromyxobacterde | Bacteria | Proteobacteria | delta/epsilon s | Deltaproteobacteria | | Myxococcales | Cystobacterineae | Anaeromyxobacteraceae | Anaeromyxobacter | | | Anaeromyxobacterde | | Anaeromyxobacterde | 2.1.2.-(100%) | Hydroxymethyl- | formyl-and related tr | | | GO:0000287(100%);GO:0004644(10 | magnesium ion binding (100%);phosphori | GO:0006189(100%) | 'de novo' IMP biosynth | 1.00E+00 | 1.00E+00 | 1.19E+05 | 0.12816 | 1.381E-13 | 0.0021081 | 0.9 | NS | NS | NS | NS | 38.565 | 3.103E-10 | 0.0026029 |

| | | | | | | | | | | | | | | | | | | | | | | | | | | | | | | | | | | | | | |
|---|---|---|---|---|---|---|---|---|---|---|---|---|---|---|---|---|---|---|---|---|---|---|---|---|---|---|---|---|---|---|---|---|---|---|---|---|---|
| | | | halogenans2CP-C | | | ubdivisions | | | | | | | | | halogenans | | halogenans2CP-C | | | ansferases(100%) | | | 0%);GO:0005524(100%) | bosylglycinamideformyltransferase activity(100%);ATP binding(100%) | | etic process(100%) | | | | | | | | | | | |
| 461 | Variable 513 | LKPLLLEGAALAAEK | Syntrophobacter fumar | Bacteria | Proteobacteria | delta/epsilon su | Deltaproteobacteria | | Syntrophobacterales | | Syntrophobacteraceae | Syntrophobacter | | | Syntrophobacter fumar | | Syntrophobacter fumar | 2.7.7.4(100%) | Sulfate adenylyltransferase(100%) | | | GO:0004781(100%); | sulfate adenylyltransferase (ATP) activity(10 | GO:0000103(100%);GO:0070814(1 | sulfate assimilation(100%);hydrog | IPR002650(100%);I | 1.00E+00 | 9.69E+04 | 1.00E+00 | NS | NS | NS | NS | NS | NS | NS | NS |

| | | | | | | | | | | | | | | | | | | | | | | | | | | | | | | | | | | | | | | | | |
|---|---|---|---|---|---|---|---|---|---|---|---|---|---|---|---|---|---|---|---|---|---|---|---|---|---|---|---|---|---|---|---|---|---|---|---|---|---|---|---|---|
| | | | oxidansMPOB | | | bdivisions | | | | | | | | | oxidans | | oxidansMPOB | | | | | GO:0005524(100%) | 0%);ATP binding (100%) | 00%) | ensulfide biosynthetic process (100%) | PR014729(100%);IPR015947(100%) | | | | | | | | | | | | | | |
| 463 | Variable 515 | HLMLCTGDLGFSAAK | Campylobacter | Bacteria | Proteobacteria | delta/epsilon | Epsilonproteobacteria | | Campylobacterales | | Campylobacteraceae | Campylobacter | | | | | | 6.1.1.11(100%) | Serine--tRNA ligase (100%) | GO:0005737(99% | cytoplasm (99%) | GO:0004828(100 | serine-tRNA ligase activity (100%);A | GO:0006434(100%);GO:001626 | seryl-tRNA aminoacylation (1 | IPR002314(100% | 1.00E+00 | 1.00E+00 | 6.81E+04 | 0.20858 | 1.2614E-12 | 0.0030462 | 0.9 | NS | NS | NS | NS | 44.065 | 4.8558E-11 | 0.015384 |

| | | | | | | | | | | | | | | | | | | | | | | | | | | |
|---|---|---|---|---|---|---|---|---|---|---|---|---|---|---|---|---|---|---|---|---|---|---|---|---|---|---|
| | | | | | | subdivisions | | | | | | | | | | | | | | ) | | %);GO:0005524(100%) | TP binding (100%) | 0 (96%);GO:0097056(78%) | 00%);selenocysteine biosynthetic process(96%);selenocysteinyl-tRNA(Sec) biosynthetic proc | );IPR002317(100%);IPR033729(99%) |

| | | | | | | | | | | | | | | | | | | | | | | | | | | | | | | | | | | | | | | | | |
|---|---|---|---|---|---|---|---|---|---|---|---|---|---|---|---|---|---|---|---|---|---|---|---|---|---|---|---|---|---|---|---|---|---|---|---|---|---|---|---|---|
| | | | | | | | | | | | | | | | | | | | | | | | | | ess (78%) | | | | | | | | | | | | | | | |
| 464 | Variable 516 | DPSSATSNALAK | Campylobacter | Bacteria | Proteobacteria | delta/epsilon subdivisions | Epsilonproteobacteria | | Campylobacterales | | Campylobacteraceae | Campylobacter | | | | | | 4.2.1.151 (100%) | Chorismate dehydratase (100%) | | | GO:0016836 (96%);GO:0016829 (4%) | hydro-lyase activity (96%);lyase activity (4%) | GO:0009234 (100%) | menaquinone biosynthetic process (100%) | IPR003773 (100%);IPR030868 (100%) | 1.00E+00 | 1.00E+00 | 6.65E+04 | 15.175 | 9.0462E-09 | 0.00054892 | 0.9 | NS | NS | NS | NS | 22.249 | 2.5771E-07 | 0.0050392 |
| 465 | Variabl | ESGEVLFDSLTLLDESK | Campylob | Bacteri | Proteob | delta/e | Epsilonprot | | Campyloba | | Campylobact | Campylo | | | | | | 2.7.7.6 | DNA-directed RNA | | | GO:0003 | DNA binding (10 | GO:0006351 (1 | transcription | DNA-tem | 1.00E+0 | 1.00E+0 | 7.80E+0 | 3.2251 | 0.02148 | 0.00123 | 0.9 | NS | NS | NS | NS | | | |

| | | | | | | | | | | | | | | | | | | | | | | | | | | | | | | | | | | | | | |
|---|---|---|---|---|---|---|---|---|---|---|---|---|---|---|---|---|---|---|---|---|---|---|---|---|---|---|---|---|---|---|---|---|---|---|---|---|---|
| | e 517 | | acter | a | acteria | psilon subdivisions | eobacteria | | cterales | | eraceae | bacter | | | | | | (100%) | polymerase (100%) | | | 677(100%);GO:0003899(100%);GO:0032549(99%) | 0%);DNA-directed 5'-3' RNA polymerase activity (100%);ribonucleoside binding (99%) | 00%) | | plated (100%) | 0 | 0 | 4 | | 4 | 37 | | | | | |
| 466 | Var | QVGFDY | Cam | Bac | Pro | del | Epsi | | Cam | | Cam | Cam | | | Cam | | | 3.6 | Nucleoti | GO: | cytopl | GO: | NAD | GO:00 | nucleo | IPR | 1.0 | 1.0 | 5.6 | NS | NS | NS | NS | NS | NS | NS | NS |

| | | | | | | | | | | | | | | | | | | | | | | | | | | | | | | | | | | | | | |
|---|---|---|---|---|---|---|---|---|---|---|---|---|---|---|---|---|---|---|---|---|---|---|---|---|---|---|---|---|---|---|---|---|---|---|---|---|---|
| | iable 518 | DENLDK | pylobacterjejuni | teria | teobacteria | ta/epsilon subdivisions | lonproteobacteria | | pylobacterales | | pylobacteraceae | pylobacter | | | pylobacterjejuni | | | .1.9(100%) | dediphosphatase(100%) | 0005737(100%) | asm(100%) | 0035529(80%);GO:0036218(80%);GO:0047429(20%) | Hpyrophosphatase activity(80%);dTTP diphosphatase activity(80%);nucleoside-triphosphate diphosphatase activity(20%) | 09117(100%) | tidemetabolic process(100%) | 0036997(100%);IPR029001(100%) | 0E+00 | 0E+00 | 5E+04 | | | | | | | | |
| 4 | V | NN | H | B | P | d | E | | C | | H | H | | | | | | 2 | DN | G | D | G | D | G | D | I | 1 | 1 | 8 | 2 | 0 | 0 | 0 | N | N | N | N |

| | | | | | | | | | | | | | | | | | | | | | | | | | | | | | | | | | | | | | |
|---|---|---|---|---|---|---|---|---|---|---|---|---|---|---|---|---|---|---|---|---|---|---|---|---|---|---|---|---|---|---|---|---|---|---|---|---|---|
| 67 | ariable 519 | ALFESLDSEHSETAK | elicobacter | acteria | roteobacteria | elta/epsilon subdivisions | psilonproteobacteria | | ampylobacterales | | elicobacteraceae | elicobacter | | | | | | .7.7.7(100%) | A-directed DNA polymerase (100%) | O:0009360(100%);GO:0005737(100%) | NA polymerase III complex(100%);cytoplasm(100%) | O:0003677(100%);GO:0003887(100%);GO:0008408(1 | NA binding(100%);DNA-directed DNA polymerase activity(100%);3'-5' exonuclease activity(100%) | O:0006260(100%);GO:0006271(0%) | NA replication(100%);DNA strand elongation involved in DNA replication(0%) | PR001001(100%);IPR022635(100%);IPR022637(100% | .00E+00 | .00E+00 | .32E+04 | .8087 | .046392 | | .9 | S | S | S | S |

| | | | | | | | | | | | | | | | | | | | | | | | | | | | | | | | | | | | | | |
|---|---|---|---|---|---|---|---|---|---|---|---|---|---|---|---|---|---|---|---|---|---|---|---|---|---|---|---|---|---|---|---|---|---|---|---|---|---|
| | | | | | | | | | | | | | | | | | | | | | | 00%) | | | | ) | | | | | | | | | | | |
| 468 | Variable 52 | EYSDDVAEGK | Bifidobacterium longum | Bacteria | Actinobacteria | | Actinomycetia | | Bifidobacteriales | | Bifidobacteriaceae | Bifidobacterium | | | Bifidobacterium longum | | | 2.7.11.1(100%) | Non-specific serine/threonine protein kinase (100%) | GO:0016021(98%) | integral component of membrane (98%) | GO:0005524(100%);GO:0106310(100%);GO:0106 | ATP binding (100%);protein serine kinase activity (100%);protein threonine kinase activity (100%) | | | IPR000719(100%);IPR005543(100%);IPR008271 | 1.00E+00 | 1.00E+00 | 5.11E+05 | 85.657 | 0.005071 | 0.00016429 | 0.9 | NS | NS | NS | NS |

| | | | | | | | | | | | | | | | | | | | | | | | | | | | | | | | | | | | | | | | | |
|---|---|---|---|---|---|---|---|---|---|---|---|---|---|---|---|---|---|---|---|---|---|---|---|---|---|---|---|---|---|---|---|---|---|---|---|---|---|---|---|---|
| | | | | | | | | | | | | | | | | | | | | | | 311(100%) | | | | (100%) | | | | | | | | | | | | | | |
| 469 | Variable 520 | VTMPYLELLWNQFPK | Sphaerochaeta | Bacteria | Spirochaetes | | Spirochaetia | | Spirochaetales | | Sphaerochaetaceae | Sphaerochaeta | | | | | | | | | | GO:0003677(100%) | DNA binding(100%) | GO:0006355(100%) | regulation of transcription | DNA-templated(100%) | 1.00E+00 | 1.00E+00 | 2.16E+05 | 0.5439 | 9.0306E-07 | 0.0010643 | 0.9 | NS | NS | NS | NS | 6.3837 | 0.0037956 | 0.00044756 |
| 470 | Variable 521 | GYSPEFLAR | Treponema | Bacteria | Spirochaetes | | Spirochaetia | | Spirochaetales | | Treponemataceae | Treponema | | | | | | 2.7.7.-(75%);2.7.7. | Nucleotidyltransferases(75%);DNA primase DnaG | GO:1990077(80%) | primosome complex(80%) | GO:0003677(100%);G | DNA binding(100%);DNA primase activity | | | IPR002694(100%);IP | 1.00E+00 | 1.00E+00 | 4.31E+04 | NS | NS | NS | NS | NS | NS | NS | NS | | | |

| | | | | | | | | | | | | | | | | | | | | | | | | | | | | | | | | | | | | | | | |
|---|---|---|---|---|---|---|---|---|---|---|---|---|---|---|---|---|---|---|---|---|---|---|---|---|---|---|---|---|---|---|---|---|---|---|---|---|---|---|---|
| | | | | | | | | | | | | | | | | | 101(25%) | (25%) | | | O:0003896(100%);GO:0008270(100%) | (100%);zinc ion binding(100%) | | | R013264(100%);IPR036977(100%) | | | | | | | | | | | | | | |
| 471 | Variable 522 | EYDALAATYETDK | Treponema | Bacteria | Spirochaetes | | Spirochaetia | | Spirochaetales | | Treponemataceae | Treponema | | | | | | | | | | | | | IPR011004(10 | 1.00E+00 | 1.00E+00 | 3.77E+04 | 0.24195 | 3.9965E-12 | 0.0013564 | 0.9 | NS | NS | NS | NS | 37.473 | 4.576E-10 | 0.0070767 |

| | | | | | | | | | | | | | | | | | | | | | | | | | | | | | | | | | | | | | |
|---|---|---|---|---|---|---|---|---|---|---|---|---|---|---|---|---|---|---|---|---|---|---|---|---|---|---|---|---|---|---|---|---|---|---|---|---|---|
| | | | | | | | | | | | | | | | | | | | | | | | | | | 0%);IPR032533(100%) | | | | | | | | | | | |
| 472 | Variable 523 | EYYAGEEK | Treponema | Bacteria | Spirochaetes | | Spirochaetia | | Spirochaetales | | Treponemataceae | Treponema | | | | | | 5.2.1.8(100%) | Peptidylprolyl isomerase(100%) | GO:0005737(100%) | cytoplasm(100%) | GO:0003755(100%) | peptidyl-prolyl cis-trans isomerase activity(100%) | GO:0006457(100%);GO:0007049(100%);GO:0015031(1 | protein folding(100%);cell cycle(100%);protein trans | IPR005215(100%);IPR008880( | 1.00E+00 | 1.00E+00 | 4.04E+04 | NS | NS | NS | NS | NS | NS | NS | NS |

| | | | | | | | | | | | | | | | | | | | | | | | | | | | | | | | | | | | | | |
|---|---|---|---|---|---|---|---|---|---|---|---|---|---|---|---|---|---|---|---|---|---|---|---|---|---|---|---|---|---|---|---|---|---|---|---|---|---|
| | | | | | | | | | | | | | | | | | | | | | | | | 00%) | port(100%) | 100%);IPR008881(100%) | | | | | | | | | | | |
| 473 | Variable 524 | LCGDVDFDAVAHK | Treponema pallidum | Bacteria | Spirochaetes | | Spirochaetia | | Spirochaetales | | Treponemataceae | Treponema | | | Treponema pallidum | | | 1.5.1.5(100%);3.5.4.9(10 | Methylenetetrahydrofolate dehydrogenase (NADP(+))(100%);Methenyl | | | GO:0004477(100%);GO:0004 | methenyltetrahydrofolate cyclohydrolase activity(100%);methylenet | GO:0000105(100%);GO:0006164(100%);GO:000908 | histidine biosynthetic process(100%);purine nucl | IPR000672(100%);IPR020663 | 1.00E+00 | 1.00E+00 | 2.64E+04 | NS | NS | NS | NS | NS | NS | NS | NS |

| | | | | | | | | | | | | | | | | | | | | | | | | | | | | | | | | | | | | | |
|---|---|---|---|---|---|---|---|---|---|---|---|---|---|---|---|---|---|---|---|---|---|---|---|---|---|---|---|---|---|---|---|---|---|---|---|---|---|
| | | | | | | | | | | | | | | | | | | 0%) | tetrahydrofolate cyclohydrolase (100%) | | | 488(100%) | etrahydrofolate dehydrogenase (NADP+) activity (100%) | 6(100%) | eotide biosynthetic process (100%); methionine biosynthetic process (100%) | 0(100%); IPR020631(100%) | | | | | | | | | | | |
| 475 | Variab | FQGSVQAGR | Mycopla | Bacter | Teneric | | Mollicutes | | Mycoplasm | | Mycoplasm | Mycopla | | | | | | | | GO:000 | ribosome (8 | GO:000 | structural const | GO:0006412 | translation (1 | IPR000 | 1.00E+ | 1.00E+ | 5.51E+ | NS | NS | NS | NS | NS | NS | NS | NS |

| | | | | | | | | | | | | | | | | | | | | | | | | | | | | | | | | | | | | | |
|---|---|---|---|---|---|---|---|---|---|---|---|---|---|---|---|---|---|---|---|---|---|---|---|---|---|---|---|---|---|---|---|---|---|---|---|---|---|
| | le 526 | | sma | ia | utes | | | | atales | | ataceae | sma | | | | | | | | 5840(89%);GO:0022625(11%) | 9%);cytosolic large ribosomal subunit(11%) | 3735(100%);GO:0019843(100%) | ituent of ribosome(100%);rRNA binding(100%) | (100%) | 00%) | 597(100%);IPR009000(100%);IPR019927(100%) | 00 | 00 | 04 | | | | | | | | |
| 476 | Vari | VEELVLDL | Myco | Bact | Tene | | Mollic | | Mycopl | | Mycopla | Myco | | | | | | 7.3. | ABC-type | GO:0 | plasma | GO:0 | inorganic ph | GO:0035 | phosphat | IPR0 | 1.00 | 1.00 | 1.52 | NS | NS | NS | NS | NS | NS | NS | NS |

| | | | | | | | | | | | | | | | | | | | | | | | | | | | | | |
|---|---|---|---|---|---|---|---|---|---|---|---|---|---|---|---|---|---|---|---|---|---|---|---|---|---|---|---|---|---|
| | able 527 | K | plasma | eria | ricutes | | utes | | asmatales | | smataceae | plasma | | | | | | 2.1(100%) | phosphate transporter(100%) | 0058886(94%);GO:0016020(6%) | membrane(94%);membrane(6%) | 005315(100%);GO:0005524(100%);GO:0015415(94%) | osphate transmembrane transporter activity(100%);ATP binding(100%);ATPase-coupled phosphate ion transmembrane | 435(6%) | eion transmembrane transport(6%) | 03439(100%);IPR003593(100%);IPR005670(100%) | E+00 | E+00 | E+06 |

| | | | | | | | | | | | | | | | | | | | | | | | | | | | | | | | | | | | | | |
|---|---|---|---|---|---|---|---|---|---|---|---|---|---|---|---|---|---|---|---|---|---|---|---|---|---|---|---|---|---|---|---|---|---|---|---|---|---|
| | | | | | | | | | | | | | | | | | | | | | | | transporter activity (94%) | | | | | | | | | | | | | | |
| 477 | Variable 528 | FFLENPEEAAK | Mycoplasma | Bacteria | Tenericutes | | Mollicutes | | Mycoplasmatales | | Mycoplasmataceae | Mycoplasma | | | | | | 5.6.2.2(100%) | DNA topoisomerase (ATP-hydrolyzing) (100%) | GO:0005694(29%);GO:0005737(29%) | chromosome (29%);cytoplasm (29%) | GO:0003677(100%);GO:0003918(100%);GO | DNA binding (100%);DNA topoisomerase type II (double strand cut | ATP-hydrolyzing) activity (100%);ATP binding (100%) | GO:0006265(100%);GO:0006261(29%) | DNA topological change (100%);DNA-dependent | 1.00E+00 | 1.00E+00 | 1.10E+05 | NS | NS | NS | NS | NS | NS | NS | NS |

| | | | | | | | | | | | | | | | | | | | | | | | | | | | | | | | | | | | | | | | | |
|---|---|---|---|---|---|---|---|---|---|---|---|---|---|---|---|---|---|---|---|---|---|---|---|---|---|---|---|---|---|---|---|---|---|---|---|---|---|---|---|---|
| | | | | | | | | | | | | | | | | | | | | | | :0005524(100%) | | | | DNA replication (29%) | | | | | | | | | | | | | | |
| 478 | Variable 529 | AHLVEVAVEYDEELLMK | Mycoplasma | Bacteria | Tenericutes | | Mollicutes | | Mycoplasmatales | | Mycoplasmataceae | Mycoplasma | | | | | | | | GO:0005737(100%) | cytoplasm(100%) | GO:0003746(100%);GO:0003924(10 | translation elongation factor activity(100%);GTPase activity(100%);GTP binding(10 | | | IPR000640(100%);IPR000795(100% | 1.00E+00 | 1.00E+00 | 4.88E+04 | 9.1615 | 0.000010409 | 0.0001 | 0.9 | NS | NS | NS | NS | 7.8567 | 0.0012631 | 0.0031364 |

| | | | | | | | | | | | | | | | | | | | | | | | | | | | | | | | | | | | | | |
|---|---|---|---|---|---|---|---|---|---|---|---|---|---|---|---|---|---|---|---|---|---|---|---|---|---|---|---|---|---|---|---|---|---|---|---|---|---|
| | | | | | | | | | | | | | | | | | | | | | | 0%);GO:0005525(100%) | 0%) | | | );IPR004161(100%) | | | | | | | | | | | |
| 479 | Variable 53 | YFDTVLAR | Corynebacteriales | Bacteria | Actinobacteria | | Actinomycetia | | Corynebacteriales | | | | | | | | | 1.4.4.2(100%) | Glycine dehydrogenase (aminomethyl-transferring) (100%) | GO:0005618(1%);GO:0005829 | cell wall(1%);cytosol(1%);plasma membrane(1% | GO:0004375(100%);GO:00165 | glycine dehydrogenase (decarboxylating) activity (100%);glycine bin | GO:0019464(86%);GO:0006546(14%) | glycine decarboxylation via glycine cleavage syste | IPR015421(100%);IPR015424 | 1.00E+00 | 1.00E+00 | 4.41E+05 | 11.432 | 0.0471 | 0.00625 | 0.9 | NS | NS | NS | NS |

| | | | | | | | | | | | | | | | | | | | | | | | | | | | | | | | | | | | | | |
|---|---|---|---|---|---|---|---|---|---|---|---|---|---|---|---|---|---|---|---|---|---|---|---|---|---|---|---|---|---|---|---|---|---|---|---|---|---|
| | | | | | | | | | | | | | | | | | | | | (1%);GO:0005886(1%) | ) | 94(1%);GO:0030170(1%) | ding(1%);pyridoxal phosphate binding(1%) | | m(86%);glycine catabolic process(14%) | (100%);IPR020581(100%) | | | | | | | | | | | |
| 480 | Variable 530 | QASGSFGTR | Mycoplasma | Bacteria | Tenericutes | | Mollicutes | | Mycoplasmatales | | Mycoplasmataceae | Mycoplasma | | | | | | | | GO:0005840(75%);GO:0022 | ribosome(75%);cytosolic large ribosomal subu | GO:0003735(100%);GO:001 | structural constituent of ribosome(100%);rRNA binding(100% | GO:0000027(100%);GO:0006412(100%) | ribosomal large subunit assembly(100%);translat | IPR005813(100%);IPR0355 | 1.00E+00 | 4.97E+04 | 1.00E+00 | NS | NS | NS | NS | NS | NS | NS | NS |

| | | | | | | | | | | | | | | | | | | | | | | | | | | | | | | | | | | | | |
|---|---|---|---|---|---|---|---|---|---|---|---|---|---|---|---|---|---|---|---|---|---|---|---|---|---|---|---|---|---|---|---|---|---|---|---|---|
| | | | | | | | | | | | | | | | | | | | 625(25%) | nit(25%) | 9843(100%) | ) | | ion(100%) | 66(100%) | | | | | | | | | | | |
| 481 | Variable 531 | GALEGKPEWEAK | Mycoplasma | Bacteria | Tenericutes | | Mollicutes | | Mycoplasmatales | | Mycoplasmataceae | Mycoplasma | | | | | | | GO:0005737(80%) | cytoplasm(80%) | GO:0003746(100%);GO:0003924(100%);G | translation elongation factor activity(100%);GTPase activity(100%);GTP binding(100%) | | | IPR000795(100%);IPR004161(100%);IPR | 1.00E+00 | 2.32E+06 | 1.00E+00 | NS | NS | NS | NS | NS | NS | NS | NS |

| | | | | | | | | | | | | | | | | | | | | | | | | | | | | | | | | | | | | | |
|---|---|---|---|---|---|---|---|---|---|---|---|---|---|---|---|---|---|---|---|---|---|---|---|---|---|---|---|---|---|---|---|---|---|---|---|---|---|
| | | | | | | | | | | | | | | | | | | | | | | O:0005525(100%) | | | | 0090000(100%) | | | | | | | | | | | |
| 482 | Variable 532 | STVETVFTVLHAGGK | Mycoplasma | Bacteria | Tenericutes | | Mollicutes | | Mycoplasmatales | | Mycoplasmataceae | Mycoplasma | | | | | | 5.6.2.2(100%) | DNA topoisomerase (ATP-hydrolyzing) (100%) | GO:0005694(100%);GO:0005737(1 | chromosome(100%);cytoplasm(100%) | GO:0003677(100%);GO:0003918(1 | DNA binding(100%);DNA topoisomerase type II (double strand cut | ATP-hydrolyzing) activity(100%);ATP binding(100%) | GO:0006261(100%);GO:0006265(100%) | DNA-dependent DNA replication(100%);D | 3.91E+05 | 1.00E+00 | 1.00E+00 | NS | NS | NS | NS | NS | NS | NS | NS |

| | | | | | | | | | | | | | | | | | | | | | | | | | | | | | | | | | | | | | |
|---|---|---|---|---|---|---|---|---|---|---|---|---|---|---|---|---|---|---|---|---|---|---|---|---|---|---|---|---|---|---|---|---|---|---|---|---|---|
| | | | | | | | | | | | | | | | | | | | | 00%) | | 00%);GO:0005524(100%) | | | | NAtopological change(100%) | | | | | | | | | | | |
| 483 | Variable 533 | VAKPAAEDTK | Mycoplasma arthritidis 158L3-1 | Bacteria | Tenericutes | | Mollicutes | | Mycoplasmatales | | Mycoplasmataceae | Mycoplasma | | | Mycoplasma arthritidis | | Mycoplasma arthritidis 158L3- | | | GO:0015934(100%) | large ribosomal subunit (100%) | GO:0003735(100%);GO:001 | structural constituent of ribosome (100%);rRNA binding (100% | GO:0006412 (100%) | translation (100%) | IPR001063(100%);IPR0057 | 1.00E+00 | 1.00E+00 | 3.50E+04 | NS | NS | NS | NS | NS | NS | NS | NS |

| | | | | | | | | | | | | | | | | | | | | | | | | | | | | | | | | | | | | | |
|---|---|---|---|---|---|---|---|---|---|---|---|---|---|---|---|---|---|---|---|---|---|---|---|---|---|---|---|---|---|---|---|---|---|---|---|---|---|
| | | | | | | | | | | | | | | | | | 1 | | | | | 9843(100%) | ) | | | 27(100%);IPR036394(100%) | | | | | | | | | | | |
| 484 | Variable 534 | GAEGYAGFMR | Mycoplasmagallisepticum | Bacteria | Tenericutes | | Mollicutes | | Mycoplasmatales | | Mycoplasmataceae | Mycoplasma | | | Mycoplasmagallisepticum | | | | | GO:0005886(100%);GO:0 | plasmamembrane(100%);proton-transpo | catalyticcoreF(1)(100%) | GO:0005524(100%);GO:0046933(100%) | ATPbinding(100%);proton-transportingATPsynth | rotationalmechanism(100%) | | 1.00E+00 | 1.00E+00 | 7.38E+05 | NS | NS | NS | NS | NS | NS | NS | NS |

| | | | | | | | | | | | | | | | | | | | | | | | | | | | | | | | | | | | | | |
|---|---|---|---|---|---|---|---|---|---|---|---|---|---|---|---|---|---|---|---|---|---|---|---|---|---|---|---|---|---|---|---|---|---|---|---|---|---|
| | | | | | | | | | | | | | | | | | | | | 045261(100%) | rting ATP synthase complex | | | ase activity | | | | | | | | | | | | | |
| 485 | Variable 535 | WSNSATAASR | Mycoplasma gallisepticum | Bacteria | Tenericutes | | Mollicutes | | Mycoplasmatales | | Mycoplasmataceae | Mycoplasma | | | Mycoplasma gallisepticum | | | | | GO:0005840(100%) | ribosome(100%) | GO:0003735(100%);GO:0019843(100 | structural constituent of ribosome(100%);rRNA binding(100%) | GO:0006412(100%) | translation(100%) | IPR002583(100%);IPR036510(100%) | 1.00E+00 | 2.71E+05 | 1.00E+00 | NS | NS | NS | NS | NS | NS | NS | NS |

| | | | | | | | | | | | | | | | | | | | | | | | | | | | | | | | | | | | | | |
|---|---|---|---|---|---|---|---|---|---|---|---|---|---|---|---|---|---|---|---|---|---|---|---|---|---|---|---|---|---|---|---|---|---|---|---|---|---|
| | | | | | | | | | | | | | | | | | | | | | | %) | | | | | | | | | | | | | | | |
| 486 | Variable 536 | VEFADNSVLQAR | Mycoplasmagenitalium | Bacteria | Tenericutes | | Mollicutes | | Mycoplasmatales | | Mycoplasmataceae | Mycoplasma | | | Mycoplasmagenitalium | | | | | GO:0016021(100%);GO:0005886(6%) | integral component of membrane (100%);plasma membrane (6%) | | | GO:0007155 (6%) | cell adhesion (6%) | IPR007885 (100%) | 1.00E+00 | 1.00E+00 | 2.16E+04 | NS | NS | NS | NS | NS | NS | NS | NS |
| 487 | Variable 5 | ALQAGFDLSNK | Mycoplasmag | Bacteria | Tenericutes | | Mollicutes | | Mycoplasmatales | | Mycoplasmataceae | Mycoplasma | | | Mycoplasmag | | | | | GO:0016021 | integral component of m | | | GO:0020035 (5%) | cytoadherence to micr | mediated bys | 2.68E+05 | 1.00E+00 | 1.00E+00 | NS | NS | NS | NS | NS | NS | NS | NS |

| | | | | | | | | | | | | | | | | | | | | | | | | | | | | | | | | | | | | | |
|---|---|---|---|---|---|---|---|---|---|---|---|---|---|---|---|---|---|---|---|---|---|---|---|---|---|---|---|---|---|---|---|---|---|---|---|---|---|
| | 37 | | enitalium | | | | | | | | | | | | enitalium | | | | | (100%);GO:0005886(5%) | embrane(100%);plasmamembrane(5%) | | | | ovasculature | ymbiontprotein(5%) | | | | | | | | | | | |
| 489 | Variable 539 | QHDNANVLCLSSR | MycoplasmagenitaliumG37 | Bacteria | Tenericutes | | Mollicutes | | Mycoplasmatales | | Mycoplasmataceae | Mycoplasma | | | Mycoplasmagenitalium | | Mycoplasmagenitalium G37 | 5.3.1.6(100%) | Ribose-5-phosphate isomerase(100%) | | | GO:0004751(100%);GO:001 | ribose-5-phosphate isomerase activity(100%);isomerase activit | GO:0009052(100%);GO:0019316(100%) | pentose-phosphate shunt | non-oxidative branch(100%); | 1.00E+00 | 1.00E+00 | 7.31E+05 | NS | NS | NS | NS | NS | NS | NS | NS |

| | | | | | | | | | | | | | | | | | | | | | | | | | | | | | | | | | | | | | |
|---|---|---|---|---|---|---|---|---|---|---|---|---|---|---|---|---|---|---|---|---|---|---|---|---|---|---|---|---|---|---|---|---|---|---|---|---|---|
| | | | | | | | | | | | | | | | | | | | | | | 6853(100%) | y(100%) | | | D-allose catabolic process(100%) | | | | | | | | | | | |
| 490 | Variable 54 | SLFAMFLEAAAAESASR | Corynebacterium | Bacteria | Actinobacteria | | Actinomycetia | | Corynebacteriales | | Corynebacteriaceae | Corynebacterium | | | | | | | | GO:0005886(100%);GO: | plasma membrane(100%);proton-trans | catalytic core F(1)(100% | GO:0005524(100%);GO:0046933(100%) | ATP binding(100%);proton-transporting ATP sy | rotational mechanism(100%) | GO:0042777(100%) | 1.00E+00 | 1.00E+00 | 1.26E+05 | NS | NS | NS | NS | NS | NS | NS | NS |

| | | | | | | | | | | | | | | | | | | | | | | | | | | | | | | | | | | | | | | | | |
|---|---|---|---|---|---|---|---|---|---|---|---|---|---|---|---|---|---|---|---|---|---|---|---|---|---|---|---|---|---|---|---|---|---|---|---|---|---|---|---|---|
| | | | | | | | | | | | | | | | | | | | | 0045261(100%) | porting ATP synthase complex | ) | | nthase activity | | | | | | | | | | | | | | | | |
| 491 | Variable 540 | YCDSLGEFEK | Mycoplasma genitalium G37 | Bacteria | Tenericutes | | Mollicutes | | Mycoplasmatales | | Mycoplasmataceae | Mycoplasma | | | Mycoplasma genitalium | | Mycoplasma genitalium G37 | | | GO:0005737(100%) | cytoplasm(100%) | | | GO:0007049(100%);GO:0007059(100%);GO:0051301(100%) | cell cycle(100%);chromosome segregation(100%);cell divisi | IPR003768(100%) | 1.00E+00 | 1.00E+00 | 7.40E+04 | 17.712 | 1.2597E-07 | 0.00041493 | 0.9 | NS | NS | NS | NS | 14.517 | 0.00016124 | 0.0019885 |

| | | | | | | | | | | | | | | | | | | | | | | | | | | | | | | | | | | | | | |
|---|---|---|---|---|---|---|---|---|---|---|---|---|---|---|---|---|---|---|---|---|---|---|---|---|---|---|---|---|---|---|---|---|---|---|---|---|---|
| | | | | | | | | | | | | | | | | | | | | | | | | | on (100%) | | | | | | | | | | | | |
| 492 | Variable 541 | NQNFTESNK | Mycoplasmagenitalium G37 | Bacteria | Tenericutes | | Mollicutes | | Mycoplasmatales | | Mycoplasmataceae | Mycoplasma | | | Mycoplasmagenitalium | | Mycoplasmagenitalium G37 | 3.1.-.-(100%) | Acting on ester bonds (100%) | GO:0005886(100%);GO:0016021(100%) | plasma membrane (100%);integral component of membrane (100%) | GO:0003723(100%);GO:0004521(100%) | RNA binding (100%);endoribonuclease activity (100%) | GO:0006402 (100%) | mRNA catabolic process (100%) | IPR003607(100%);IPR006674(100%);IPR006677 | 1.00E+00 | 1.00E+00 | 5.03E+04 | NS | NS | NS | NS | NS | NS | NS | NS |

| | | | | | | | | | | | | | | | | | | | | | | | | | | | | | | | | | | | | | |
|---|---|---|---|---|---|---|---|---|---|---|---|---|---|---|---|---|---|---|---|---|---|---|---|---|---|---|---|---|---|---|---|---|---|---|---|---|---|
| | | | | | | | | | | | | | | | | | | | | | | | | | | 5(100%) | | | | | | | | | | | |
| 494 | Variable 543 | YENEGFPGYHWAK | Mycoplasmagenitalium G37 | Bacteria | Tenericutes | | Mollicutes | | Mycoplasmatales | | Mycoplasmataceae | Mycoplasma | | | Mycoplasmagenitalium | | Mycoplasmagenitalium G37 | | | GO:0005886(100%);GO:0016021(100%) | plasma membrane(100%);integral component of membrane(100%) | | | | | | 1.00E+00 | 1.00E+00 | 4.74E+04 | NS | NS | NS | NS | NS | NS | NS | NS |
| 495 | Var | SLNSFEAK | Myc | Bac | Ten | | Molli | | Myco | | Mycop | Myc | | | Myc | | Myc | | | | | GO: | ATP bin | | | IPR | 1.0 | 1.0 | 5.0 | 3.7 | 0.0 | 0.00 | 0.9 | NS | NS | NS | NS |

| | | | | | | | | | | | | | | | | | | | | | | | | | | | | | | | | | | | | | |
|---|---|---|---|---|---|---|---|---|---|---|---|---|---|---|---|---|---|---|---|---|---|---|---|---|---|---|---|---|---|---|---|---|---|---|---|---|---|
| | iable 544 | | oplasmagenitaliumG37 | teria | ericutes | | cutes | | plasmatales | | lasmataceae | oplasma | | | oplasmagenitalium | | oplasmagenitalium G37 | | | | | 0005524(100%);GO:0016887(100%) | ding(100%);ATPase(100%) | | | 003439(100%);IPR003593(100%);IPR017871(100%) | 0E+00 | 0E+00 | 7E+04 | 169 | 052126 | 012941 | | | | | |
| 4 | V | LM | M | B | T | | M | | M | | M | M | | | M | | | | | | | G | AT | G | pr | I | 1 | 5 | 1 | N | N | N | N | N | N | N | N |

| | | | | | | | | | | | | | | | | | | | | | | | | | | | | | | | | | | | | | |
|---|---|---|---|---|---|---|---|---|---|---|---|---|---|---|---|---|---|---|---|---|---|---|---|---|---|---|---|---|---|---|---|---|---|---|---|---|---|
| 96 | ariable 545 | GTSQTVHLNNK | ycoplasmahominis | acteria | enericutes | | ollicutes | | ycoplasmatales | | ycoplasmataceae | ycoplasma | | | ycoplasmahominis | | | | | | | O:0005524(100%);GO:0016887(83%);GO:0051082(33 | P binding(100%);ATPase(83%);unfolded protein binding(33%) | O:0006457(33%) | otein folding(33%) | PR013126(100%);IPR018181(100%);IPR029047(100% | .00E+00 | .81E+04 | .00E+00 | S | S | S | S | S | S | S | S |

| | | | | | | | | | | | | | | | | | | | | | | | | | | | | | | | | | | | | | | | | |
|---|---|---|---|---|---|---|---|---|---|---|---|---|---|---|---|---|---|---|---|---|---|---|---|---|---|---|---|---|---|---|---|---|---|---|---|---|---|---|---|---|
| | | | | | | | | | | | | | | | | | | | | | | %) | | | | ) | | | | | | | | | | | | | | |
| 497 | Variable 546 | VFNEVQNCK | Mycoplasmahyopneumoniae | Bacteria | Tenericutes | | Mollicutes | | Mycoplasmatales | | Mycoplasmataceae | Mycoplasma | | | Mycoplasmahyopneumoniae | | | 2.1.1.-(100%) | Methyltransferases (100%) | GO:0005737(100%) | cytoplasm (100%) | GO:0070043(83%);GO:0008649(17%) | rRNA (guanine-N7-)-methyltransferase activity (83%);rRNA methyltransferase activity (17%) | | | IPR003682(100%);IPR029063(100%) | 1.00E+00 | 1.00E+00 | 6.62E+05 | NS | NS | NS | NS | NS | NS | NS | NS | | | |
| 499 | Variable | GVYQDLEFLDENR | Mycoplasm | Bacteria | Tenericut | | Mollicutes | | Mycoplasmatal | | Mycoplasmatac | Mycoplasm | | | Mycoplasm | | Mycoplasm | 3.6.5.0( | -1 | GO:00058 | plasmamembr | GO:00037 | translationelongation | GO:0045727(100 | positive regulatio | IPR00064 | 1.00E+00 | 1.00E+00 | 7.99E+04 | 0.36415 | 3.048E- | 0.001456 | 0.9 | NS | NS | NS | NS | 15.219 | 0.00010 | 0.00909 |

| | | | | | | | | | | | | | | | | | | | | | | | | | | | | | | | | | | | | | | | |
|---|---|---|---|---|---|---|---|---|---|---|---|---|---|---|---|---|---|---|---|---|---|---|---|---|---|---|---|---|---|---|---|---|---|---|---|---|---|---|---|
| | 548 | | amobile163K | | es | | | | es | | eae | a | | | amobile | | amobile163K | 100%) | | 86(100%) | ane(100%) | 46(100%);GO:0003924(100%);GO:0005525(100%) | factor activity (100%);GTPase activity (100%);GTP binding (100%) | %) | n of translation (100%) | 0(100%);IPR000795(100%);IPR004161(100%) | | | | | 09 | | | | | | | 686 | 69 |
| 500 | Var | YSDLFFLE | Myc | Bac | Ten | | Molli | | Myco | | Mycop | Myc | | | Myc | | Myc | 6.5 | DNAlig | | | GO: | DNA | GO:00 | DNA | IPR | 1.0 | 1.0 | 1.0 | 6.0 | 0.0 | 0.00 | 0.9 | NS | NS | NS | NS | | |

| | | | | | | | | | | | | | | | | | | | | | | | | | | | | | | | | | | | | | |
|---|---|---|---|---|---|---|---|---|---|---|---|---|---|---|---|---|---|---|---|---|---|---|---|---|---|---|---|---|---|---|---|---|---|---|---|---|---|
| | iable 549 | K | oplasmamobile163K | teria | ericutes | | cutes | | plasmatales | | lasmataceae | oplasma | | | oplasmamobile | | oplasmamobile163K | .1.2(100%) | ase(NAD(+))(100%) | | | 0003911(100%);GO:0046872(100%) | ligase(NAD+) activity(100%);metal ion binding(100%) | 06260(100%);GO:0006281(100%) | replication(100%);DNA repair(100%) | 001357(100%);IPR001679(100%);IPR004150(100%) | 0E+00 | 0E+00 | 4E+05 | 956 | 0073652 | 025336 | | | | | |
| 5 | V | CQ | C | B | A | | A | | C | | C | C | | | | | | 2 | Thi | G | cy | G | thi | G | thi | I | 1 | 1 | 4 | 1 | 0 | 0 | 0 | N | N | N | N |

| | | | | | | | | | | | | | | | | | | | | | | | | | | | | | | | | | | | | | |
|---|---|---|---|---|---|---|---|---|---|---|---|---|---|---|---|---|---|---|---|---|---|---|---|---|---|---|---|---|---|---|---|---|---|---|---|---|---|
| 01 | ariable 55 | DPVAMATAMR | orynebacterium | acteria | ctinobacteria | | ctinomycetia | | orynebacteriales | | orynebacteriaceae | orynebacterium | | | | | | .8.1.10(100%) | azole synthase (100%) | O:0005737(100%) | toplasm (100%) | O:1990107(100%) | azole synthase activity (100%) | O:0009229 (100%) | amine diphosphate biosynthetic process (100%) | PR008867(100%); IPR013785(100%); IPR033983(100% | .00E+00 | .00E+00 | .78E+05 | 3.551 | .012796 | | .9 | S | S | S | S |

| | | | | | | | | | | | | | | | | | | | | | | | | | | | | | | | | | | | | | | | | |
|---|---|---|---|---|---|---|---|---|---|---|---|---|---|---|---|---|---|---|---|---|---|---|---|---|---|---|---|---|---|---|---|---|---|---|---|---|---|---|---|---|
| | | | | | | | | | | | | | | | | | | | | | | | | | | ) | | | | | | | | | | | | | | |
| 502 | Variable 550 | TSLQYLFASVGPK | Mycoplasmamobile163K | Bacteria | Tenericutes | | Mollicutes | | Mycoplasmatales | | Mycoplasmataceae | Mycoplasma | | | Mycoplasmamobile | | Mycoplasmamobile163K | | | GO:0005840(100%) | ribosome(100%) | GO:0003735(100%) | structural constituent of ribosome(100%) | GO:0006412(100%) | translation(100%) | IPR000456(100%);IPR036373(100%) | 1.00E+00 | 1.00E+00 | 2.12E+05 | NS | NS | NS | NS | NS | NS | NS | NS | 10.612 | 0.0018604 | 0.0007441 |
| 503 | Variable 551 | AGHSQEMLK | Mycoplasmamobi | Bacteria | Tenericutes | | Mollicutes | | Mycoplasmatales | | Mycoplasmataceae | Mycoplasma | | | Mycoplasmamob | | Mycoplasmamob | 2.1.1.199(10 | 16S rRNA(cytosine(1402)-N(4)) | GO:0005737(1 | cytoplasm(100%) | GO:0071424(1 | rRNA(cytosine-N4-)-methyltra | GO:0070475(100%) | rRNA base methylation(1 | IPR002903(10 | 1.00E+00 | 1.00E+00 | 9.27E+04 | 5.3309 | 0.00531 | 0.001257 | 0.9 | NS | NS | NS | NS | | | |

| | | | | | | | | | | | | | | | | | | | | | | | | | | | | | | | | | | | | | |
|---|---|---|---|---|---|---|---|---|---|---|---|---|---|---|---|---|---|---|---|---|---|---|---|---|---|---|---|---|---|---|---|---|---|---|---|---|---|
| | | | le163K | | | | | | | | | | | | ile | | ile163K | 0%) | -methyltransferase (100%) | 00%) | | 00%) | nsferase activity (100%) | | 00%) | 0%); IPR023397 (100%); IPR029063 (100%) | | | | | | | | | | | |
| 504 | Variable 5 | LGHEMGALLK | Mycoplasmam | Bacteria | Tenericutes | | Mollicutes | | Mycoplasmatales | | Mycoplasmataceae | Mycoplasma | | | Mycoplasmam | | Mycoplasmam | | | GO:0005840 | ribosome (100%) | GO:0003735 | structural constituent of rib | GO:0006412 (100%) | translation (100%) | IPR004389( | 1.00E+00 | 1.00E+00 | 1.40E+06 | NS | NS | NS | NS | NS | NS | NS | NS |

| | | | | | | | | | | | | | | | | | | | | | | | | | | | | | | | | | | | | | |
|---|---|---|---|---|---|---|---|---|---|---|---|---|---|---|---|---|---|---|---|---|---|---|---|---|---|---|---|---|---|---|---|---|---|---|---|---|---|
| | 52 | | obile 163K | | | | | | | | | | | | obile | | obile 163K | | | (100%) | | (100%);GO:0019843(100%) | osome(100%);rRNA binding(100%) | | | 100%);IPR005484(100%) | | | | | | | | | | | |
| 505 | Variable 553 | NNVASEPGSFNAK | Mycoplasma mobile 163K | Bacteria | Tenericutes | | Mollicutes | | Mycoplasmatales | | Mycoplasmataceae | Mycoplasma | | | Mycoplasma mobile | | Mycoplasma mobile 163K | | | GO:0005737(100%);GO:00 | cytoplasm(100%);excinuclease repair compl | GO:0003677(100%);GO:00 | DNA binding(100%);ATP binding(100%);excinuclease AB | GO:0006289(100%);GO:0009432(100%) | nucleotide-excision repair(100%);SOS resp | IPR001650(100%);IPR001 | 1.00E+00 | 1.00E+00 | 7.92E+04 | NS | NS | NS | NS | NS | NS | NS | NS |

| | | | | | | | | | | | | | | | | | | | | | | | | | | | | | | | | | | | | | |
|---|---|---|---|---|---|---|---|---|---|---|---|---|---|---|---|---|---|---|---|---|---|---|---|---|---|---|---|---|---|---|---|---|---|---|---|---|---|
| | | | | | | | | | | | | | | | | | | | | 09380(100%) | ex(100%) | 05524(100%);GO:0009381(100%) | C activity(100%) | | onse(100%) | 943(100%);IPR004807(100%) | | | | | | | | | | | |
| 506 | Variable 554 | TPAEYLAEK | Mycoplasmapneumoni | Bacteria | Tenericutes | | Mollicutes | | Mycoplasmatales | | Mycoplasmataceae | Mycoplasma | | | Mycoplasmapneumon | | | | | GO:0005886(100%) | plasmamembrane(100%) | | | | | IPR004890(100%);I | 1.00E+00 | 1.00E+00 | 1.02E+06 | 1.5497 | 0.0002666 | 0.0004271 | 0.9 | NS | NS | NS | NS |

| | | | | | | | | | | | | | | | | | | | | | | | | | | | | | | | | | | | | | | | | |
|---|---|---|---|---|---|---|---|---|---|---|---|---|---|---|---|---|---|---|---|---|---|---|---|---|---|---|---|---|---|---|---|---|---|---|---|---|---|---|---|---|
| | | | ae | | | | | | | | | | | | iae | | | | | | | | | | | PR004984(100%) | | | | | | | | | | | | | | |
| 507 | Variable 555 | YWVNSPAVKPTAQPSR | Mycoplasmapneumoniae | Bacteria | Tenericutes | | Mollicutes | | Mycoplasmatales | | Mycoplasmataceae | Mycoplasma | | | Mycoplasmapneumoniae | | | | | GO:0005886(100%);GO:0016021(10 | plasmamembrane(100%);integralcomponentofmembrane | | | | | IPR004477(100%) | 1.00E+00 | 1.00E+00 | 1.26E+05 | 0.28369 | 2.4227E-10 | 0.0013569 | 0.9 | NS | NS | NS | NS | 17.95 | 2.3229E-06 | 0.00081275 |

| | | | | | | | | | | | | | | | | | | | | | | | | | | | | | | | | | | | | | | | | |
|---|---|---|---|---|---|---|---|---|---|---|---|---|---|---|---|---|---|---|---|---|---|---|---|---|---|---|---|---|---|---|---|---|---|---|---|---|---|---|---|---|
| | | | | | | | | | | | | | | | | | | | | 0%) | (100%) | | | | | | | | | | | | | | | | | | | |
| 508 | Variable 556 | QVVFDSGDR | Mycoplasmapneumoniae | Bacteria | Tenericutes | | Mollicutes | | Mycoplasmatales | | Mycoplasmataceae | Mycoplasma | | | Mycoplasmapneumoniae | | | 2.4.2.-(100%) | Pentosyltransferases (100%) | GO:0005576(100%);GO:0005886(33%);GO:0009 | extracellular region (100%);plasma membrane (33%);cell surface (33%) | GO:0003950(100%);GO:0090729(33%) | NAD+ ADP-ribosyltransferase activity (100%);toxin activity (33%) | GO:0009405(100%);GO:0044075(33%) | obsolete pathogenesis(100%);modulation by symbiont of host vacuole organization (3 | IPR003898(100%) | 1.00E+00 | 1.00E+00 | 1.50E+05 | 8.5973 | 0.000082861 | 0.00013333 | 0.9 | NS | NS | NS | NS | 5.3703 | 0.083793 | 0.00064642 |

| | | | | | | | | | | | | | | | | | | | | | | | | | | | | | | | | | | | | | | | | |
|---|---|---|---|---|---|---|---|---|---|---|---|---|---|---|---|---|---|---|---|---|---|---|---|---|---|---|---|---|---|---|---|---|---|---|---|---|---|---|---|---|
| | | | | | | | | | | | | | | | | | | | | 86(33%) | | | | | 3%) | | | | | | | | | | | | | | | |
| 509 | Variable 557 | FNASQDCGVLK | Mycoplasmapneumoniae | Bacteria | Tenericutes | | Mollicutes | | Mycoplasmatales | | Mycoplasmataceae | Mycoplasma | | | Mycoplasmapneumoniae | | | | | | | GO:0003677(100%) | DNA binding (100%) | GO:0006304(60%);GO:0009307(40%) | DNA modification (60%);DNA restriction-modification system (40%) | IPR000055(100%) | 1.00E+00 | 1.00E+00 | 1.33E+05 | NS | NS | NS | NS | NS | NS | NS | NS | | | |
| 5 | V | VL | M | B | T | | M | | M | | M | M | | | M | | | | | G | pl | | | | | I | 1 | 1 | 4 | 4 | 0 | 0. | 0 | N | N | N | N | 6 | 0 | 0 |

| | | | | | | | | | | | | | | | | | | | | | | | | | | | | | | | | | | | | | | | | |
|---|---|---|---|---|---|---|---|---|---|---|---|---|---|---|---|---|---|---|---|---|---|---|---|---|---|---|---|---|---|---|---|---|---|---|---|---|---|---|---|---|
| 11 | ariable 559 | GVQSQTVK | ycoplasmapneumoniae | acteria | enericutes | | ollicutes | | ycoplasmatales | | ycoplasmataceae | ycoplasma | | | ycoplasmapneumoniae | | | | | O:0005886(100%) | asmamembrane(100%) | | | | | PR001595(100%) | .00E+00 | .00E+00 | .14E+04 | .1059 | .00027374 | 0001818 | .9 | S | S | S | S | .8312 | .0027004 | .00602 |
| 513 | Variable 560 | SGLNLGGQAAR | Mycoplasmapneumoniae | Bacteria | Tenericutes | | Mollicutes | | Mycoplasmatales | | Mycoplasmataceae | Mycoplasma | | | Mycoplasmapneumoniae | | | 2.8.1.13(100%) | tRNA-uridine 2-sulfurtransferase(100%) | GO:0005737(100%) | cytoplasm(100%) | GO:0000049(100%);GO:0005524 | tRNA binding(100%);ATP binding(100%);sulfurtransferase activity(10 | GO:0006400(100%);GO:0032259(33%) | tRNA modification(100%); methylation(33%) | IPR004506(100%);IPR014729(1 | 1.00E+00 | 1.00E+00 | 4.64E+05 | 5.9753 | 0.043249 | 0.00063944 | 0.9 | NS | NS | NS | NS | | | |

| | | | | | | | | | | | | | | | | | | | | | | | | | | | | | | | | | | | | | |
|---|---|---|---|---|---|---|---|---|---|---|---|---|---|---|---|---|---|---|---|---|---|---|---|---|---|---|---|---|---|---|---|---|---|---|---|---|---|
| | | | | | | | | | | | | | | | | | | | | | | (100%);GO:0016783(100%) | 0%) | | | 00%);IPR023382(100%) | | | | | | | | | | | |
| 514 | Variable 561 | TLDQANLQLWTGAGWR | Mycoplasmapneumoniae | Bacteria | Tenericutes | | Mollicutes | | Mycoplasmatales | | Mycoplasmataceae | Mycoplasma | | | Mycoplasmapneumoniae | | | | | GO:0016021(100%);GO:00 | integral component of membrane (100%);plasm | | | GO:0020035 (4%) | cytoadherence to microvasculature | mediated by symbiont protein ( | 1.00E+00 | 1.00E+00 | 1.89E+05 | NS | NS | NS | NS | NS | NS | NS | NS |

| | | | | | | | | | | | | | | | | | | | | | | | | | | | | | | | | | | | | | | | | |
|---|---|---|---|---|---|---|---|---|---|---|---|---|---|---|---|---|---|---|---|---|---|---|---|---|---|---|---|---|---|---|---|---|---|---|---|---|---|---|---|---|
| | | | | | | | | | | | | | | | | | | | | 05886(4%);GO:0016020(4%) | amembrane(4%);membrane(4%) | | | | | 4%) | | | | | | | | | | | | | | |
| 515 | Variable 562 | GVQSLEWAK | Mycoplasmapneumoniae | Bacteria | Tenericutes | | Mollicutes | | Mycoplasmatales | | Mycoplasmataceae | Mycoplasma | | | Mycoplasmapneumoniae | | | 3.6.1.1(100%) | Inorganic diphosphatase (100%) | GO:0005737(100%) | cytoplasm(100%) | GO:0000287(100%);GO:0 | magnesium ion binding(100%);inorganic diphosphatase act | GO:0006796(100%) | phosphate-containing compound metabolic | IPR008162(100%);IPR03 | 1.00E+00 | 1.00E+00 | 1.39E+05 | 0.42625 | 3.344E-10 | 0.001738 | 0.9 | NS | NS | NS | NS | 19.396 | 1.0804E-06 | 0.0027649 |

| | | | | | | | | | | | | | | | | | | | | | | | | | | | | | | | | | | | | | |
|---|---|---|---|---|---|---|---|---|---|---|---|---|---|---|---|---|---|---|---|---|---|---|---|---|---|---|---|---|---|---|---|---|---|---|---|---|---|
| | | | | | | | | | | | | | | | | | | | | | | 0044427(100%) | ivity (100%) | | process (100%) | 6649(100%) | | | | | | | | | | | |
| 516 | Variable 563 | VVGAFNQAK | Mycoplasma pneumoniae | Bacteria | Tenericutes | | Mollicutes | | Mycoplasmatales | | Mycoplasmataceae | Mycoplasma | | | Mycoplasma pneumoniae | | | | | GO:0005737(100%) | cytoplasm(100%) | | | GO:0030490(100%) | maturation of SSU-rRNA(100%) | IPR000238(100%);IPR015946(100%);I | 1.00E+00 | 1.00E+00 | 1.18E+05 | 7.4785 | 0.0052775 | 0 | 0.9 | NS | NS | NS | NS |

| | | | | | | | | | | | | | | | | | | | | | | | | | | | | | | | | | | | | | |
|---|---|---|---|---|---|---|---|---|---|---|---|---|---|---|---|---|---|---|---|---|---|---|---|---|---|---|---|---|---|---|---|---|---|---|---|---|---|
| | | | | | | | | | | | | | | | | | | | | | | | | | | PR020053(100%) | | | | | | | | | | | |
| 518 | Variable 565 | MCPDLNLPR | Mycoplasmapneumoniae | Bacteria | Tenericutes | | Mollicutes | | Mycoplasmatales | | Mycoplasmataceae | Mycoplasma | | | Mycoplasmapneumoniae | | | | | | | GO:0005524(100%);GO:0046872(10 | ATP binding(100%);metal ion binding(100%) | | | IPR011761(100%);IPR013651(100% | 1.00E+00 | 9.35E+05 | 1.00E+00 | NS | NS | NS | NS | NS | NS | NS | NS |

| | | | | | | | | | | | | | | | | | | | | | | | | | | | | | | | | | | | | | |
|---|---|---|---|---|---|---|---|---|---|---|---|---|---|---|---|---|---|---|---|---|---|---|---|---|---|---|---|---|---|---|---|---|---|---|---|---|---|
| | | | | | | | | | | | | | | | | | | | | | | 0%) | | | | );IPR013815(100%) | | | | | | | | | | | |
| 519 | Variable 566 | AMTANYPPSWR | Mycoplasma pneumoniae | Bacteria | Tenericutes | | Mollicutes | | Mycoplasmatales | | Mycoplasmataceae | Mycoplasma | | | Mycoplasma pneumoniae | | | | | GO:0016021(100%);GO:0005886( | integral component of membrane (100%);plasma membran | | | GO:0020035 (6%) | cytoadherence to microvasculature | mediated by symbiont protein (6%) | 1.00E+00 | 4.28E+05 | 1.00E+00 | NS | NS | NS | NS | NS | NS | NS | NS |

| | | | | | | | | | | | | | | | | | | | | | | | | | | | | | | | | | | | | | |
|---|---|---|---|---|---|---|---|---|---|---|---|---|---|---|---|---|---|---|---|---|---|---|---|---|---|---|---|---|---|---|---|---|---|---|---|---|---|
| | | | | | | | | | | | | | | | | | | | | 6%);GO:0016020(6%) | e(6%);membrane(6%) | | | | | | | | | | | | | | | | |
| 520 | Variable 567 | LNLQDQQDR | Mycoplasmapneumoniae | Bacteria | Tenericutes | | Mollicutes | | Mycoplasmatales | | Mycoplasmataceae | Mycoplasma | | | Mycoplasmapneumoniae | | | | | GO:0016021(100%);GO:0005886 | integral component of membrane(100%);plasma membr | | | | | | 1.00E+00 | 1.83E+05 | 1.00E+00 | NS | NS | NS | NS | NS | NS | NS | NS |

| | | | | | | | | | | | | | | | | | | | | | | | | | | | | | | | | | | | | | |
|---|---|---|---|---|---|---|---|---|---|---|---|---|---|---|---|---|---|---|---|---|---|---|---|---|---|---|---|---|---|---|---|---|---|---|---|---|---|
| | | | | | | | | | | | | | | | | | | | | (50%) | ane(50%) | | | | | | | | | | | | | | | | |
| 521 | Variable 568 | LTASGHALFAK | Mycoplasmapneumoniae M129 | Bacteria | Tenericutes | | Mollicutes | | Mycoplasmatales | | Mycoplasmataceae | Mycoplasma | | | Mycoplasmapneumoniae | | Mycoplasmapneumoniae M129 | | | GO:0016021(100%) | integral component of membrane(100%) | | | | | IPR007422(100%);IPR036764(100%) | 1.00E+00 | 1.00E+00 | 4.93E+04 | NS | NS | NS | NS | NS | NS | NS | NS |
| 522 | Variabl | QLNQNAHQLFLELLK | Mycoplas | Bacteri | Tenericu | | Mollicutes | | Mycoplasmat | | Mycoplasmat | Mycoplas | Mycopla | | Mycoplas | Mycopla | | 6.1.1.1 | Glycine-tRNA lig | GO:0005 | cytoplasm(100 | GO:0004 | glycine-tRNA lig | GO:0006426(1 | glycyl-tRNA | IPR0023 | 1.00E+0 | 7.94E+0 | 1.00E+0 | NS | NS | NS | NS | NS | NS | NS | NS |

| | | | | | | | | | | | | | | | | | | | | | | | | | | | | | | | | | | | | | | | | |
|---|---|---|---|---|---|---|---|---|---|---|---|---|---|---|---|---|---|---|---|---|---|---|---|---|---|---|---|---|---|---|---|---|---|---|---|---|---|---|---|---|
| | e 569 | | mamycoidessubsp.mycoides | a | tes | | | | ales | | aceae | ma | smamycoidesgroup | | mamycoides | smamycoidessubsp.mycoides | | 4(100%) | ase(100%) | 737(100%) | %) | 820(100%);GO:0005524(100%);GO:0046983(50%) | aseactivity(100%);ATPbinding(100%);proteindimerizationactivity(50%) | 00%) | aminoacylation(100%) | 14(100%);IPR002315(100%);IPR004154(100%) | 0 | 4 | 0 | | | | | | | | | | | |
| 523 | Var | MSEAVG | Cor | Bac | Acti | | Actin | | Cory | | Coryn | Cor | | | | | | 6.1 | Isoleucin | GO: | cytopl | GO: | tRNA | GO:00 | isoleu | IPR | 1.0 | 1.0 | 6.4 | 0.4 | 4.9 | 0.00 | 0.9 | NS | NS | NS | NS | 10. | 0.0 | 0.0 |

| | | | | | | | | | | | | | | | | | | | | | | | | | | | | | | | | | | | | | | | | |
|---|---|---|---|---|---|---|---|---|---|---|---|---|---|---|---|---|---|---|---|---|---|---|---|---|---|---|---|---|---|---|---|---|---|---|---|---|---|---|---|---|
| | iable 57 | GVYPQVDLSGGSSR | ynebacterium | teria | nobacteria | | omycetia | | nebacteriales | | ebacteriaceae | ynebacterium | | | | | | .1.5(100%) | e--tRNA ligase(100%) | 0005737(100%) | asm(100%) | 0000049(100%);GO:0002161(100%);GO:0004822(100 | binding(100%);aminoacyl-tRNA editing activity(100%);isoleucine-tRNA ligase activity(100%) | 06428(100%) | cyl-tRNA aminoacylation(100%) | 001412(100%);IPR002300(100%);IPR002301(100%) | 0E+00 | 0E+00 | 7E+04 | 7617 | 901E-08 | 13518 | | | | | | 699 | 0017561 | 0047792 |

| | | | | | | | | | | | | | | | | | | | | | | | | | | | | | | | | | | | | | |
|---|---|---|---|---|---|---|---|---|---|---|---|---|---|---|---|---|---|---|---|---|---|---|---|---|---|---|---|---|---|---|---|---|---|---|---|---|---|
| | | | | | | | | | | | | | | | | | | | | | | %) | | | | | | | | | | | | | | | |
| 524 | Variable 570 | MSSNFDNMPR | Ureaplasma | Bacteria | Tenericutes | | Mollicutes | | Mycoplasmatales | | Mycoplasmataceae | Ureaplasma | | | | | | | | | | | | | | | 1.00E+00 | 1.00E+00 | 3.01E+04 | NS | NS | NS | NS | NS | NS | NS | NS |
| 525 | Variable 571 | VVDDYNSSSVEQAK | Ureaplasma parvum | Bacteria | Tenericutes | | Mollicutes | | Mycoplasmatales | | Mycoplasmataceae | Ureaplasma | | | Ureaplasma parvum | | | | | GO:0005737 (33%) | cytoplasm (33%) | | | GO:0007059 (100%);GO:0007049 (33%);GO:0051301 (33%) | chromosome segregation (100%);cell cycle (33%);cell division (33%) | IPR003768 (100%) | 1.00E+00 | 1.00E+00 | 3.04E+04 | 2.1342 | 0.00058078 | 0.00021538 | 0.9 | NS | NS | NS | NS |

| | | | | | | | | | | | | | | | | | | | | | | | | | | | | | | | | | | | | | |
|---|---|---|---|---|---|---|---|---|---|---|---|---|---|---|---|---|---|---|---|---|---|---|---|---|---|---|---|---|---|---|---|---|---|---|---|---|---|
| | | | | | | | | | | | | | | | | | | | | | | | | | (33%) | | | | | | | | | | | | |
| 526 | Variable 572 | SSDSVFGYTDLGFLPK | Thermotoga | Bacteria | Thermotogae | | Thermotogae | | Thermotogales | | Thermotogaceae | Thermotoga | | | | | | 2.7.1.30(100%) | Glycerol kinase (100%) | GO:0005829(25%) | cytosol (25%) | GO:0004370(100%);GO:0005524(100%) | glycerol kinase activity (100%);ATP binding (100%) | GO:0006072(100%);GO:0019563(100%);GO:0006071(25%) | glycerol-3-phosphate metabolic process (100%);glycerol catabolic process (100% | IPR000577(100%);IPR005999(100%);IPR018483 | 1.00E+00 | 1.00E+00 | 3.46E+04 | NS | NS | NS | NS | NS | NS | NS | NS |

| | | | | | | | | | | | | | | | | | | | | | | | | | | | | | | | | | | | | | | | | |
|---|---|---|---|---|---|---|---|---|---|---|---|---|---|---|---|---|---|---|---|---|---|---|---|---|---|---|---|---|---|---|---|---|---|---|---|---|---|---|---|---|
| | | | | | | | | | | | | | | | | | | | | | | | | | ); glycerol metabolic process (25%) | (100%) | | | | | | | | | | | | | | |
| 529 | Variable 575 | CTLDHFEEDR | Thermotoga maritima MSB8 | Bacteria | Thermotogae | | Thermotogae | | Thermotogales | | Thermotogaceae | Thermotoga | | | Thermotoga maritima | | Thermotoga maritima MSB8 | 5.1.2.7 (100%) | Tagaturonate epimerase (100%) | | | GO:0016856 (100%); GO:004687 | racemase and epimerase activity | acting on hydroxy acids and derivatives (100%);metal ion bindin | | | 1.00E+00 | 1.00E+00 | 3.46E+04 | 0.389 | 1.234E-10 | 0.0021348 | 0.9 | NS | NS | NS | NS | 21.829 | 3.1637E-07 | 0.003213 |

| | | | | | | | | | | | | | | | | | | | | | | | | | | | | | | | | | | | | | | | | |
|---|---|---|---|---|---|---|---|---|---|---|---|---|---|---|---|---|---|---|---|---|---|---|---|---|---|---|---|---|---|---|---|---|---|---|---|---|---|---|---|---|
| | | | | | | | | | | | | | | | | | | | | | | 2(100%) | | g(100%) | | | | | | | | | | | | | | | | |
| 530 | Variable 576 | FGYVGEQPVYQEAETGGSFFEDVFR | Thermotoganeapolitana DSM 4359 | Bacteria | Thermotogae | | Thermotogae | | Thermotogales | | Thermotogaceae | Thermotoga | | | Thermotoganeapolitana | | Thermotoganeapolitana DSM 4359 | | | GO:0005737(100%) | cytoplasm(100%) | GO:0005524(100%);GO:0008270(100%);GO:0 | ATP binding(100%);zinc ion binding(100%);heat shock protein binding(100%) | GO:0006260(100%);GO:0006457(100%);GO:0009408(100%) | DNA replication(100%);protein folding(100%);response to heat(100%) | IPR001305(100%);IPR001623(100%);IPR002 | 1.00E+00 | 1.00E+00 | 2.70E+04 | 5.3271 | 1.19E-07 | 0.00020327 | 0.9 | NS | NS | NS | NS | 15.64 | 8.3861E-06 | 0.0019612 |

| | | | | | | | | | | | | | | | | | | | | | | | | | | | | | | | | | | | | | |
|---|---|---|---|---|---|---|---|---|---|---|---|---|---|---|---|---|---|---|---|---|---|---|---|---|---|---|---|---|---|---|---|---|---|---|---|---|---|
| | | | | | | | | | | | | | | | | | | | | | | 0310072(100%) | | | | 939(100%) | | | | | | | | | | | |
| 531 | Variable 577 | LLVDESTGVTFADVAGCDEAK | Methylacidiphilum | Bacteria | Verrucomicrobia | | Methylacidiphilae | | Methylacidiphilales | | Methylacidiphilaceae | Methylacidiphilum | | | | | | 3.4.24.-(100%) | Metalloendopeptidases(100%) | GO:0005886(100%);GO:0016021(100% | plasma membrane(100%);integral component of membrane(100 | GO:0004176(100%);GO:0004222(100% | ATP-dependent peptidase activity(100%);metalloendopeptidase activity(100%);ATP | GO:0030163(100%);GO:0051301(67%) | protein catabolic process(100%);cell division(67%) | IPR000642(100%);IPR003593(100%);I | 1.00E+00 | 1.00E+00 | 1.06E+05 | NS | NS | NS | NS | NS | NS | NS | NS |

| | | | | | | | | | | | | | | | | | | | | | | | | | | | | | | | | | | | | | |
|---|---|---|---|---|---|---|---|---|---|---|---|---|---|---|---|---|---|---|---|---|---|---|---|---|---|---|---|---|---|---|---|---|---|---|---|---|---|
| | | | | | | | | | | | | | | | | | | | | ) | %) | );GO:0005524(100%) | binding(100%) | | | PR003959(100%) | | | | | | | | | | | |
| 532 | Variable 578 | EDAADLTQESFLK | CandidatusMoanabactertarae | Bacteria | Verrucomicrobia | | Opitutae | | Puniceicoccales | | Puniceicoccaceae | CandidatusMoanabacter | | | CandidatusMoanabactertarae | | | | | | | GO:0003677(100%);GO:0003700 | DNA binding(100%);DNA-binding transcription factor activity(100%);si | GO:0006352(100%) | DNA-templated transcription | initiation(100%) | 6.78E+05 | 1.00E+00 | 1.00E+00 | NS | NS | NS | NS | NS | NS | NS | NS |

| | | | | | | | | | | | | | | | | | | | | | | | | | | | | | | | | | | | | | |
|---|---|---|---|---|---|---|---|---|---|---|---|---|---|---|---|---|---|---|---|---|---|---|---|---|---|---|---|---|---|---|---|---|---|---|---|---|---|
| | | | | | | | | | | | | | | | | | | | | | | (100%);GO:0016987(100%) | gma factor activity (100%) | | | | | | | | | | | | | | |
| 533 | Variable 579 | LNLNPLVEGR | Akkermansiamuciniphila | Bacteria | Verrucomicrobia | | Verrucomicrobiae | | Verrucomicrobiales | | Akkermansiaceae | Akkermansia | | | Akkermansiamuciniphila | | | | | GO:0005737(100%) | cytoplasm(100%) | | | GO:0006415(100%) | translational termination (100%) | IPR002661(100%);IPR023 | 1.00E+00 | 1.00E+00 | 7.80E+04 | NS | NS | NS | NS | NS | NS | NS | NS |

| | | | | | | | | | | | | | | | | | | | | | | | | | | | | | | | | | | | | | |
|---|---|---|---|---|---|---|---|---|---|---|---|---|---|---|---|---|---|---|---|---|---|---|---|---|---|---|---|---|---|---|---|---|---|---|---|---|---|
| | | | | | | | | | | | | | | | | | | | | | | | | | | 584(100%);IPR036191(100%) | | | | | | | | | | | |
| 534 | Variable 58 | QSAELSSFMG | Corynebacterium | Bacteria | Actinobacteria | | Actinomycetia | | Corynebacteriales | | Corynebacteriaceae | Corynebacterium | | | | | | | | | | GO:0000287(100%);GO: | magnesium ion binding(100%);dGTPase activity(100% | GO:0006203(100%) | dGTP catabolic process(100%) | IPR003607(100%);IPR0 | 1.00E+00 | 1.00E+00 | 2.97E+04 | NS | NS | NS | NS | NS | NS | NS | NS |

| | | | | | | | | | | | | | | | | | | | | | | | | | | | | | | | | | | | | | |
|---|---|---|---|---|---|---|---|---|---|---|---|---|---|---|---|---|---|---|---|---|---|---|---|---|---|---|---|---|---|---|---|---|---|---|---|---|---|
| | | | | | | | | | | | | | | | | | | | | | | 0008832(100%) | ) | | | 06261(100%);IPR006674(100%) | | | | | | | | | | | |
| 535 | Variable 59 | FAEMLPLMADGR | Corynebacterium | Bacteria | Actinobacteria | | Actinomycetia | | Corynebacteriales | | Corynebacteriaceae | Corynebacterium | | | | | | 1.1.5.4(100%) | Malate dehydrogenase (quinone) (100%) | | | GO:0008924(100%);G | malate dehydrogenase (quinone) activity (100%) | GO:0006099(100%) | tricarboxylic acid cycle (100%) | IPR006231(100%);IP | 1.00E+00 | 9.24E+04 | 1.00E+00 | NS | NS | NS | NS | NS | NS | NS | NS |

| | | | | | | | | | | | | | | | | | | | | | | | | | | | | | | | | | | | | | |
|---|---|---|---|---|---|---|---|---|---|---|---|---|---|---|---|---|---|---|---|---|---|---|---|---|---|---|---|---|---|---|---|---|---|---|---|---|---|
| | | | | | | | | | | | | | | | | | | | | | | O:0052589(100%) | );malate dehydrogenase (menaquinone) activity (100%) | | | R0361888(100%) | | | | | | | | | | | |
| 537 | Variable 60 | FALSLLAAAK | Corynebacterium | Bacteria | Actinobacteria | | Actinomycetia | | Corynebacteriales | | Corynebacteriaceae | Corynebacterium | | | | | | 6.1.1.19(100%) | Arginine--tRNA ligase (100%) | GO:0005737(100%) | cytoplasm (100%) | GO:0004814(100%);GO:0005 | arginine-tRNA ligase activity (100%);ATP binding (100%) | GO:0006420 (100%) | arginyl-tRNA aminoacylation (100%) | IPR001278(100%);IPR00141 | 1.00E+00 | 5.61E+05 | 1.00E+00 | NS | NS | NS | NS | NS | NS | NS | NS |

| | | | | | | | | | | | | | | | | | | | | | | | | | | | | | | | | | | | | | |
|---|---|---|---|---|---|---|---|---|---|---|---|---|---|---|---|---|---|---|---|---|---|---|---|---|---|---|---|---|---|---|---|---|---|---|---|---|---|
| | | | | | | | | | | | | | | | | | | | | | | 524(100%) | | | | 2(100%);IPR005148(100%) | | | | | | | | | | | |
| 538 | Variable 61 | VSGAPEAPYAVVLGGSK | Corynebacterium urealyticum | Bacteria | Actinobacteria | | Actinomycetia | | Corynebacteriales | | Corynebacteriaceae | Corynebacterium | | | Corynebacterium urealyticu | | | 2.7.2.3(100%) | Phosphoglycerate kinase (100%) | GO:0005737(100%) | cytoplasm(100%) | GO:0004618(100%);GO:00 | phosphoglycerate kinase activity(100%);ATP binding(100%) | GO:0006096(100%) | glycolytic process(100%) | IPR001576(100%);IPR015 | 1.00E+00 | 1.00E+00 | 7.89E+04 | 3.0855 | 0.02411 | 0 | 0.9 | NS | NS | NS | NS |

| | | | | | | | | | | | | | | | | | | | | | | | | | | | | | | | | | | | | | | | | |
|---|---|---|---|---|---|---|---|---|---|---|---|---|---|---|---|---|---|---|---|---|---|---|---|---|---|---|---|---|---|---|---|---|---|---|---|---|---|---|---|---|
| | | | | | | | | | | | | | | | m | | | | | | | 05524(100%) | | | | 824(100%);IPR015911(100%) | | | | | | | | | | | | | | |
| 539 | Variable 62 | AADLFAAAQGDSQQVMAK | Mycobacteriaceae | Bacteria | Actinobacteria | | Actinomycetia | | Corynebacteriales | | Mycobacteriaceae | | | | | | | | | GO:0005886(100%);GO: | plasma membrane (100%);integral com | coupling factor F(o) (10 | GO:0046933 (100%) | proton-transporting ATP synthase activity | rotational mechanism (100%) | | 1.00E+00 | 1.00E+00 | 3.93E+04 | 5.643 | 1.655E-08 | 0.00052623 | 0.9 | NS | NS | NS | NS | 20.994 | 4.7818E-07 | 0.00534334 |

| | | | | | | | | | | | | | | | | | | | | | | | | | | | | | | | | | | | | | | | | |
|---|---|---|---|---|---|---|---|---|---|---|---|---|---|---|---|---|---|---|---|---|---|---|---|---|---|---|---|---|---|---|---|---|---|---|---|---|---|---|---|---|
| | | | | | | | | | | | | | | | | | | | | 0016021(100%);GO:0045263(100%) | ponent of membrane (100%);proton-transporting ATP synthase complex | 0%) | | | | | | | | | | | | | | | | | | |
| 540 | Variable 6 | FAEAAAQTVR | Mycobacteriac | Bacteria | Actinobacte | | Actinomycetia | | Corynebacteriales | | Mycobacteriaceae | | | | | | | 3.6.4.-(100 | Acting on ATP; involved in cell | GO:1990077 | primosome complex | GO:0003677 | DNA binding (100%);ATP | GO:0006268(90%);G | DNA unwinding in vo | synthesis of | 1.00E+00 | 1.00E+00 | 1.93E+05 | 0.29679 | 1.5671E-09 | 0.00176 | 0.9 | NS | NS | NS | NS | 12.092 | 0.0007119 | 0.0012747 |

| | | | | | | | | | | | | | | | | | | | | | | | | | | | | | | | | | | | | | | |
|---|---|---|---|---|---|---|---|---|---|---|---|---|---|---|---|---|---|---|---|---|---|---|---|---|---|---|---|---|---|---|---|---|---|---|---|---|---|---|
| | 3 | | eae | | ria | | | | | | | | | | | | | %) | ular and subcellular movement (100%) | (90%); GO:0005886(1%) | (90%);plasmamembrane(1%) | (100%);GO:0005524(90%);GO:0008270(90%) | binding(90%);zinc ion binding(90%) | O:0006269(87%);GO:0006270(1%) | lved in DNA replication(90%);DNA replication | RNA primer(87%);DNA replication initiation(1%) | | | | | | | | | | | | 5 |
| 541 | Variabl | DLFASDPGECCR | Mycobact | Bacteri | Actinoba | | Actinomyce | | Corynebacter | | Mycobacteriac | | | | | | | 1.8.4.8 | Phosphoadenylyl-sul | GO:0005 | cytoplasm(93 | GO:0004 | phosphoadenylyl-sul | GO:0019344(9 | cysteinebiosyn | phosph | 1.00E+0 | 1.00E+0 | 2.58E+0 | NS | NS | NS | NS | NS | NS | NS | NS | |

| | | | | | | | | | | | | | | | | | | | | | | | | | | | | | |
|---|---|---|---|---|---|---|---|---|---|---|---|---|---|---|---|---|---|---|---|---|---|---|---|---|---|---|---|---|---|
| | e 64 | | eriaceae | a | cteria | | tia | | iales | | eae | | | | | | | (100%) | fate reductase (thioredoxin) (100%) | 737 (93%) | %) | 604 (100%) | fate reductase (thioredoxin) activity (100%) | 3%);GO:0019379 (93%);GO:0070814 (93%) | thetic processes (93%);sulfate assimilation | oadenylyl sulfate reduction by phosphoadenylyl-sulfate reductase | 0 | 0 | 4 |

| | | | | | | | | | | | | | | | | | | | | | | | | | | |
|---|---|---|---|---|---|---|---|---|---|---|---|---|---|---|---|---|---|---|---|---|---|---|---|---|---|---|
| | | | | | | | | | | | | | | | | | | | | | | | | | | e (thioredoxin) (93%); hydrogen sulfide biosynthetic process ( |

| | | | | | | | | | | | | | | | | | | | | | | | | | | | | | | | | | | | | | | | | |
|---|---|---|---|---|---|---|---|---|---|---|---|---|---|---|---|---|---|---|---|---|---|---|---|---|---|---|---|---|---|---|---|---|---|---|---|---|---|---|---|---|
| | | | | | | | | | | | | | | | | | | | | | | | | | | 93%) | | | | | | | | | | | | | | |
| 542 | Variable 65 | DVDDDALNVLAQK | Mycobacteriaceae | Bacteria | Actinobacteria | | Actinomycetia | | Corynebacteriales | | Mycobacteriaceae | | | | | | | | | | | GO:0003677(100%);GO:0005524(100%);GO:0008 | DNA binding(100%);ATP binding(100%);zinc ion binding(100%) | GO:0045892(100%) | negative regulation of transcription | DNA-templated(100%) | 1.00E+00 | 1.00E+00 | 7.24E+04 | 16.095 | 1.017E-08 | 0.0008985 | 0.9 | NS | NS | NS | NS | 22.151 | 2.7023E-07 | 0.0065843 |

| | | | | | | | | | | | | | | | | | | | | | | | | | | | | | | | | | | | | | |
|---|---|---|---|---|---|---|---|---|---|---|---|---|---|---|---|---|---|---|---|---|---|---|---|---|---|---|---|---|---|---|---|---|---|---|---|---|---|
| | | | | | | | | | | | | | | | | | | | | | | 270(100%) | | | | | | | | | | | | | | | |
| 543 | Variable 66 | GPGGGGGRPGGPGGGGGGNYR | Mycobacteriaceae | Bacteria | Actinobacteria | | Actinomycetia | | Corynebacteriales | | Mycobacteriaceae | | | | | | | | | GO:0005737(100%) | cytoplasm(100%) | GO:0003743(100%);GO:0003924(100%);GO | translation initiation factor activity(100%);GTPase activity(100%);GTP binding(100%) | | | IPR000178(100%);IPR000795(100%);IPR0 | 1.00E+00 | 2.03E+06 | 1.00E+00 | NS | NS | NS | NS | NS | NS | NS | NS |

| | | | | | | | | | | | | | | | | | | | | | | | | | | | | | | | | | | | | | |
|---|---|---|---|---|---|---|---|---|---|---|---|---|---|---|---|---|---|---|---|---|---|---|---|---|---|---|---|---|---|---|---|---|---|---|---|---|---|
| | | | | | | | | | | | | | | | | | | | | | | :0005525(100%) | | | | 05225(100%) | | | | | | | | | | | |
| 545 | Variable 68 | LAEVGVAALHR | Mycobacteriaceae | Bacteria | Actinobacteria | | Actinomycetia | | Corynebacteriales | | Mycobacteriaceae | | | | | | | 2.5.1.75(100%) | tRNA dimethylallyltransferase(100%) | | | GO:0005524(100%);GO:0052381(10 | ATP binding(100%);tRNA dimethylallyltransferase activity(100%) | GO:0008033(100%) | tRNA processing(100%) | IPR018022(100%);IPR027417(100% | 3.50E+06 | 1.00E+00 | 1.00E+00 | NS | NS | NS | NS | NS | NS | NS | NS |

| | | | | | | | | | | | | | | | | | | | | | | | | | | | | | | | | | | | | | |
|---|---|---|---|---|---|---|---|---|---|---|---|---|---|---|---|---|---|---|---|---|---|---|---|---|---|---|---|---|---|---|---|---|---|---|---|---|---|
| | | | | | | | | | | | | | | | | | | | | | | 0%) | | | | );IPR039657(100%) | | | | | | | | | | | |
| 550 | Variable 72 | SVVFASSGR | Mycobacterium | Bacteria | Actinobacteria | | Actinomycetia | | Corynebacteriales | | Mycobacteriaceae | Mycobacterium | | | | | | 6.1.1.6(100%);2.3.2.3(36%) | Lysine--tRNA ligase (100%);Lysyltransferase (36%) | GO:0005886(100%);GO:0016021( | plasma membrane (100%);integral component of membr | GO:0004824(96%);GO:0005524(9 | lysine-tRNA ligase activity (96%);ATP binding (92%);lysyltransferase act | GO:0006430(92%);GO:0046677(88%);GO:0006629(85%) | lysyl-tRNA aminoacylation (92%);response to antibiotic (8 | IPR024320(100%);IPR031553(10 | 1.00E+00 | 1.00E+00 | 3.26E+04 | NS | NS | NS | NS | NS | NS | NS | NS |

| | | | | | | | | | | | | | | | | | | | | | | | | | | | | | | | | | | | | | |
|---|---|---|---|---|---|---|---|---|---|---|---|---|---|---|---|---|---|---|---|---|---|---|---|---|---|---|---|---|---|---|---|---|---|---|---|---|---|
| | | | | | | | | | | | | | | | | | | | | 100%);GO:0005737(88%) | ane(100%);cytoplasm(88%) | 2%);GO:0050071(92%) | ivity (92%) | | 8%);lipid metabolic process (85%) | 0%);IPR004365(96%) | | | | | | | | | | | |
| 552 | Variable 74 | AFQELGFDSLTAVEMR | Mycobacterium | Bacteria | Actinobacteria | | Actinomycetia | | Corynebacteriales | | Mycobacteriaceae | Mycobacterium | | | | | | 2.3.1.41(76%);2.3.1.94(9 | Beta-ketoacyl-[acyl-carrier-protein] synthase I (76%); 6-deoxyerythro | GO:0016021(1%) | integral component of membrane (1%) | GO:0031177(100%);GO:0016 | phosphopantetheine binding (100%);oxidoreductase activity (89%) | GO:0006633(77%);GO:0006631(9%);GO:0008152 | fatty acid biosynthetic process (77%);fatty acid m | IPR009081(100%);IPR02080 | 1.00E+00 | 1.00E+00 | 2.27E+04 | NS | NS | NS | NS | NS | NS | NS | NS |

| | | | | | | | | | | | | | | | | | | | | | | | | | | | | | | | | | | | | | |
|---|---|---|---|---|---|---|---|---|---|---|---|---|---|---|---|---|---|---|---|---|---|---|---|---|---|---|---|---|---|---|---|---|---|---|---|---|---|
| | | | | | | | | | | | | | | | | | | %);1.-.-.-(6%) | nolide-B synthase (9%); Oxidoreductases (6%) | | | 491(89%);GO:0004315(82%) | ;3-oxoacyl-[acyl-carrier-protein] synthase activity (82%) | (0%) | etabolic processes (9%); metabolic processes (0%) | 6(100%);IPR036736(100%) | | | | | | | | | | | |
| 553 | Variable 75 | AAQDSFFTAR | Mycobacterium | Bacteria | Actinobacteria | | Actinomycetia | | Corynebacteriales | | Mycobacteriaceae | Mycobacterium | | | | | | | | GO:0005618(100%);GO:0 | cell wall (100%);cytosol (100%);plasmame | | | | | IPR007139(100%) | 1.00E+00 | 1.00E+00 | 9.17E+05 | 51.169 | 0.016469 | 0.000094538 | 0.9 | NS | NS | NS | NS |

| | | | | | | | | | | | | | | | | | | | | | | | | | | | | | | | | | | | | | |
|---|---|---|---|---|---|---|---|---|---|---|---|---|---|---|---|---|---|---|---|---|---|---|---|---|---|---|---|---|---|---|---|---|---|---|---|---|---|
| | | | | | | | | | | | | | | | | | | | | 005829(100%);GO:0005886(100%) | mbrane(100%) | | | | | | | | | | | | | | | | |
| 554 | Variable 76 | VWGQMDSAR | Mycobacterium | Bacteria | Actinobacteria | | Actinomycetia | | Corynebacteriales | | Mycobacteriaceae | Mycobacterium | | | | | | 6.3.2.2(100%) | Glutamate--cysteine ligase (100%) | GO:0005886(2%) | plasma membrane (2%) | GO:0004357(100% | glutamate-cysteine ligase activity (100%);A | GO:0006750(100%);GO:0052699 | glutathione biosynthetic process | IPR006336(100%); | 1.00E+00 | 1.00E+00 | 4.14E+04 | NS | NS | NS | NS | NS | NS | NS | NS |

| | | | | | | | | | | | | | | | | | | | | | | | | | | | | | | | | | | | | | |
|---|---|---|---|---|---|---|---|---|---|---|---|---|---|---|---|---|---|---|---|---|---|---|---|---|---|---|---|---|---|---|---|---|---|---|---|---|---|
| | | | | | | | | | | | | | | | | | | | | | | ");GO:0005524(90%) | TP binding (90%) | (98%) | s (100%);ergothioneine biosynthetic process (98%) | IPR011556(100%);IPR014746(100%) | | | | | | | | | | | |
| 555 | Variable 77 | MTNSTDGR | Mycobacterium | Bacteria | Actinobacteria | | Actinomycetia | | Corynebacteriales | | Mycobacteriaceae | Mycobacterium | | | | | | 1.1.1.267(100 | 1-deoxy-D-xylulose-5-phosphate red | | | GO:0070402(10 | NADPH binding (100%);1-deox | GO:0019288(85%);GO:0016 | isopentenyl diphosphate bi | methylerythritol 4- | 1.00E+00 | 1.00E+00 | 3.32E+04 | NS | NS | NS | NS | NS | NS | NS | NS |

| | | | | | | | | | | | | | | | | | | | | | | | | | | |
|---|---|---|---|---|---|---|---|---|---|---|---|---|---|---|---|---|---|---|---|---|---|---|---|---|---|---|
| | | | | | | | | | | | | | | | | | | %) | uctoisomerase (100%) | | | 0%);GO:0030604(92%);GO:0046872(85%) | y-D-xylulose-5-phosphate reductoisomerase activity (92%);metal ion binding (85%) | 114(77%);GO:0051483(8%) | osynthetic process | phosphate pathway(85%);terpenoid biosynthetic process(77 |

| | | | | | | | | | | | | | | | | | | | | | | | | | | | | | | | | | | | | | | | | |
|---|---|---|---|---|---|---|---|---|---|---|---|---|---|---|---|---|---|---|---|---|---|---|---|---|---|---|---|---|---|---|---|---|---|---|---|---|---|---|---|---|
| | | | | | | | | | | | | | | | | | | | | | | | | | | %);terpenoid biosynthetic process | | | | | | | | | | | | | | |
| 556 | Variable 78 | VYESTEQAR | Mycobacterium | Bacteria | Actinobacteria | | Actinomycetia | | Corynebacteriales | | Mycobacteriaceae | Mycobacterium | | | | | | 6.1.1.3(100%) | Threonine--tRNA ligase (100%) | GO:0005737(95%);GO:00 | cytoplasm(95%);cytosol(5%);plasmame | GO:0000049(100%);GO:0 | tRNA binding(100%);threonine-tRNA ligase activity | GO:0006435(100%) | threonyl-tRNA aminoacylation (100%) | IPR002320(100%);IPR01 | 1.00E+00 | 1.00E+00 | 4.45E+04 | 0.27058 | 2.882E-11 | 0.0019609 | 0.9 | NS | NS | NS | NS | 29.5 | 9.941E-09 | 0.0020057 |

| | | | | | | | | | | | | | | | | | | | | | | | | | | | | | | | | | | | | | |
|---|---|---|---|---|---|---|---|---|---|---|---|---|---|---|---|---|---|---|---|---|---|---|---|---|---|---|---|---|---|---|---|---|---|---|---|---|---|
| | | | | | | | | | | | | | | | | | | | | 05829(5%);GO:0005886(5%) | membrane(5%) | 004829(100%);GO:0005524(100%) | (100%);ATP binding(100%) | | | 2947(100%);IPR018163(100%) | | | | | | | | | | | |
| 557 | Variable 79 | VVPQTNSLTYR | Mycobacterium | Bacteria | Actinobacteria | | Actinomycetia | | Corynebacteriales | | Mycobacteriaceae | Mycobacterium | | | | | | 2.3.3.16(100%);2 | Citrate synthase (unknown stereospecificity) | GO:0005737(100% | cytoplasm(100%) | GO:0036440(100% | citrate synthase activity(100%);2-methylcit | GO:0006099(100%);GO:0019679 | tricarboxylic acid cycle(100%); | methylcitrate cycle(1 | 1.00E+00 | 5.30E+04 | 1.00E+00 | NS | NS | NS | NS | NS | NS | NS | NS |

| | | | | | | | | | | | | | | | | | | | | | | | | | | | | | | | | | | | | | |
|---|---|---|---|---|---|---|---|---|---|---|---|---|---|---|---|---|---|---|---|---|---|---|---|---|---|---|---|---|---|---|---|---|---|---|---|---|---|
| | | | | | | | | | | | | | | | | | | .3.3.5(67%) | (100%);2-methylcitrate synthase(67%) | ) | | );GO:0050440(18%);GO:0004108(6%) | rate synthase activity(18%);citrate(Si)-synthase activity(6%) | (18%);GO:0005975(6%) | propionate metabolic process | 8%);carbohydrate metabolic process(6%) | | | | | | | | | | | |
| 559 | Variable 80 | FPS NT ESR | Mycobacterium | Bacteria | Actinobacteria | | Actinomycetia | | Corynebacteriales | | Mycobacteriaceae | Mycobacterium | | | | | | | | GO:0016021(10 | integral component of membr | | | GO:0042783(3%) | evasion of host immune resp | IPR021403(100 | 1.00E+00 | 1.15E+05 | 1.00E+00 | NS | NS | NS | NS | NS | NS | NS | NS |

| | | | | | | | | | | | | | | | | | | | | | | | | | | | | | | | | | | | | | |
|---|---|---|---|---|---|---|---|---|---|---|---|---|---|---|---|---|---|---|---|---|---|---|---|---|---|---|---|---|---|---|---|---|---|---|---|---|---|
| | | | | | | | | | | | | | | | | | | | | 0%);GO:0005886(8%) | ane(100%);plasmamembrane(8%) | | | | onse(3%) | %) | | | | | | | | | | | |
| 561 | Variable 82 | EQVTSPQCQR | Mycobacterium | Bacteria | Actinobacteria | | Actinomyceta | | Corynebacteriales | | Mycobacteriaceae | Mycobacterium | | | | | | 2.4.2.-(78%);2.4.2.34(67%) | Pentosyltransferases(78%);Indolylacetylinositol arabinosyltransferase(67%) | GO:0016021(98%);GO:0005886 | integral component of membrane(98%);plasmamem | GO:0052636(84%);GO:0050409 | arabinosyltransferase activity(84%);indolylacetylinositol arabinosyltransfera | GO:0071766(94%);GO:0071555(71%);GO:0046677(3 | Actinobacterium-type cell wall biogenesis(94%);cell | IPR027451(100%);IPR040920( | 1.00E+00 | 4.08E+05 | 1.00E+00 | NS | NS | NS | NS | NS | NS | NS | NS |

| | | | | | | | | | | | | | | | | | | | | | | | | | | | | | | | | | | | | | |
|---|---|---|---|---|---|---|---|---|---|---|---|---|---|---|---|---|---|---|---|---|---|---|---|---|---|---|---|---|---|---|---|---|---|---|---|---|---|
| | | | | | | | | | | | | | | | | | | | | (71%);GO:0005618(2%) | brane(71%);cell wall(2%) | (10%);GO:0016740(3%) | se activity(10%);transferase activity(3%) | %) | wall organization(71%);response to antibiotic(3%) | 100%);IPR007680(95%) | | | | | | | | | | | |
| 562 | Variable 83 | NDQGMDPEDPEMLAEK | Mycobacterium | Bacteria | Actinobacteria | | Actinomycetia | | Corynebacteriales | | Mycobacteriaceae | Mycobacterium | | | | | | 3.1.1.-(100%) | Carboxylic ester hydrolases (100%) | GO:0005615(7%);GO:00 | extracellular space(7%);cell wall(7%) | GO:0052689(100%) | carboxylic ester hydrolase activity(100%) | GO:0016042(7%) | lipid catabolic process(7%) | IPR013830(100%);IPR0 | 1.00E+00 | 5.10E+04 | 1.00E+00 | NS | NS | NS | NS | NS | NS | NS | NS |

| | | | | | | | | | | | | | | | | | | | | | | | | | | | | | | | | | | | | | |
|---|---|---|---|---|---|---|---|---|---|---|---|---|---|---|---|---|---|---|---|---|---|---|---|---|---|---|---|---|---|---|---|---|---|---|---|---|---|
| | | | | | | | | | | | | | | | | | | | | 05618(7%) | | | | | | 36514(100%) | | | | | | | | | | | |
| 563 | Variable 84 | NFLPAPGPVTK | Mycobacterium | Bacteria | Actinobacteria | | Actinomycetia | | Corynebacteriales | | Mycobacteriaceae | Mycobacterium | | | | | | 6.3.4.14(100%) | Biotin carboxylase (100%) | GO:0005618(3%);GO:0005886(3%);GO:0 | cell wall(3%);plasma membrane(3%);acetyl-CoA carboxylase co | GO:0005524(100%);GO:0046872(69%);G | ATP binding(100%);metal ion binding(69%);ligase activity(52%) | GO:0015977(3%) | carbon fixation(3%) | IPR005479(100%);IPR011054(100%);IP | 1.00E+00 | 1.64E+05 | 1.00E+00 | NS | NS | NS | NS | NS | NS | NS | NS |

| | | | | | | | | | | | | | | | | | | | | | | | | | | | | | | | | | | | | | |
|---|---|---|---|---|---|---|---|---|---|---|---|---|---|---|---|---|---|---|---|---|---|---|---|---|---|---|---|---|---|---|---|---|---|---|---|---|---|
| | | | | | | | | | | | | | | | | | | | | 0093317(3%) | mplex(3%) | O:0016874(52%) | | | | R005482(97%) | | | | | | | | | | | |
| 564 | Variable 85 | LNLLADLLPPLSHKPR | Mycobacterium | Bacteria | Actinobacteria | | Actinomycetia | | Corynebacteriales | | Mycobacteriaceae | Mycobacterium | | | | | | | | GO:0005618(100%);GO:0005829(10 | cell wall(100%);cytosol(100%);plasma membrane(100%) | | | | | IPR036188(100%) | 1.00E+00 | 4.45E+05 | 1.00E+00 | NS | NS | NS | NS | NS | NS | NS | NS |

| | | | | | | | | | | | | | | | | | | | | | | | | | | | | | | | | | | | | | |
|---|---|---|---|---|---|---|---|---|---|---|---|---|---|---|---|---|---|---|---|---|---|---|---|---|---|---|---|---|---|---|---|---|---|---|---|---|---|
| | | | | | | | | | | | | | | | | | | | | 0%);GO:0005886(100%) | | | | | | | | | | | | | | | | | |
| 565 | Variable 86 | LVADGVEQFGR | Mycobacterium | Bacteria | Actinobacteria | | Actinomycetia | | Corynebacteriales | | Mycobacteriaceae | Mycobacterium | | | | | | 1.-.-.-(100%);1.1.99.-(25%) | Oxidoreductases(100%); With other acceptors(25%) | | | GO:0016491(100%) | oxidoreductase activity(100%) | | | IPR002347(100%);IPR020904 | 1.66E+06 | 1.00E+00 | 1.00E+00 | NS | NS | NS | NS | NS | NS | NS | NS |

| | | | | | | | | | | | | | | | | | | | | | | | | | | | | | | | | | | | | | |
|---|---|---|---|---|---|---|---|---|---|---|---|---|---|---|---|---|---|---|---|---|---|---|---|---|---|---|---|---|---|---|---|---|---|---|---|---|---|
| | | | | | | | | | | | | | | | | | | | | | | | | | | (100%);IPR023985(100%) | | | | | | | | | | | |
| 566 | Variable 87 | AVLGEASEVR | Mycobacterium | Bacteria | Actinobacteria | | Actinomycetia | | Corynebacteriales | | Mycobacteriaceae | Mycobacterium | | | | | | 2.3.1.-(89%);2.3.1.111(1 | Transferring groups other than amino-acyl groups (89%);Mycocero | GO:0005618(5%);GO:00058 | cell wall(5%);cytosol(5%);plasma membrane | GO:0016491(86%);GO:0031 | oxidoreductase activity(86%);phosphopantetheine binding(82%) | GO:0006633(59%);GO:0006631(18%);GO:0008 | fatty acid biosynthetic process(59%);fatty acid | IPR020807(100%);IPR0421 | 3.60E+05 | 1.00E+00 | 1.00E+00 | NS | NS | NS | NS | NS | NS | NS | NS |

| | | | | | | | | | | | | | | | | | | | | | | | | | | | | | | | | | | | | | |
|---|---|---|---|---|---|---|---|---|---|---|---|---|---|---|---|---|---|---|---|---|---|---|---|---|---|---|---|---|---|---|---|---|---|---|---|---|---|
| | | | | | | | | | | | | | | | | | | 1%) | sate synthase (11%) | 29(5%);GO:0005886(5%) | (5%) | 177(82%);GO:0004315(59%) | ;3-oxoacyl-[acyl-carrier-protein] synthase activity (59%) | 610 (5%) | metabolic process (18%);lipid biosynthetic process (5%) | 04(100%);IPR014043(88%) | | | | | | | | | | | |
| 567 | Variable 88 | FMNAVDWAHGAR | Mycobacterium leprae | Bacteria | Actinobacteria | | Actinomycetia | | Corynebacteriales | | Mycobacteriaceae | Mycobacterium | | | Mycobacterium lepra | | | 2.1.2.9(100%) | Methionyl-tRNA formyltransferase (100%) | | | GO:0004479(100% | methionyl-tRNA formyltransferase activ | | | IPR002376(100%); | 1.00E+00 | 1.00E+00 | 1.80E+05 | NS | NS | NS | NS | NS | NS | NS | NS |

| | | | | | | | | | | | | | | | | | | | | | | | | | | | | | | | | | | | | | | | | |
|---|---|---|---|---|---|---|---|---|---|---|---|---|---|---|---|---|---|---|---|---|---|---|---|---|---|---|---|---|---|---|---|---|---|---|---|---|---|---|---|---|
| | | | | | | | | | | | | | | | e | | | | | | | | ) | y (100%) | | | IPR005793(100%);IPR005794(100%) | | | | | | | | | | | | | |
| 571 | Variable 91 | MTSVTDHSAHSMER | Mycobacterium lepra | Bacteria | Actinobacteria | | Actinomycetia | | Corynebacteriales | | Mycobacteriaceae | Mycobacterium | | | Mycobacterium le | | | 2.1.3.-(50%);2. | Carboxy-and carbamoyltransferases (50 | | | GO:0004658(67 | propionyl-CoA carboxylase activity | | | IPR011762(100 | 1.00E+00 | 1.00E+00 | 4.27E+04 | 4.7154 | 0.0001635 | 0.000083333 | 0.9 | NS | NS | NS | NS | 7.3388 | 0.0018475 | 0.00016776 |

| | | | | | | | | | | | | | | | | | | | | | | | | | | | | | | | | | | | | | |
|---|---|---|---|---|---|---|---|---|---|---|---|---|---|---|---|---|---|---|---|---|---|---|---|---|---|---|---|---|---|---|---|---|---|---|---|---|---|
| | | | e | | | | | | | | | | | | prae | | | 1.3.1.5(50%);6.4.1.3(50%) | %);Acetyl-CoA carboxytransferase(50%);Propionyl-CoA carboxylase(50%) | | | %);GO:0005524(33%);GO:0016740(33%) | (67%);ATP binding(33%);transferase activity(33%) | | | %);IPR011763(100%);IPR029045(100%) | | | | | | | | | | | |
| 573 | Variable 93 | MSLETSTTPAGSPLSDR | Mycobacteriuma | Bacteria | Actinobacteri | | Actinomycetia | | Corynebacteriales | | Mycobacteriaceae | Mycobacterium | Mycobacteriu | | | | | 4.1.2.-(100% | Aldehyde-lyases(100%) | | | GO:0016832( | aldehyde-lyase activity(100% | GO:0005975(100%) | carbohydrate metabolic | IPR005593(1 | 1.00E+00 | 1.00E+00 | 2.59E+04 | 2.7046 | 0.00073571 | 0.00016935 | 0.9 | NS | NS | NS | NS |

| | | | | | | | | | | | | | | | | | | | | | | | | | | | | | | | | | | | | | |
|---|---|---|---|---|---|---|---|---|---|---|---|---|---|---|---|---|---|---|---|---|---|---|---|---|---|---|---|---|---|---|---|---|---|---|---|---|---|
| | | | viumcomplex(MAC) | | a | | | | | | | | m avium complex(MAC) | | | | | ) | | | | 100%) | ) | | process(100%) | 00%);IPR009014(100%);IPR018969(100%) | | | | | | | | | | | |
| 574 | Variable | VDGWPGQQFALLFVDWLTAD | Mycobacteriu | Bacteria | Actinobact | | Actinomycetia | | Corynebacteriale | | Mycobacteriaceae | Mycobacteri | Mycobacte | | | | | 2.7.1.24( | Dephospho-CoA kinase(10 | GO:000573 | cytoplasm(100%) | GO:000414 | dephospho-CoA kinase act | GO:0015937(100%) | coenzyme A biosyn | IPR001977 | 1.00E+00 | 1.00E+00 | 1.83E+05 | 1.6326 | 0.0002239 | 0.00054736 | 0.9 | NS | NS | NS | NS |

| | | | | | | | | | | | | | | | | | | | | | | | | | | | | | | | | | | | | | | | | |
|---|---|---|---|---|---|---|---|---|---|---|---|---|---|---|---|---|---|---|---|---|---|---|---|---|---|---|---|---|---|---|---|---|---|---|---|---|---|---|---|---|
| | 94 | PDAR | maviumcomplex(MAC) | | eria | | | | s | | | um | rium avium complex(MAC) | | | | | 100%) | 0%) | 7(100%) | | 0(100%);GO:0005524(100%) | ivity(100%);ATPbinding(100%) | | thetic process(100%) | (100%);IPR007344(100%);IPR027417(100%) | | | | | | | | | | | | | | |
| 575 | Variabl | SLFPEFSELFAAFPSFAGL | Mycobact | Bacteri | Actinoba | | Actinomyce | | Corynebacter | | Mycobacteriac | Mycobact | Mycobac | | | | | | | GO:0005 | extracellularregi | GO:0044 | proteinfoldingcha | GO:0001666(2 | responsetohypo | IPR0020 | 1.00E+0 | 1.00E+0 | 7.31E+0 | 0.1086 | 1.949E | 0.002094 | 0.9 | NS | NS | NS | NS | 25.792 | 4.9316E | 0.00048 |

| | | | | | | | | | | | | | | | | | | | | | | | | | | | | | | | | | | | | | | | |
|---|---|---|---|---|---|---|---|---|---|---|---|---|---|---|---|---|---|---|---|---|---|---|---|---|---|---|---|---|---|---|---|---|---|---|---|---|---|---|---|
| | e 95 | RPTFDTR | eriumtuberculosiscomplex | a | cteria | | tia | | iales | | eae | erium | terium tuberculosiscomplex | | | | | | | 576(75%);GO:0005618(75%);GO:0005737(50%) | on(75%);cell wall(75%);cytoplasm(50%) | 183(25%) | perone(25%) | 5%);GO:0006457(25%);GO:0009267(25%) | xia(25%);protein folding(25%);cellular response to starvation(25%) | 68(100%);IPR008978(100%);IPR031107(100%) | 0 | 0 | 5 | | -13 | 2 | | | | | | -08 | 192 |
| 576 | Varia | GSQTTGPK | Mycob | Bacte | Actino | | Actinom | | Corynеba | | Mycobacte | Mycob | Mycob | | | | | | | GO:00 | integralcom | | | GO:004278 | evasion of | IPR02 | 1.00E | 1.00E | 7.06E | NS | NS | NS | NS | NS | NS | NS | NS | | |

| | | | | | | | | | | | | | | | | | | | | | | | | | | | | | | | | | | | | | | | | |
|---|---|---|---|---|---|---|---|---|---|---|---|---|---|---|---|---|---|---|---|---|---|---|---|---|---|---|---|---|---|---|---|---|---|---|---|---|---|---|---|---|
| | ble 96 | | acterium tuberculosis complex | ria | bacteria | | ycetia | | cteriales | | riaceae | acterium | acterium tuberculosis complex | | | | | | | 16021(100%);GO:0005886(23%) | ponent of membrane(100%);plasma membrane(23%) | | | 3(8%) | host immune response(8%) | 1403(100%) | +00 | +00 | +04 | | | | | | | | | | | |
| 579 | Variable 99 | QYGPLTPAR | Mycobacterium tuberculo | Bacteria | Actinobacteria | | Actinomycetia | | Corynebacteriales | | Mycobacteriaceae | Mycobacterium | Mycobacterium tub | | | | | 2.7.11.1(100%);2. | Non-specific serine/threonine protein kinase (10 | GO:0005886(83%);G | plasma membrane(83%);integ | GO:0005524(100%); | ATP binding(100%);protein serine kinase activit | GO:0009405(11%);GO:0016036(6 | obsolete pathogenesis(11%);cellul | IPR000719(100%);I | 1.00E+00 | 1.00E+00 | 5.64E+04 | 8.9121 | 1.2011E-06 | 0.00038592 | 0.9 | NS | NS | NS | NS | 10.759 | 0.00016883 | 0.00016843 |

| | | | | | | | | | | | | | | | | | | | | | | | | | | |
|---|---|---|---|---|---|---|---|---|---|---|---|---|---|---|---|---|---|---|---|---|---|---|---|---|---|---|
| | | | siscomplex | | | | | | | | | | erculosiscomplex | | | | | 7.1.-(17%) | 0%);Phosphotransferases with an alcohol group as acceptor (17%) | O:0016021(78%);GO:0005576(6%) | ral component of membrane (78%);extracellular region (6%) | GO:0106310(100%);GO:0106311(100%) | y (100%);protein threonine kinase activity (100%) | %);GO:0032091(6%) | ar response to phosphate starvation (6%); negative regulation of protein binding (6%) | PR011009(100%);IPR008271(83%) |

### Supplementary Table 3- Correlation analysis of Bacterial phylum and metabolic pathway

| | Actinobacteria_Grey | Acidobacteria_Turquoise | Actinobacteria_Turquoise | Aquificae_Turquoise | Bacteroidetes_Turquoise | Chlamydiae_Turquoise | Chlorobi_Turquoise | Chloroflexi_Turquoise | Cyanobacteria_Turquoise | Deinococcus_Turquoise | Dictyoglomi_Turquoise | Elusimicrobia_Turquoise | Firmicutes_Turquoise | Proteobacteria_Turquoise | Spirochaetes_Turquoise | Tenericutes_Turquoise | Thermotogae_Turquoise | Verrucomicrobia_Turquoise | Actinobacteria_Yellow | Chlorobi_Yellow | Cyanobacteria_Yellow | Deinococcus_Yellow | Firmicutes_Yellow | Nitrospirae_Yellow | Proteobacteria_Yellow | Tenericutes_Yellow |
|---|---|---|---|---|---|---|---|---|---|---|---|---|---|---|---|---|---|---|---|---|---|---|---|---|---|---|
| Histidine metabolism | 0.303 | 0.420 | 0.461 | 0.410 | 0.354 | 0.377 | 0.403 | 0.353 | 0.374 | 0.553 | 0.397 | 0.335 | 0.354 | 0.406 | 0.459 | 0.414 | 0.448 | 0.353 | 0.254 | 0.276 | 0.299 | 0.311 | 0.356 | 0.227 | 0.612 | 0.252 |
| Arginine and proline metabolism | 0.262 | 0.286 | 0.314 | 0.317 | 0.328 | 0.502 | 0.268 | 0.395 | 0.314 | 0.518 | 0.254 | 0.233 | 0.385 | 0.277 | 0.487 | 0.274 | 0.217 | 0.277 | 0.293 | 0.326 | 0.361 | 0.242 | 0.276 | 0.331 | 0.247 | 0.370 |
| Aminoacyl-tRNA biosynthesis | 0.355 | 0.356 | 0.403 | 0.306 | 0.368 | 0.299 | 0.341 | 0.307 | 0.299 | 0.385 | 0.306 | 0.326 | 0.464 | 0.371 | 0.408 | 0.366 | 0.299 | 0.312 | 0.230 | 0.316 | 0.339 | 0.233 | 0.286 | 0.378 | 0.292 | 0.412 |
| Tryptophan metabolism | 0.436 | 0.393 | 0.403 | 0.426 | 0.382 | 0.342 | 0.402 | 0.441 | 0.396 | 0.575 | 0.454 | 0.447 | 0.427 | 0.650 | 0.406 | 0.525 | 0.434 | 0.462 | 0.332 | 0.349 | 0.229 | 0.435 | 0.257 | 0.434 | 0.359 | 0.250 |
| Tyrosine metabolism | 0.310 | 0.209 | 0.346 | 0.272 | 0.207 | 0.215 | 0.220 | 0.244 | 0.331 | 0.233 | 0.302 | 0.271 | 0.247 | 0.311 | 0.428 | 0.313 | 0.262 | 0.290 | 0.237 | 0.345 | 0.226 | 0.316 | 0.317 | 0.372 | 0.257 | 0.321 |
| Sphingolipid | 0.304 | 0.555 | 0.541 | 0.541 | 0.541 | 0.541 | 0.541 | 0.541 | 0.541 | 0.544 | 0.541 | 0.573 | 0.541 | 0.718 | 0.541 | 0.541 | 0.541 | 0.541 | 0.404 | 0.321 | 0.265 | 0.305 | 0.351 | 0.240 | 0.303 | 0.274 |
| Primary bile | 0.260 | 0.252 | 0.221 | 0.252 | 0.332 | 0.275 | 0.412 | 0.290 | 0.296 | 0.244 | 0.209 | 0.249 | 0.244 | 0.303 | 0.309 | 0.210 | 0.281 | 0.180 | 0.354 | 0.231 | 0.388 | 0.246 | 0.238 | 0.312 | 0.454 | 0.306 |

| | | | | | | | | | | | | | | | | | | | | | | | | | | |
|---|---|---|---|---|---|---|---|---|---|---|---|---|---|---|---|---|---|---|---|---|---|---|---|---|---|---|
| acid synthesis | | | | | | | | | | | | | | | | | | | | | | | | | | |
| Cysteine and methionine metabolism | 0.341 | 0.476 | 0.376 | 0.189 | 0.366 | 0.287 | 0.306 | 0.219 | 0.358 | 0.294 | 0.323 | 0.303 | 0.206 | 0.226 | 0.333 | 0.354 | 0.207 | 0.183 | 0.272 | 0.407 | 0.260 | 0.252 | 0.293 | 0.220 | 0.288 | 0.475 |

Supplementary Table 4-**:** List of primers of bile acid transport

| S.no | | Gene name | | |
|---|---|---|---|---|
| 1 | Mouse | FXR | F | ACTTCCGTCTGGGCATTCTGAC |
| | | | R | GCTGTAAGCAGAGCA TACTCCTC |
| 2 | Mouse | CYP8B1 | F | ACGCTTCCTCTATCGCCTGAA |
| | | | R | GTG CCTCAGACGCAGAGGAT |
| 3 | Mouse | OSTα/β | F | ATGCATCTGGGTGAACAGAA |
| | | | R | GAGTAGGGAGGTGAGCAAGC |
| 4 | Mouse | FGF15 | F | GAGGACCAAAACGAACGAAATT |
| | | | R | ACGTCCTTGATGGCAATCG |
| 5 | Mouse | GAPDH | F | CATCACTGCCACCCAGAAGACTG |
| | | | R | GAGCTTCCCGTTCAG |
| 6 | Mouse | SLC10A1 | F | TATGTTGCTGCTCATCATGC |
| | | | R | AATGCTGAGGTTCATGTCCC |
